# Novel Synthetic Promoter Armed Oncolytic Herpes Simplex Virus For Treatment of PAX3-FOXO1 Positive Rhabdomyosarcoma

**DOI:** 10.64898/2026.06.09.731216

**Authors:** Cole Peters, Miriam Valenzuela Cardenas, Conner Kidd, Analynn Lechien, Andrew Dinh, Moe Kawakami, Jaden Nguyen, Allison Flores, Stephen Ma, Lance Heady, Giorgia Del Vecchio, Matteo Pellegrini, Theodore S. Nowicki

## Abstract

**Background:** Due to the difficulty in targeting oncogenic transcription factors, there are no targeted treatments available for pediatric patients afflicted with PAX3-FOXO1 fusion positive alveolar rhabdomyosarcoma. Additionally, alveolar rhabdomyosarcomas are immunologically cold tumors inherently resistant to immune checkpoint blockade. Viral vectors present a unique opportunity to address this gap in treatment strategies as their presence causes immune infiltration, and a virus can be engineered to rely on tumor specific transcriptional networks for its own replication.

**Methodology:** We engineered a synthetic promoter which relies on PAX3-FOXO1 to drive expression of a downstream gene. We then placed the HSV-1 RL1 gene encoding ICP34.5 under this promoter within the G47Δ oncolytic herpes simplex virus (oHSV), generating a novel oHSV targeting cells harboring the PAX3-FOXO1 fusion gene, which we name oRP3Fus (“Orpheus;” oncolytic recombinant G47Δ targeting PAX3-FOXO1 virus). PAX3-FOXO1 specificity was determined by qRT-PCR, protein expression, and chromatin immunoprecipitation. Antitumor efficacy of oncolytic virus and anti-PD1 antibody was determined using immuno-competent mice harboring syngeneic PAX3-FOXO1 expressing rhabdomyosarcoma tumors. Bulk RNA sequencing was performed on infected and mock infected fusion positive and negative cells for differential gene expression analysis.

**Results:** oRP3Fus kills PAX3-FOXO1 positive human rhabdomyosarcoma lines, while remaining non-toxic to normal human skeletal muscle cells *in vitro*. In immunocompetent mouse models, both novel and parental viruses improved tumor control when combined with anti-PD1, while oRP3Fus treatment increased T cell infiltration within tumors. RNA and DNA extraction from tumors demonstrate viral expansion within oRP3Fus treated tumors up to three weeks post treatment. Meanwhile oRP3Fus treated mice displayed decreased amounts of necroptosis-associated gene expression within normal organs as compared to groups treated with parental virus. Bulk RNA sequencing of fusion positive and negative human rhabdomyosarcoma lines demonstrated PAX3-FOXO1 increases expression of genes involved in the TNFa-NFκB signaling axis, as well as innate and adaptive immune signaling. Additionally, ICP34.5 expression from oRP3Fus reduces RIPK1 activation.

**Conclusions:** ICP34.5-armed oHSV (oRP3Fus) combined with anti-PD1 provides effective antitumor activity against PAX3-FOXO1-positive alveolar rhabdomyosarcoma *in vivo* and should be further explored for fusion oncogene-driven tumors.

**What is already known on this topic?:** PAX3-FOXO1 fusion-positive and negative rhabdomyosarcoma (RMS) is highly resistant to immune checkpoint blockade. Previously, a PAX3-FOXO1 driven oncolytic adenovirus showed efficacy in an immuno-compromised fusion-positive RMS model. In fusion-negative, immuno-competent RMS models, oncolytic herpes viruses have been shown to improve survival when combined with anti-PD1 therapy, but combinational efficacy was only observed with sex-mismatched models and was never tested on fusion-positive RMS tumors in immunocompetent mice.

**What this study adds?:** We improved the specificity of the synthetic PAX3-FOXO1 targeted promoter. We also show, for the first time, efficacy of oncolytic herpes simplex virus-1 in immunocompetent mice harboring orthotopic PAX3-FOXO1-expressing RMS tumors. Our study also demonstrates improvement in survival efficacy in sex-matched tumors, whereas previous reports did not observe complete responses except when using sex-mismatched tumor models.

**How this study might effect research, practice, or policy?:** The synthetic promoter generated here can be used for future gene therapy or targeting efforts to study PAX3-FOXO1 or target PAX3-FOXO1 tumors. We observe anti-PD1 therapy benefits survival when paired with an oHSV, demonstrating oncolytic virus can sensitize PAX3-FOXO1 expressing RMS tumors to immunotherapy.

## BACKGROUND

Rhabdomyosarcoma (RMS) is the most common soft tissue sarcoma in children, with alveolar rhabdomyosarcomas (aRMS) accounting for 40% of these tumors^1^. The majority of aRMS harbor a translocation derived fusion gene which drives tumor spread, most notably PAX3-FOXO1 (P3F). The presence of P3F is strongly associated with more aggressive disease and worse prognosis, making this fusion protein extremely relevant to translational cancer research^2–4^. P3F acts as an oncogenic transcription factor, remodeling tumor gene expression to resemble a developmental state between embryonic and fetal muscle tissue^5^. This alteration relies on continuous PAX3-FOXO1 expression, and its knockdown ablates or attenuates cell growth in fusion positive tumor cells^6^. Unfortunately, oncogenic transcription factors have been notoriously difficult to treat with small molecule inhibitors or other chemical compounds^7,8^. Paired with the low overall mutational burden of RMS tumors and their resistance to checkpoint inhibitors^9,10^, patients with aRMS have few options for treatment in relapsed or refractory disease not responsive to standard of care therapies. While the PAX3-FOXO1 protein may be difficult to target with a small molecule, its role as a unique transcription factor could be harnessed to direct a therapeutic viral vector to tumor cells.

G47Δ is a third-generation oncolytic herpes simplex virus (oHSV) which was developed to treat glioblastoma and has been safely used in mice and human patients^11–14^. It was generated from two previous viruses which inserted the LacZ gene within the viral ribonucleotide reductase (ICP6/U_L_49) and deleted the neurovirulence RL1 genes encoding the γ34.5/ICP34.5 protein. A further deletion of the U_s_12/ICP47 ORF complements the loss of ICP34.5 and promotes MHCI expression in infected cells^11^.

Multiple groups have determined PAX3-FOXO1 drives transcription via physical binding to several promoters and enhancer sequences active in fetal muscle and aRMS tissue^15–19^, with one group already generating an oncolytic adenovirus driven by a minimal myogenin promoter to target RMS cells in an immuno-compromised mouse model^20^. Here we combine several paired recognition sites and enhancer sequence motifs bound by PAX3-FOXO1 with a myogenin promoter region to make a synthetic promoter which allows downstream transgene expression only in the presence of PAX3-FOXO1 protein. We then modified the G47Δ oHSV to express ICP34.5 within fusion positive RMS cell lines, which we refer to as oRP3Fus (“Orpheus;” oncolytic recombinant G47Δ targeting PAX3-FOXO1 virus). This novel virus is tolerated in both human skeletal muscle and immunocompetent mice and improved survival and efficacy of anti-PD1 antibody treatment in mice harboring orthotopic PAX3-FOXO1+ aRMS tumors.

## METHODS

### Cell lines, plasmids, and viruses

All human and murine RMS cell lines were cultured in RPMI1640 supplemented with 10% heat inactivated serum, 1x Penicillin/Streptomycin, and 1x Glutamax. HEK293T cells were cultured in DMEM with the same supplementation. Human RMS lines RH2, RH4, RH28, RH41, and JR1, were generously donated by Gerard Grosveld - St. Jude Children’s Research Hospital. Murine eRMS M3-9-M cells were generously donated by the laboratories of Timmothy Cripe (Nationwide Children’s Hospital) and Crystal Mackall (Stanford University). Vero (CCL-81), Skeletal muscle (950-010), HEK293T (CRL-3216), RH30 (CRL-2061), and RD (CCL-136) were purchased from ATCC. Cell lines were grown as directed by ATCC. Plasmids harboring the: PAX3-FOXO1 cDNA were generously donated by Frederic Barr – NCI; minimal myogenin promoter was generously donated by Hideki Yoshida (Kyoto Prefectural University of Medicine) and Masato Yamamoto (University of Minnesota Medical School).

Lentivirus DNA were purchased from System Biosciences and used to generate RMS cells with empty or PAX3-FOXO1 expressing lentiviruses. F-strain herpes virus G47Δ as well as the shuttle vector svICP6 plasmid were kind gifts from Samuel D. Rabkin (Harvard University). PGL3 plasmid backbone was a gift from Debrya Groskreutz (Addgene plasmid # 212936). Enhancers and promoter sequences were PCR amplified using primers **(Supplemental Table 1)**. All cell lines used to overexpress human PAX3-FOXO1 or EMPTY lentivirus were transduced, selected with puromycin (5-10ug/mL) for three passages, and then tested via Western blot **(Supplemental Figure 1A,B).**

### *In Vivo* and histological assays

All experiments were performed in accordance with the NIH *Guidelines for the care and use of laboratory animals*, and all procedures were approved by the UCLA Chancellor’s Animal Research Committee (ARC-2022-070). For tumor implantation, male 6wk old C57/Bl6J mice were implanted with 5 x10^5^ M3-9-M:PAX3-FOXO1 tumor cells in the left gastrocnemius muscle via a 22-gauge needle. Virus (1 x10^8^ PFU) was injected (1-3 times depending on the experiment) intratumorally. Anti-mouse PD1 (BioXcell BE0273 clone RMP1-14) in PBS was injected intraperitoneally three times a week for two weeks at a dose of 250ug/20g. Tumor volume was measured by caliper using the equation: 3.14*((Longer diameter) * (shorter diameter^2))/6. Body condition of mice was monitored on a BCS scale of 1 (poor) to 5 (obese) with 3 (normal) being the starting condition^21^. Additionally, we ensured mice were able to bear weight on implanted legs, those that could not were euthanized. Upon reaching a tumor volume of 1000mm^3^ mice were euthanized by isoflurane and cervical dislocation. Tissue sections taken for IHC were fixed in 10% neutral buffered saline, stored in 70% Ethanol and transferred to the UCLA Translational Pathology Core Laboratory (TPCL) for paraffin embedding, slide generation and staining with antibody against CD3 (DAKO A0452) or Myogenin (Abcam ab1835) as well as hematoxylin and eosin staining. Slides were scanned and analyzed using QuPath^22^.

### Flow Cytometry

Tumors and spleens were mechanically dissociated and passed through a 40um mesh filter for single-cell suspension. Cells in single-cell suspension were resuspended in FACS buffer containing 0.5mM EDTA and 2% FBS in phosphate-buffered saline (PBS). Prior to staining, cells in single-cell suspension were either incubated with LIVE/DEAD Fixable Yellow (ThermoFisher) or 7-AAD for cell viability. Cells were washed and then stained with fluorescence conjugated antibodies for surface markers at a 1:50 concentration for 30min at 4°C. Cells were given another wash and analyzed on an Attune NxT 6 cytometer operated by UCLA Jonsson Comprehensive Cancer Center (JCCC) and the Center for AIDS Research Flow Cytometry Core Facility. Data were analyzed using FlowJo (v10) software. For gating strategy see **(Supplemental Data 1)**.

### Quantitative PCR

RNA and DNA was isolated using the AllPrep DNA/RNA isolation or the DNeasy Blood and Tissue kits (Qiagen). RNA was treated with RQ1 DNAse (Promega) prior to cDNA synthesis with the SuperScript VILO kit (Thermo). qRT-PCR was carried out using the StepOne Plus 6 series PCR machine and SYBR PowerUp Master mix (Thermo). Primers used are listed in **Supplemental Table 1**. All calculations shown utilize primers with efficiencies measured between 85-110%, and use the ΔΔCt calculation method based on either 18sRNA or GAPDH as a calibrator gene.

### RNA Sequencing

RD:EMP or RD:P3F cells were infected with G47Δ or oRP3Fus at MOI(2) for 6 hours before total RNA was harvested via RNeasy kit (QIAGEN). RNA was assayed by NanoDrop & TapeStation for quality and then libraries generated using the Roche Kapa mRNA stranded library prep kit. Libraries were sequenced by the UCLA Technology Center for Genomics and Bioinformatics. Differential expression analysis was carried out with the DESeq2 package v1.46.0. The design formula included factors for cell type (Empty, PAX3-FOXO1) and virus (mock, G47, oRP3Fus). Significantly differentially expressed genes (DEGs) were defined as those with an adjusted p-value (padj) < 0.05 and an absolute log₂ fold change (|log₂FC|) ≥ 1. Gene Ontology (GO) and gene set enrichment analysis (GSEA) were conducted. Heat maps were generated using gene lists sourced from gsea-msigdb.org and are listed in (**Supplemental Data 2)**.

### Statistical Analysis

All analysis was performed using GraphPad PRISM ver10.6.1. Tests for significant differences are labelled within figure legends. Two-way ANOVA with Holm-Šidák or Tukey multiple comparison testing was used to access differences between groups when measuring viral titer or tumor size through time, or differences of tumor cell infiltration numbers between *in vivo* experimental groups. One-way ANOVA with Šidák or Tukey multiple comparison testing was used for assays where a single variable was compared between groups. Differences in survival were measured via long-rank (Mantel-Cox) test. p values of <0.05 were considered significant for all analyses. RNAseq analysis for differential gene expression used DESeq2 package v1.46.0. Significant DEGs were discovered after Benjamini-Hochberg FDR testing, significance was determined if adjusted p values <0.05, and a log_2_ fold change of at least 1 were observed.

For details on certain assays please consult the **supplemental methods** section.

## RESULTS

### Generation of PAX3-FOXO1 Selective Synthetic Promoter

We amplified gene promoter or enhancer fragments shown to be targeted by PAX3-FOXO1 and cloned them upstream (5’) or downstream (3’) of a firefly luciferase reporter gene. To determine which promoter endowed PAX3-FOXO1 specificity, we combined enhancer sequences with promoter gene sequences from human myogenin (MYOG), myogenic differentiation 1 (MYOD1), myogenic factor 5 (MYF5), or a nonspecific control promoter from SV40 virus **(Figure 1A)**. Plasmids were transfected into HEK293T and RD RMS cell lines stably transduced with lentivirus to express PAX3-FOXO1 (P3F) or empty (EMP) transgene **(Supplemental Figure 1A)**. Luciferase expression was compared between P3F and EMP cells in relation to a promoterless pGL3-Basic-Luciferase control plasmid to determine specificity of the synthetic promoters **(Figure 1B,C; Supplemental Figure 2A-F)**. We observed 5’ALK or 5’PRS1 enhancers paired with a MYOG promoter allowed luciferase expression in the presence but not absence of PAX3-FOXO1. We further validated our top constructs by swapping luciferase with GFP and observed the MYOG promoter paired with 5’PRS1 or 5’ALK enhancers generated a synthetic promoter which allowed expression only in the presence of PAX3-FOXO1 **(Figure 1D)**. In contrast, we observed other candidate promoters allowed leaky expression of GFP reporter **(Supplemental Figure 3A)**. To ensure our observations were due to promoter-controlled transcription, we performed qRT-PCR for luciferase expression. Comparing mRNA from RD:EMP and RD:P3F cells transfected with equal amounts pGL3-5’-PRS1 or 5’ALK showed significant increases of luciferase mRNA in the presence of P3F over promoterless Basic controls **(Figure 1E)**. To ensure that our synthetic promoter did not cross react with PAX3 or other PAX proteins we performed the same reporter assay in HEK293T cells transduced to overexpress PAX3, PAX4, or PAX7. The 5’PRS1 and 5’ALK MYOG constructs showed no increased expression of luciferase in comparison to HEK293T:EMP cell control **(Supplemental Figure 3B)**. As FOXO1 is ubiquitous in all cell lines, we did not test our constructs in cells overexpressing it. To ensure PAX3-FOXO1 was binding our construct we performed chromatin immunoprecipitation (ChIP) for PAX3-FOXO1 following transfection of HEK293T:P3F cells with pGL3- 5’PRS1-MYOG plasmid. QPCR of ChIP eluates demonstrated the presence of P3F at our synthetic PRS1 enhancer, MYOG promoter, and Luciferase transgene region over control IgG **(Figure 1F)**. We deemed the synthetic 5’PRS1-MYOG promoter allows controlled expression in the presence of PAX3-FOXO1, and incorporated it into our novel oHSV.

**Figure 1.**
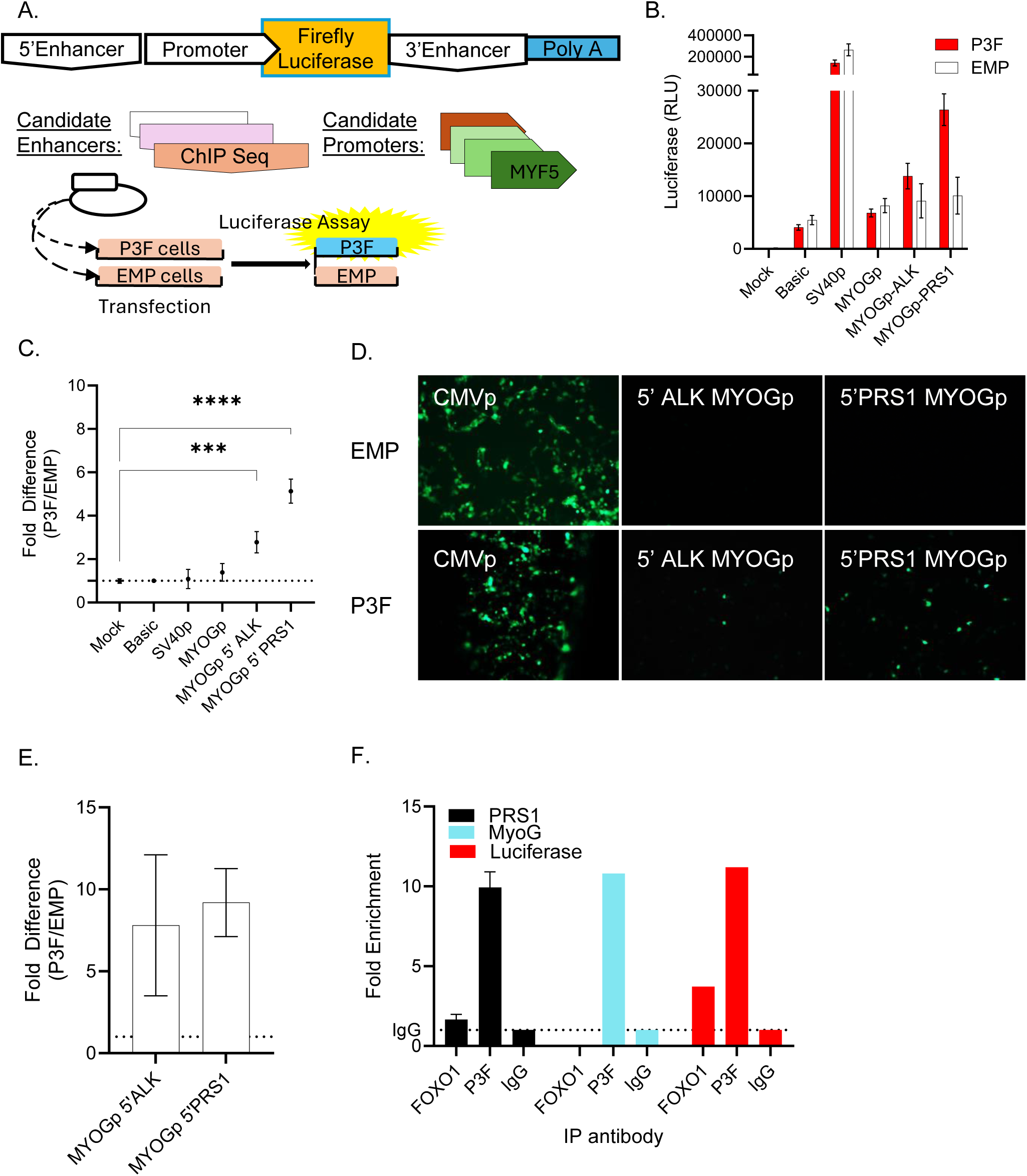
Generation of a PAX3-FOXO1 specific promoter system. (A) Diagram of plasmid constructs and methodology to determine promoter activity in the presence or absence of PAX3-FOXO1. (B) Relative light units (RLU) after transfection of candidate constructs, and (C) fold expression detected in P3F/EMP cells in relation to the promoterless Basic control. One-way ANOVA with Šídák post-hoc test (*** p<0.005, **** p<0.001), error bars shown represent mean SEM, (n>48). (D) GFP expression under synthetic promoters in P3F or EMP transduced HEK293T cells. (E) RTqPCR to determine reporter mRNA in relation to pGL3-basic via ΔΔCt to18sRNA (n=3). (F) QPCR of DNA fragments collected by ChIP with indicated antibodies, controlled for %input.

### Generation of oRP3Fus/rG47-PRS1 to target PAX3-FOXO1 RMS tumors

To the pGL3-5’PRS1-MYOG reporter plasmid, we replaced the luciferase gene with the F strain HSV-1 RL1 gene encoding then ICP34.5/γ34.5 protein, added a human ubiquitin promoter driven GFP, and inserted this into the ICP6-LacZ gene of oncolytic herpes virus G47Δ, generating rG47-5’PRS1-MYOGpro-ICP34.5;hUBQpro-GFP (rG47-PRS1/oRP3Fus). We generated a similar virus utilizing the 5’ALK enhancer as well (rG47-ALK) **(Supplemental Figure 4A)**, however only oRP3Fus will be described herein. Insertion of the synthetic promoter was verified by restriction digest and GFP plaque selection **(Supplemental Figure 4B-C)**, as well as sanger sequencing **(Supplemental Data 3)**. We determined that oHSV G74Δ effectively infects human P3F expressing RMS cell lines RH2, RH4, RH30, and RH41 as well as fusion negative lines RD and JR1 **(Figure 2A/ Supplemental Figure 4D)**. Similarly viral true-late gene expression of glycoprotein C was similar between P3F expressing and negative RMS lines (**Figure 2B)**. We did detect reduction of viral growth in PAX3-FOXO1 positive RH30 cells **(Figure 2C-D)**, suggesting differences of permissiveness within RMS lines to oHSV. Supplying ICP34.5 expression in trans increased G47Δ growth within RH30 cells suggesting ICP34.5 improves viral potency within fusion positive RMS lines **(Figure 2E)**. We next compared our novel virus oRP3Fus, armed with 5’PRS1-MYOG controlled ICP34.5 expression, against parental G47Δ. We see oRP3Fus kills PAX3-FOXO1-expressing RMS cell lines faster than G47Δ, while retaining similar cell killing kinetics in fusion negative cells **(Figure 2F)**. Additionally, oRP3Fus grows to higher titers than G47Δ in RH30 cells, and similar titers in several other RMS lines **(Figure 2G; Supplemental Figure 4E-H)**. Western blotting for ICP34.5 confirmed protein expression is confined to fusion positive RMS cell lines during oRP3Fus infection (**Supplemental Figure 4I).** Lastly, infection of normal human skeletal muscle cells with either virus did not lead to cell death after forty-eight hours in culture, suggesting oRP3Fus would maintain the same safety profile as its parental G47Δ virus **(Figure 2H)**.

**Figure 2.**
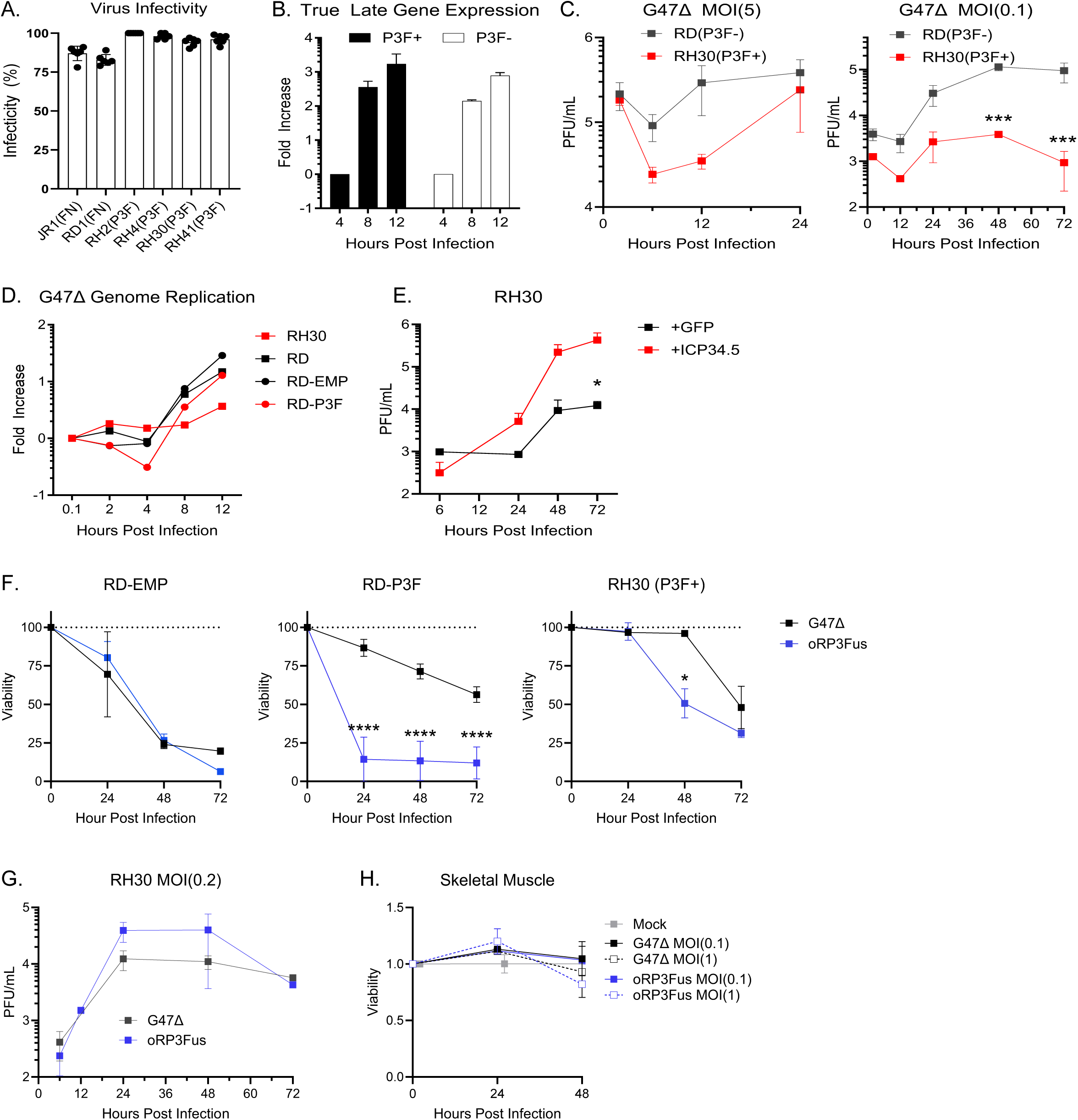
Oncolytic HSV G47Δ and oRP3Fus effectively infect and kill RMS lines. (A) RMS cells infected with G47Δ and observed for LacZ expression by Xgal staining (n=5). (B) RTqPCR from infected cell lysates for true late gene Glycoprotein C (n=2). (C) Plaque assay of RMS cell lysates after infection at MOI(5) or MOI(0.1) (n=3). (D) QPCR of viral genomes during infection of RMS cell lines. Fold increase compared to 10min infected lysate (ΔΔCt:GAPDH). (E) Plaque assay of RH30 cells transfected with 5’PRS1-MYOG driven GFP or ICP34.5 and then infected with G47Δ at MOI(0.1) (n=3). (F) Viability testing of RMS cells infected with oRP3Fus or G47Δ (n=3). (G) Plaque assay of RH30 cells infected with oRP3Fus or G47Δ. (H) Viability assay of human skeletal muscle cells infected with oHSV (n=3). Error bars represent mean and standard deviation, Two-way ANOVA with Holm-Šidák multiple comparisons test between oRP3Fus and G47Δ (*p<0.05; **p<0.001;****p<0.0001).

### Combination of oncolytic herpes virus with anti-PD1 antibody improves survival of mice harboring PAX3-FOXO1 RMS tumors

To determine the safety of oRP3Fus virus, we injected escalating doses of virus into the gastrocnemius muscle of 6-week-old male C57/Bl6J mice, and observed no lethality or decline in body condition score **(Supplemental Figure 5A-B)**. We then transduced M3-9-M murine RMS cells with PAX3-FOXO1 or empty lentivirus and assayed cell, virus, and tumor growth potential, finding that PAX3-FOXO1 expression led to slightly faster cell growth **(Supplemental Figure 1B; Supplemental Figure 5C-D)**. P3F-positive RMS cells formed tumors when injected into the gastrocnemius muscle or flanks of male C57/Bl6J mice with similar kinetics as control RMS cells **(Supplemental Figure 5E-F)**. Like human RMS tumors, P3F-positive RMS leg tumors are myogenin positive and low in T cell infiltration **(Supplemental Figure 5G)**. We observed consistent implantation in male mice, whereas implantation into female mice was inconsistent (**Supplemental Figure 5H**). Virus growth within M3-9-M:P3F cells was similar between oRP3Fus and parental virus *in vitro*, while oRP3Fus grew slightly slower in the control RMS cells **(Supplemental Figure 4F)**.

To determine whether either oHSV provided therapeutic benefit in P3F expressing tumors, we implanted M3-9-M:P3F cells into C57/Bl6 mice gastrocnemius muscle, with and waited until leg volume increased to 50-100mm^3^ before injecting one or three doses of 1E8 PFU of G47Δ or oRP3Fus into the tumor followed by intraperitoneal injection of anti-mouse PD1 antibody three times a week **(Figure 3A)**. Treatment with single agent virus or in combination with anti-PD1 significantly delayed tumor growth in mice receiving a single or triple-dose of virus **(Figure 3B-C; Supplemental Figure 6A-D)**. When combined with anti-PD1 antibody treatment, we observed significant increases in mice survival with either single or triple-dose virus. However, only triple-dose virus injection resulted in significant increases in survival without combination with anti-PD1 treatment **(Figure 3D-E)**. Similar to other reports using muscle or flank RMS tumor models, no benefit of anti-PD1 therapy alone was observed. Notably before takedown for analysis, we achieved two complete tumor regressions in mice treated with a triple injection of oRP3Fus virus and anti-PD1 and stable disease in the G47Δ combination group (**Supplemental Figure 6B)**. IHC of the gastrocnemius implantation site of these oRP3Fus-treated mice revealed possible tertiary lymphoid structures and almost no tumor presence **(Figure 3F**).

**Figure 3.**
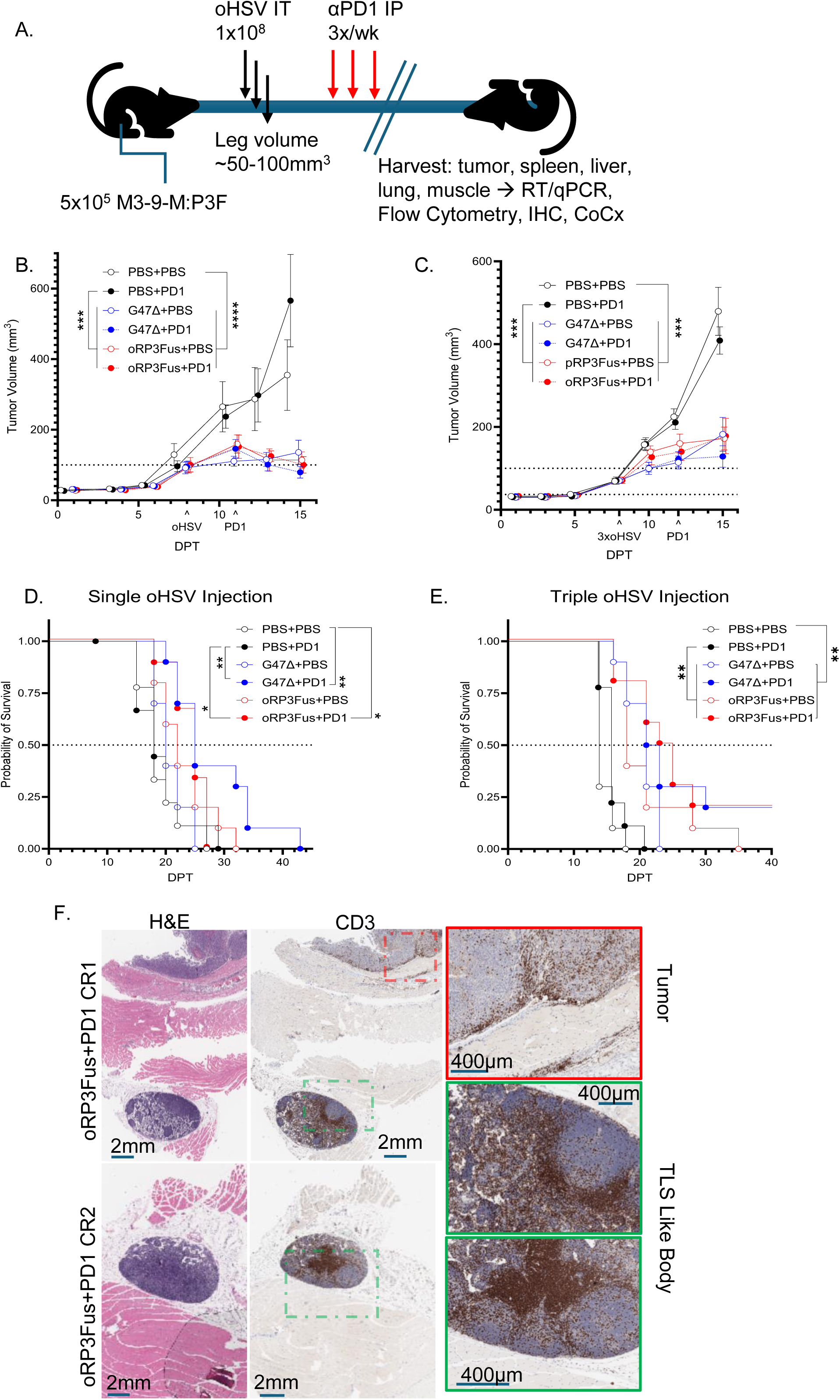
Oncolytic HSV G47Δ and oRP3Fus combine with anti-PD1 therapy to improve survival in immunocompetent RMS. (A) Diagram of study design for treatment of orthotopic M3-9-M:P3F tumors in male C57/Bl6J mice. Tumor volume measurements during treatment with (B) single (C) or triple virus injection. Survival graphs from mice treated with (D) single virus injection or (E) triple virus injection. (F) H&E and CD3 staining of long term survivors from the group treated with oRP3Fus triple injection and anti-PD1; green box indicate TLS, red box indicate residual tumor. For all in vivo models; n=9-10 per group; Error bars represent mean and SEM. Tumor analysis uses two-way with Holm-Šidák multiple comparisons test (***p<0.001; ****p<0.0001). Survival differences measured via Mantle-Cox testing (*p<0.05; **p<0.01).

### Oncolytic virus treatment leads to influx of lymphocytes into RMS tumors and splenocyte activation

Flow cytometry of tumor mass upon euthanasia for tumor infiltrating lymphocytes (TILs) did not correlate significantly with survival. We observed increases in CD45+ cells within combination treated tumor masses over other groups **(Figure 4A-B)**. However, there were no significant differences between groups. Similarly, when measuring infiltration by CD4+ or CD8+ we saw no significant differences between groups. However, when measuring total T cells (CD3+/CD4+/CD8+) we observe significant increases in T cell infiltration in the combination treatment groups over PD1 treatment alone in the single injection virus cohort, and over anti-PD1 and mock treatment groups in the triple injected oRP3Fus cohort **(Figure 4C)**, while not observing significance differences between the combination G47Δ and control treatment groups in the triple injection cohort. Additionally, we observe higher T cell infiltration within oRP3Fus-treated tumors as compared to those treated with parental G47Δ in the triple injection cohort. We observed significant decreases in PD1+CD8+ TILs in single-dose virus combination treated mice compared to mock treatments. Paradoxically, we did not observe this phenotype in triple dose virus combination cohorts **(Figure 4D)**.

**Figure 4.**
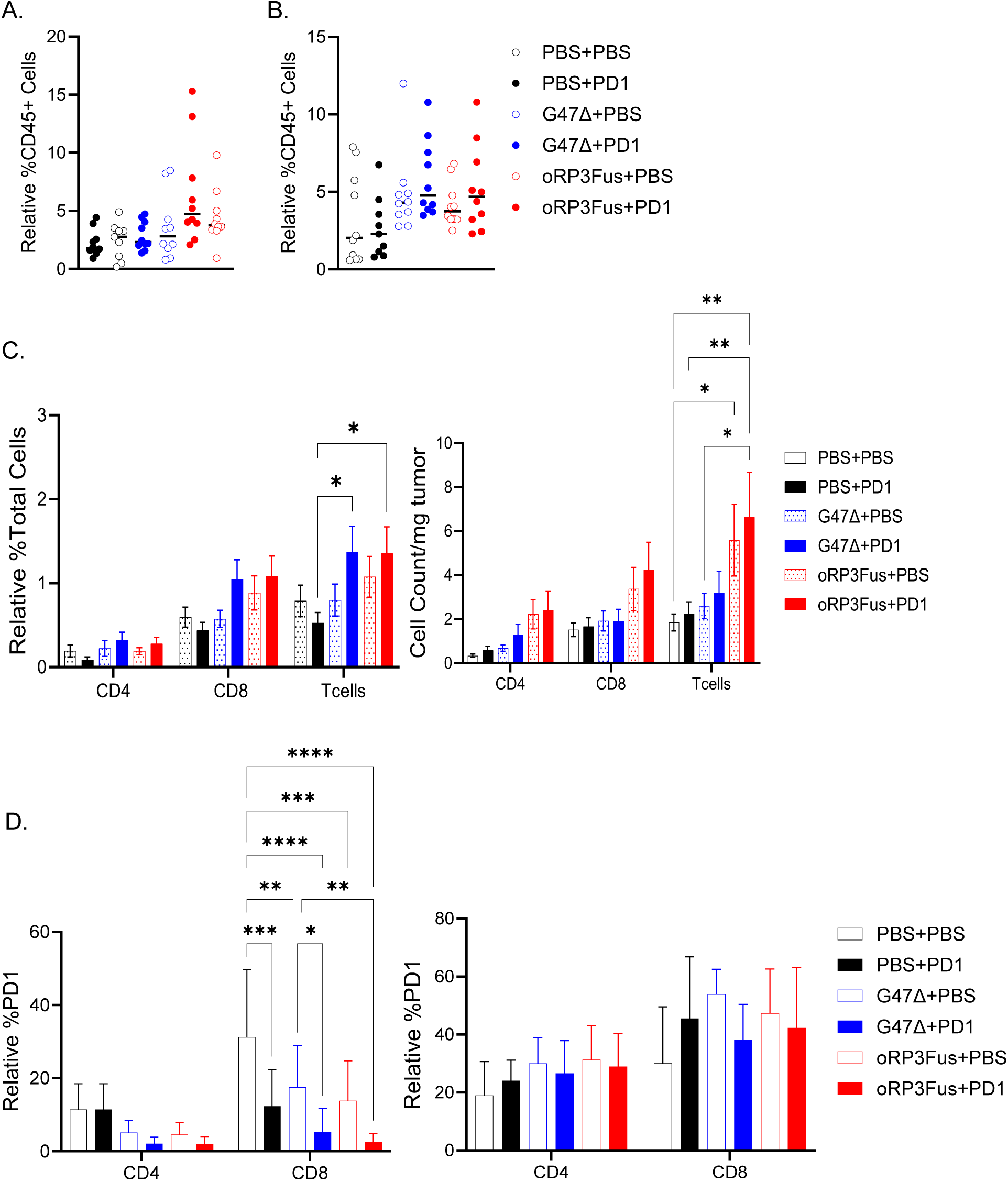
Infiltration of M3-9-M:P3F orthotopic tumors by leukocytes. Quantification of CD45+ cells by flow cytometry of tumors isolated from (A) single virus or (B) triple virus injected cohorts. (C) T cell infiltration of tumors as measured by flow cytometry of (left) single and (right) triple injected cohorts. (D) PD1 expressing CD4 or CD8 T cells observed via flow cytometry from (left) single virus or (right) triple virus injected cohorts. (E) ELISA for murine IFNγ from splenocyte and M3-9-M:P3F co-cultures in the presence or absence of low dose virus. Error bars indicate Mean and standard deviation. One way ANOVA with Tukey multiple comparisons test (A,B) or Two-way ANOVA (C,D) with Holm-Šidák tests were used to compare group means (*p<0.05; **p<0.01; ***p<0.005; ****p<0.001).

To determine whether treatment was generating adaptive memory responses, we performed co-cultures with splenocytes harvested upon euthanasia with either M3-9-M:P3F mock infected or G47Δ infected cells, and tested supernatants for mouse IFNγ after 20 hours in co-culture. While not statistically significant, we observed an increase in IFNγ production from splenocytes from all virus treated groups over mock or anti-PD1 alone treated **(Figure 5A)**. In contrast, we observe significant increases in IFNγ secretion when splenocytes from triple-dose G47Δ or oRP3Fus treated mice were compared mice treated with PBS or anti-PD1 alone. In parallel, control naïve splenocytes cultured with virus did not increase IFNγ secretion over naïve splenocyte culture alone, suggesting the increases in splenocytes activation was not due to acute virus infection **(Figure 5A)**. Additionally, we incubated splenocytes from the complete responders with CFSE and measured their division when exposed to P3F-positive RMS cells used to propagate tumors or to control RMS isogenic cells not expressing PAX3-FOXO1. We observed a significant increase in splenocyte division when co-cultured with M3-9-M:P3F but not when co-cultured with M3-9-M:EMP **(Figure 5B)**, suggesting a possible PAX3-FOXO1-associated antigen T cell response may have been responsible for tumor clearance in these mice. Flow cytometry of mock or virus infected M3-9-M and human RD fusion positive and negative cells revealed virus infection does not alter PD-L1 expression, which was expressed on both cell lines **(Figure 5C)**.

**Figure 5.**
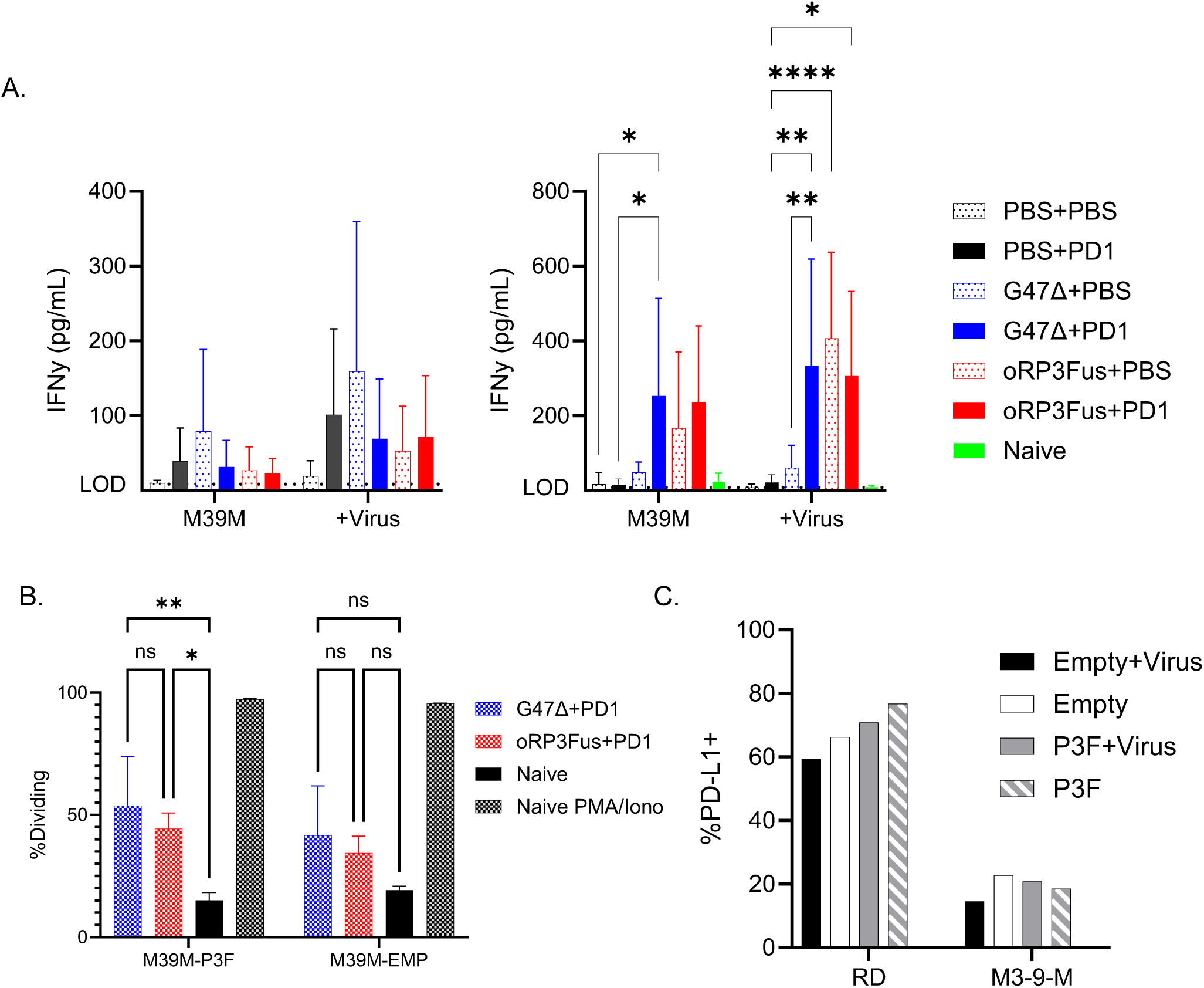
Memory response after treatment to M3-9-M cell co-culture. (A) ELISA for murine IFNγ from splenocyte and M3-9-M:P3F co-cultures in the presence or absence of low dose virus. (B) CFSE labelled splenocyte co-culture for 3 days using long term survivor G47Δ or G47-PRS1 splenocytes with M3-9-M:P3F or M3-9-M:EMP cells. (C) PD-L1 surface staining of human RD or murine M3-9-M cell lines. Error bars indicate Mean and standard deviation. Two-way ANOVA with Holm-Šidák multiple comparisons test were used to compare group means (*p<0.05; **p<0.01; ***p<0.005; ****p<0.001).

### PAX3-FOXO1 expression drives NFκB signaling in RMS

To better characterize whether PAX3-FOXO1 expression within RMS cells affects herpes virus infection, we performed bulk RNA sequencing on P3F-positive and negative human RMS cells infected with G47Δ or oRP3Fus at several times post infection. Gene set enrichment analysis (GSEA) of differentially expressed genes (DEGs) demonstrated PAX3-FOXO1 expression primarily increased gene expression associated with muscle function and development in RD cells **(Figure 6A; Supplemental Figure 7A)**. The other notable increases in gene expression due to PAX3-FOXO1 expression were associated with TNFa-NFκB signaling axis, which was even more pronounced during virus infection **(Figure 6B-C; Supplemental Figure 7B-C)**. To explore this and the effects oHSV infection has on NFκB signaling in RMS lines, we performed western blotting on protein lysates from mock or virus infected P3F -positive or negative human and murine RMS cell lines. In multiple RMS cell lines, we observed an increase in canonical NFκB signaling via phosphorylated NFκB p65 and MAPK p38 **(Figure 6D; Supplemental Figure 8)**. We observed that P3F-positive RMS cells displayed increased NFκB p100 activation marked by phosphorylation and cleavage into p52 subunit as compared to to P3F-negative RMS cells **(Figure 6D)**. Paired with our RNA sequencing data, this suggests that PAX3-FOXO1 increases NFκB signaling via activation or sensitization to the non-canonical NFκB pathway in addition to the canonical pathway, and that oHSV infection exacerbates this activation. Quantitative RT-PCR for AP1 transcription factor components and NFκB related targets also demonstrated high expression of cFOS, JUNB, and REL in human RMS lines during PAX3-FOXO1 over-expression and during G47Δ or oRP3Fus infection **(Figure 6E)**. Similarly, M3-9-M cells increased cFOS and RHOB after PAX3-FOXO1 overexpression and during infection with both viruses. Protein levels of cFOS were increased in western blotting corroborating the increased mRNA we observed was actually being translated during infection (**Supplemental Figure 8, Supplemental Figure 9**). This was also corroborated by increased expression of genes associated with TNFA signaling and interferon/anti-viral signaling when PAX3-FOXO1 is expressed in RMS cells and after infection with oHSV **(Supplemental Figure 7C).**

**Figure 6.**
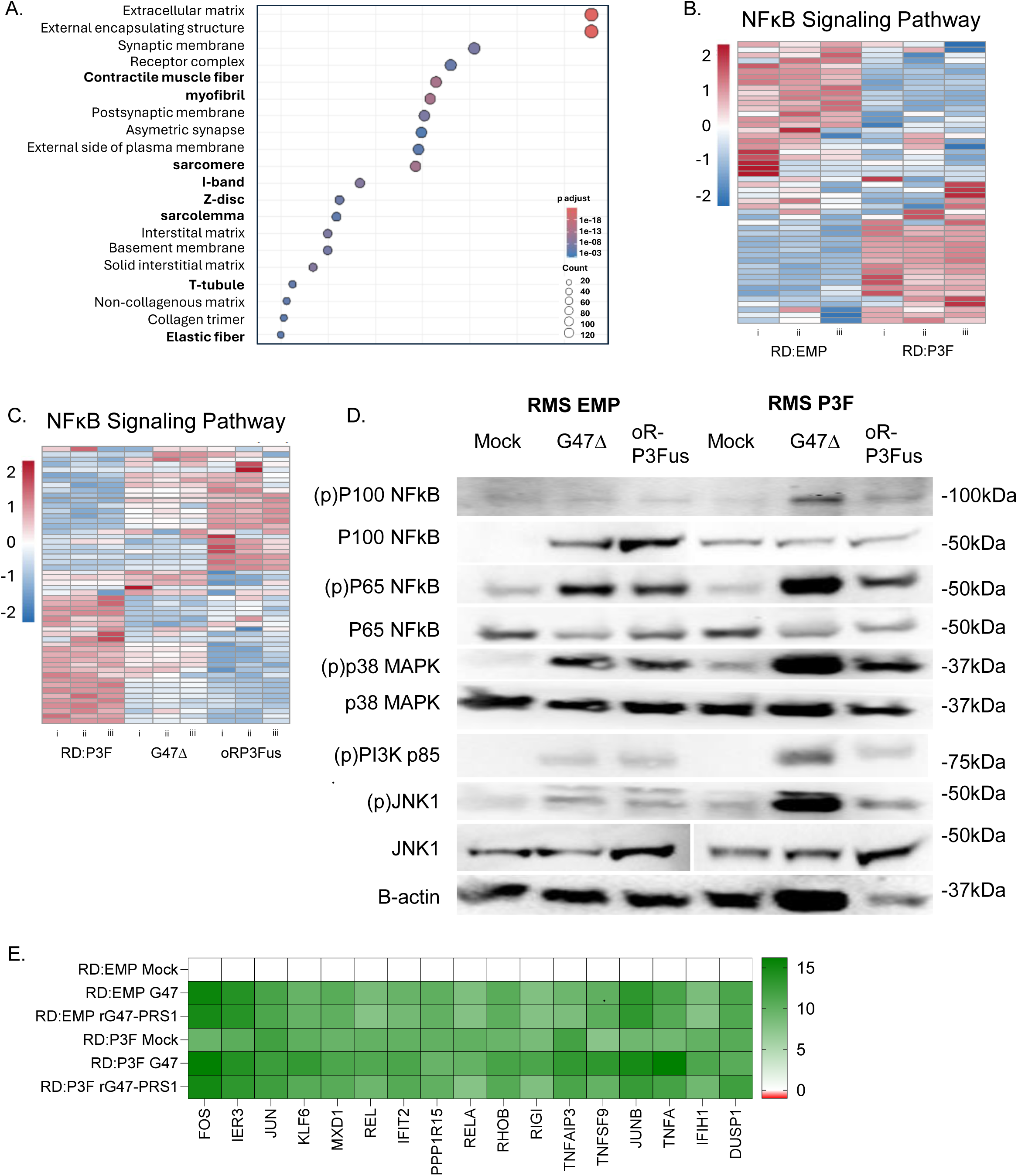
PAX3-FOXO1 and HSV infection activates NFκB signaling within RMS. (A) GSEA showing biological processes upregulated in RD:P3F vs RD:EMP cells. Heat maps of DEG relevant to NFκB signaling comparing the effect of (B) PAX3-FOXO1 expression, or (C) virus infection of P3F cells, i-iii represent sample replicates. (D) Western blot of EMP or P3F RMS cells during infection. (E) qRT-PCR for TNFA or NFκB pathway involved genes in cells 6-hour post infection at MOI(2) as compared to Mock RD:EMPTY cells. Gene lists for heat maps in supplemental data.

### ICP34.5 reduces hyperphosphorylation of RIPK and necrosis associated gene expression *in vivo*

Differential gene expression analysis of RNA sequencing revealed PAX3-FOXO1 caused increases in expression of genes involved with adaptive and innate immunity like CD274 (PD-L1), MHCI, and DUSP family proteins **(Supplemental Data 4)**, and MHCI expression was further enhanced by viral infection **(Supplemental Data 5-6)**. In corroboration with the TNFa - NFκB signaling increase we observed, we probed infected cell lysates for phosphorylation of RIP1 kinase (RIPK1) during infection, which would indicate TNFa signaling for cell death. While RIPK1 was phosphorylated during infection with G47Δ and oRP3Fus (80kDa), only G47Δ infection led to hyperphosphorylation (110kDa) of RIPK1 **(Figure 7A; Supplemental Figure 8)**. Transfection of an ICP34.5-FLAG plasmid into RD cells followed by infection with G47Δ reduced RIPK1 hyperphosphorylation **(Figure 7B)**, suggesting ICP34.5 inhibits RIPK1 hyperphosphorylation which is linked to its pro-death activity. To determine whether this had any relevance to our *in vivo* studies, we performed qRT-PCR on tissue harvested from single-dose virus *in vivo* models. We observed a significant increase in necroptosis associated genes IL-1B, Nlrc4, Nlrp3, and TNFa in spleens from G47Δ but not oRP3Fus-treated mice **(Figure 7C)**. In contrast, livers, lungs, and tumors showed no difference between treatment groups, aside from higher MKLK in combination treated oRP3Fus from virus only treated tumors **(Figure 7D; Supplemental Figure 10)**. Additionally, we observed viral genomes in tumors treated with antiPD1 and oRP3Fus, and in off-target lung and liver for G47Δ **(Figure 7E),** suggesting our novel oRP3Fus may reduce cellular stresses to off target organs due to the inclusion of ICP34.5 and remain within the tumor microenvironment longer.

**Figure 7.**
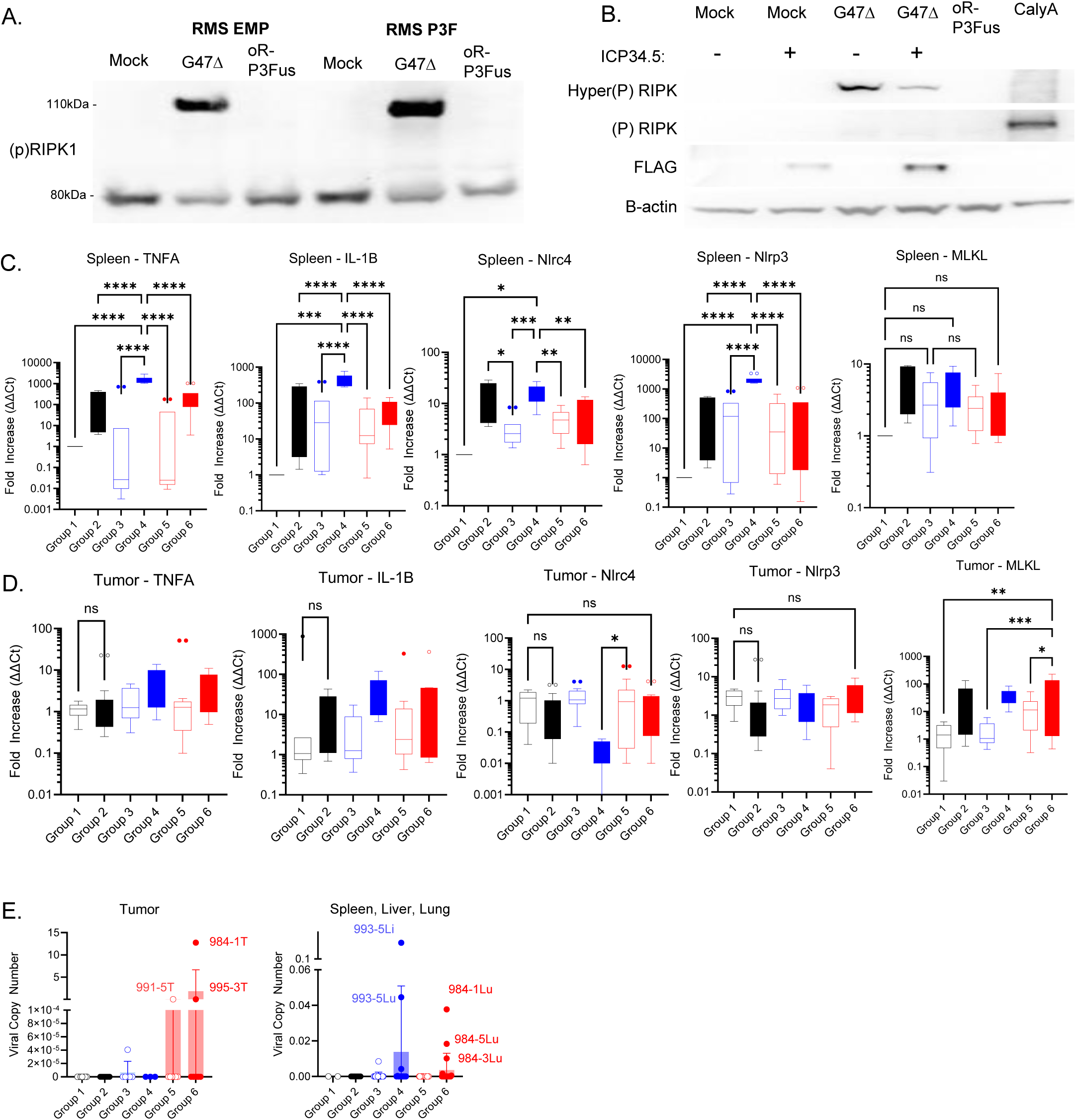
ICP34.5 expression reduces the RIPK1/necropotic like signaling *in vitro* and *in vivo*. (A) Western blot of RD:EMP infected with oHSV. (B) Western blot of RD cells transfected with or without ICP34.5-FLAG and infected with oHSV. (C) qRT-PCR of spleens from mice after treatment for M3-9-M:P3F tumors. (D) RTqPCR of M3-9-M tumors after treatment. (E) Viral copy number (VCN) measured from tumors (T) and lung (Lu), liver (Li), and spleen from treated mice. Group 1 – PBS; Group 2 – anti-PD1; Group 3 – G47Δ+PBS; Group 4 - G47Δ+PD1; Group 5 – oRP3Fus+PBS; Group 6 - oRP3Fus+PD1. Error bars represent Tukey box plot with median presented. One-way ANOVA with Tukey multiple comparison testing was performed (*p<0.05;**p<0.01; ***p<0.001; ****p<0.0001).

## DISCUSSION

The past decade has witnessed a revolution in immunotherapy for many adult cancers. However, immunotherapeutic approaches for treating pediatric sarcomas have lagged far behind. Notably, RMS is resistant to PD1 blockade both in human patients^9,10^ and in murine models^24^. This is due to a variety of factors, including an overall low tumor mutational burden (due to the driver nature of fusion oncogenes which frequently characterize pediatric tumors), as well as immunosuppressive characteristics within the tumor microenvironment of sarcomas in general. To overcome these barriers to effective T cell-based immunotherapy, we propose the use of oncolytic viruses which cause immune cell infiltration into these ‘cold’ tumors and do not carry the same risk of secondary cancer formation or long-term quality of life issues as conventional chemotherapy approaches.

Here we engineered a synthetic promoter that allows targeted expression of a gene of interest within PAX3-FOXO1-expressing tumor cells, while avoiding its expression in cells without this fusion oncogene. Our PRS1-Myogenin promoter system can be used to directly target PAX3-FOXO1 tumor cells and could be further adapted as a platform for future gene therapies or other targeted agents based on the presence or absence of a fusion oncogene. While we placed this synthetic promoter within an oHSV vector, the promoter cassette is small (455bp) meaning it could be used for other vectors such as adeno-associated viruses or in place of leader sequences for RNA vectors. Additionally, we discovered other incredibly potent promoters such as PRRX1-MYOD1 which could be used for high expression within normal and tumor tissues.

A serious weakness of OVs is their lack of long-term replication within the tumor site and their immunodominance over tumor antigens. This has dampened the excitement for their usage in immunotherapy, as checkpoint blockade is far simpler and usually more potent. However, pediatric RMS is largely resistant to immune checkpoint blockade. To increase the potency of G47Δ, which is approved for glioblastoma treatment, we restored ICP34.5 expression to allow better replication within PAX3-FOXO1 tumor cells. Restoration of ICP34.5 in trans or within the oRP3Fus (ICP34.5+) virus enhances replication kinetics and improves titer over parental G47Δ. At the same time, we show our novel virus has the same safety profile as G47Δ in skeletal muscle cell culture and after intramuscular injection into mice. While oHSVs typically are undetectable several days post injection, we see oRP3Fus genomes present within tumors up to 25 days post tumor implantation. Similar to other researchers who have restored ICP34.5 in oHSV to increase their tumor killing potency^25–27^, this study serves as a proof of principle that ICP34.5 can be utilized in herpes vectors to increases potency while maintaining safety. From this platform we intend to further modify oRP3Fus to enable better persistent tumor killing and immune activation.

For our *in vivo* studies we generated a human PAX3-FOXO1 expressing murine M3-9-M tumor line. These cells are easily passaged *in vitro* and reliably form orthotopic tumors in immunocompetent C57/Bl6 male mice. While there are other murine models to study fusion-positive RMS, they usually utilize human cells in immune-deficient mice, preventing the study of immune cell interaction. For immunocompetent models there is a report of the hU23674 line, generated by Myf6Cre driven Pax3-Foxo1 recombination in context of Tp53 loss of heterozygosity and homozygous loss of Arf^28^. However, to generate these mice requires an Agouti x C57/Bl6 cross, which means tumor cells may not faithfully implant in common laboratory mice strains. The M3-9-M tumor cell line was derived from a male C57/Bl6 mouse that transgenically expresses hepatocyte growth factor/scatter factor (HGF/SF) and is heterozygous for mutated Tp53. Studies which implant these cells into female C57/Bl6 mice observe immune clearance of cells even with mock treatment^24^, limiting their practicality to sex-restricted studies. Moving forward, there is a newly reported line generated by electroporation of human PAX3-FOXO1 and Cas9 targeting Tp53 and INK4a/Arf into C57/Bl6 skeletal muscle which forms alveolar RMS^29^. These genetically modified mouse aRMS tumors could be used for immunocompetent mouse studies. However, these were not available for our study, hence our reliance on the orthotopic implantation of the M3-9-M line in male mice.

We showed overexpression of human PAX3-FOXO1 made M3-9-M cells grow slightly faster than their parental line in culture but kept the same kinetics in forming flank and leg muscle tumors in male mice. Similar to the previous observations^24^, we observed female C57/Bl6 can reject M3-9-M implantation. In male mice we did not observe this distinction with only two failures to implant out of a hundred and nineteen implantations, hence our use of male mice to avoid confounding sex-related immunogenicity, as previously reported by others^24,30,31^. Anti-PD1 monotherapy was ineffective similar to previous reports, but when combined with virus extends mice survival, demonstrating that the influx of immune cells into the tumor after virus infection can improve receptiveness to checkpoint blockade. We additionally demonstrate splenocyte activation from treated mice when exposed to tumor and virus antigen suggesting this treatment is generating memory T cell responses, and splenic central memory. In concert with our TIL flow cytometry and IHC data, we conclude that multi-dose treatment with our oRP3Fus virus causes an influx of T cells within in the tumor, which are enhanced by anti-PD1 antibody and help clear tumor tissues. The presence of entities resembling tertiary lymphoid structures in our long-term survivor treated mice, while not being observed with mice treated with only anti-PD1 suggests that virus infection causes infiltration and establishes a niche for immune cell engagement with tumors. This study provides a foundation for expanding the role our vector can play in that engagement, with future directions to arm the virus with immune stimulating genes in place of the GFP reporter.

Our RNA sequencing shows that PAX3-FOXO1 overexpression causes human RD cells to increase gene expression involved with TNFa and NFκB signaling. In both RD:P3F and M3-9-M:P3F we observe an increase in NFκB signaling over empty controls indicating PAX3-FOXO1 expression may create a vulnerability to exploit within aRMS. The PAX3-FOXO1 upregulation of NFκB signaling could sensitize tumors to TNFa mediated killing by immune cells. Other studies have indicated disruption of the NFκB pathway in alveolar RMS could enhance treatments^32^. Studies using TAK1 or cIAP inhibitors to push TNFa signaling NFκB mediated cell death within tumors could be combined with OV to provide the final push needed to generate protective immunity against fusion positive RMS ^33,34^. In our study we observed that hyper-phosphorylation of RIPK1 was observed during G47Δ infection, but not during oRP3Fus infection. Complementation assays showed that ICP34.5 expression reduces this post-translational effect. While we do not understand how this occurs, it could relate to the dynamic of RIPK1 post-translational modification that occurs during TNFa mediated cell death signaling^34^. Whether G47Δ or oRP3Fus may synergize with TAK1 or cIAP inhibitors in the context of RMS remains to be studied, but combination of HSV1 and these inhibitors leads to rapid cell death in other cell lines^35^. This same principle could be applied to fusion positive RMS to increase treatment efficacy.

Parallel to the increased NFκB signaling, PAX3-FOXO1 has been shown to diminish JAK/STAT signaling and IFNγ expression^36^. In contrast, our RNA sequencing data indicates that PAX3-FOXO1 expression causes higher MHC-I expression during infection, and we observe phosphorylation of JAK kinase during infection via western blotting. This implies that oHSV treatment of RMS primes the TME to become more permissive for T cell infiltration and activity. From this foundation we can further arm the vector with more potent immune activators and/or test whether further antagonism of the TNFa-NFκB signaling axis can eradicate fusion positive RMS tumors.

In conclusion, our ICP34.5-armed oHSV (oRP3Fus) provides superior tumor killing activity against PAX3-FOXO1-positive alveolar rhabdomyosarcoma *in vitro*, and showed effective antitumor activity against such tumors *in vivo* when combined with anti-PD1 treatment. This synergy, whereby virus treatment sensitizes PAX3-FOXO1 positive rhabdomyosarcoma tumors to PD1 antibody treatment, represents an improvement over single-agent anti-PD1 therapy, which has historically been ineffective against these tumors. Future study of this vector may demonstrate exploitation of the enhanced NFκB signaling in PAX3-FOXO1-positive tumors to redirect immune cells to effectively destroy tumors. The presence of unique onco-transcription factors in these tumors presents a clear rationale for the design and engineering of viral vectors tailored to replicate specifically within PAX3-FOXO1 expressing alveolar rhabdomyosarcoma, as well as other fusion-driven malignancies.

## Acknowledgements

The JCCC Flow cytometry core and UCLA Translational Pathology Core Laboratory (TPCL) is supported by National Institutes of Health awards P30 CA016042 and 5P30 AI028697, and by the UCLA AIDS Institute, the David Geffen School of Medicine at UCLA, the UCLA Chancellor’s Office, and the UCLA Vice Chancellor’s Office of Research. We’d also like to thank Encarnacion Montecino-Rodriguez and Kenneth Dorshkind for use of their fluorescent microscope for plaque purification of recombinant HSV1.

## Funding

T.S.N. is supported by NIH grants R37 CA289813 and K08 CA241088, Hyundai Hope on Wheels, the Ruby Family Foundation, and Glimmers Childhood Cancer Foundation. C.W.P. is supported by NIH grant T32 AR059033. The content is solely the responsibility of the authors and does not necessarily represent the official views of the National Institutes of Health. Funding agencies had no role in the study design, collection, interpretation, or presentation of the data or writing of this report.

## Authors’ contributions

CP - conceptualization, supervision, investigation, resources, formal analysis, writing; MVC - investigation, formal analysis; AL – investigation; AD – investigation; CK – investigation; JN – investigation; AF – investigation, resources; MK – investigation; resources; SM, - investigation; LH – formal analysis; GD – formal analysis; MP – supervision; TSN – conceptualization, supervision, formal analysis, resources.

---

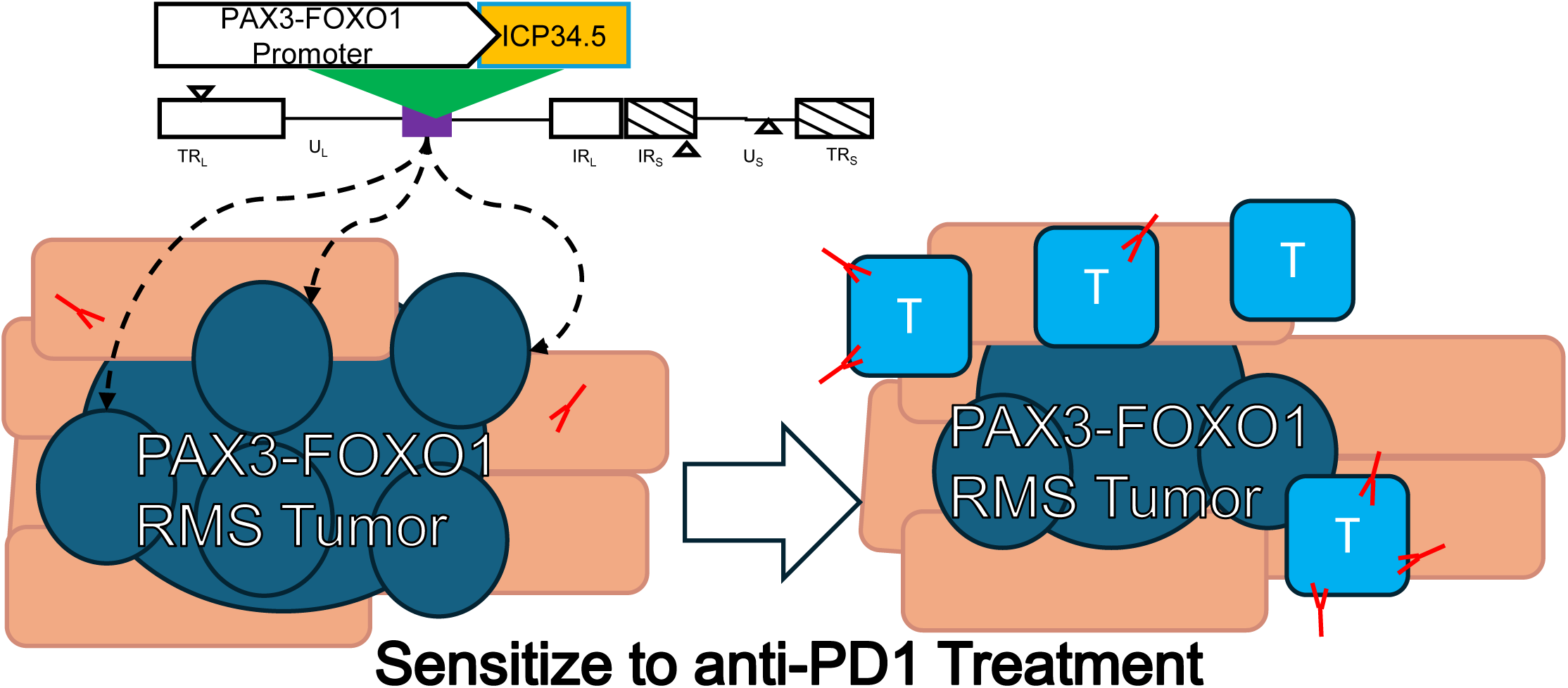

**Supplemental Figure 1.**
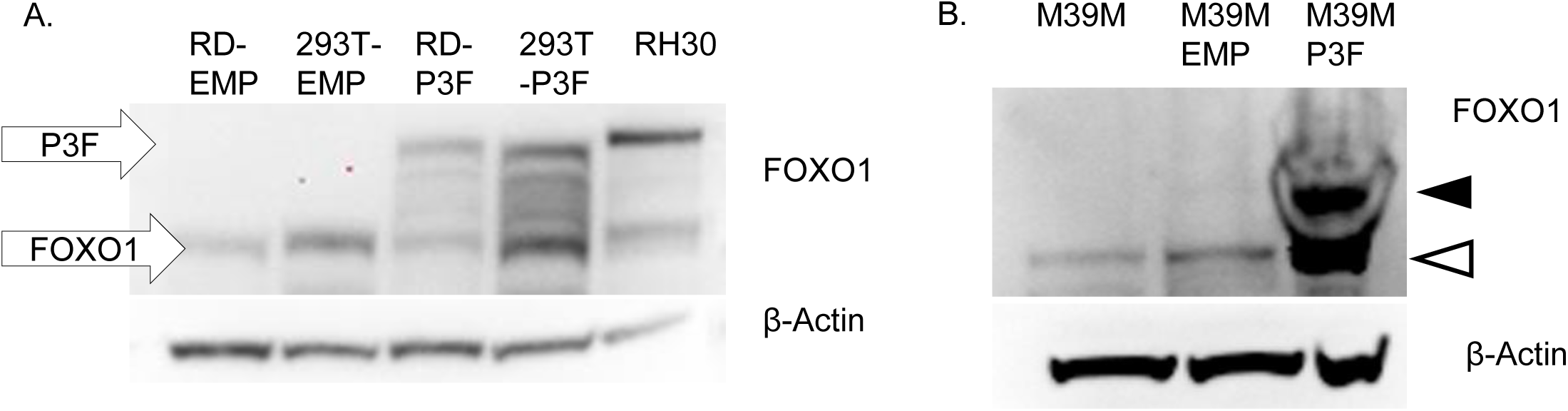
(A) Western blot of human cell lines transduced with PAX3-FOXO1 or EMPTY lentivirus. (B) Western of M3-9-M cells transduced to with PAX3-FOXO1 or EMPTY lentivirus. P3F (black arrow) and FOXO1 (white arrow).

**Supplemental Figure 2.**
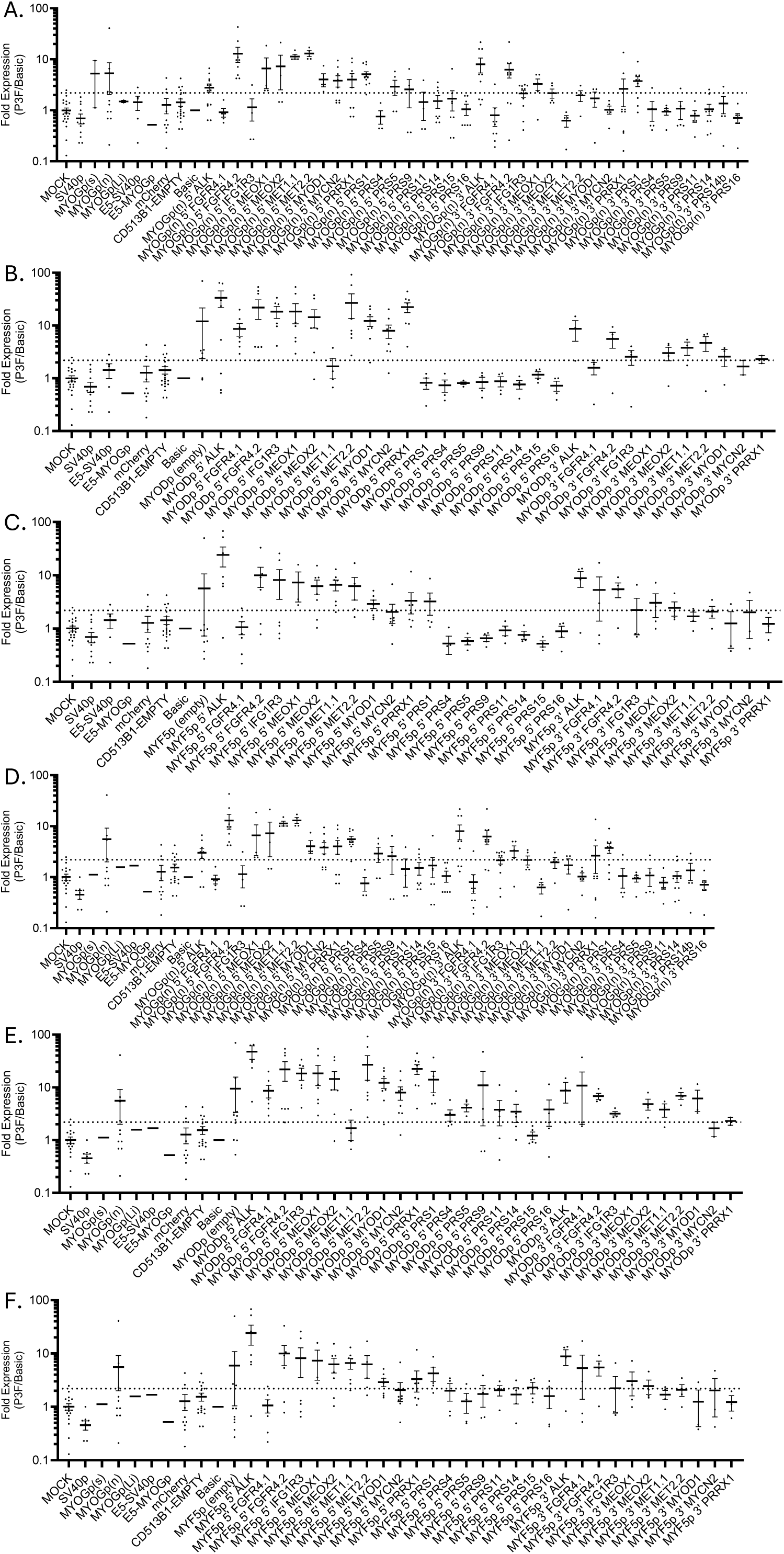
HEK293T (A,B,C) or RD (D,E,F) cell lines transduced with lentivirus to express PAX3-FOXO1 or empty control and transfected with pGL3-Luciferase constructs with promoter regions from the (A,D) myogenin (MYOG), (B,E) myogenic differentiation 1 (MYOD), or (C,F) myogenic Factor 5 (MYF5) genes supplemented with listed 5’ or 3’ enhancers. Ratio of expression vs Basic shown. Error bars represent SEM (n>5 for each construct shown).

**Supplemental Figure 3.**
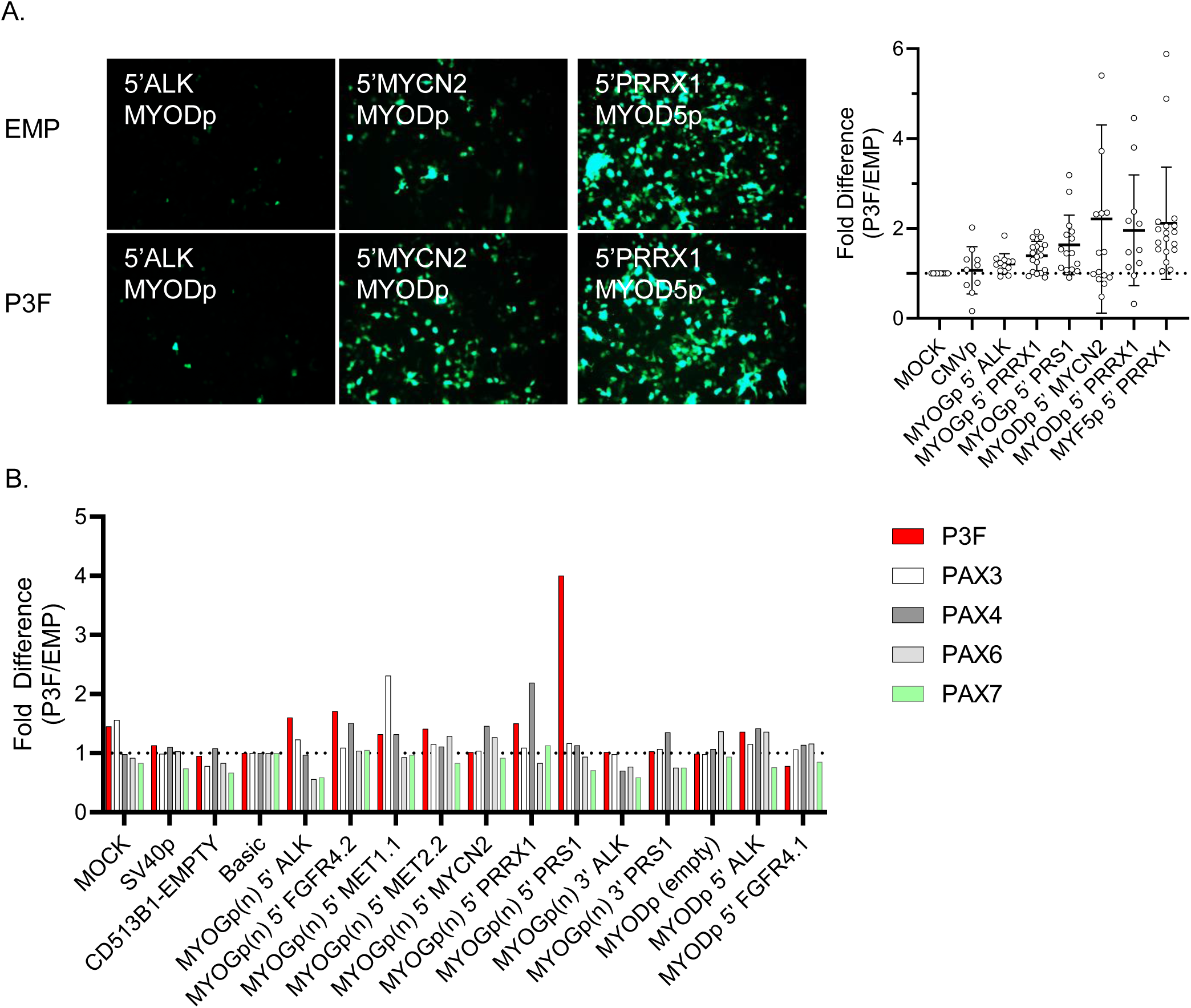
(A) GFP Reporter expression within HEK model. (B) Difference between candidate and basic luciferase reporter in HEK293T overexpressing PAX3, 4, 6, or 7.

**Supplemental Figure 4.**
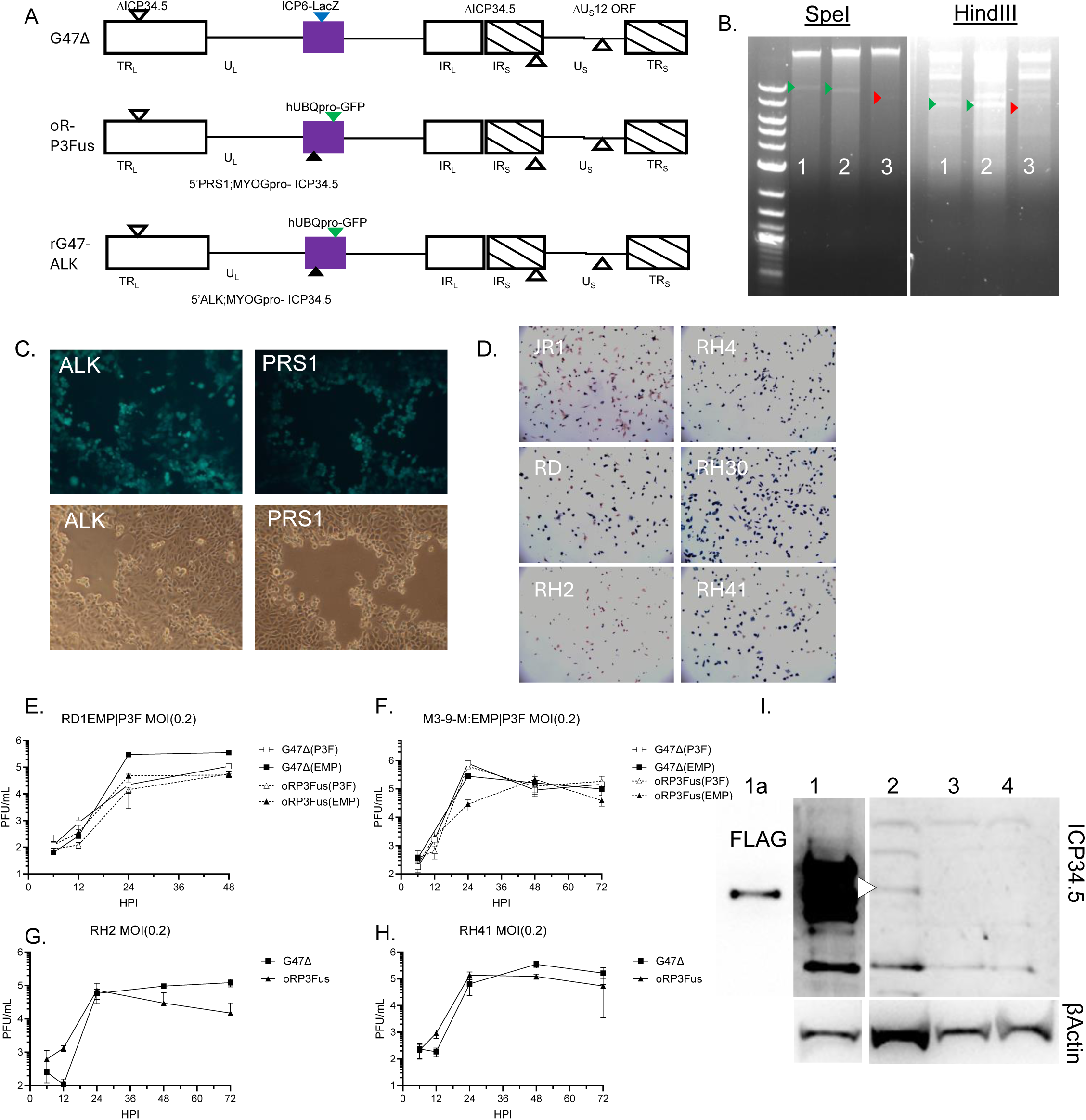
(A) Diagram of oHSVs TR - terminal repeat; IR - internal repeat; U – unique; L – long; S – short; ORF – open reading frame. (B) SpeI and HindIII digest of oHSV genomes. (C) Plaque visualization of rG47. (D) Representative photos of RMS lines infected with G47Δ were stained with Xgal solution followed by neutral. (E-H) Plaque assay of oHSV on human and murine RMS lines. (I) Western blot for ICP34.5, B-Actin, or FLAG. Lanes: 1. HEK293T transfected with FLAG tagged ICP34.5; 2. RD:P3F cells infected with oRP3Fus; 3. RD:EMP cells infected with oRP3Fus; 4. RD:EMP cells infected with G47Δ; 1a. blot was stripped and re-probed with anti-FLAG antibody. Arrowhead demarks the 32kDa ICP34.5 band. Blot has irrelevant wells removed for clarity.

**Supplemental Figure 5.**
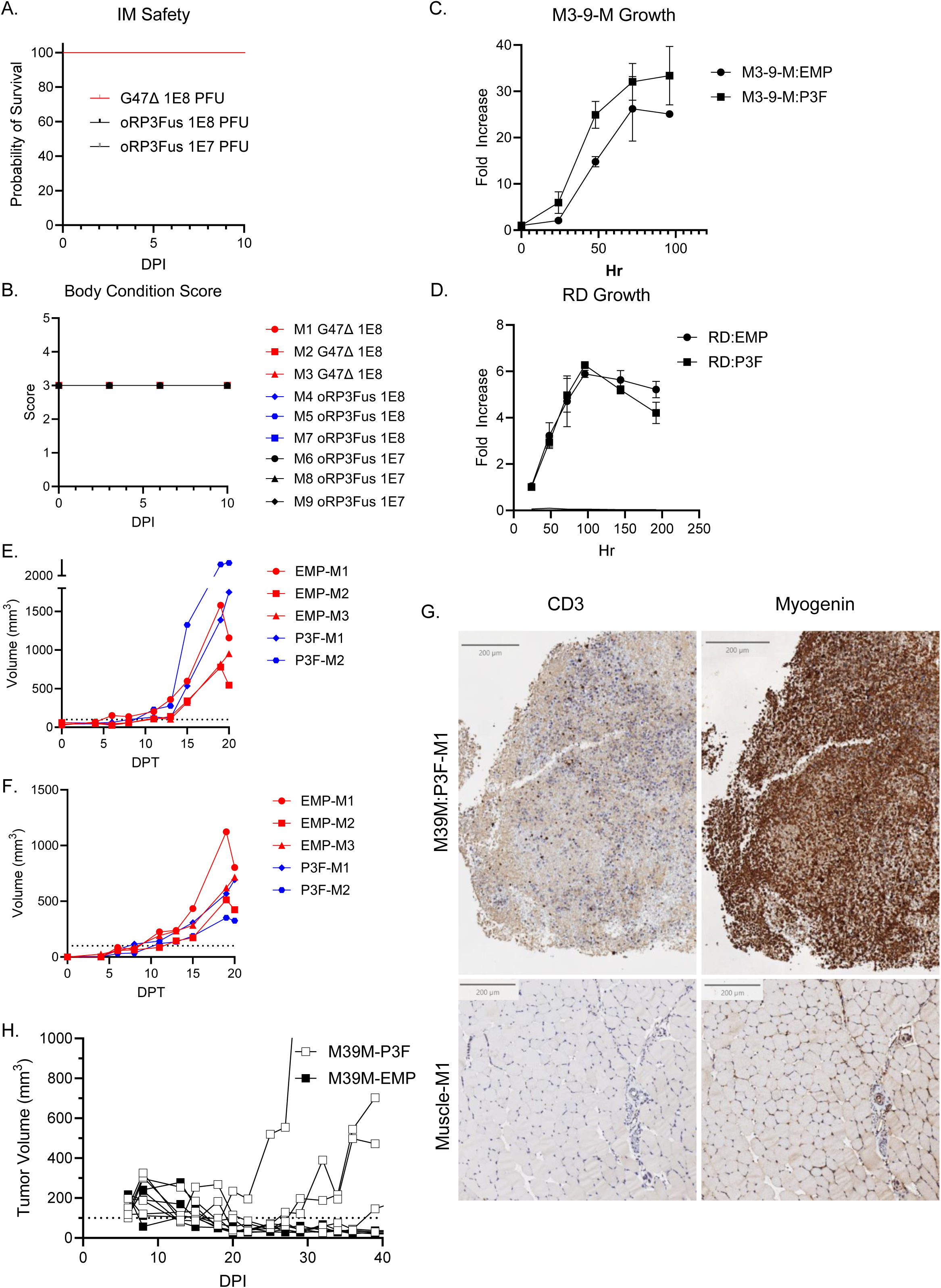
(A) Survival safety and (B) body condition score after oHSV IM injection in male C57/Bl6J mice (n=3 per group). Growth curves of (C) M3-9-M or (D) RD cell lines transduced with PAX3-FOXO1 (P3F) or Empty (EMP) lentivirus (n=3). Growth curves of M3-9-M:EMP|P3F tumors after implantation into male C57/Bl6J (E) gastrocnemius or (F) flank mice (n=3, EMP; n=2 P3F). (G) Representative images from CD3 and myogenin staining on M3-9-M:P3F tumor and normal muscle (Hematoxylin and DAB staining. (H) Implantation of M3-9-M:P3F or M3-9-M:EMP cells into gastrocnemius muscle of female C57/Bl6 mice (n=5).

**Supplemental Figure 6.**
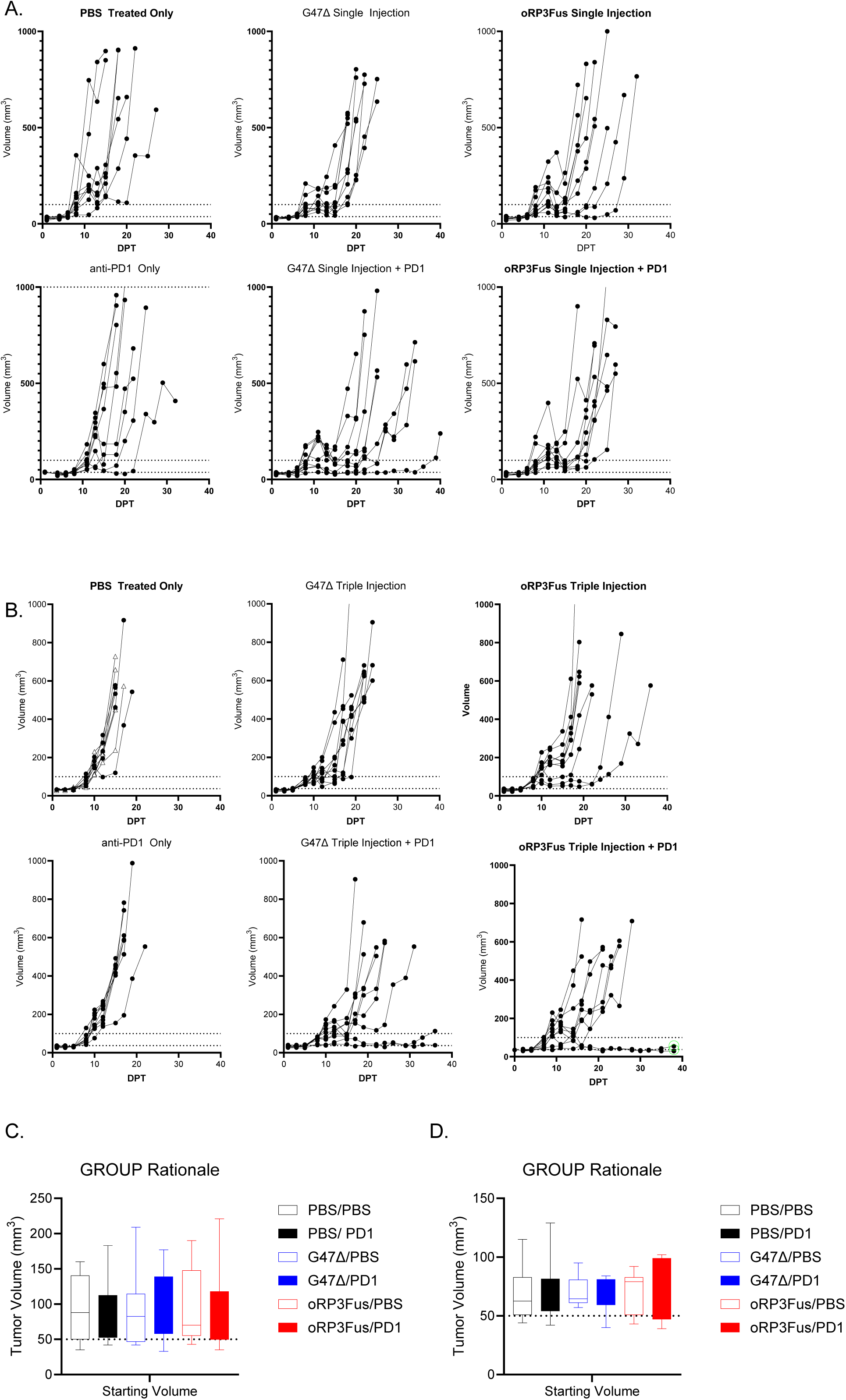
Oncolytic HSV G47Δ and oRP3Fus combine with anti-PD1 therapy to improve survival in immunocompetent RMS. (A) Tumor growth curves for mice treated with single virus injection +/- anti-PD1 antibody. (B) Tumor growth curves for mice treated with triple virus injection +/- anti-PD1 antibody. Green circles indicate mice considered responders after analysis. Starting tumor volumes of each for each treatment group for (C) single virus or (D) triple virus injected cohorts. One-way ANOVA using Holm-Šídák multiple comparison testing was used to compare group means.

**Supplemental Figure 7.**
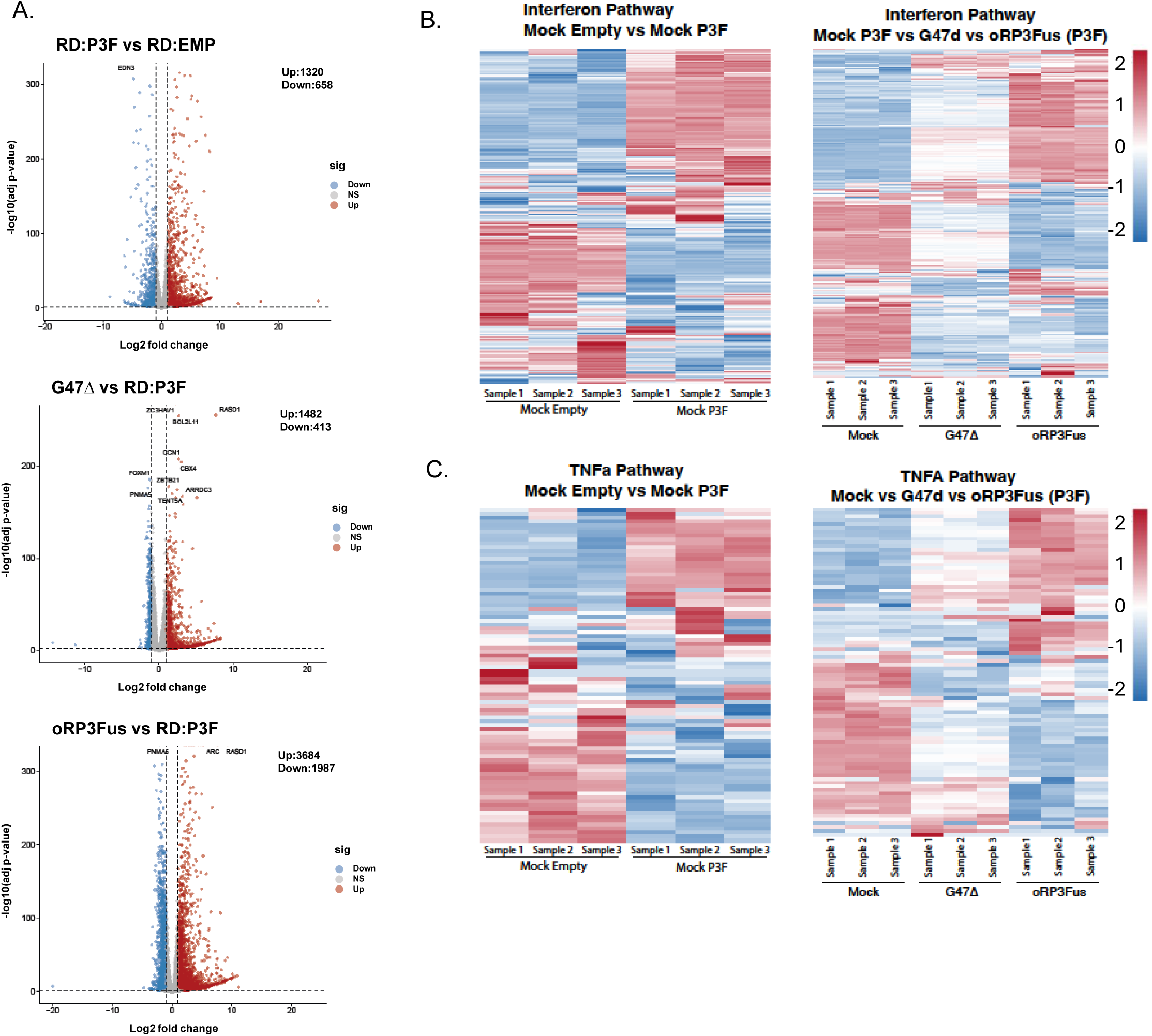
(A) Volcano plots of DEGs between mock infected RD:P3F vs RD:EMP; G47Δ infected RD:P3F; and oRP3Fus infected RD:P3F. Heat maps measuring DEG relevant to (B) interferon signaling pathways or (C) TNFA signaling pathways. Gene lists for heat maps in supplemental data.

**Supplemental Figure 8.**
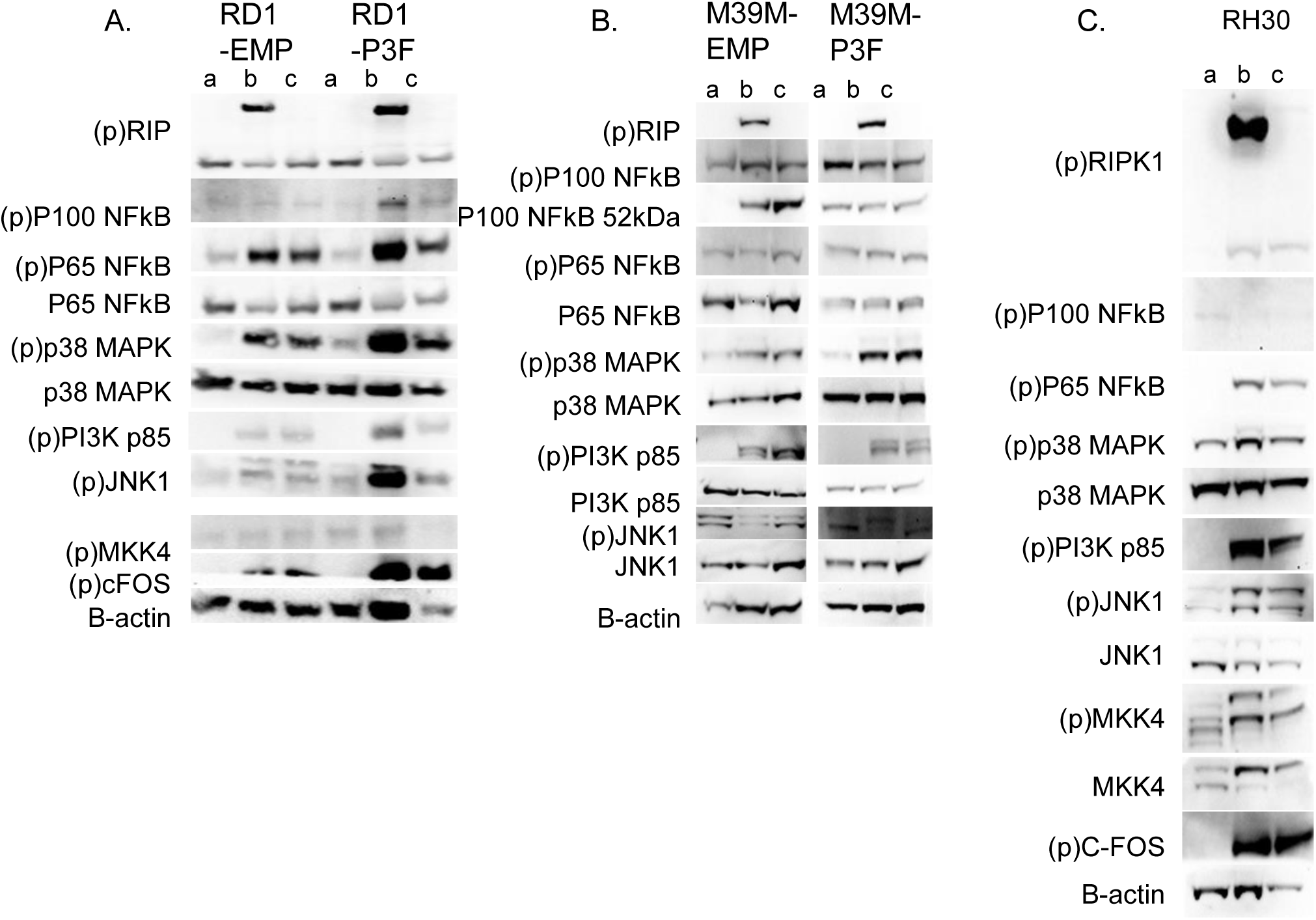
Western blotting of NFκB signaling cascade in (A) RD, (B) M3-9-M, and (C) RH30(P3F+) cell lines infected with oHSV at MOI(2) 6-hours post infection. Sample infection conditions 6hpi: a – Mock; b – G47Δ; c – oRP3Fus.

**Supplemental Figure 9.**
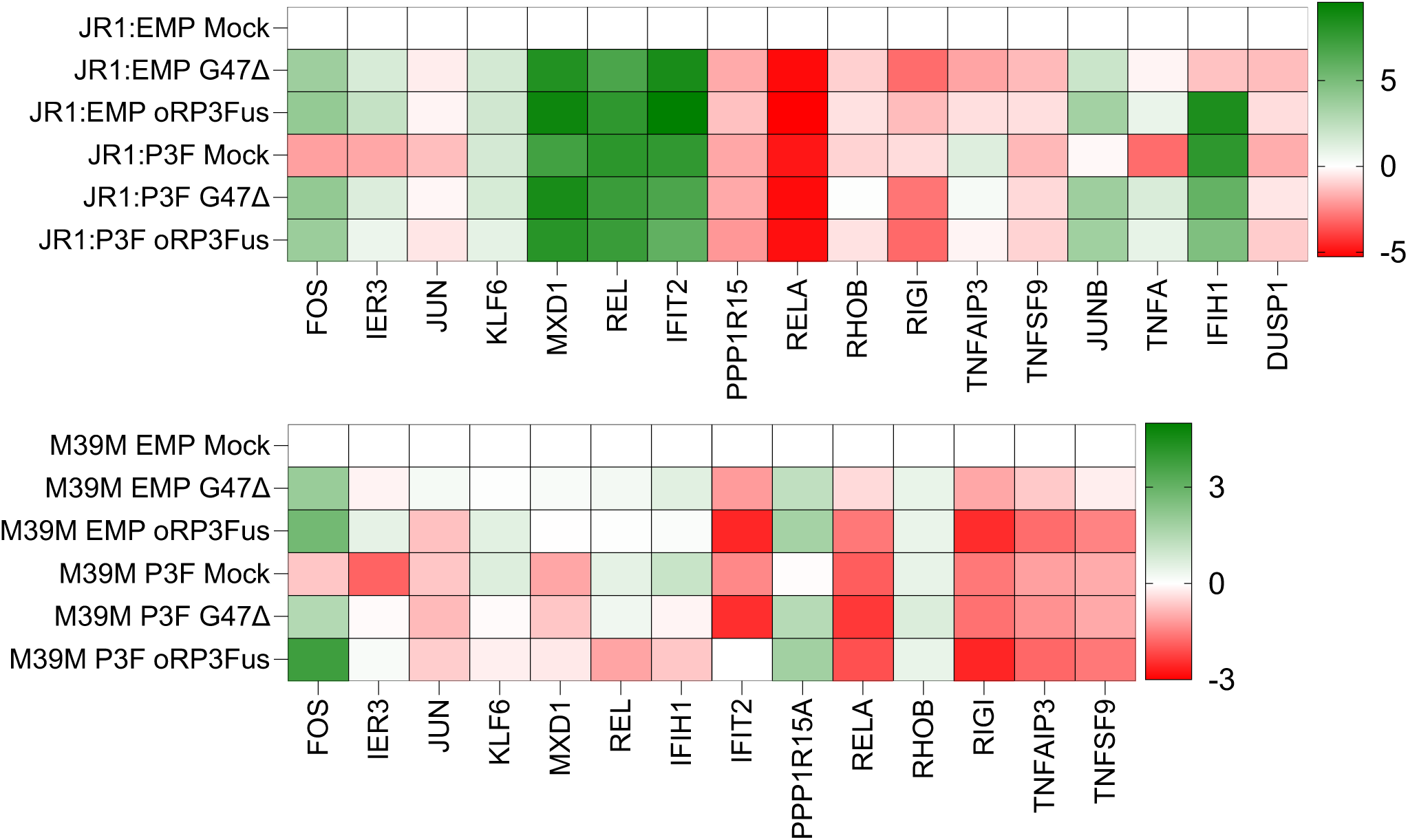
(A) qRT-PCR of M3-9-M and JR1 cells 6-hour post infection at an MOI(2) as compared to Mock empty cells.

**Supplemental Figure 10.**
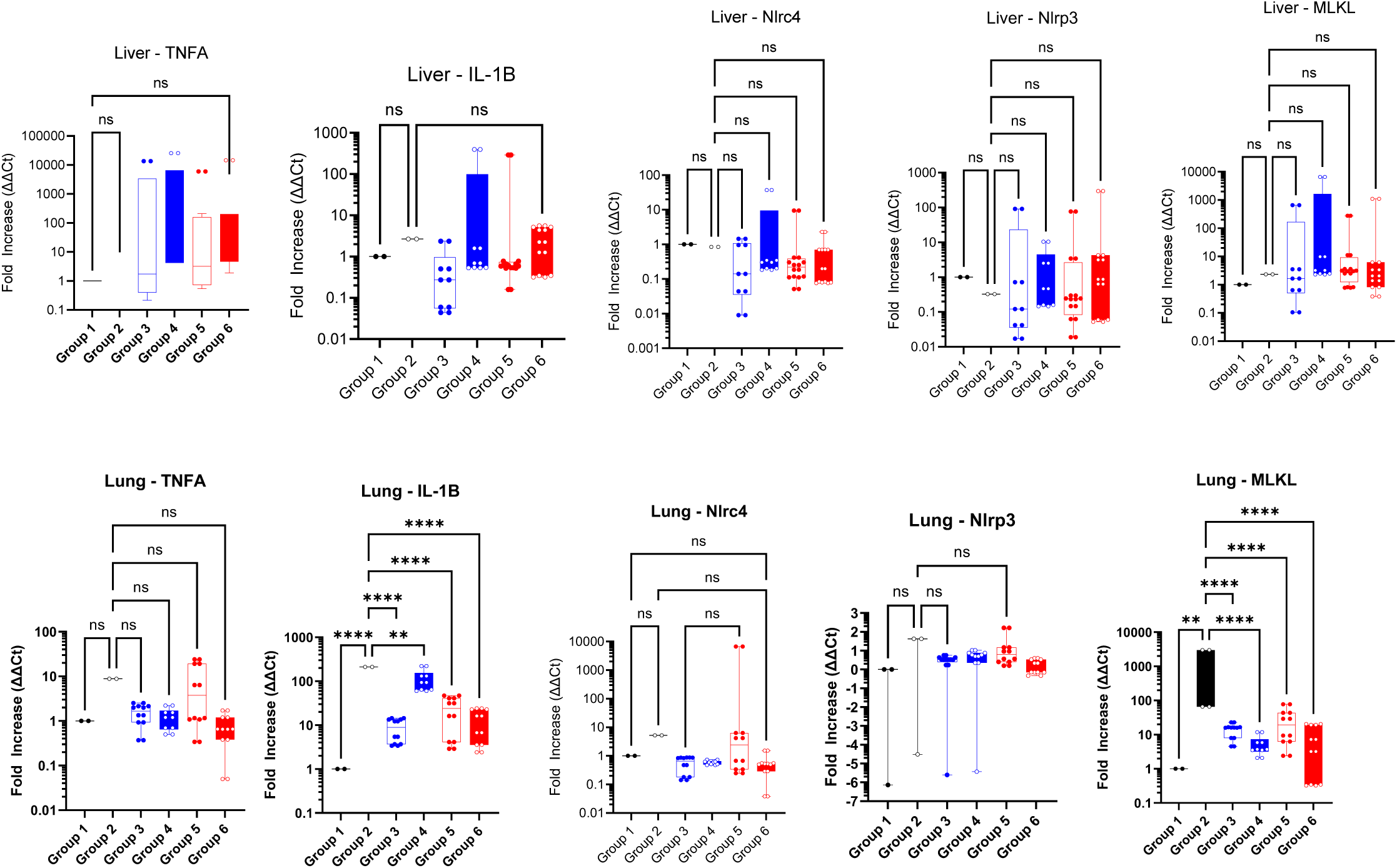
ICP34.5 expression reduces the RIPK1/necropotic like signaling in vivo. RTqPCR of M3-9-M implanted mouse liver and lungs after treatment with oHSV. Group 1 – PBS; Group 2 – anti-PD1; Group 3 – G47Δ+PBS; Group 4 - G47Δ+PD1; Group 5 – oRP3Fus+PBS; Group 6 - oRP3Fus+PD1.

**Supplemental Table 1.**
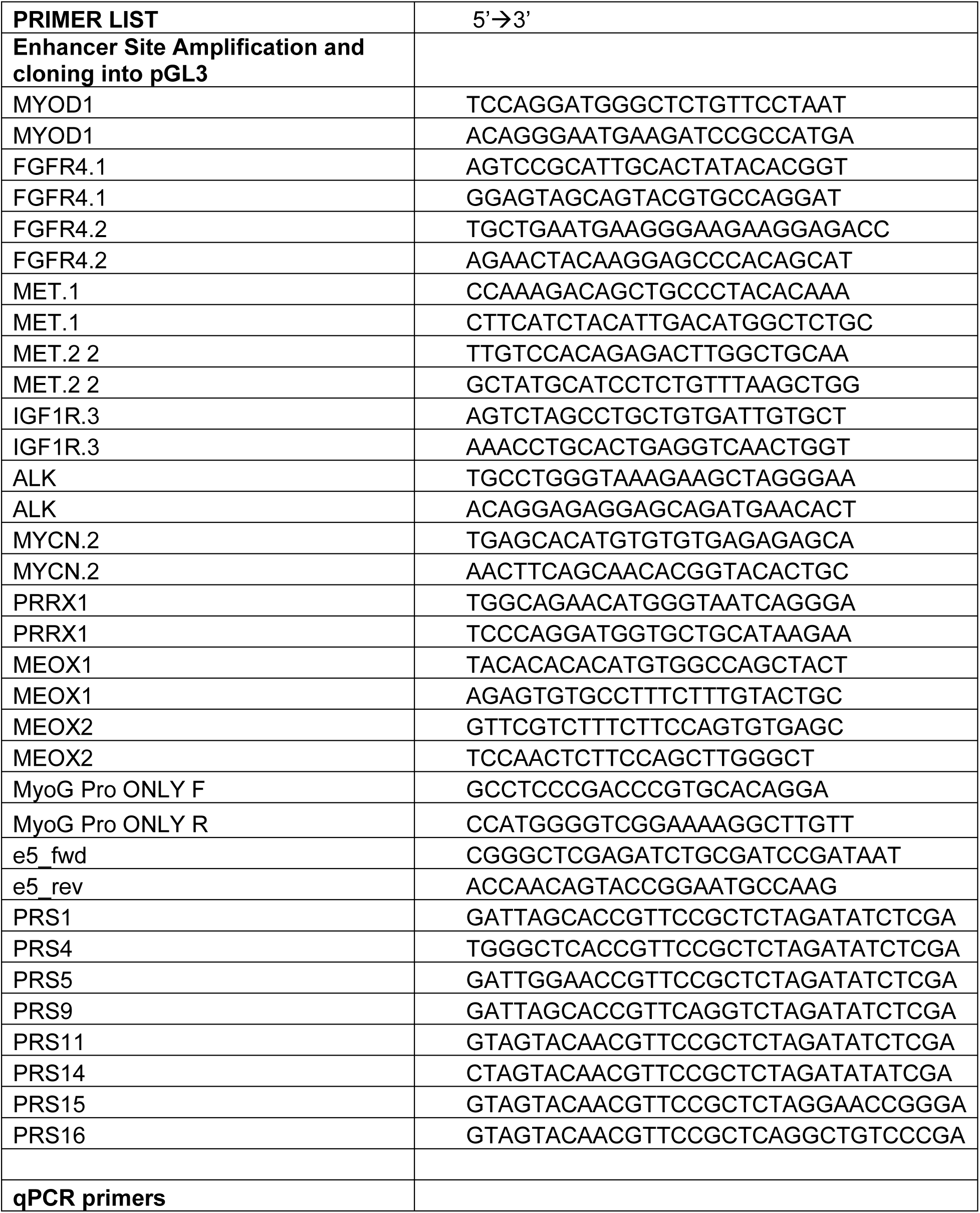
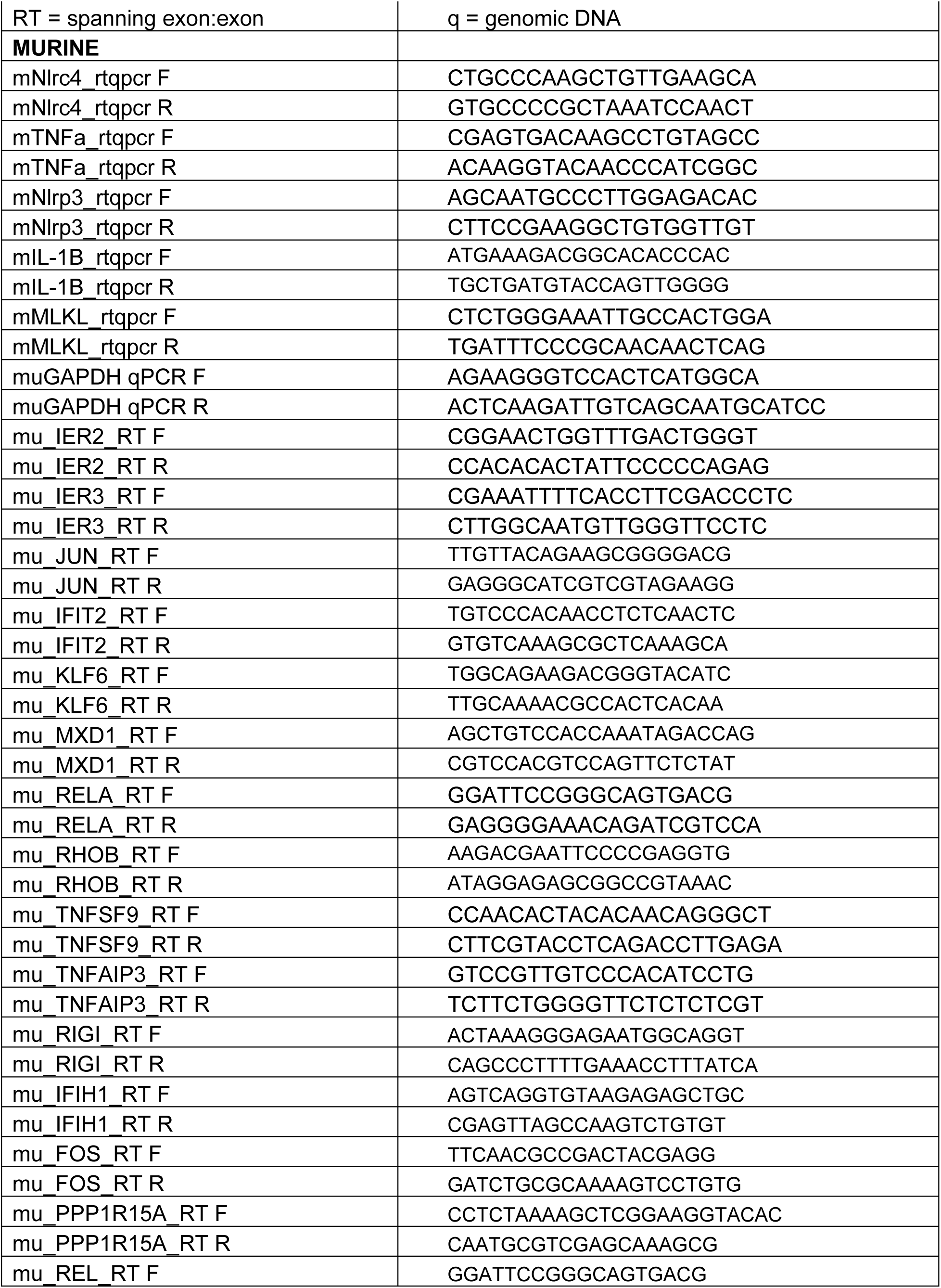
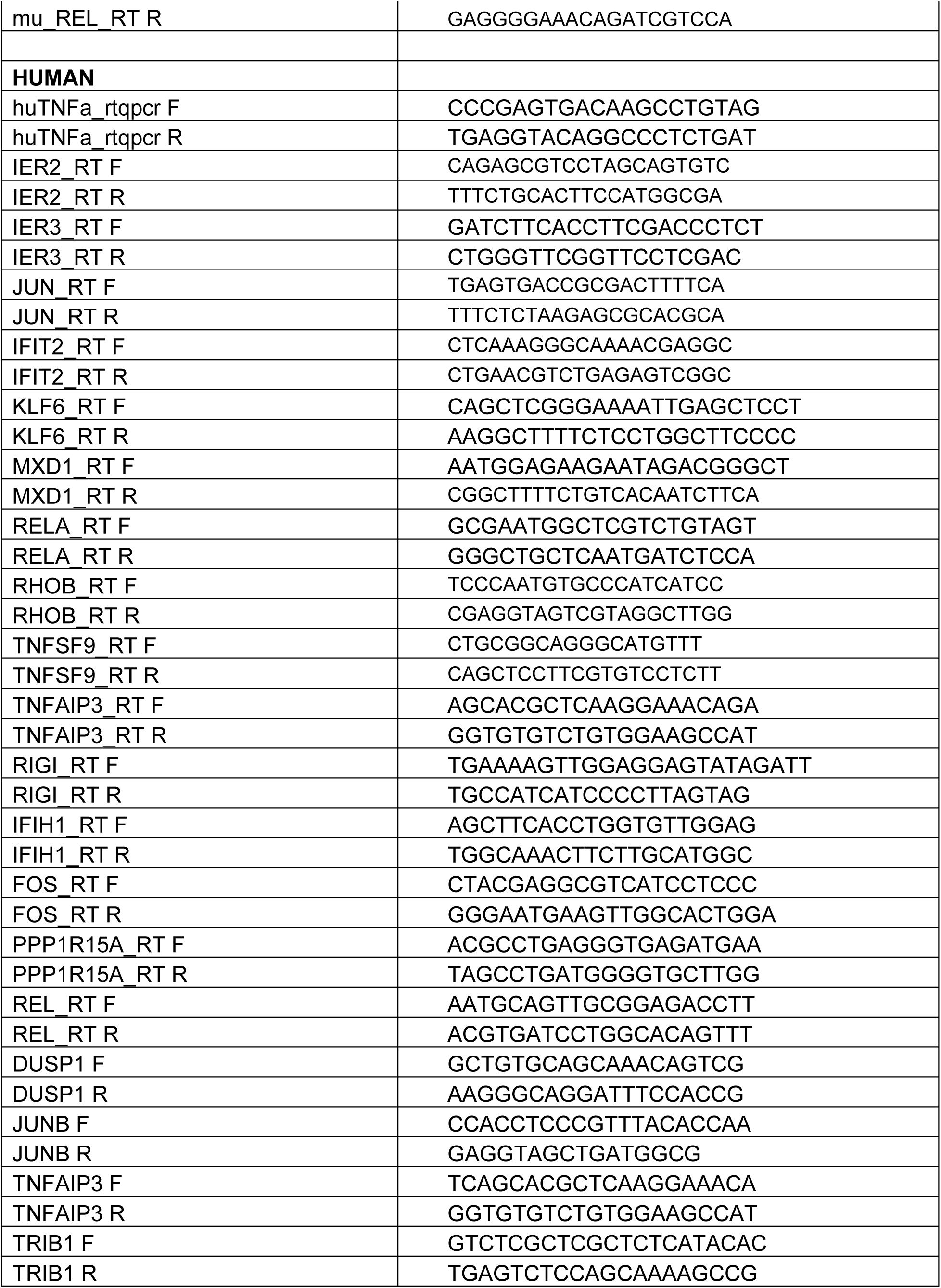
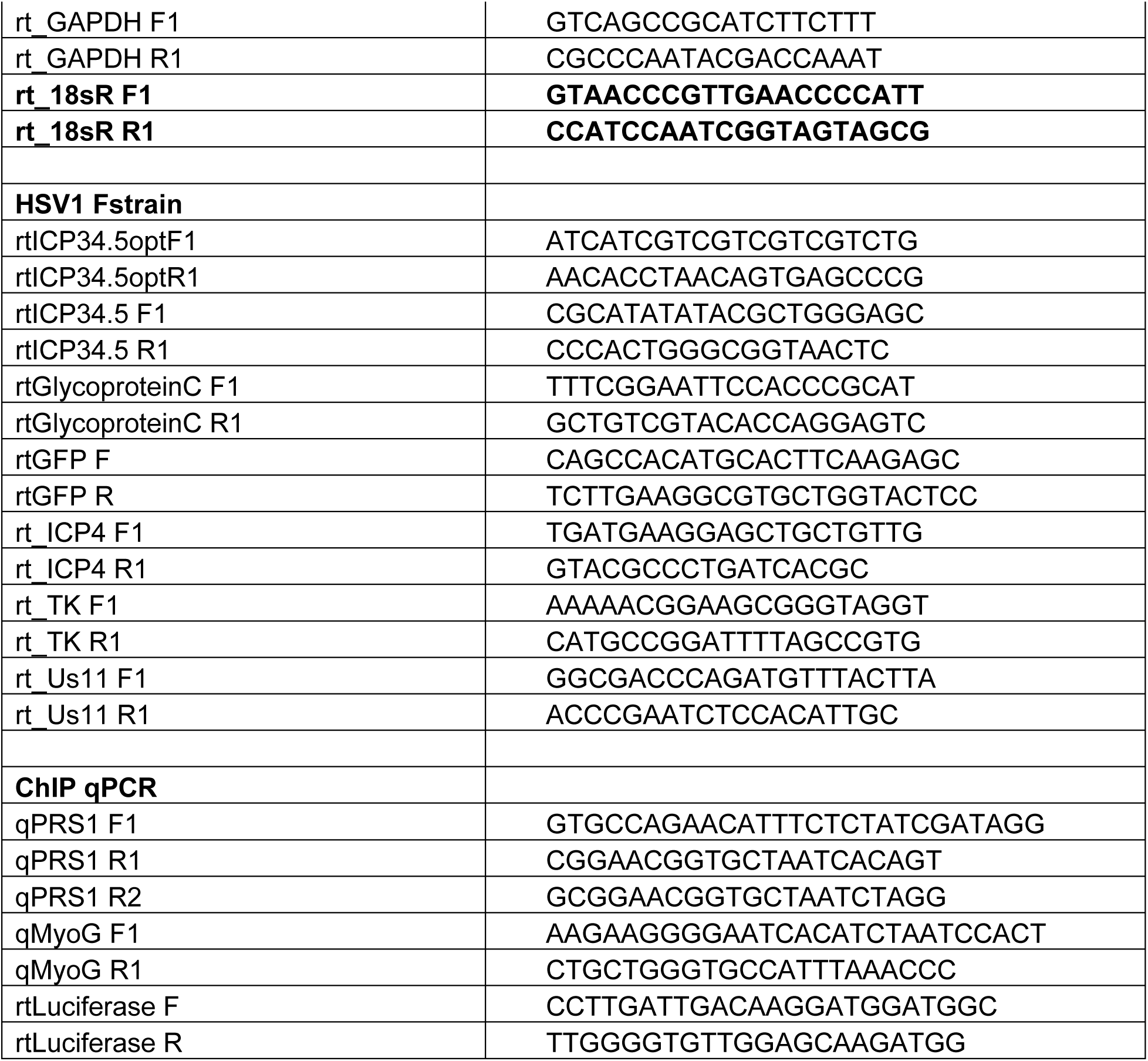
PCR primers used to generate enhancer/promoter sequences; measure quantitative PCR; or clone constructs described in this study.

**Supplemental Data 1.** Flow cytometry gating strategy. Side Scatter (SSC) vs Forward Scatter (FSC) for leukocyte gate, followed by gating for single cells via FSC-A vs FSC-H, followed by live dead stain via 7AA or Fixable/Yellow (Positive indicates dead), followed by CD45 vs FSC, followed by CD3 vs FSC, followed by CD4 vs CD8. From CD4+/CD8- or CD4-/CD8+ gates we further gated for PD1.

**Supplemental Data 2.** Genes associated with NFκB, TNFa, and Interferon stimulated pathways. Heatmaps order for Figure 6B, 6C, supplemental figure 7Bi, 7Bii, 7Ci, 7Cii.

**Supplemental Data 3.** FASTA sequence of the ICP6 insertion site of G47Δ and oRP3Fus.

**Supplemental Data 4.** DEGs when comparing Mock infected RD:P3F cells vs Mock infected RD:EMP cells. Genes shown were shown to be significantly different in expression between the two groups (padj<0.05; log2fold change>1|<-1).

**Supplemental Data 5.** DEGs when comparing G47Δ infected RD:P3F cells vs Mock infected RD:P3F cells. Genes shown were shown to be significantly different in expression between the two groups (padj<0.05; log2fold change>1|<-1).

**Supplemental Data 6.** DEGs when comparing oRP3Fus infected RD:P3F cells vs Mock infected RD:P3F cells. Genes shown were shown to be significantly different in expression between the two groups (padj<0.05; log2fold change>1|<-1).

## Supplemental Methods

### Western Blot

Protein content was acquired using RIPA buffer supplemented with cOmplete proteinase and phosphatase inhibitors (Roche). Protein was boiled in cracking buffer and loaded onto denaturing Nu-PAGE gels (Thermo). Semi-dry transfer onto PVDF membrane was performed and blotted using antibodies in 5% skim milk or BSA in 1x TBST. Secondary antibody-HRP conjugates were used to image membranes using Piere ECL substrate. Blots were stripped using stripping buffer for re-blot (BP-98-500ml).

### Antibodies

Flow cytometry include: CD45 clone 30-F11 IgG2c (BioLegend 103137); CD3 clone 17A2 IgG2b K (BioLegend 100203); CD4 clone GK1.5 IgG2A (BioLegend 553032); PD1 clone 29F.1A12 IgG2A K (BioLegend 568616); huPD-L1 (BioLegend 329705); muPD-L1 (BioLegend 155404). Western blot include: CST: B-Actin(#3700), NFkB Pp65/total (3033,8242), Pp100/Total (4810/4882), MAPK Pp38/Total (4511/9212), JNK1 P/Total (4668/3708), PI3K P/Total (17366/4257), MKK4 P/Total (4511/9152), cFOS P/Total (5348/2250), RIPK1 P/Total(58274/4926), FOXO1 (2880), and FLAG DYKDDDDK (2368). We would like to thank Julio Sanchez for generously providing the anti-PD-L1 antibodies (329705, 155404).

### Reporter Assays

pGL3 firefly luciferase was detected using the Pierce Firefly Luciferase Glow Assay Kit (Thermo 16176). Cells were seeded at 2 x10^4^ HEK293T or 3 x10^4^ RMS cells/well within a Nunc 96-Well Optical-Bottom Microplate (Thermo 165305). The following day they were transfected with 200 or 300ng of pGL3 plasmid using JetPRIME transfection reagent. Two days post transfection cells were collected, lysed, and assayed for luciferase activity, as per Glow Assay Kit Instructions via Varioskan plate reader (Thermo). GFP expression was also determined by Varioskan plate reader. Harvested splenocytes were cultured with M3-9-M:PAX3-FOXO1 or M3-9-M:Empty tumor cells at a 1:1 ratio for 20h in a U-bottom well plate before harvesting supernatants to test via mouse IFNγ ELISA (BioTechne MIF00) following manufacturer instructions.

### Chromatin Immunoprecipitation

15 x10^6^ PAX3-FOXO1 or Empty transduced HEK293T cells were seeded in a 15cm plate, transfected with relevant pGL3 plasmid and harvested after two days. Protein:DNA was crosslinked via addition of formaldehyde to a final 1% concentration, light mixing for 10min, followed by addition of glycine to a final concentration of 125mM for five minutes. DNA:Protein complexes were isolated as described^23^. Before immunoprecipitation (IP) 50uL of chromatin was kept as input controls. Remaining DNA was pre-cleared using Dynabeads (Thermo 10001D), samples were separated by volume, and incubated with anti-FLAG (CST 14793), PAX3-FOXO1 (NHGRI PaxF 160866), or FOXO1 (CST 2880) antibody. IP was performed with Dynabeads, followed by washes in low, high, and LiCl salt buffers. 20% of bead eluates were boiled for protein analysis. The remaining fraction was eluted in elution buffer (1%SDS, 100mM NaHCO3). For analysis, percent input based on volume was used. Threshold Cts were normalized to input via: ΔCt = Ct [IP] – (Ct [input] – DF); where DF was 10-fold dilution (3.32). Next yield was calculated: Yield (%) = (primer efficiency)–ΔCt x 100%. Fold enrichment was determined by normalizing to the FLAG (IgG) control: ΔΔCt = ΔCt [IP] – ΔCt [IgG]; (1+PE)^-ΔΔCt^.

### Xgal Staining

Virus plaque assays were performed by plating Vero cells at 1.2 x10^6^ cells/6-well plate overnight, followed by incubation with virus in PBS for 1.5h. Virus supernatant was then removed and cells cultured in D10+ anti-HSV1 IgG for 3 days. Cells were fixed with 2% paraformaldehyde/0.2% glutaraldehyde, incubated with X-gal stain to observe LacZ expression, and counterstained with Neutral Red. Infection of tumor or skeletal muscle lines was performed similarly, using multiplicity of infection (MOI) of 0.1, 1, or 2 based on the assay. After fixation, cells were treated with a X-gal solution (0.4mg/mL X-gal, 5mM Potassium Ferricyanide, 5mM Potassium Ferrocyanide, 2mM MgCl_2_) for 1h at 37C. Wells were rinsed with water and counterstained with Neutral red solution (0.01g/mL Neutral red dye, 0.125% MeOH) for 10min and dried.

### Viability Testing

Cell viability was tested using Alamar Blue (Invitrogen DAL1100) as manufacture specifications direct. Cells were incubated for 2h before analysis on SkanIT plate reader. Fluorescence - ex560nm | em590nm; Absorbance - 570nm | 600nm reference were used to measure growth in comparison to mock or empty cells.

### RNA Sequencing

Runs were performed on NovaSeq X+ Series 25B flow cell for 300-cycle (2 × 150 bp) read length. Raw sequencing reads were assessed for quality with FastQC v0.12.1. Adapter sequences and low-quality bases were removed using Trimmomatic v0.39 with the following parameters: LEADING:20 TRAILING:20 SLIDINGWINDOW:5:20 MINLEN:90. Trimmed reads were aligned with STAR v2.7.9a to the GRCh38 assembly reference human genome. Gene-level counts were generated during alignment using STAR’s --quantMode GeneCounts option. Prior to differential expression analysis, lowly expressed genes were filtered out in R by retaining only genes with more than 10 reads across all samples (dds <-dds[rowSums(counts(dds)) > 10,]). Significantly differentially expressed genes (DEGs) were defined as those with an adjusted p-value (padj) < 0.05 and an absolute log₂ fold change (|log₂FC|) ≥ 1, corresponding to at least a two-fold change in expression (log₂FC ≥ 1 for upregulated genes, log₂FC ≤ –1 for downregulated genes). Gene Ontology (GO) enrichment was performed with the clusterProfiler v4.14.6 R package using the gseGO function with org.Hs.eg.db, ontology set to *ALL.* Gene set enrichment analysis was conducted with the GSEA desktop application v4.3.1 (Broad Institute). Input included normalized expression counts from DESeq2 and phenotype class files. The software ranked genes using the Signal2Noise statistic and enrichment was tested against the Hallmark gene sets (MSigDB). Heat maps were generated using gene lists sourced from gsea-msigdb.org and are listed in

---

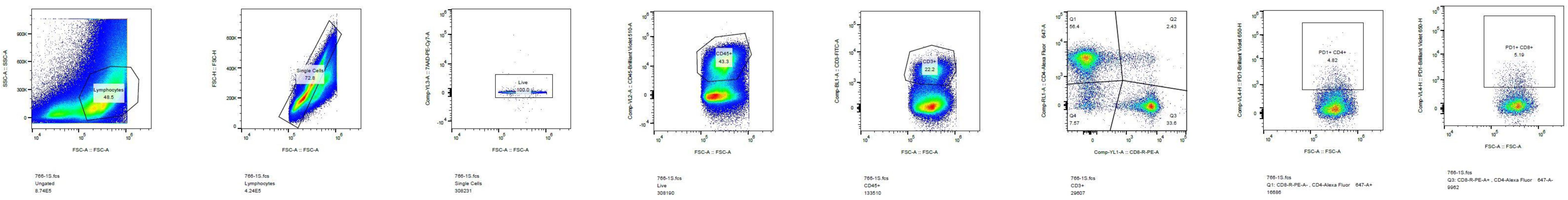

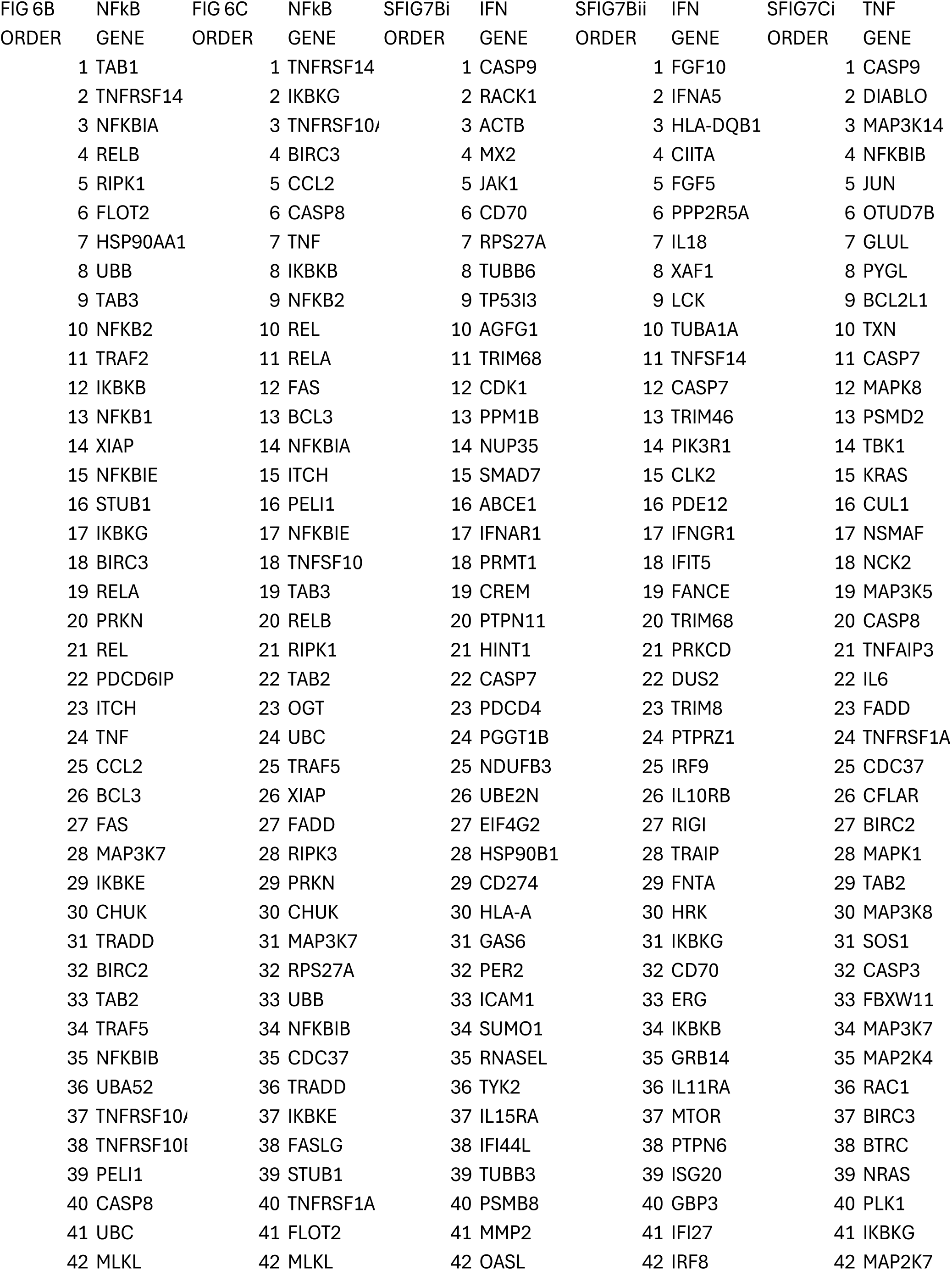
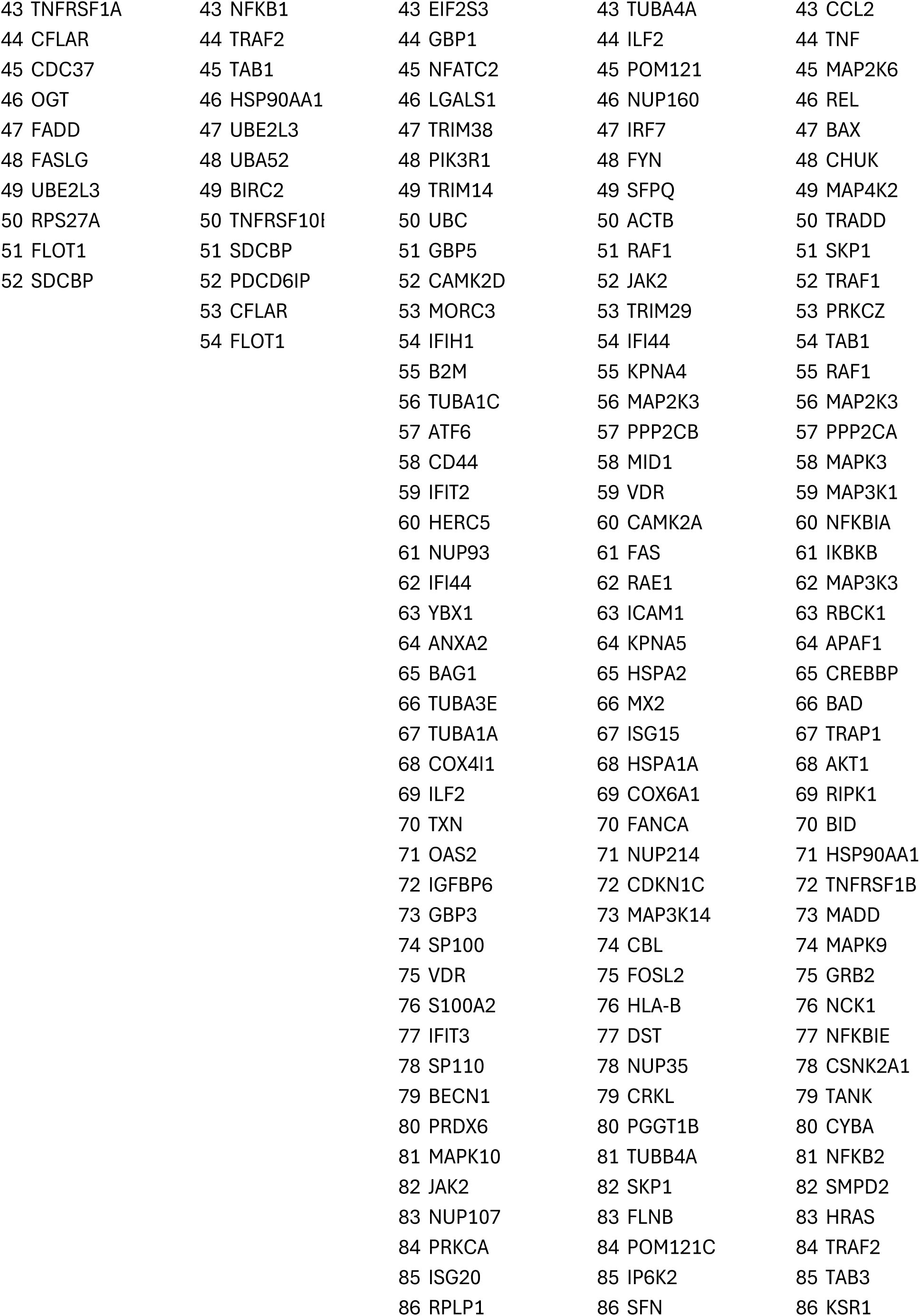
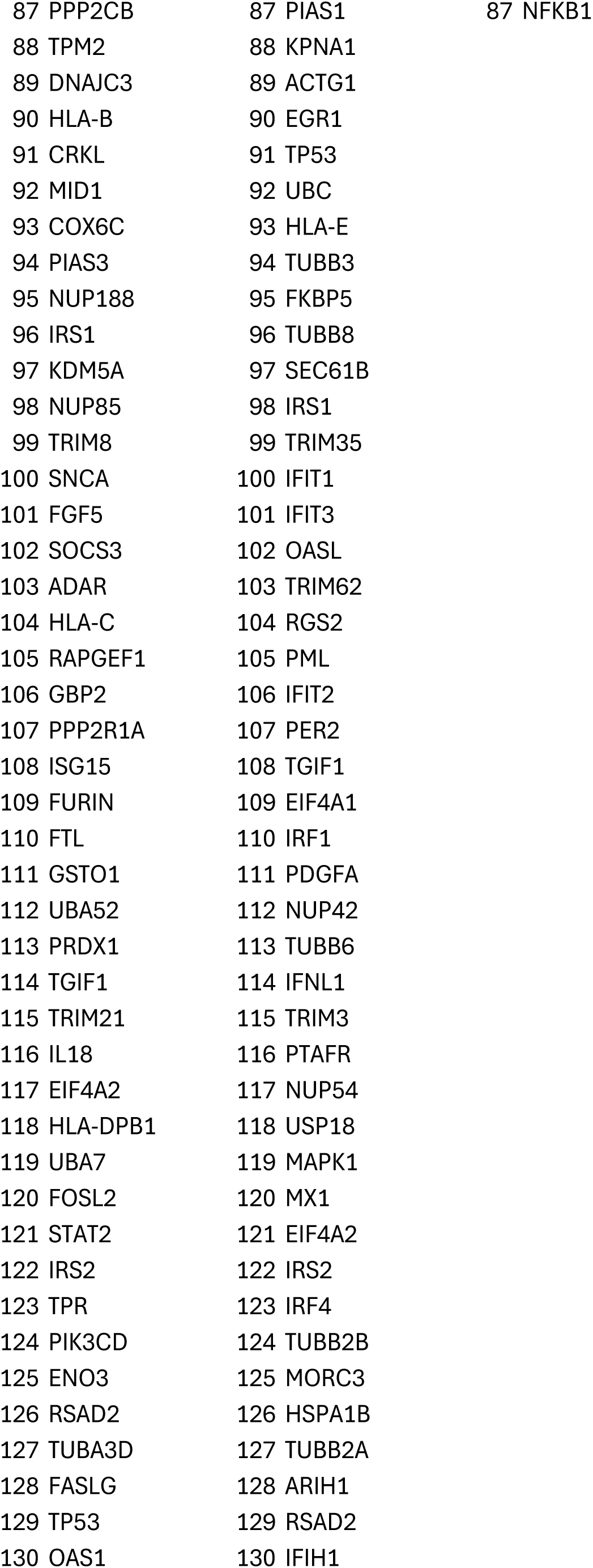
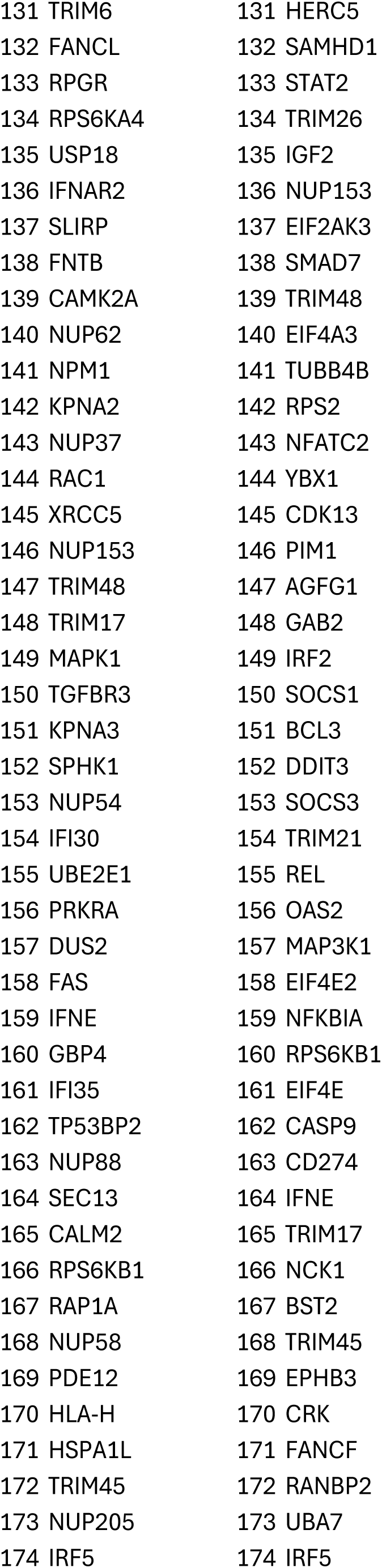
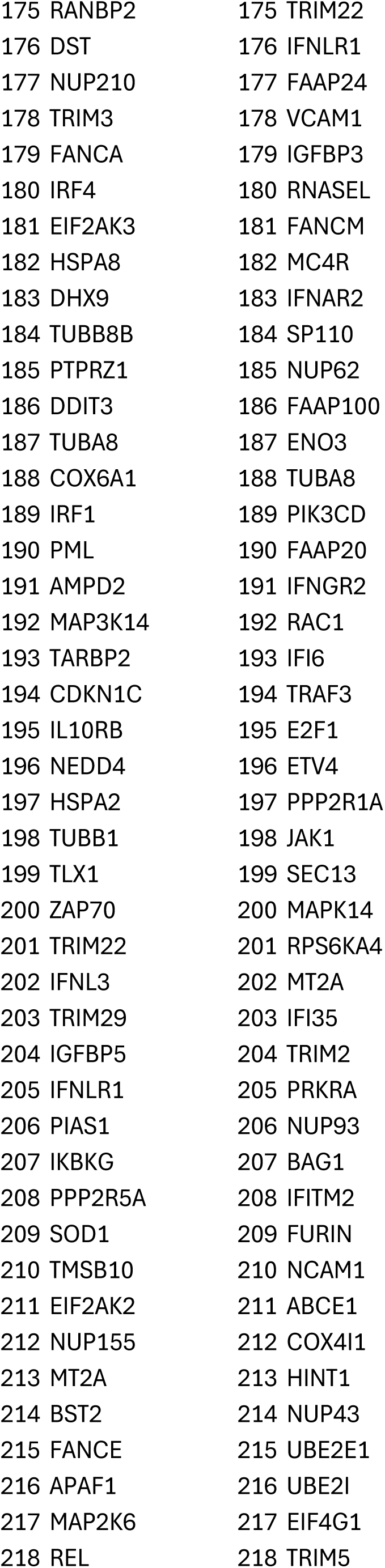
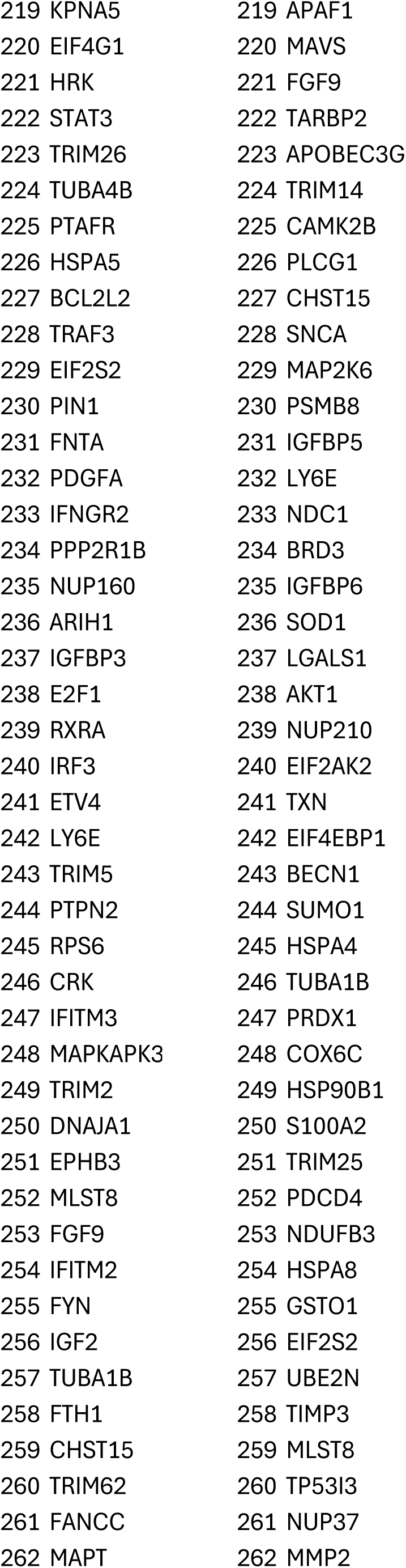
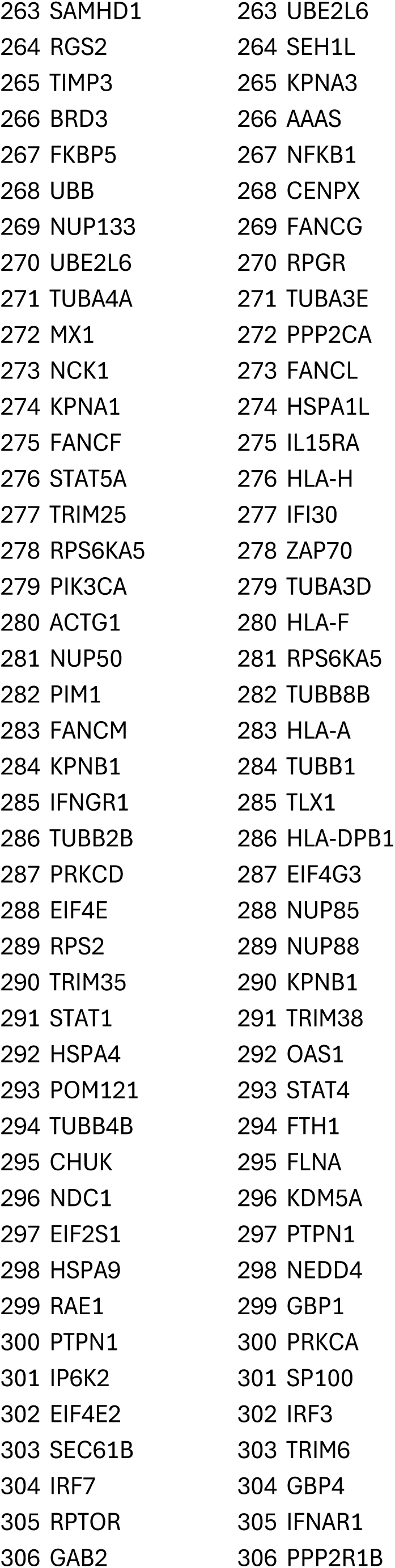
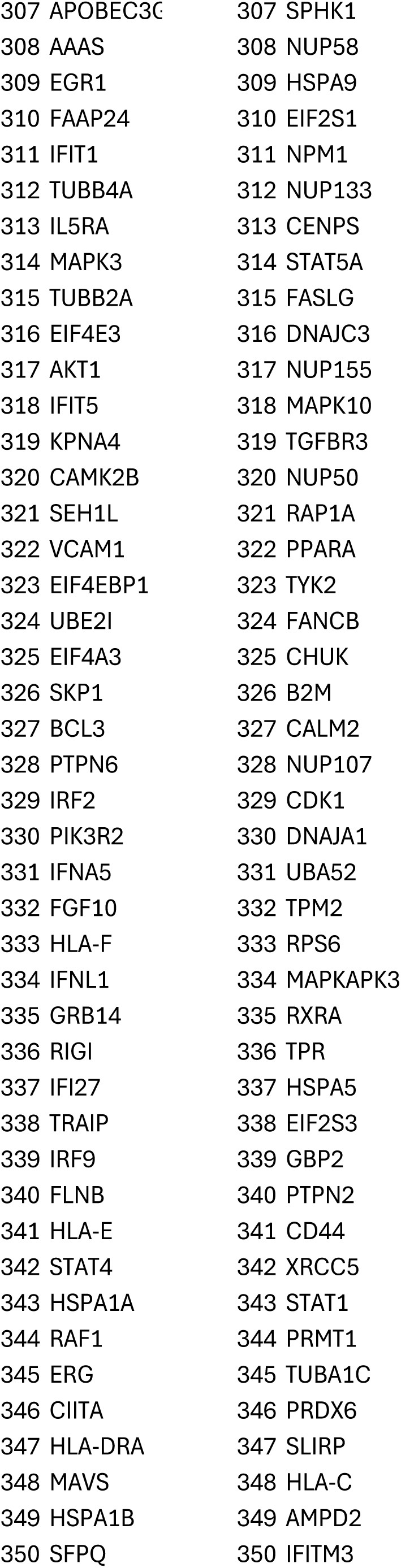
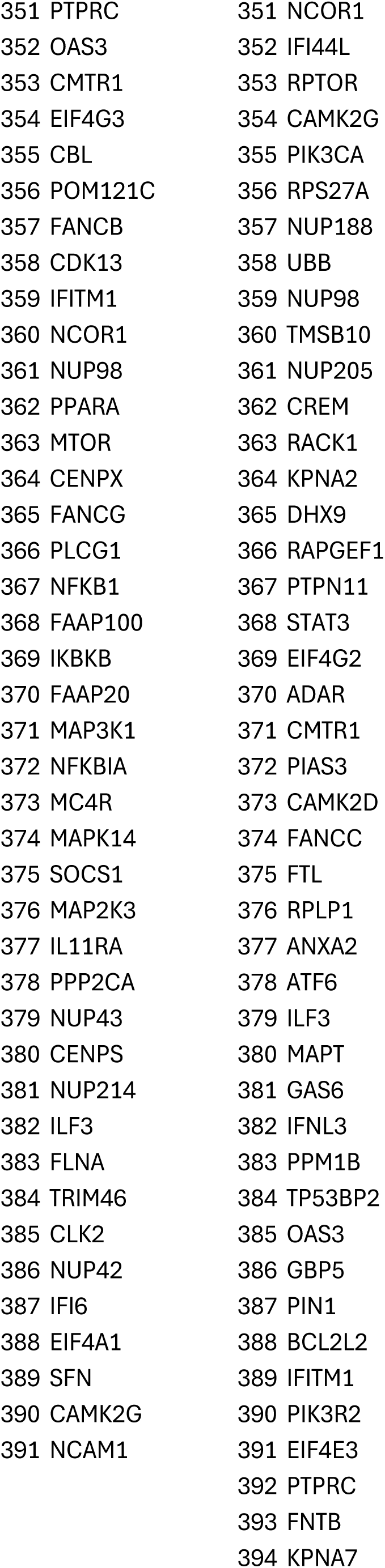
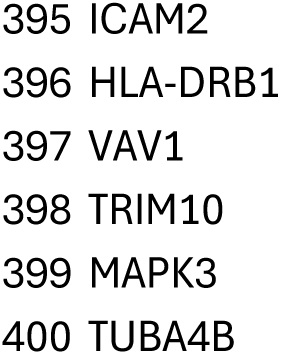
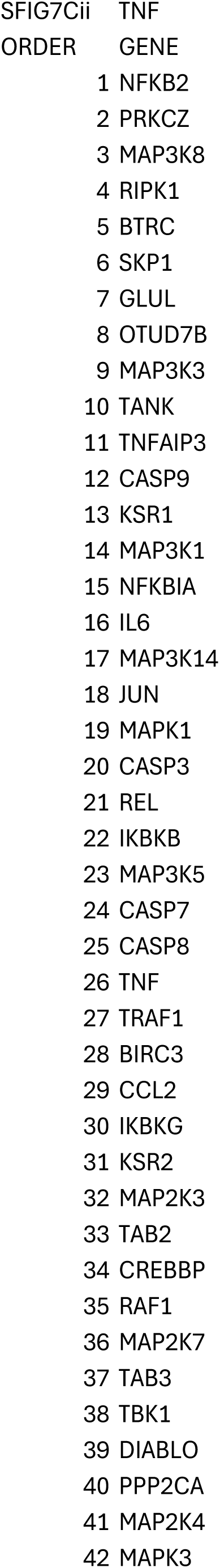
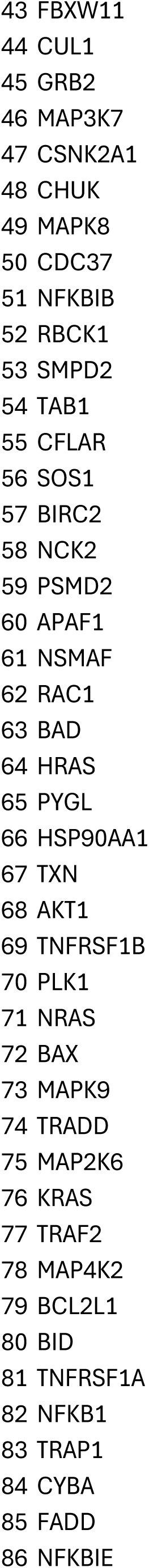
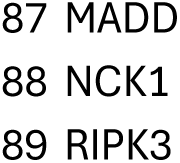

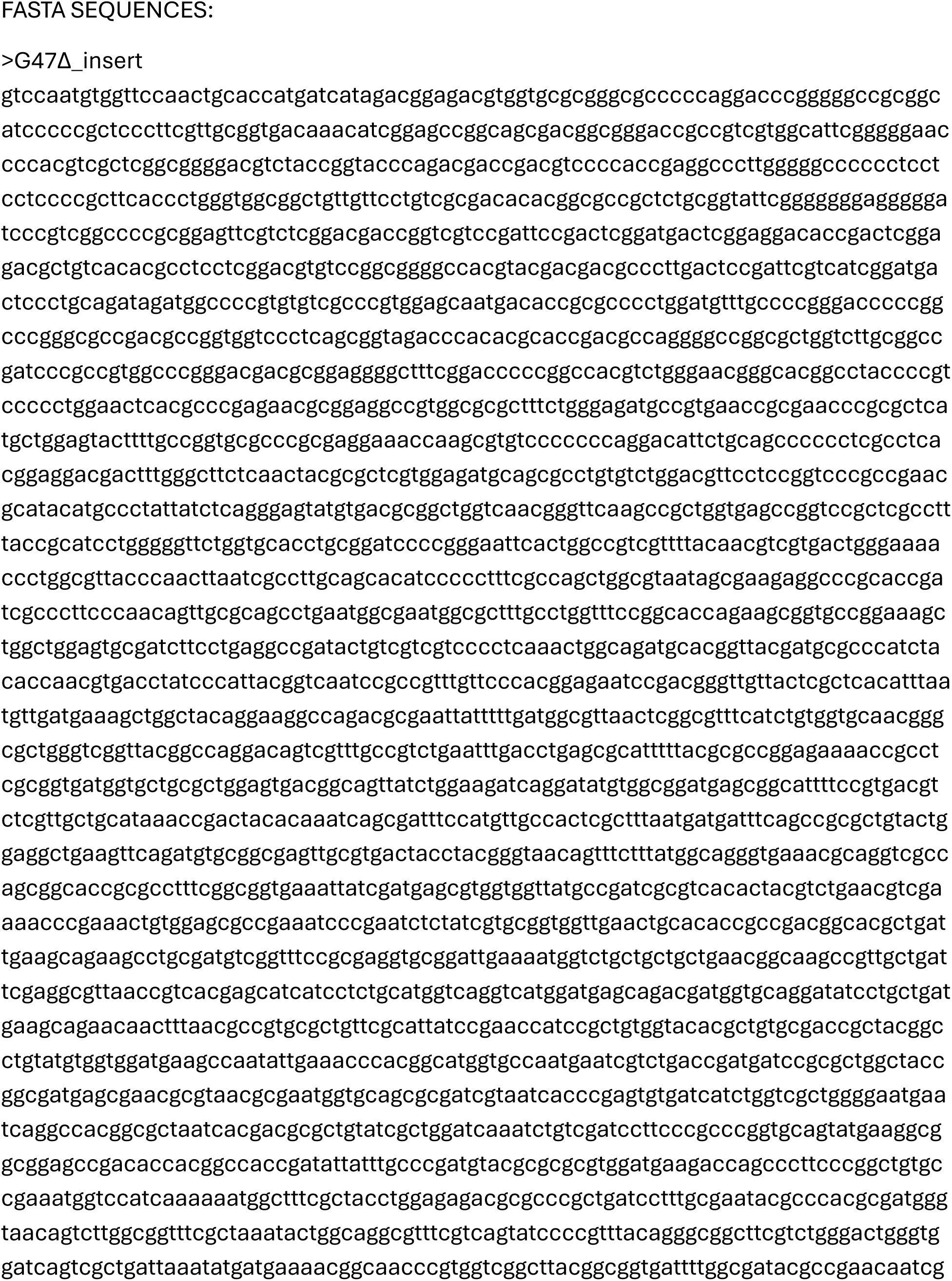
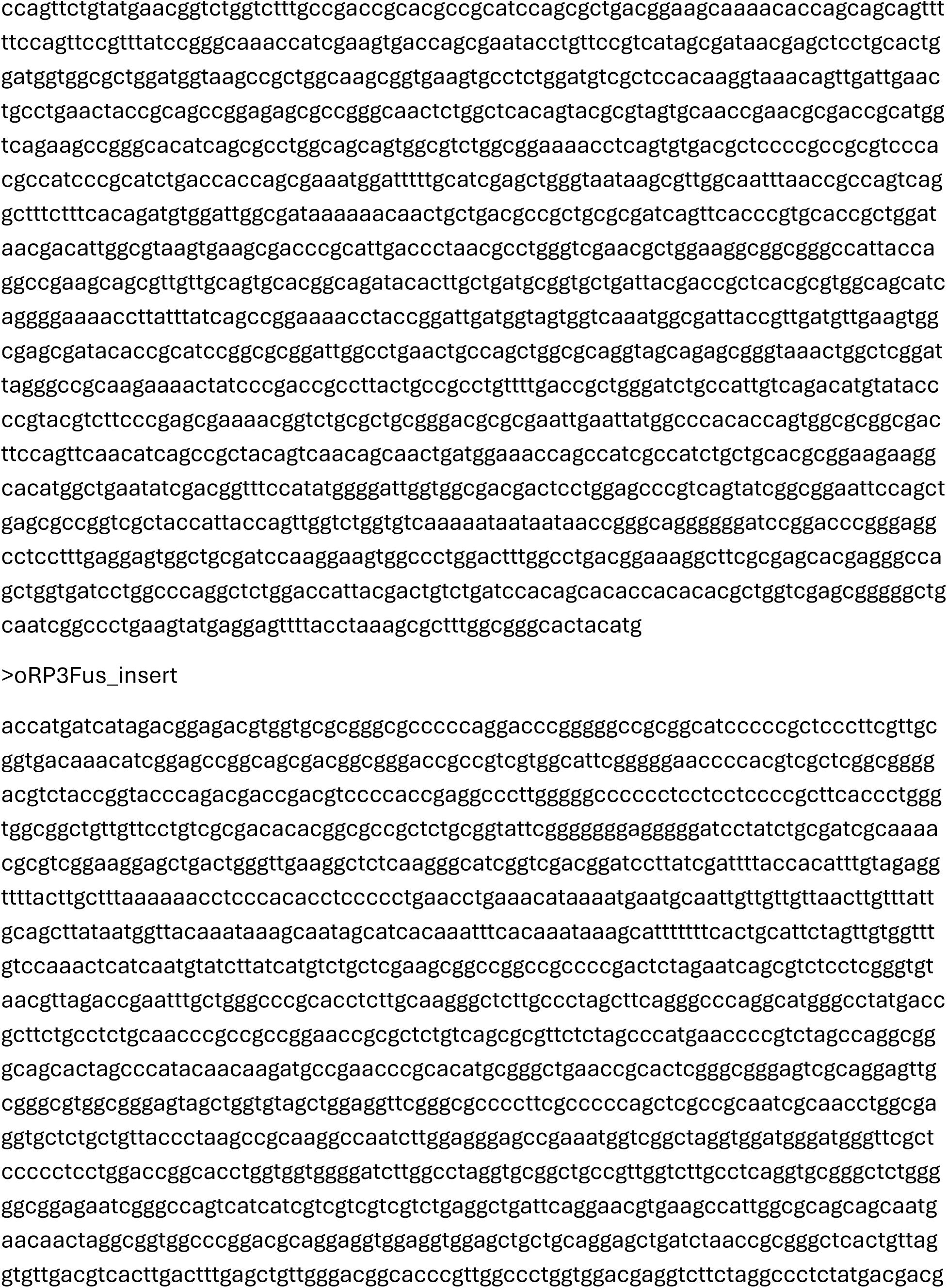
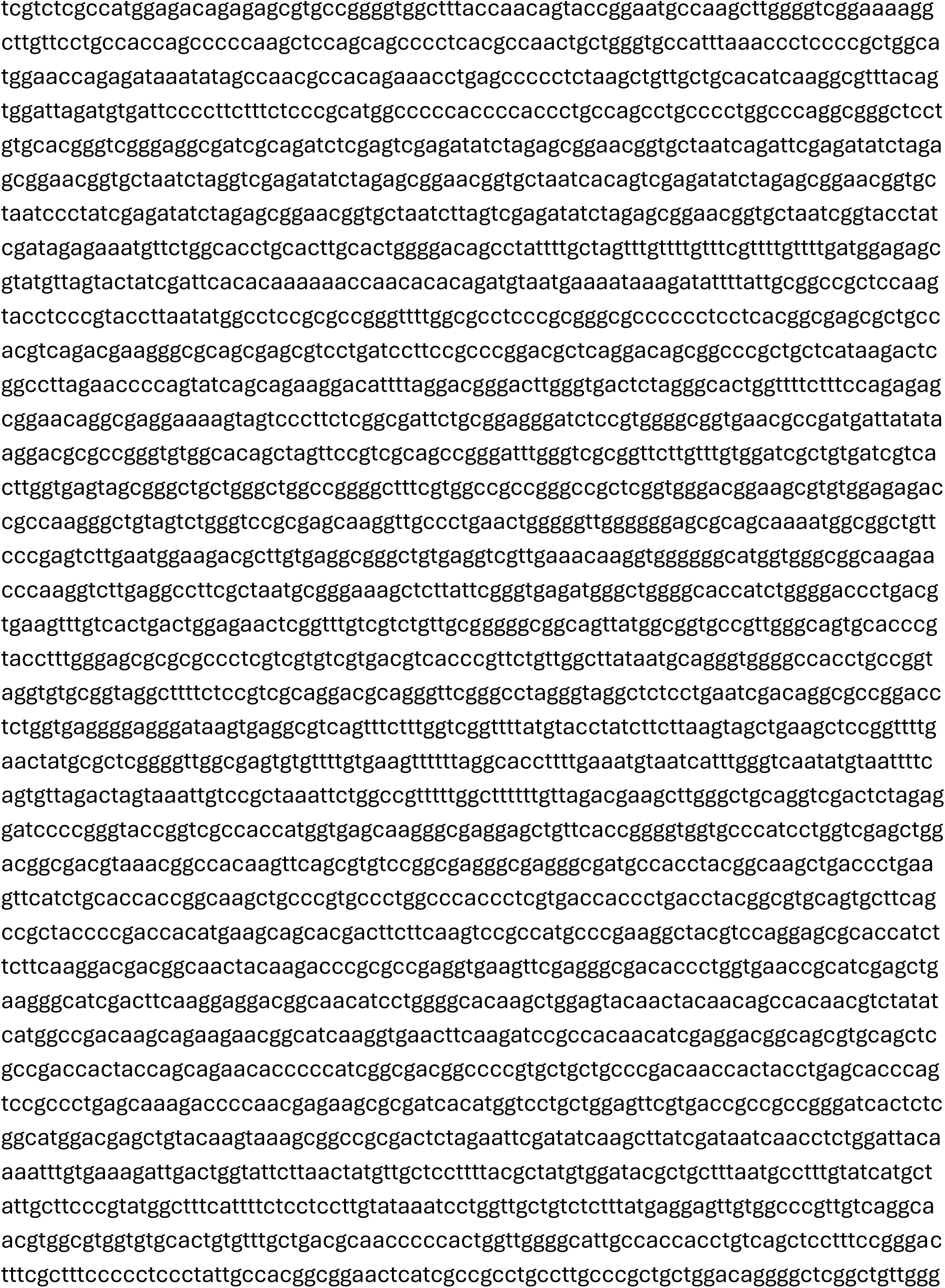
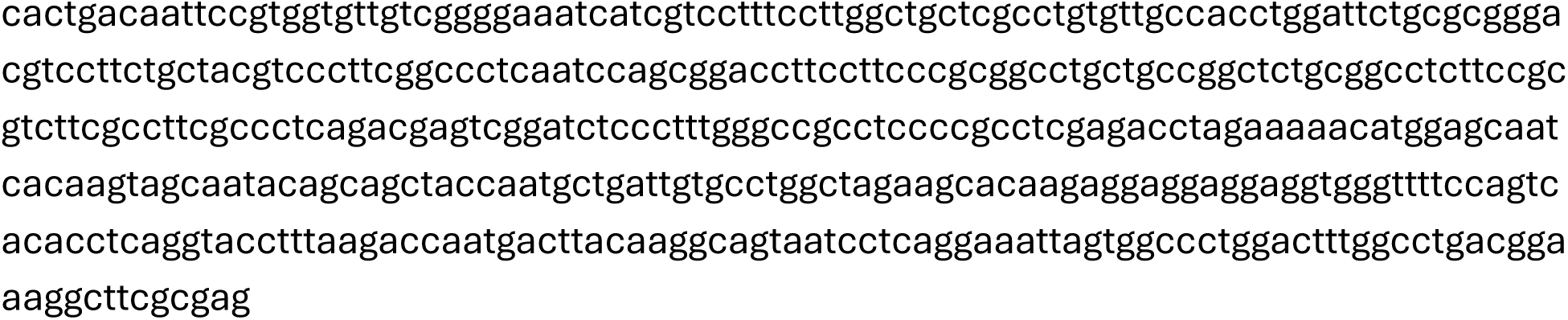

Significant DEGs during expression of PAX3-FOXO1 between Mock RD:P3F versus Mock RD:EMP

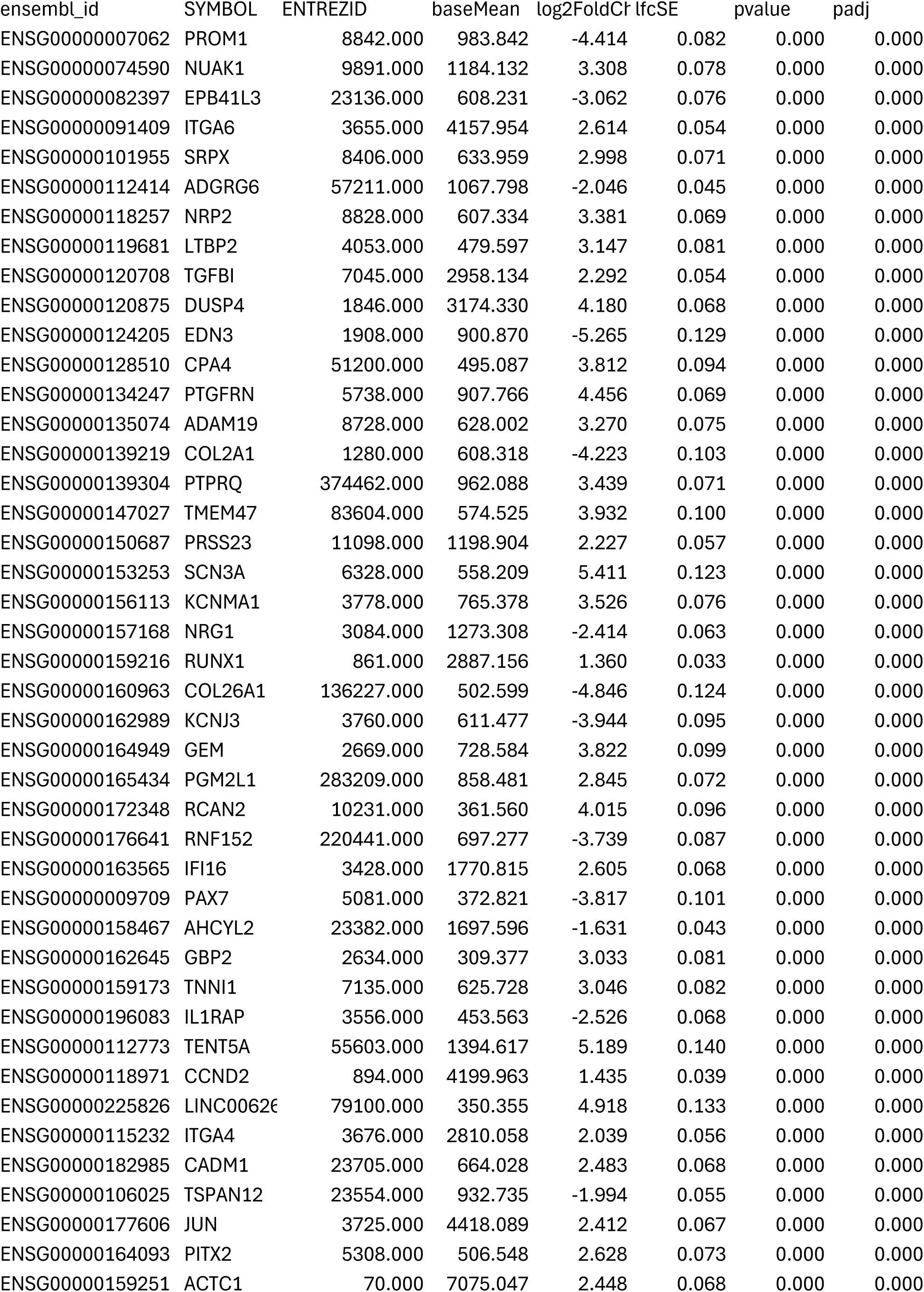
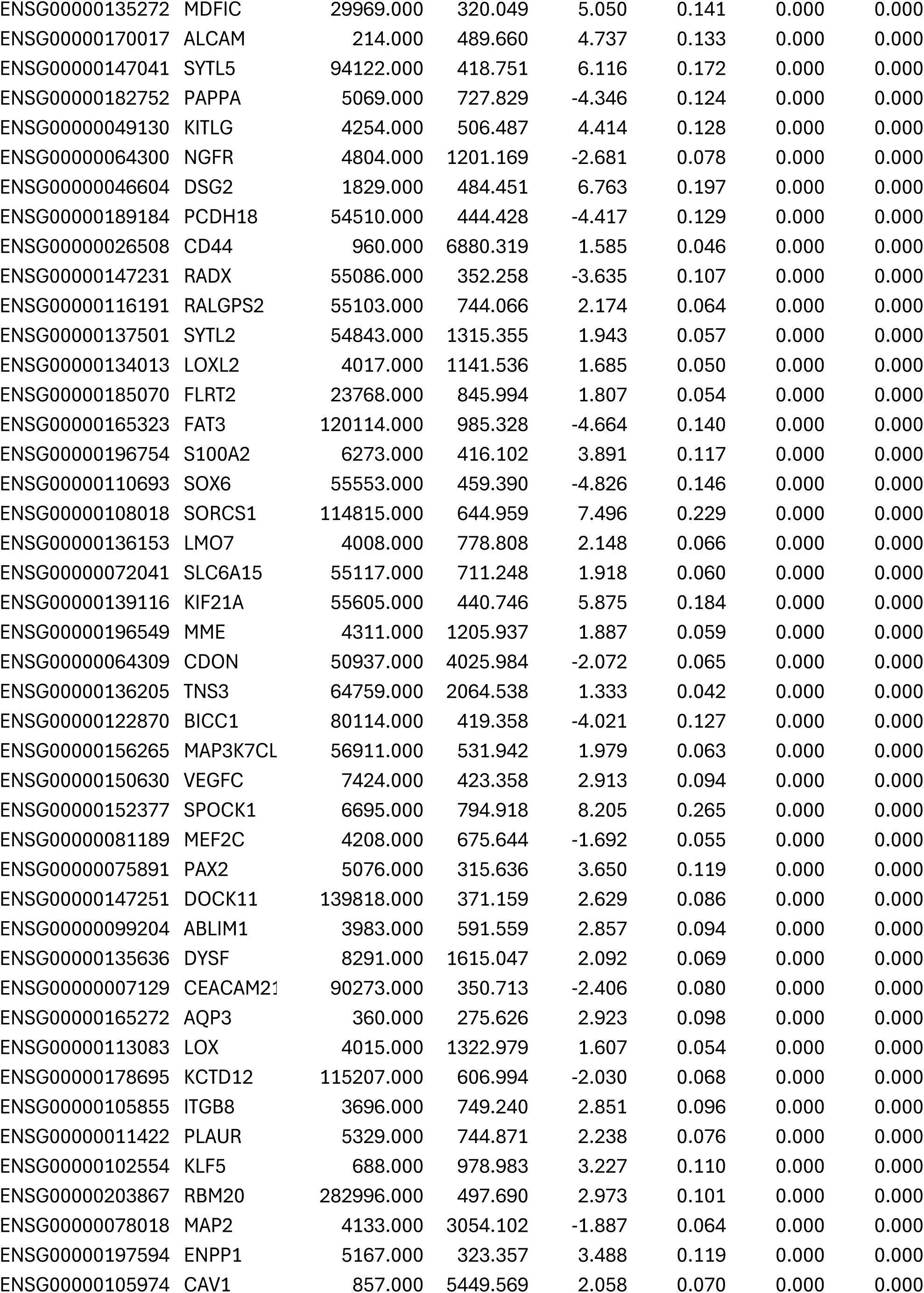
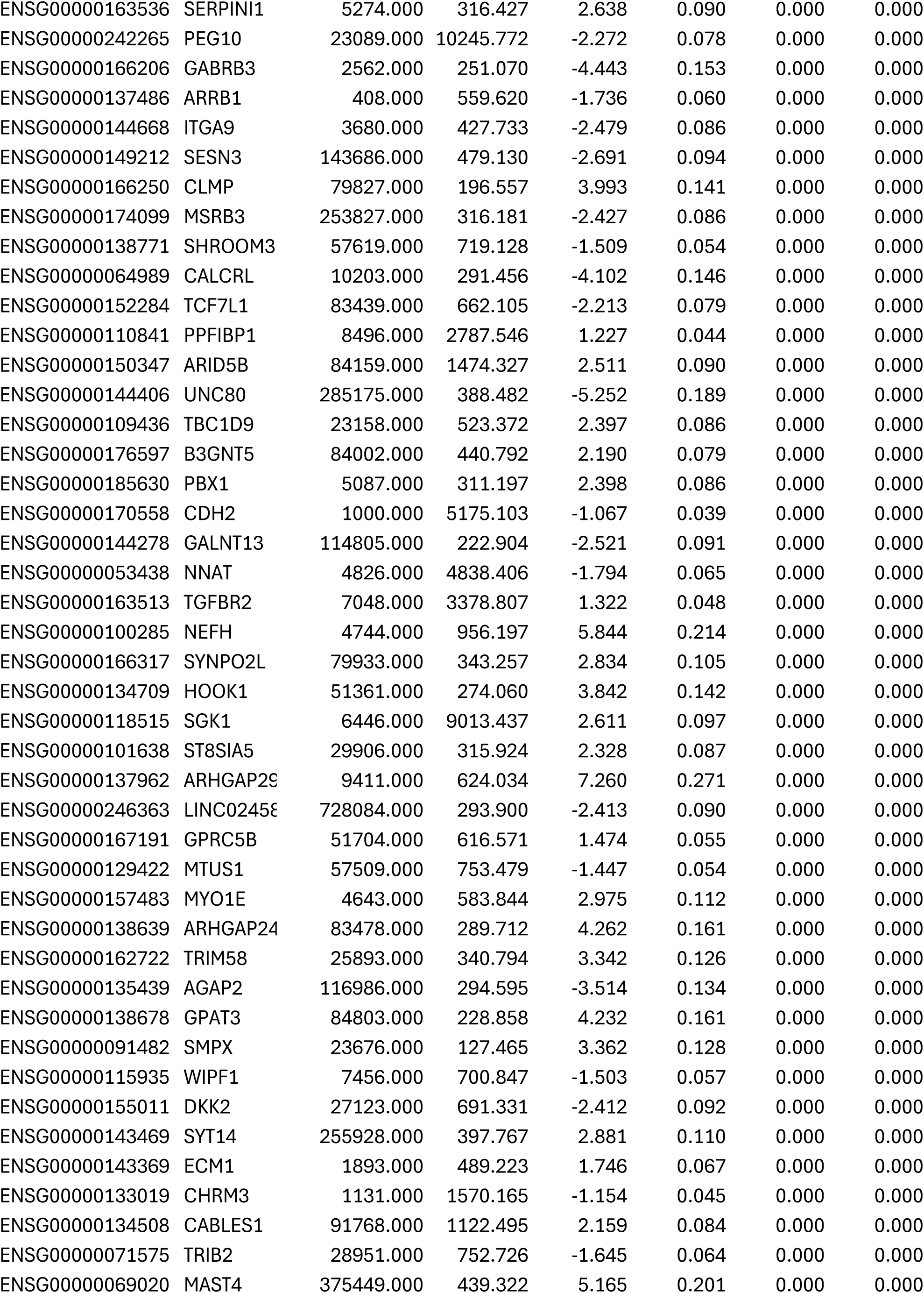
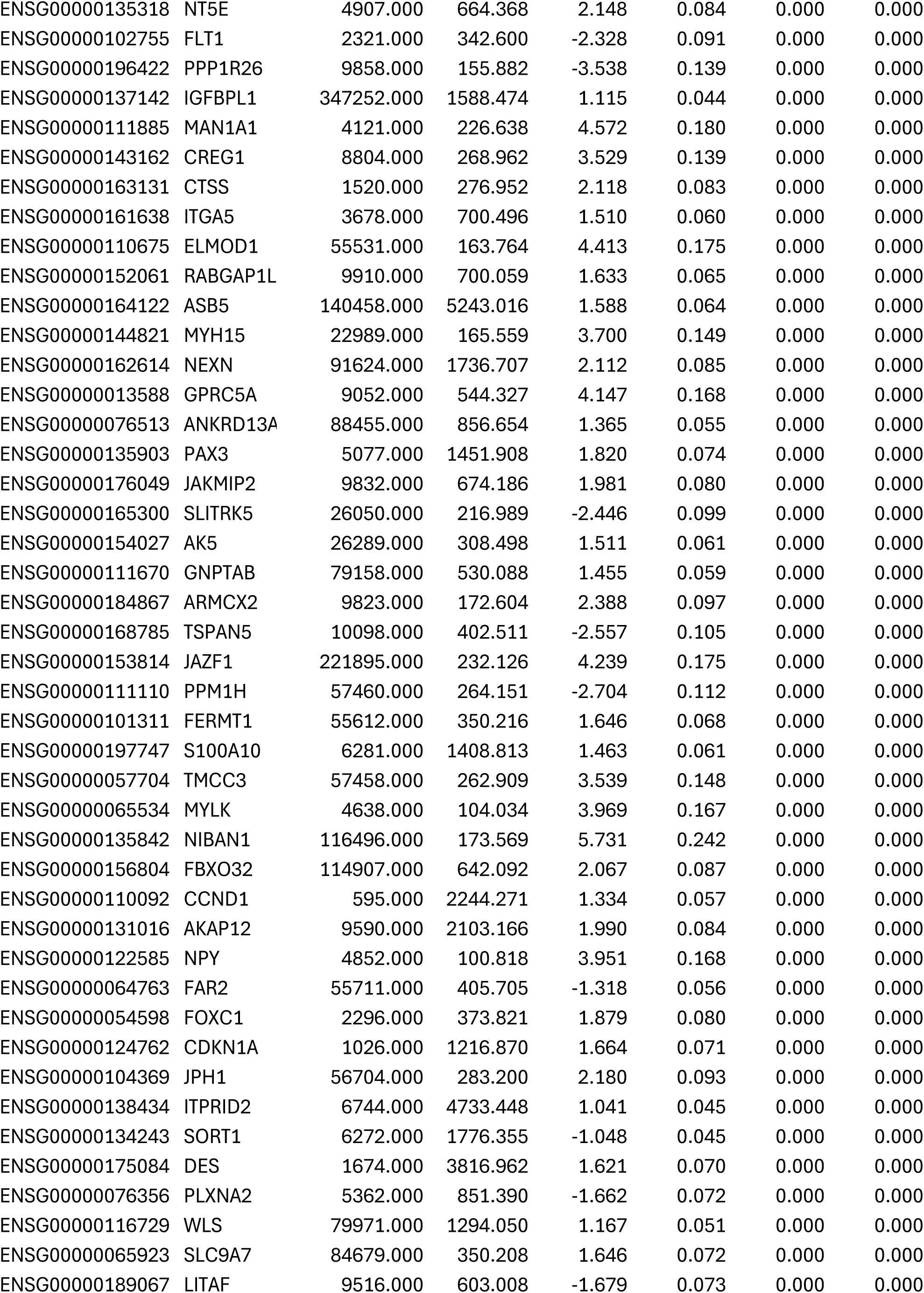
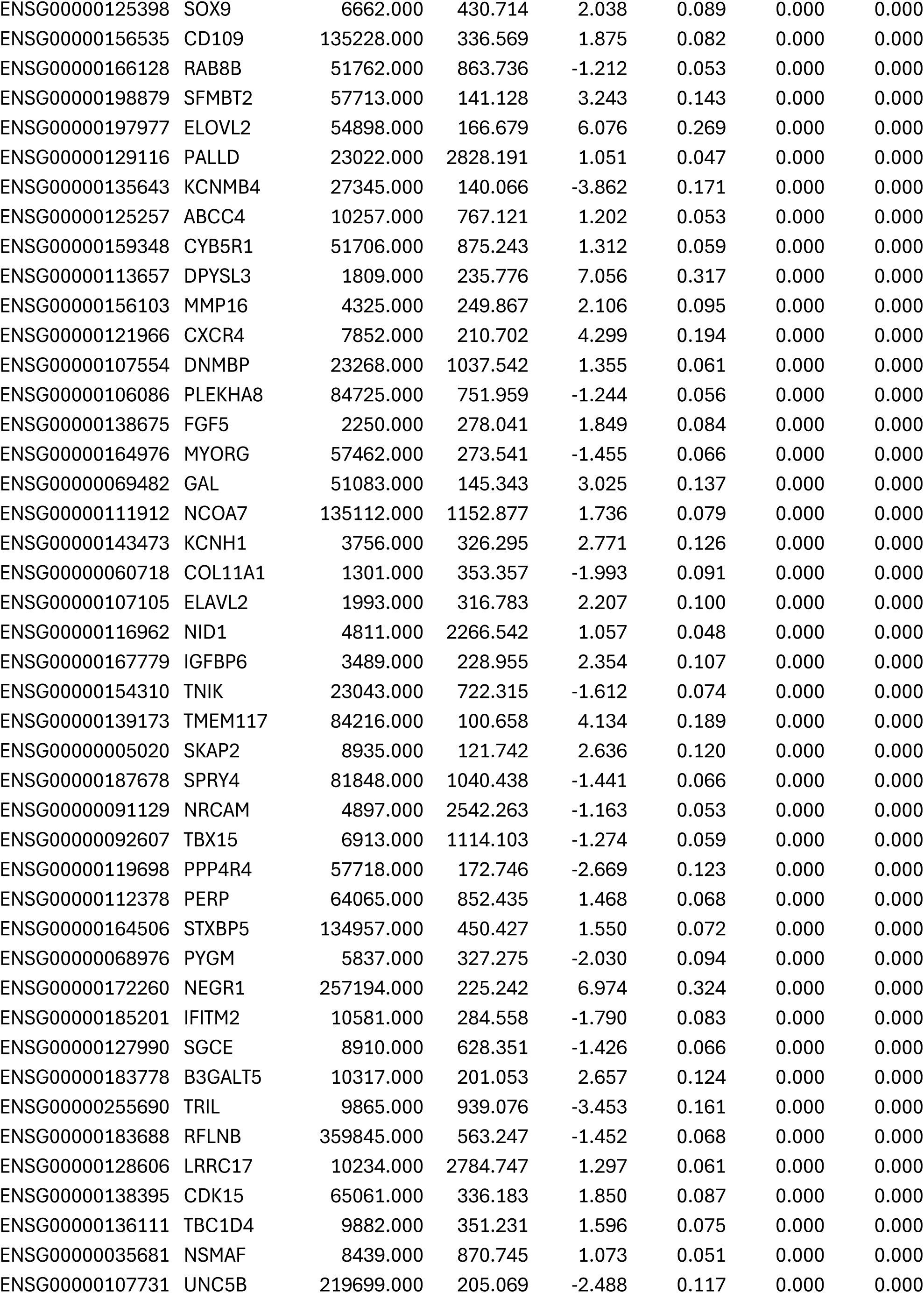
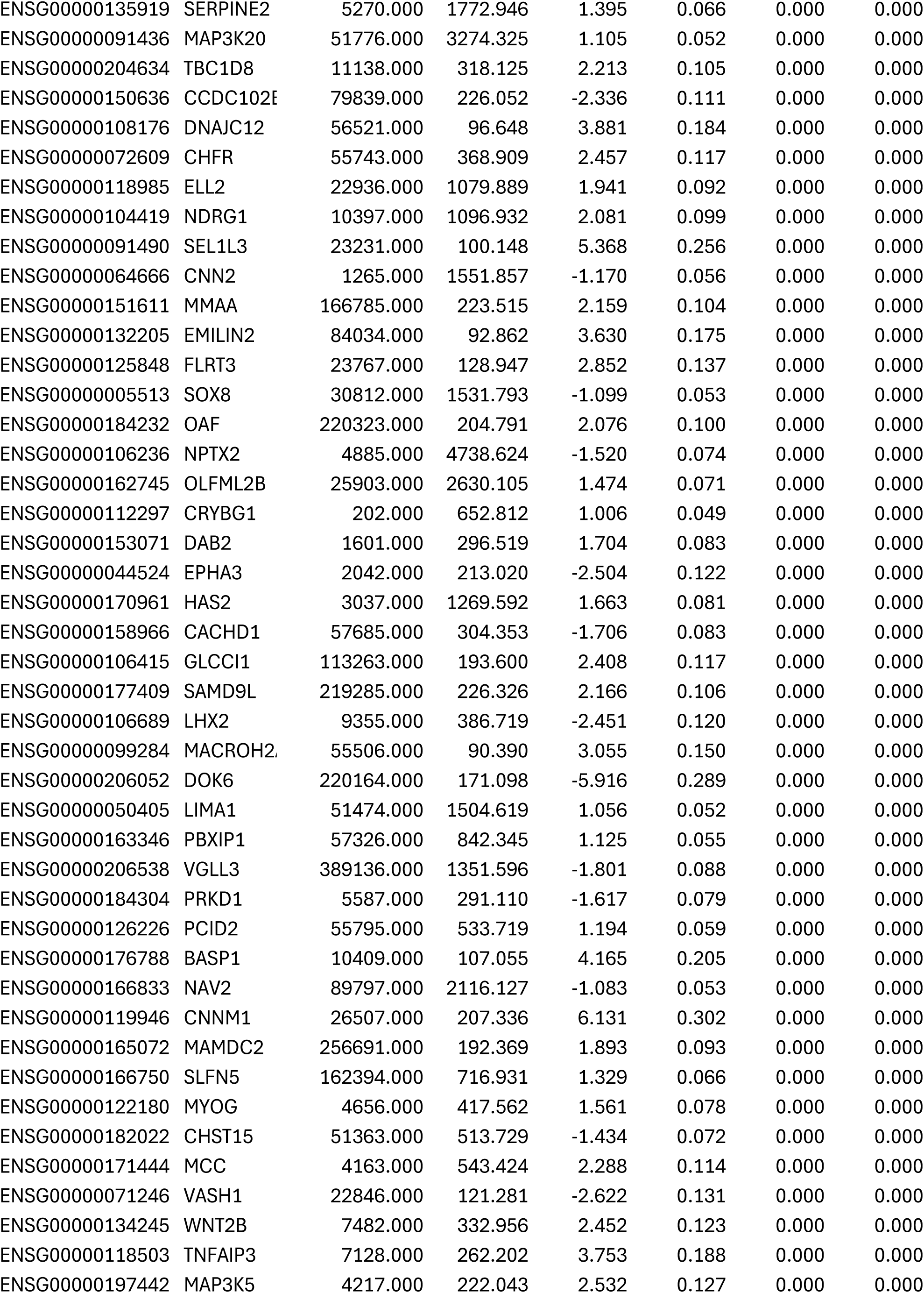
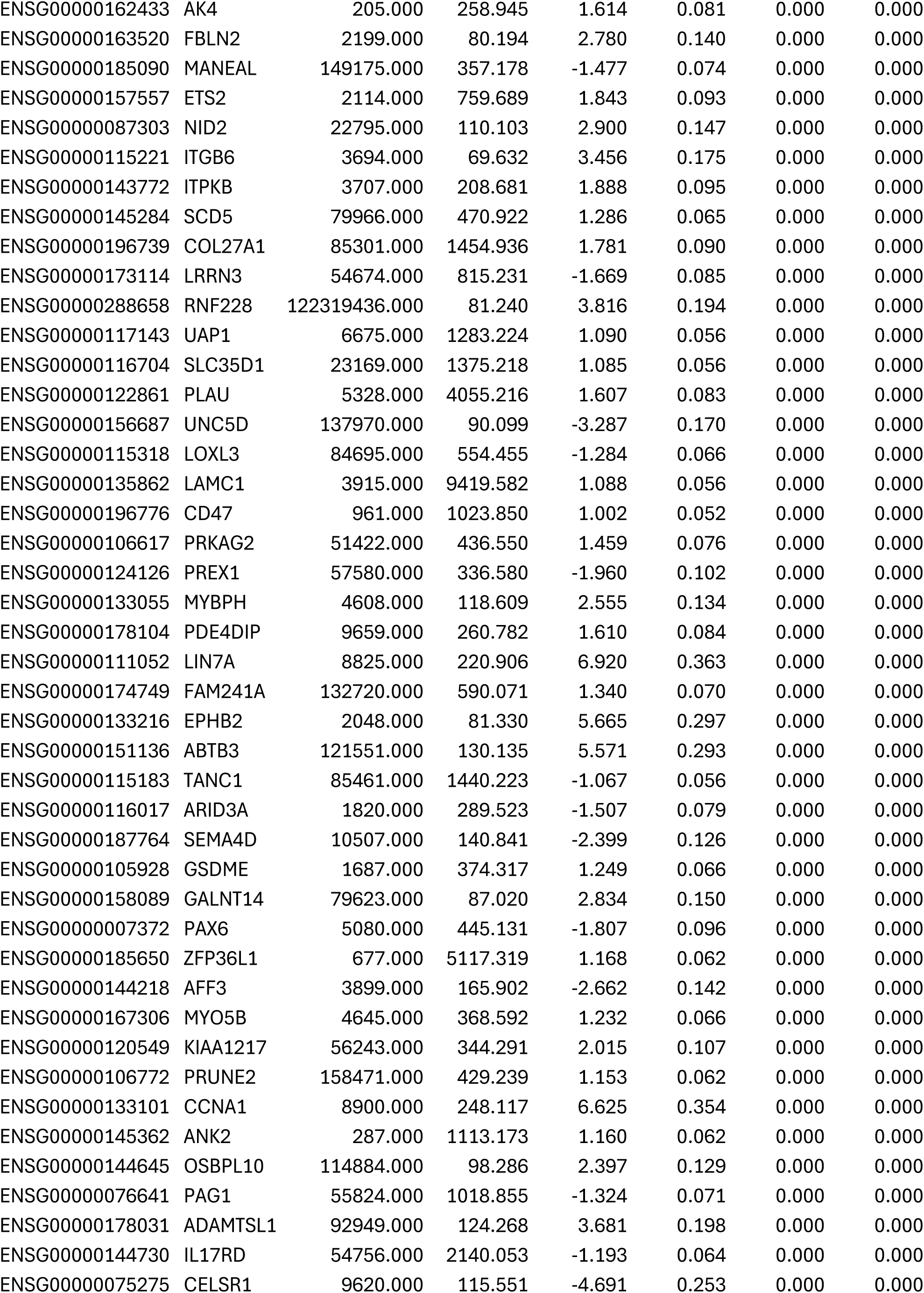
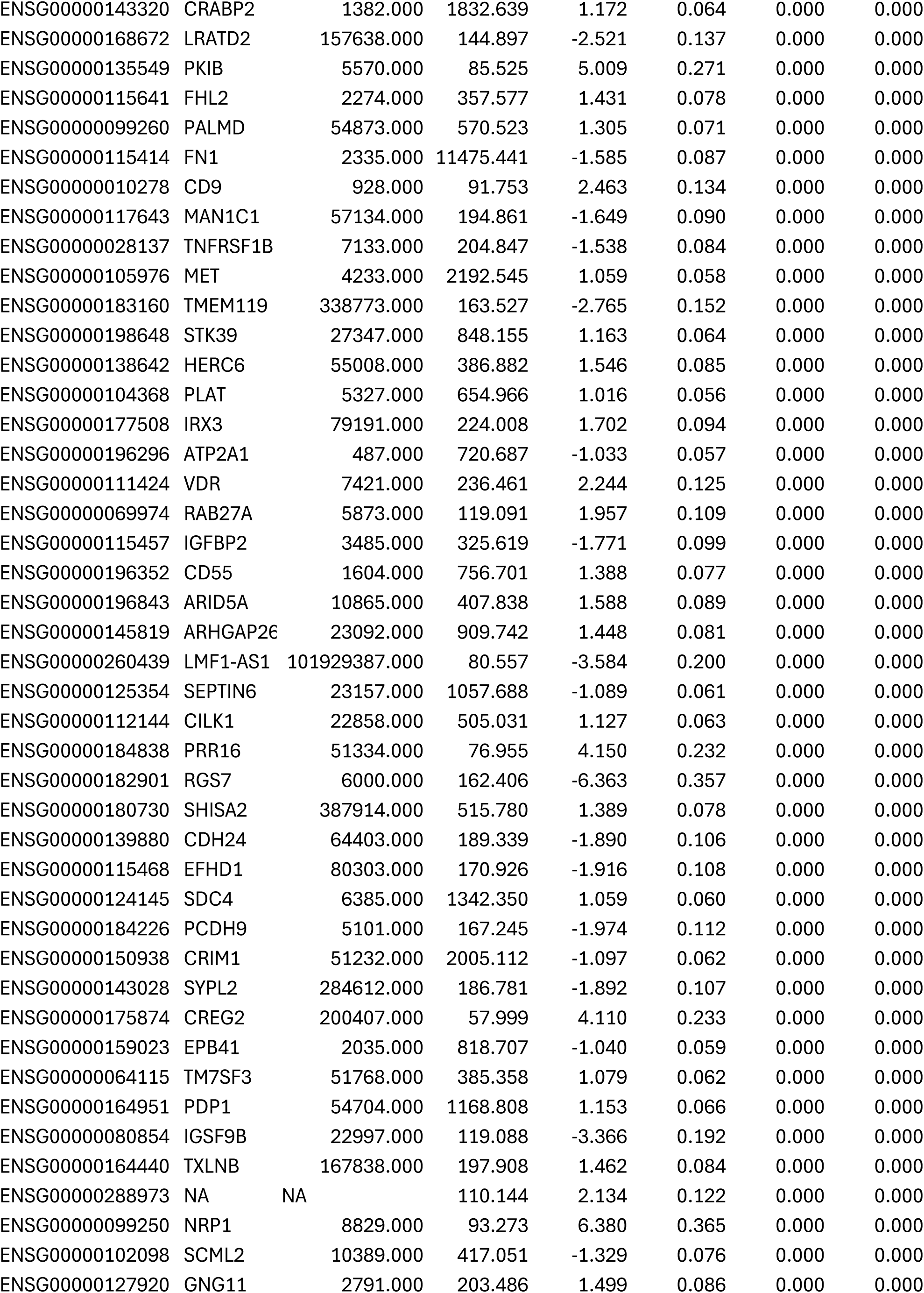
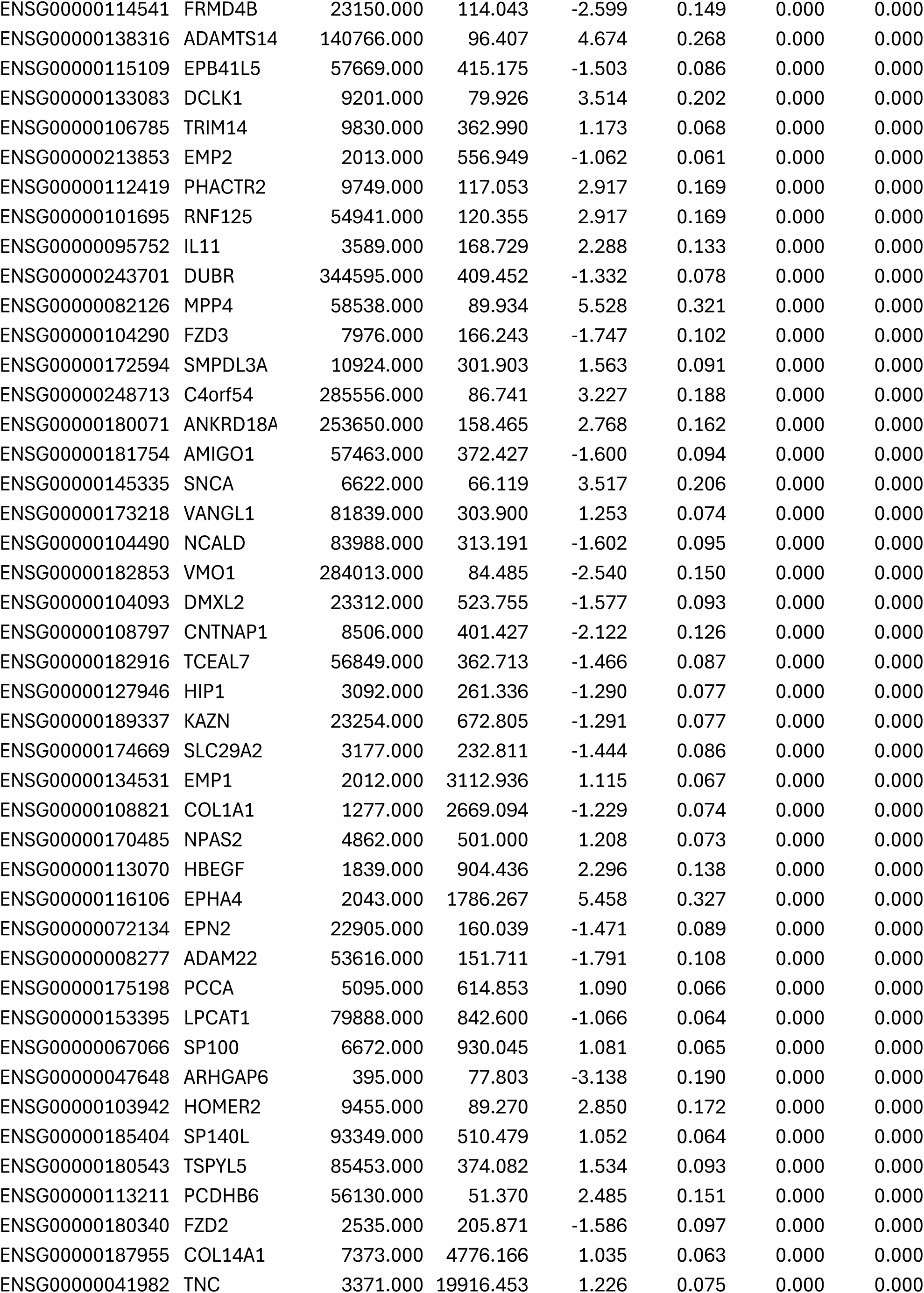
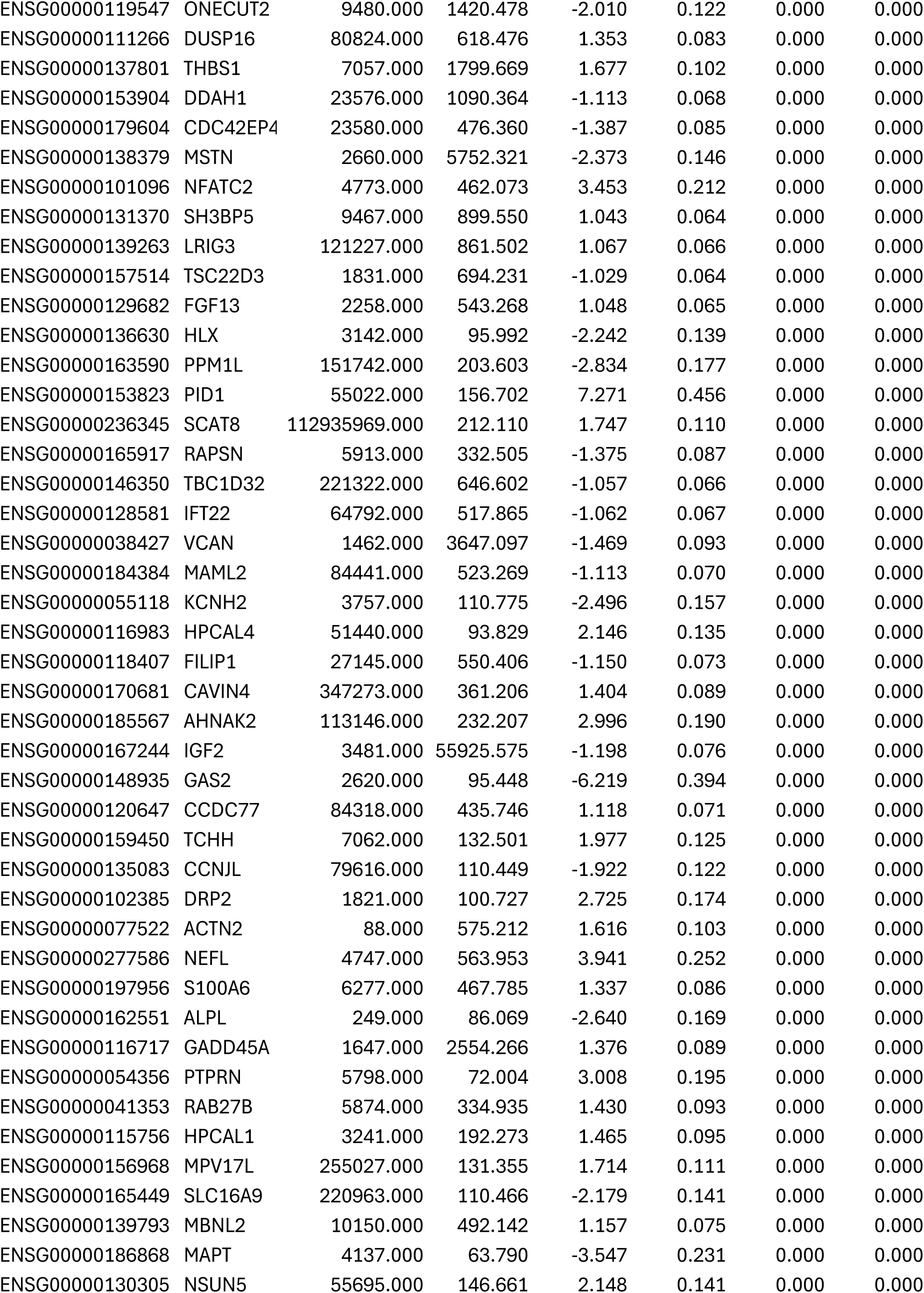
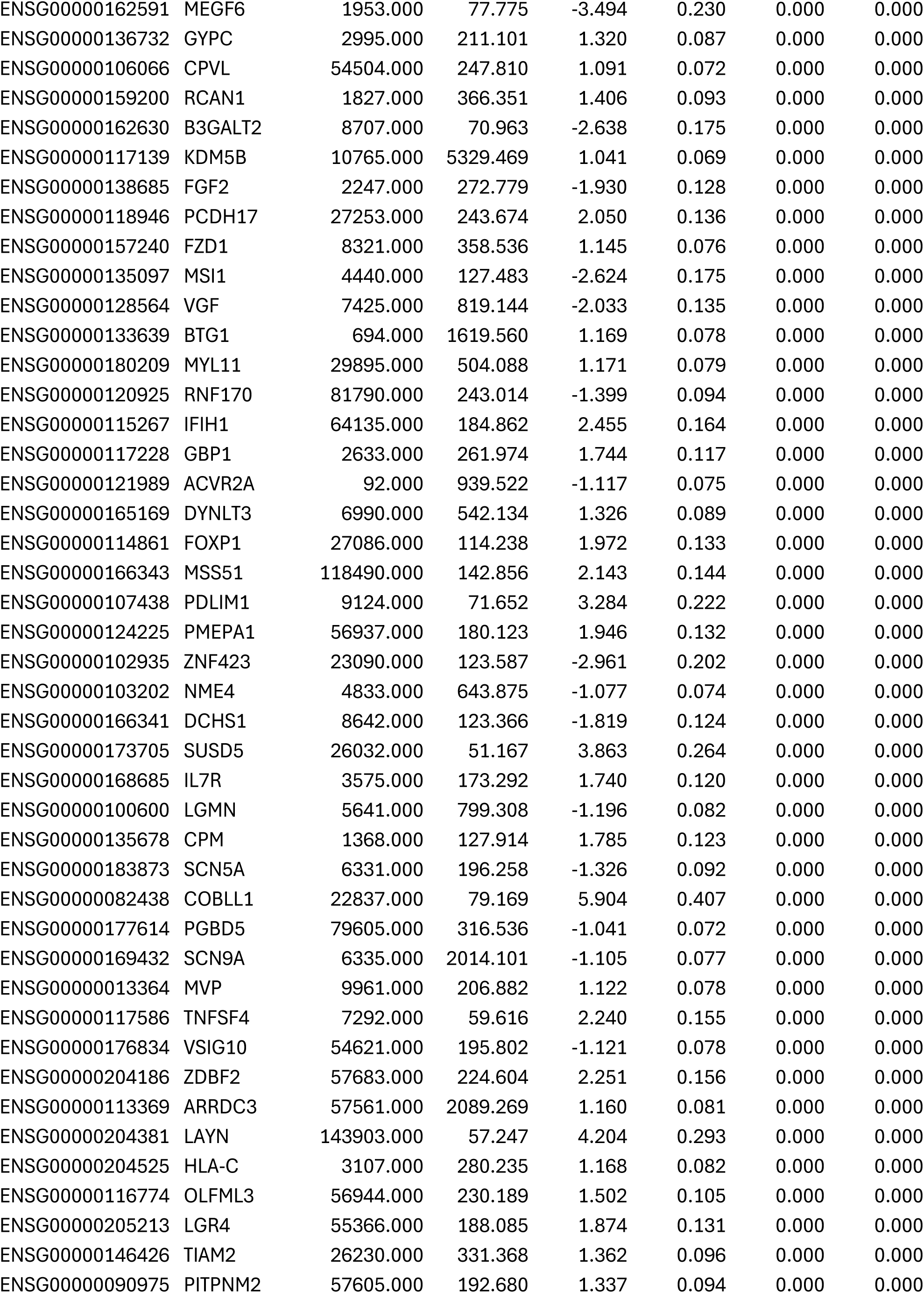
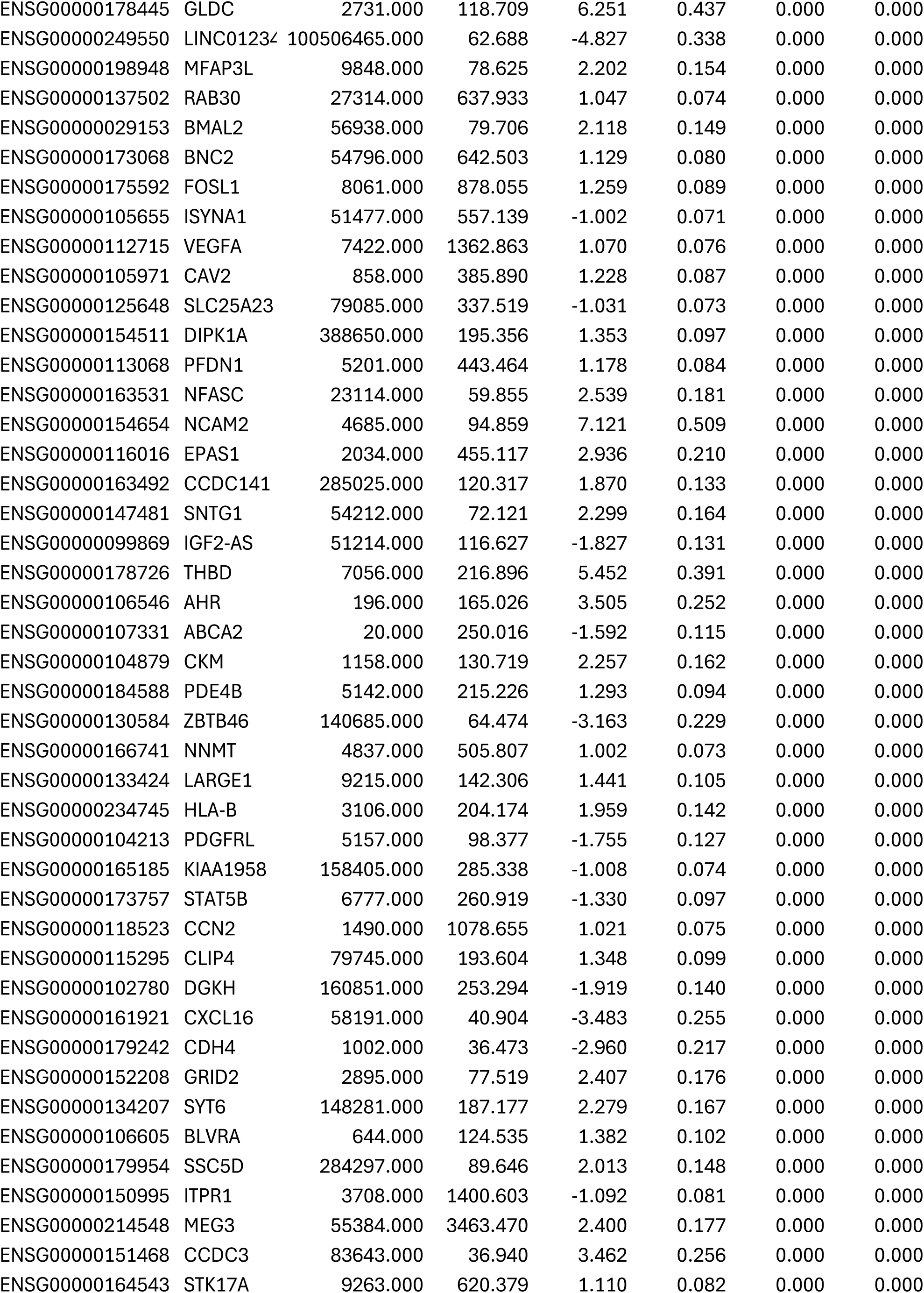
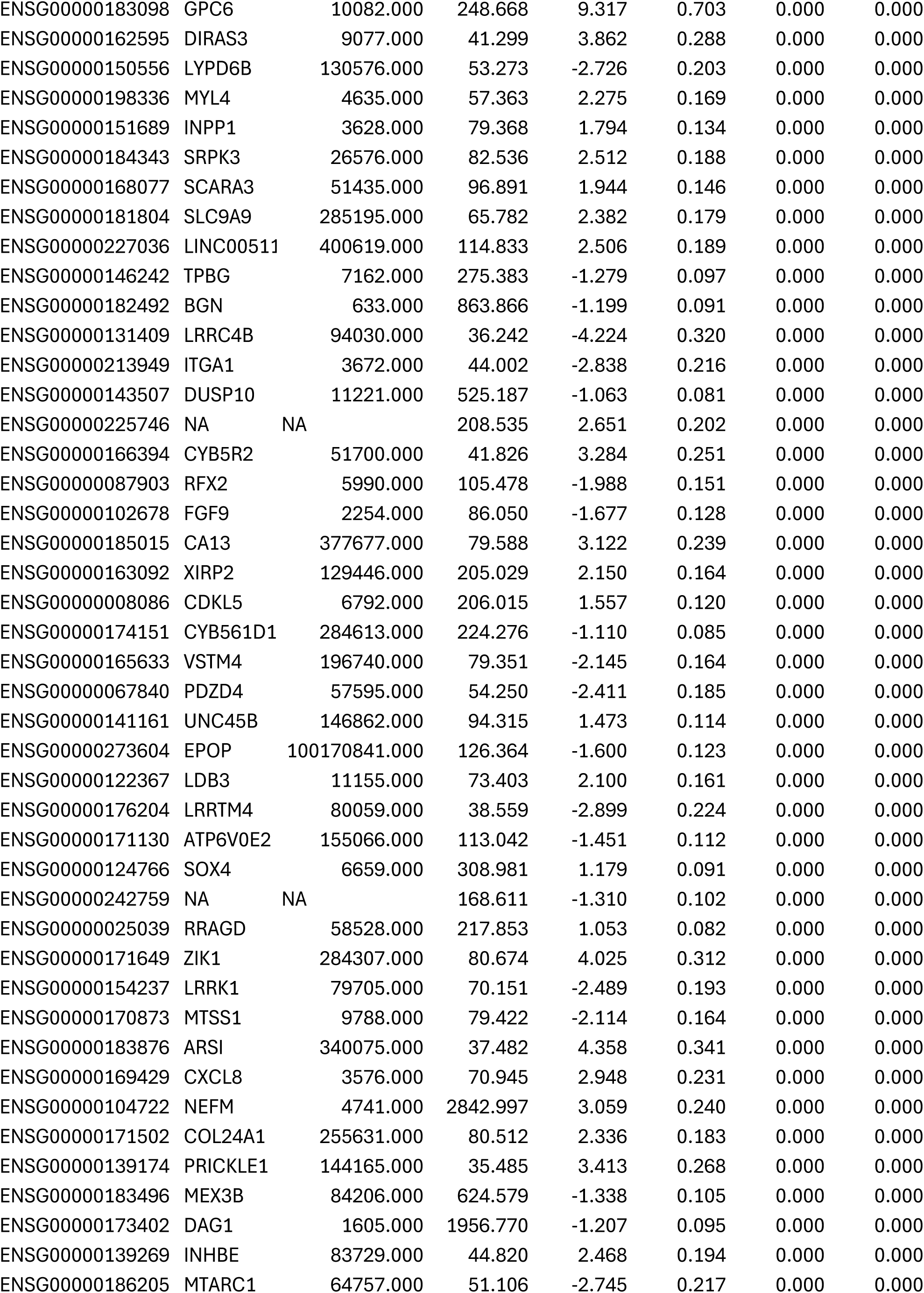
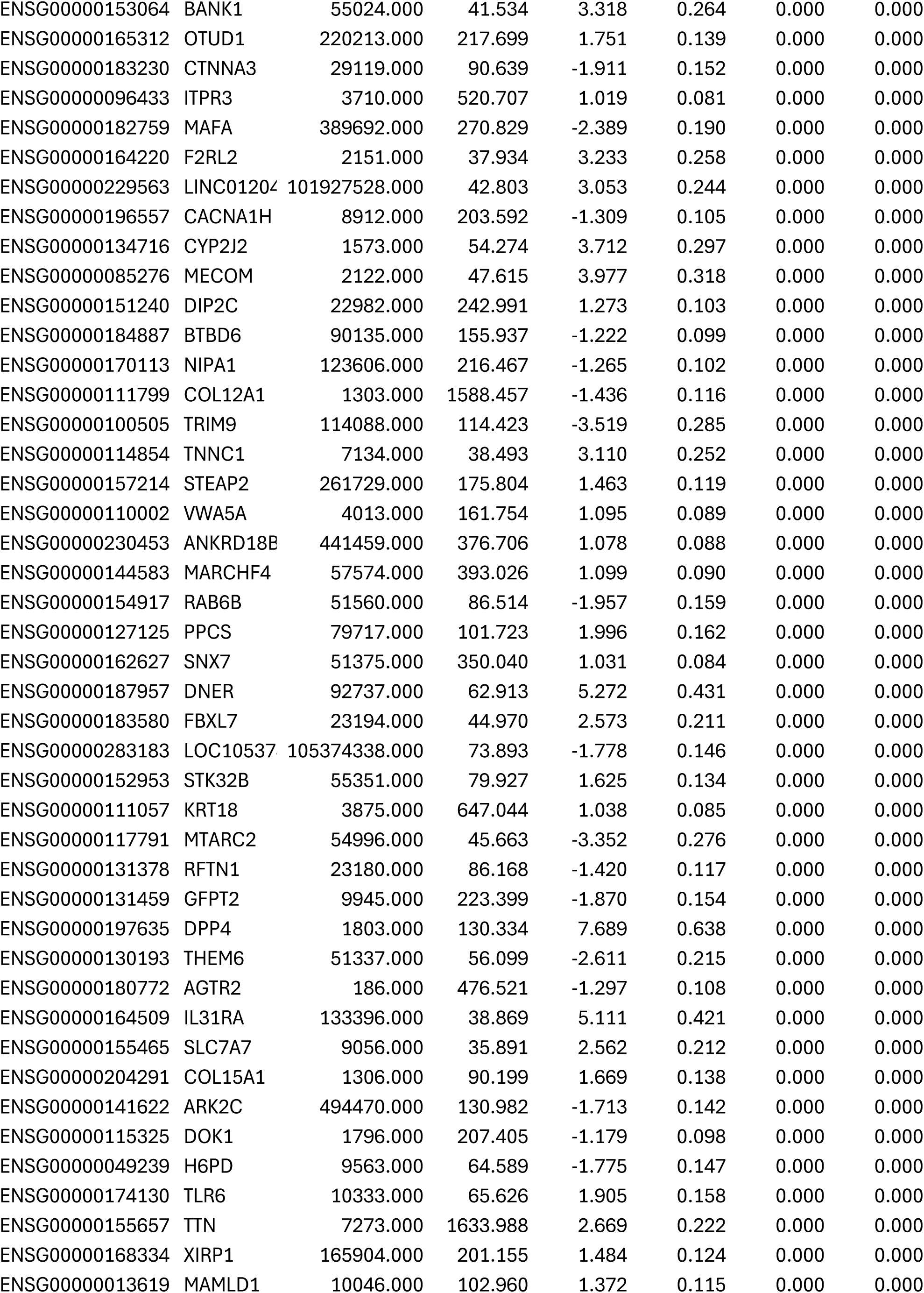
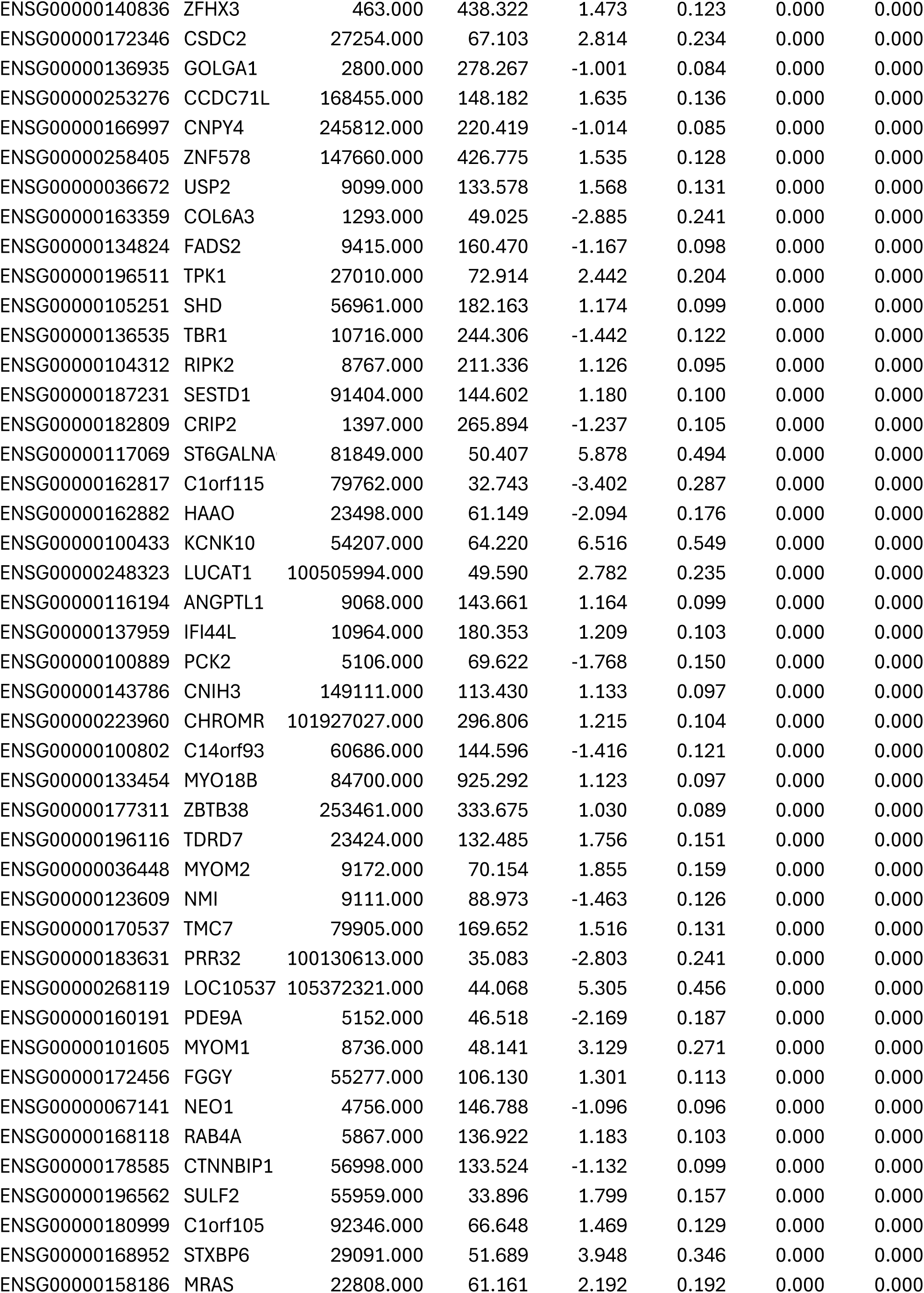
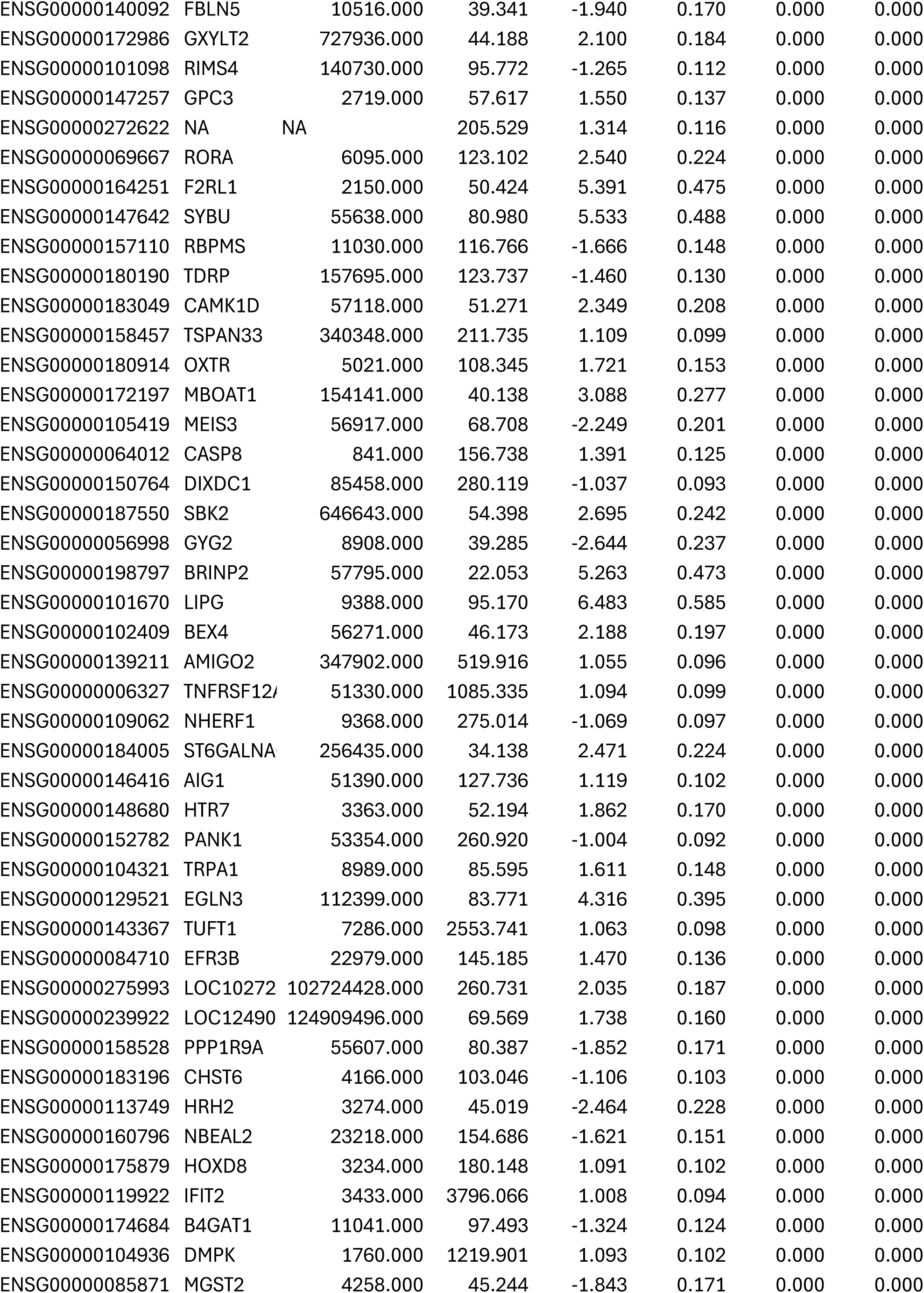
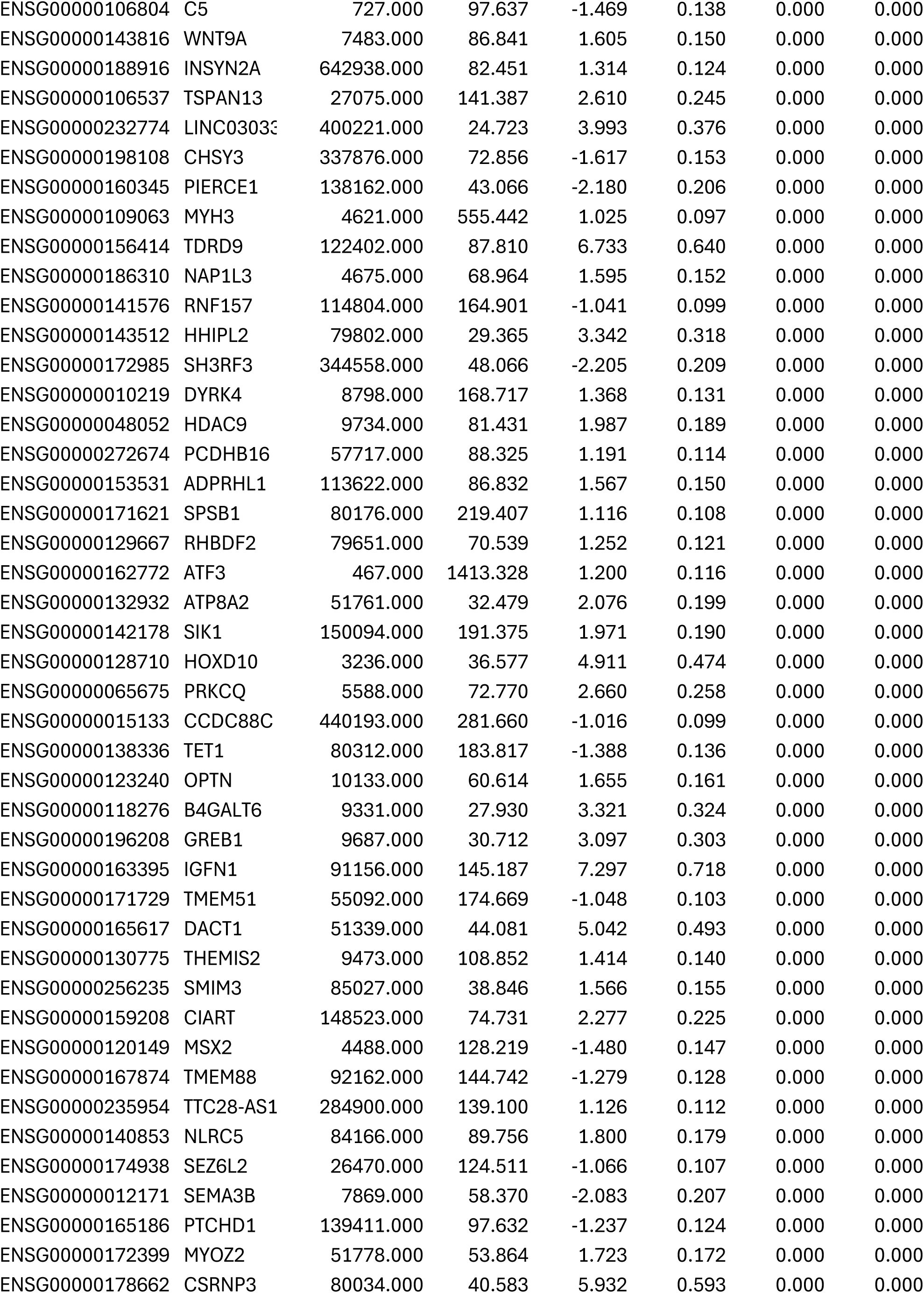
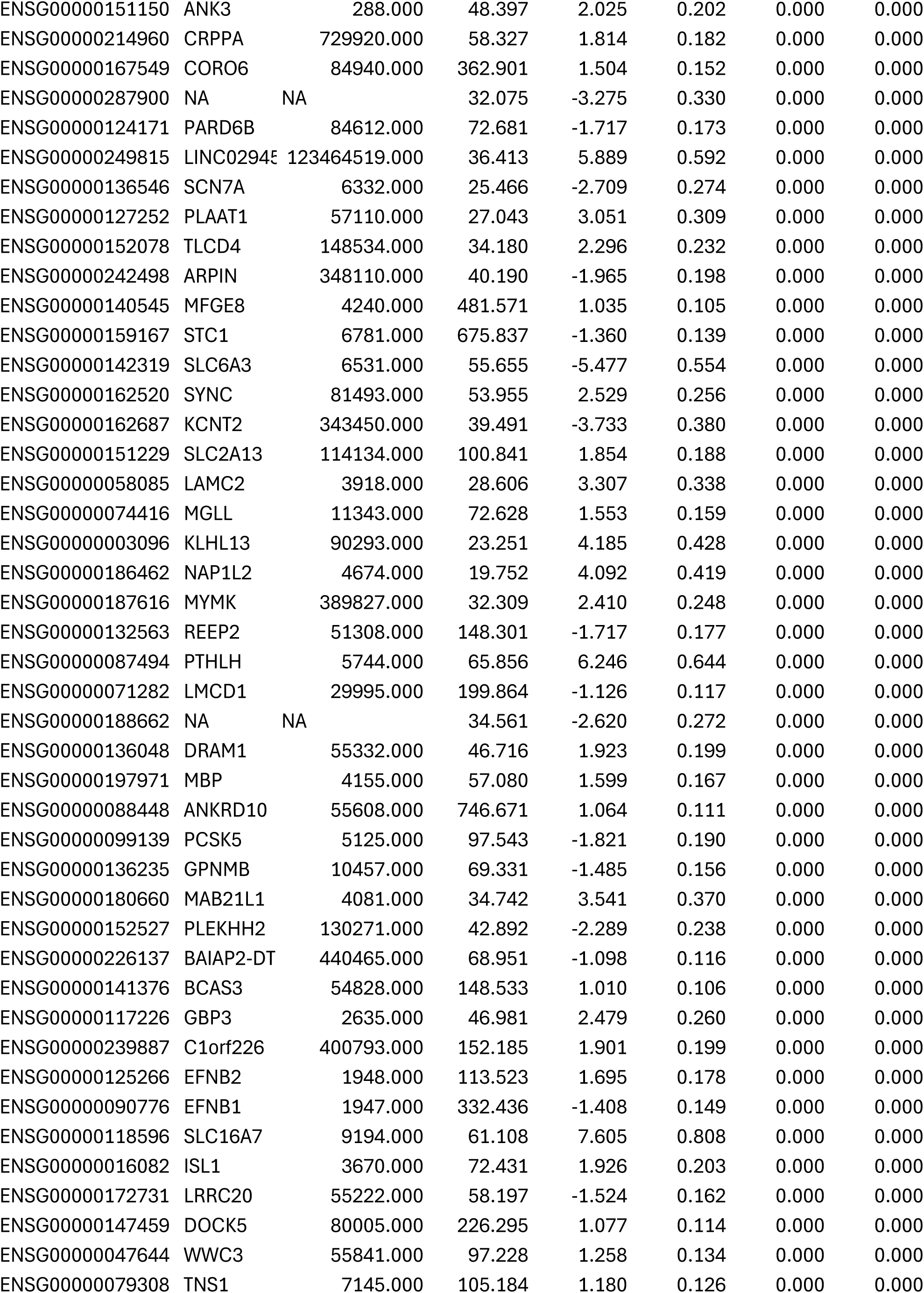
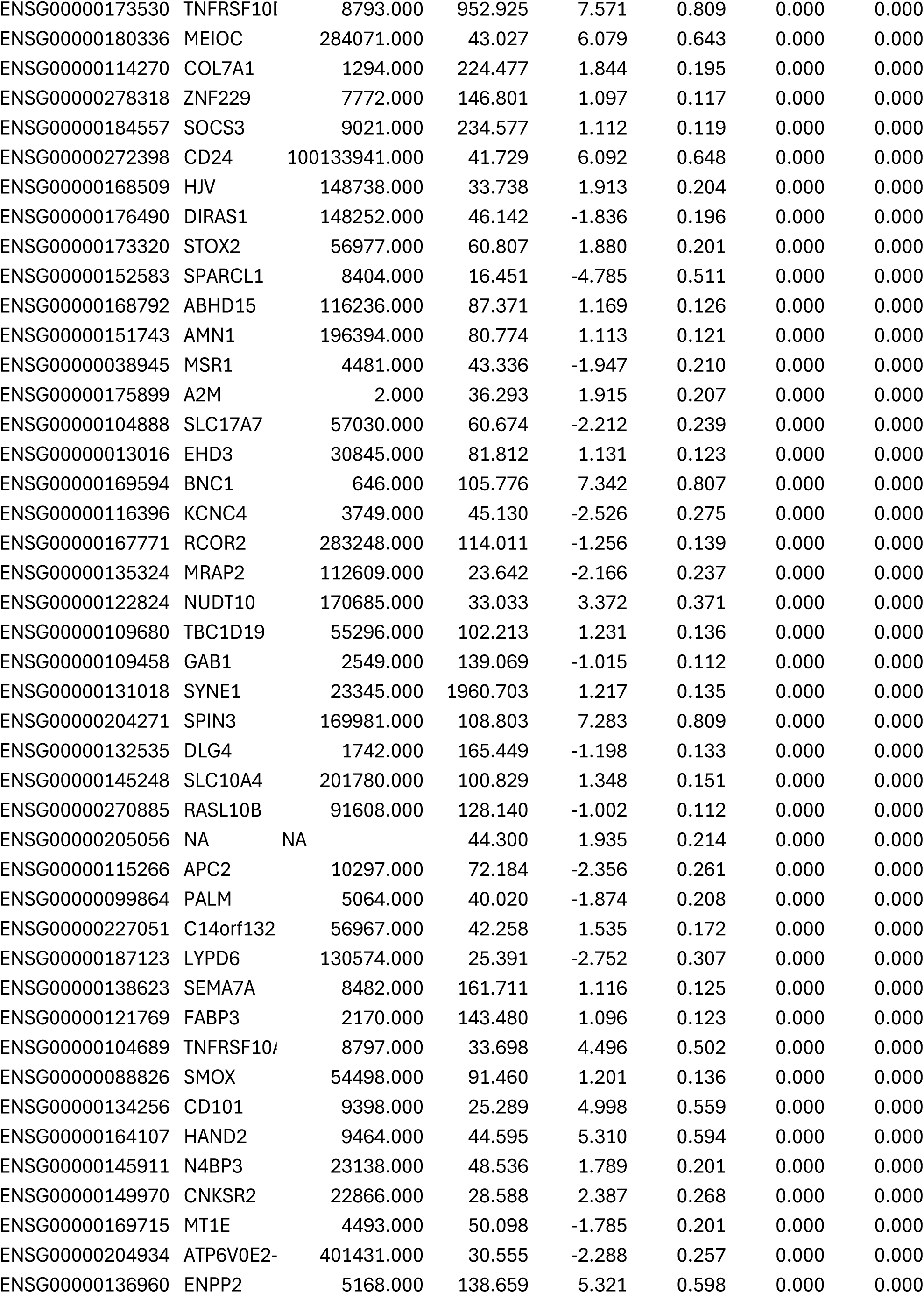
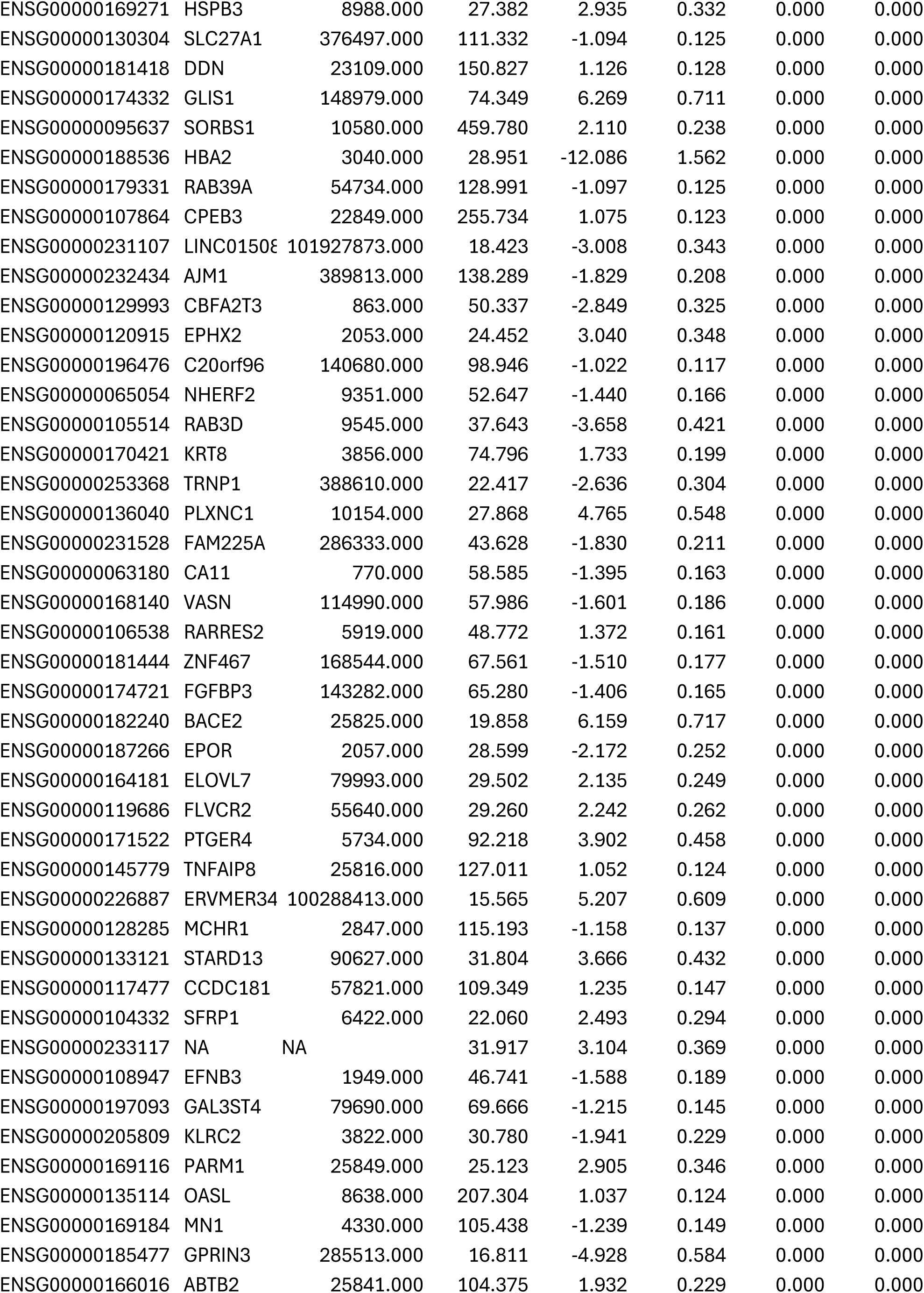
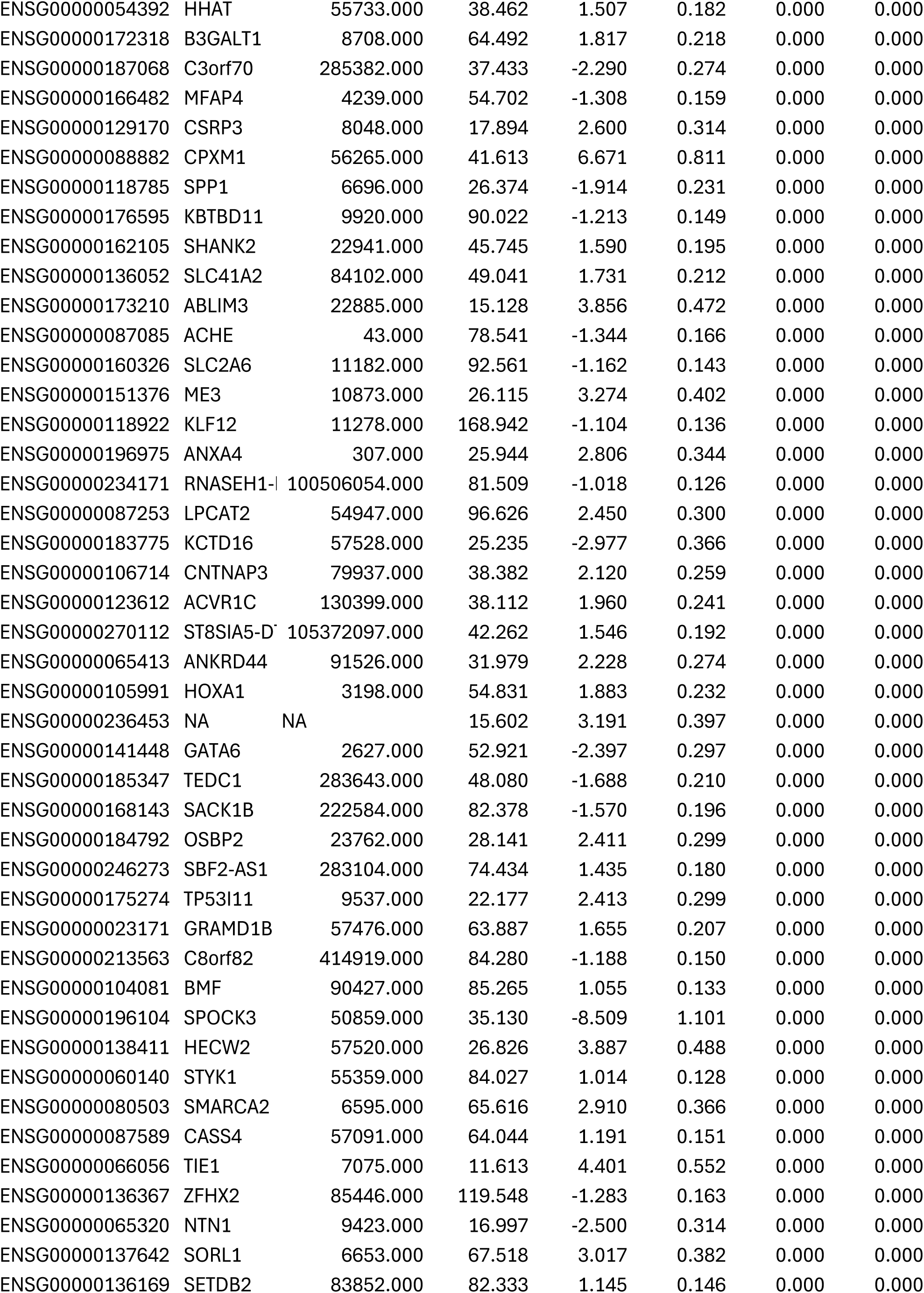
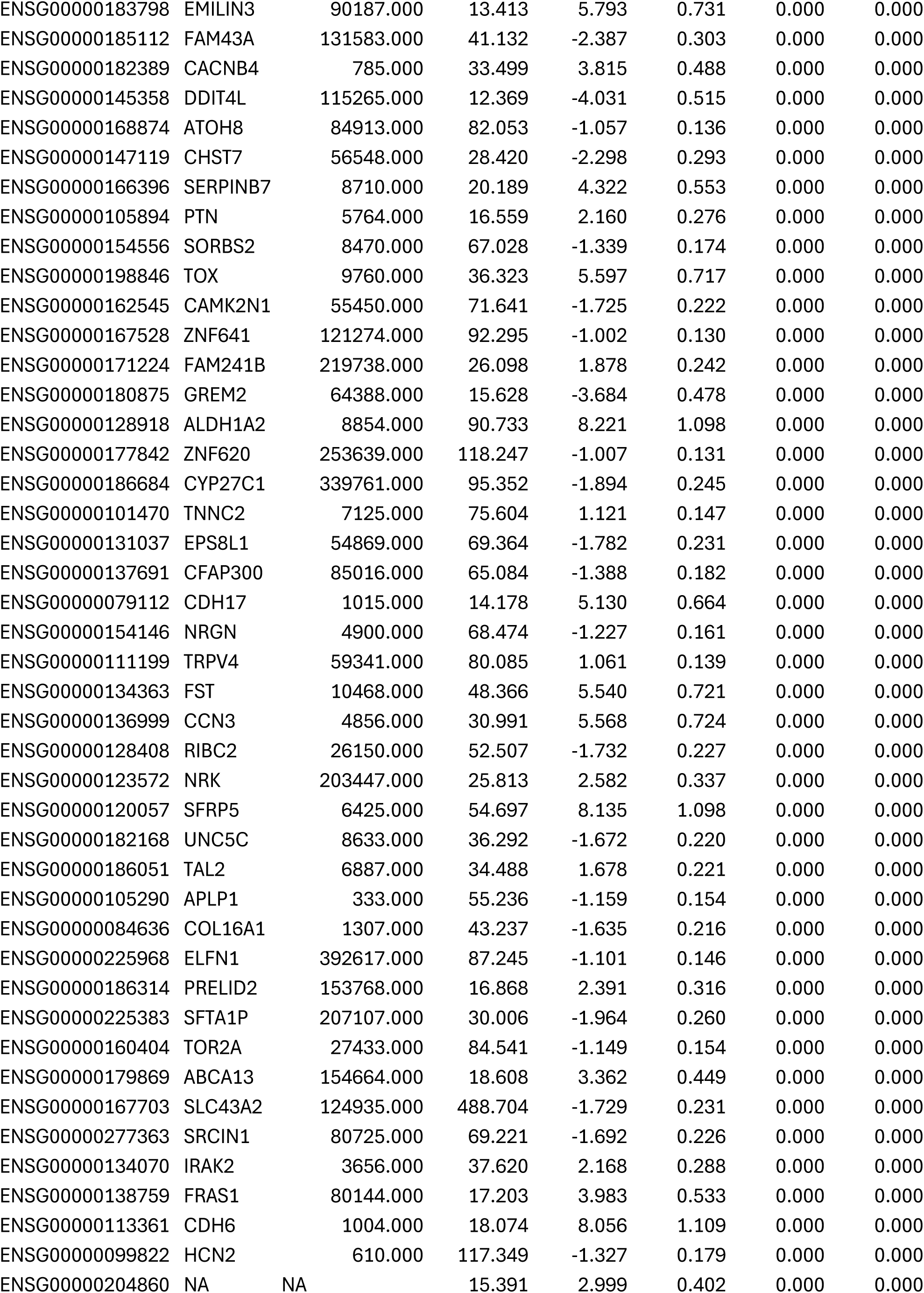
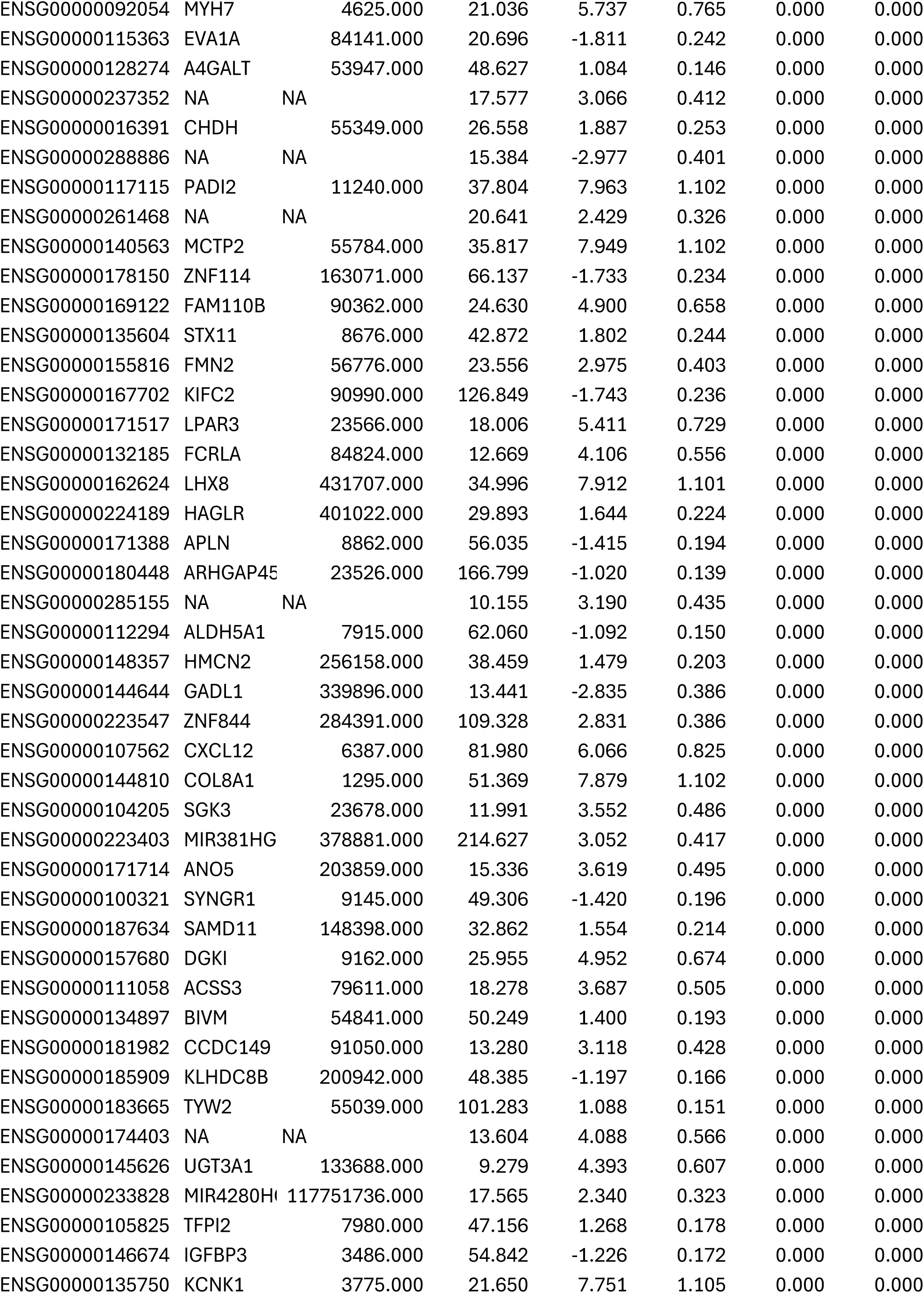
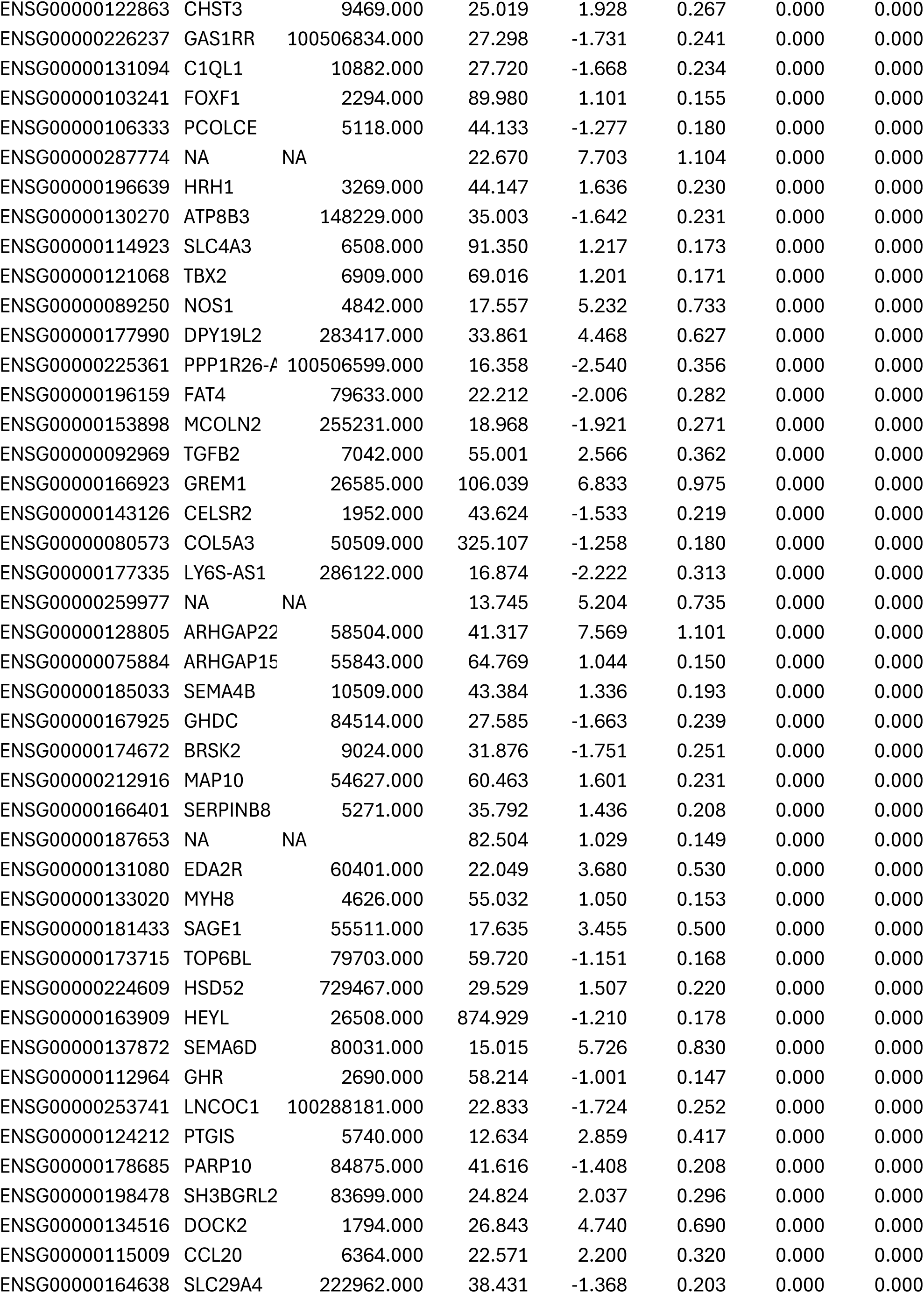
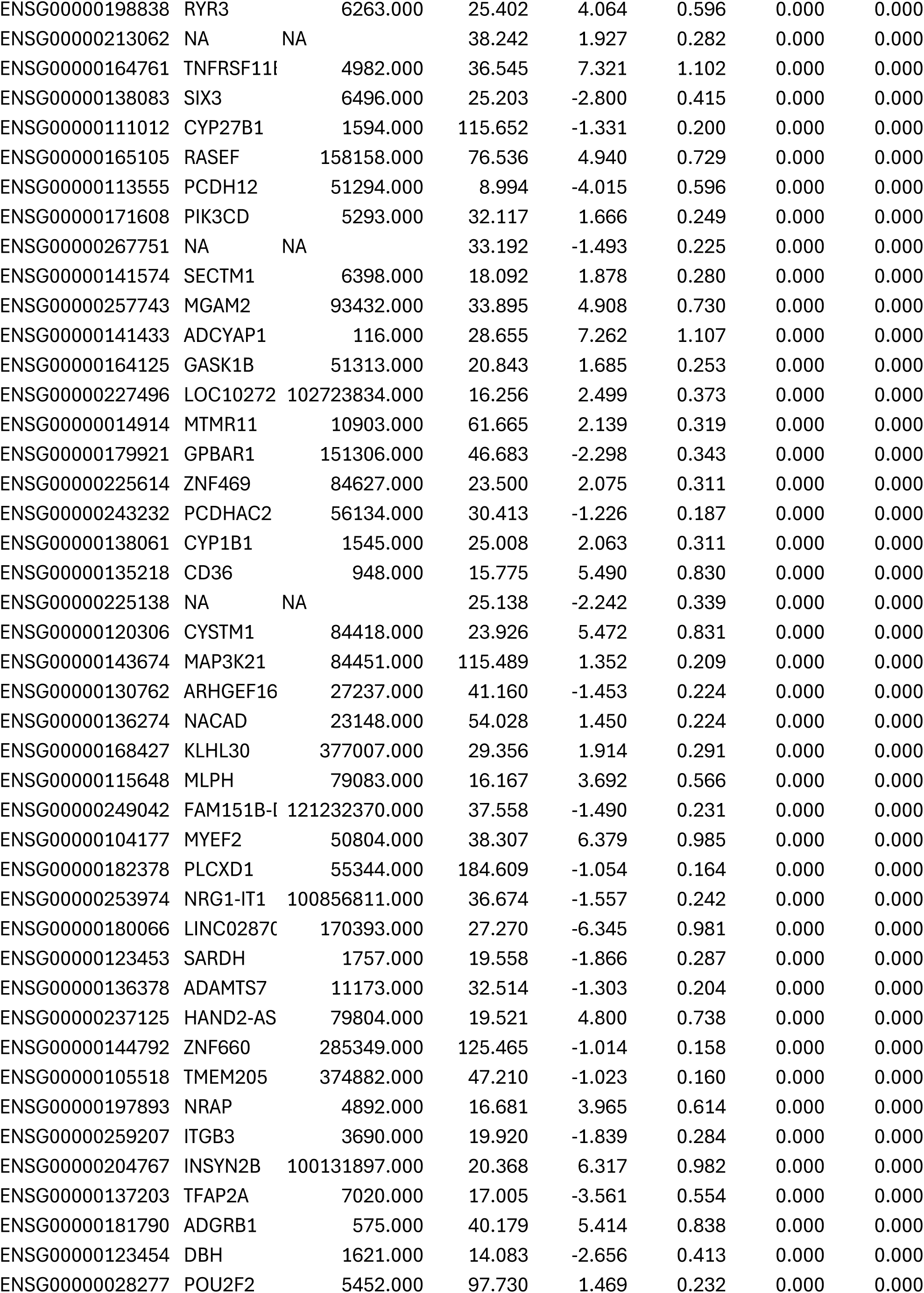
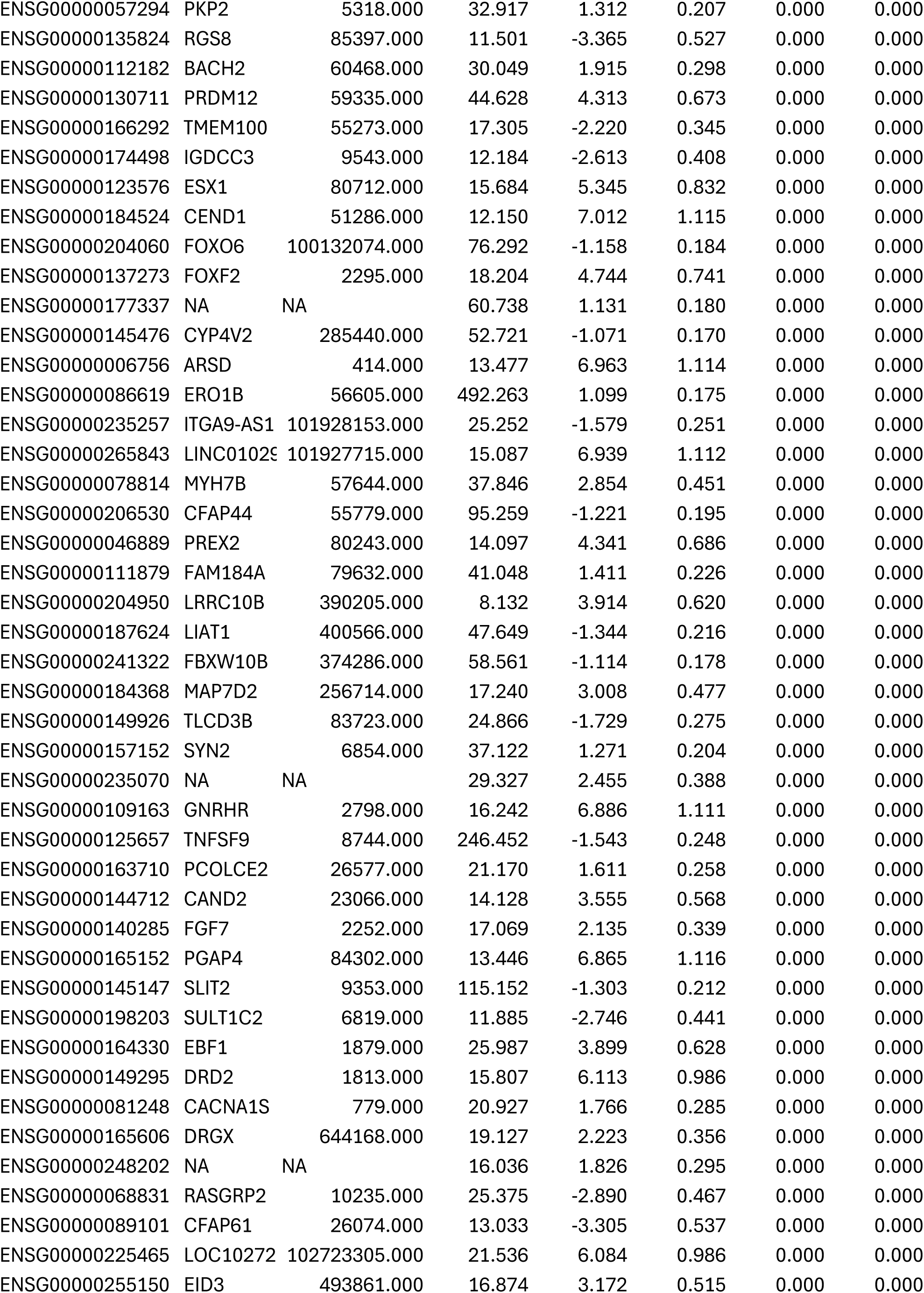
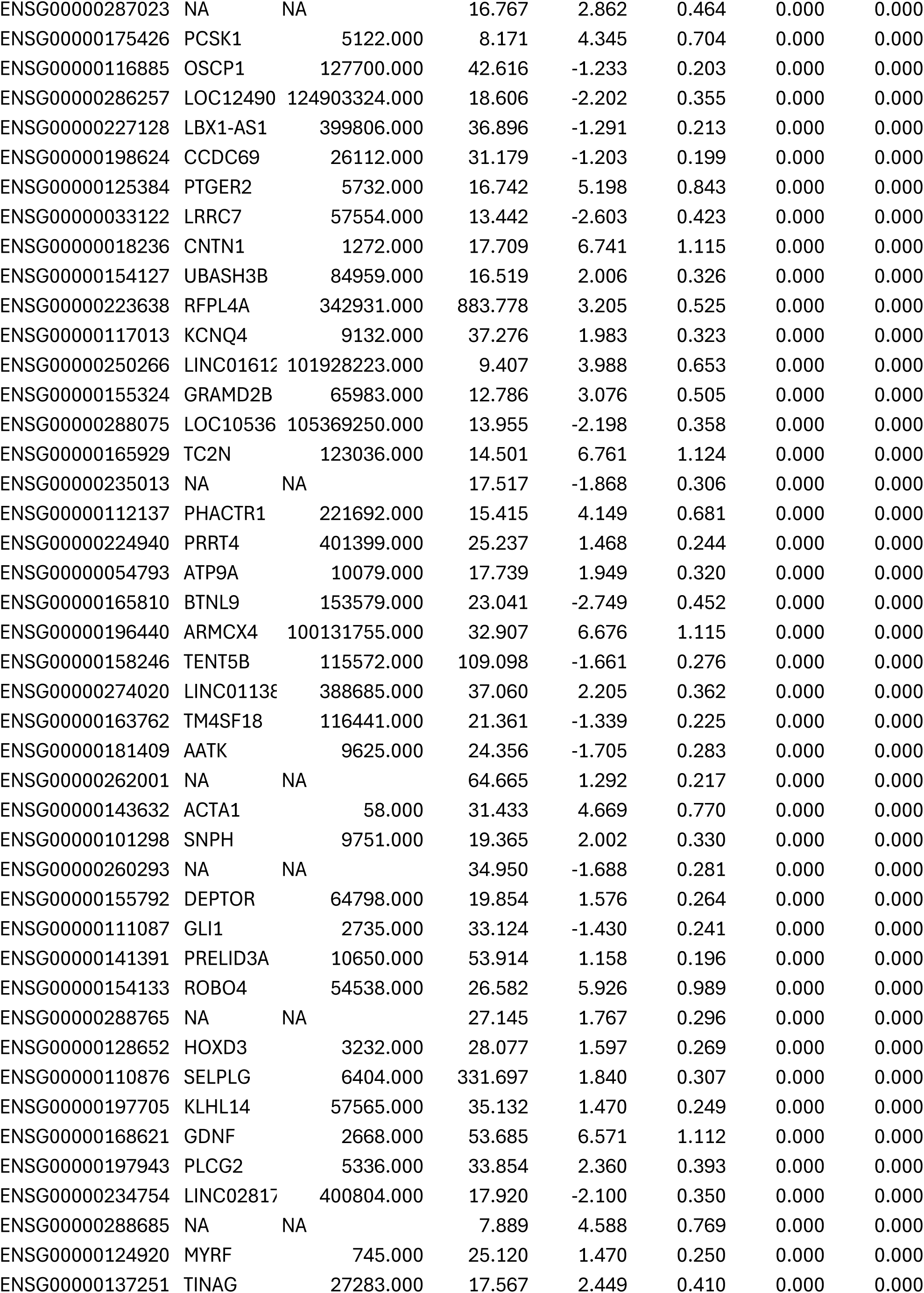
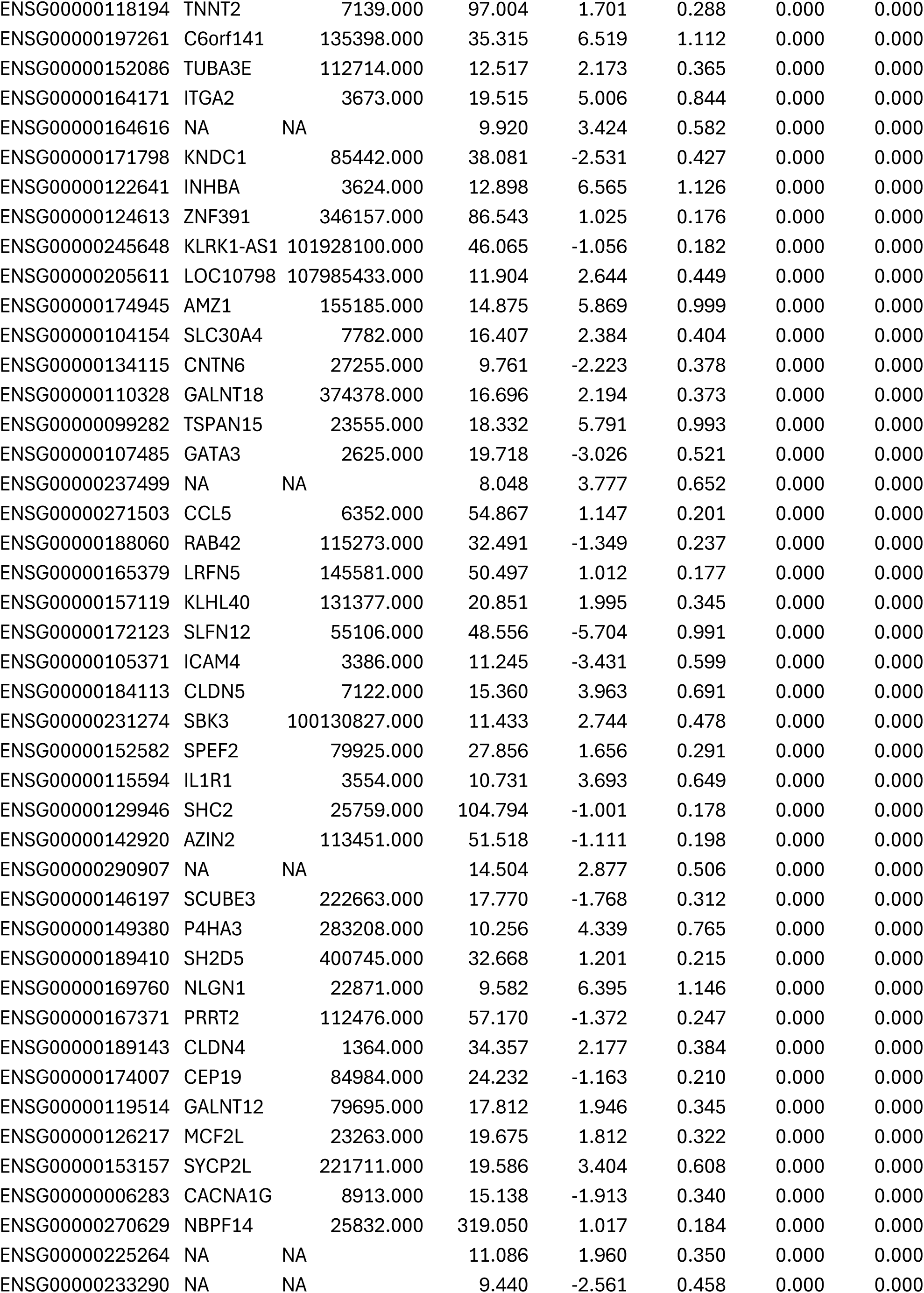
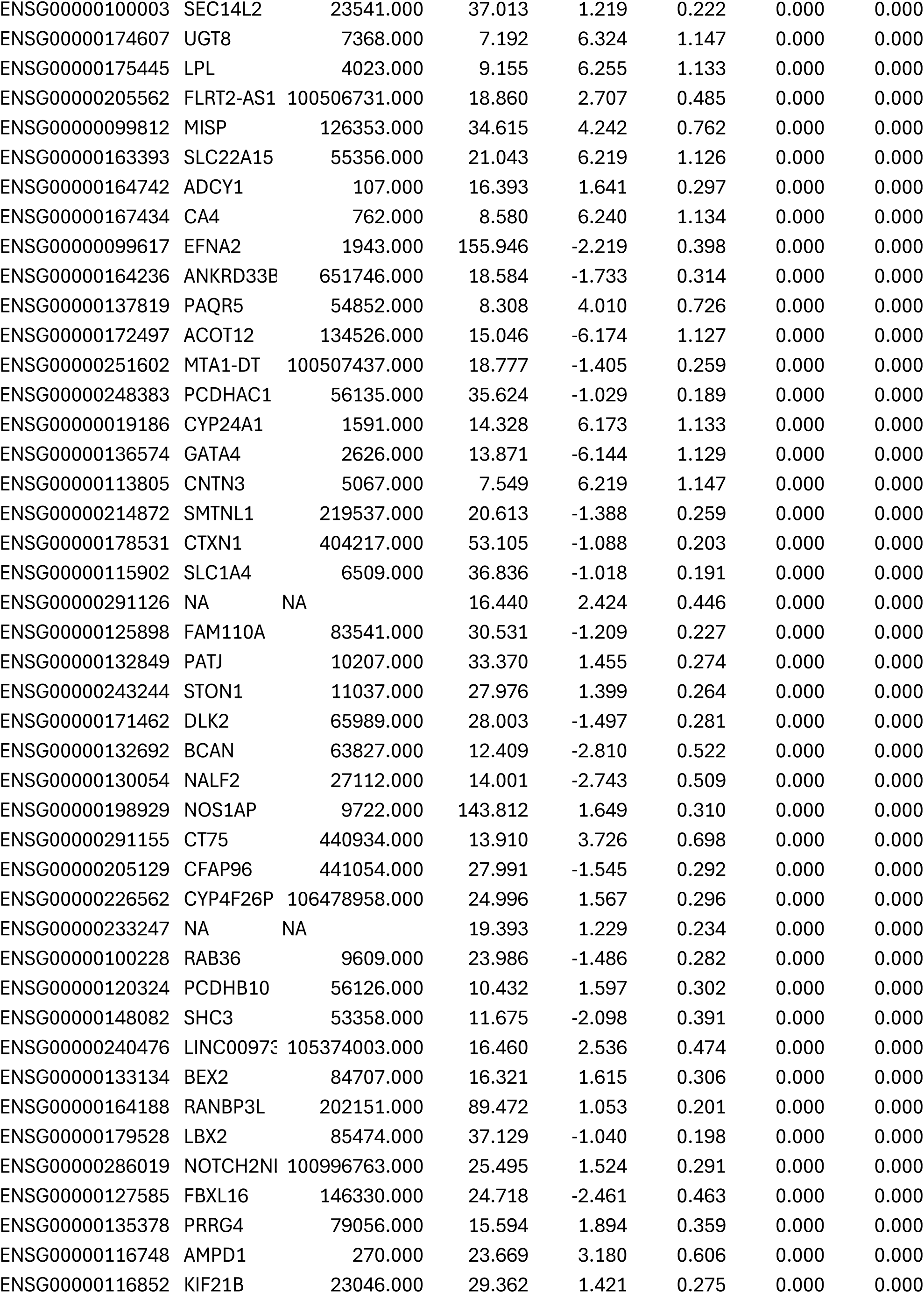
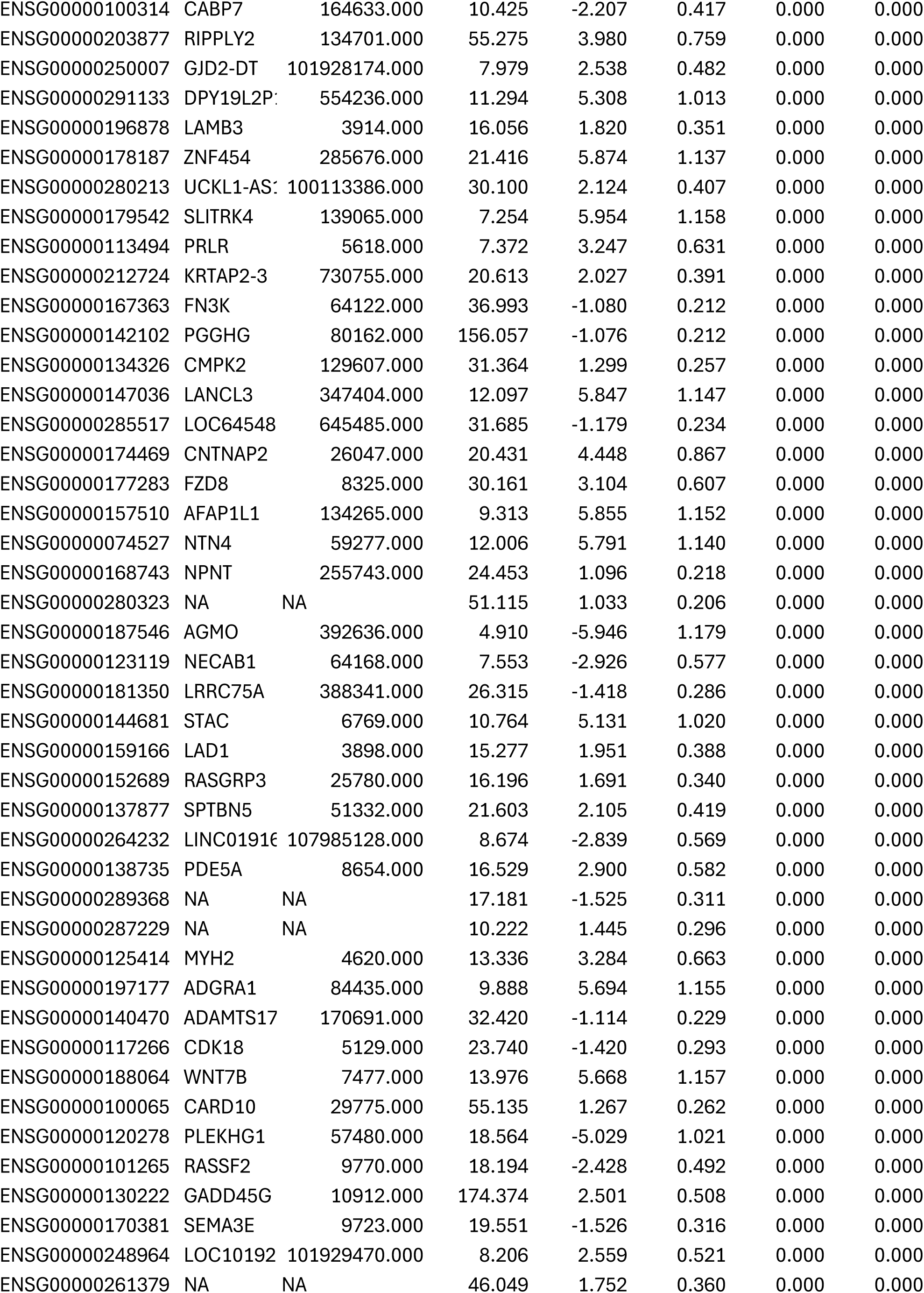
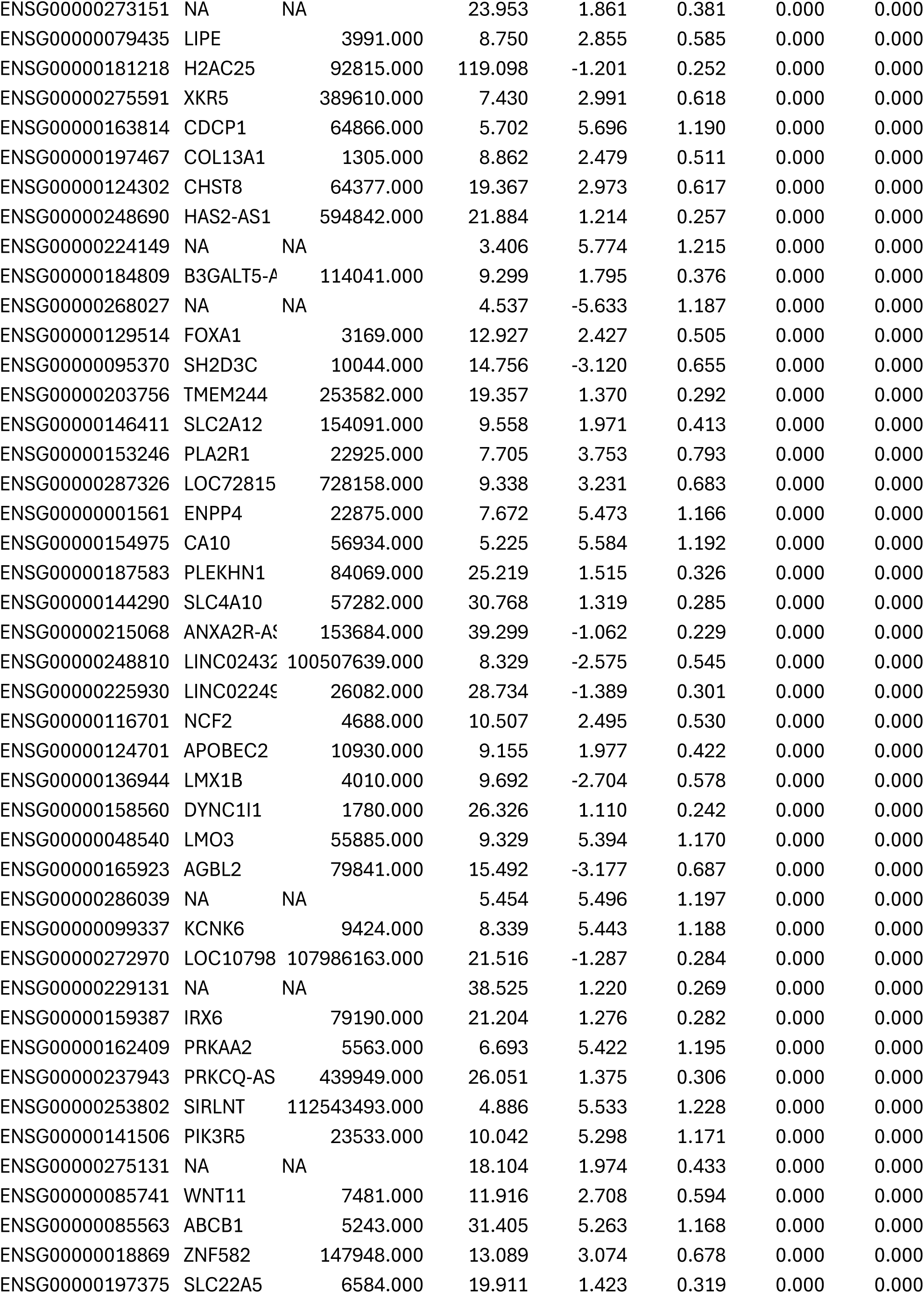
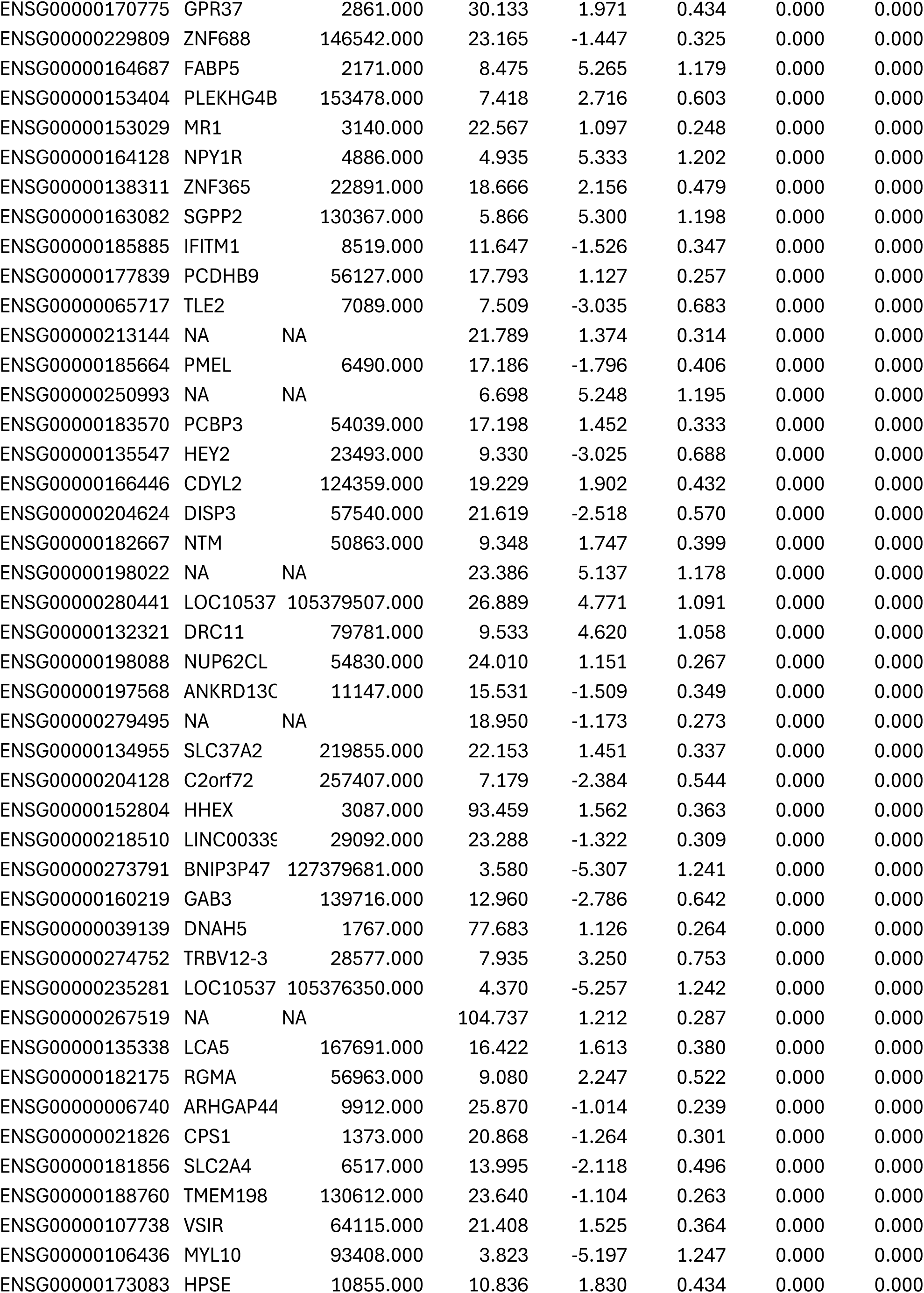
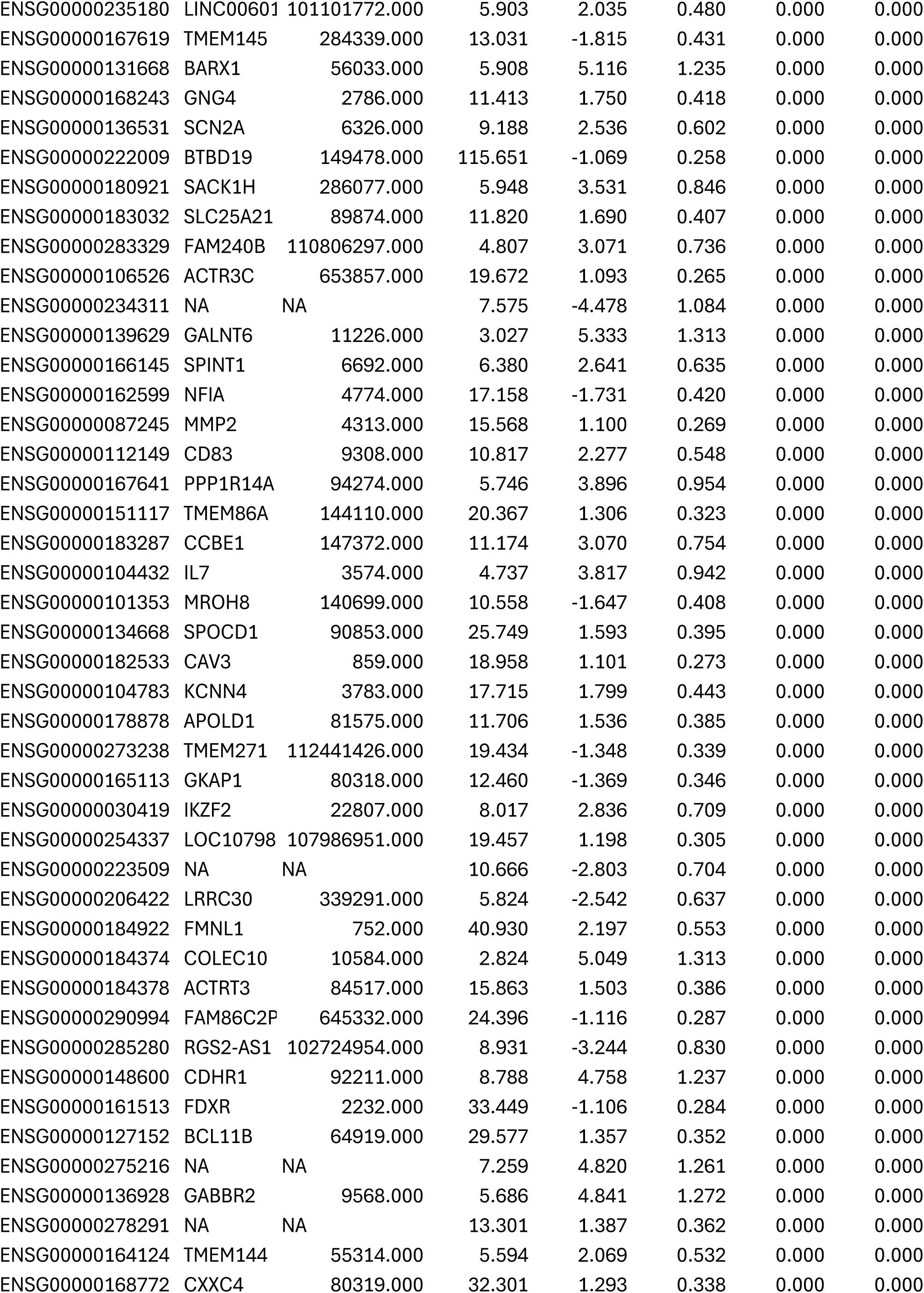
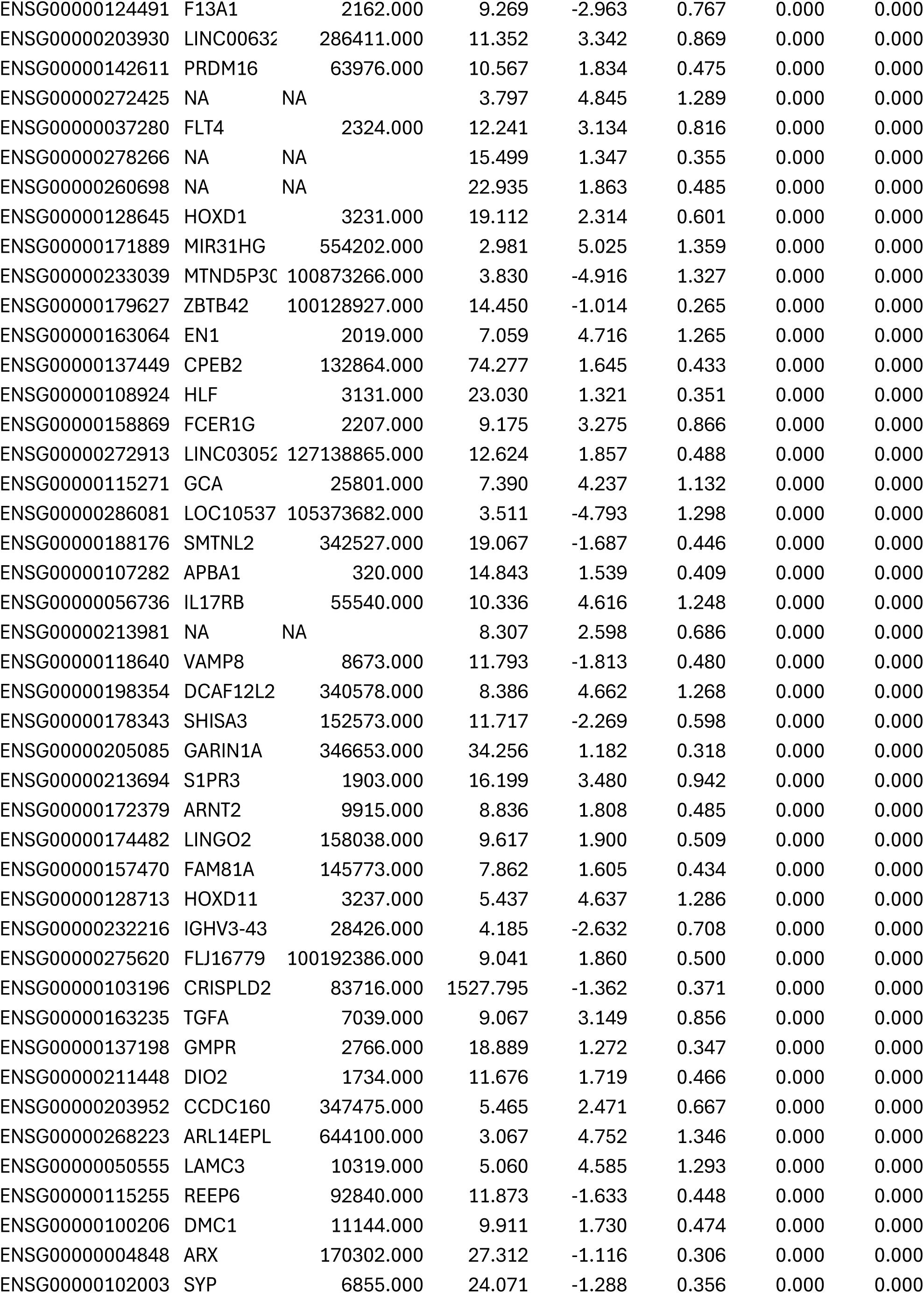
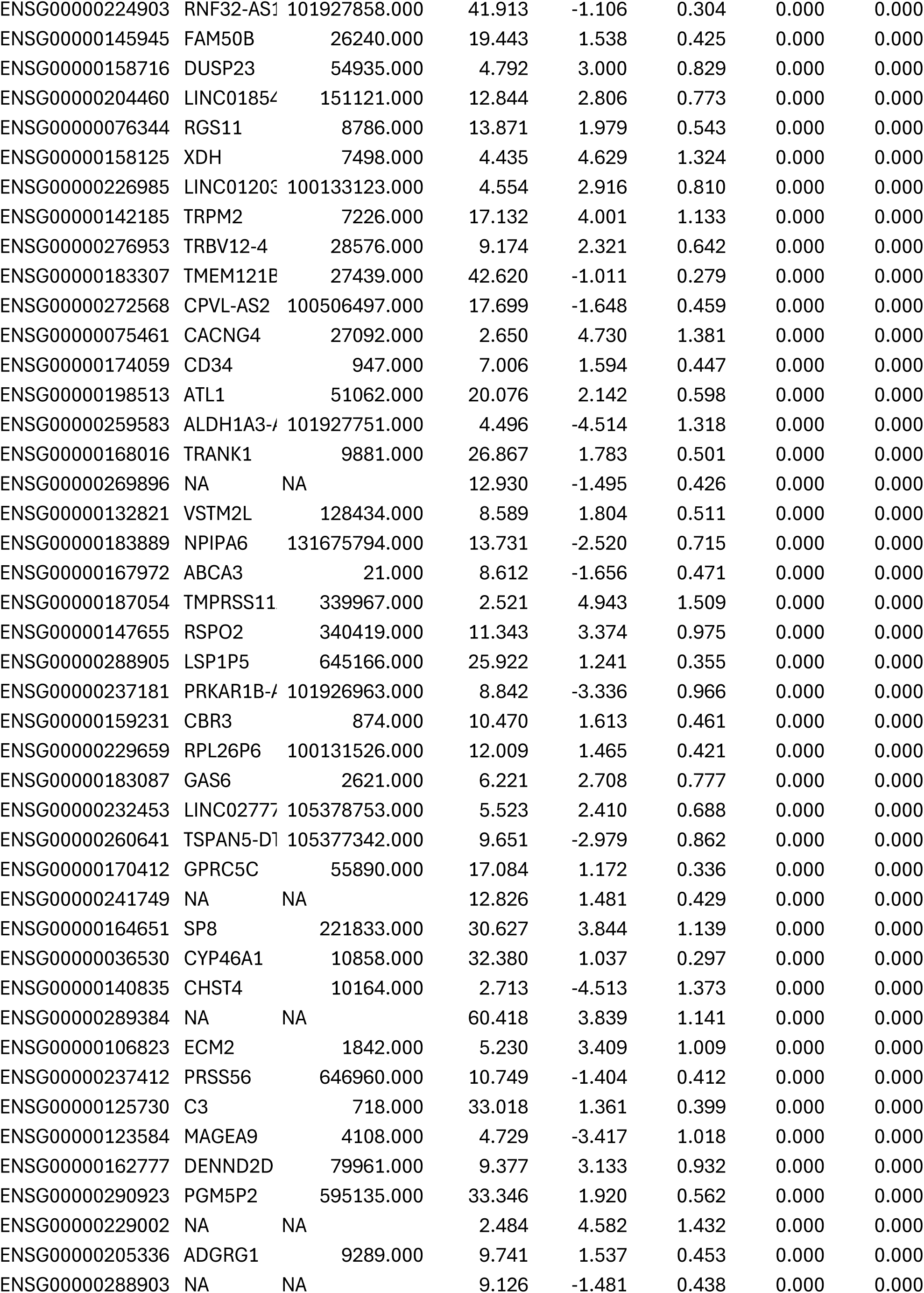
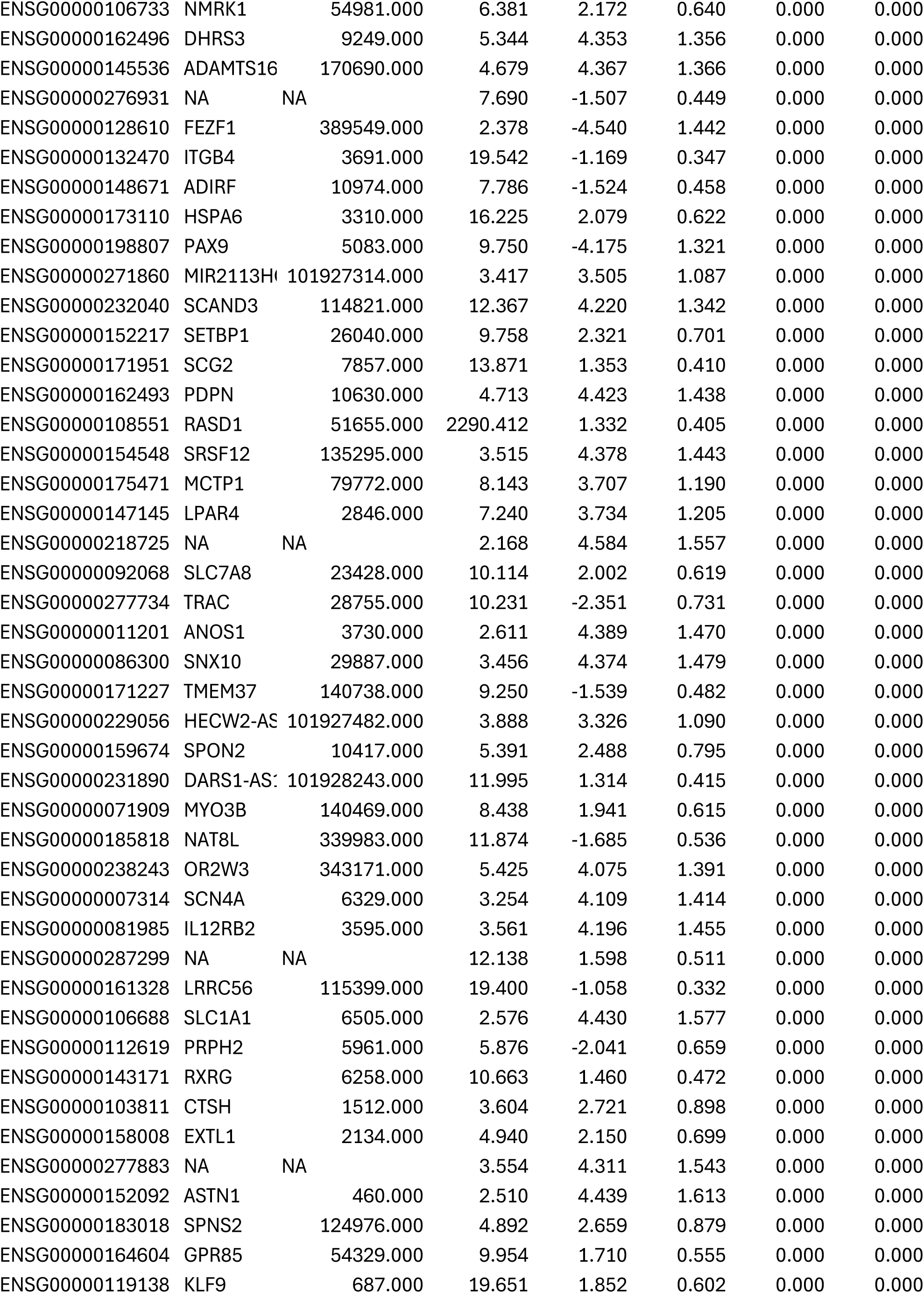
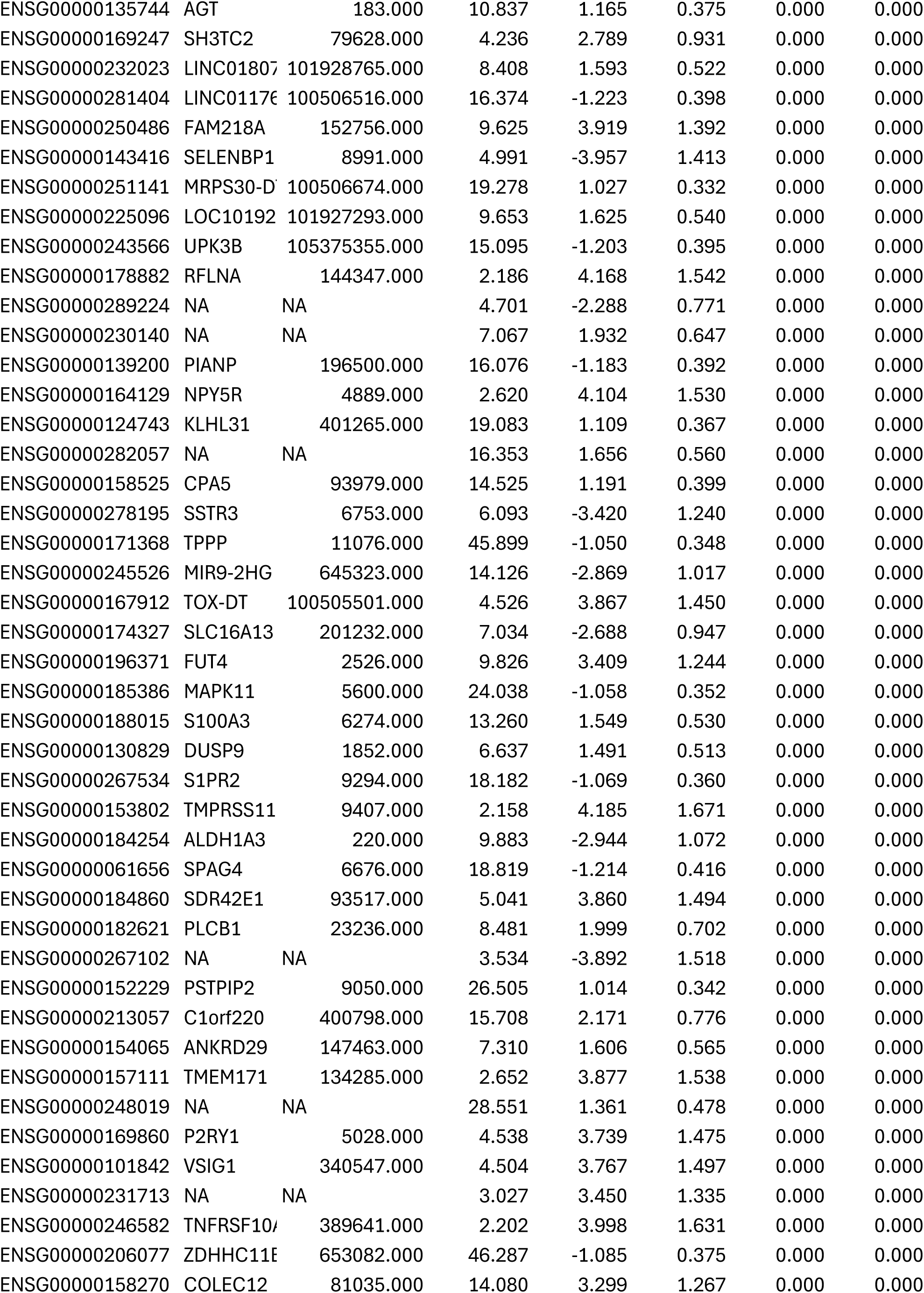
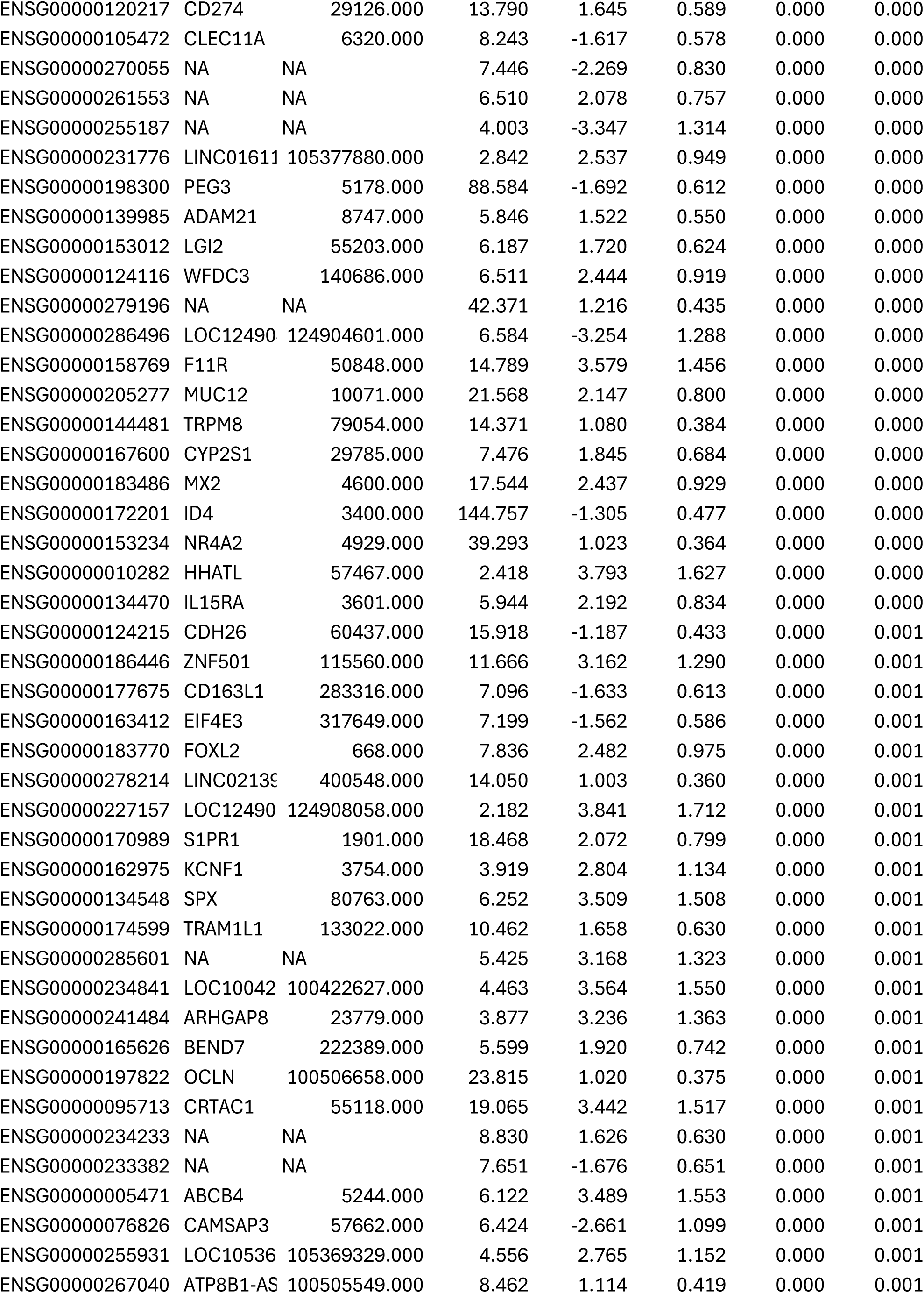
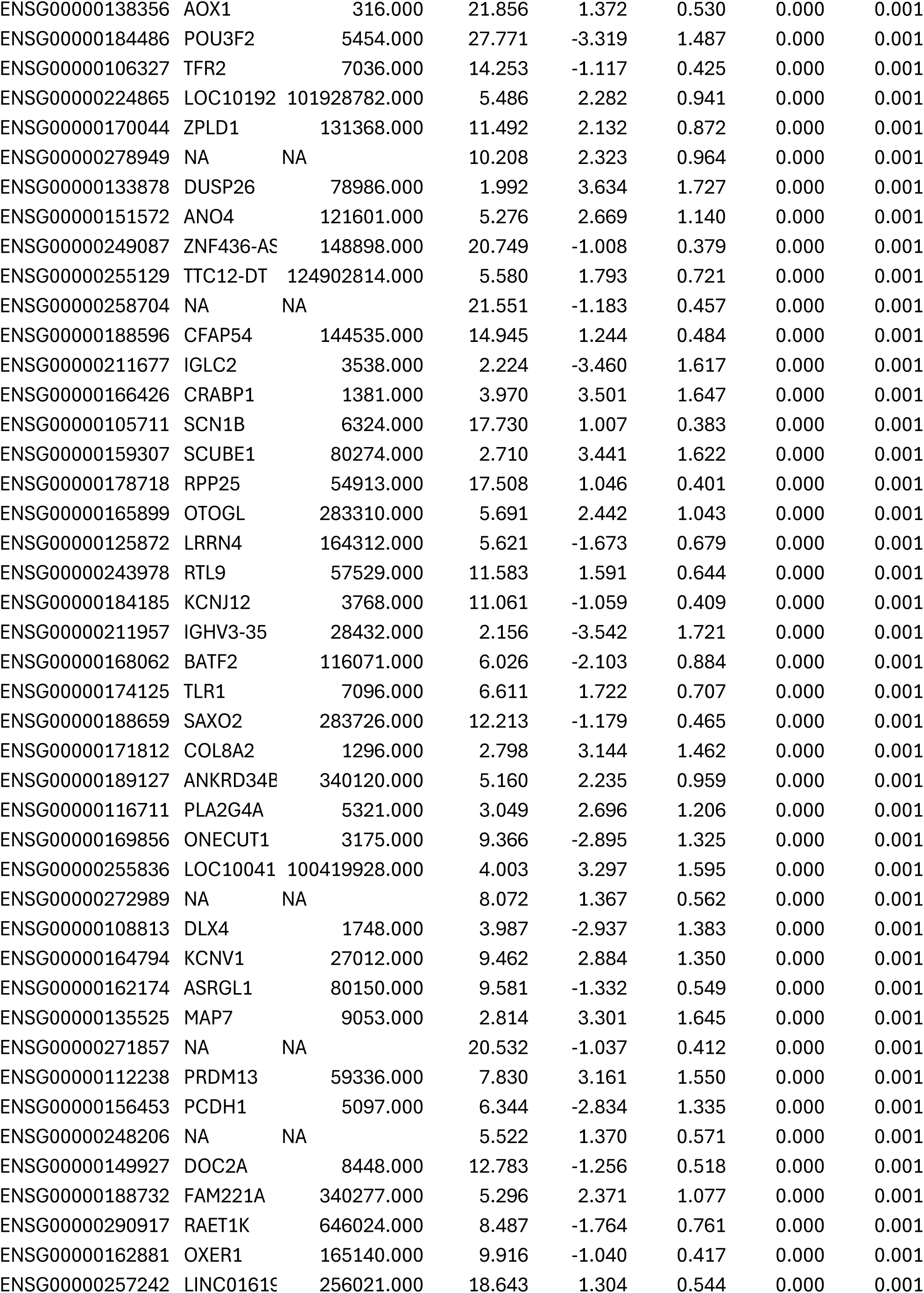
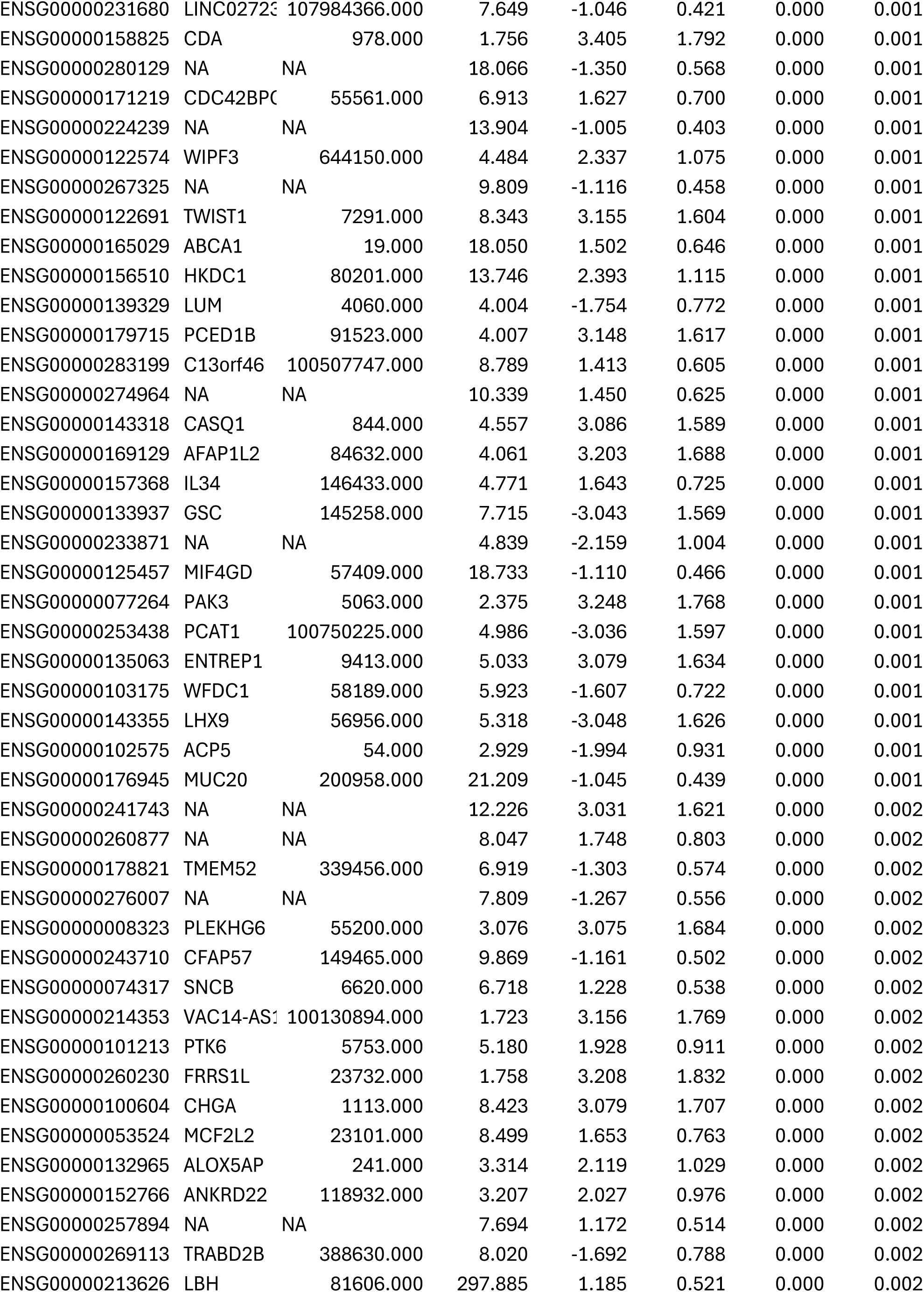
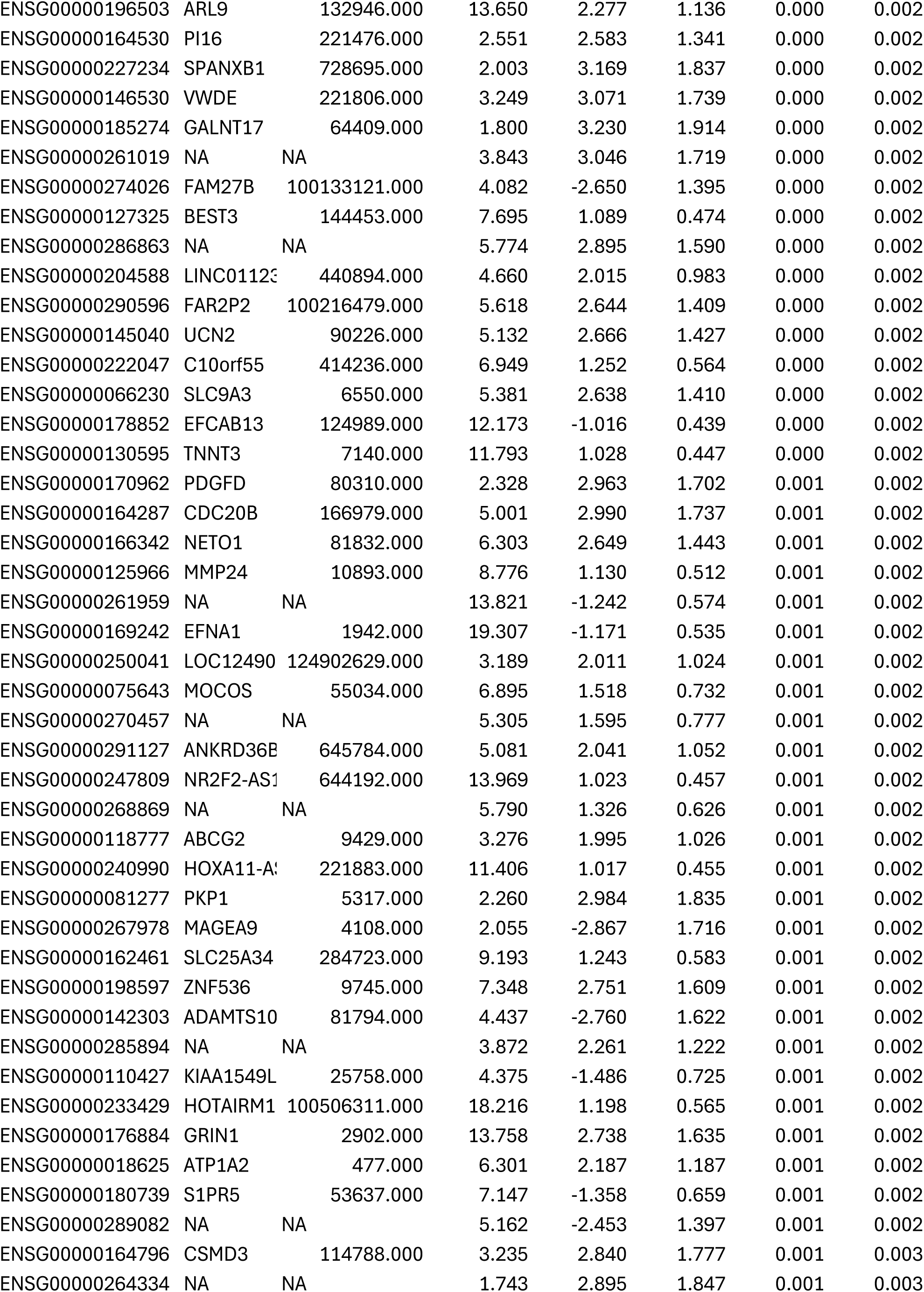
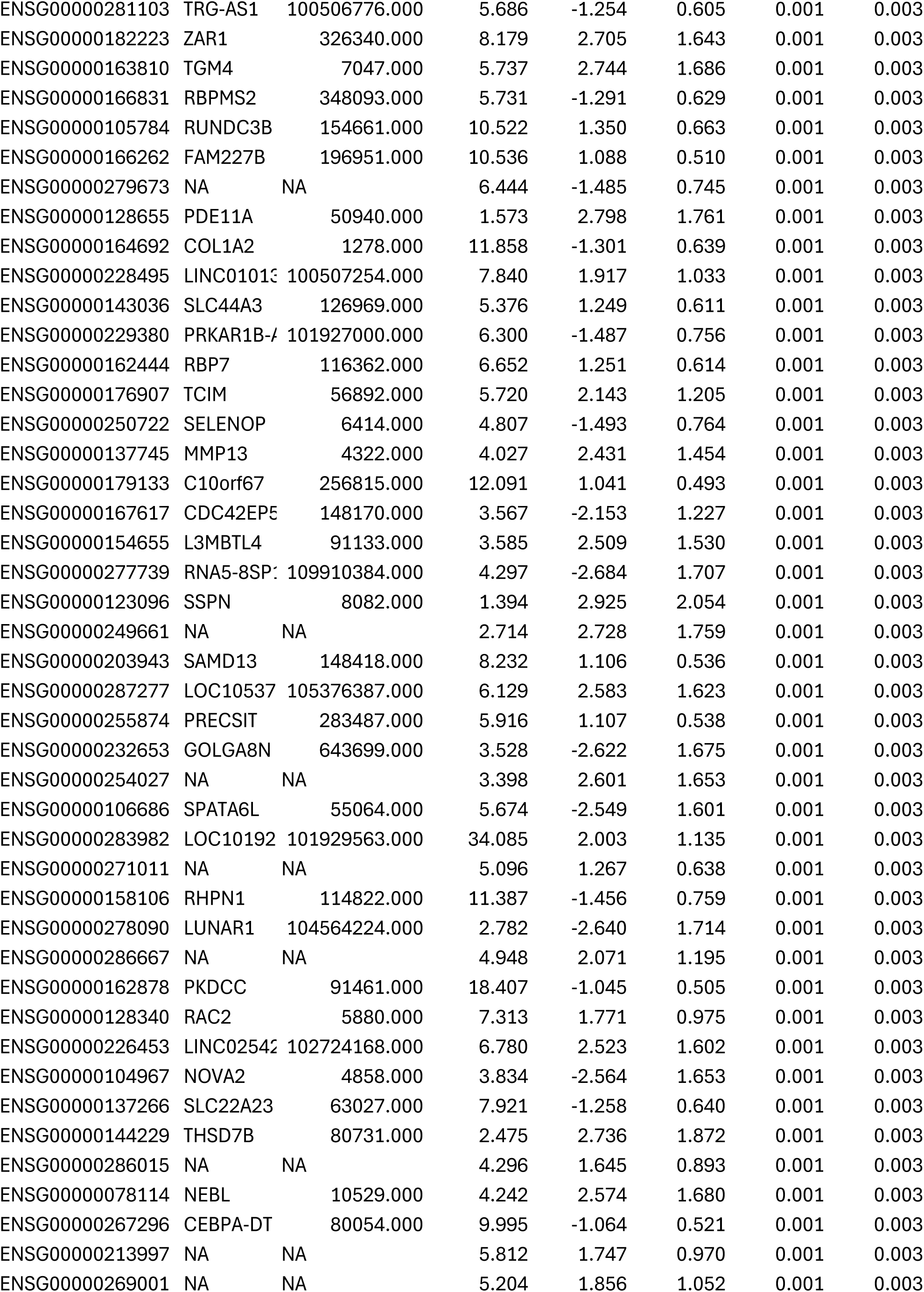
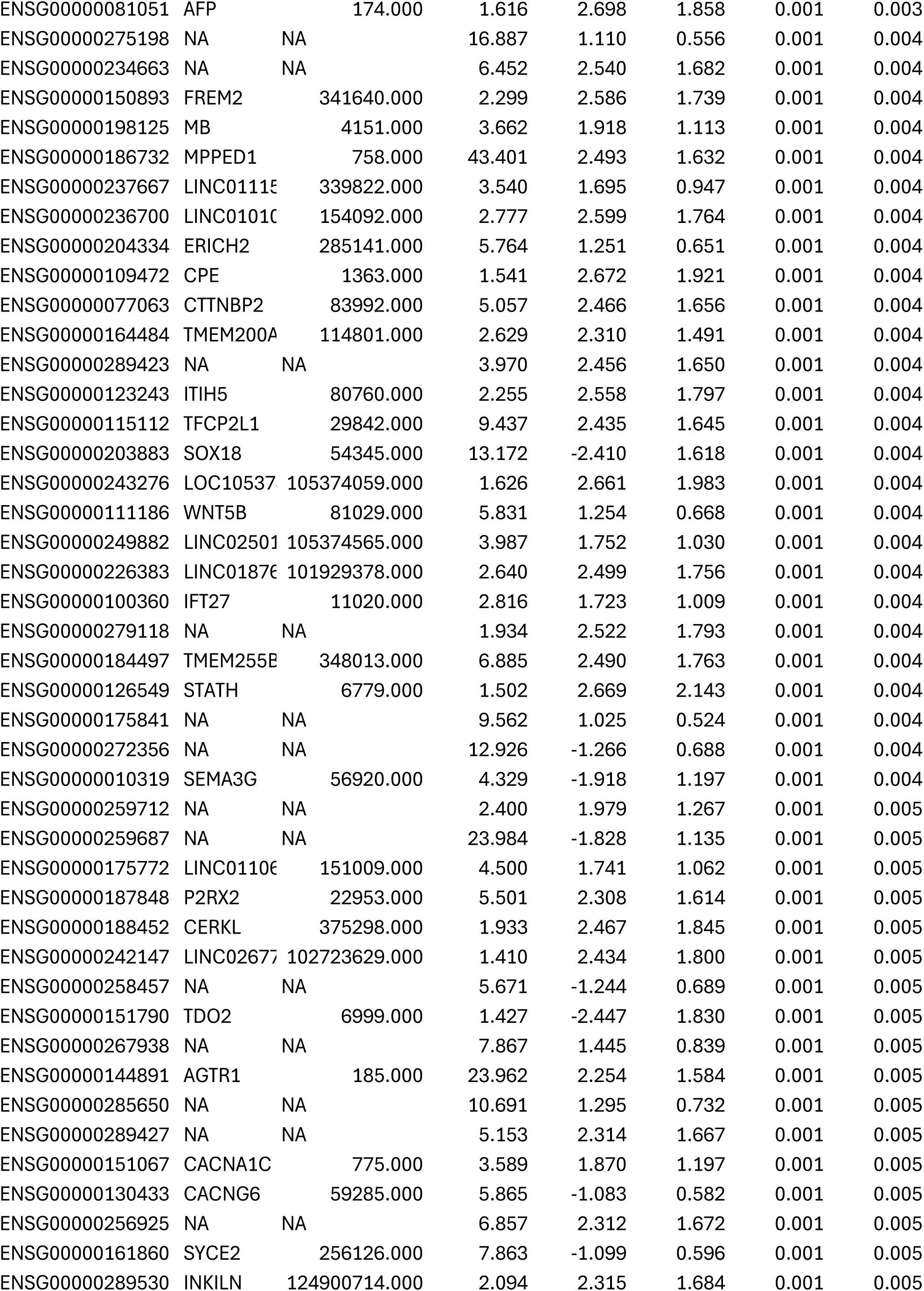
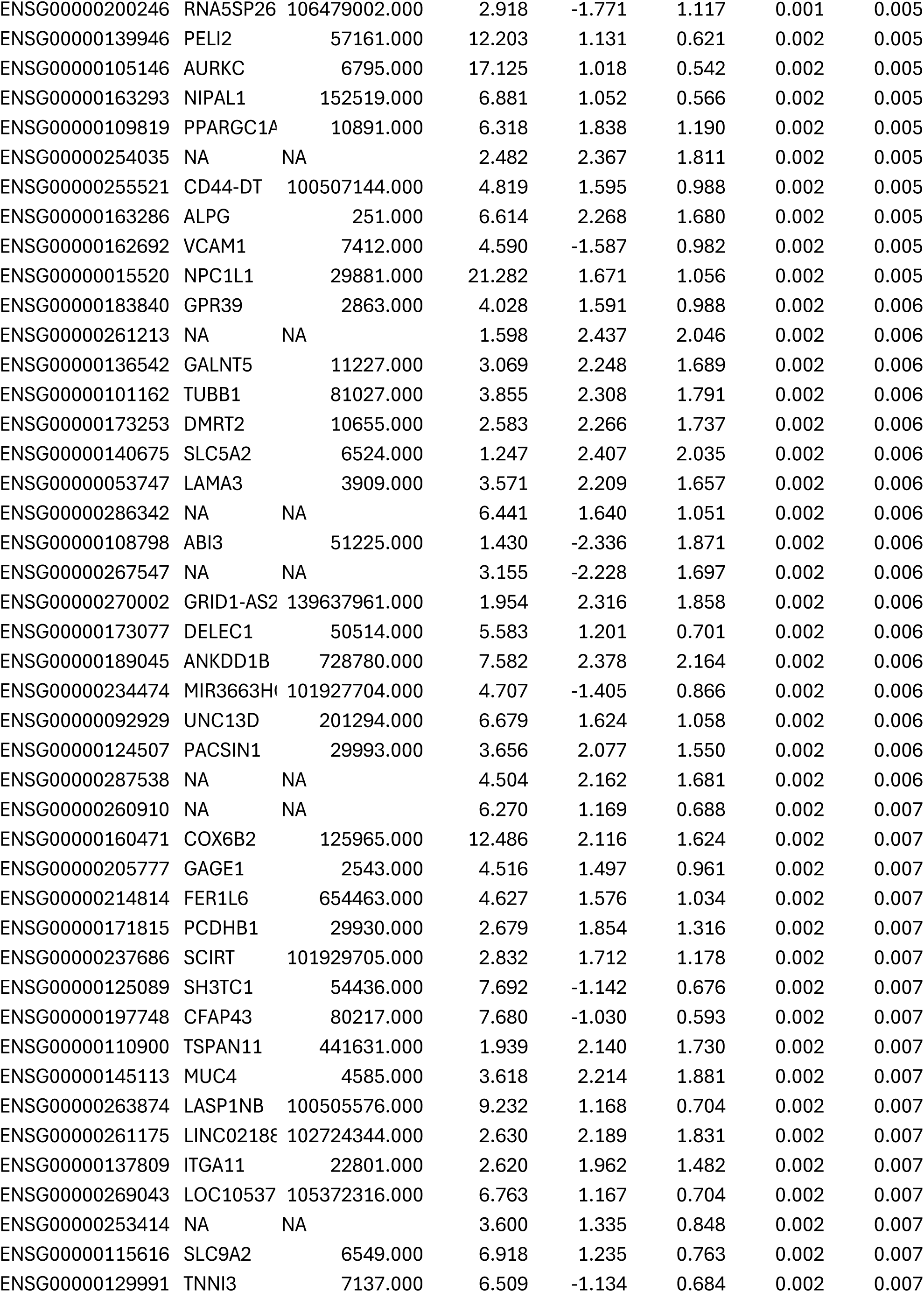
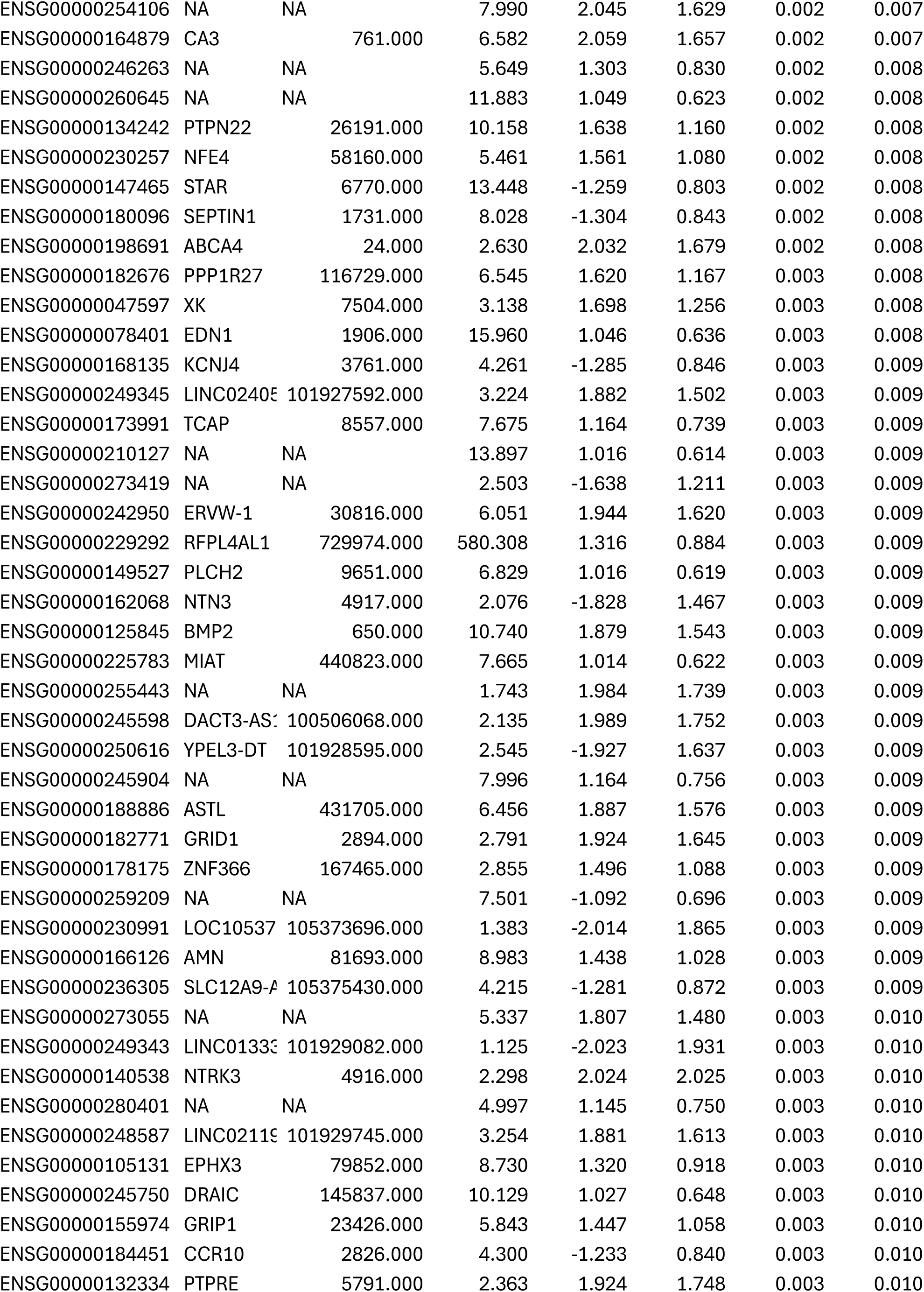
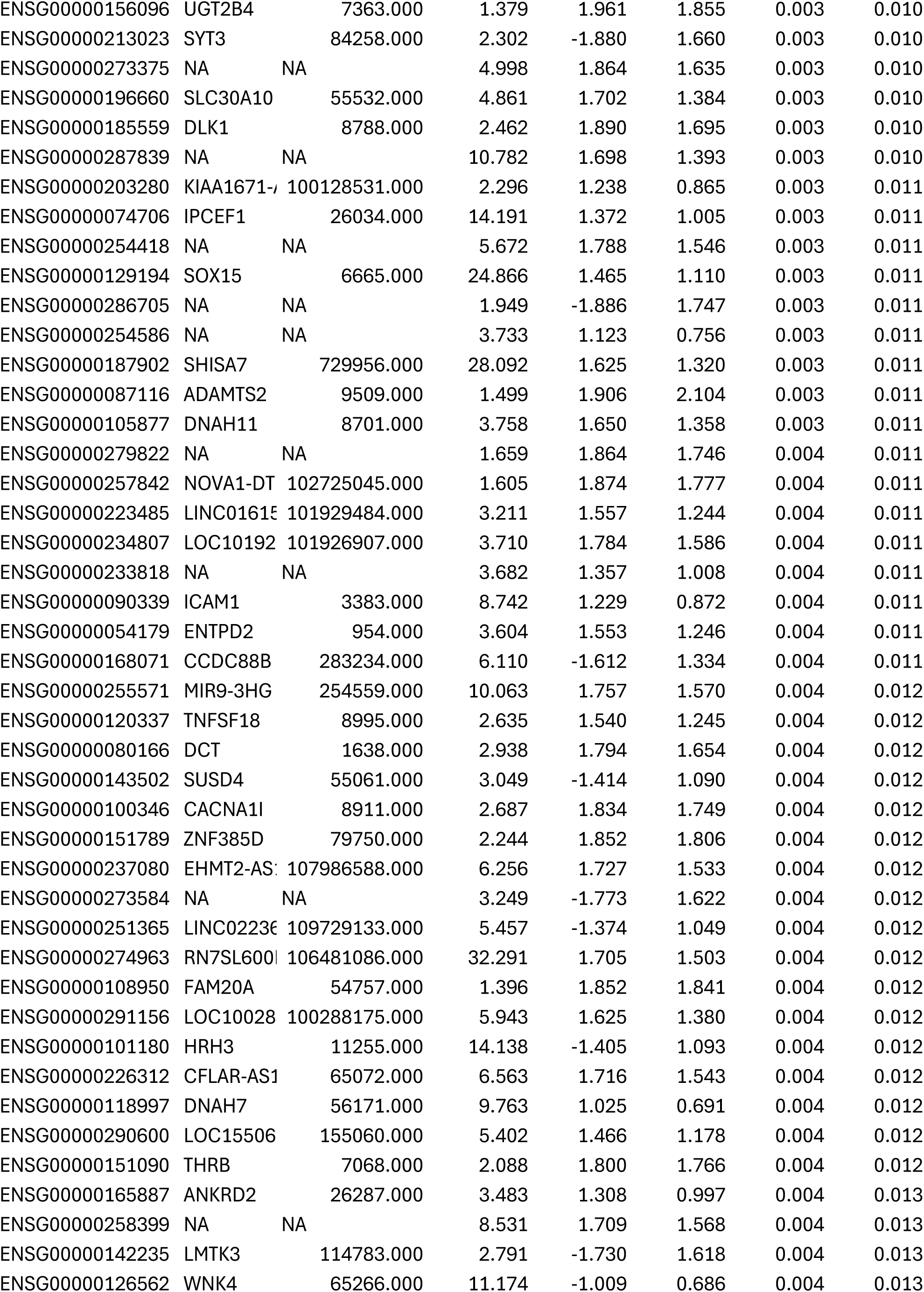
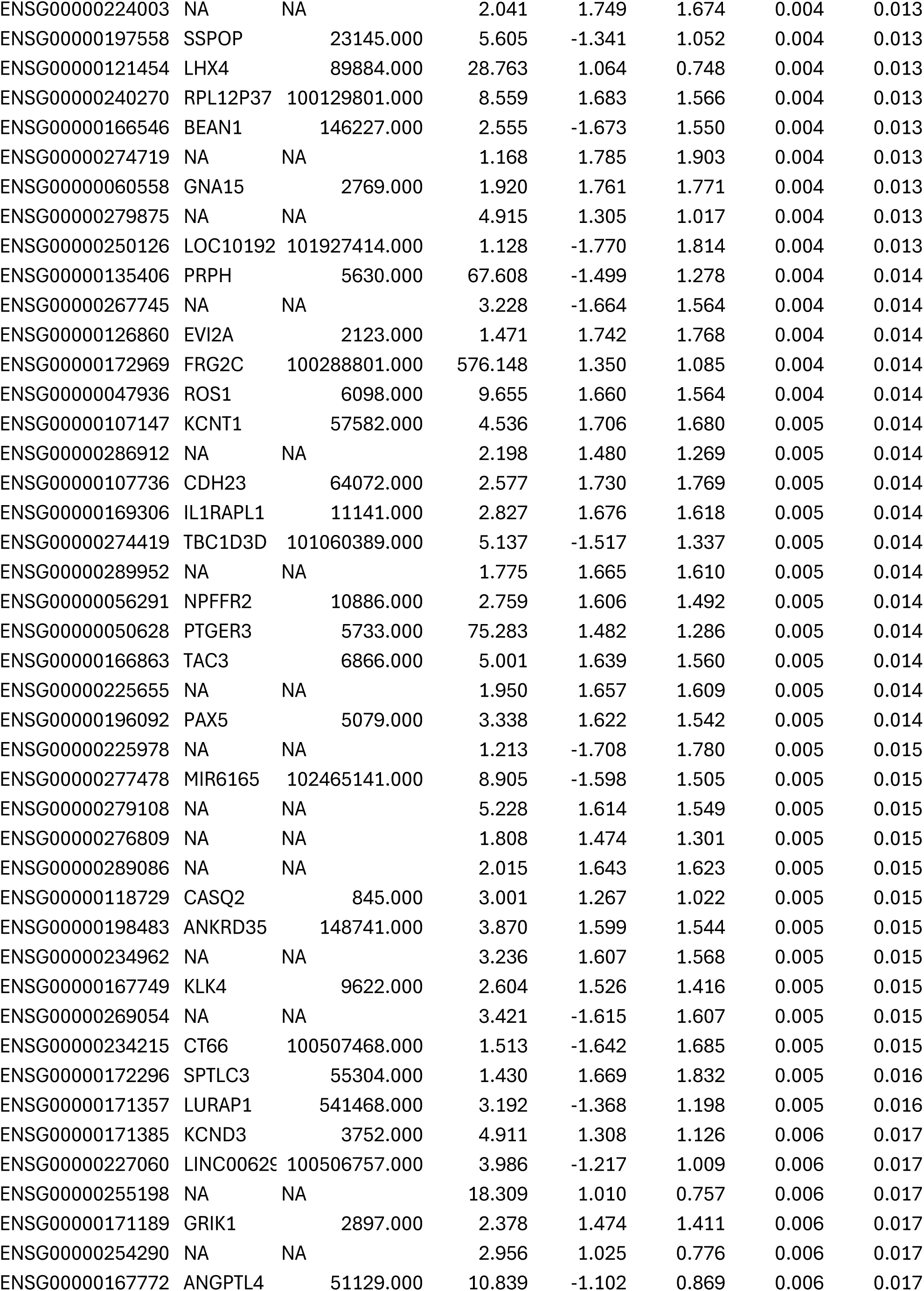
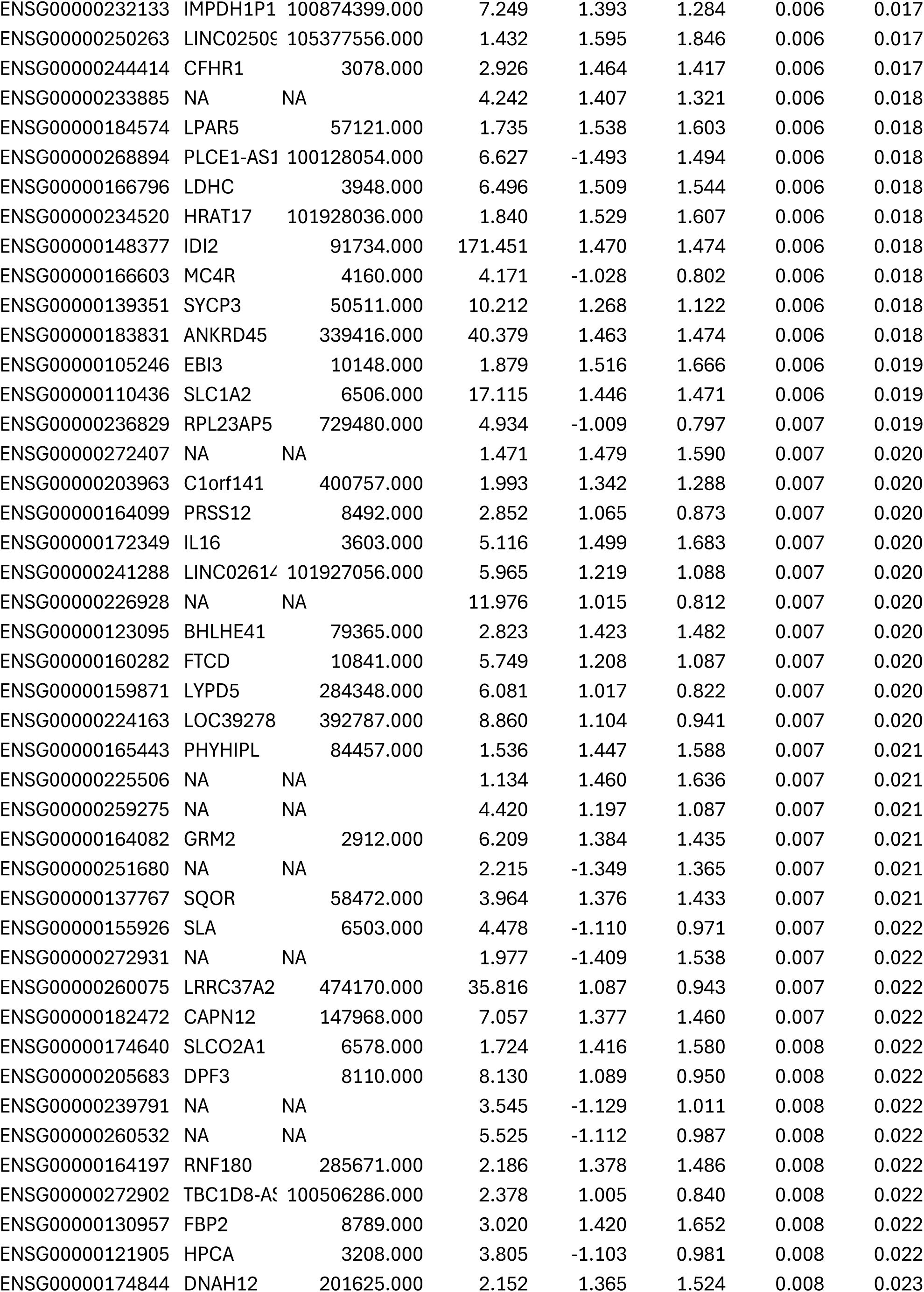
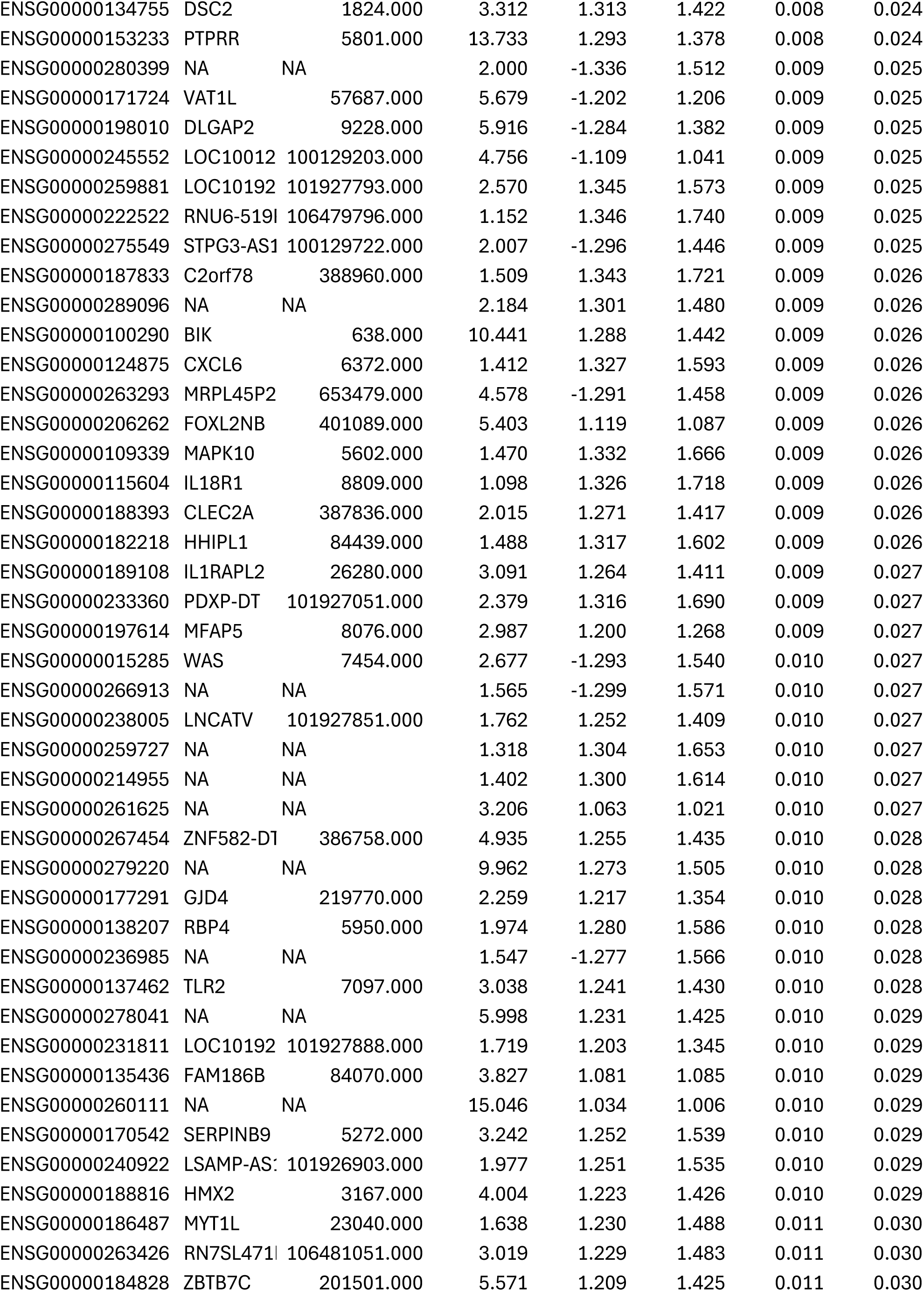
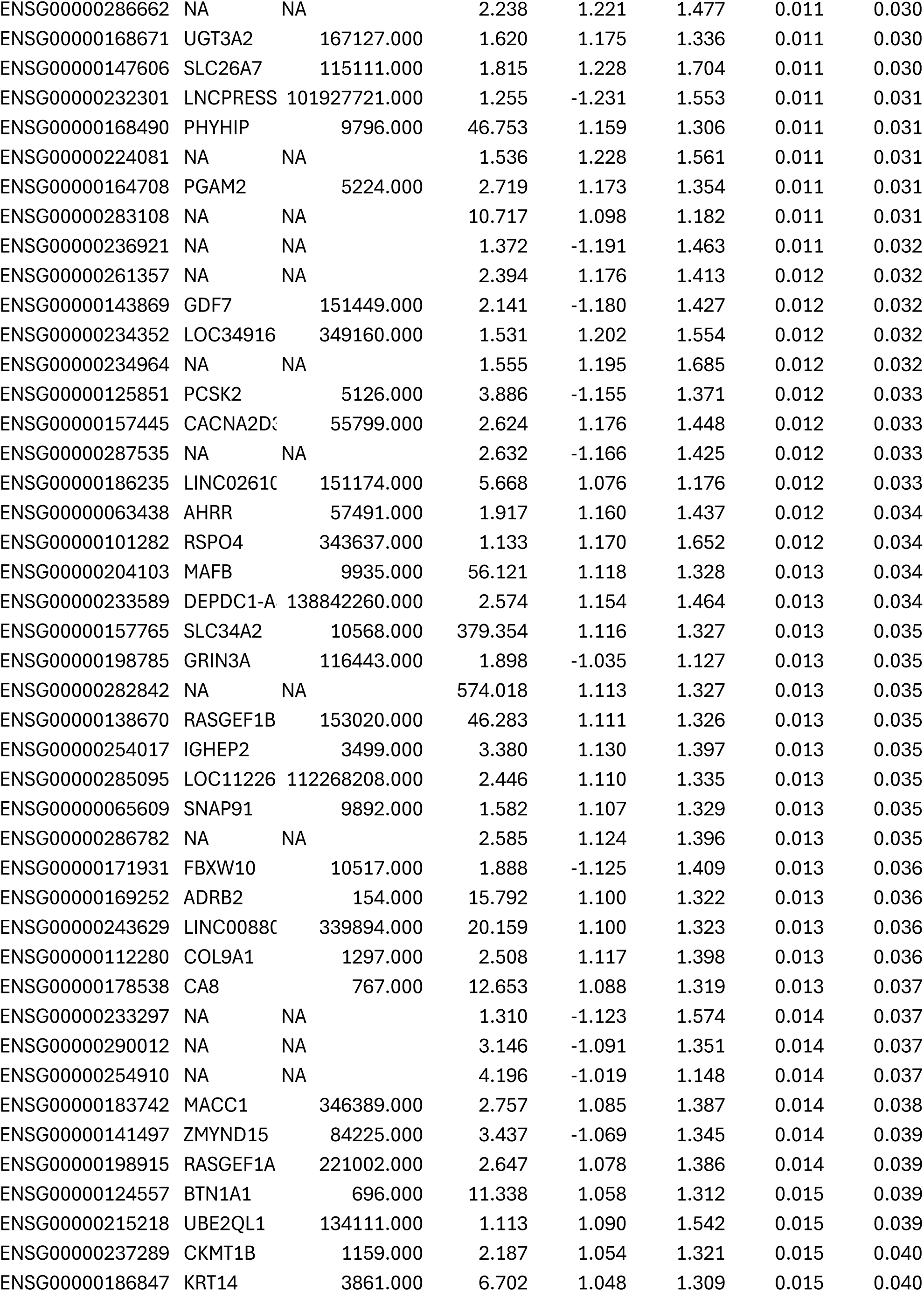
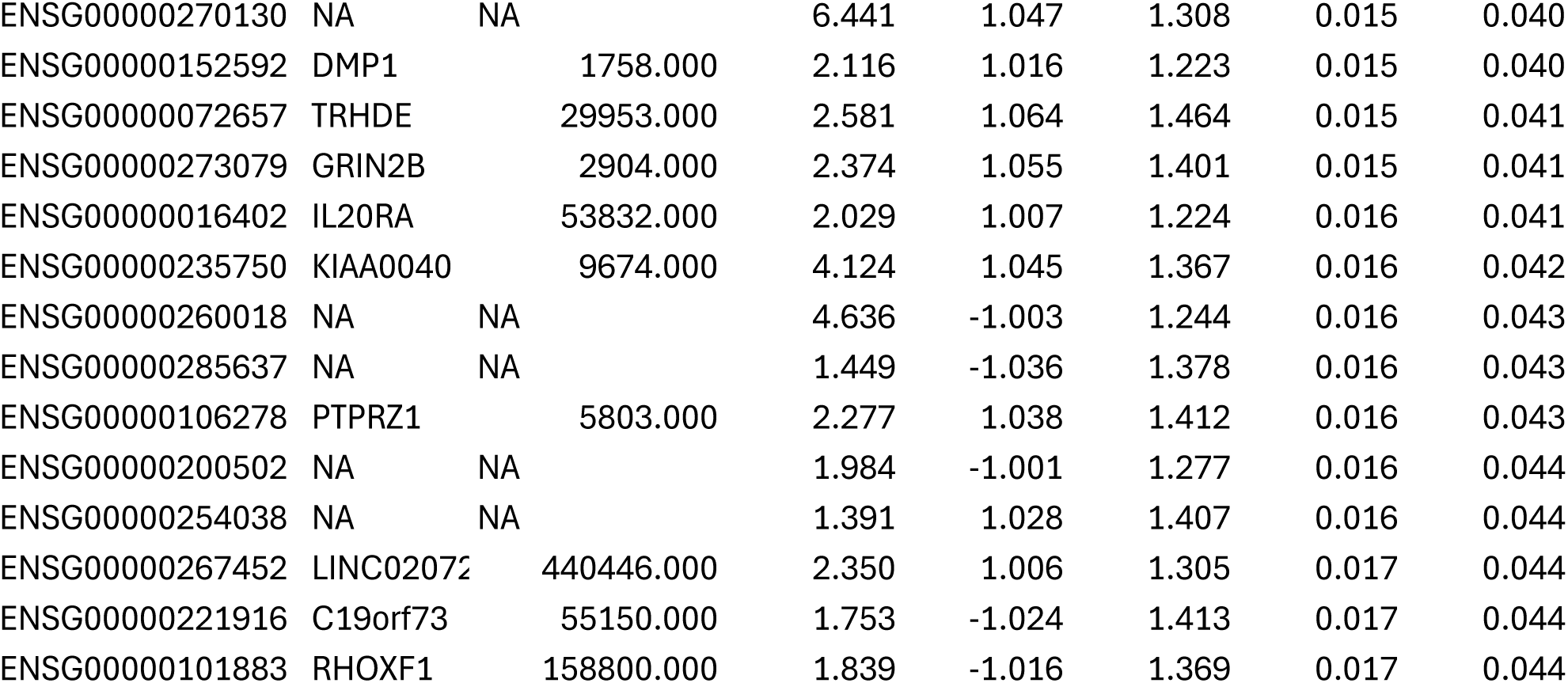

Significant DEGs during infection of RD:P3F with G47Δ at 6 hours post infection versus Mock RD:P3F.

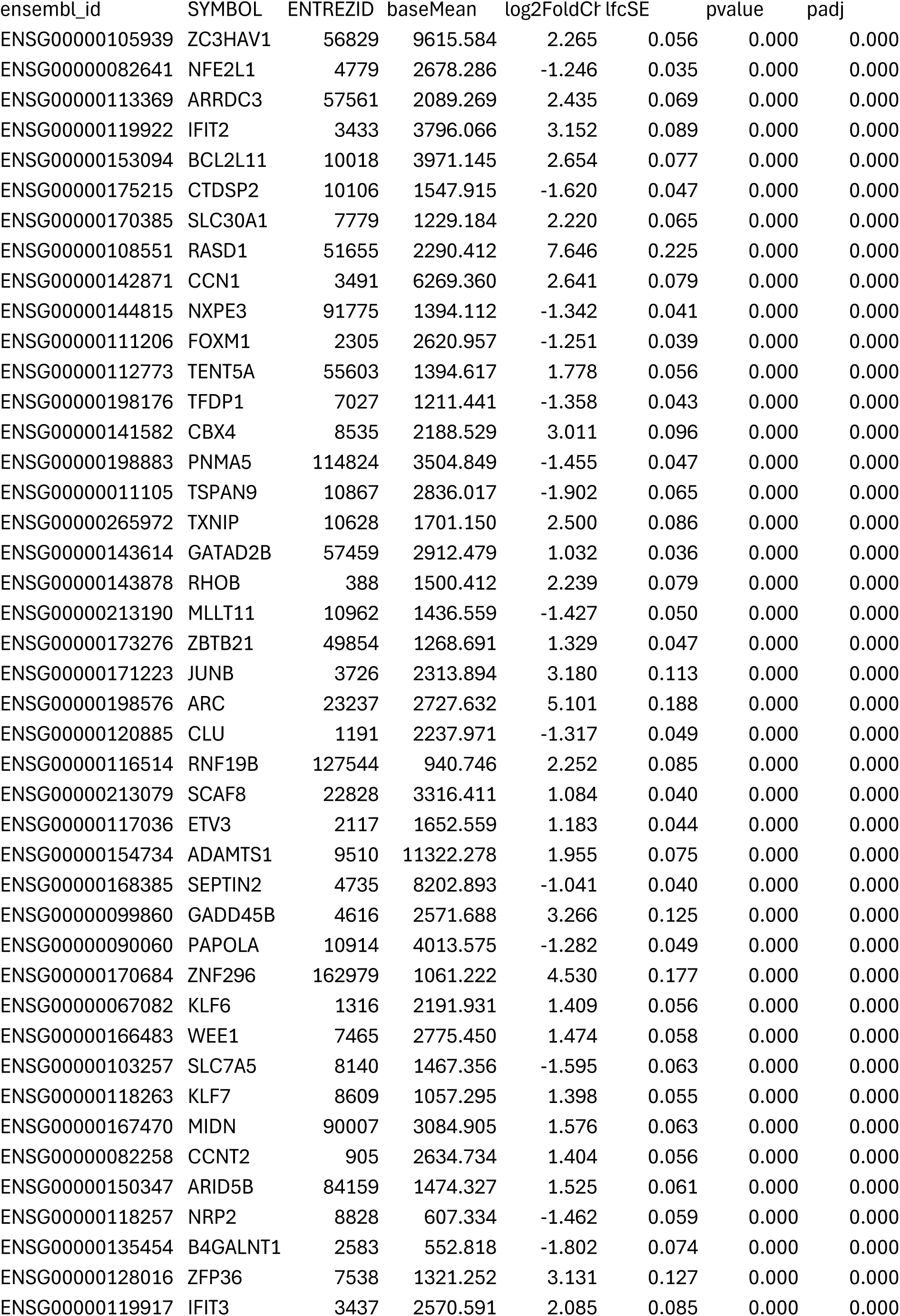
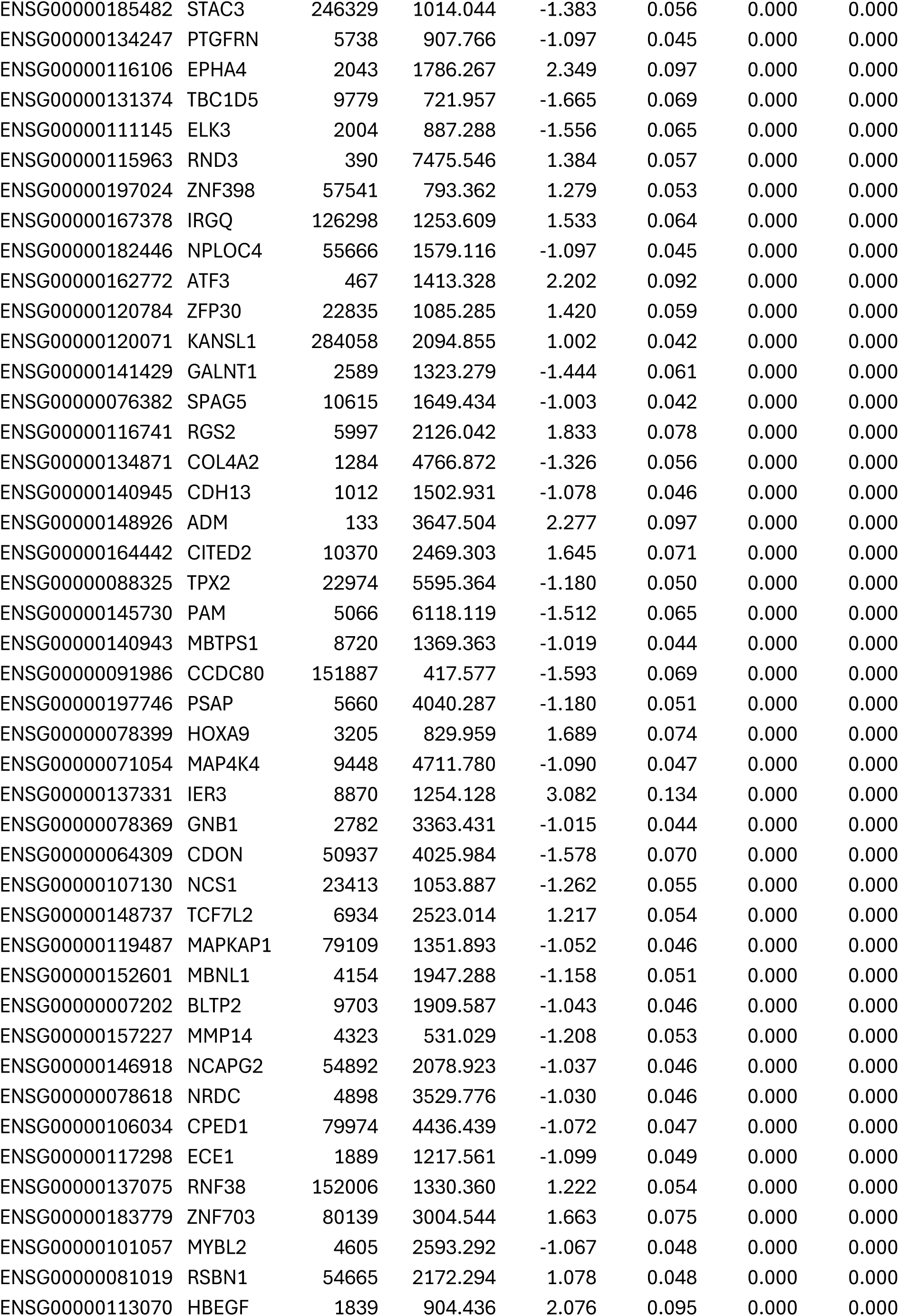
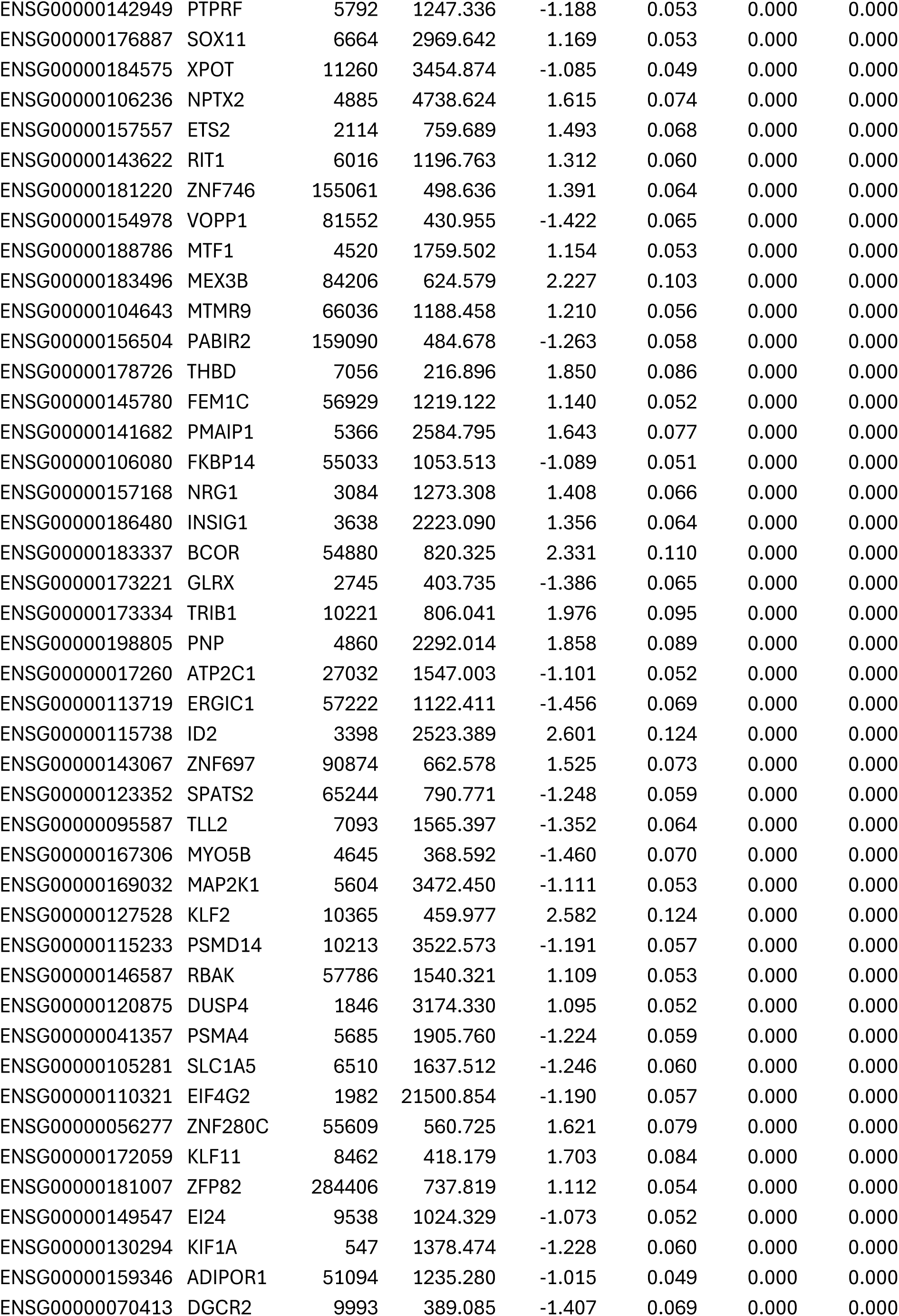
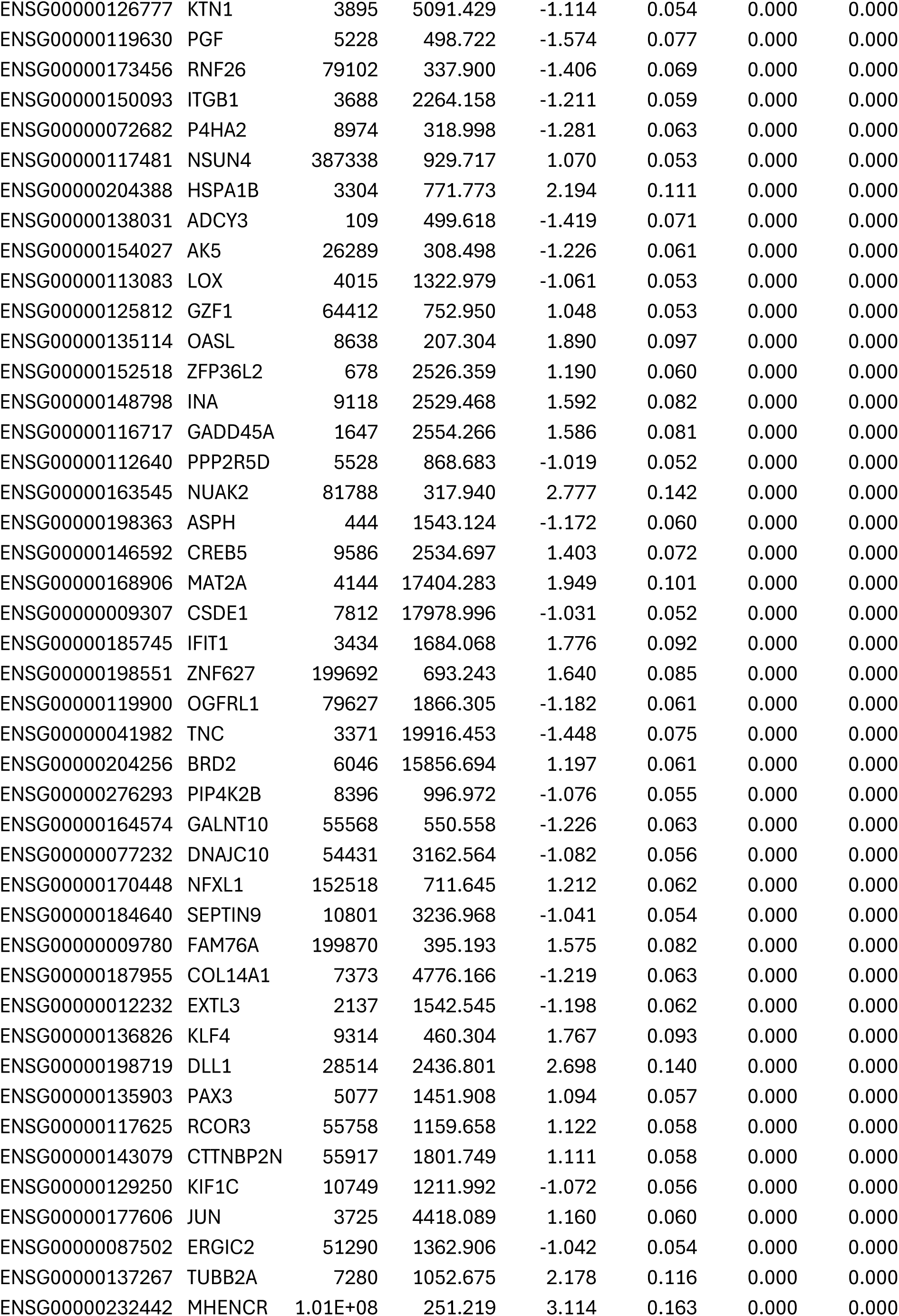
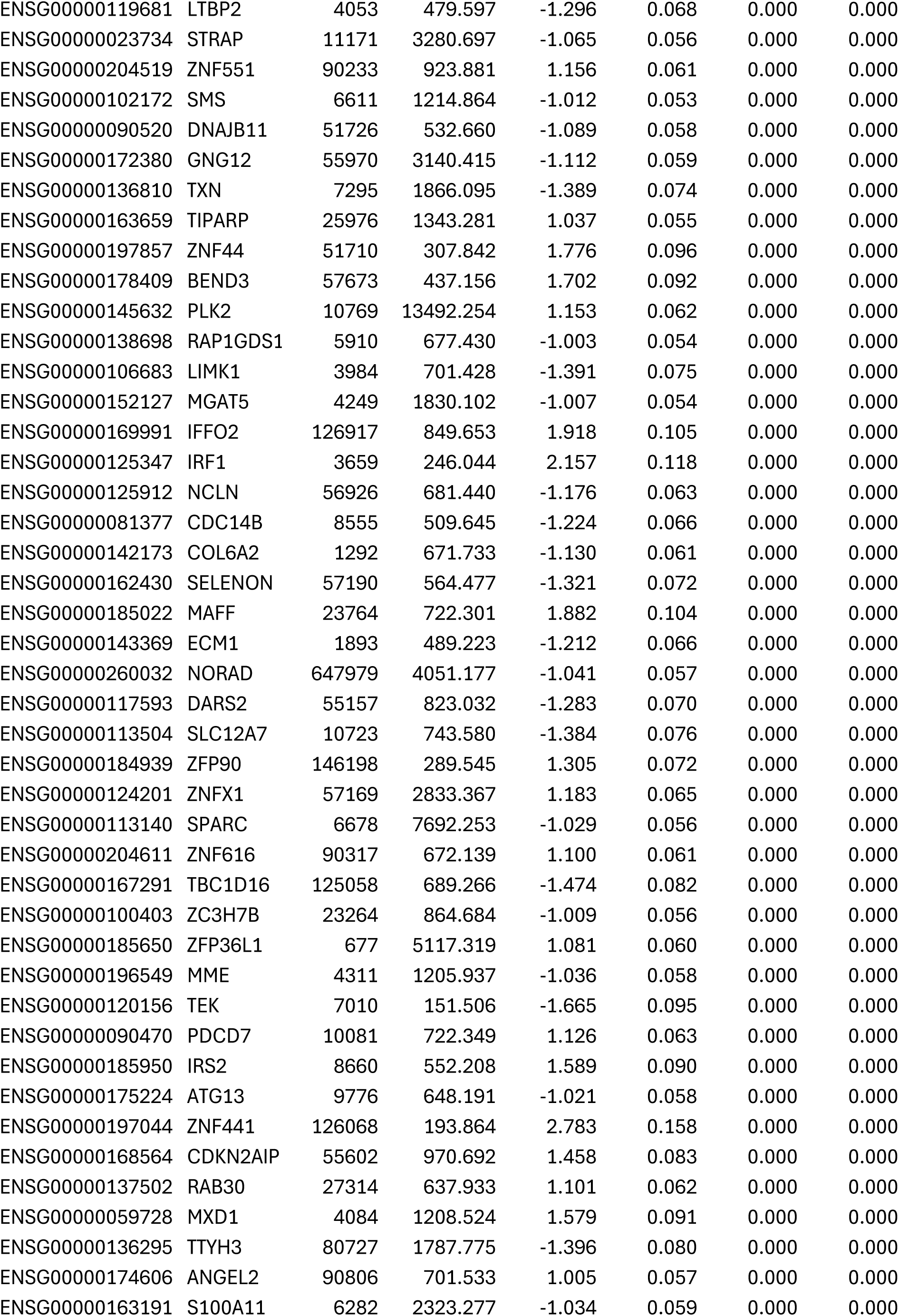
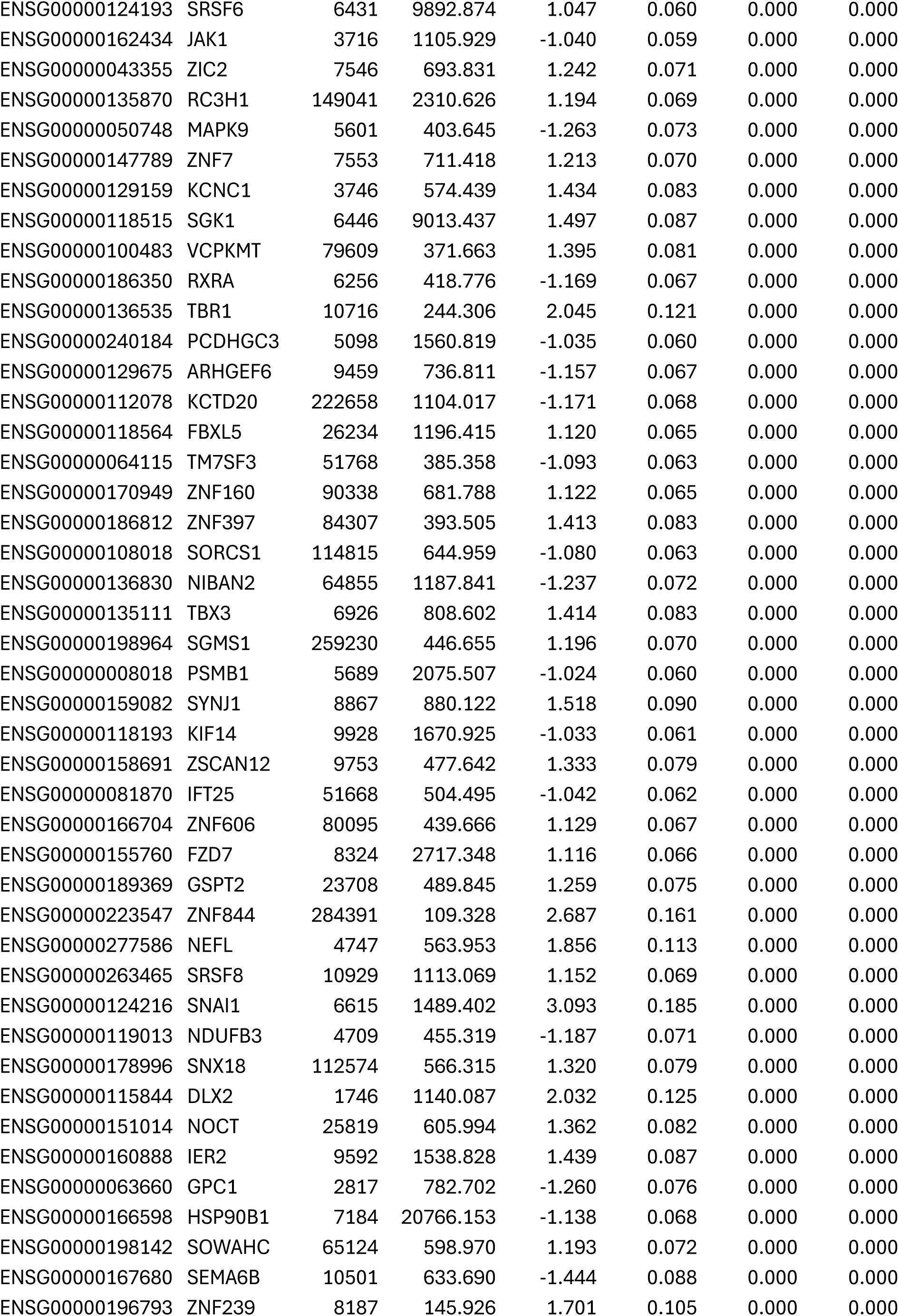
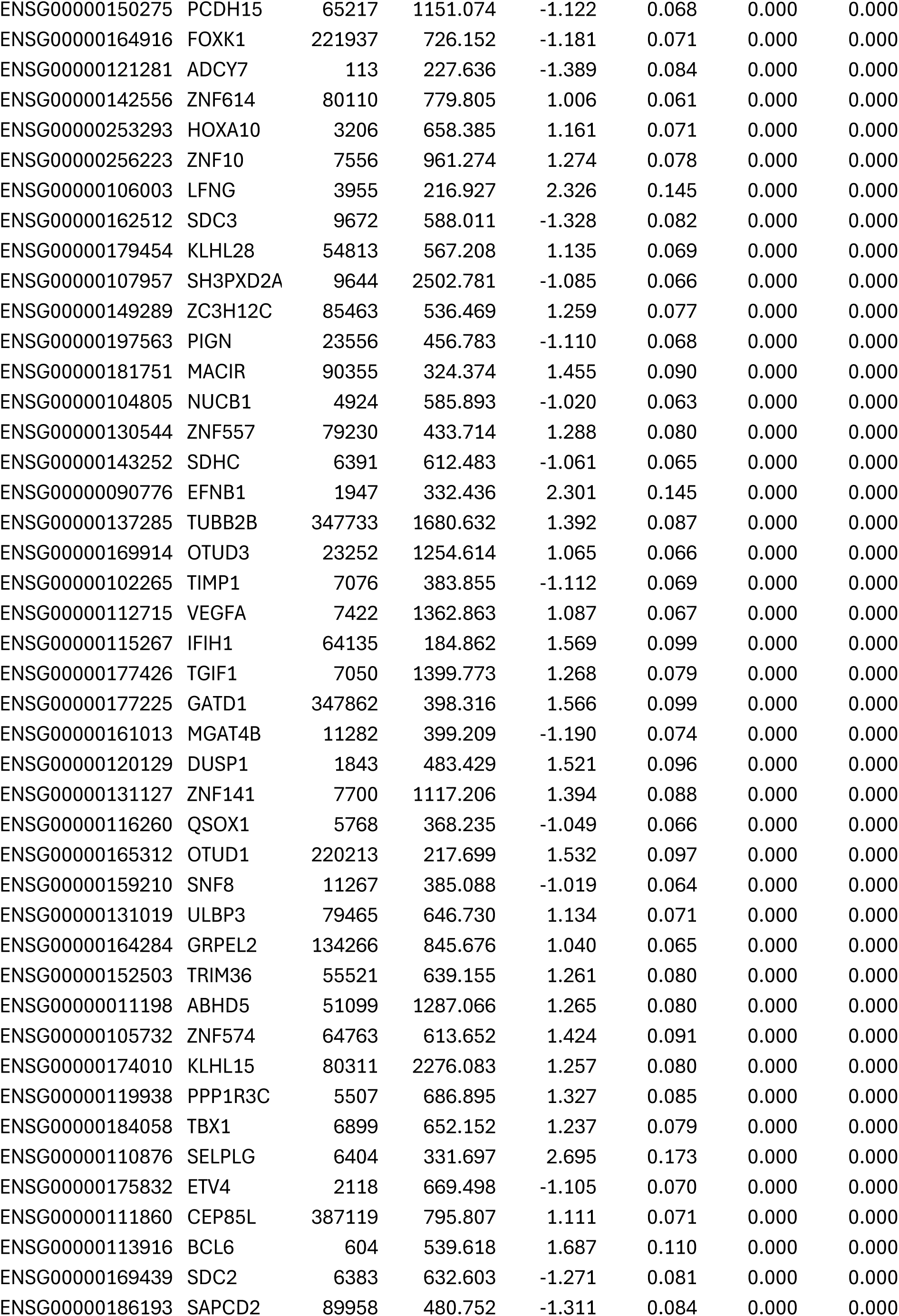
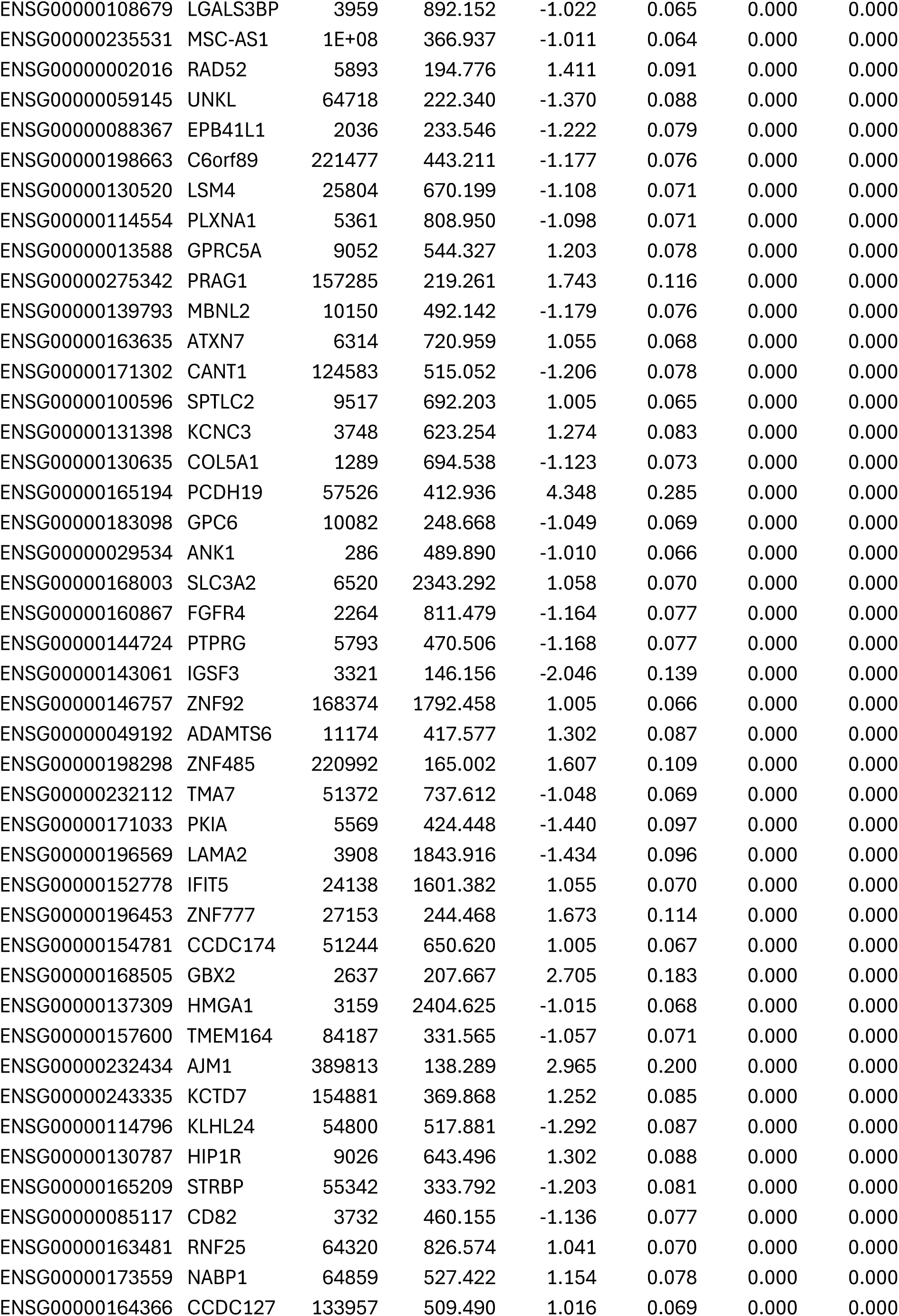
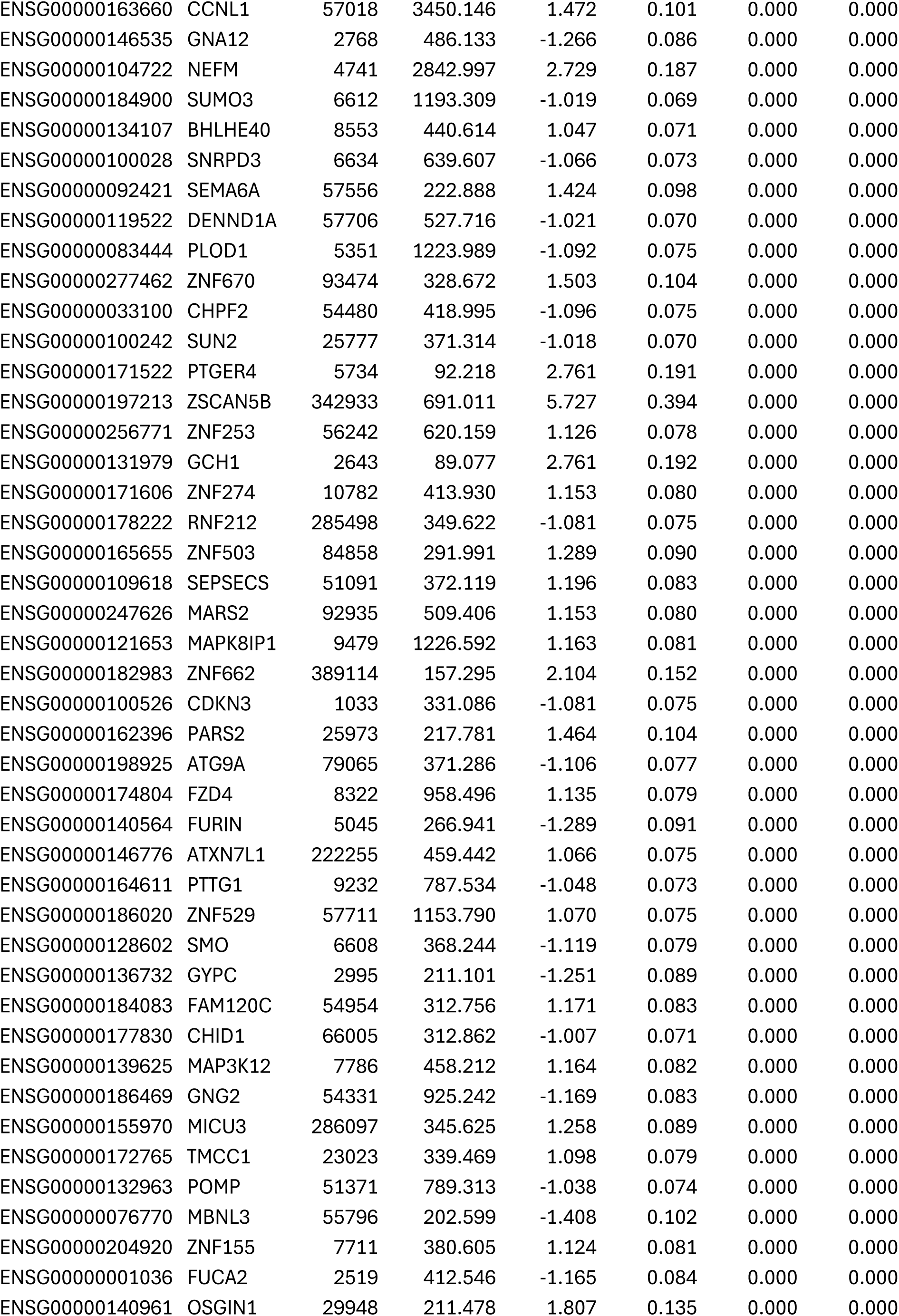
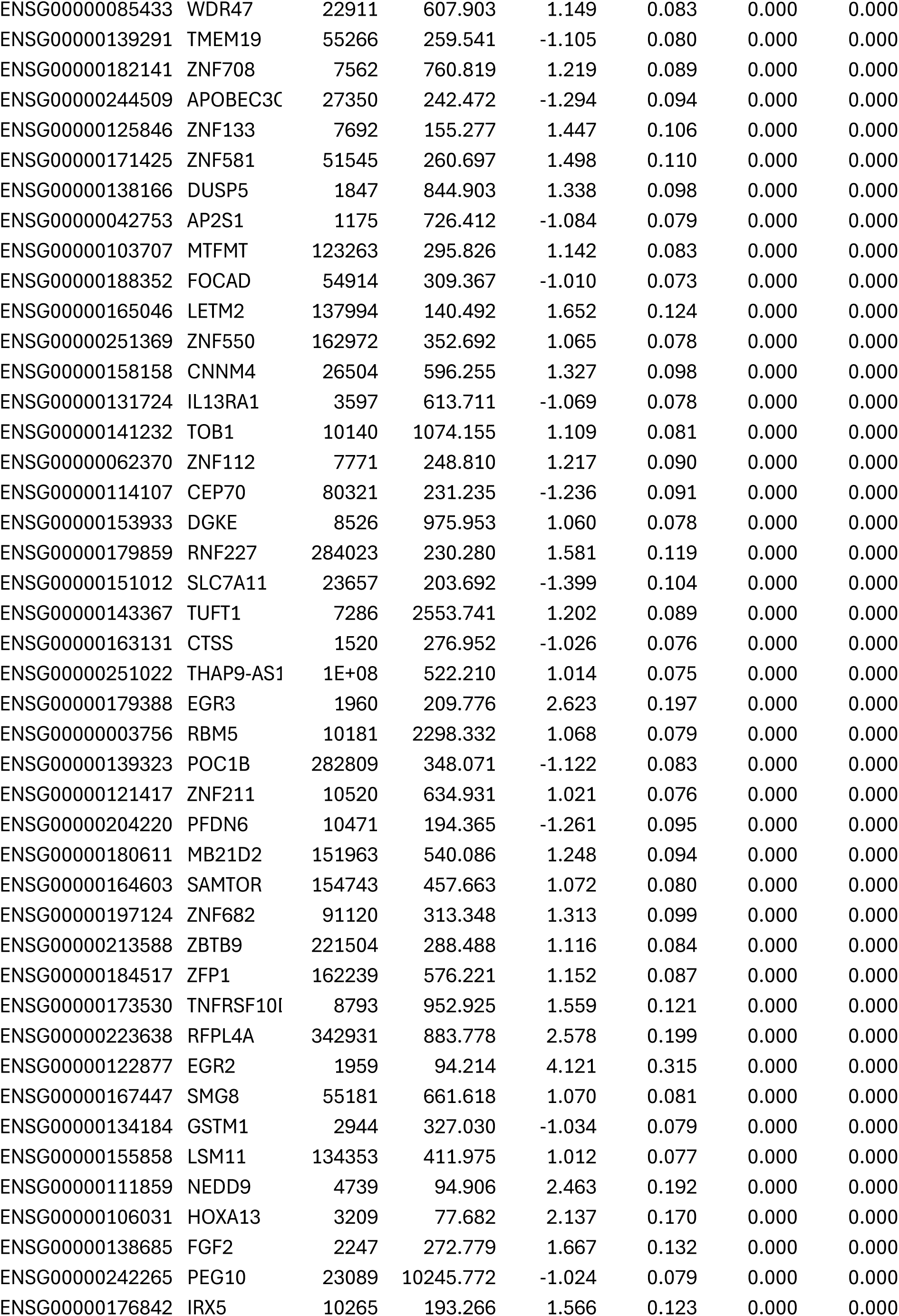
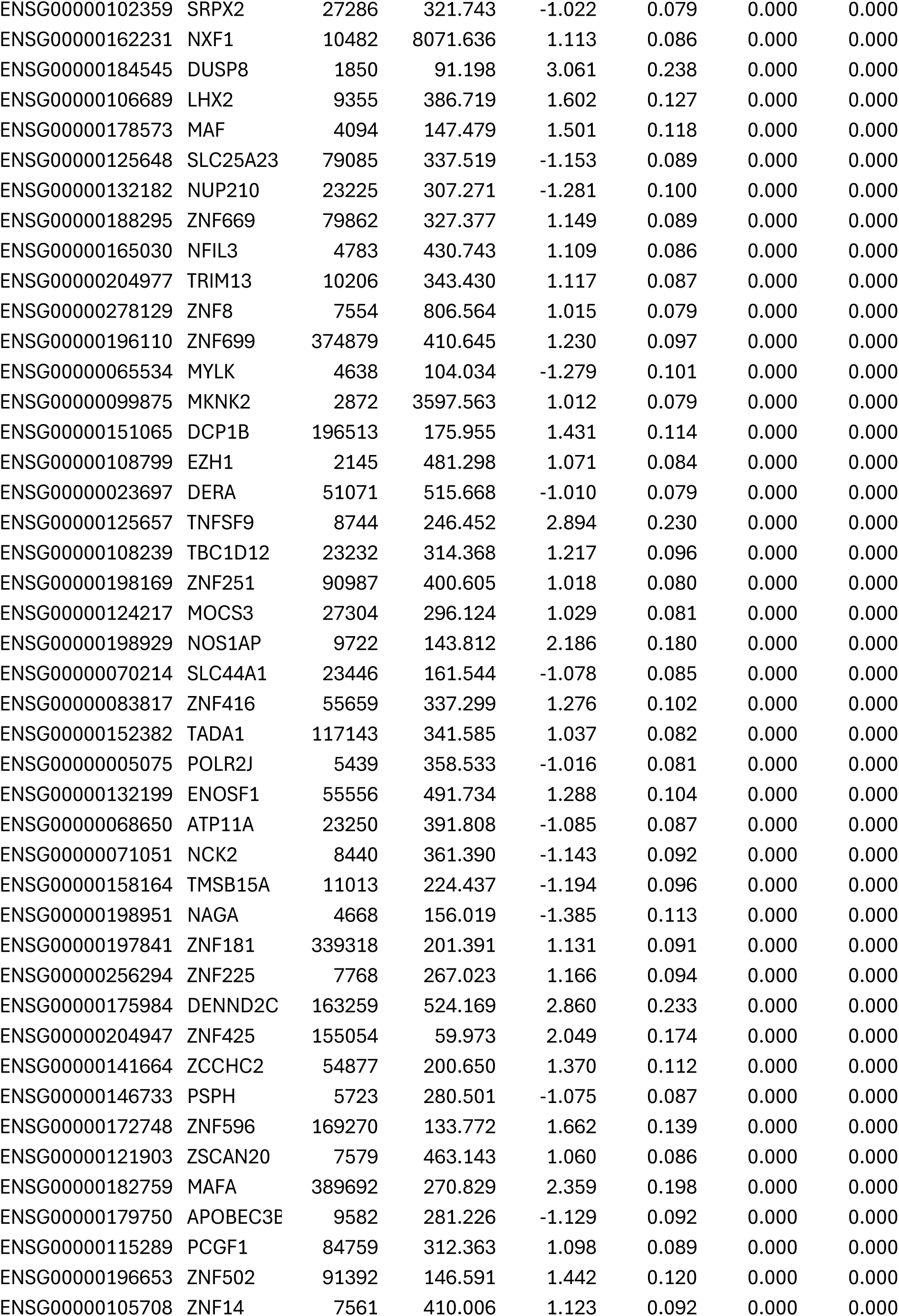
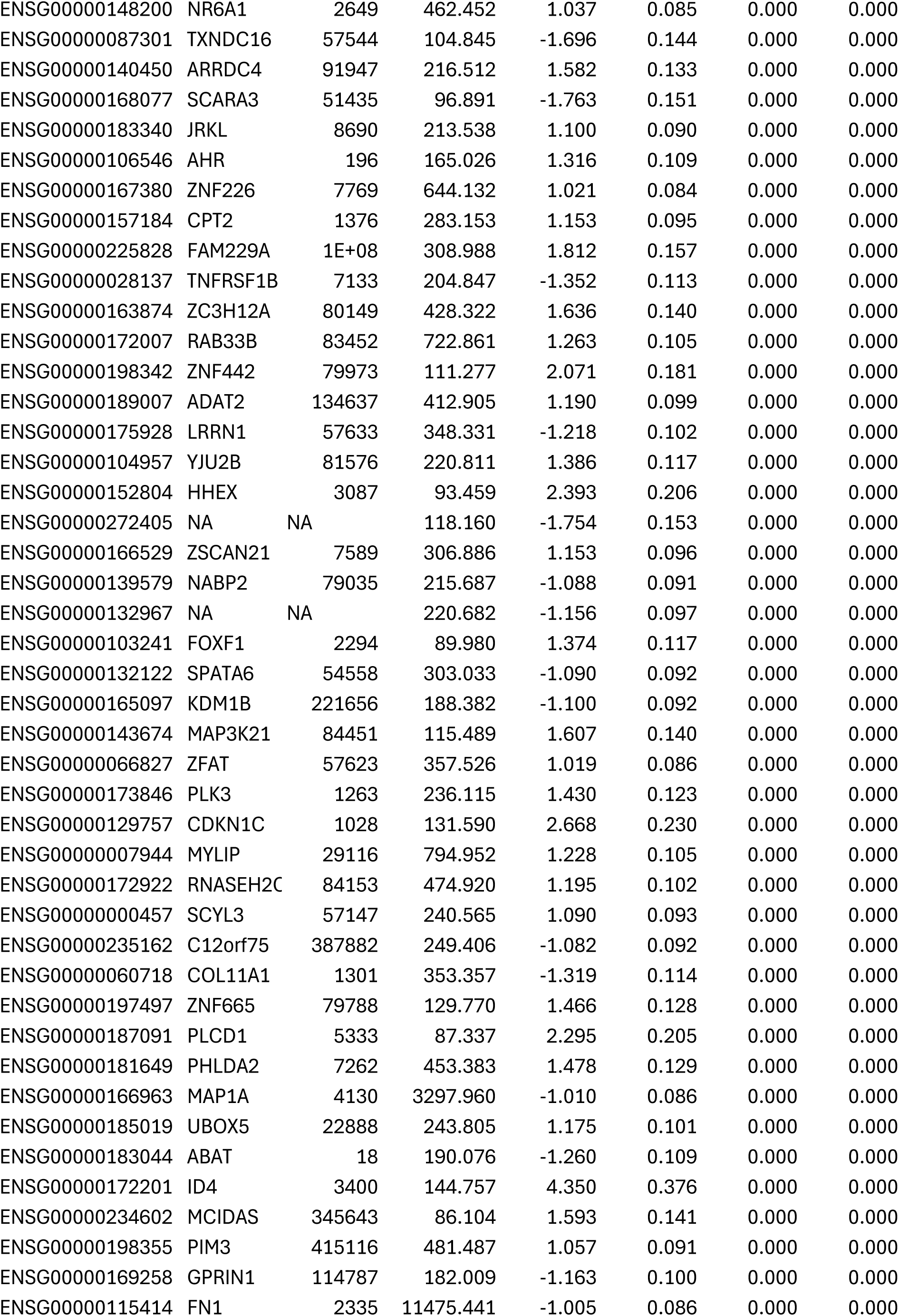
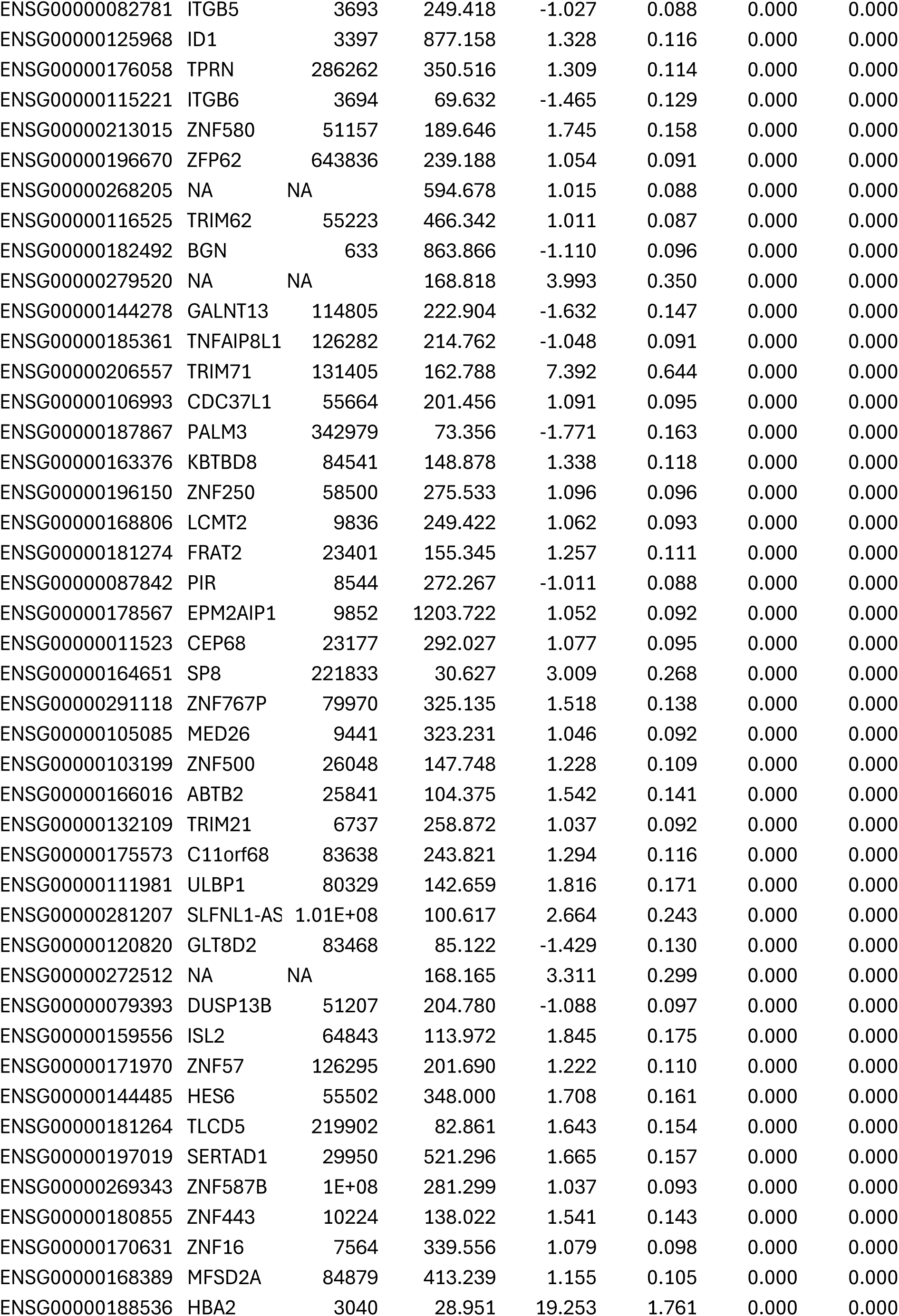
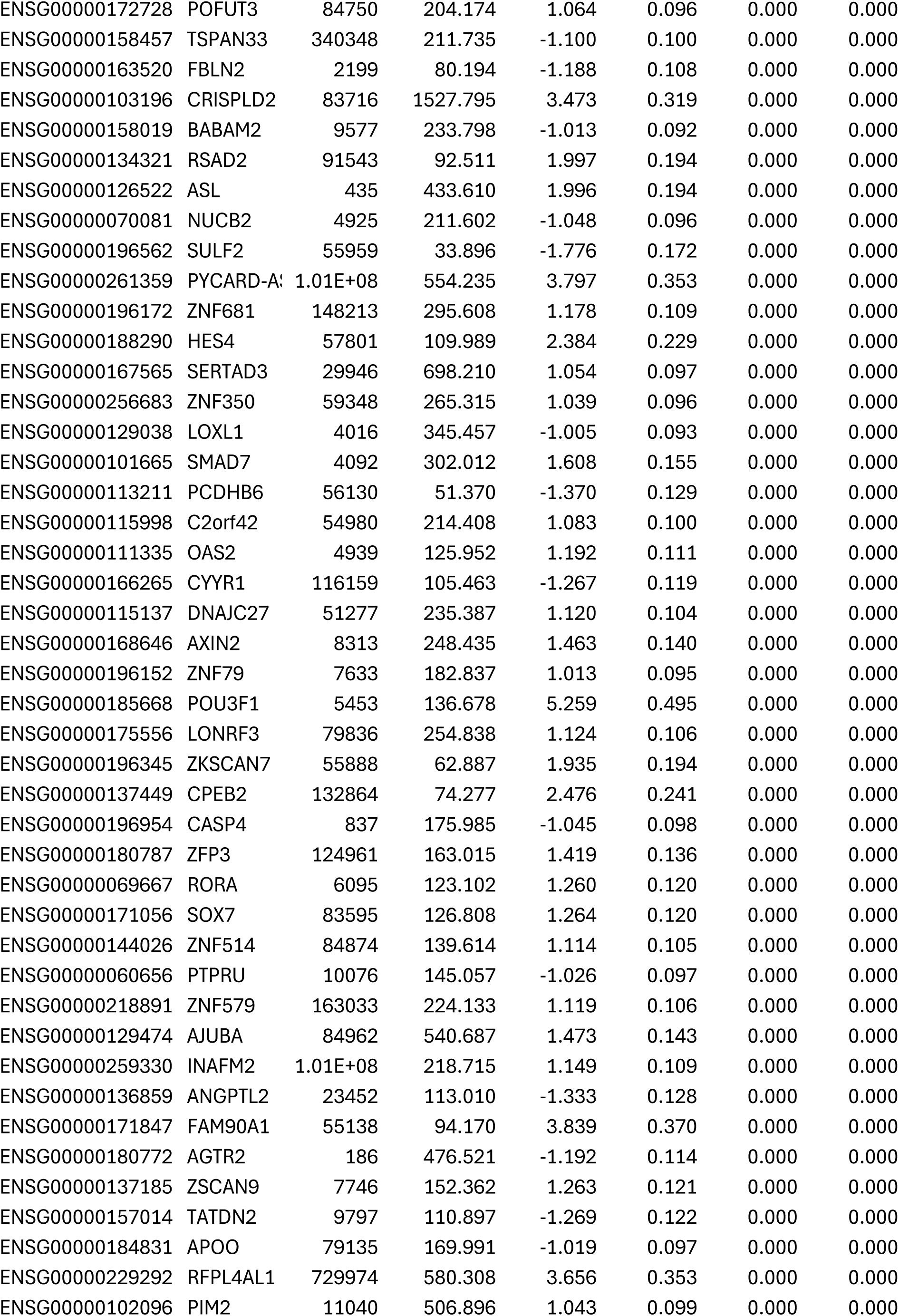
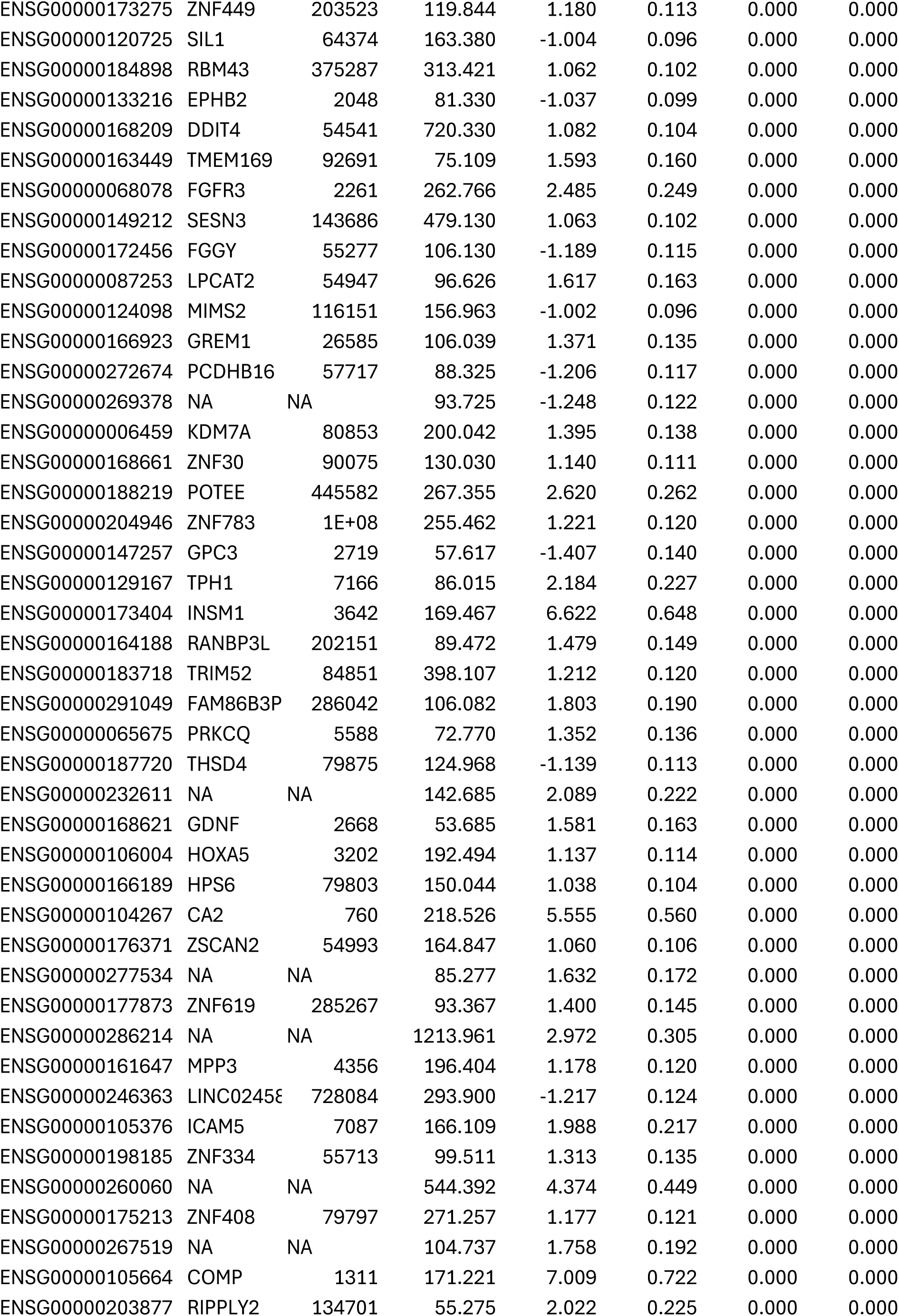
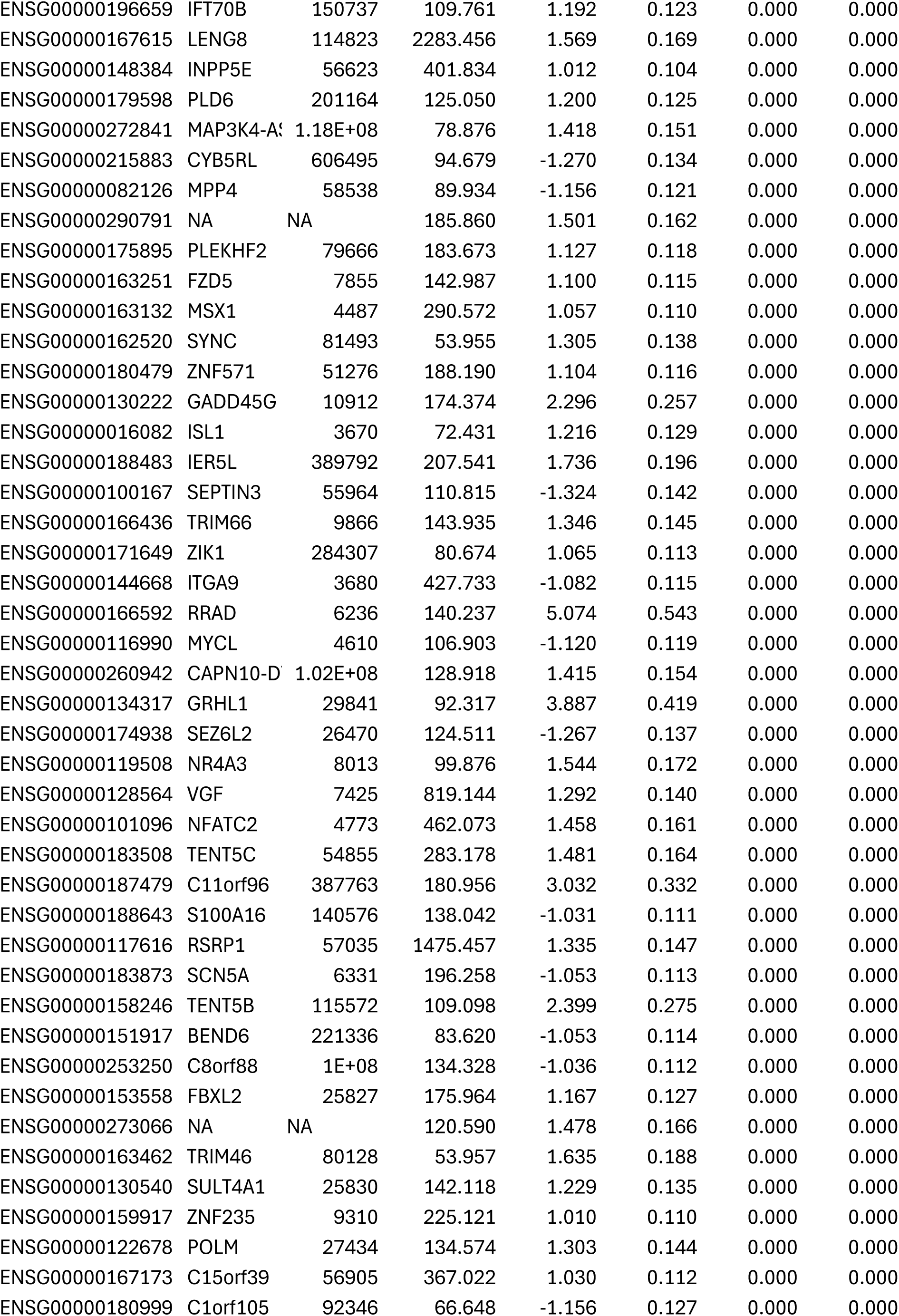
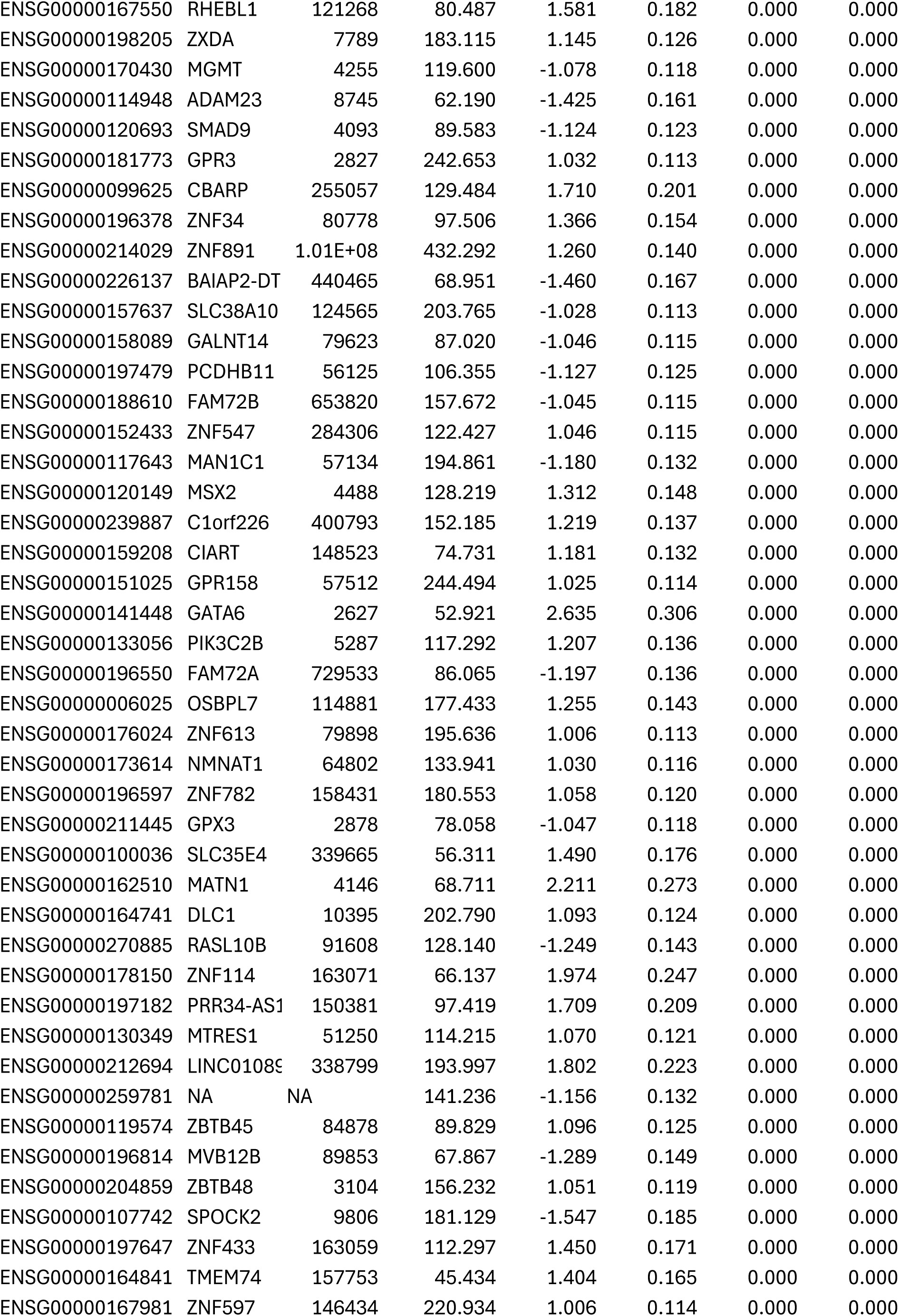
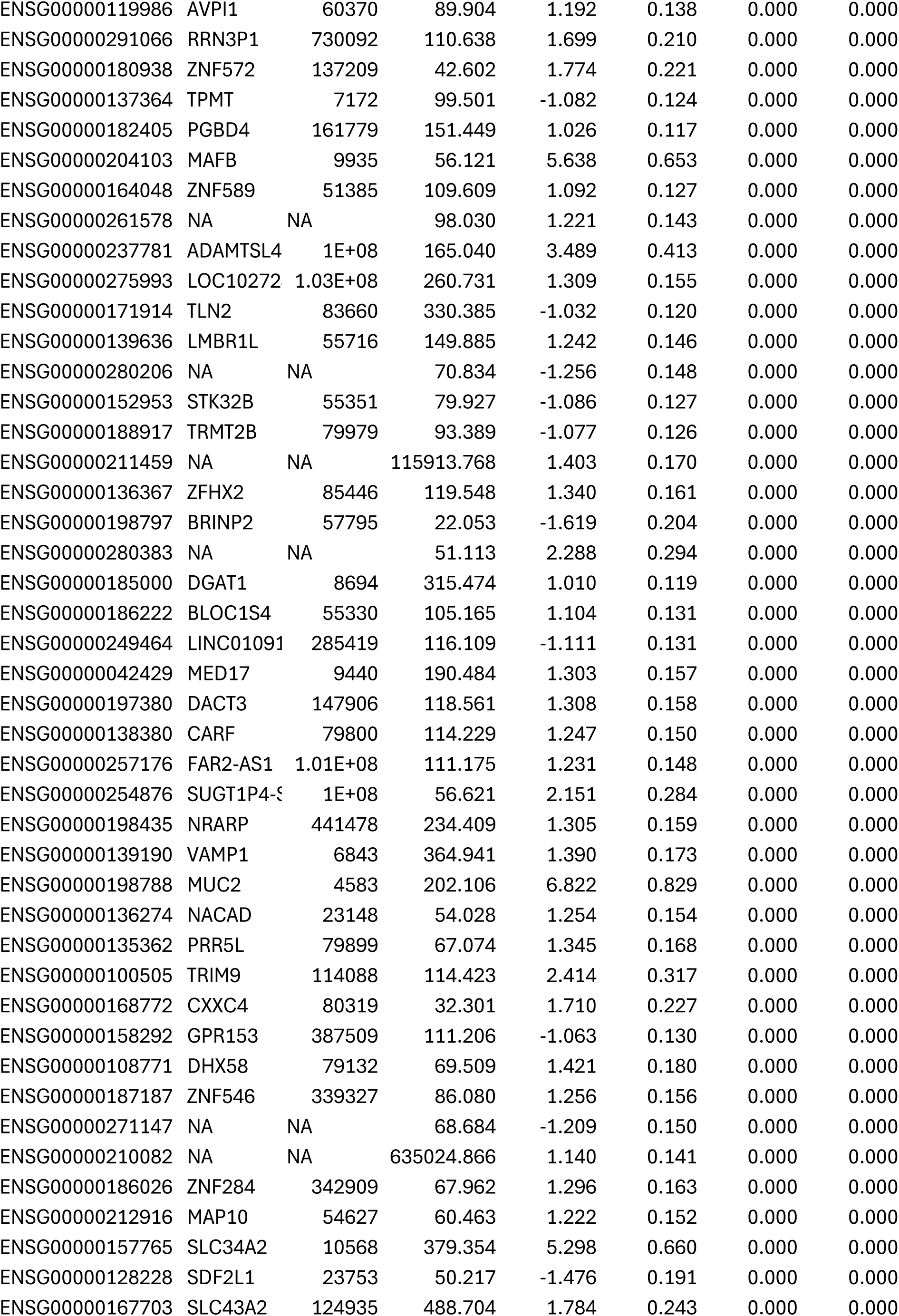
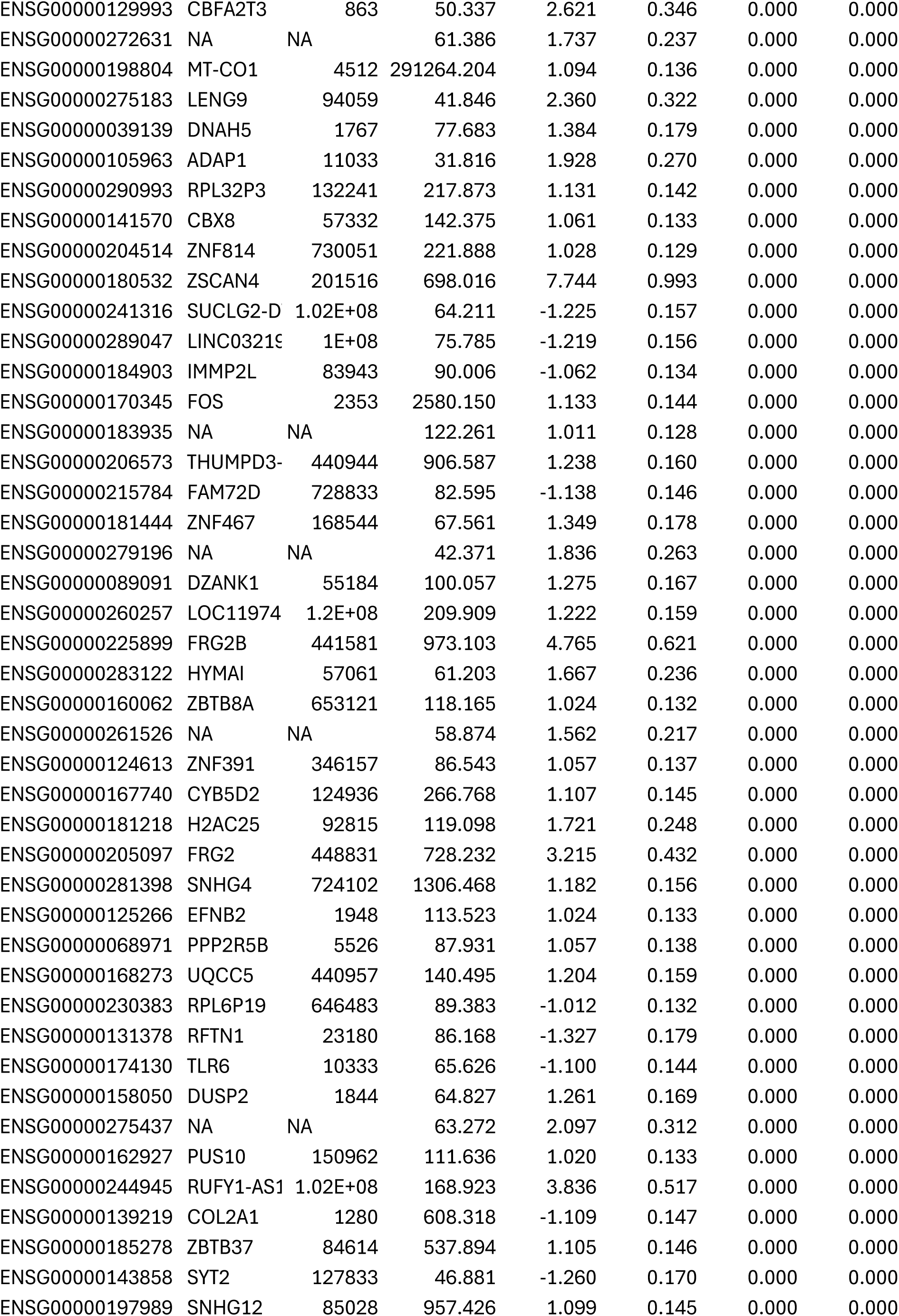
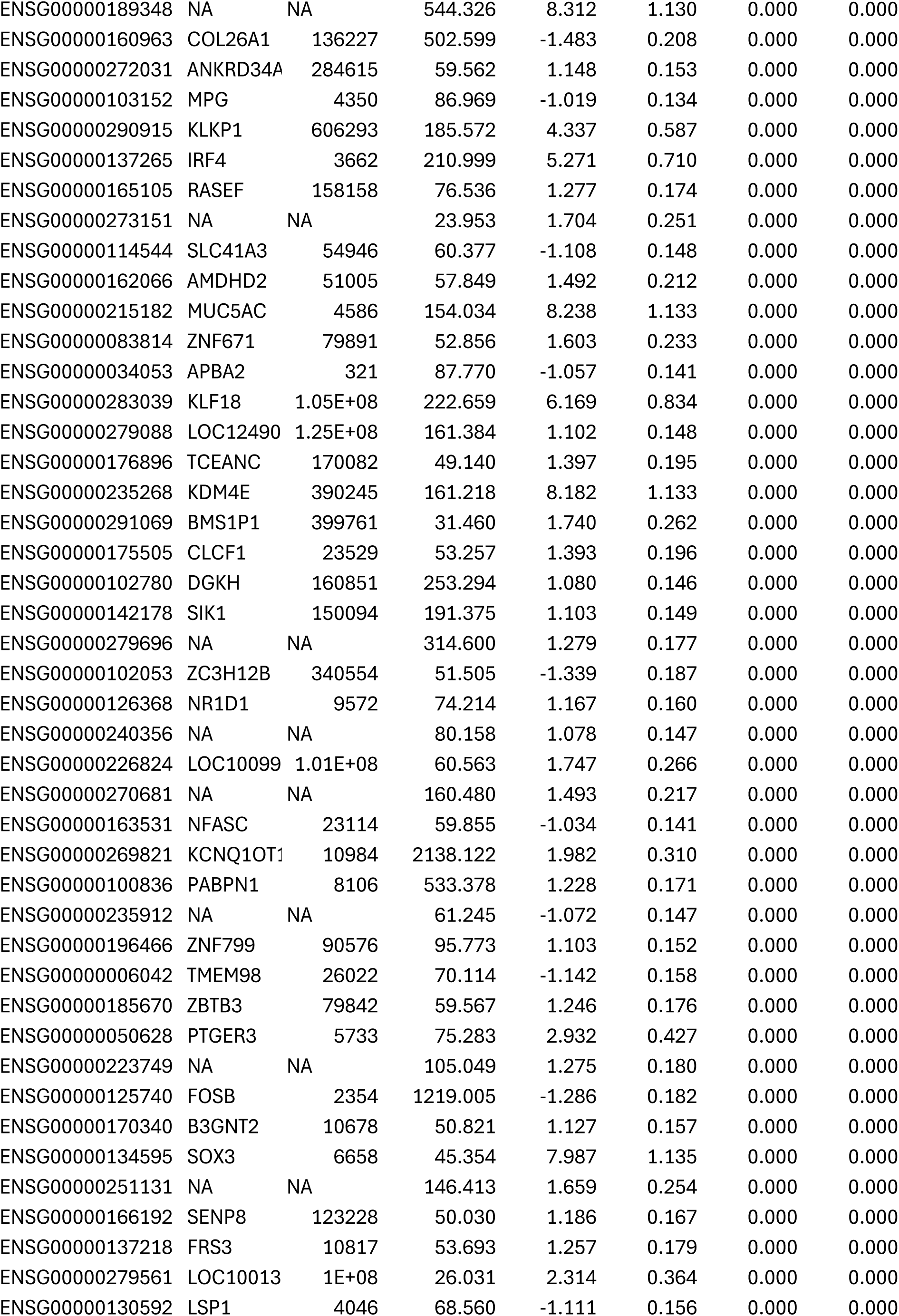
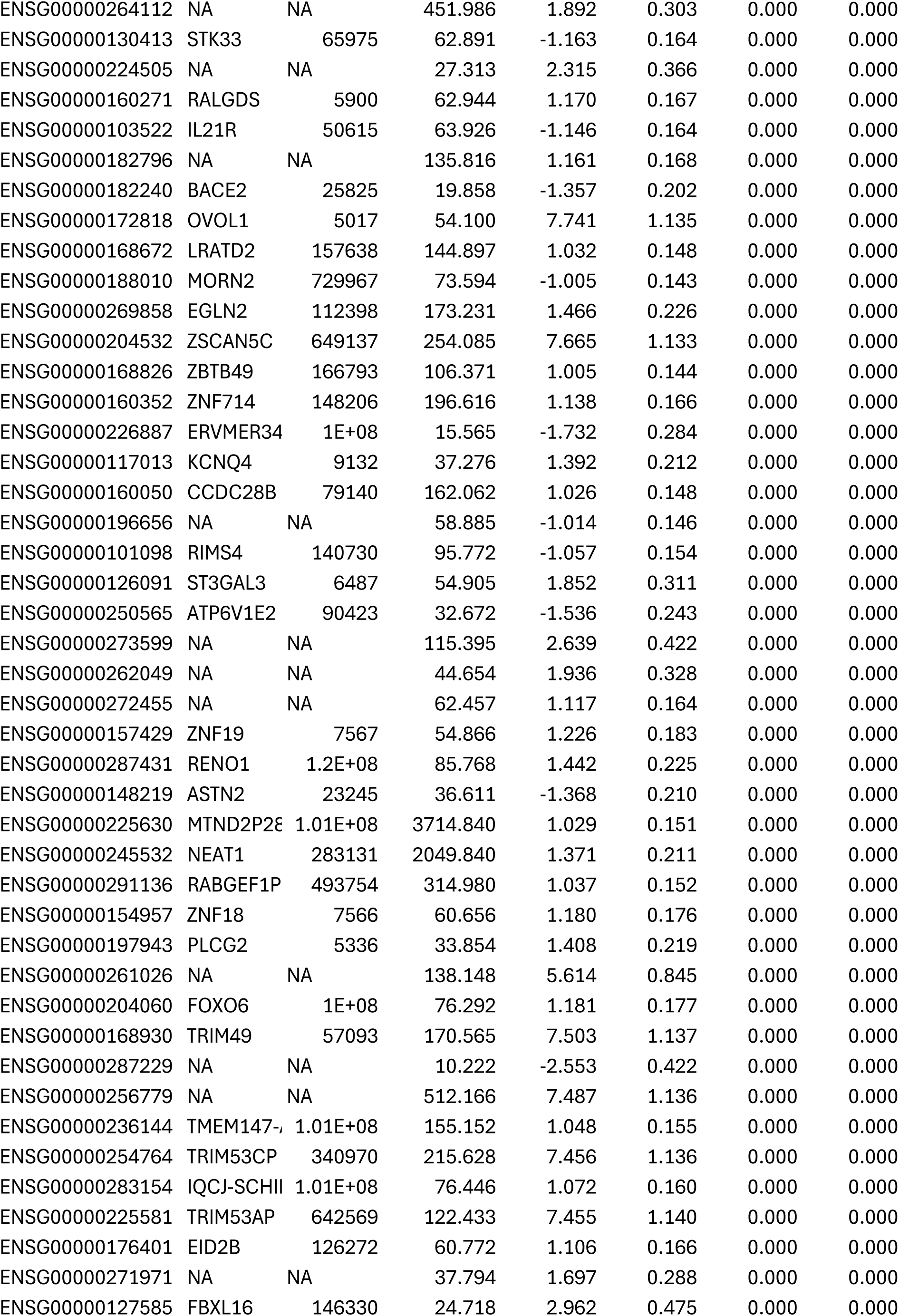
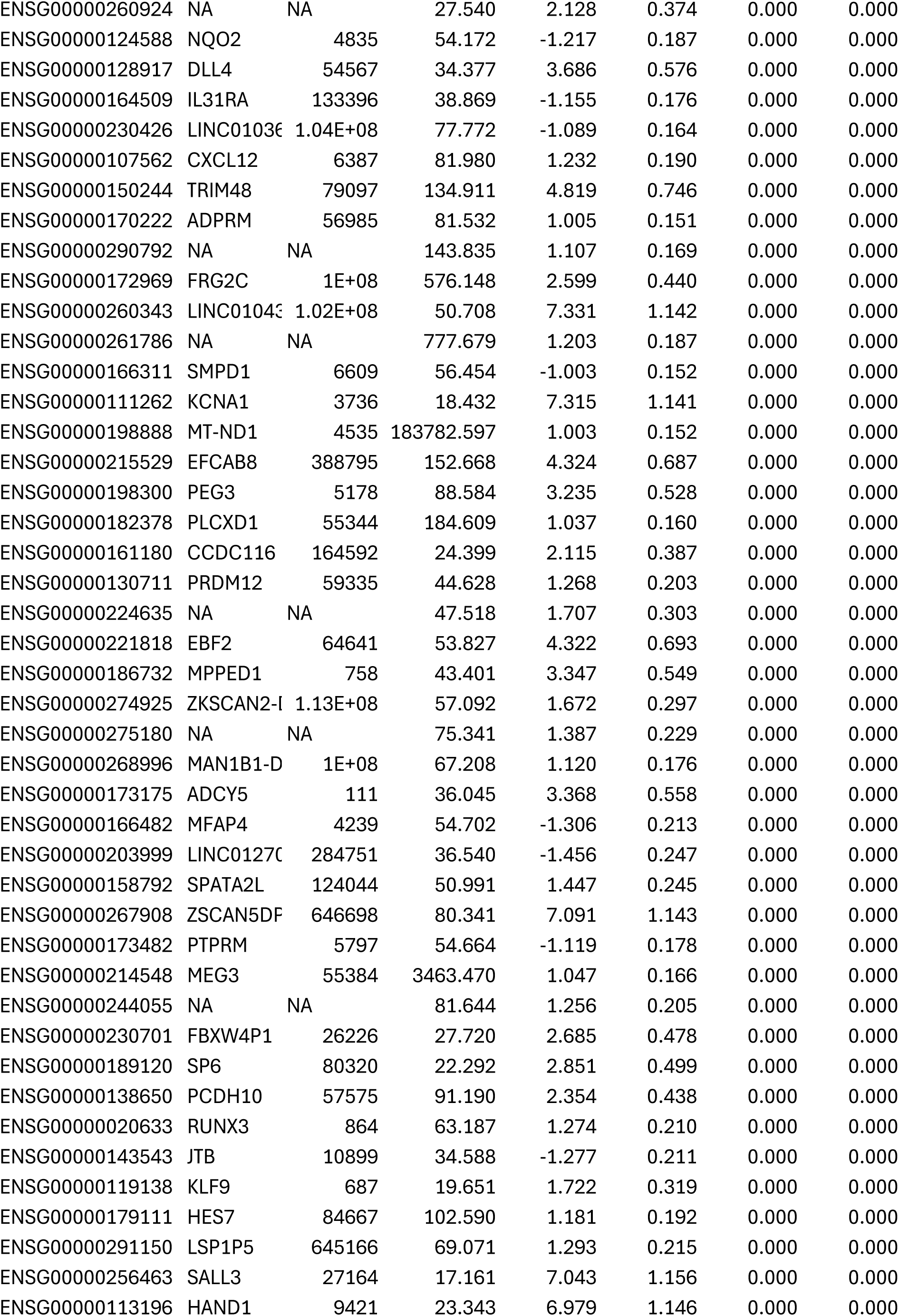
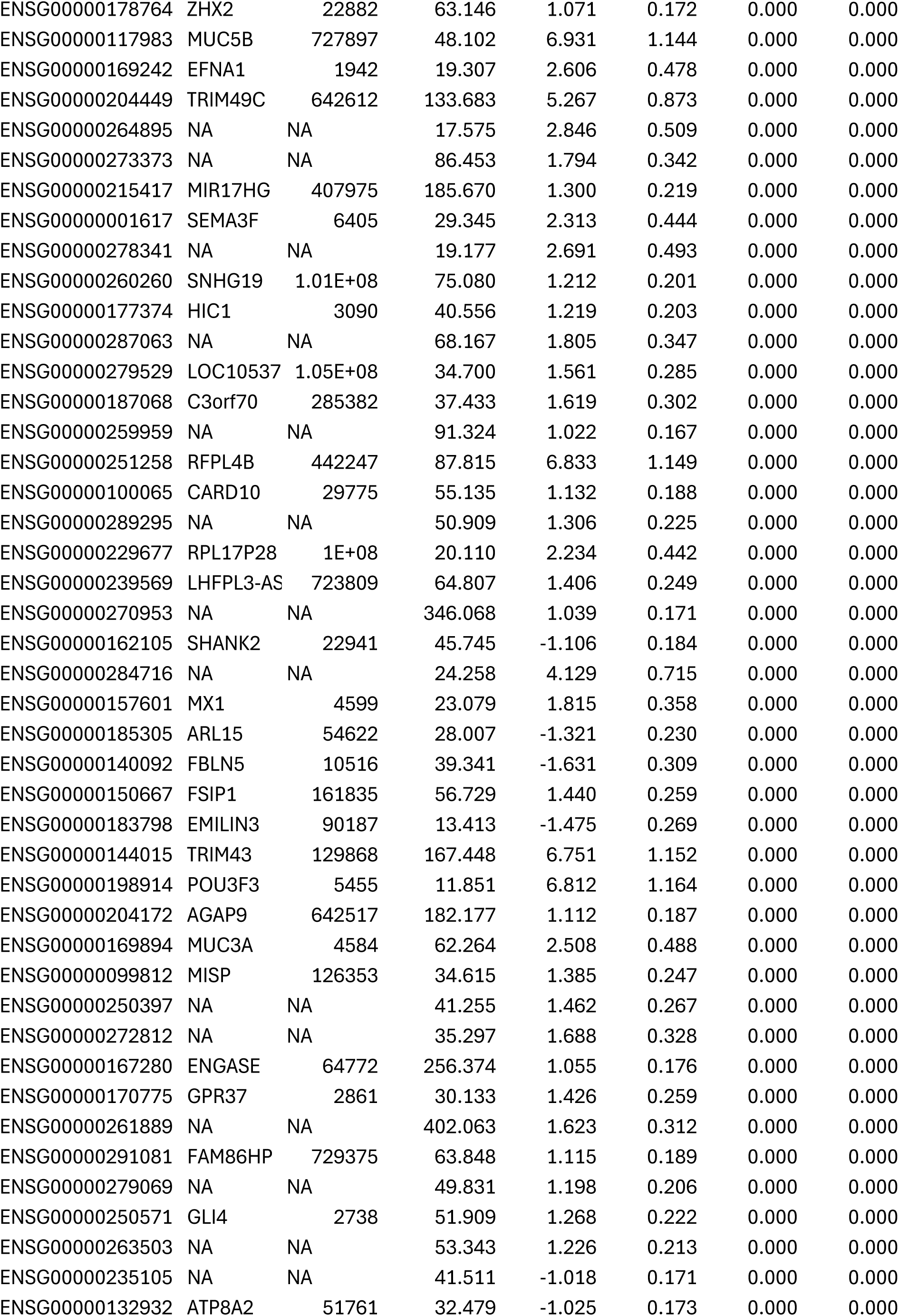
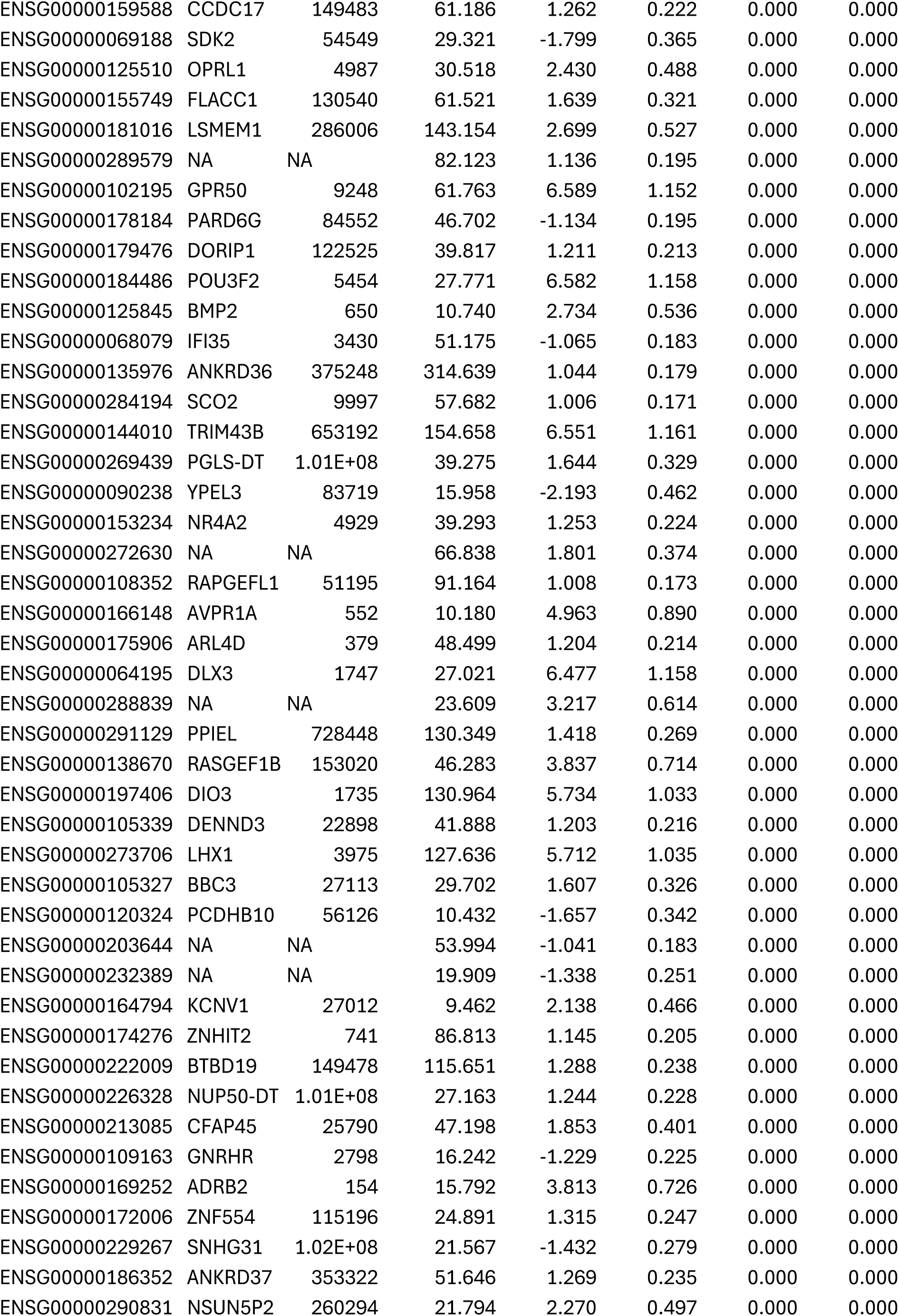
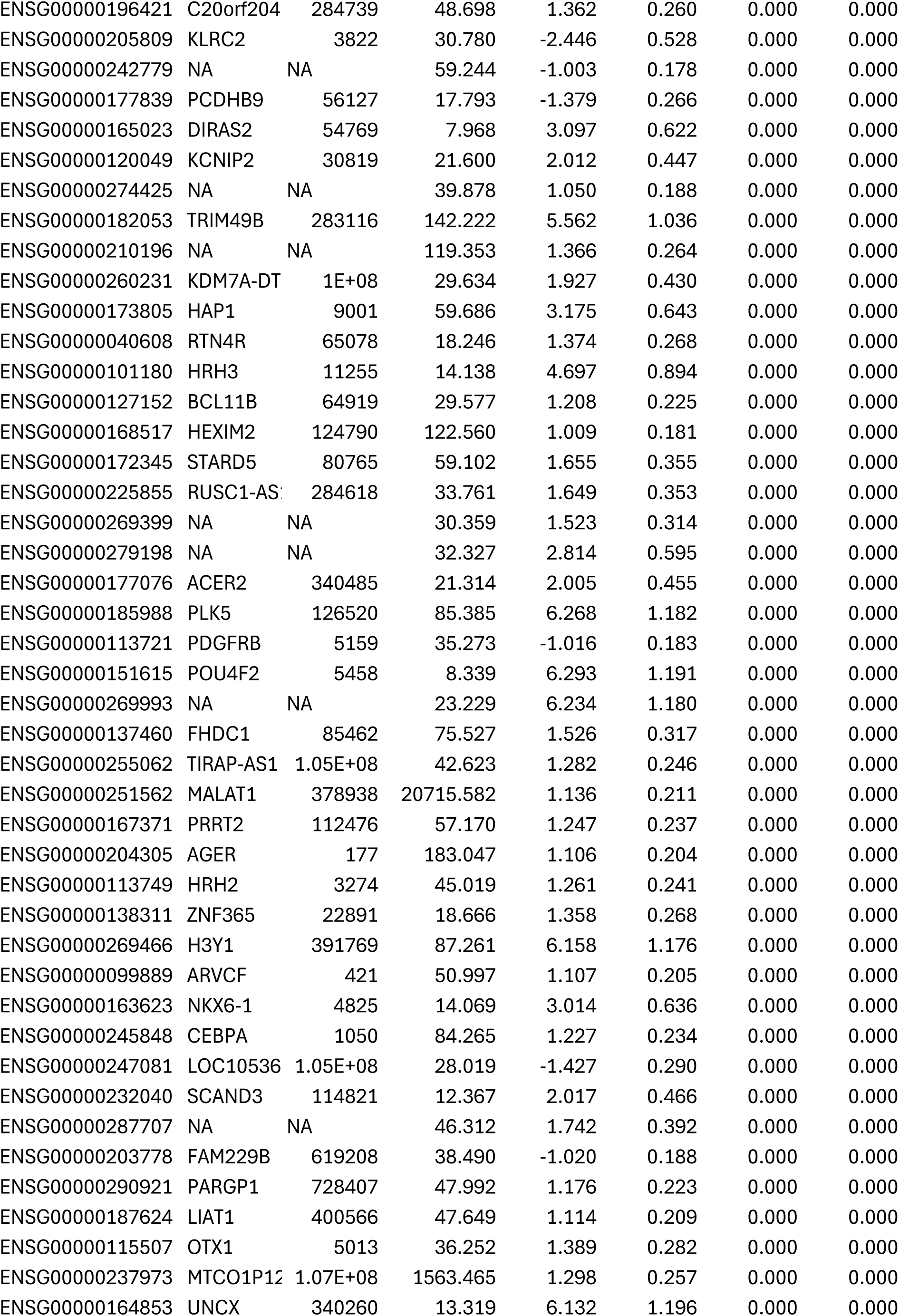
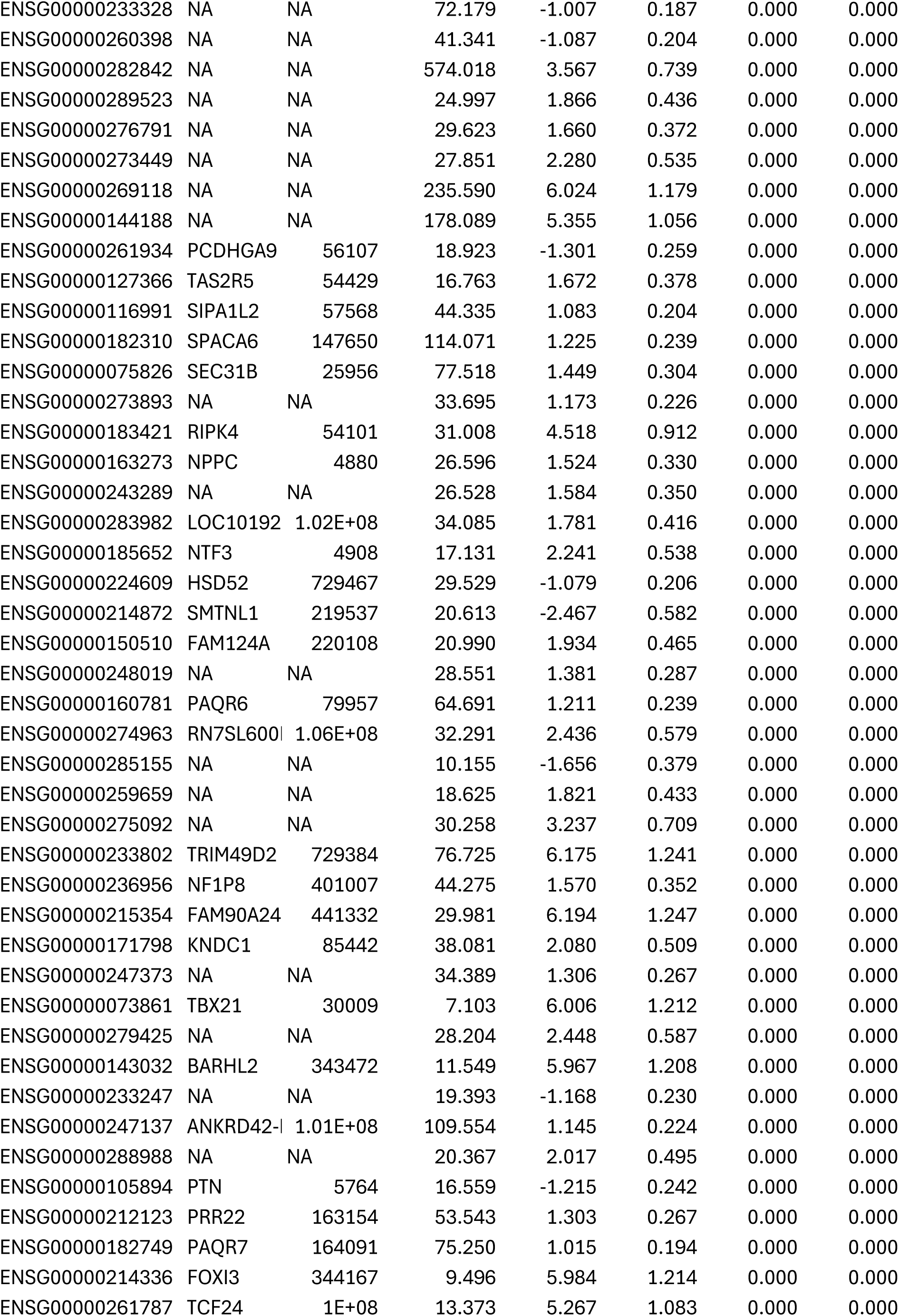
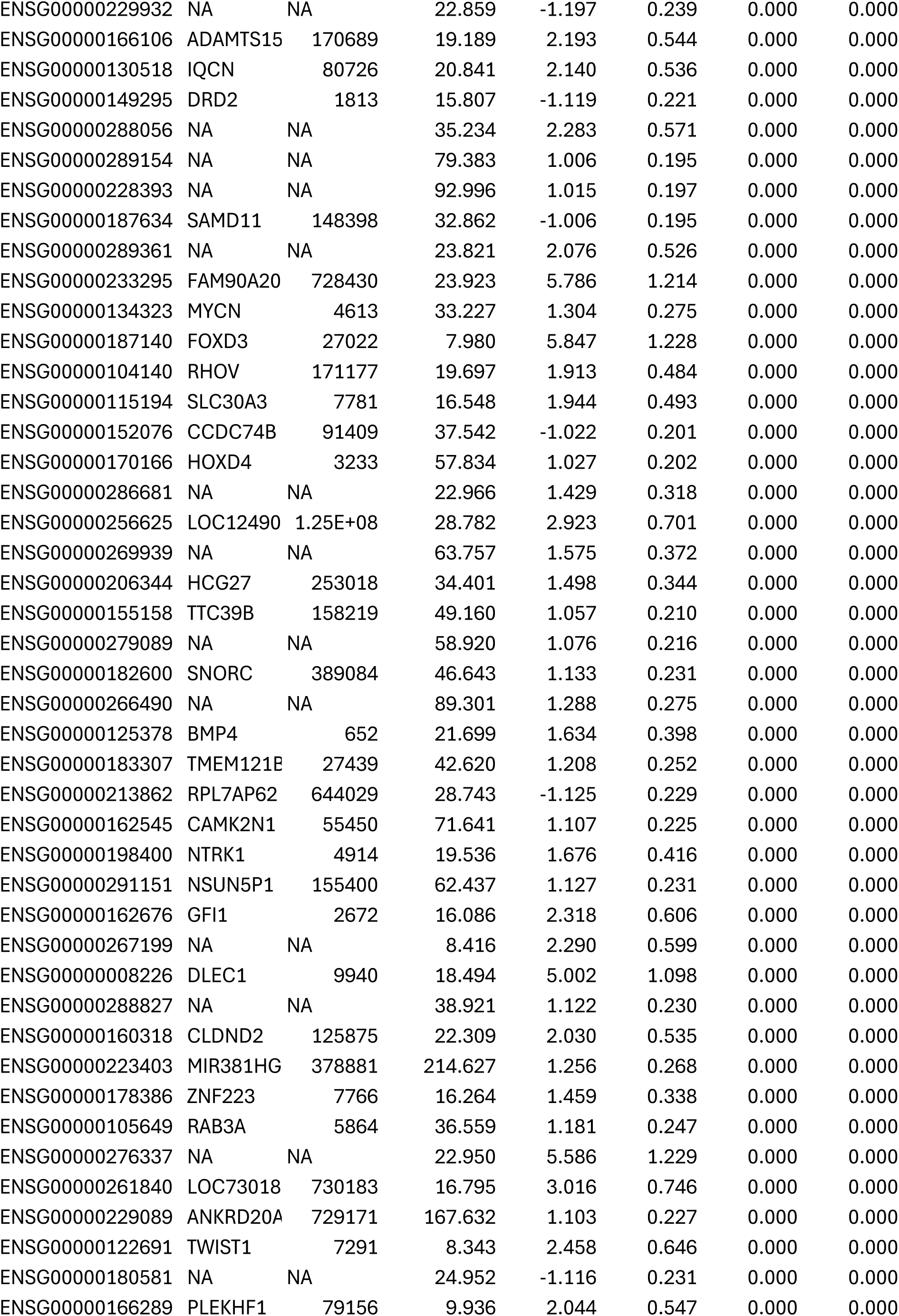
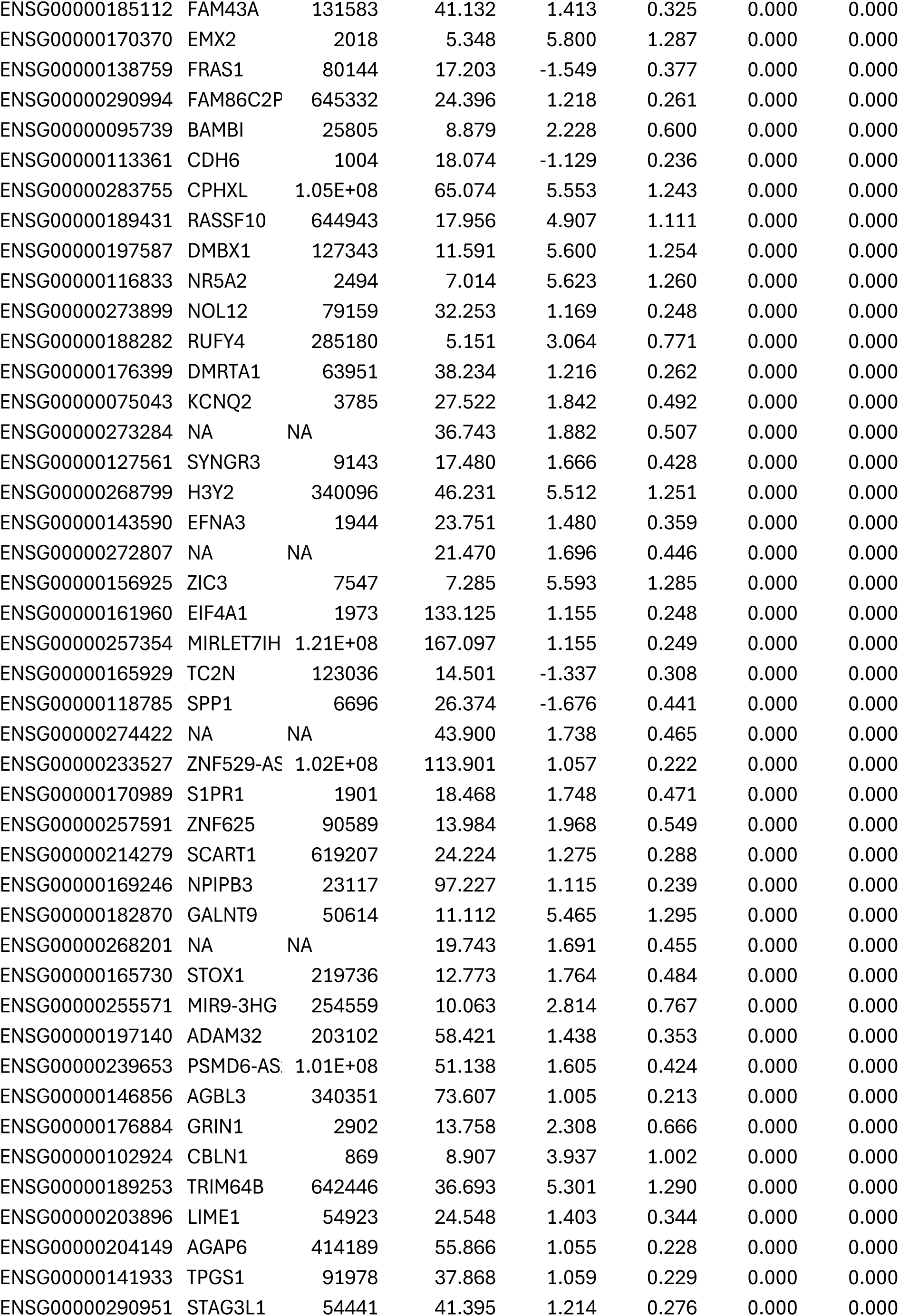
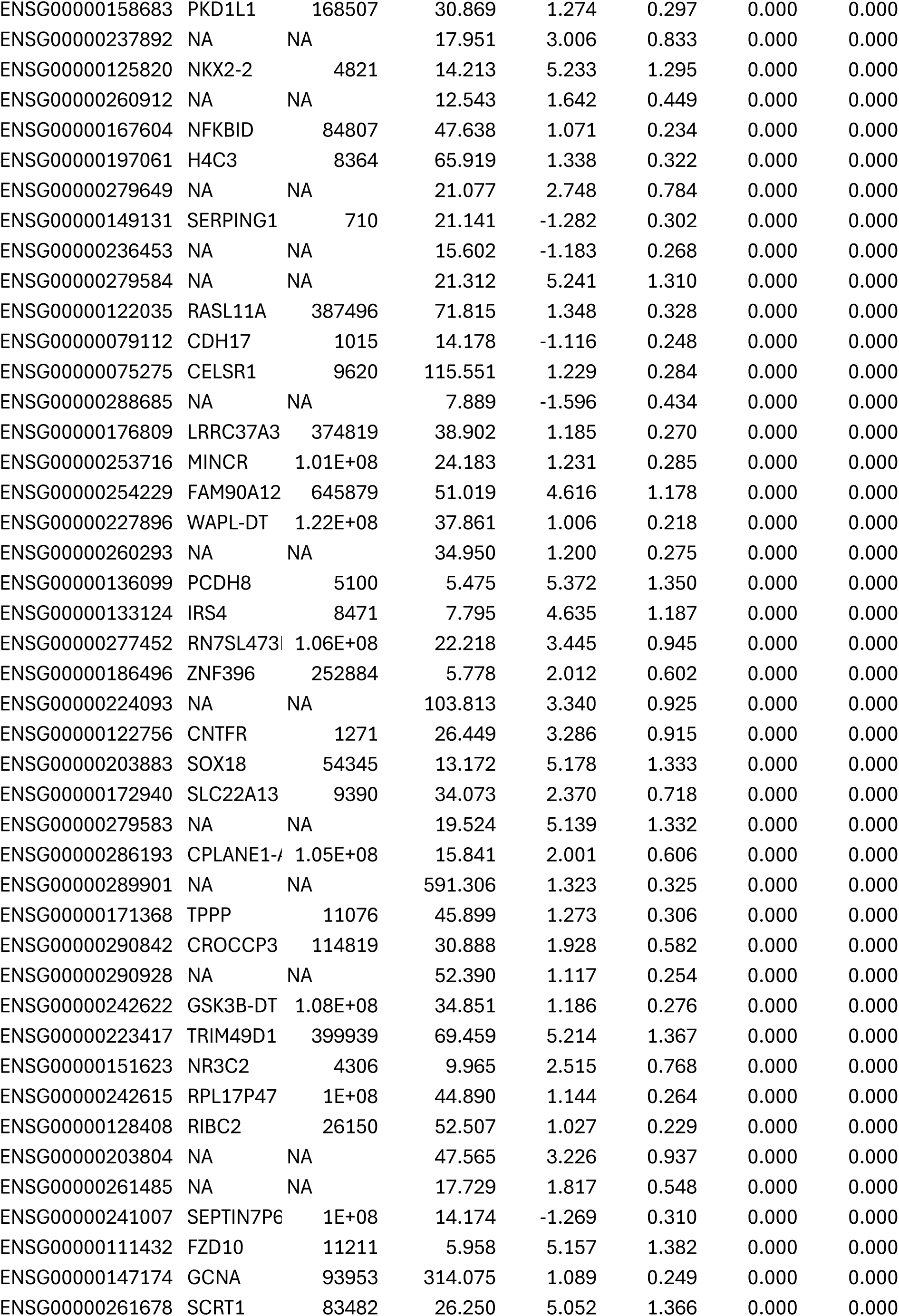
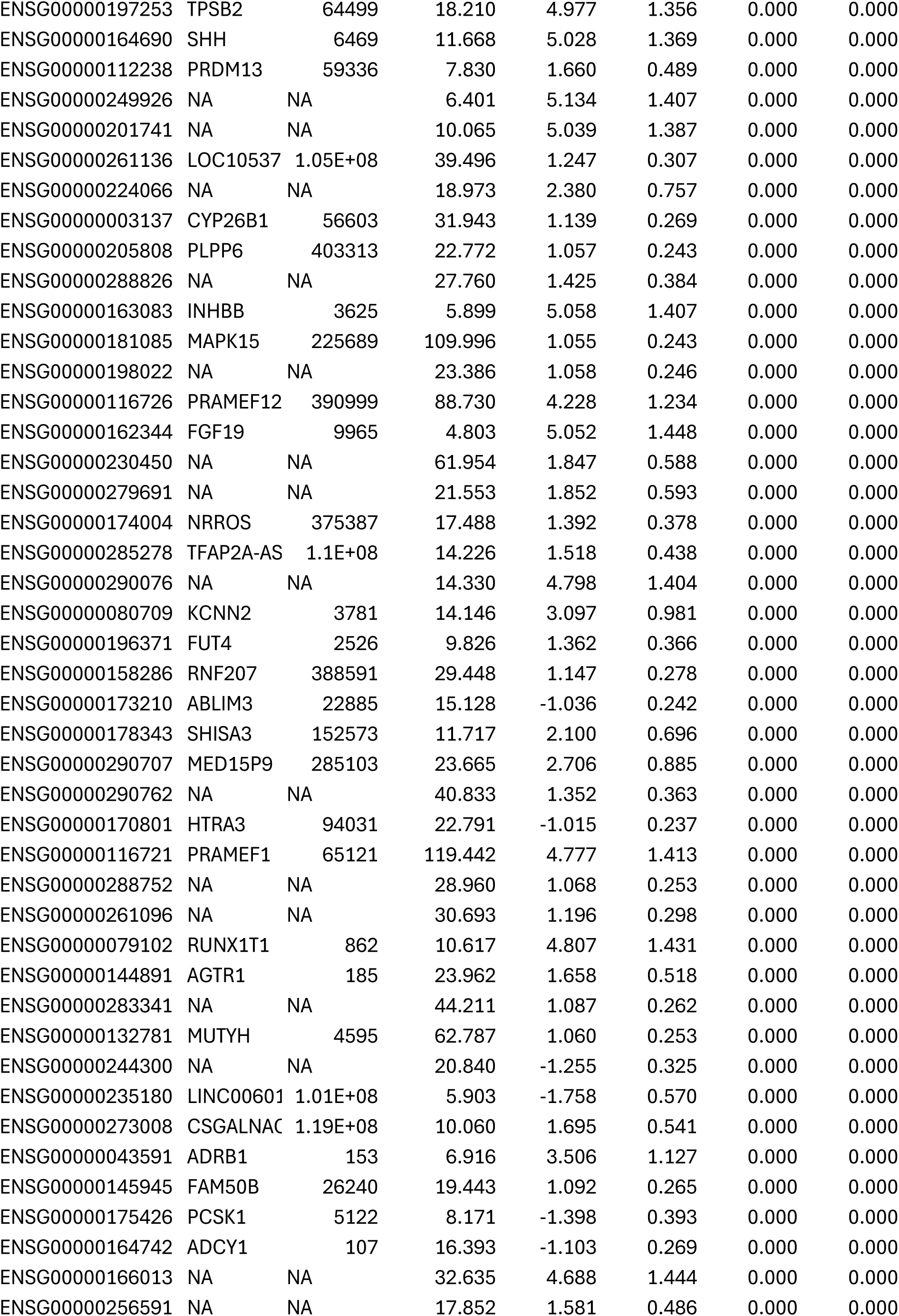
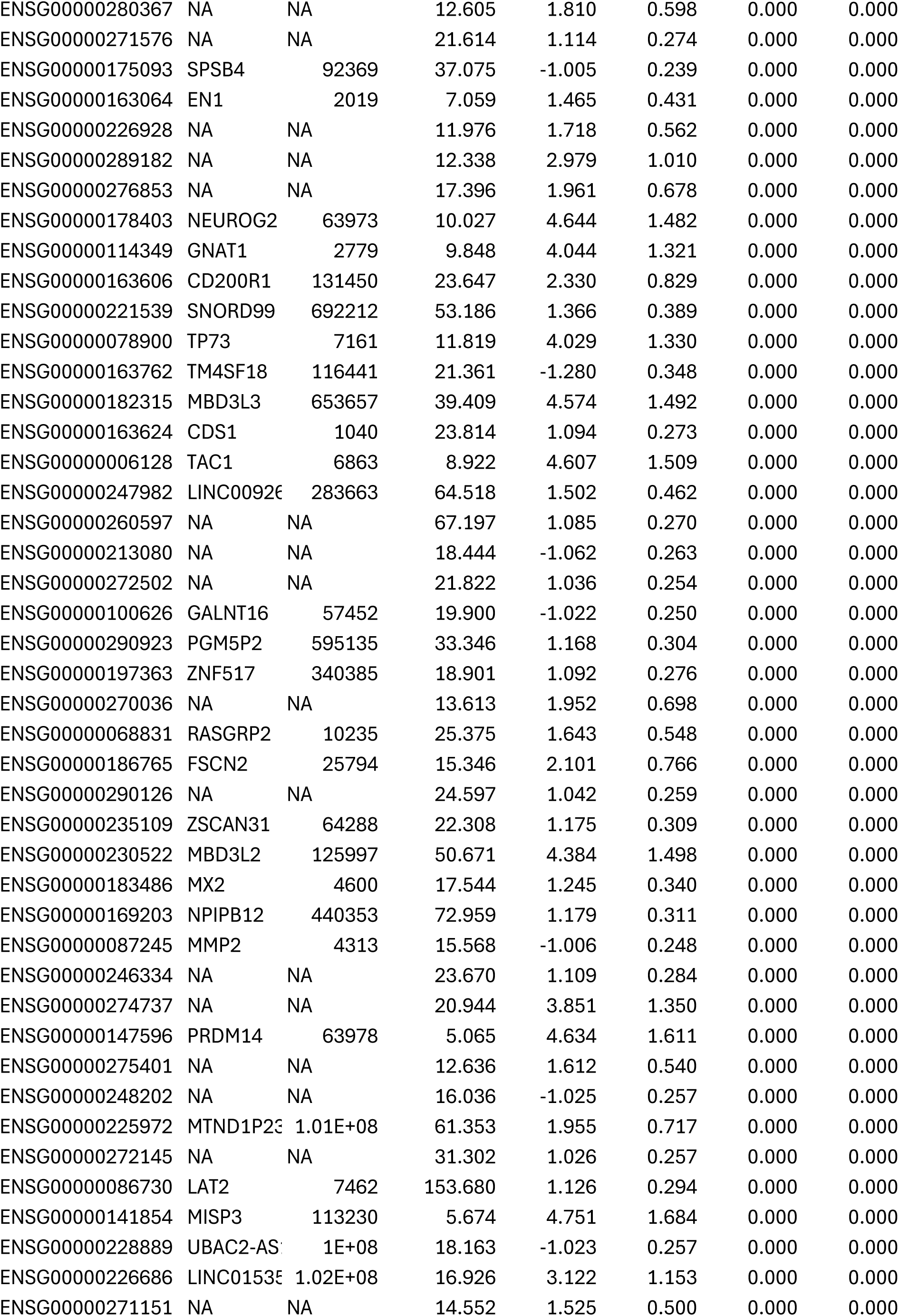
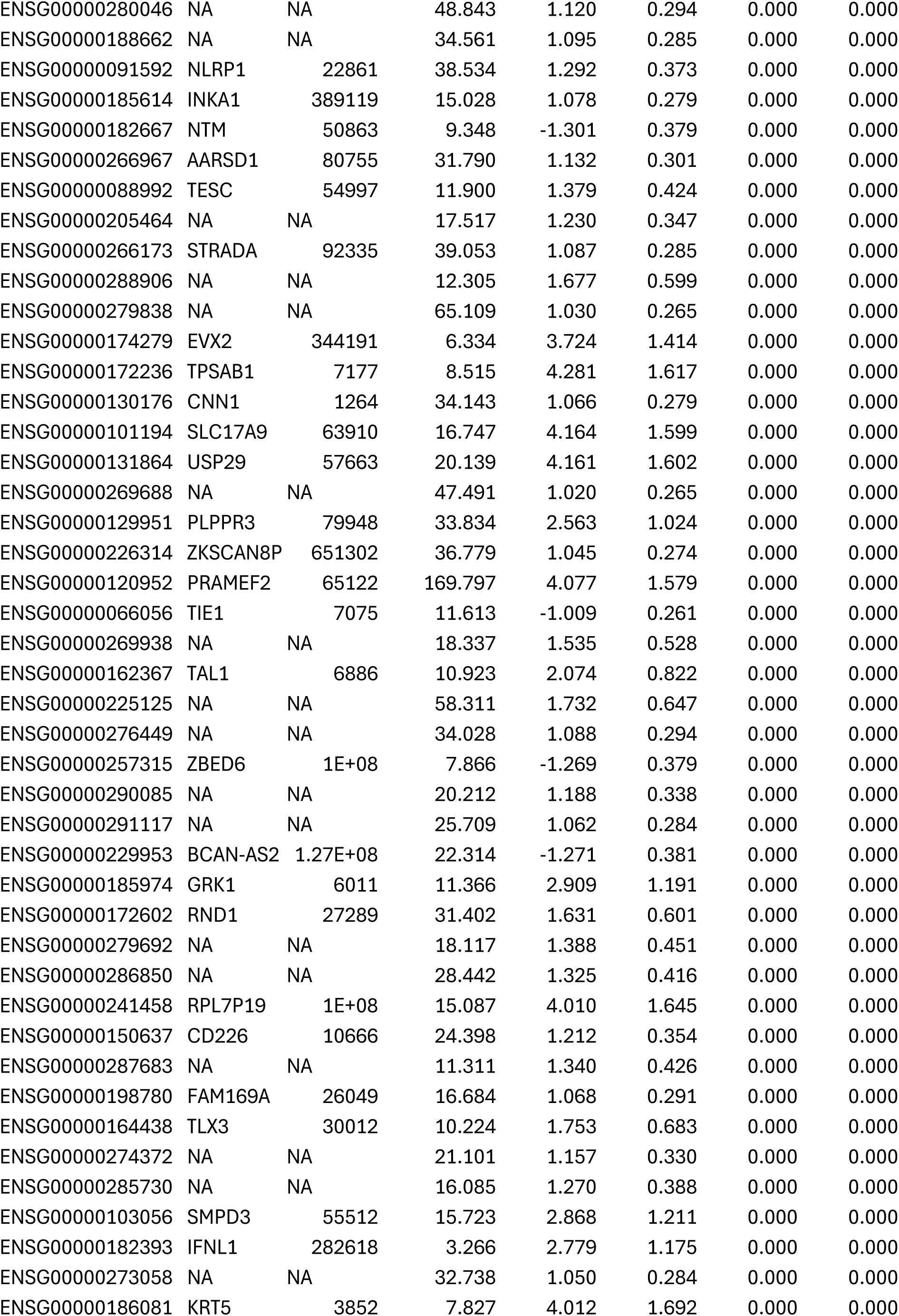
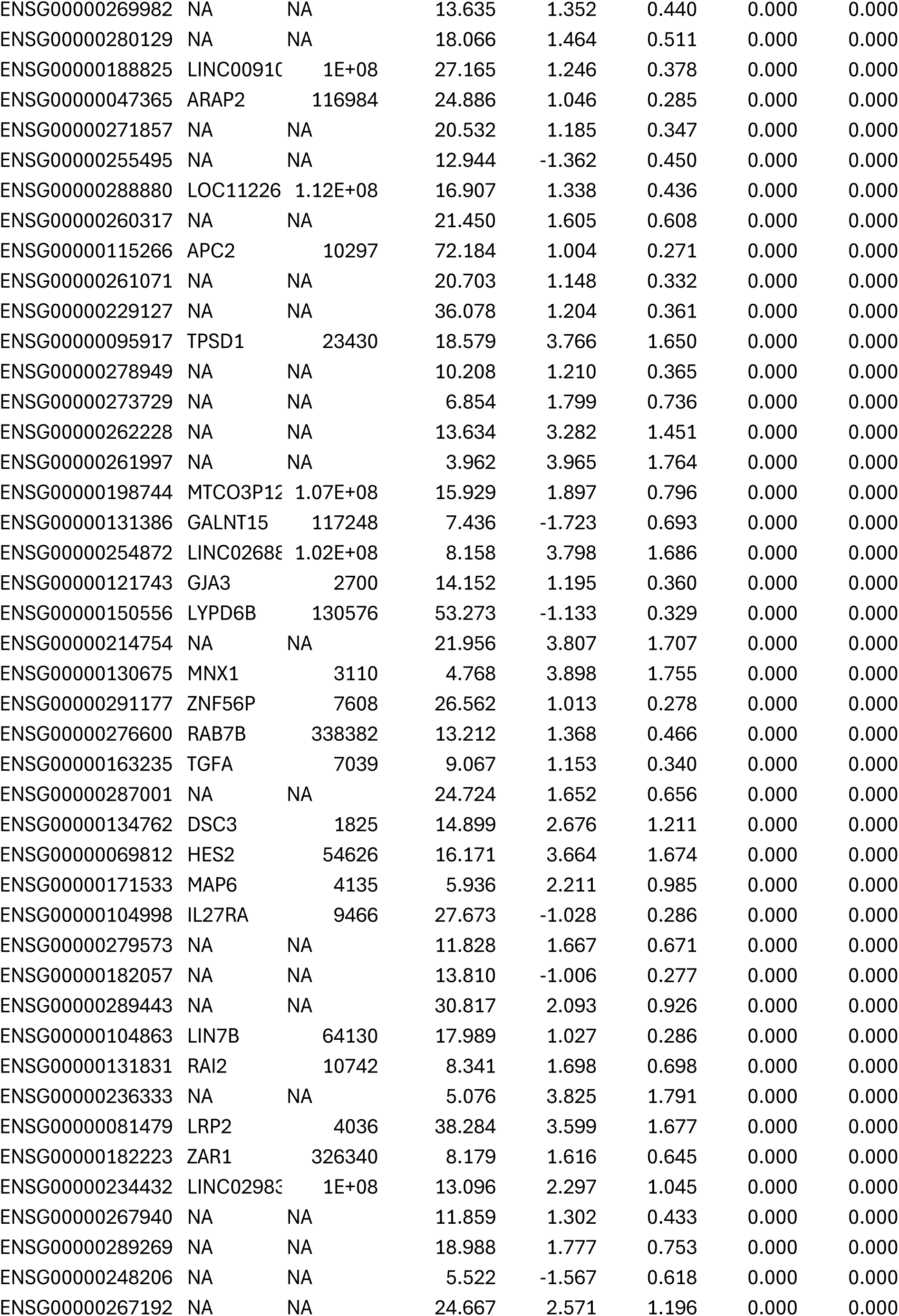
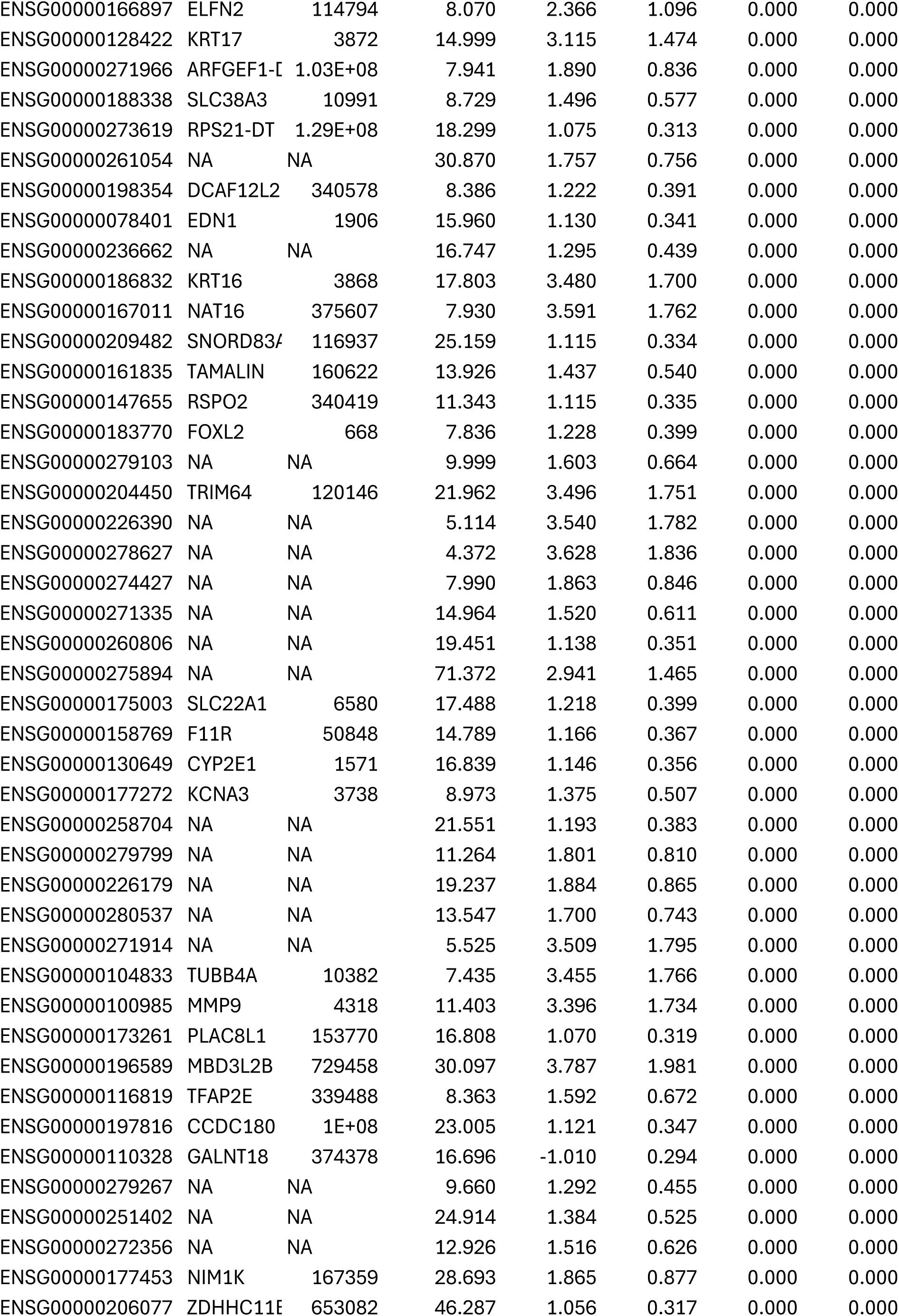
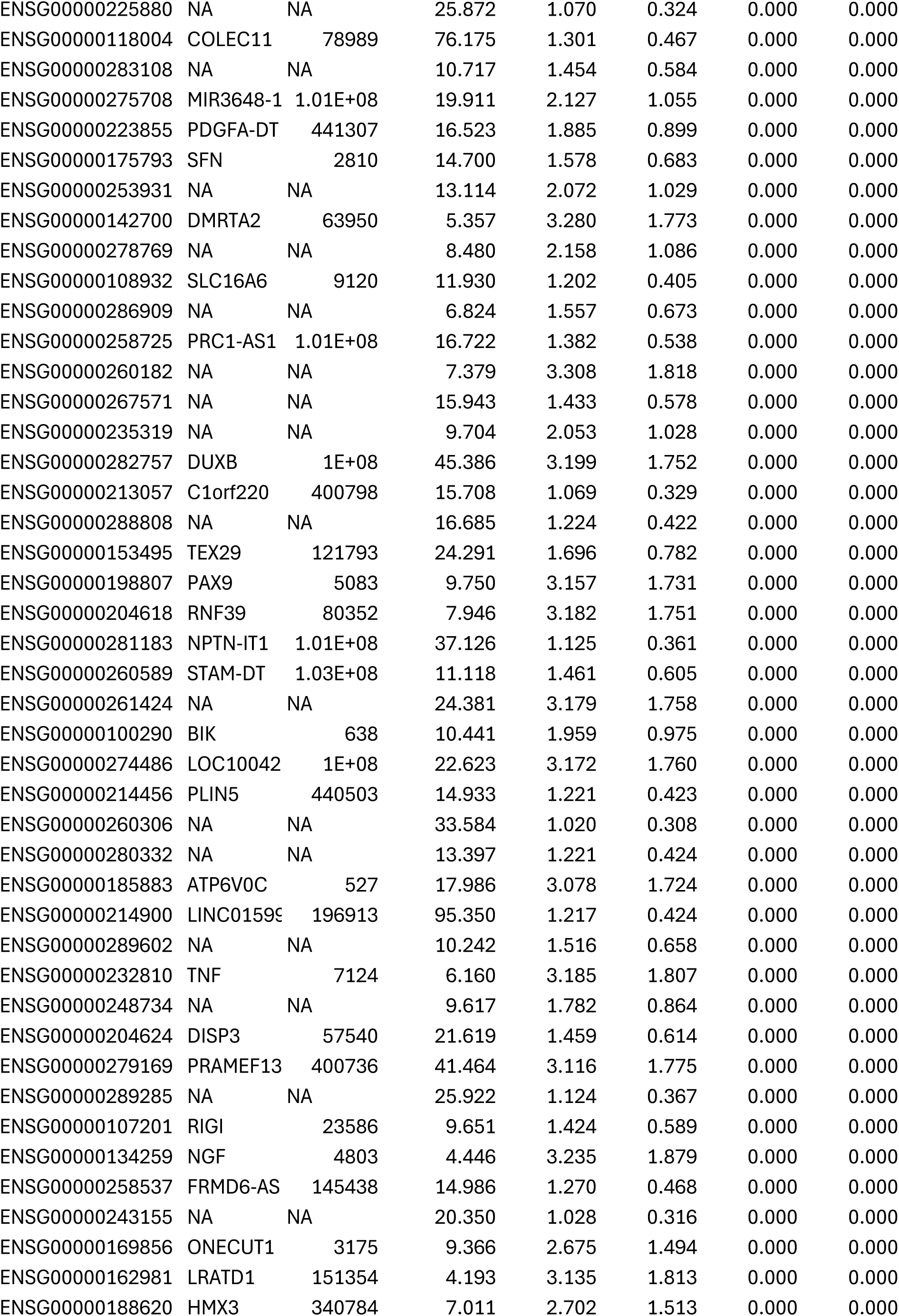
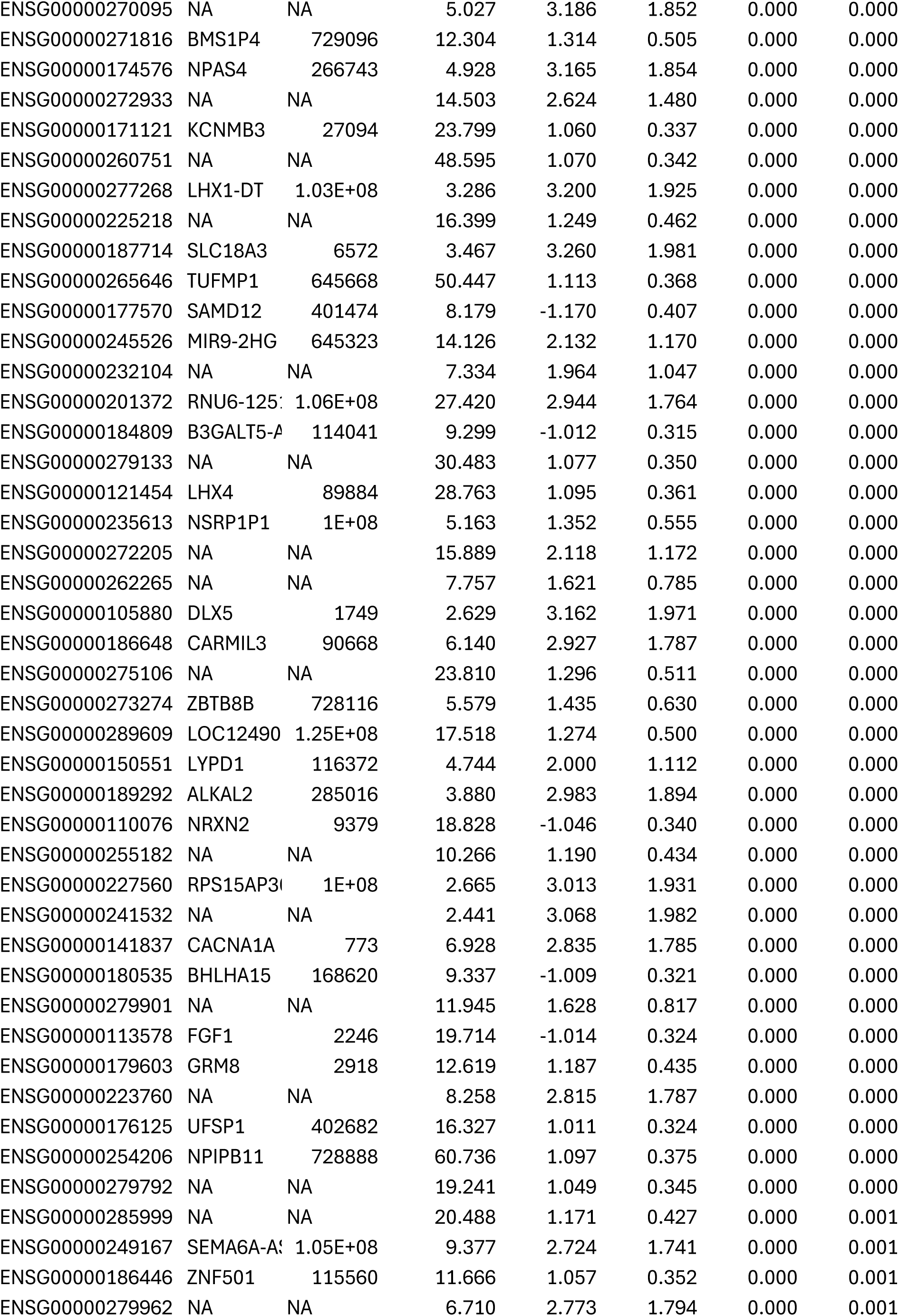
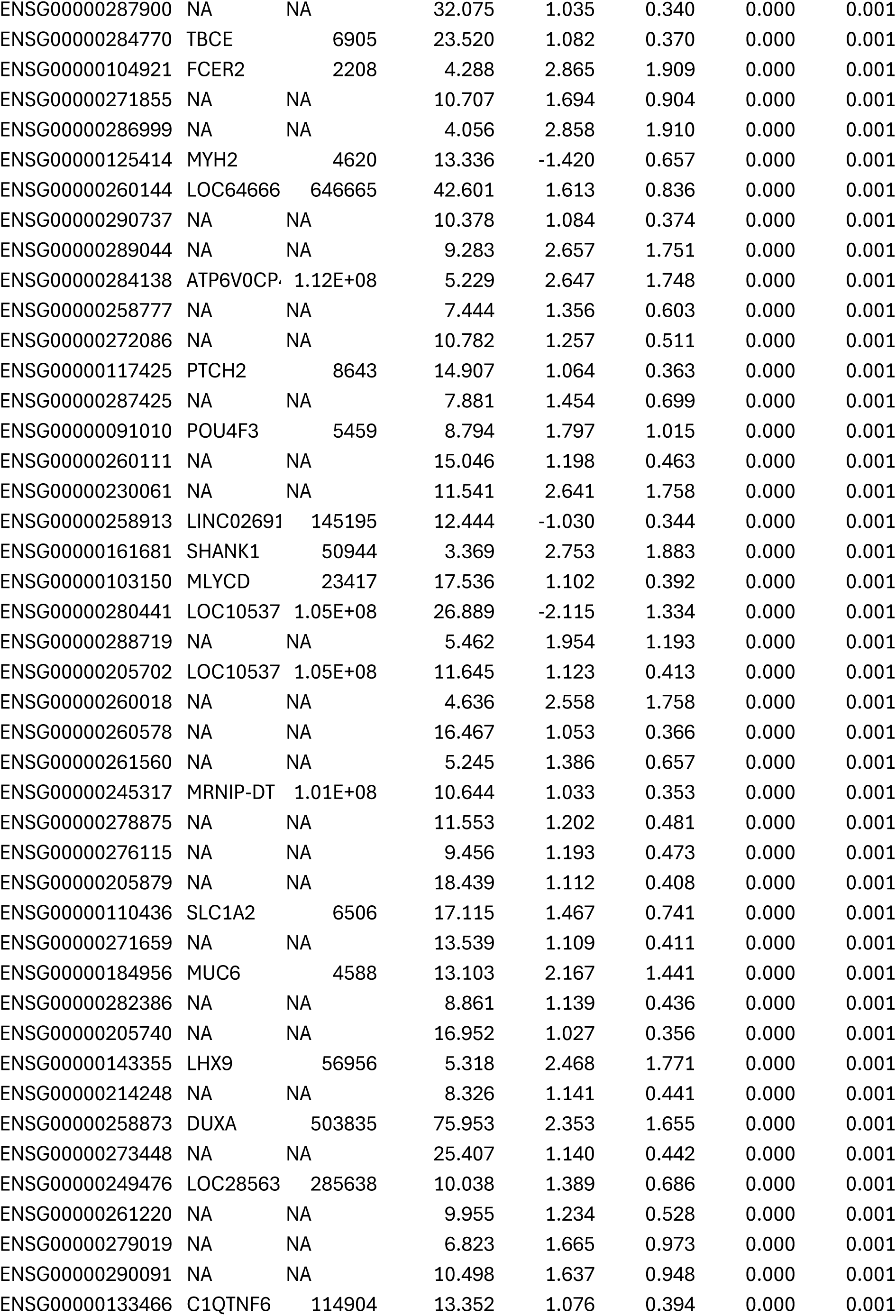
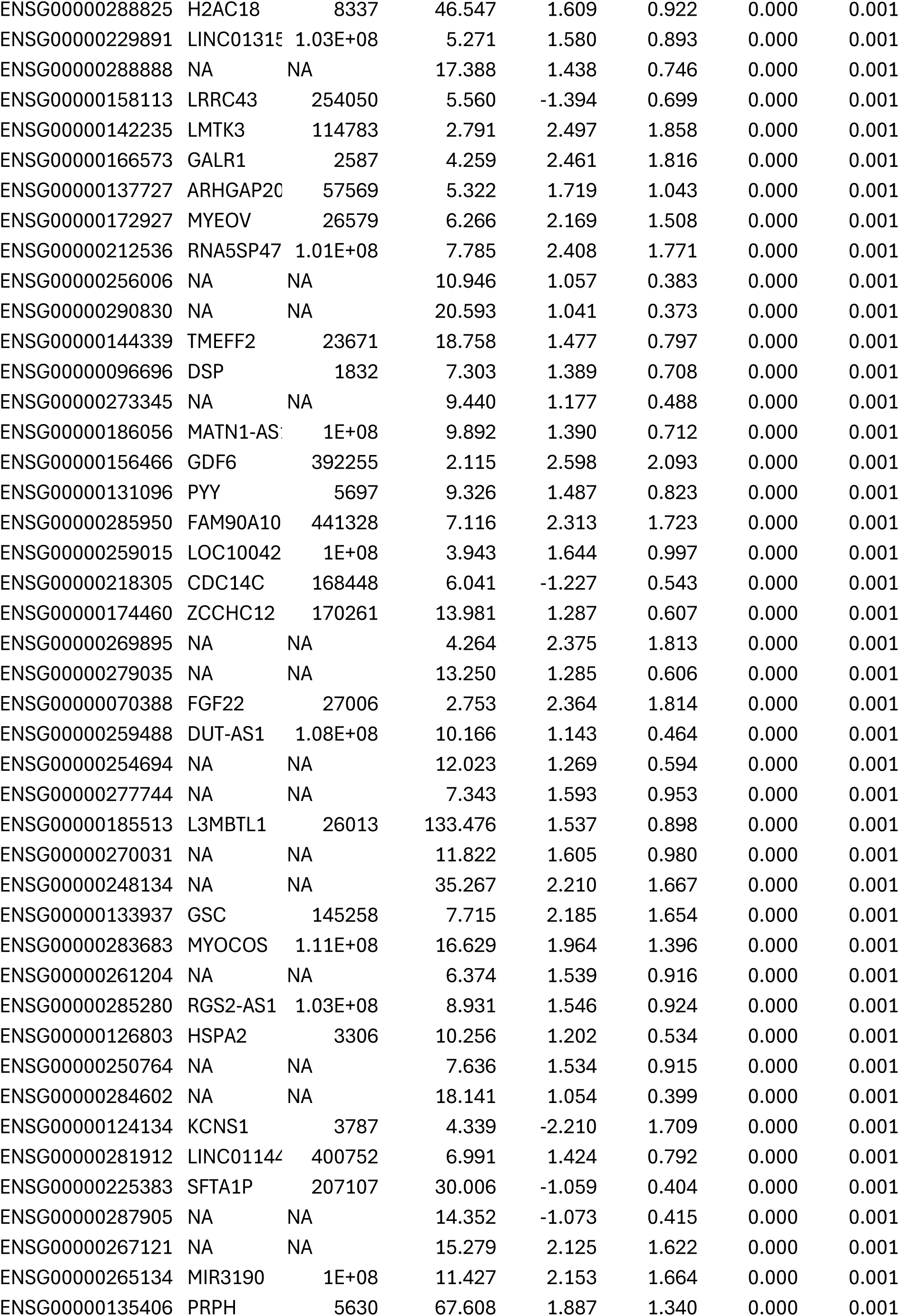
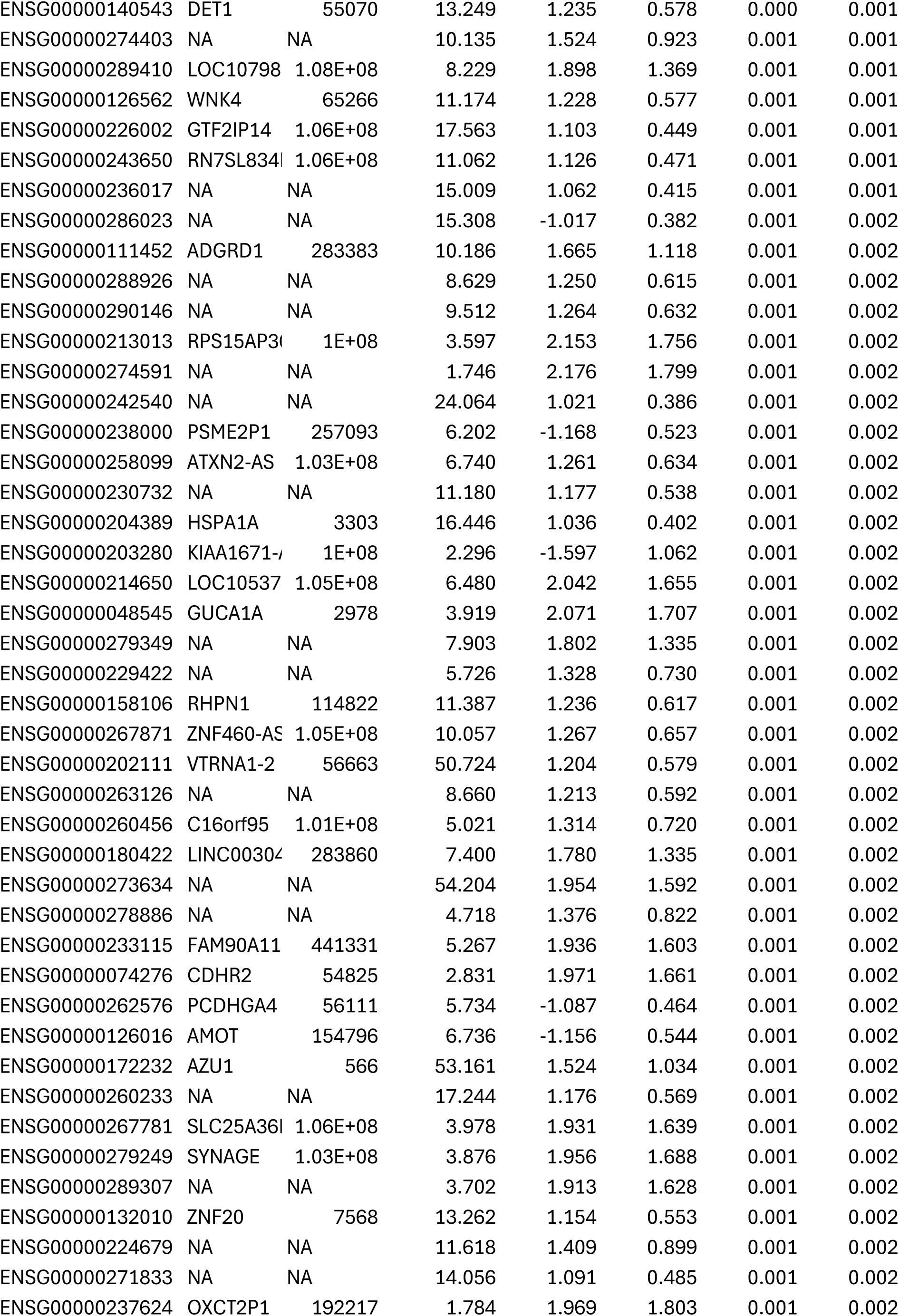
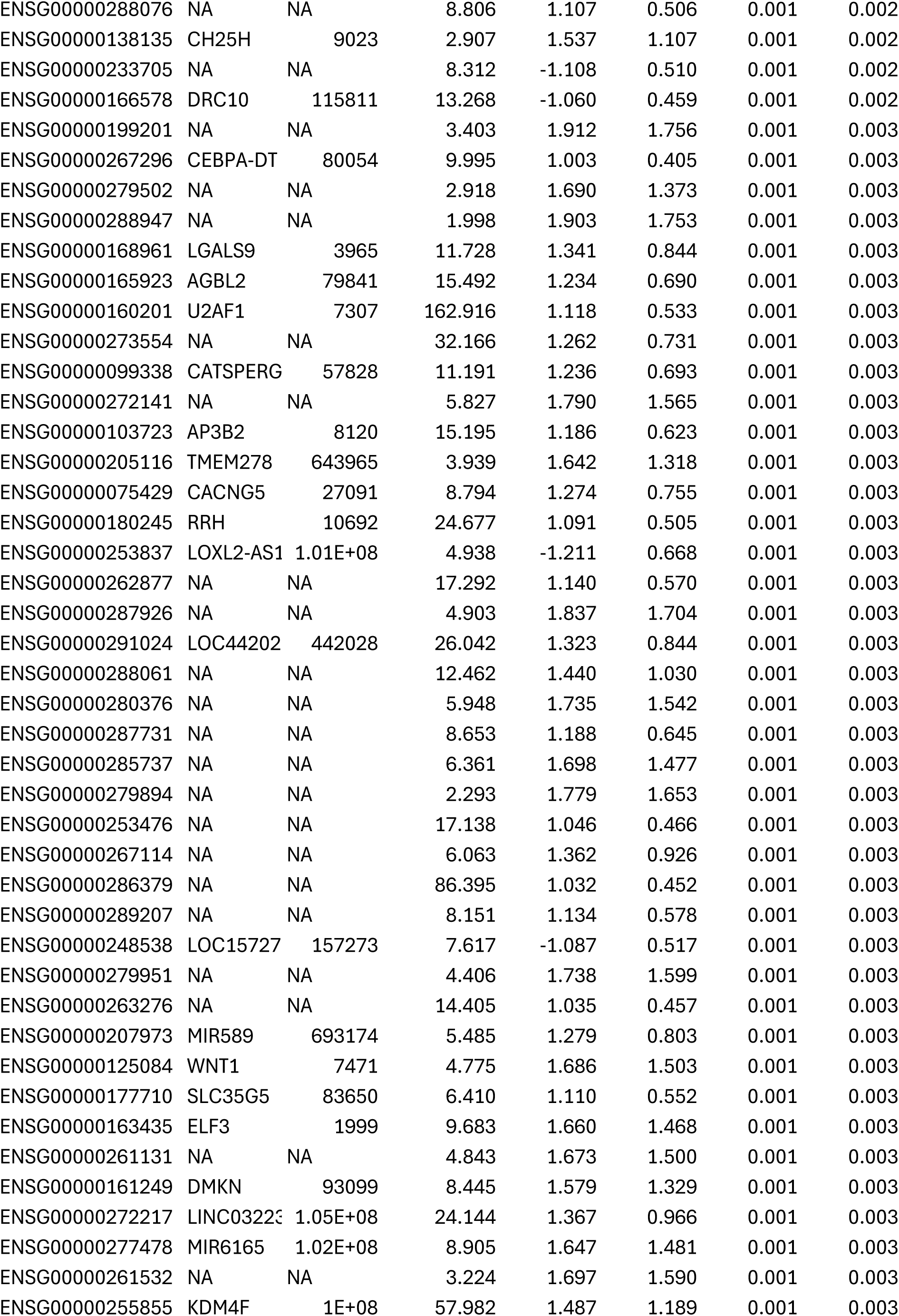
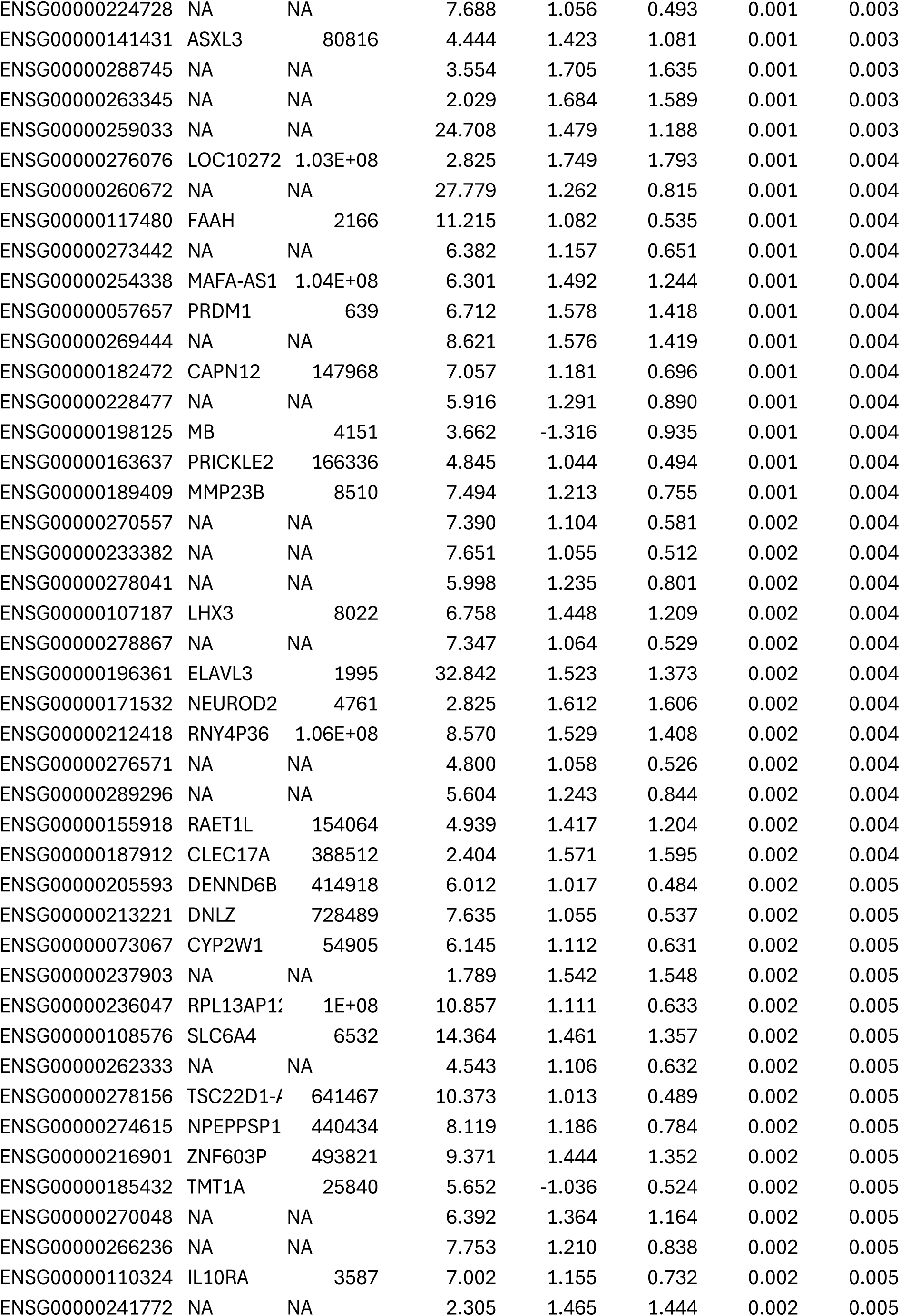
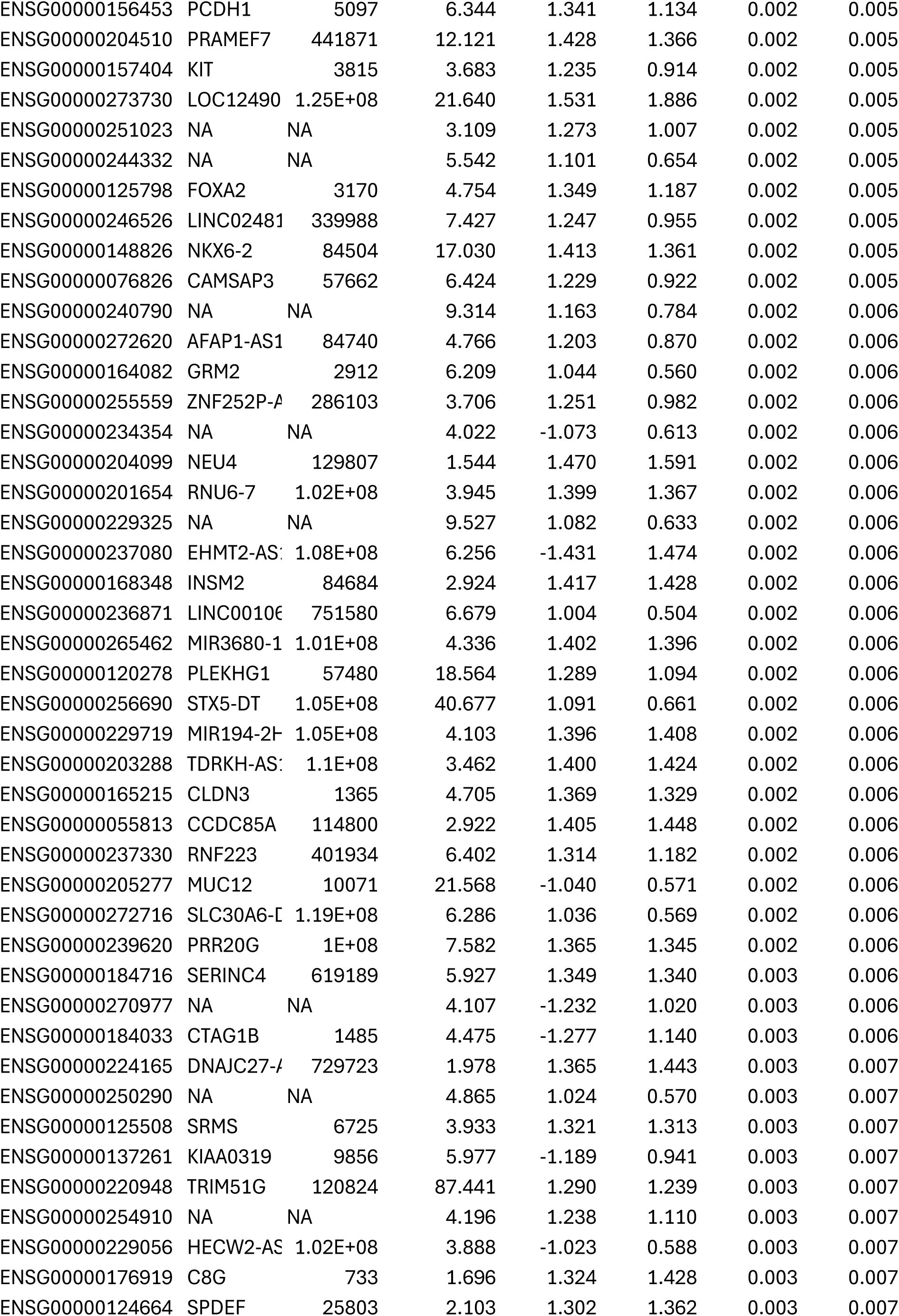
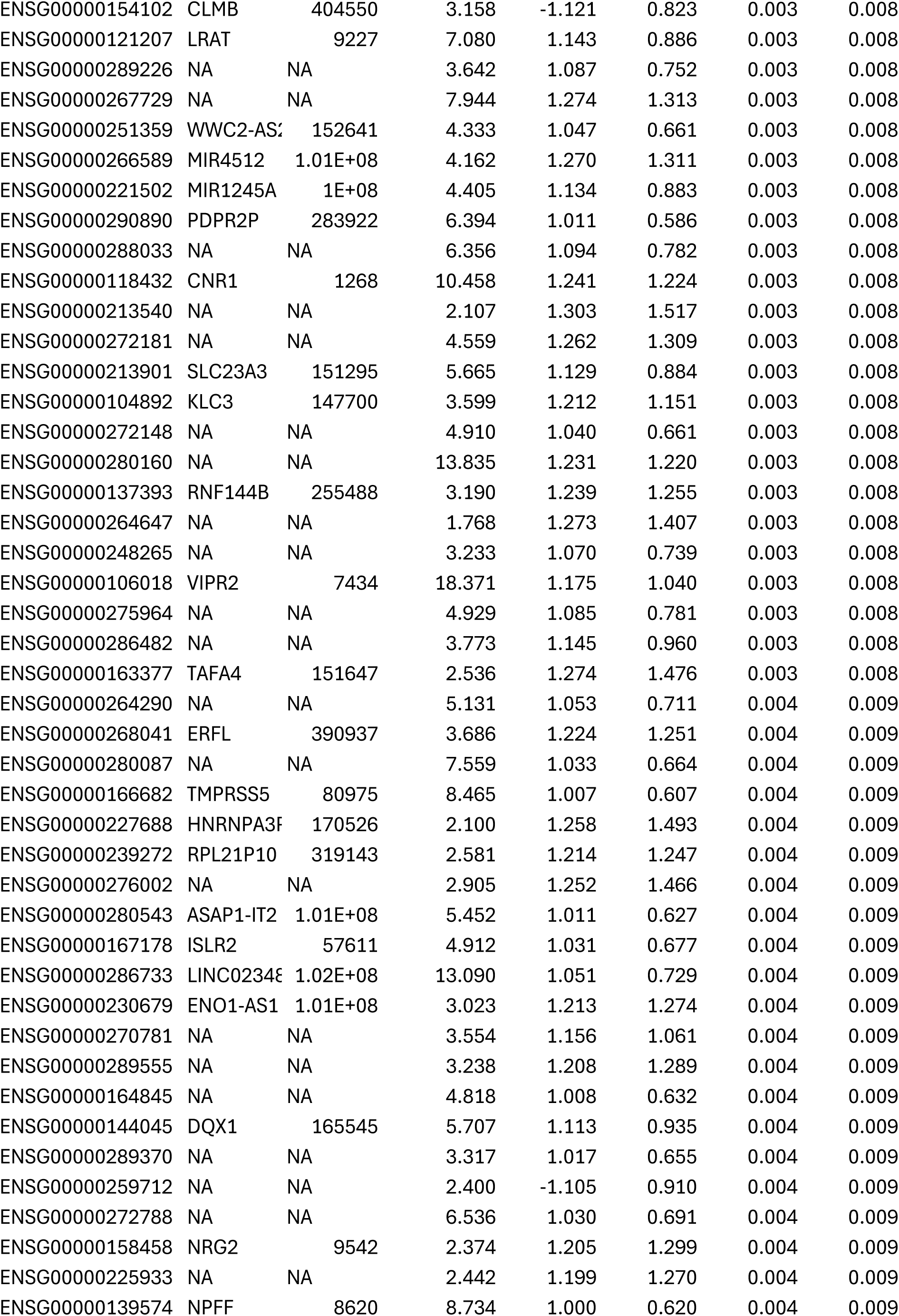
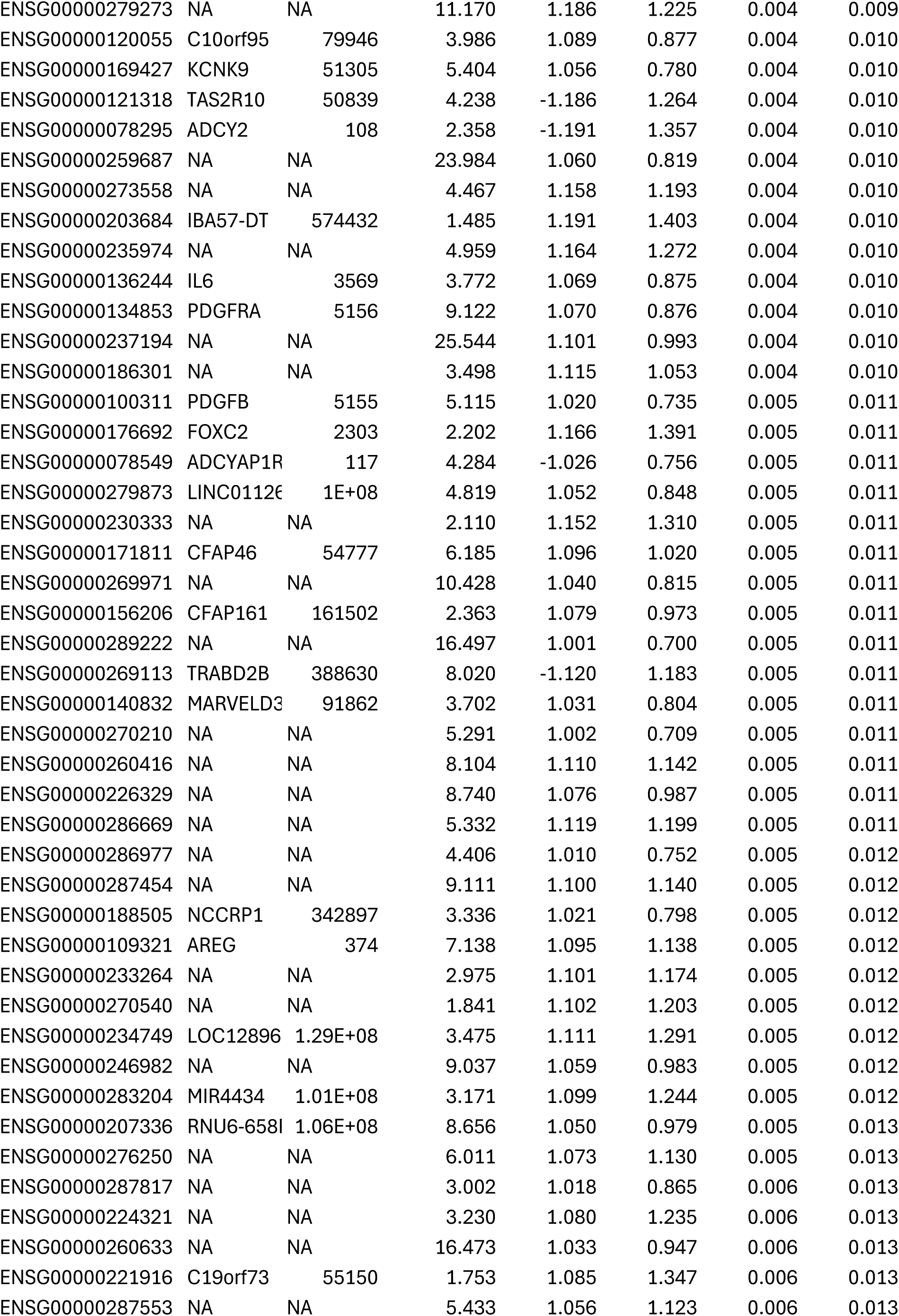
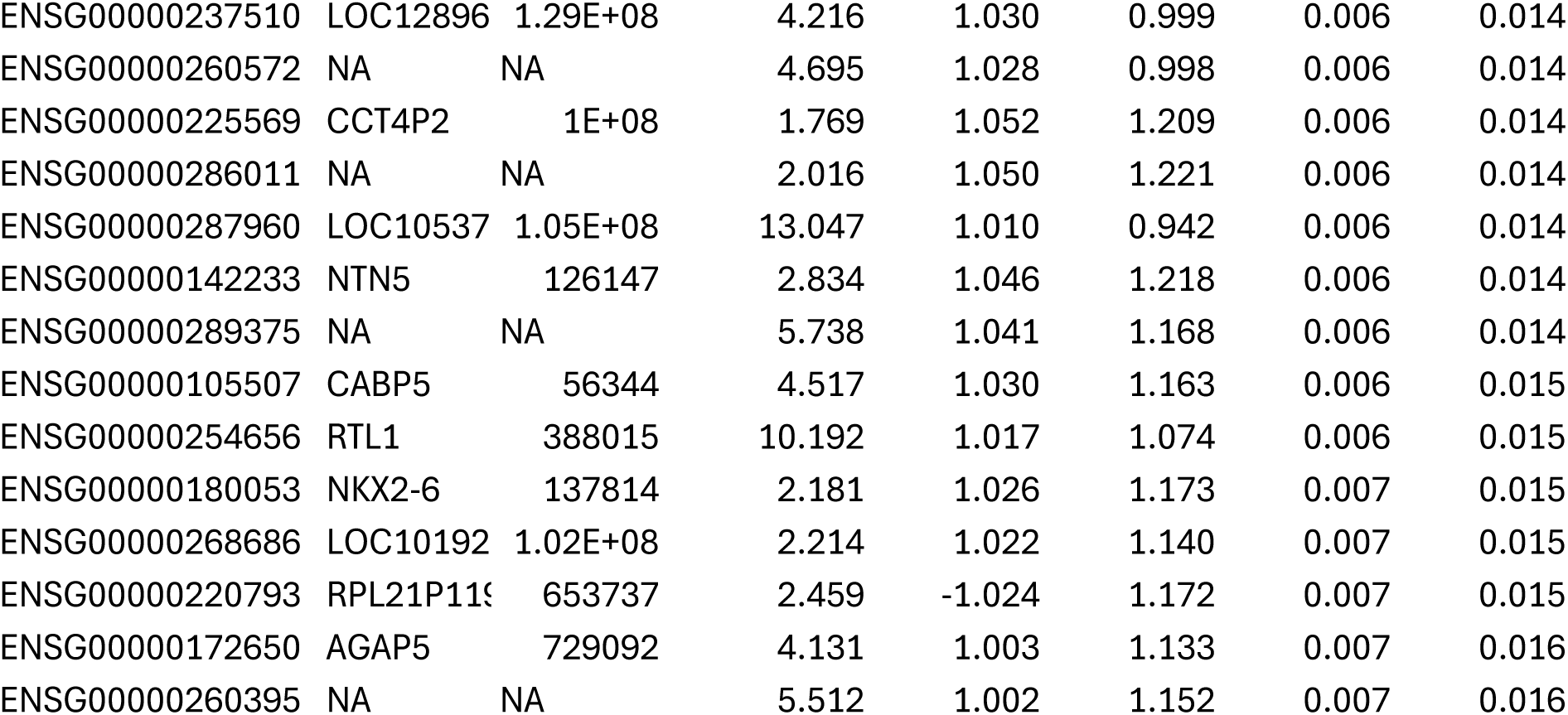

Significant DEGs during infection of RD:P3F with oRP3Fus at 6 hours post infection versus Mock RD:P3F.

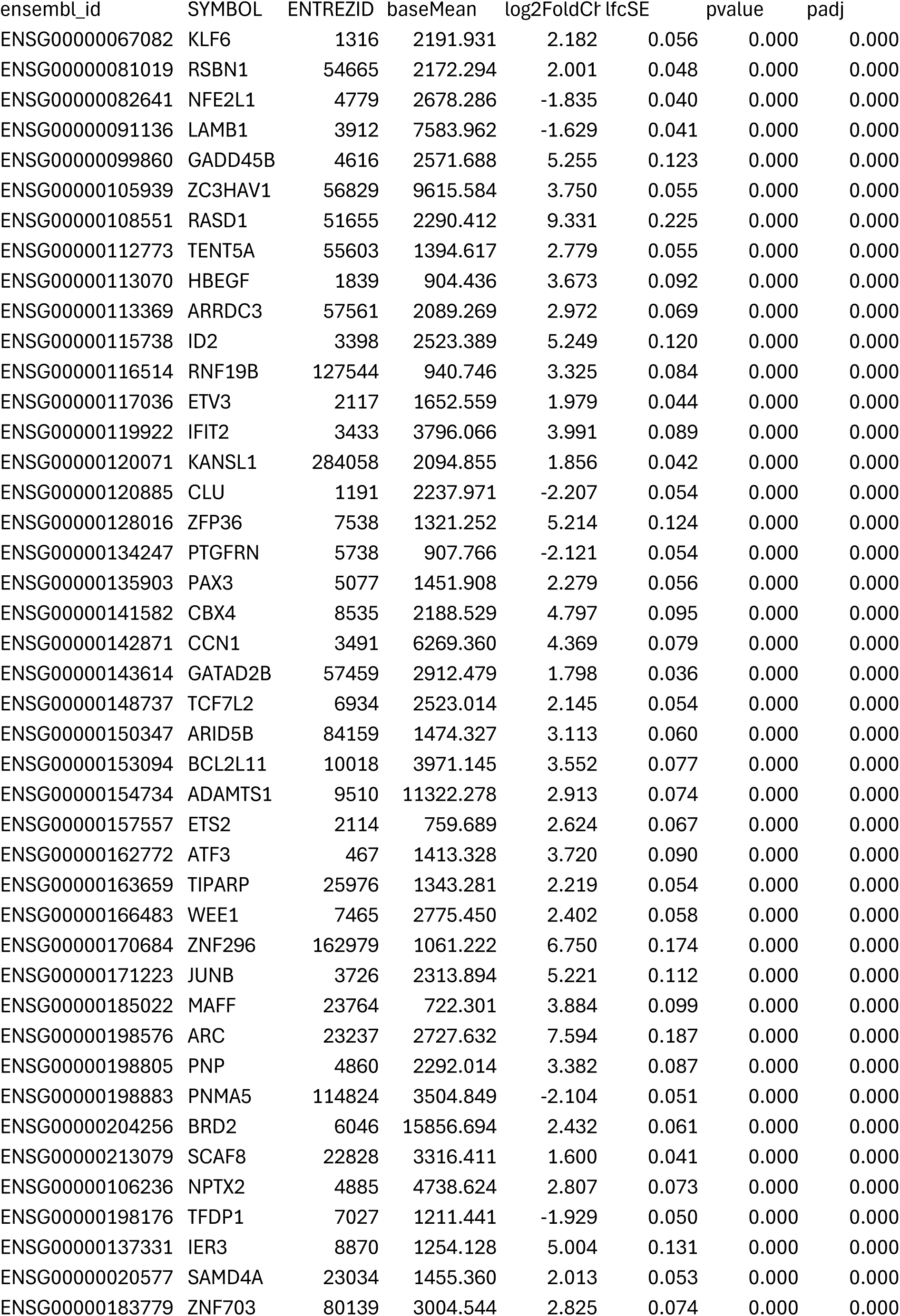
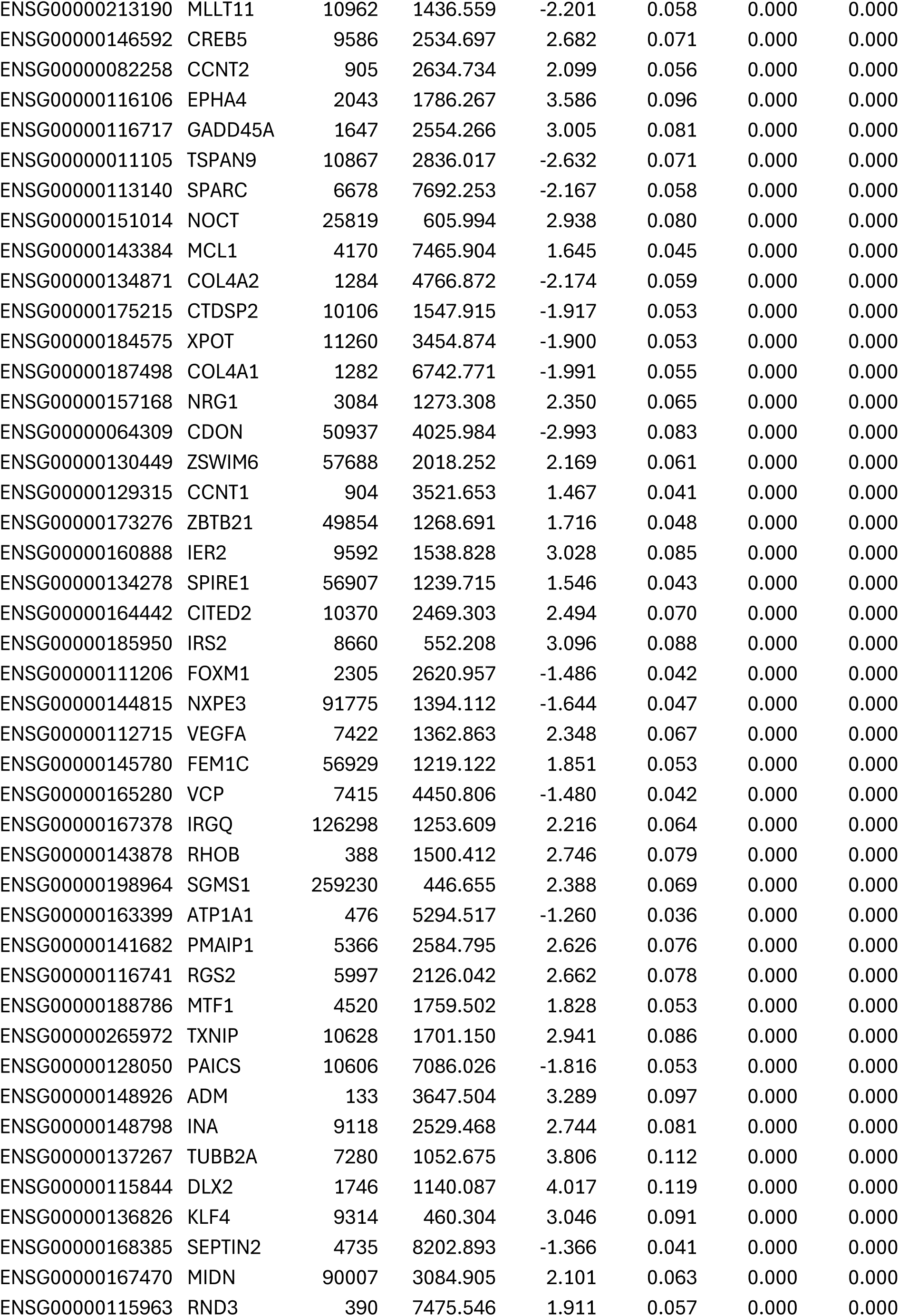
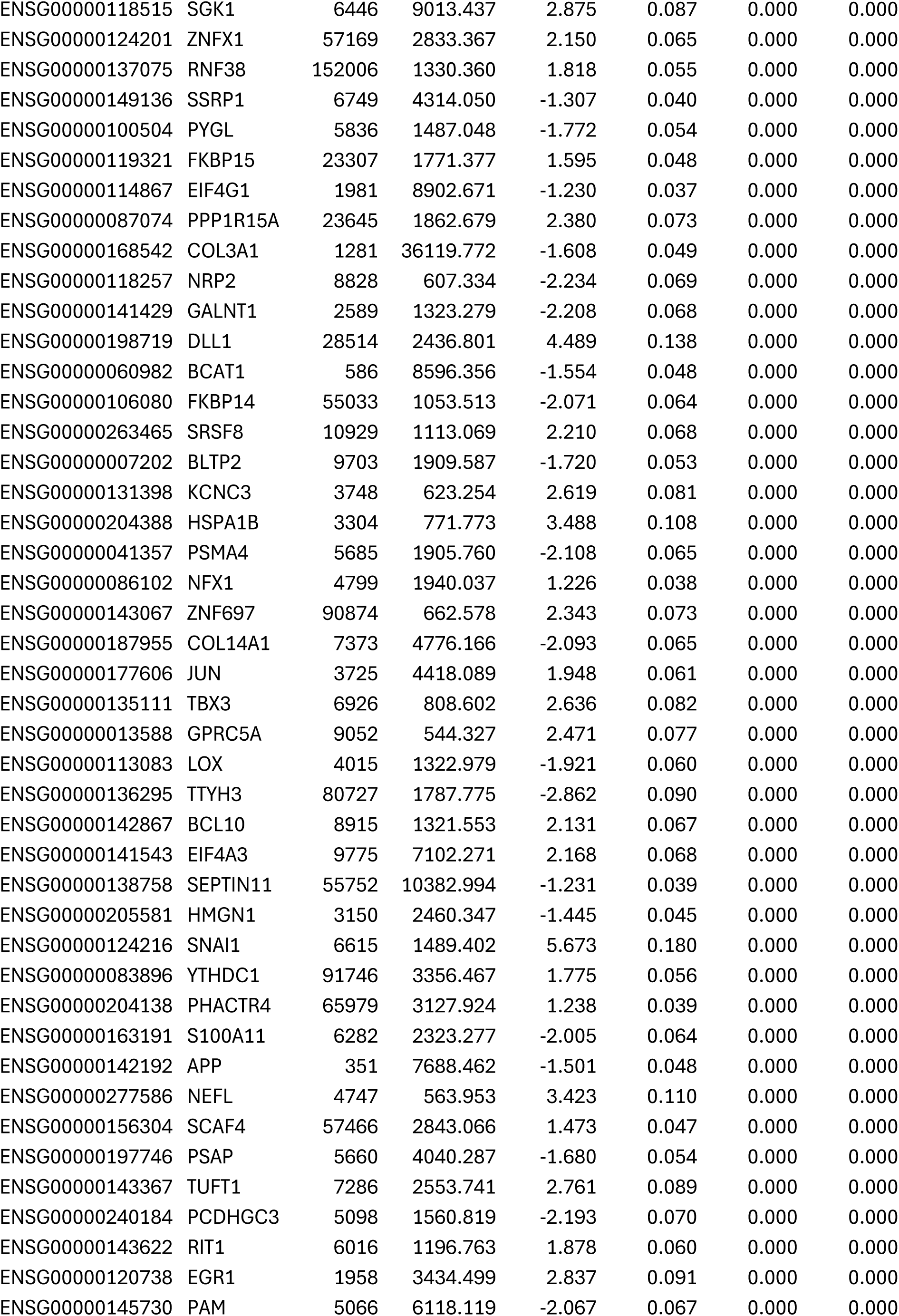
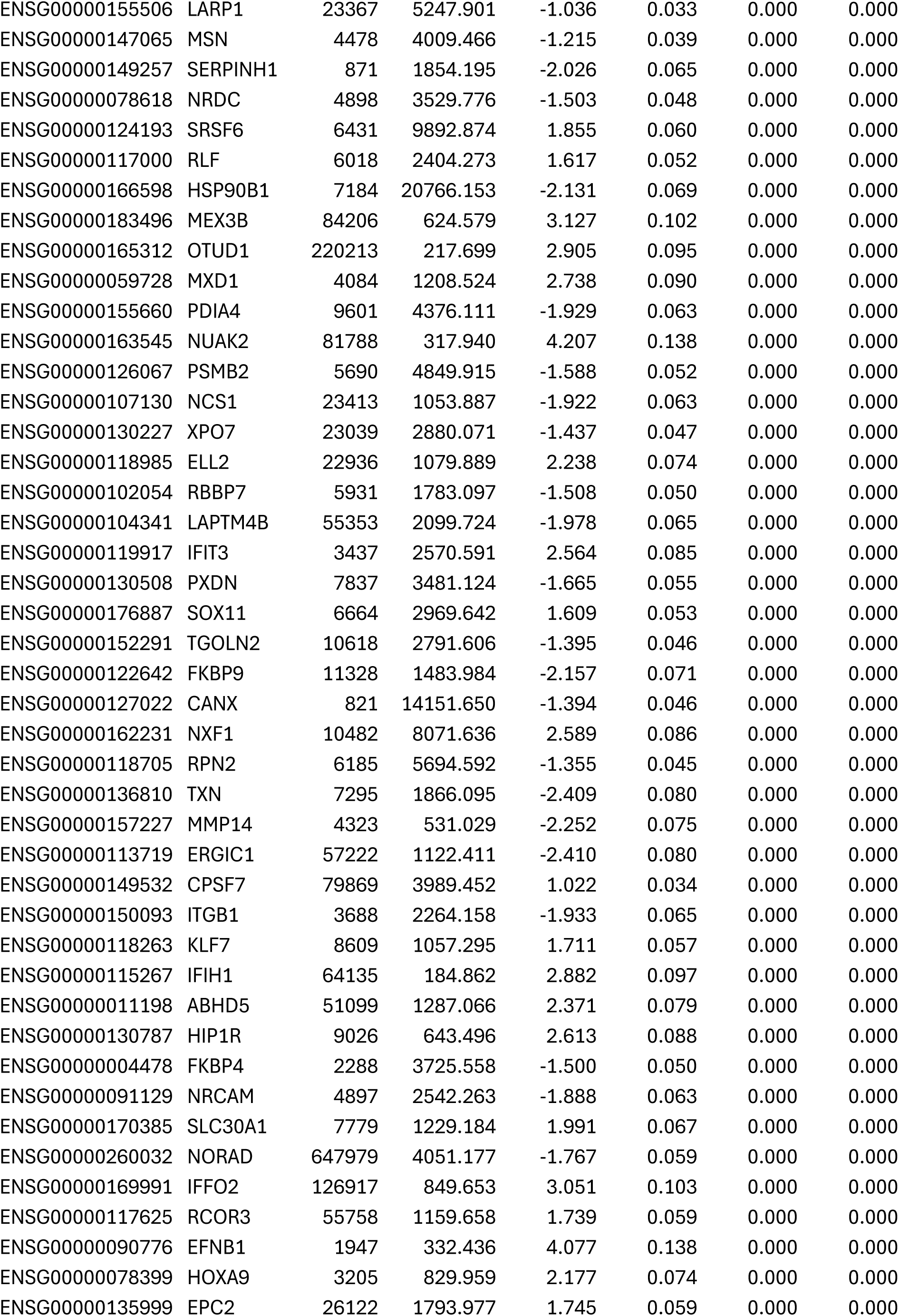
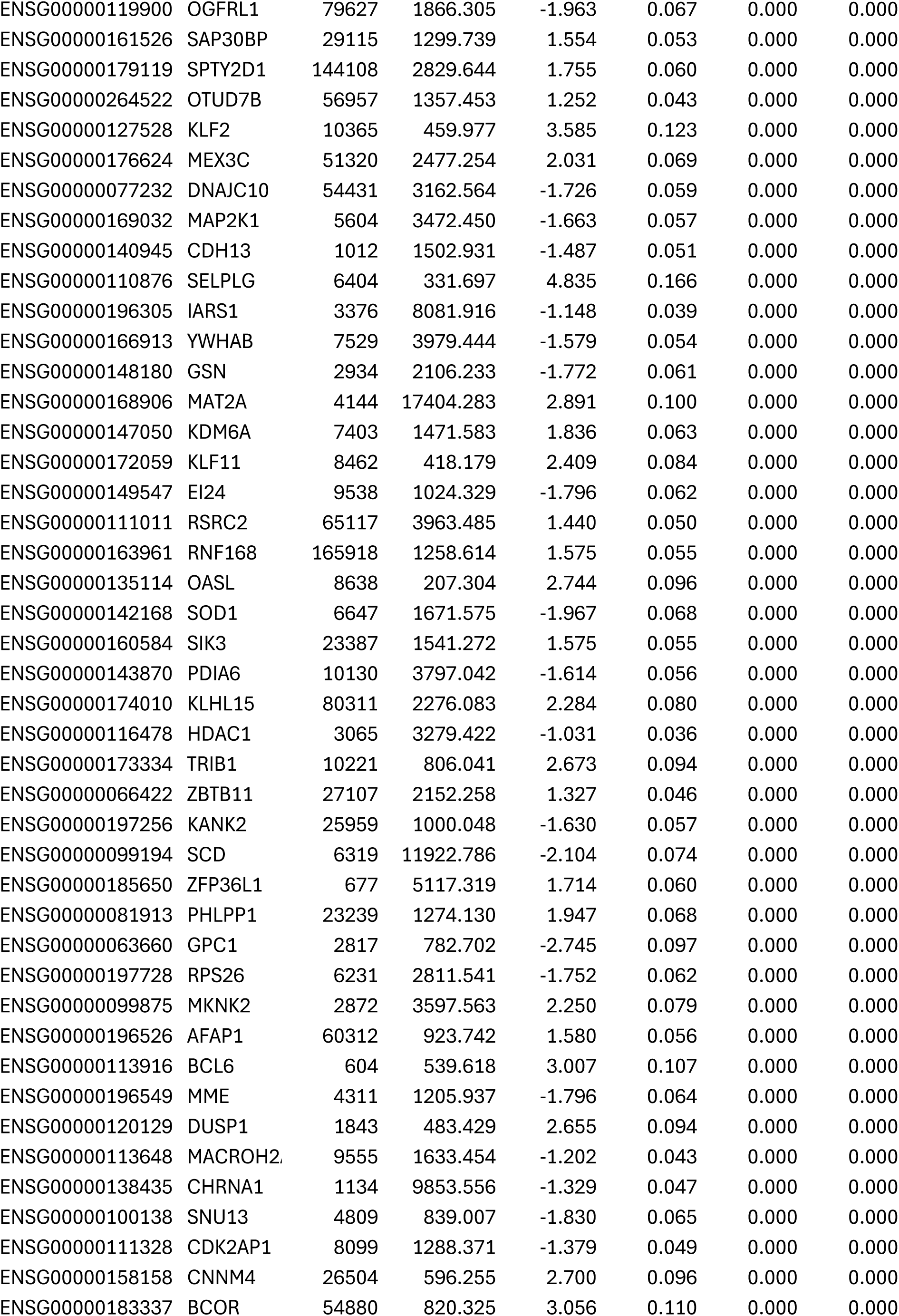
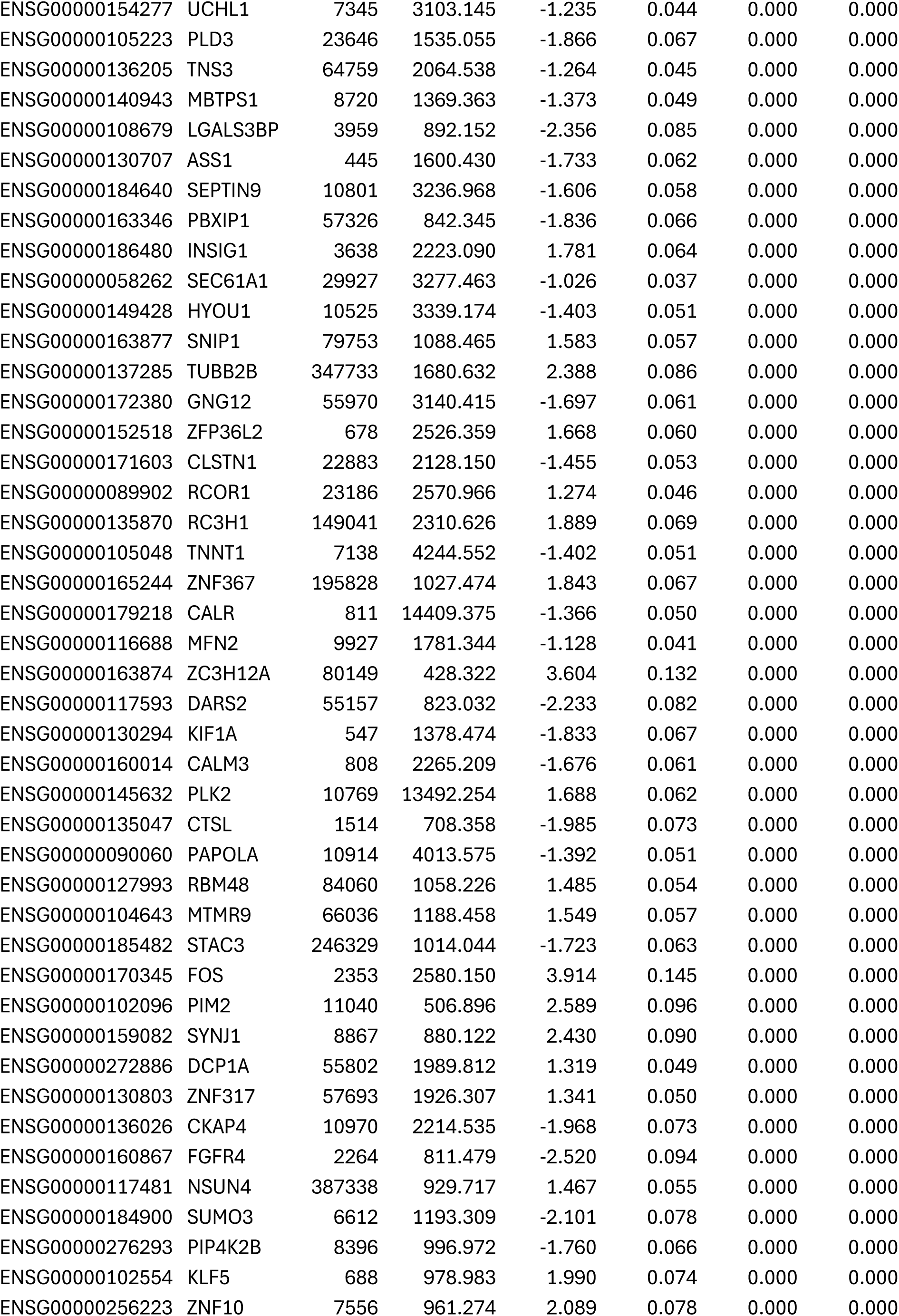
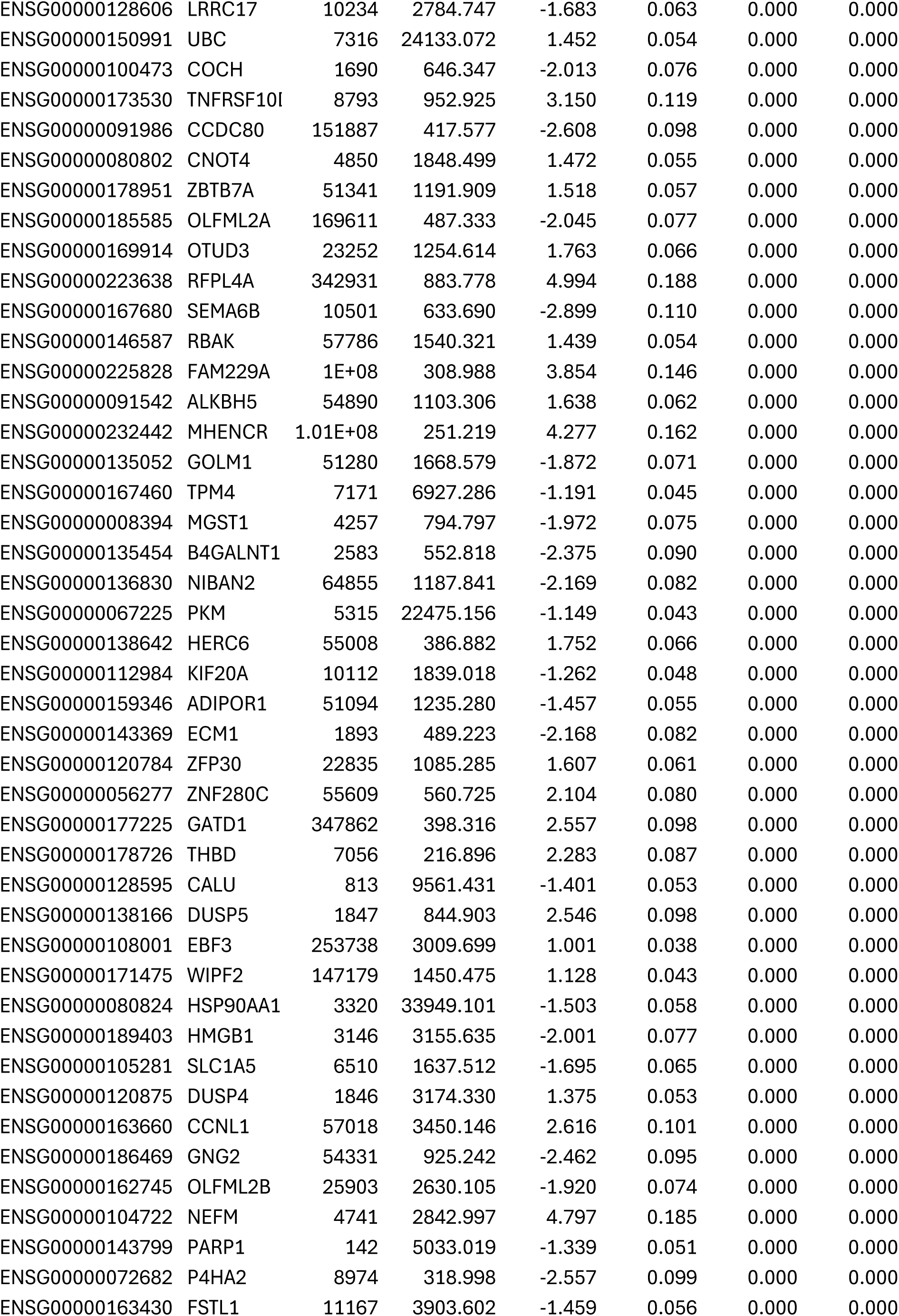
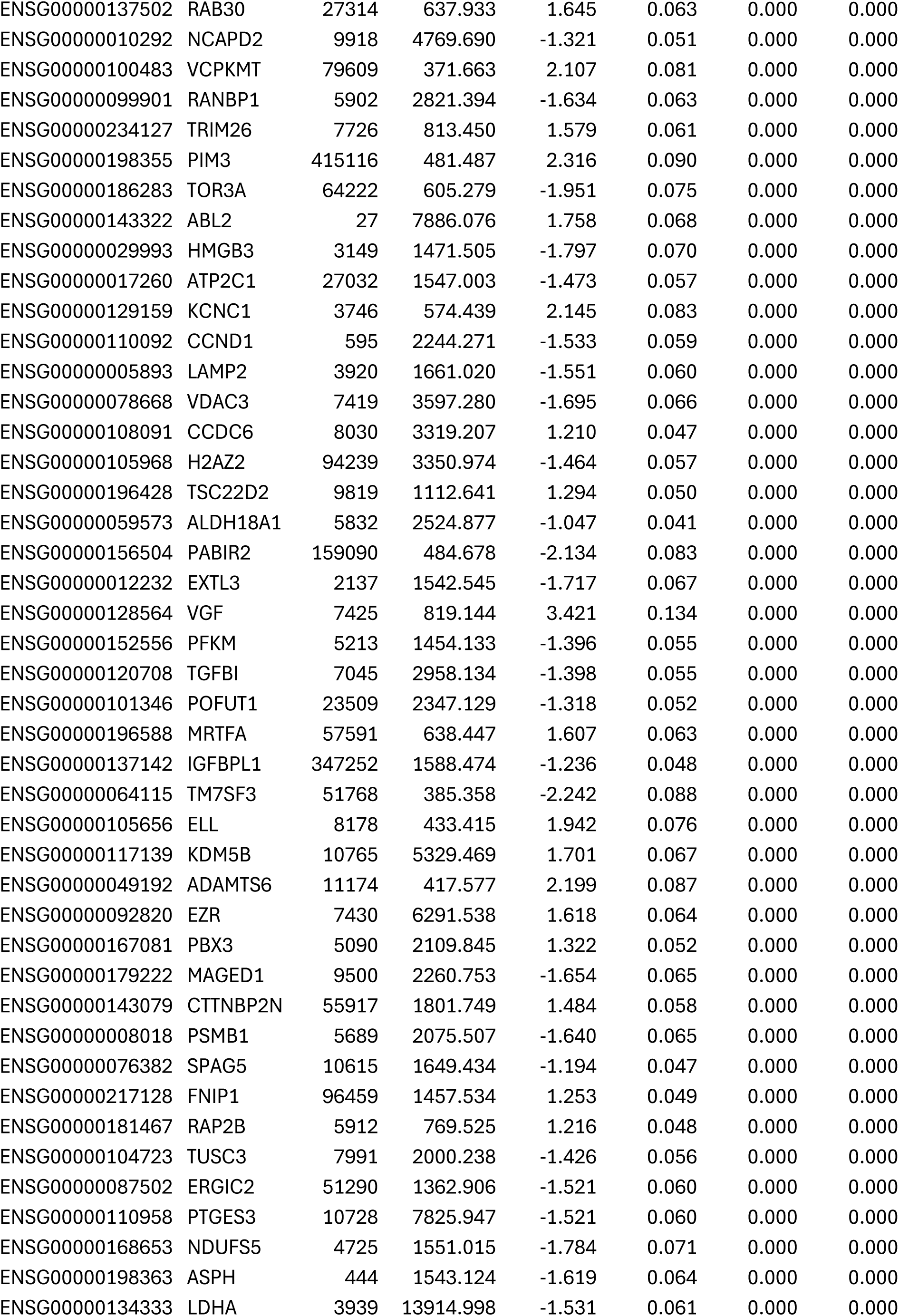
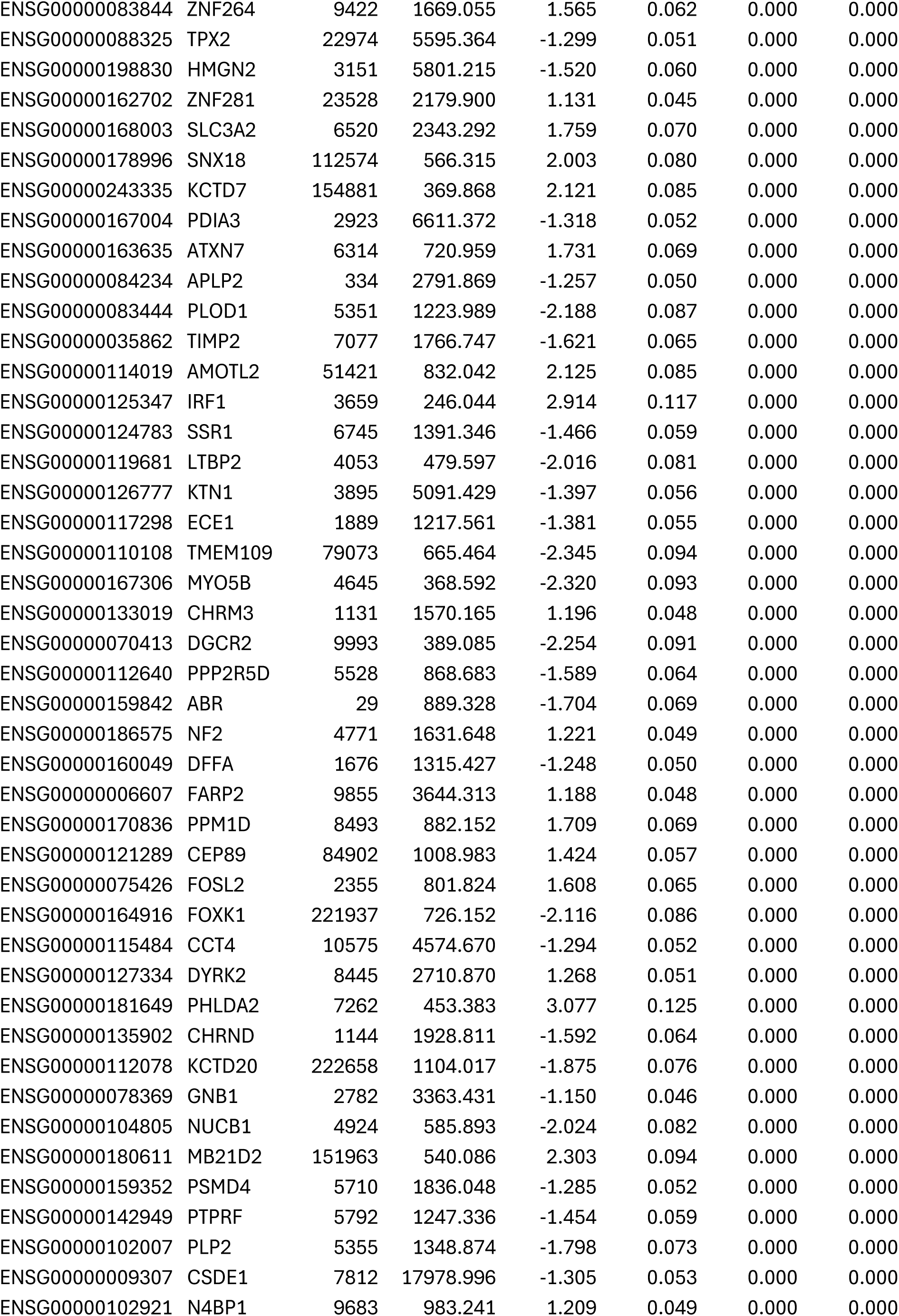
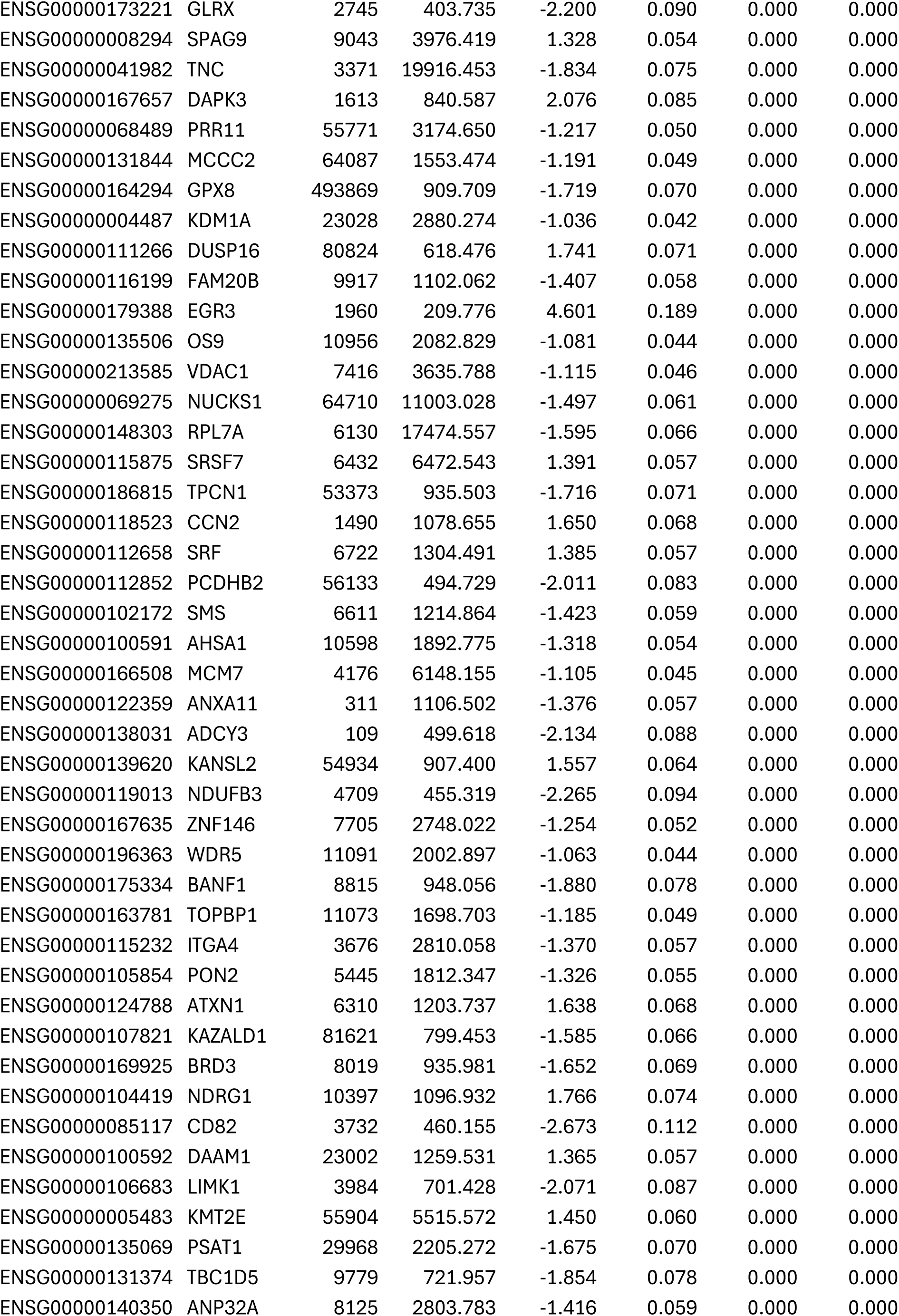
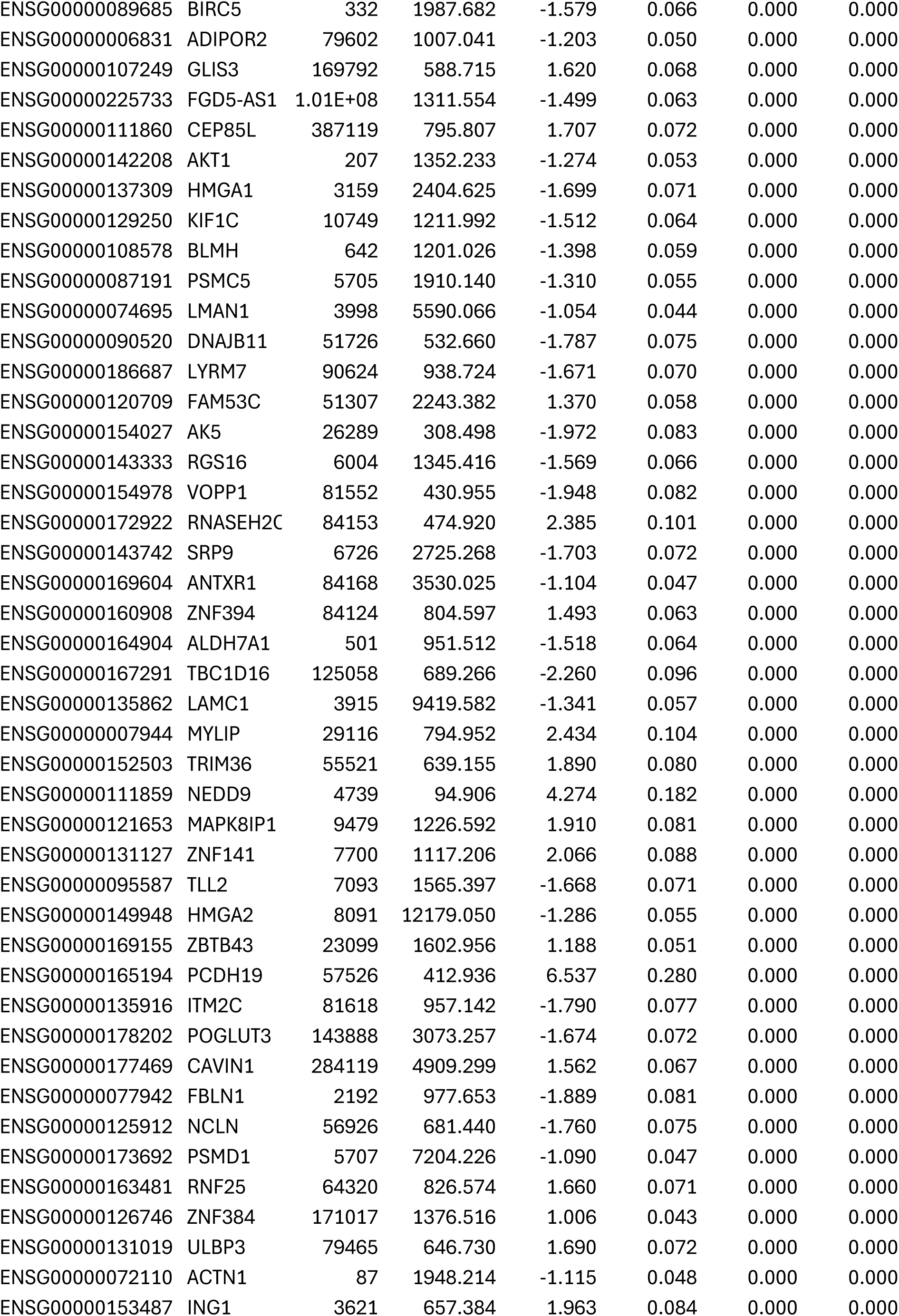
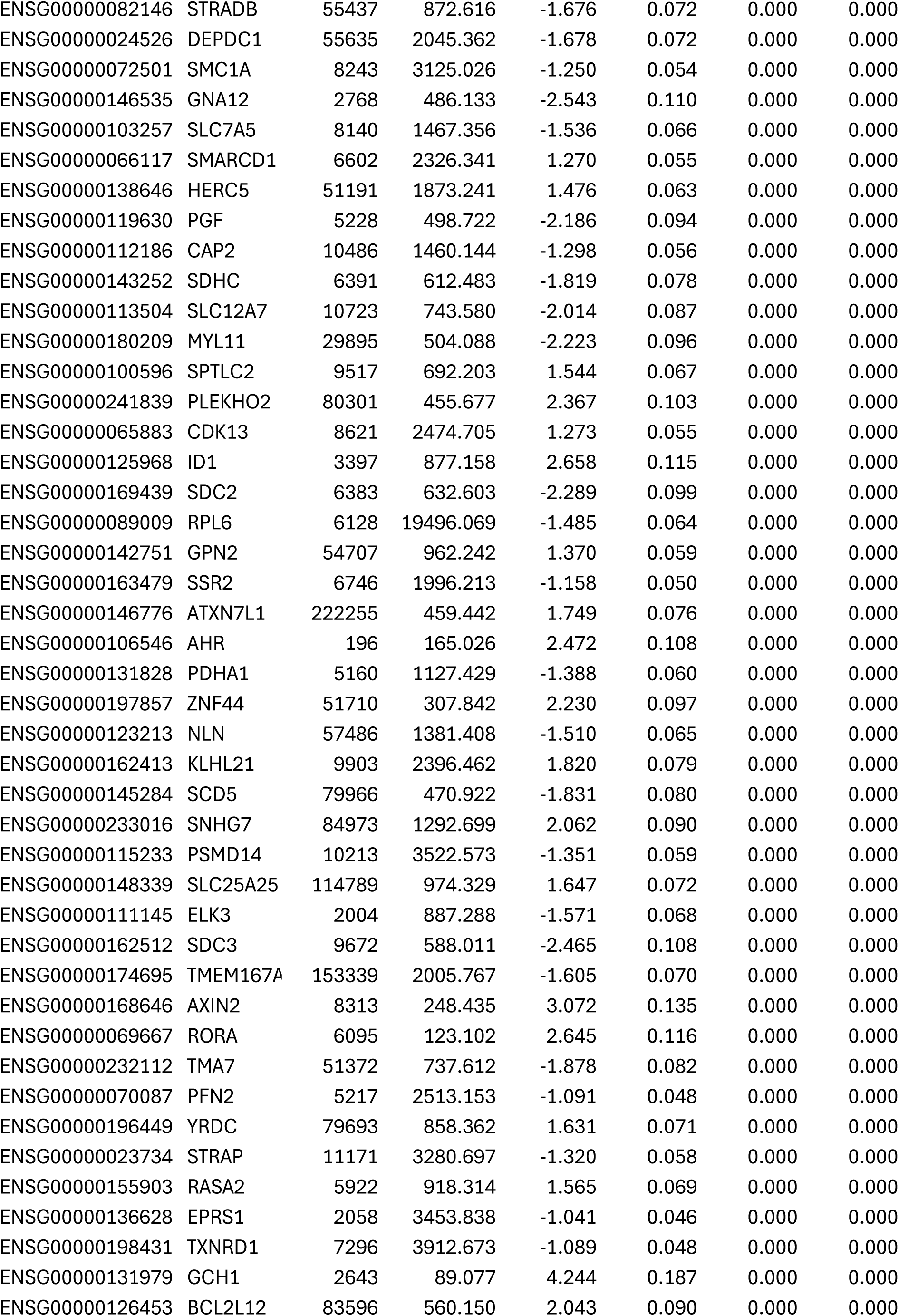
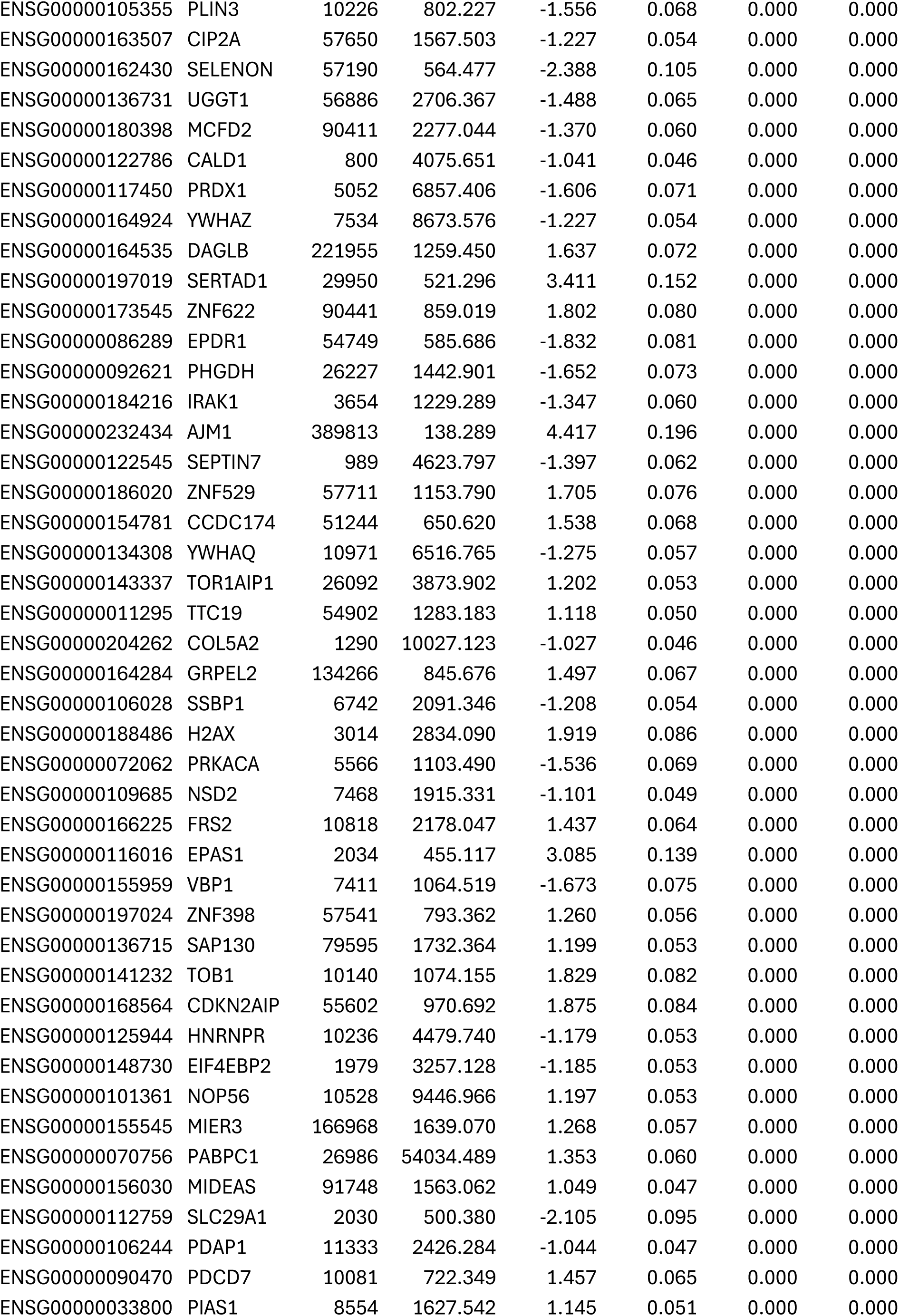
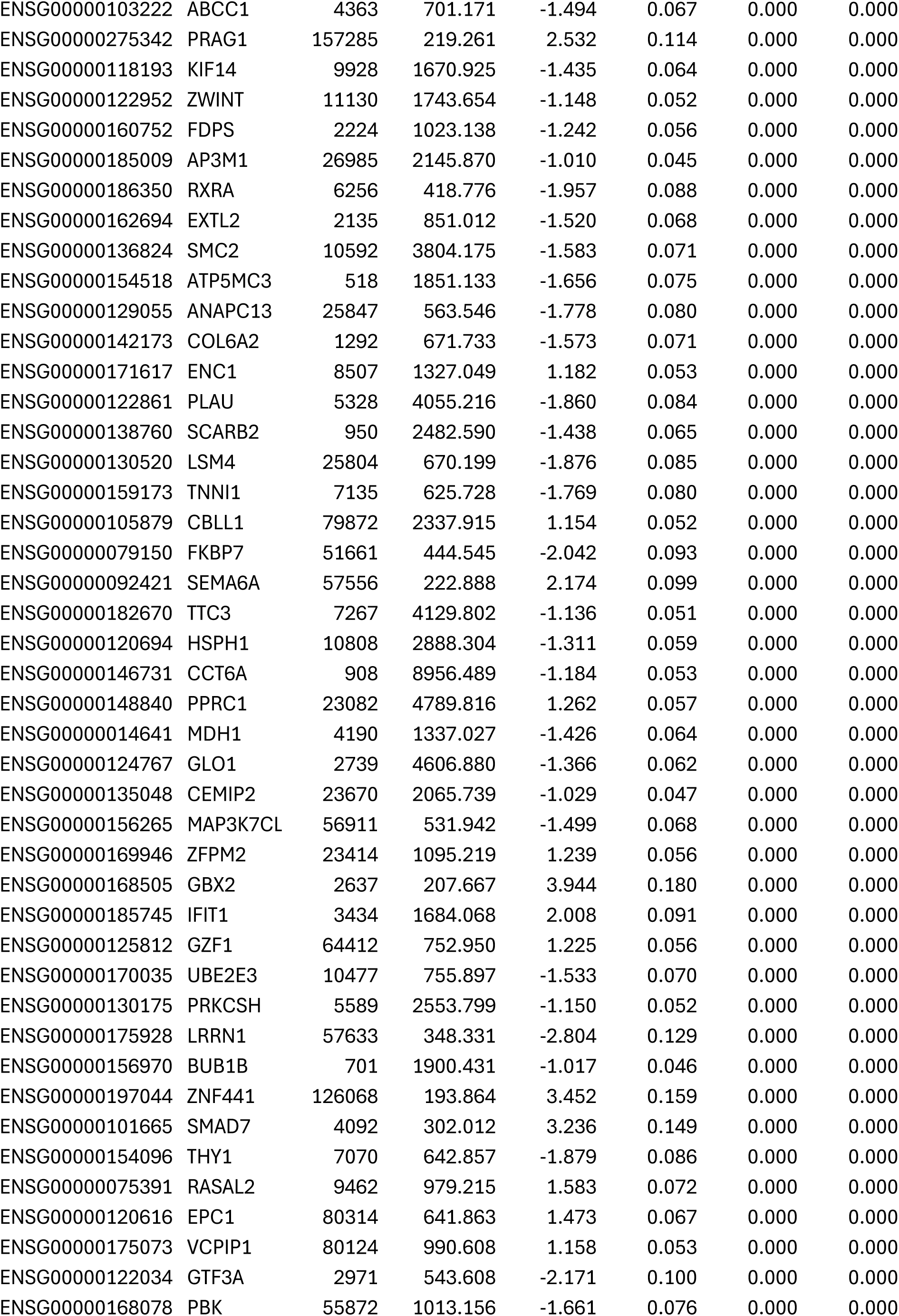
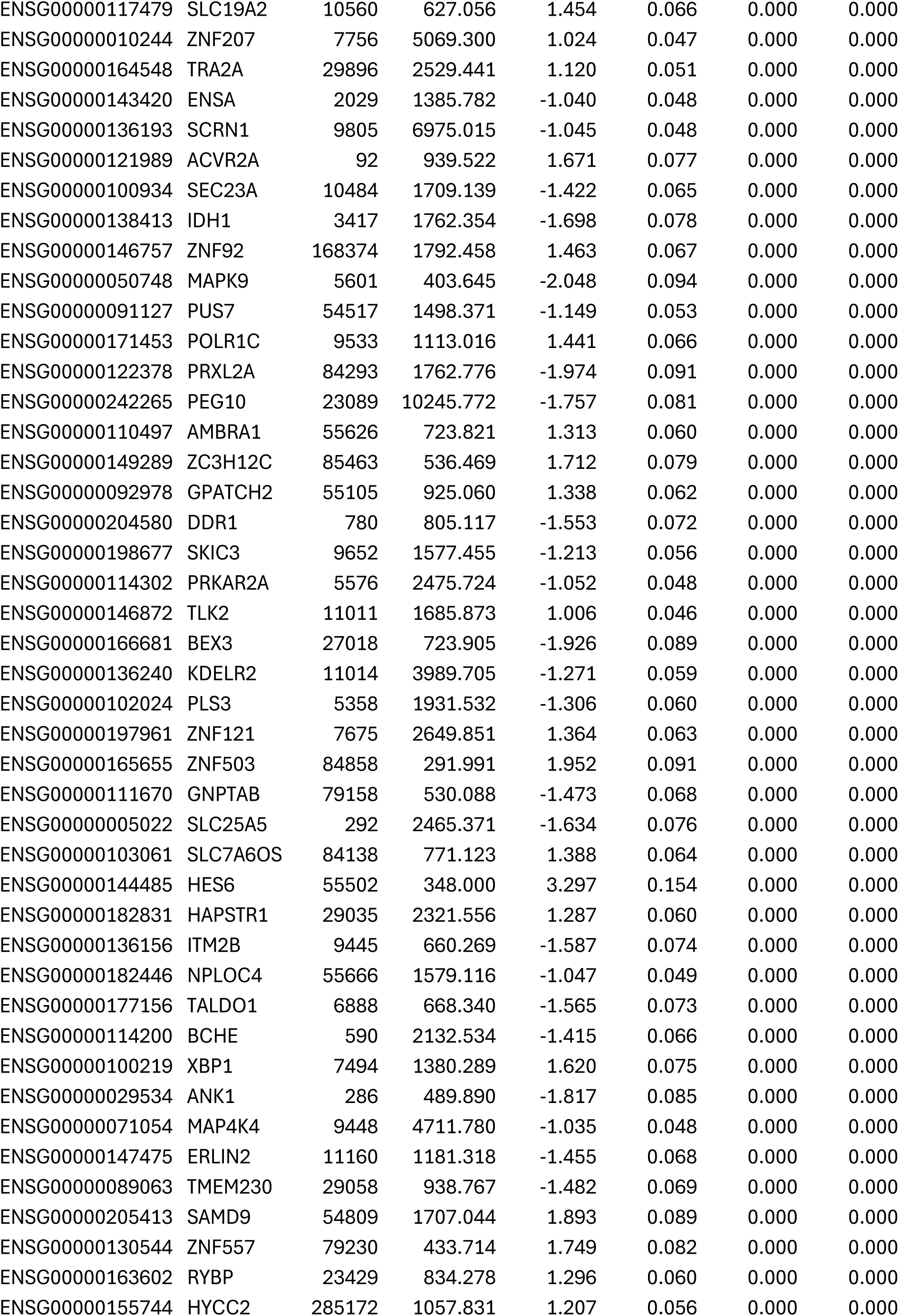
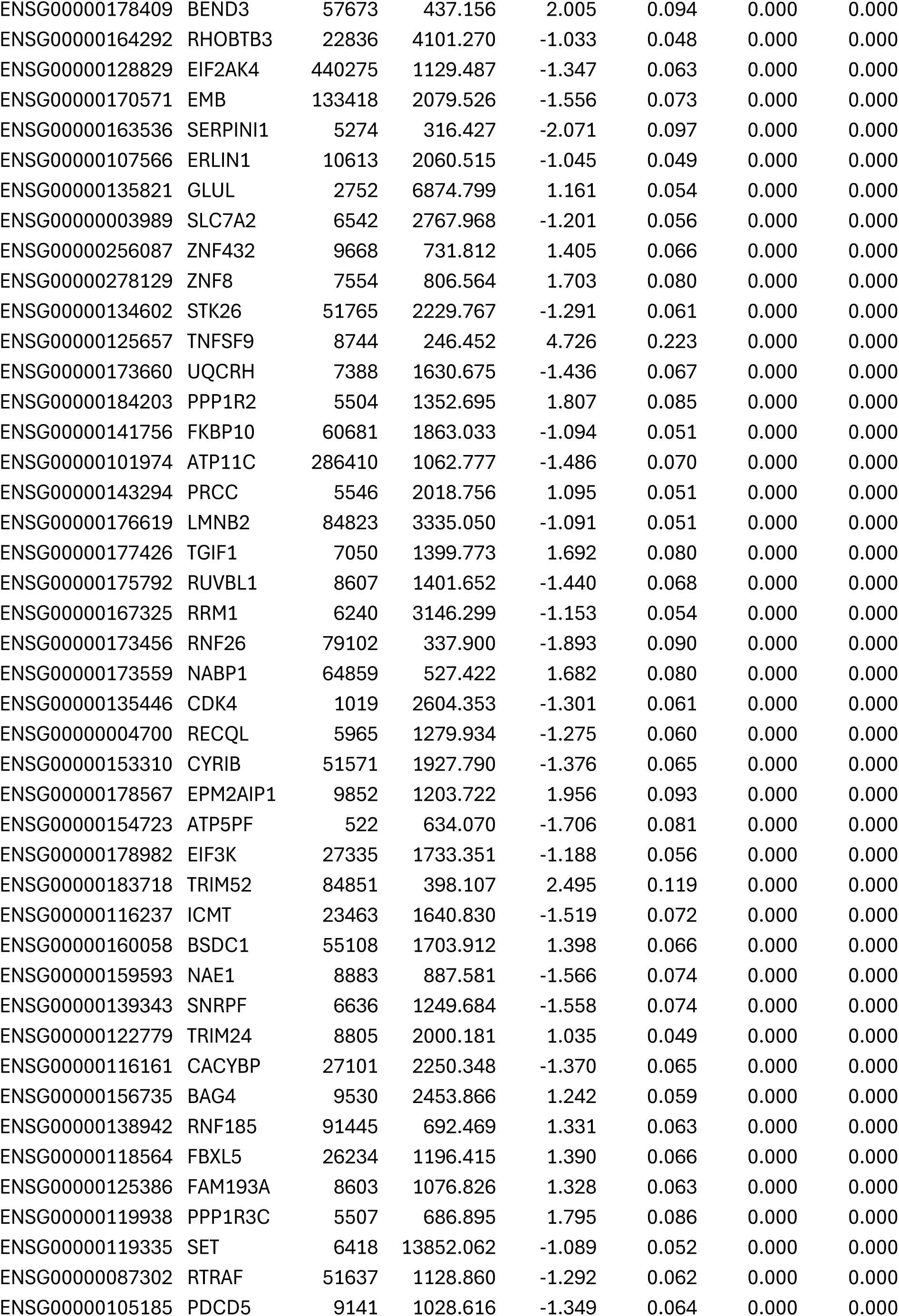
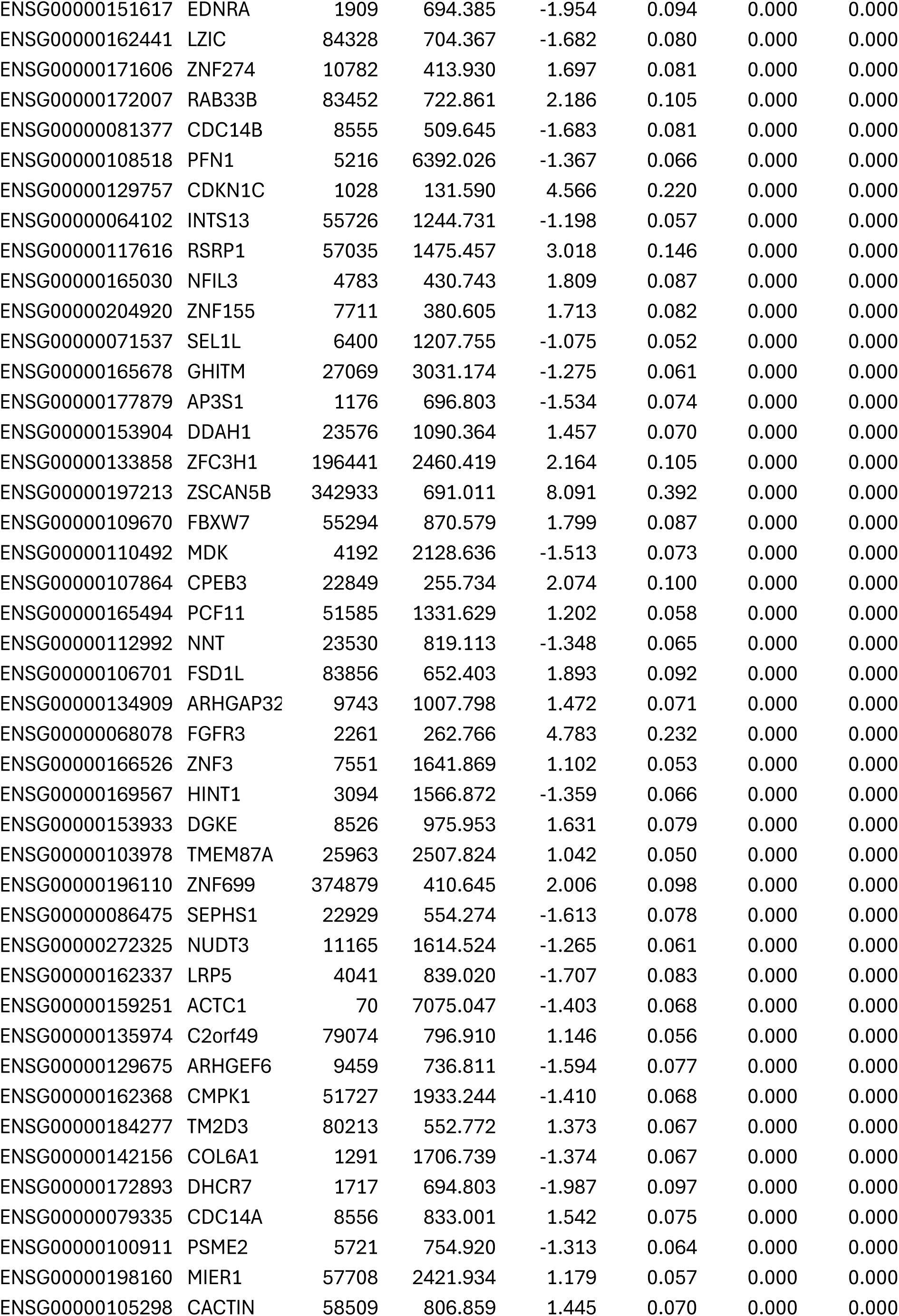
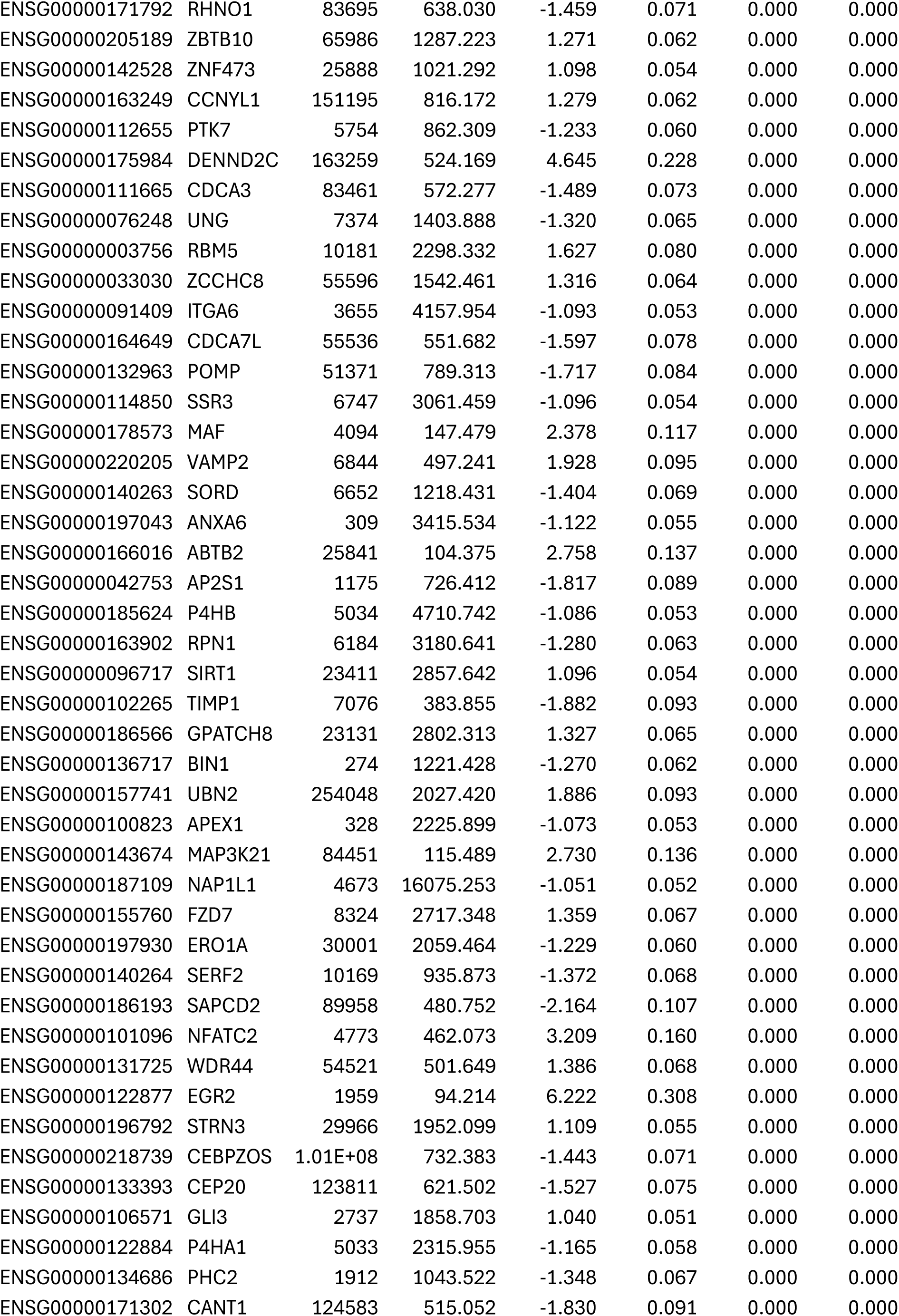
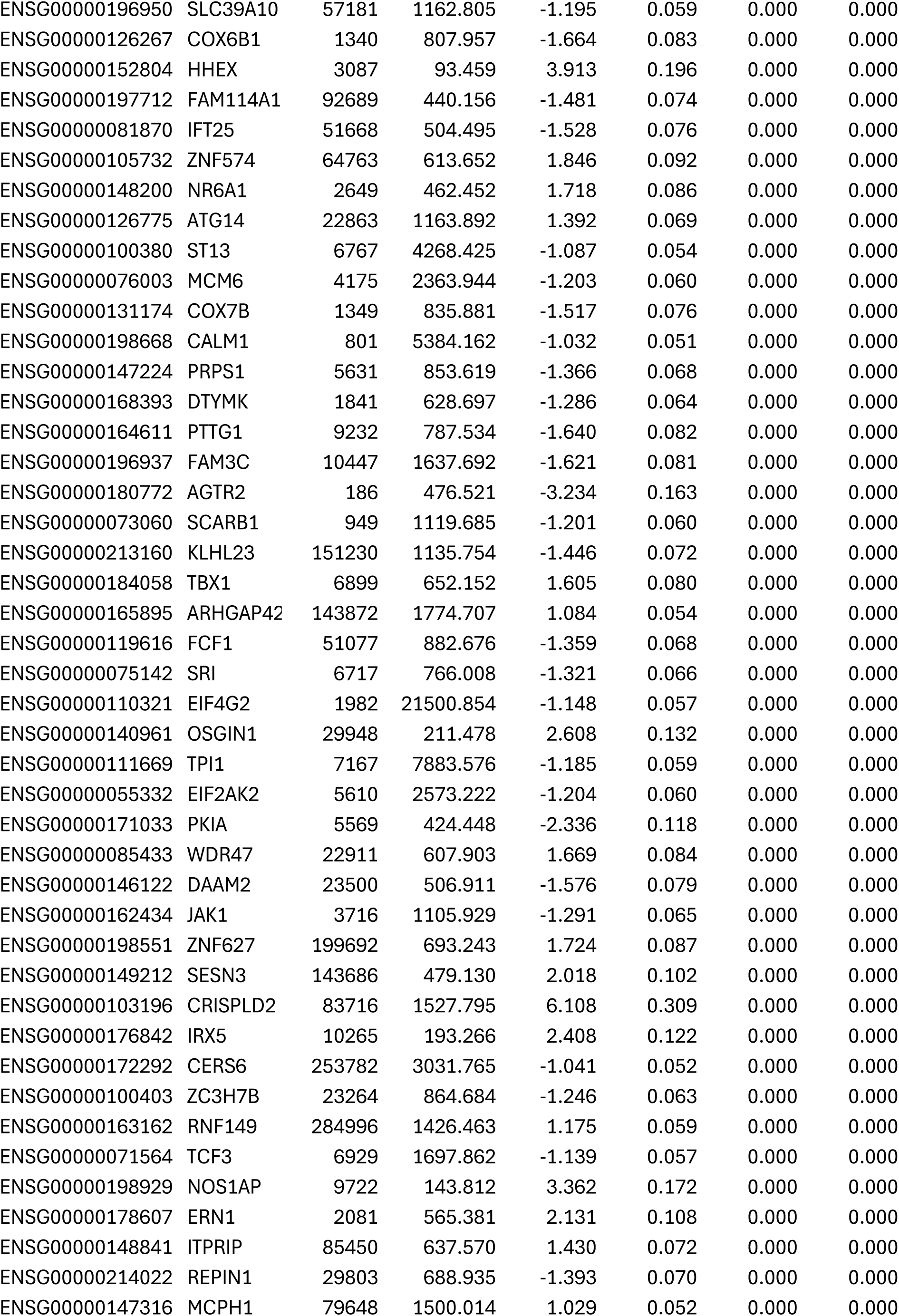
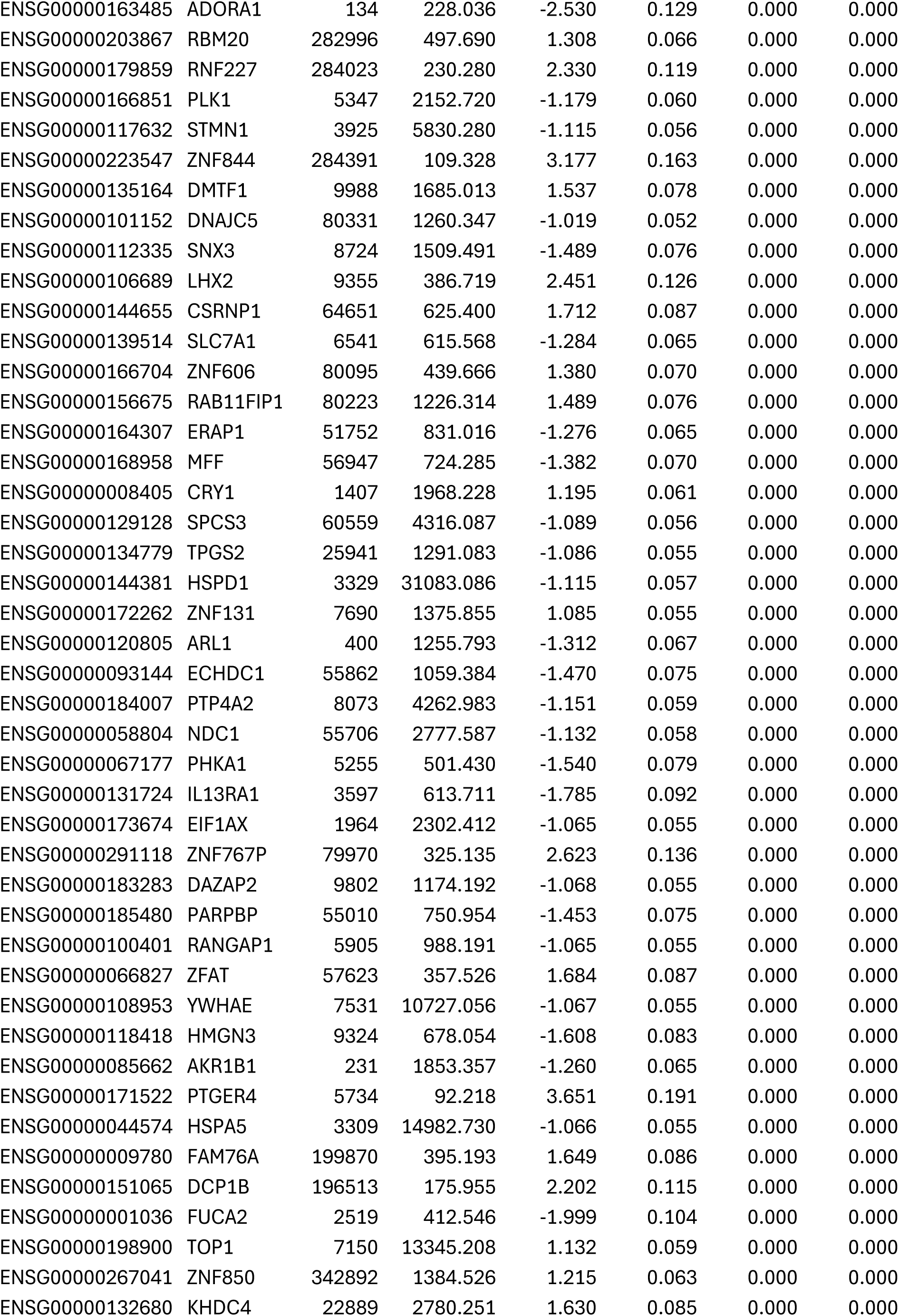
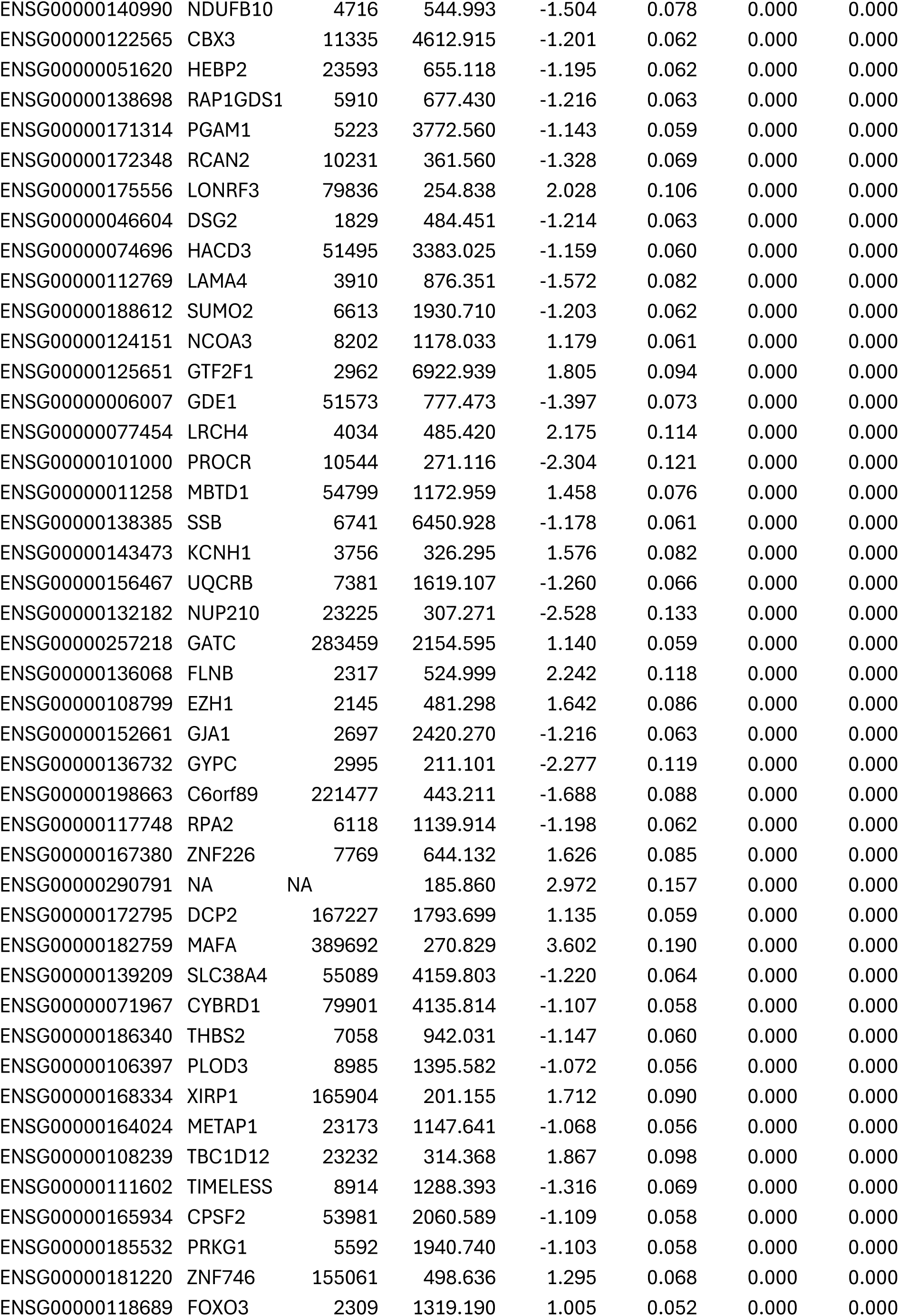
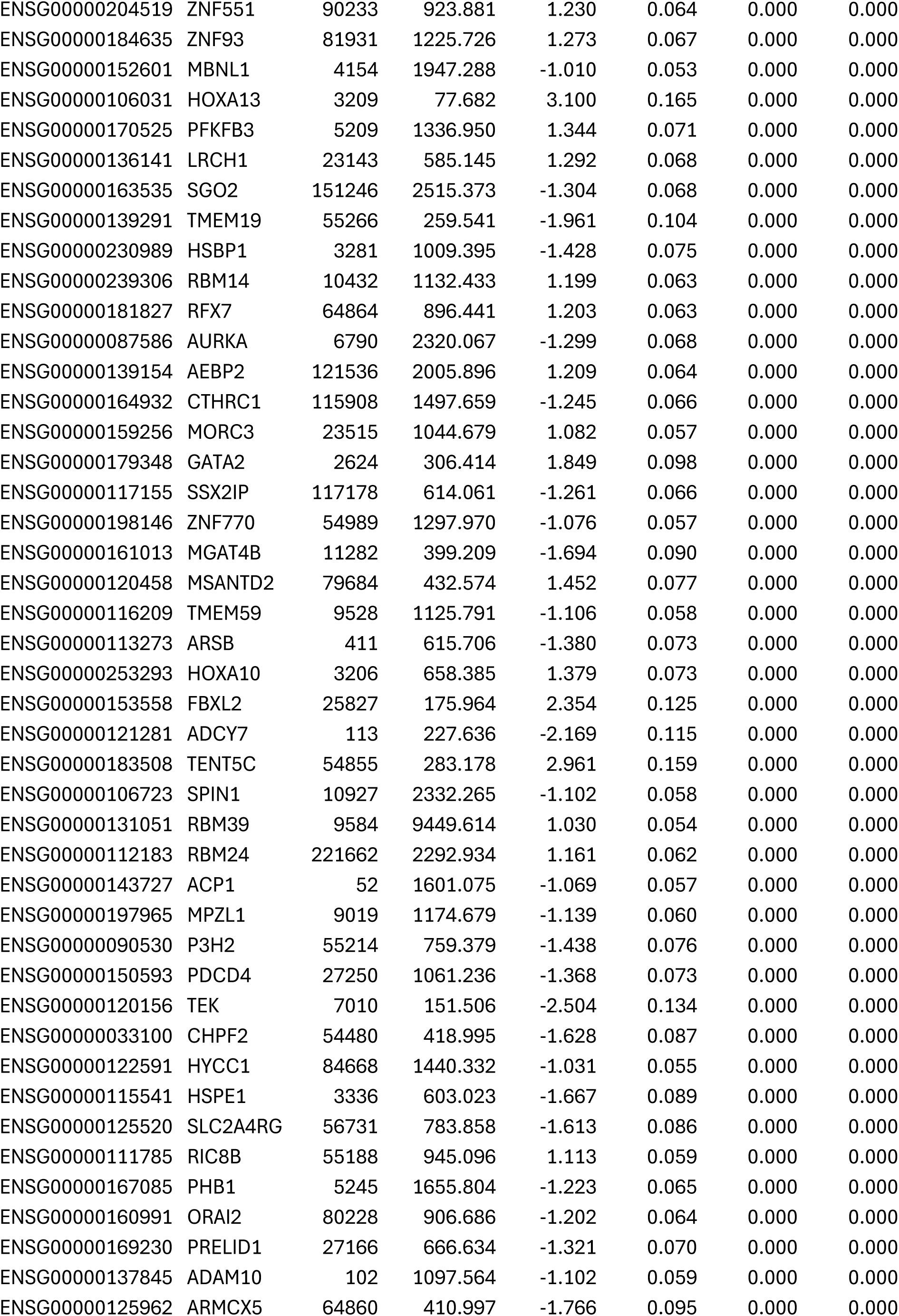
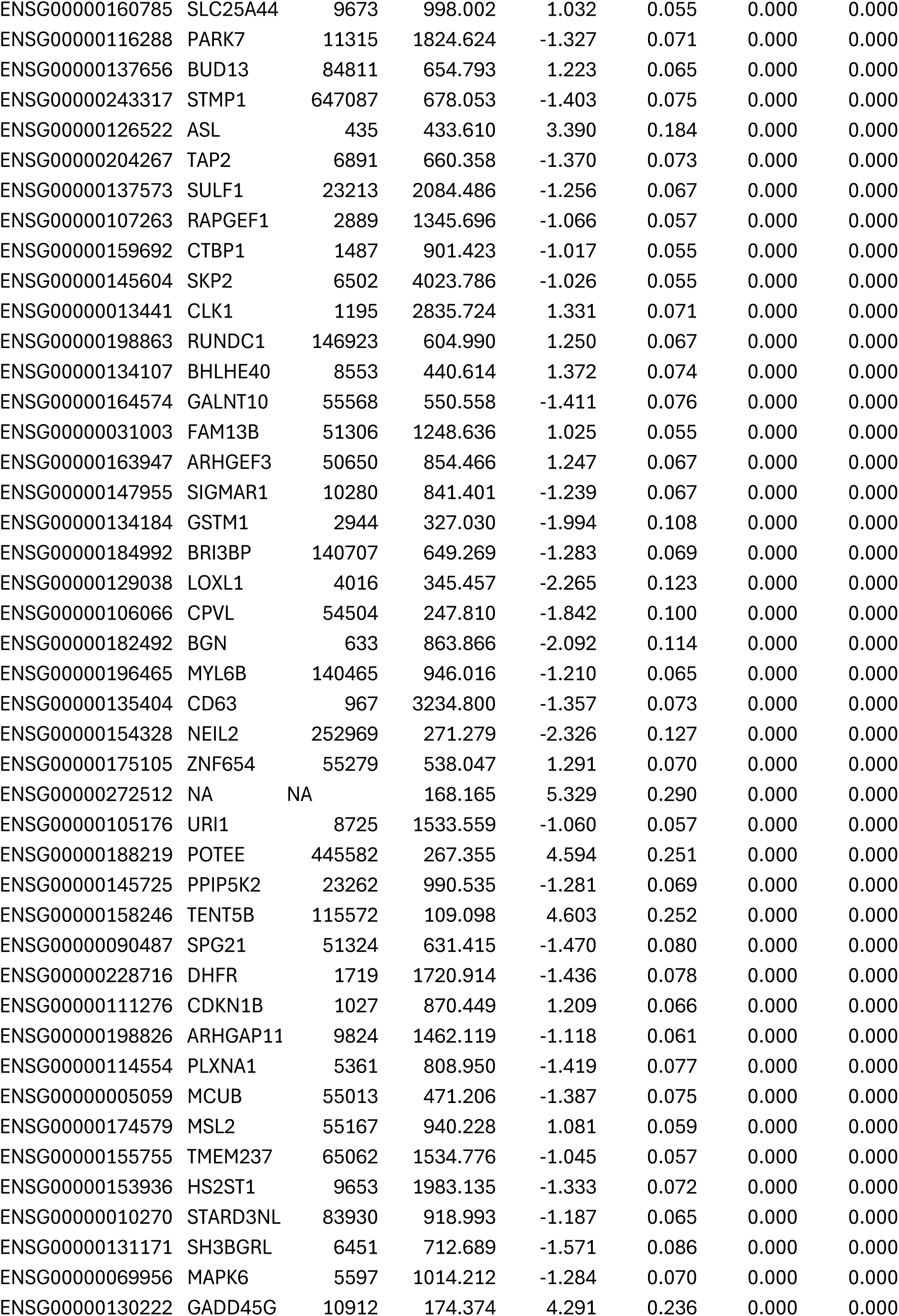
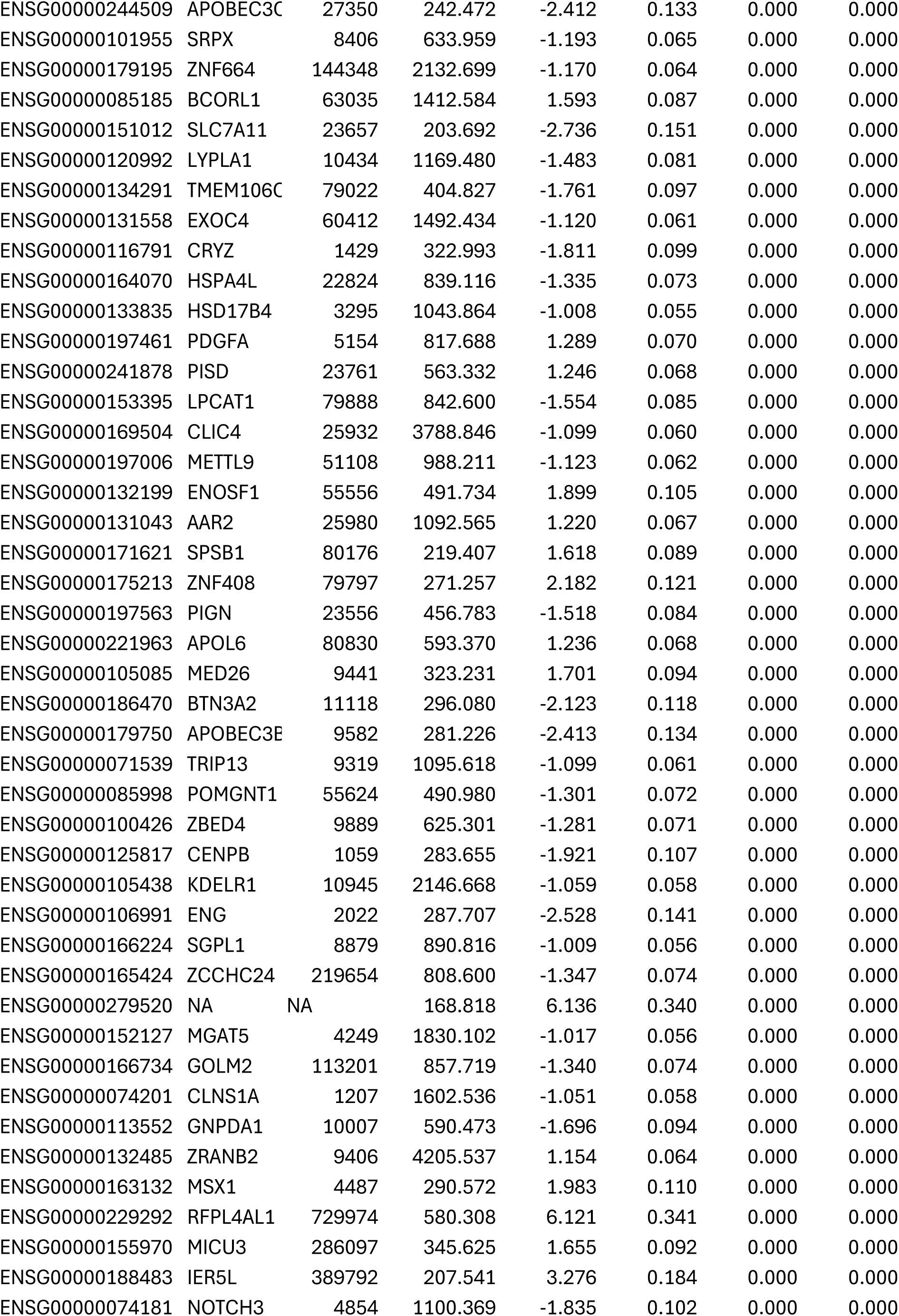
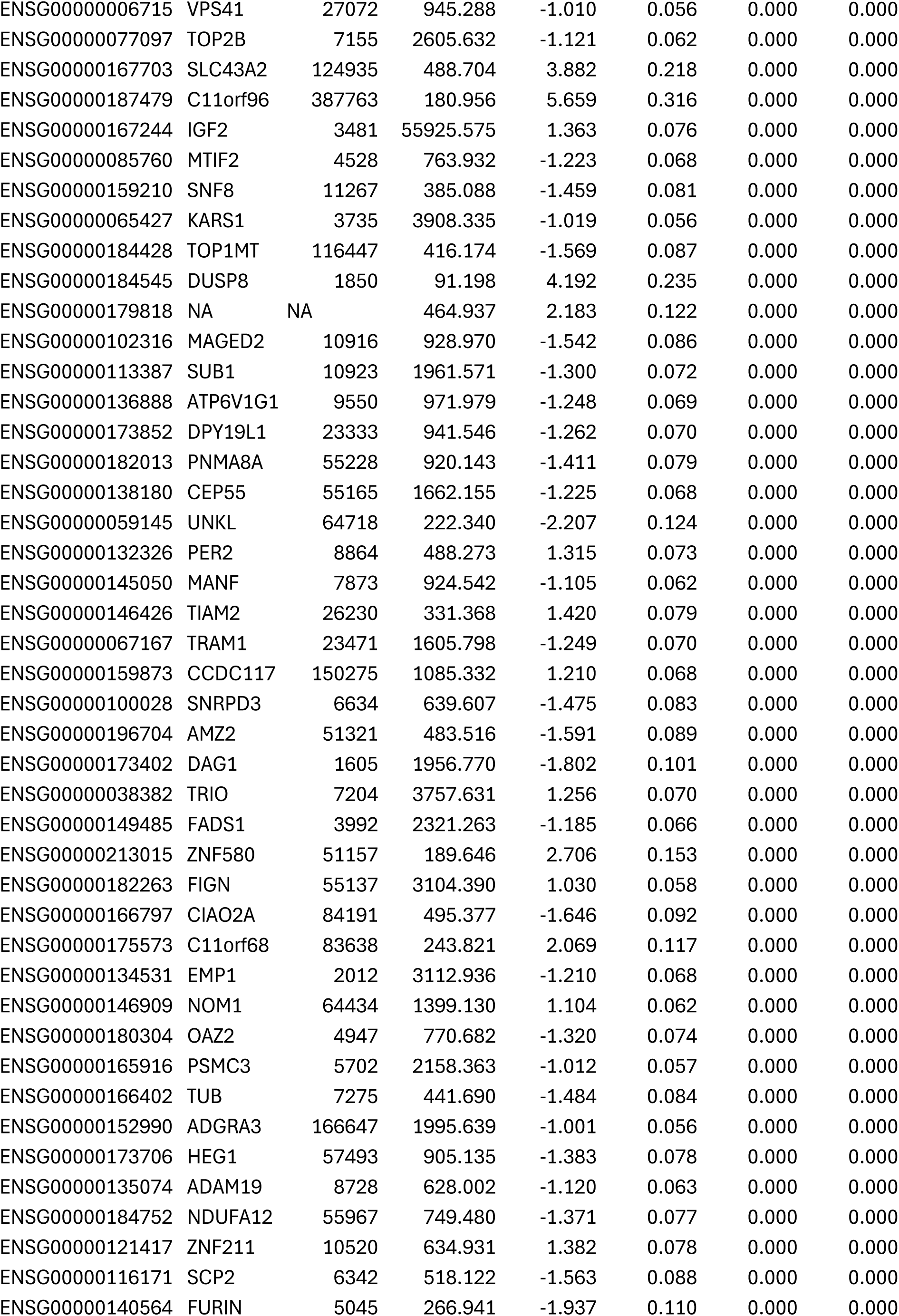
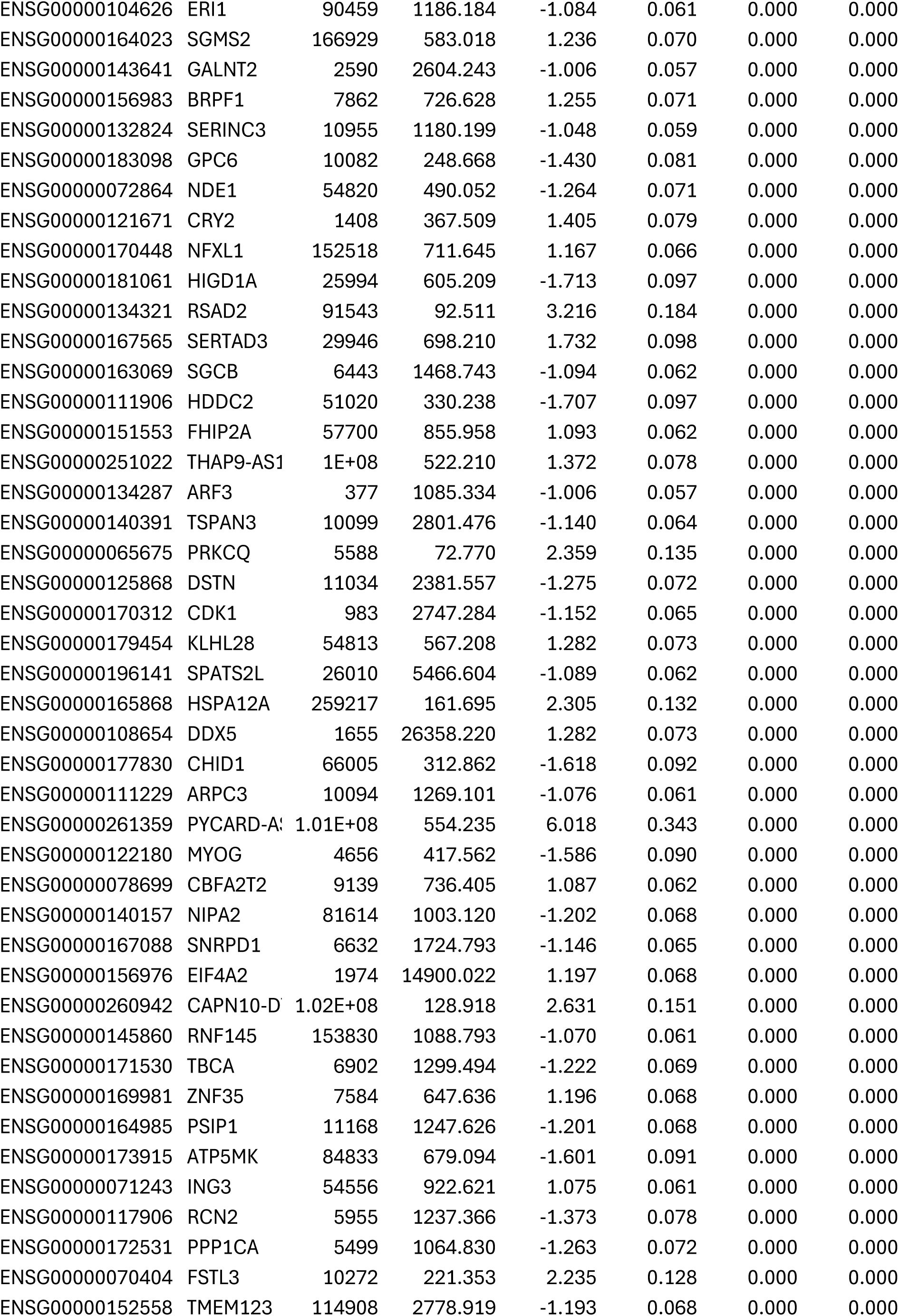
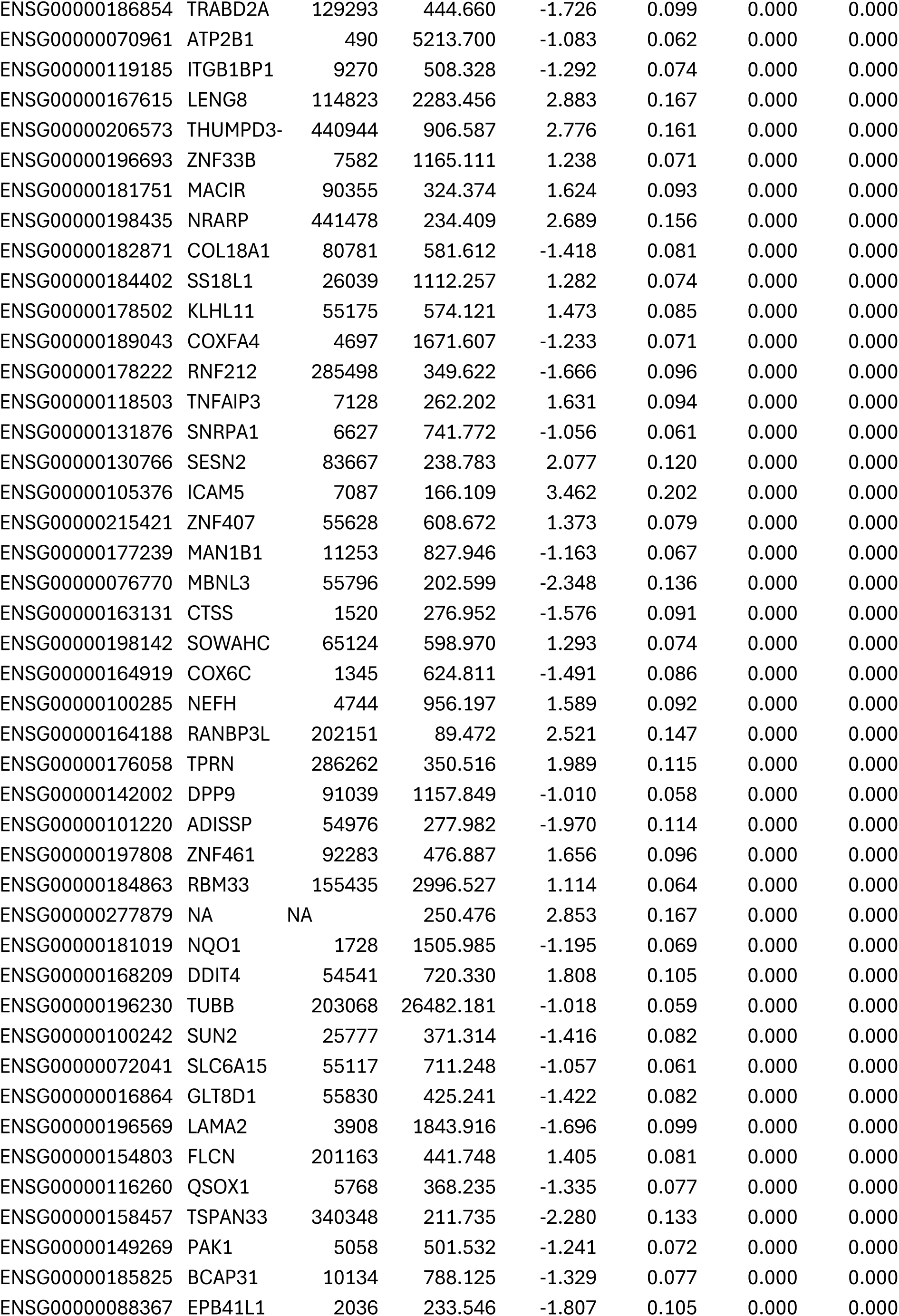
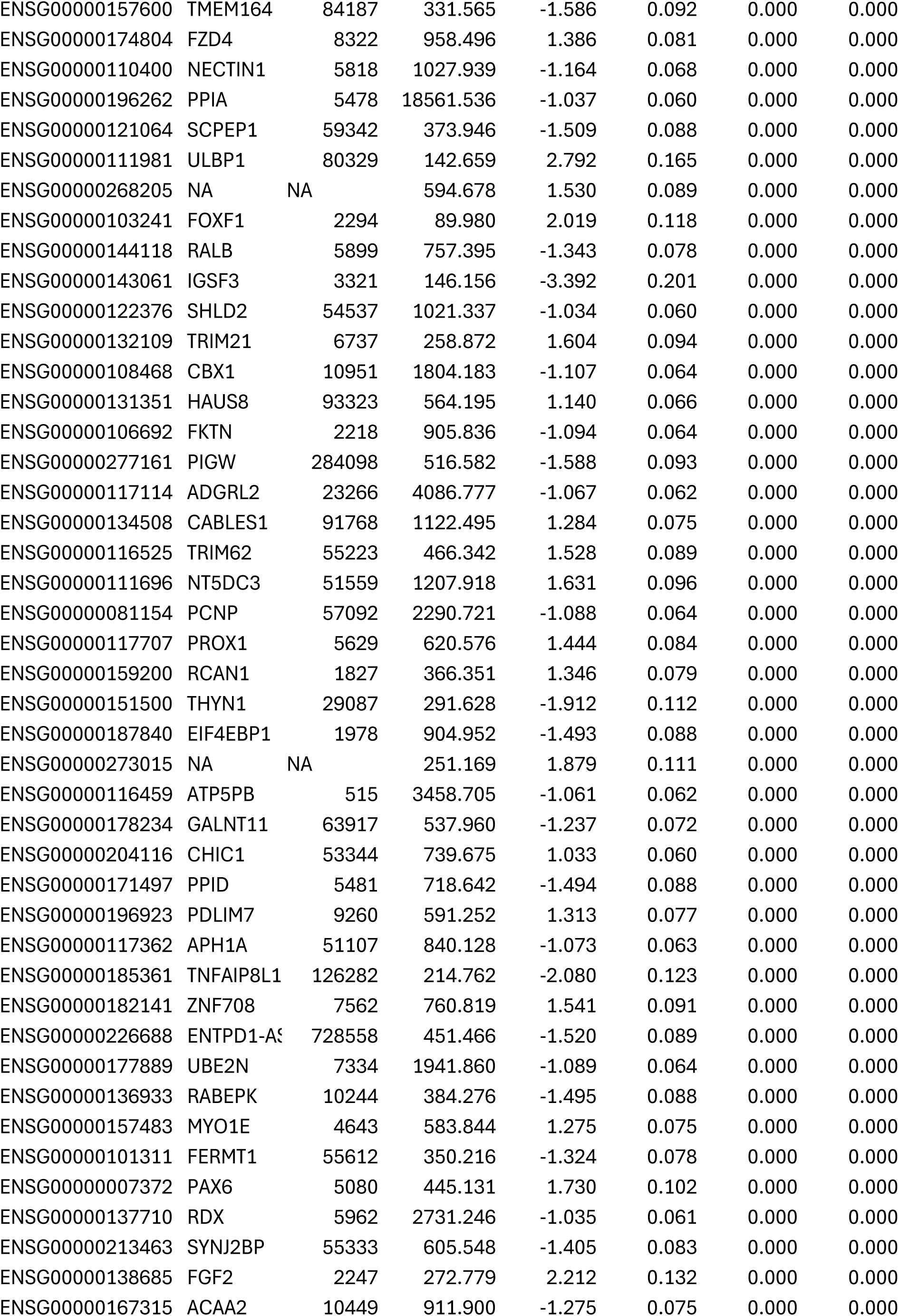
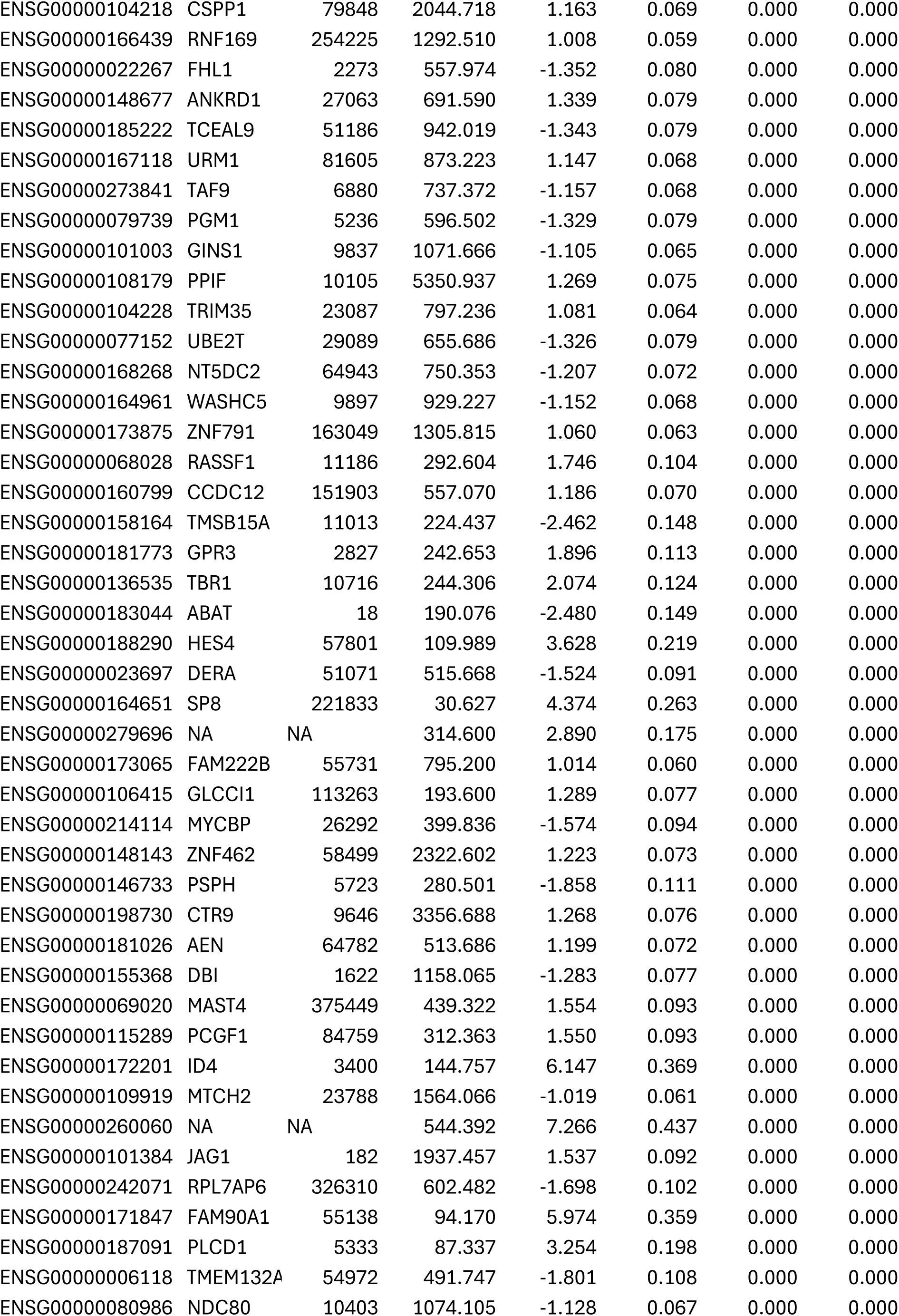
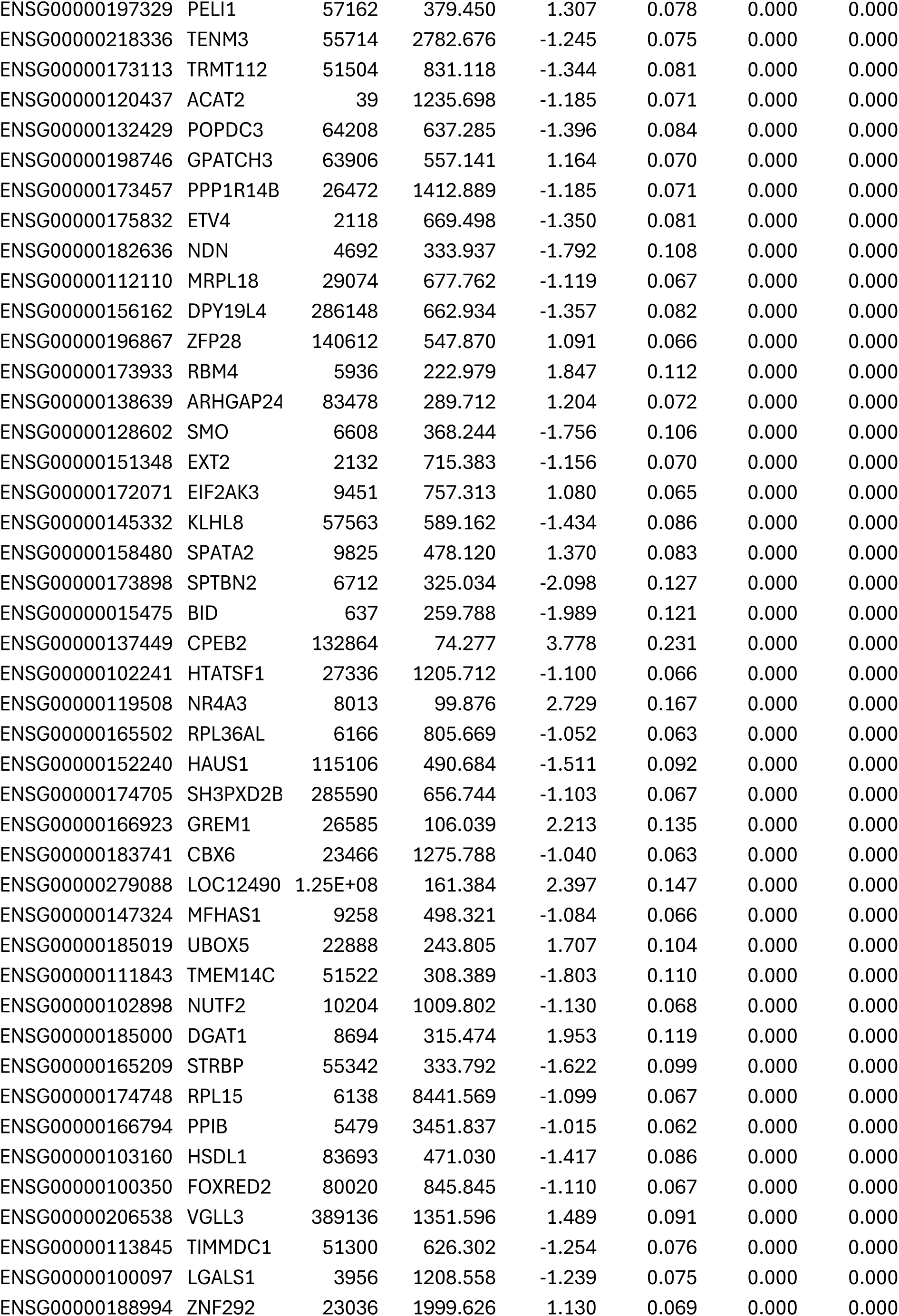
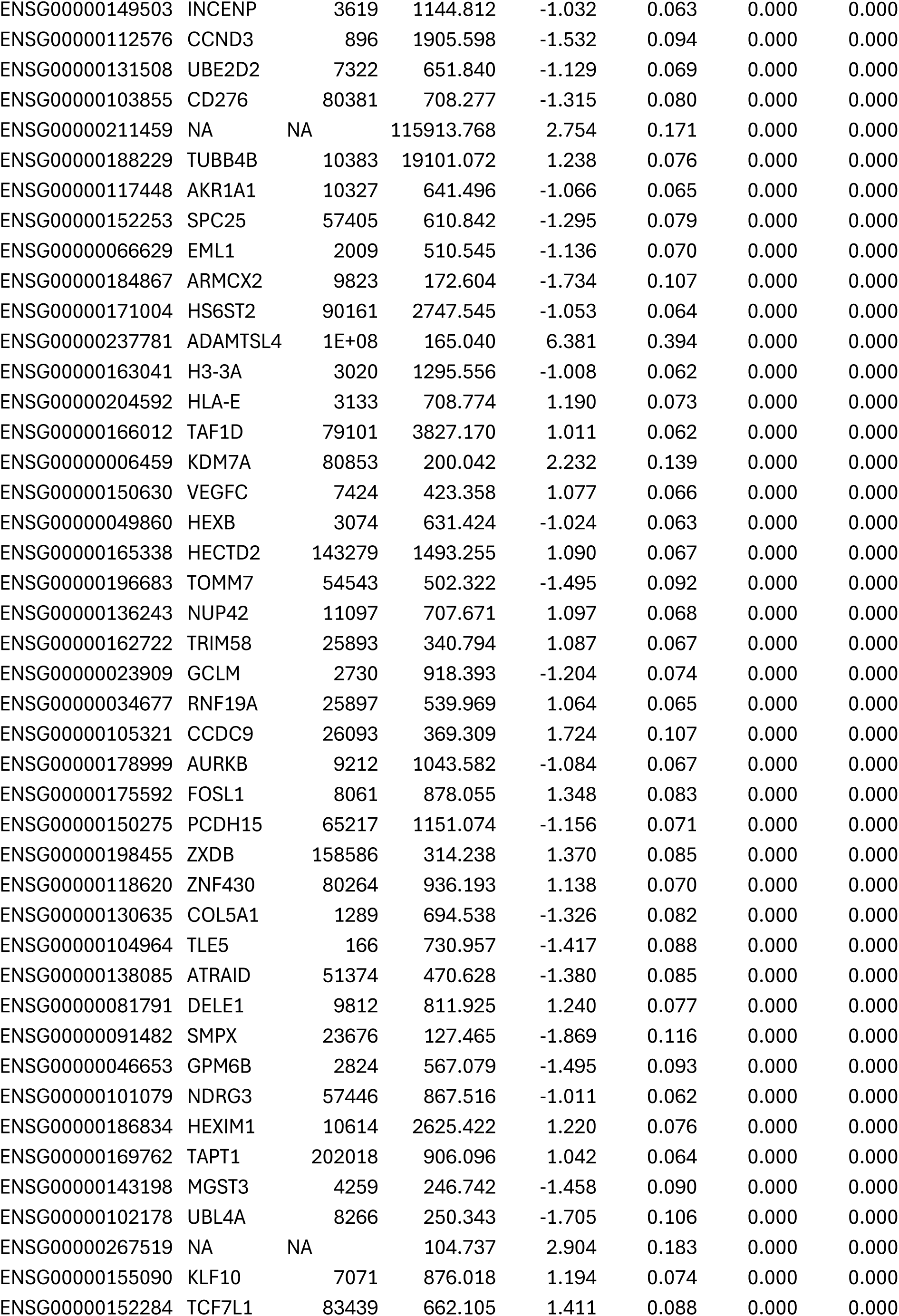
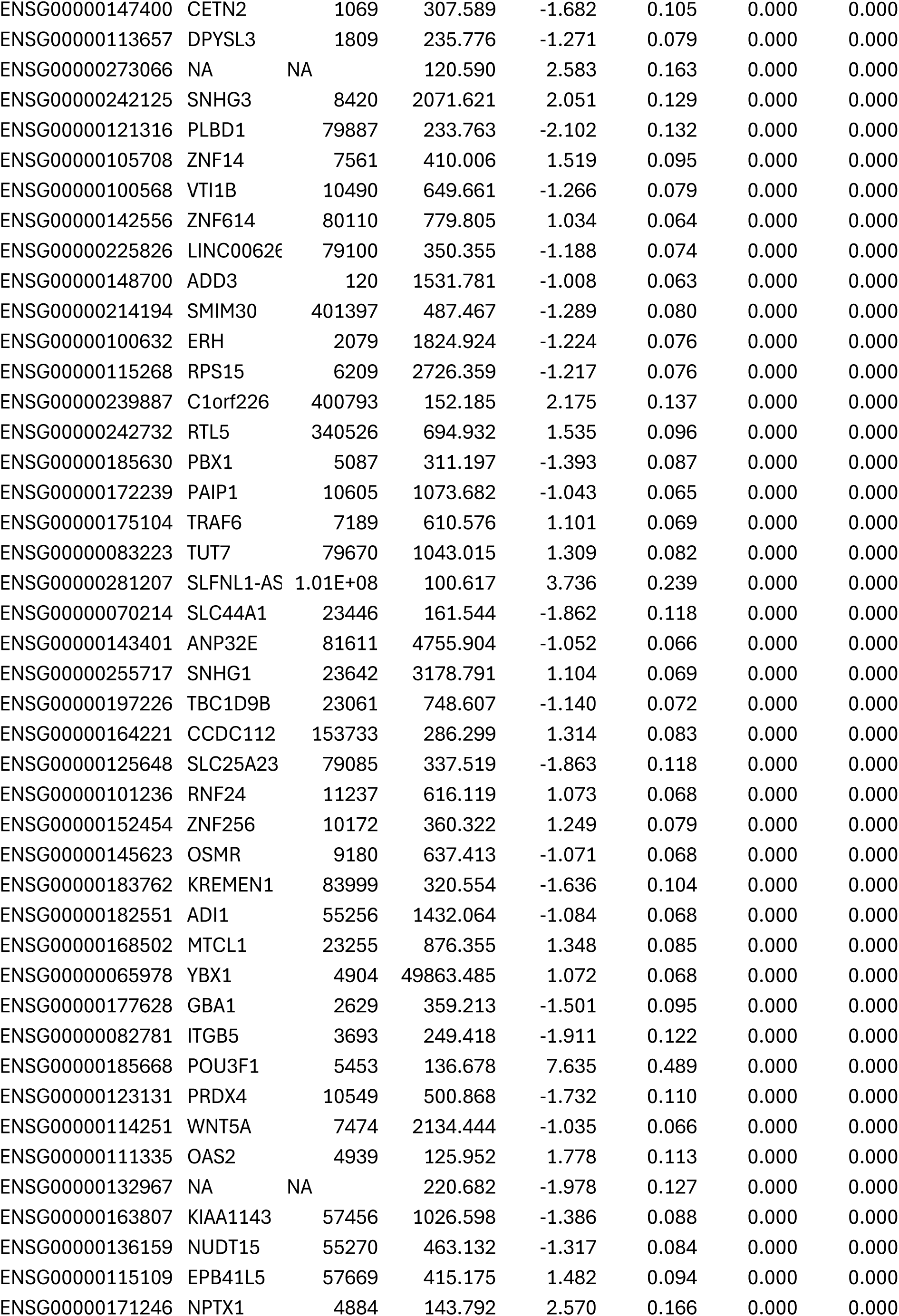
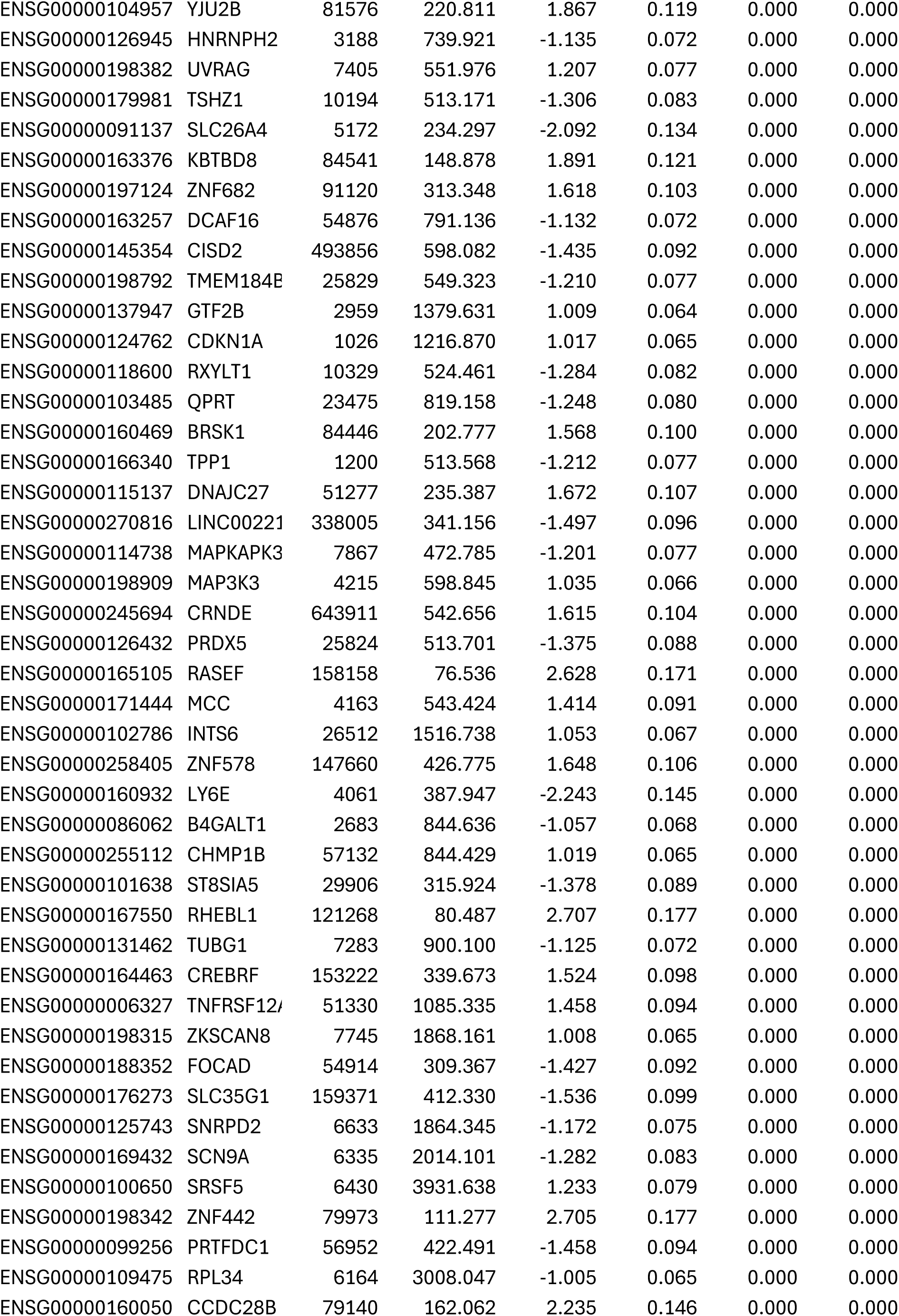
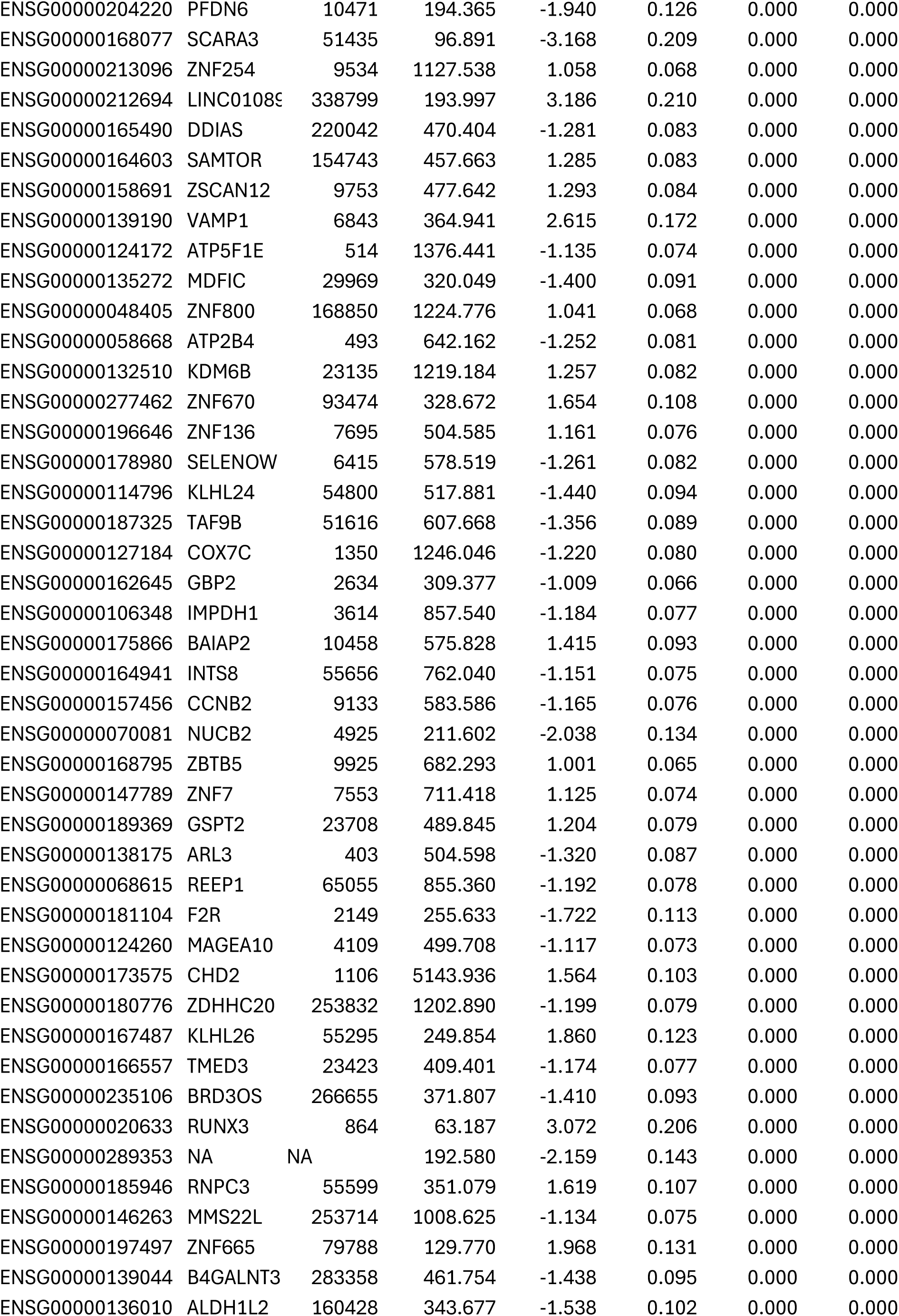
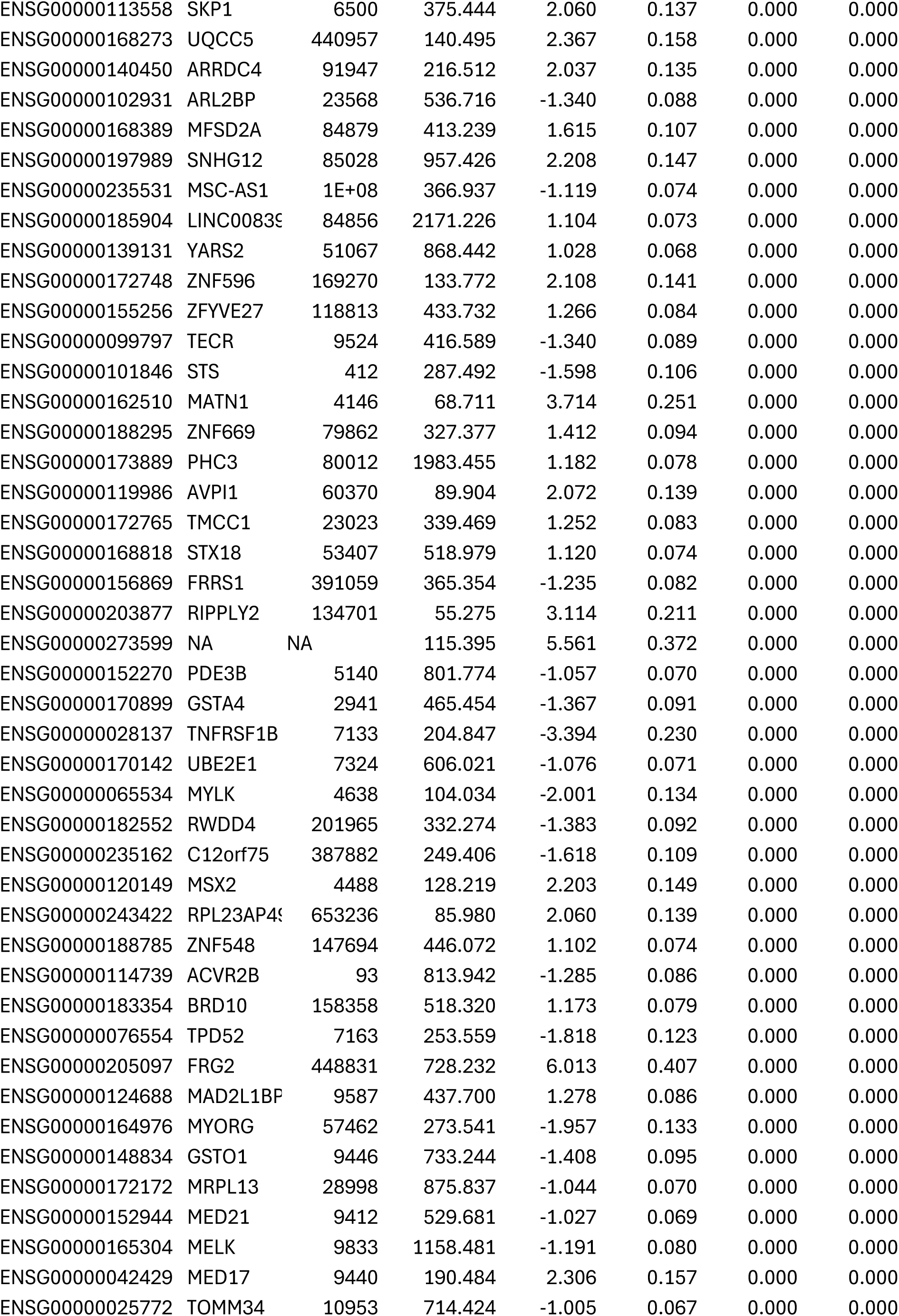
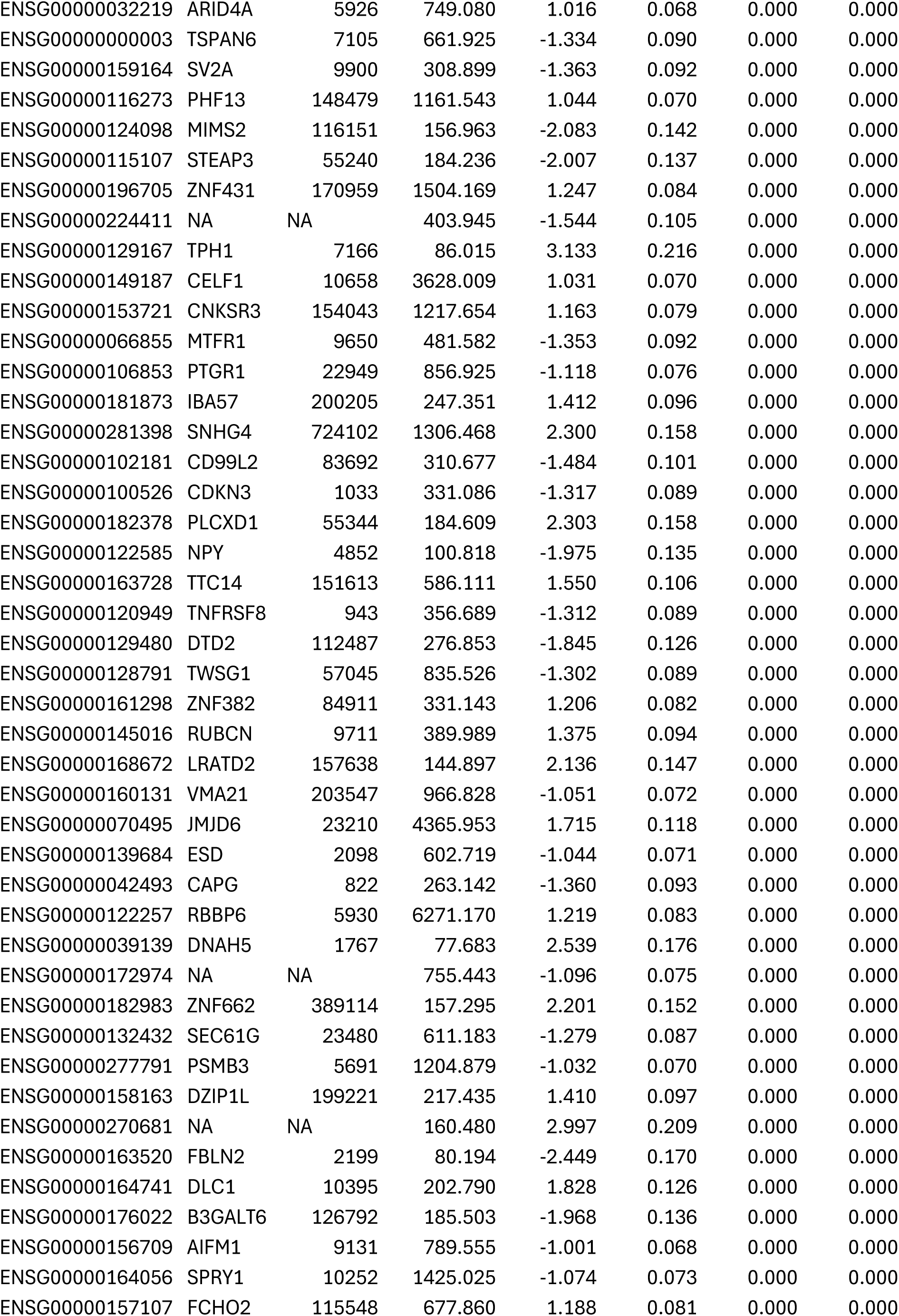
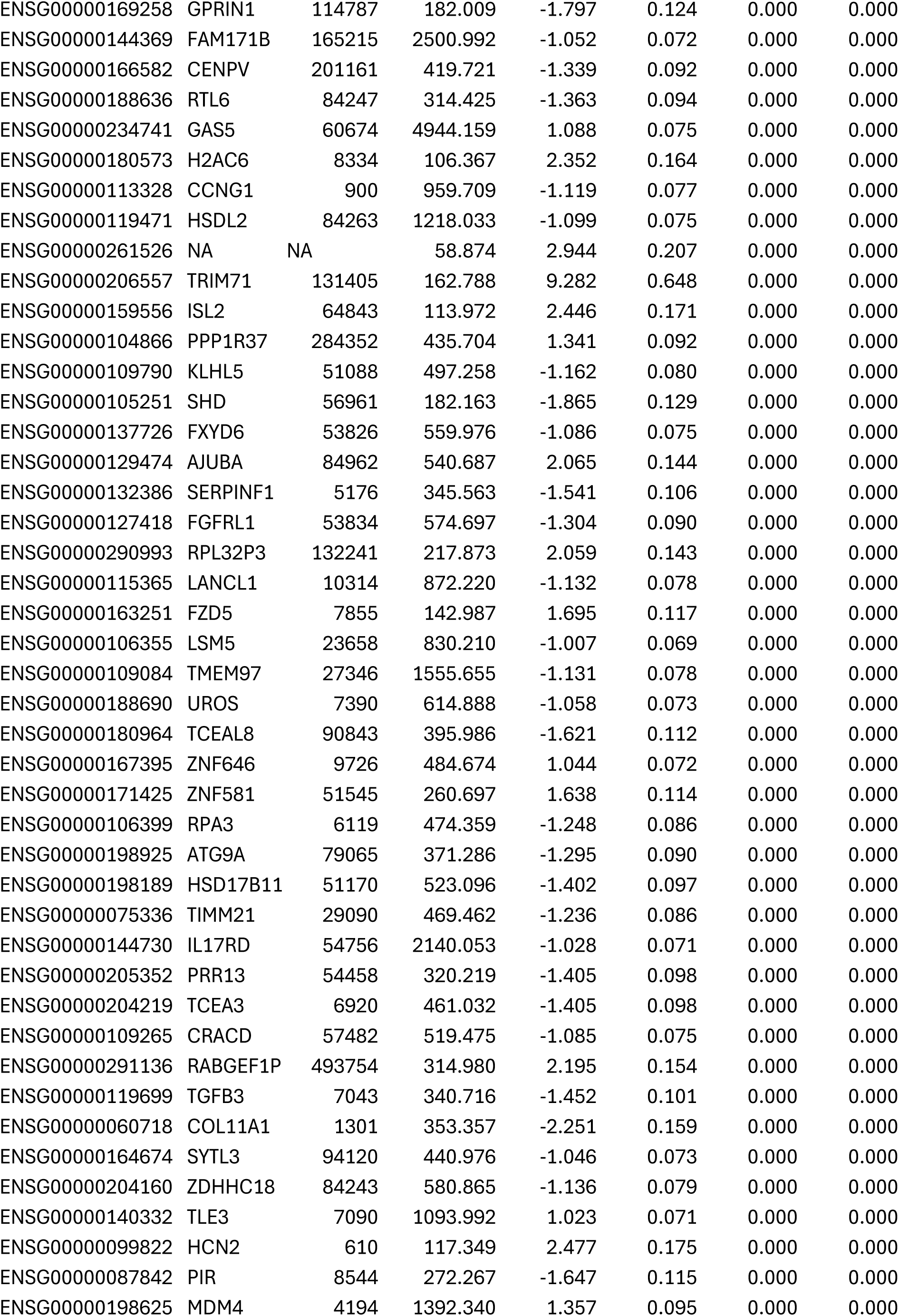
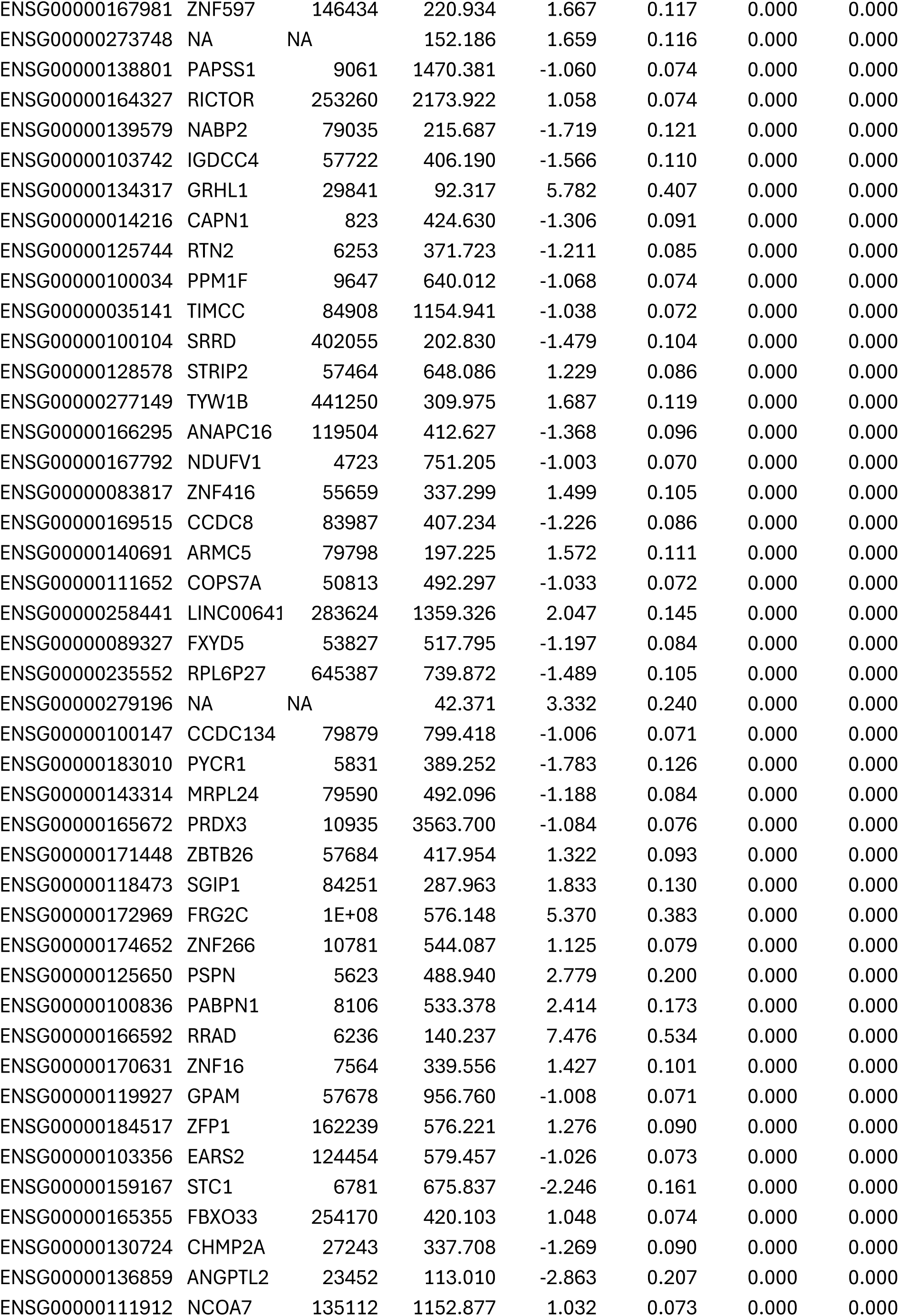
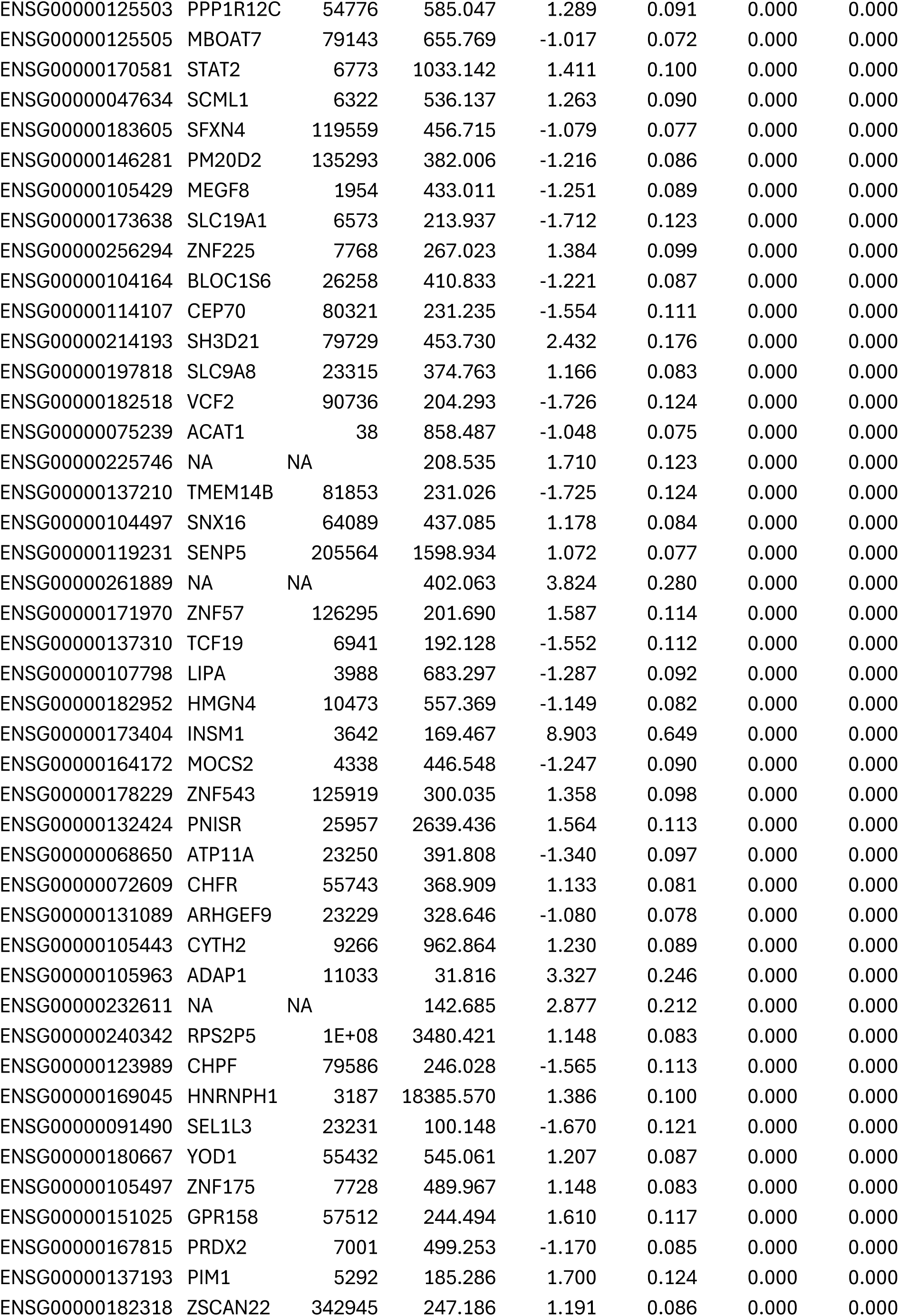
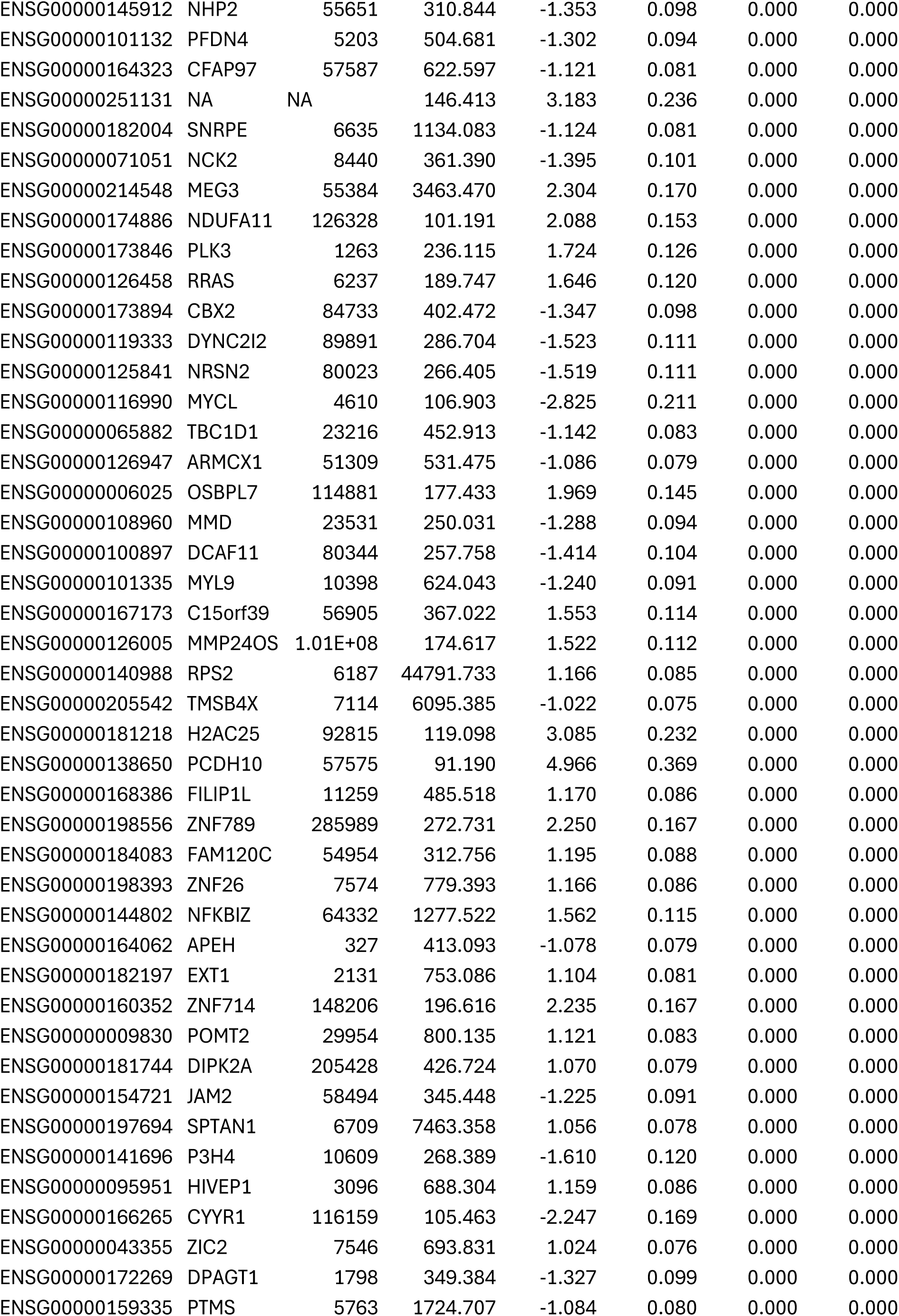
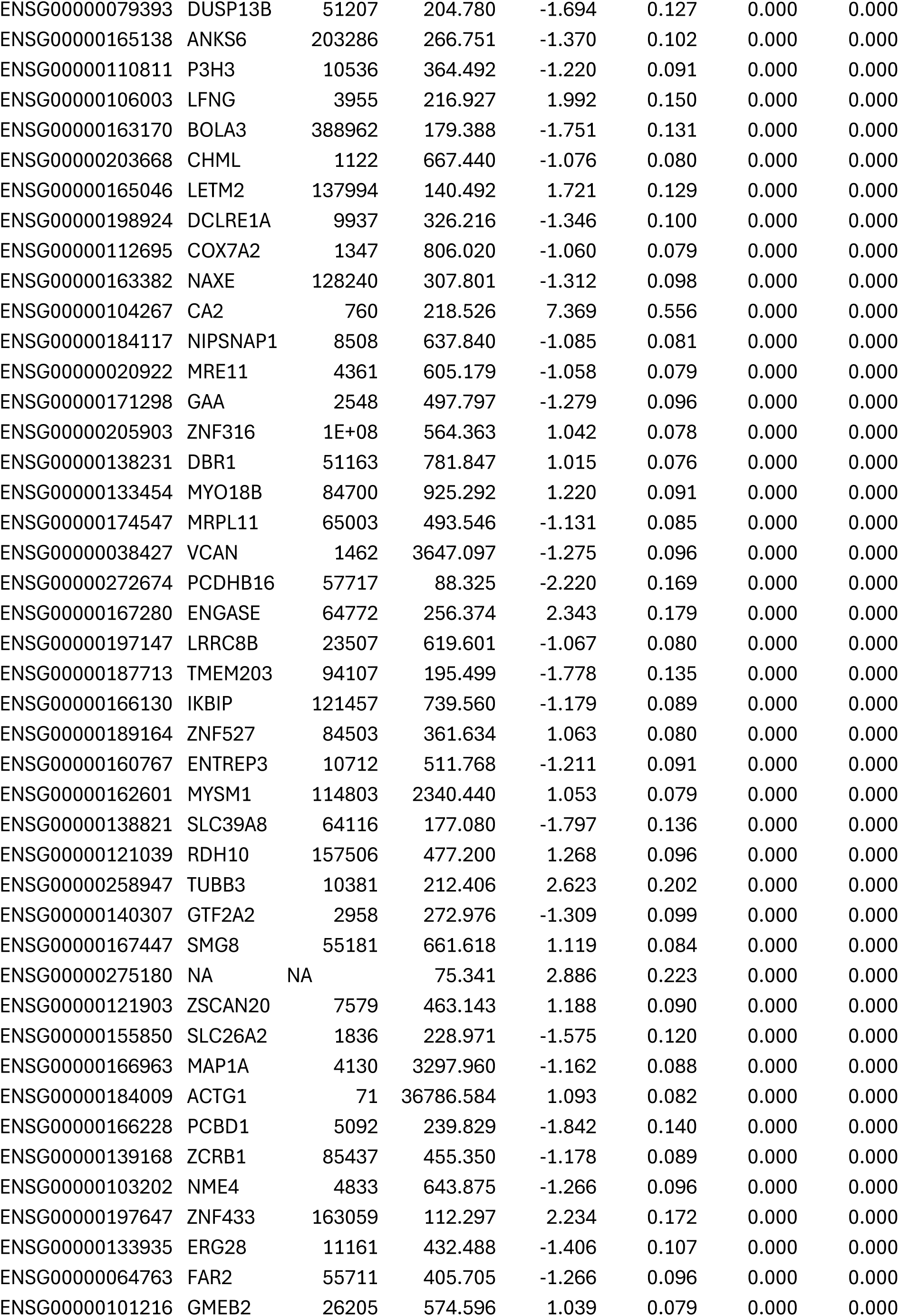
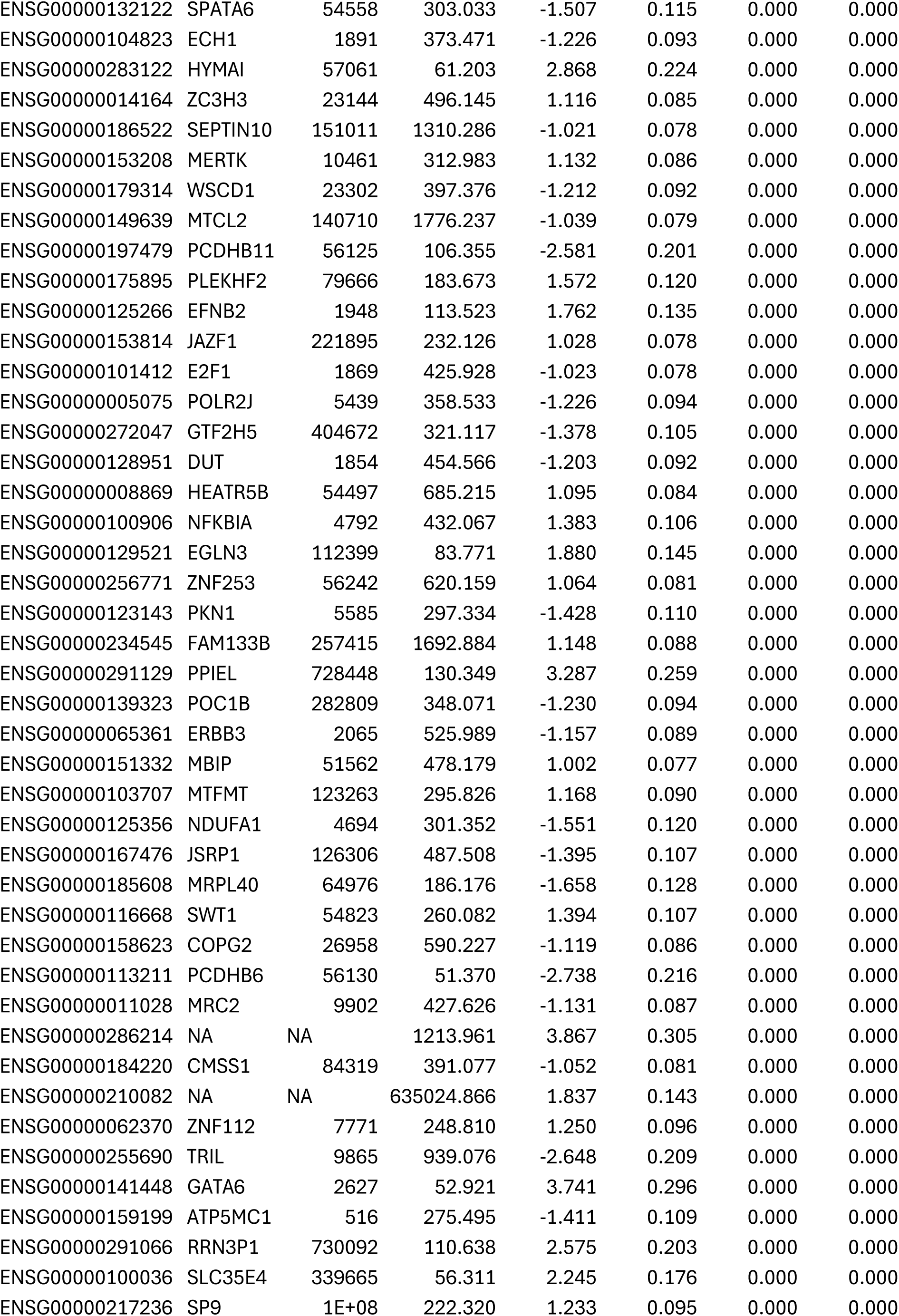
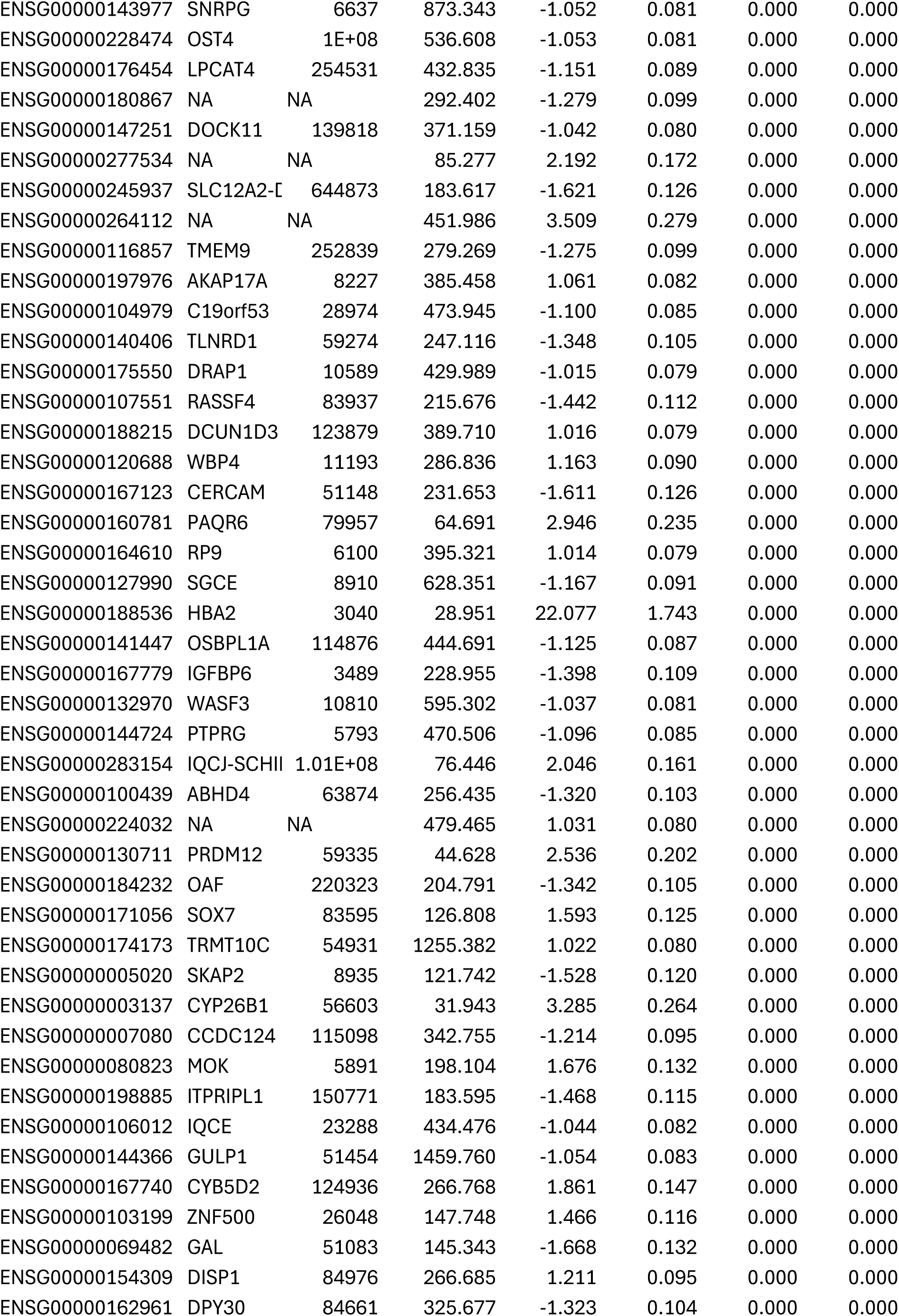
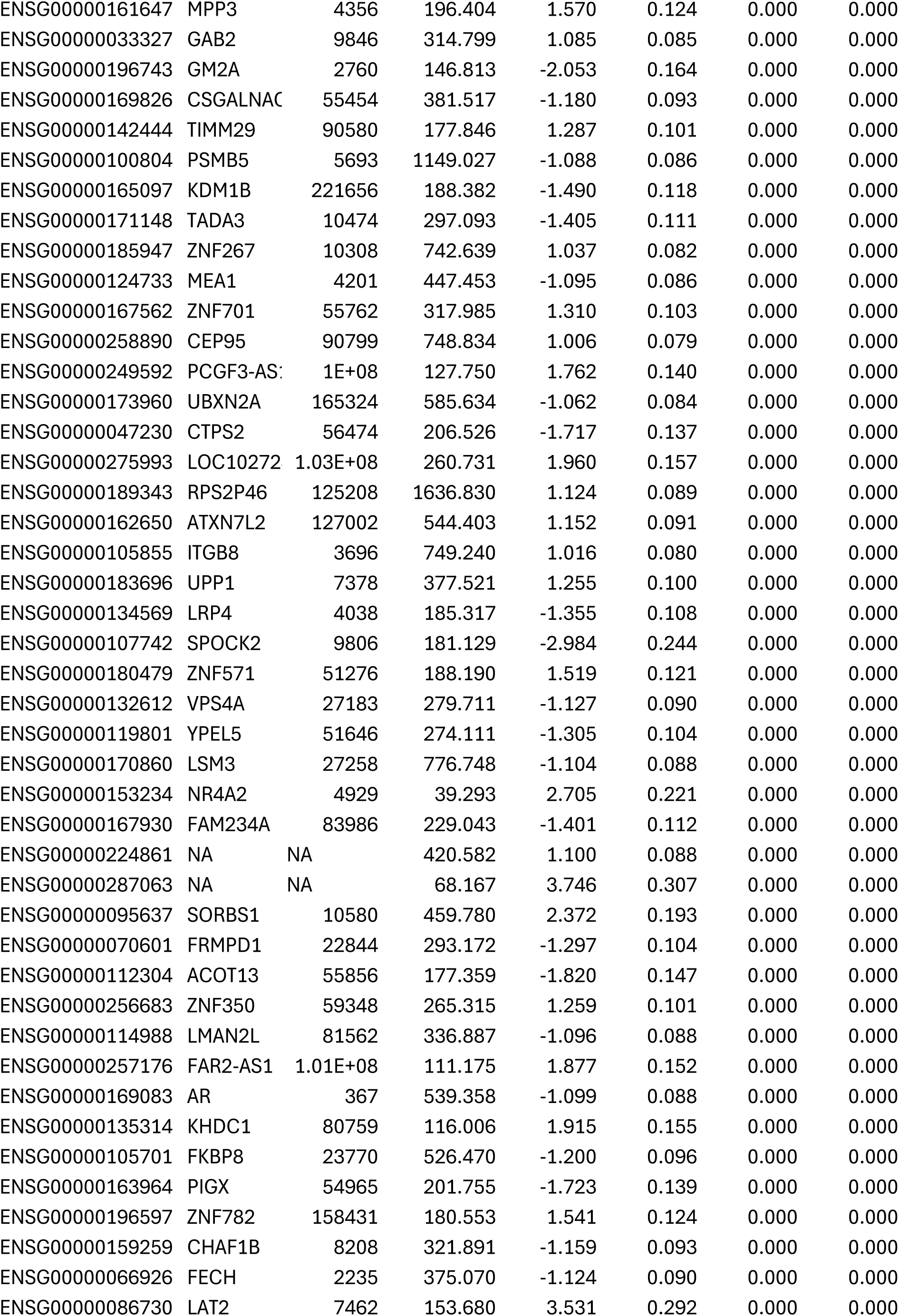
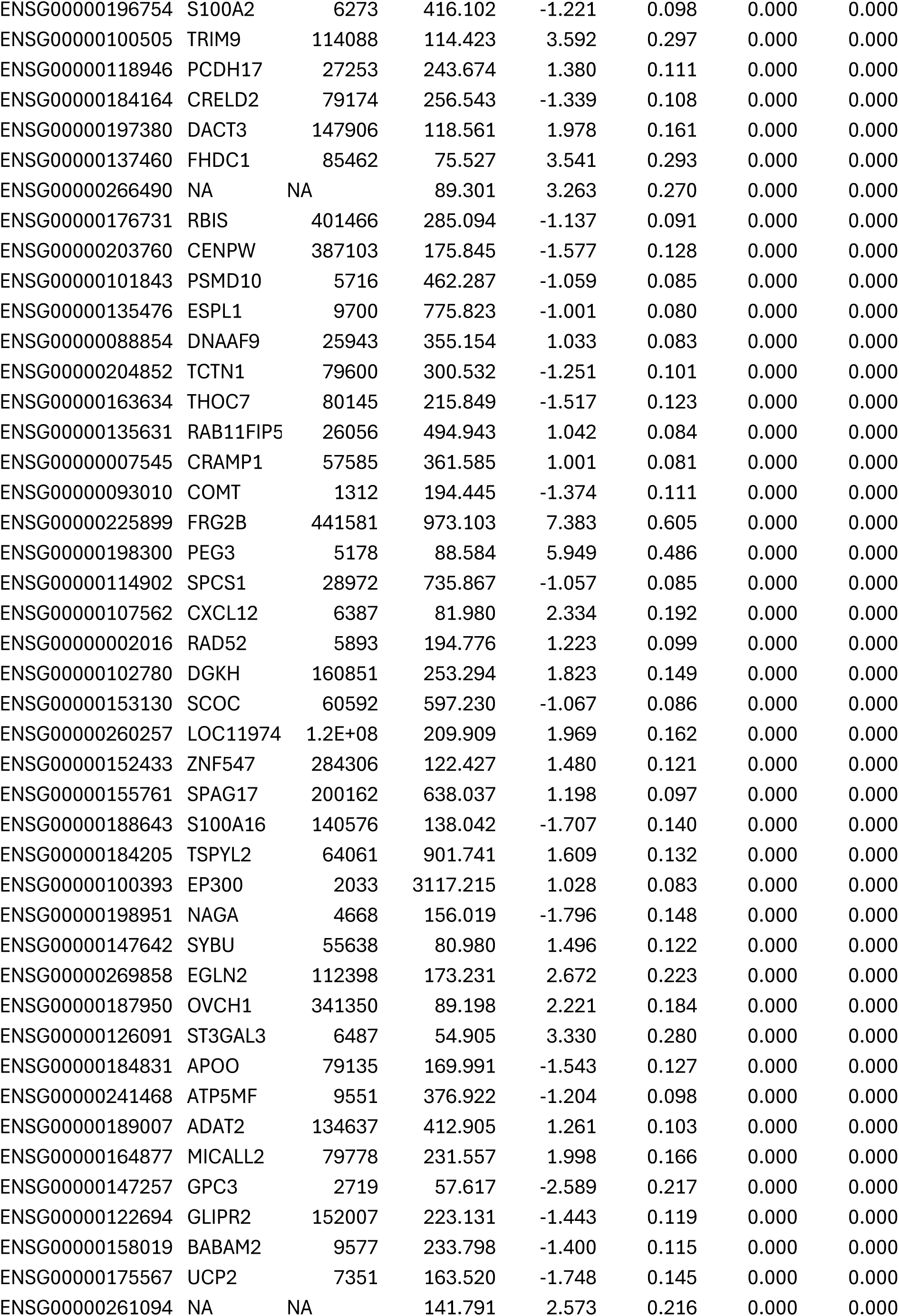
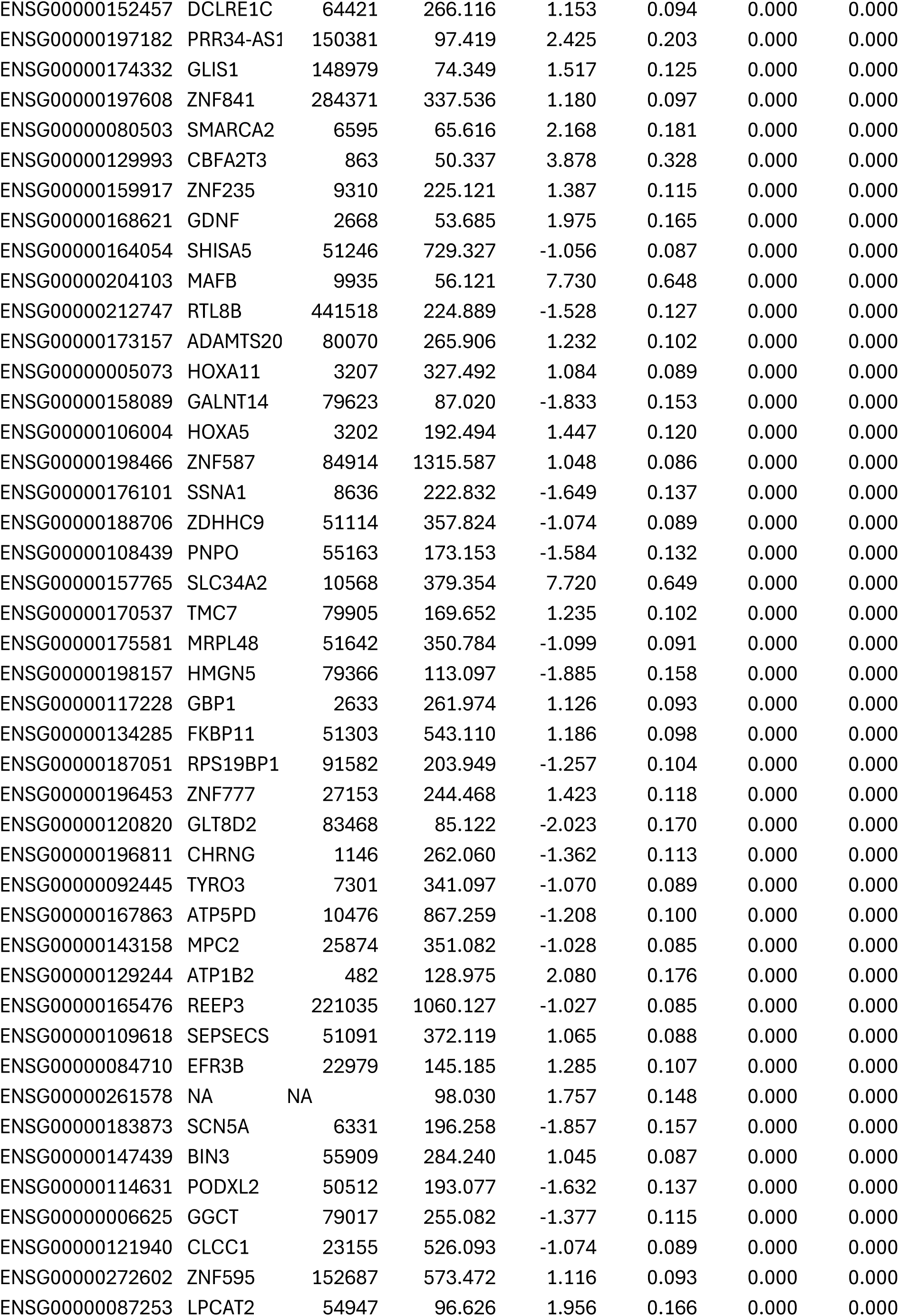
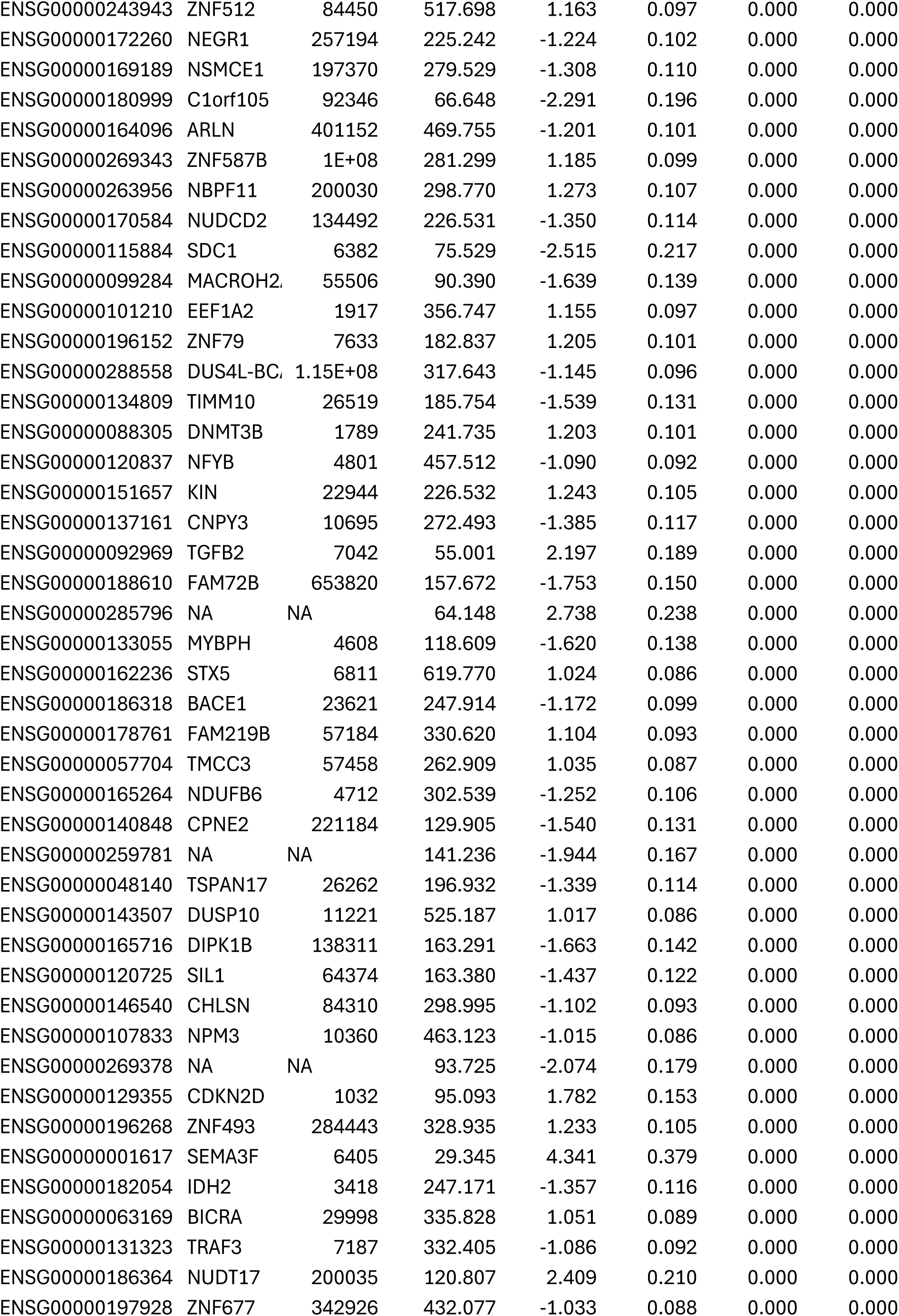
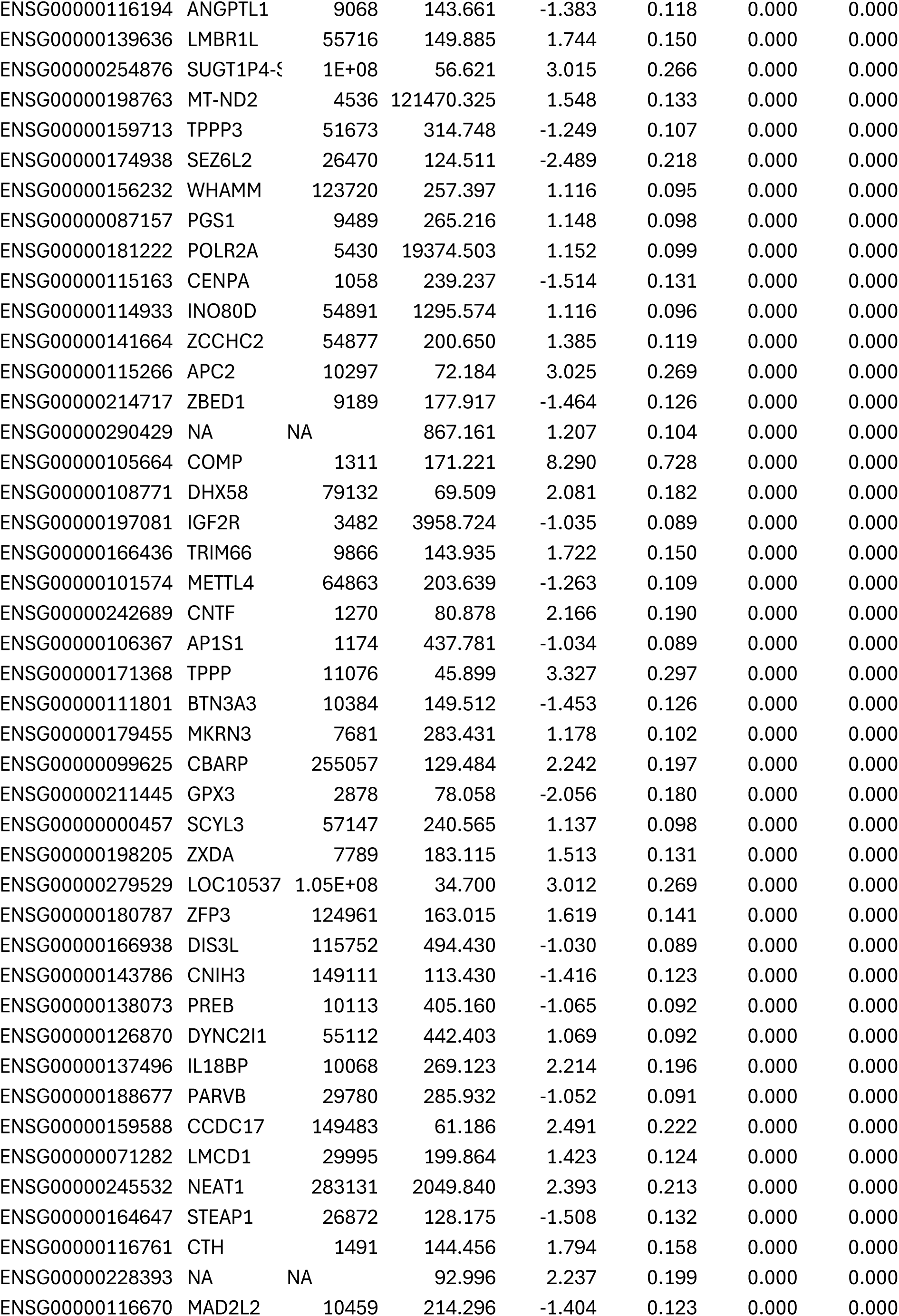
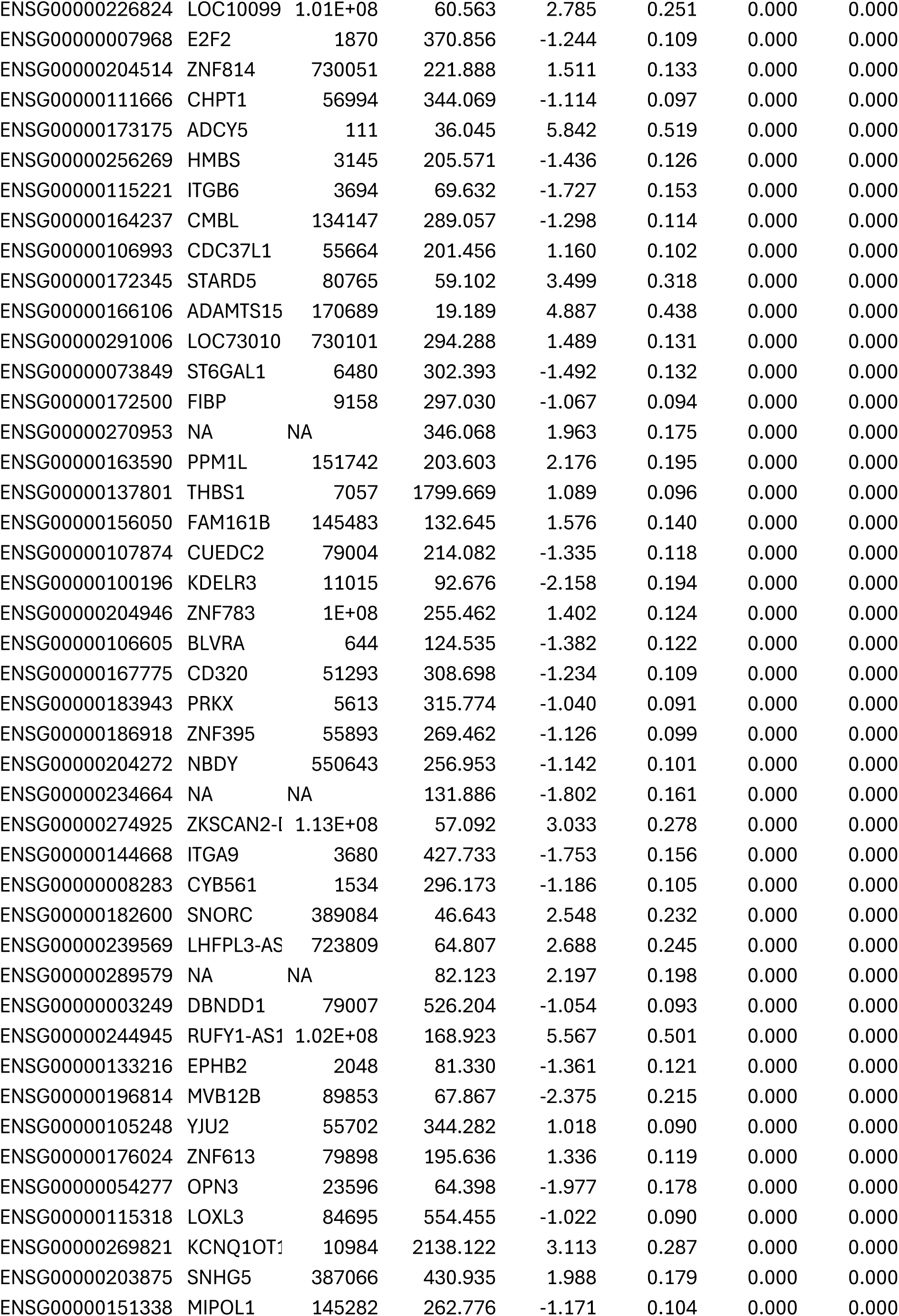
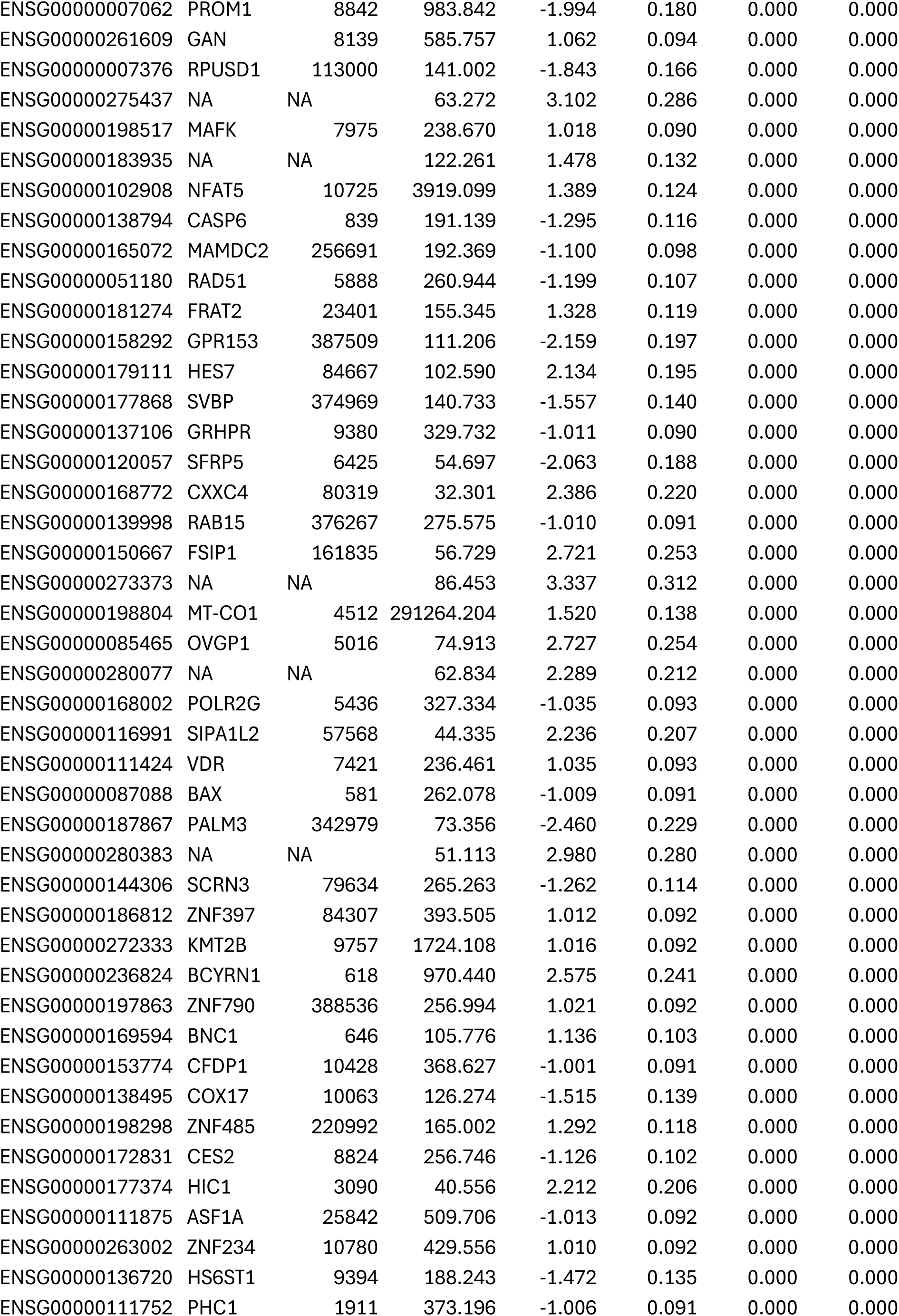
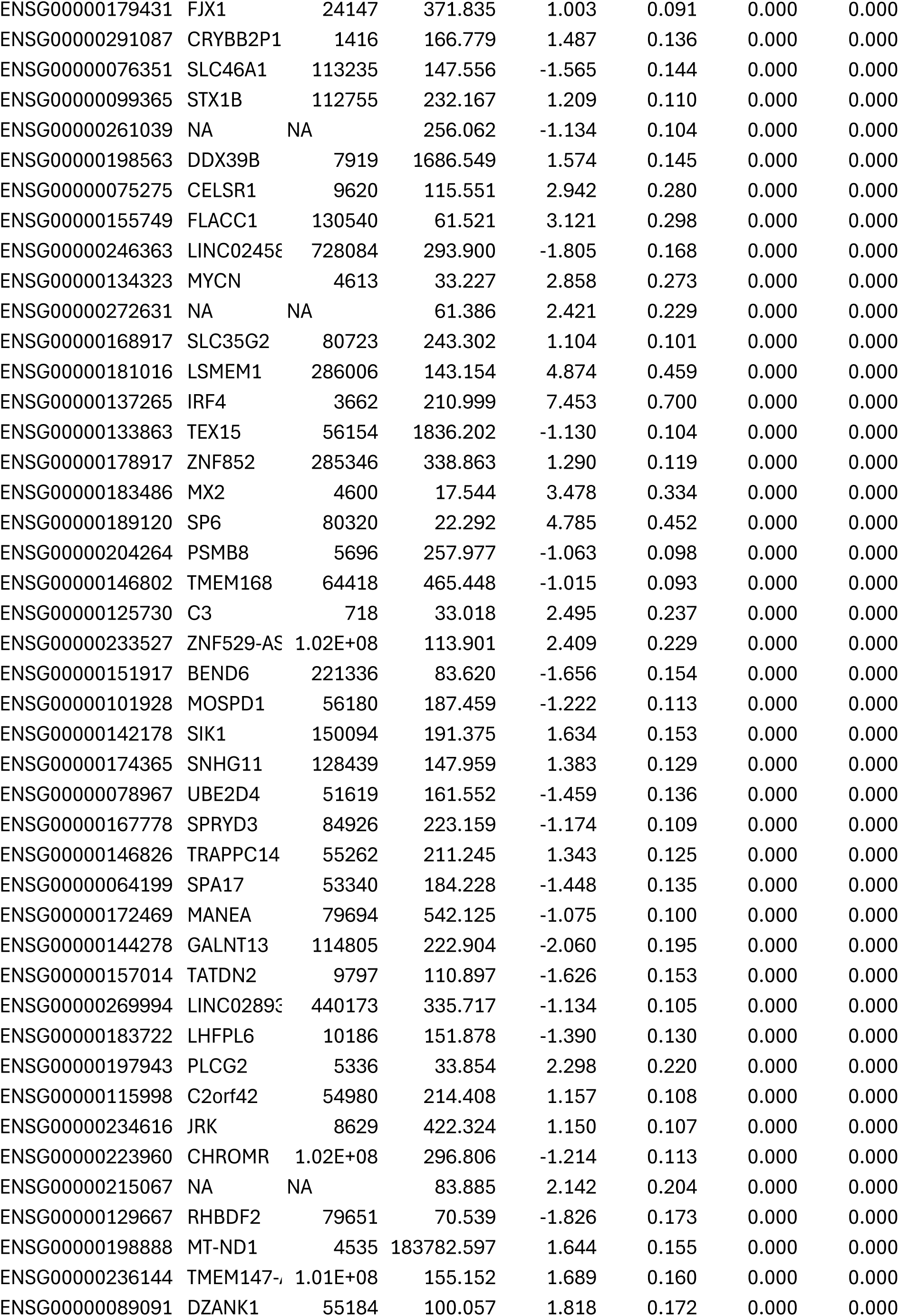
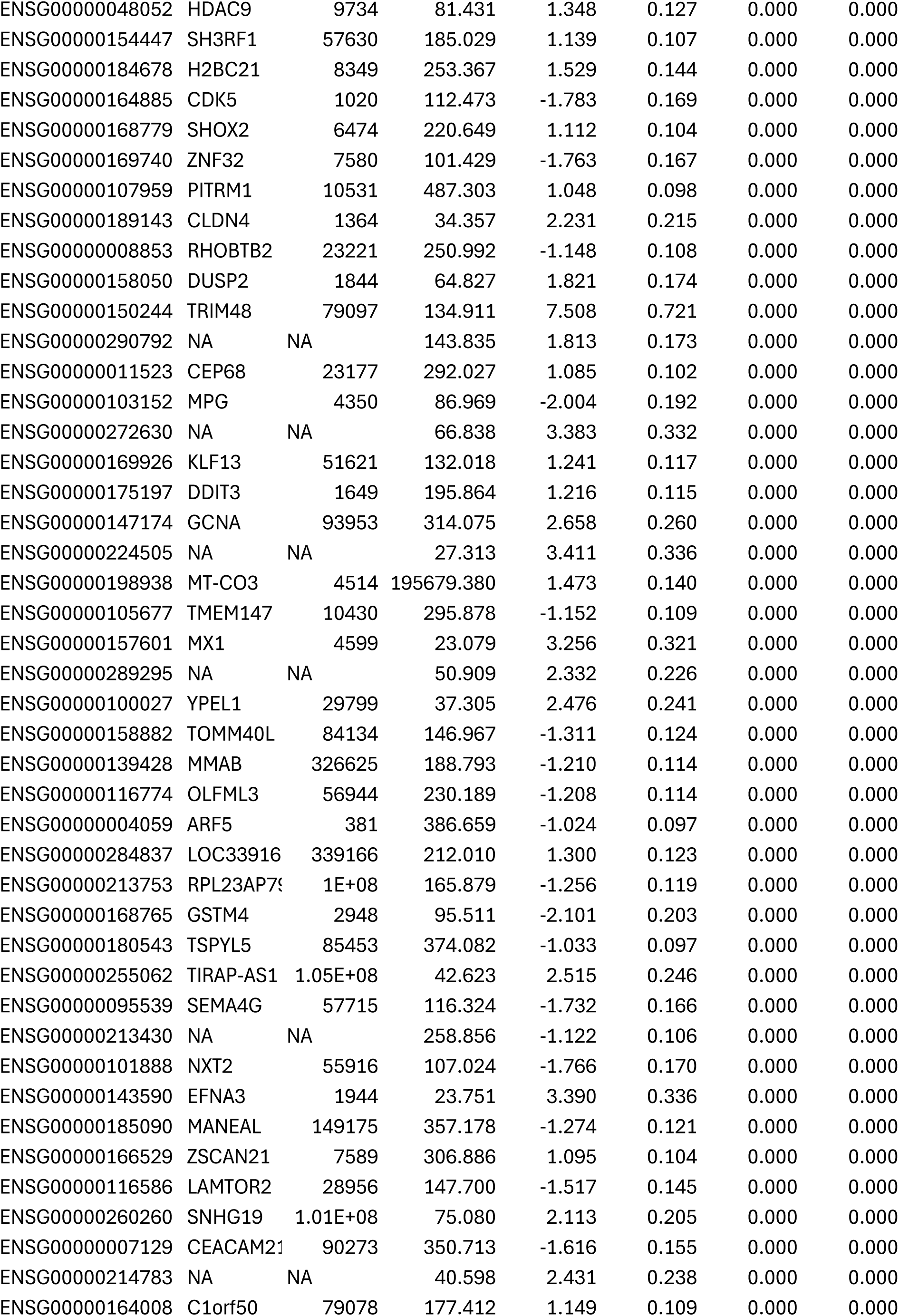
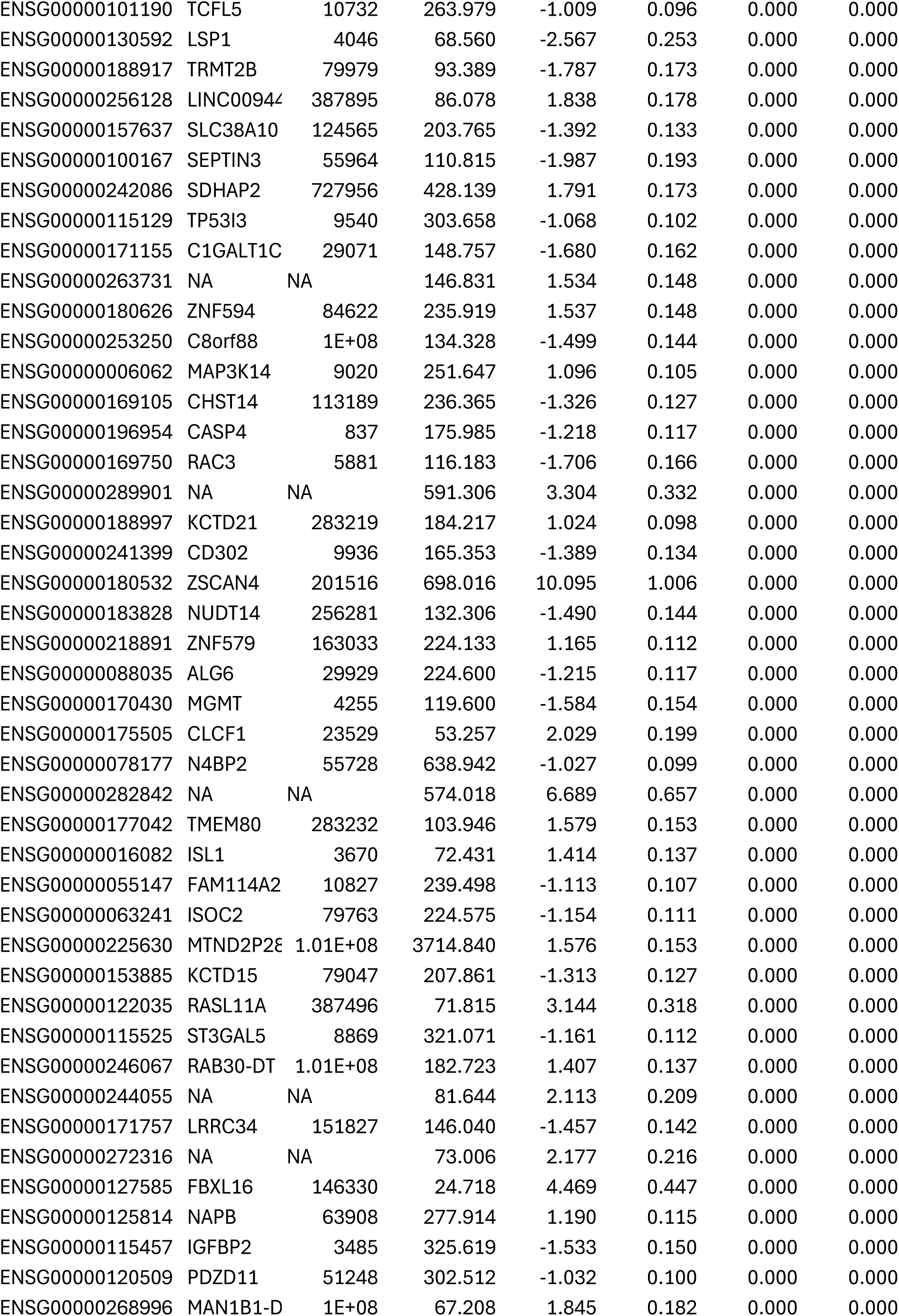
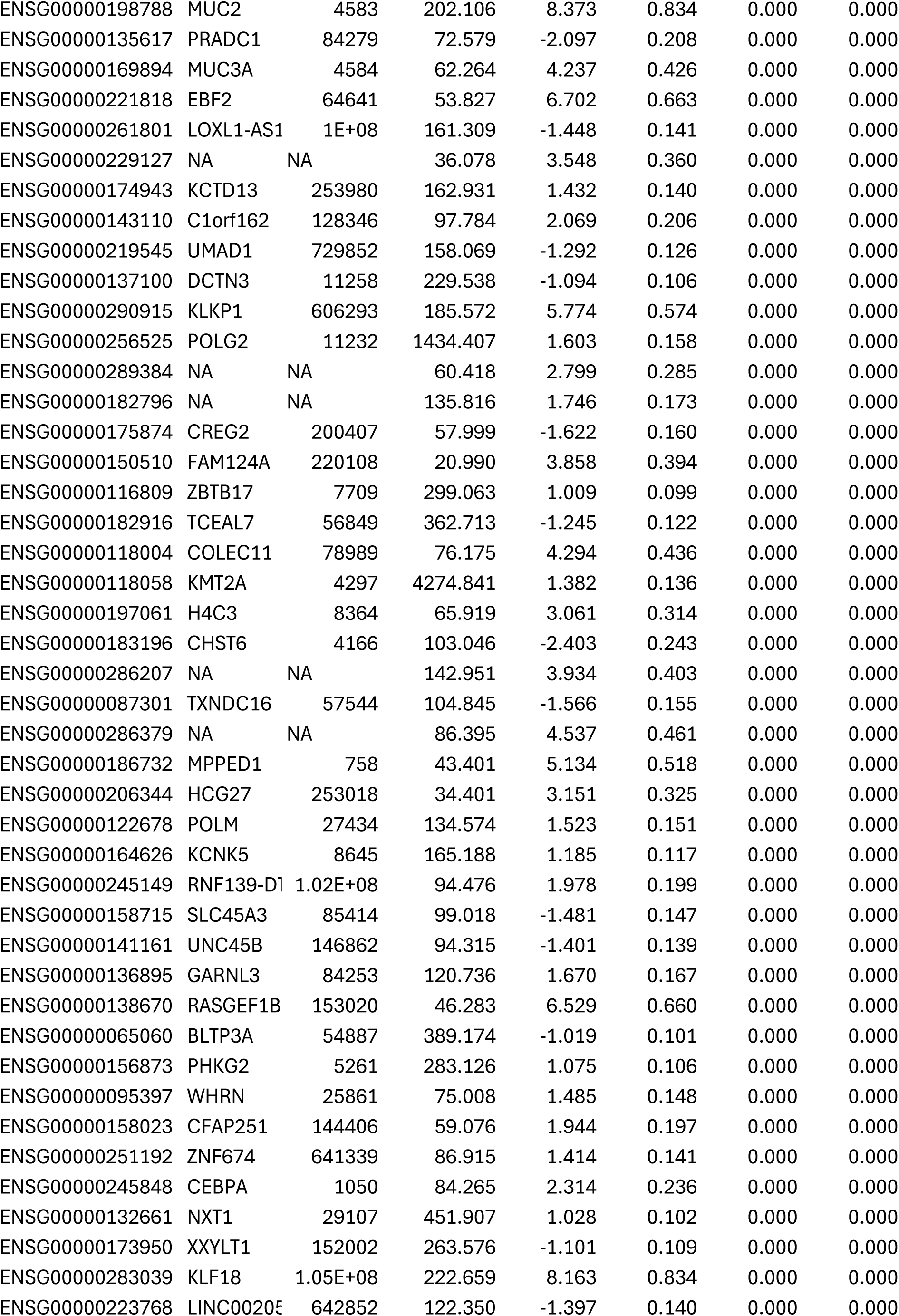
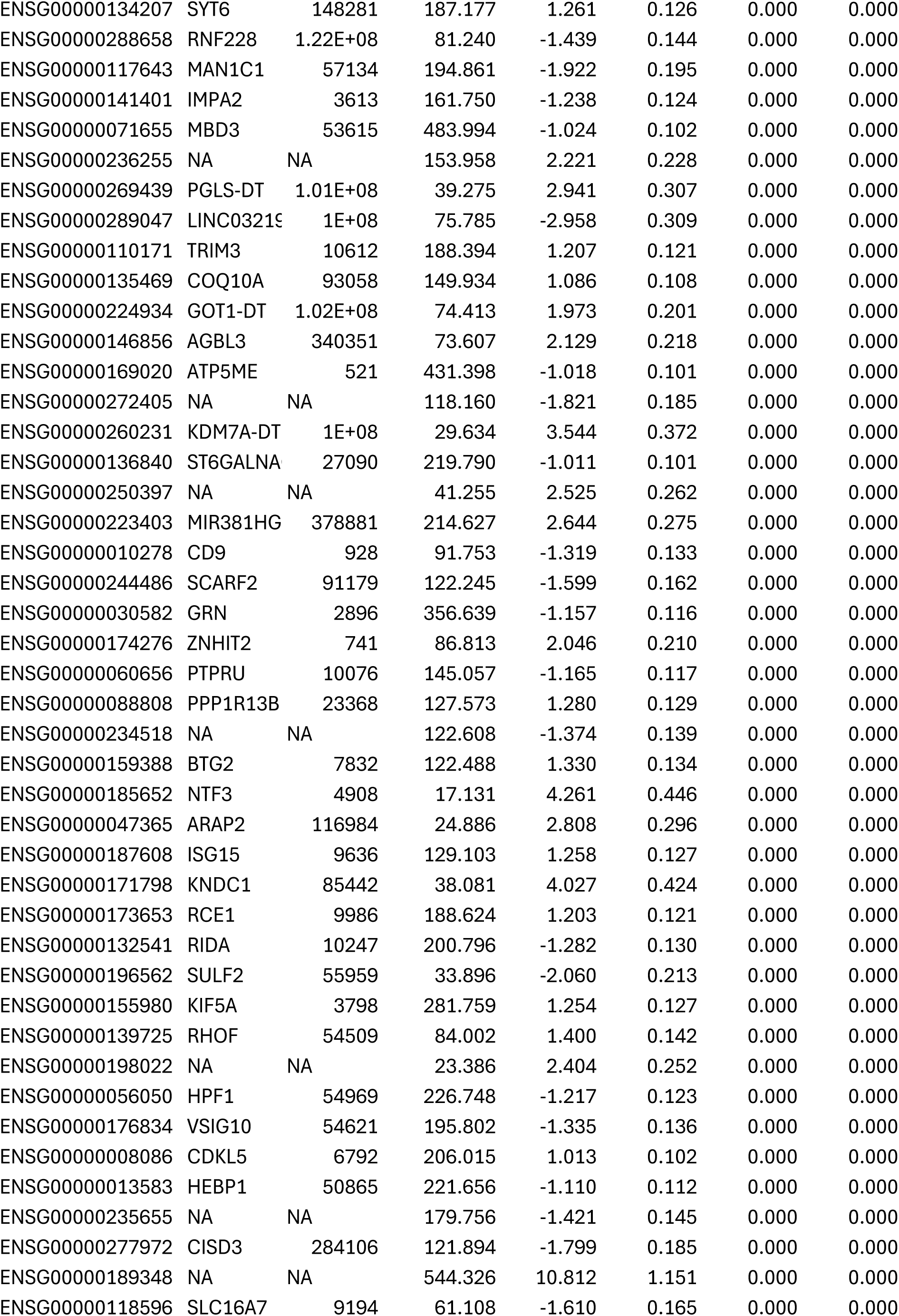
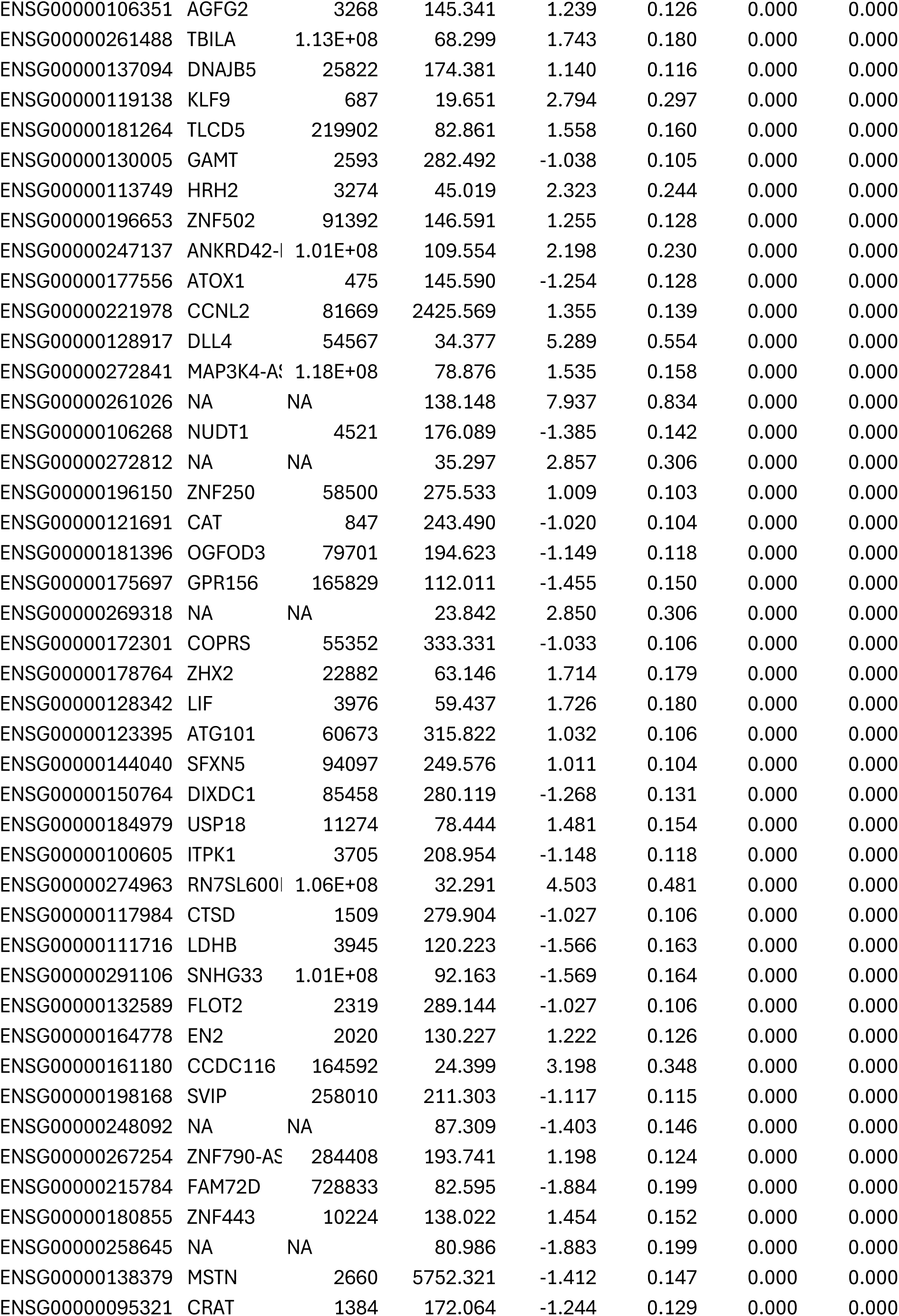
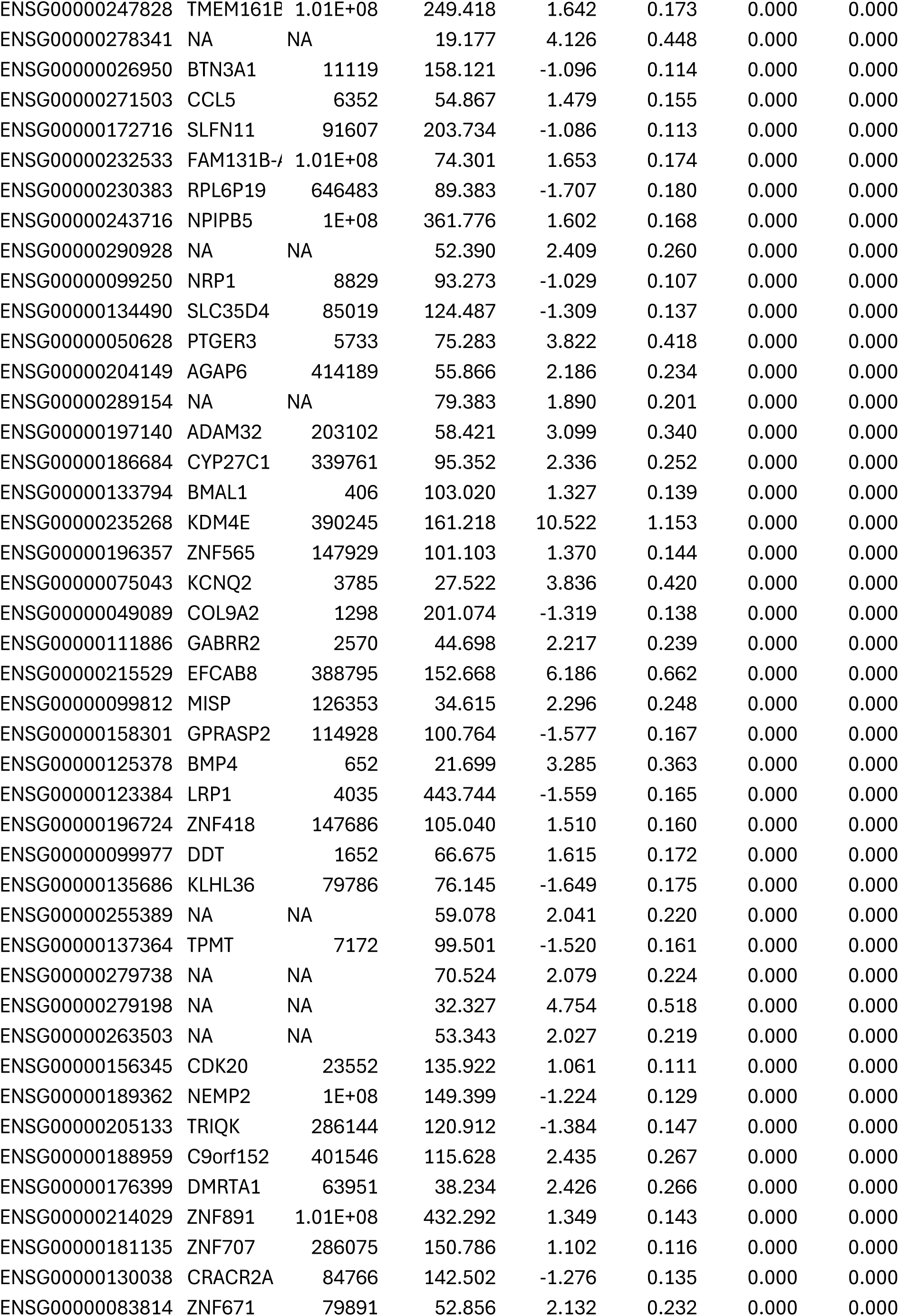
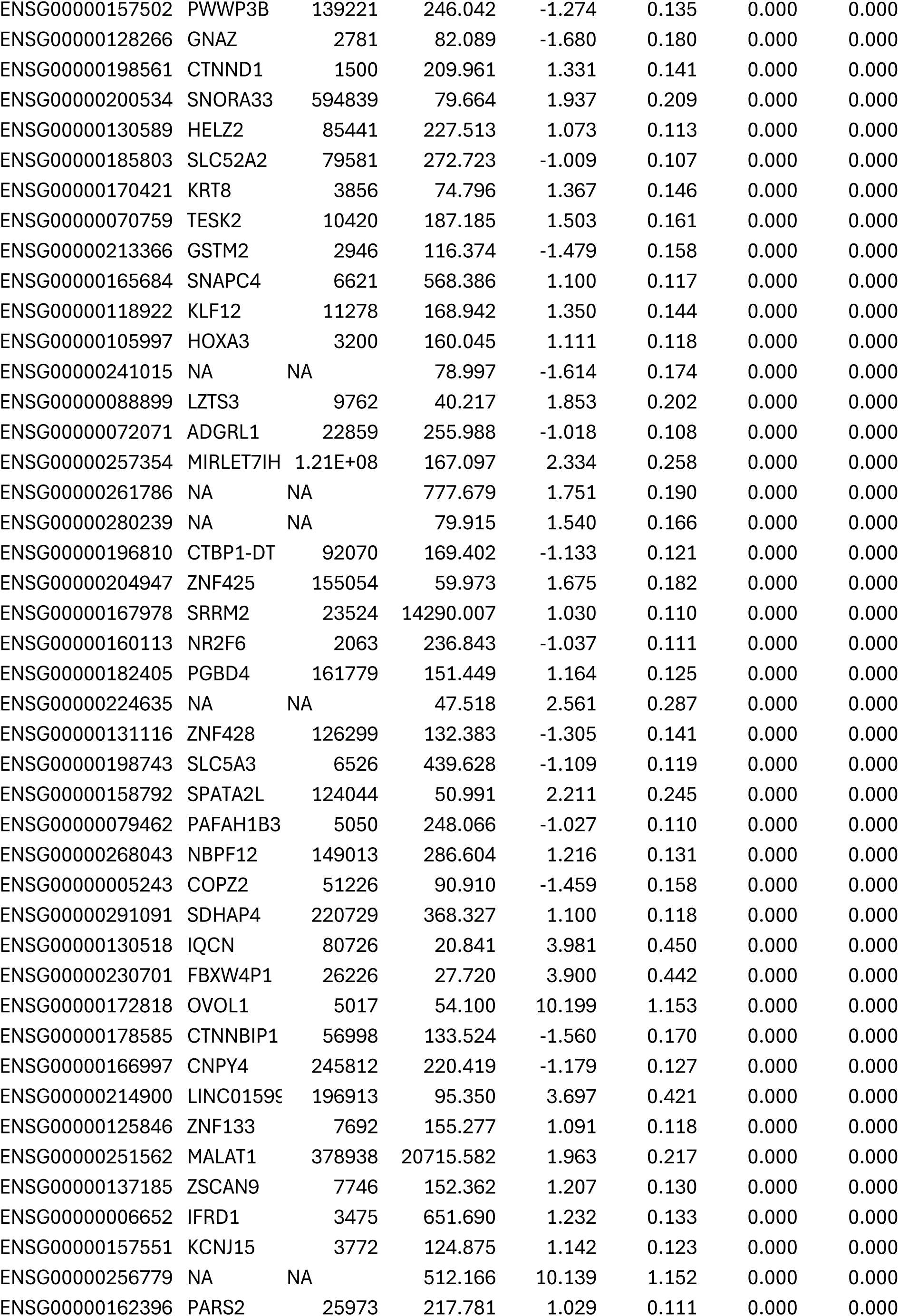
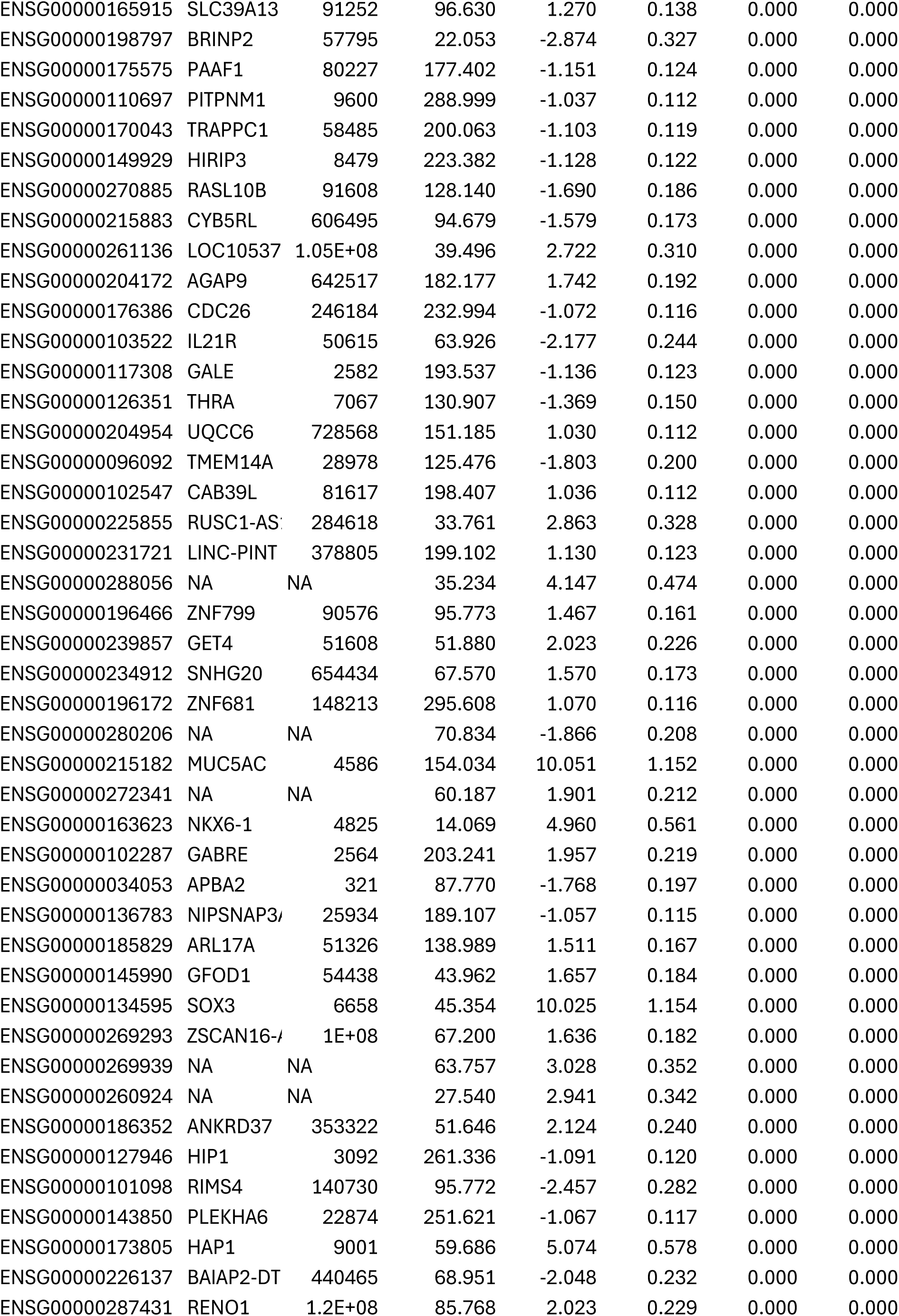
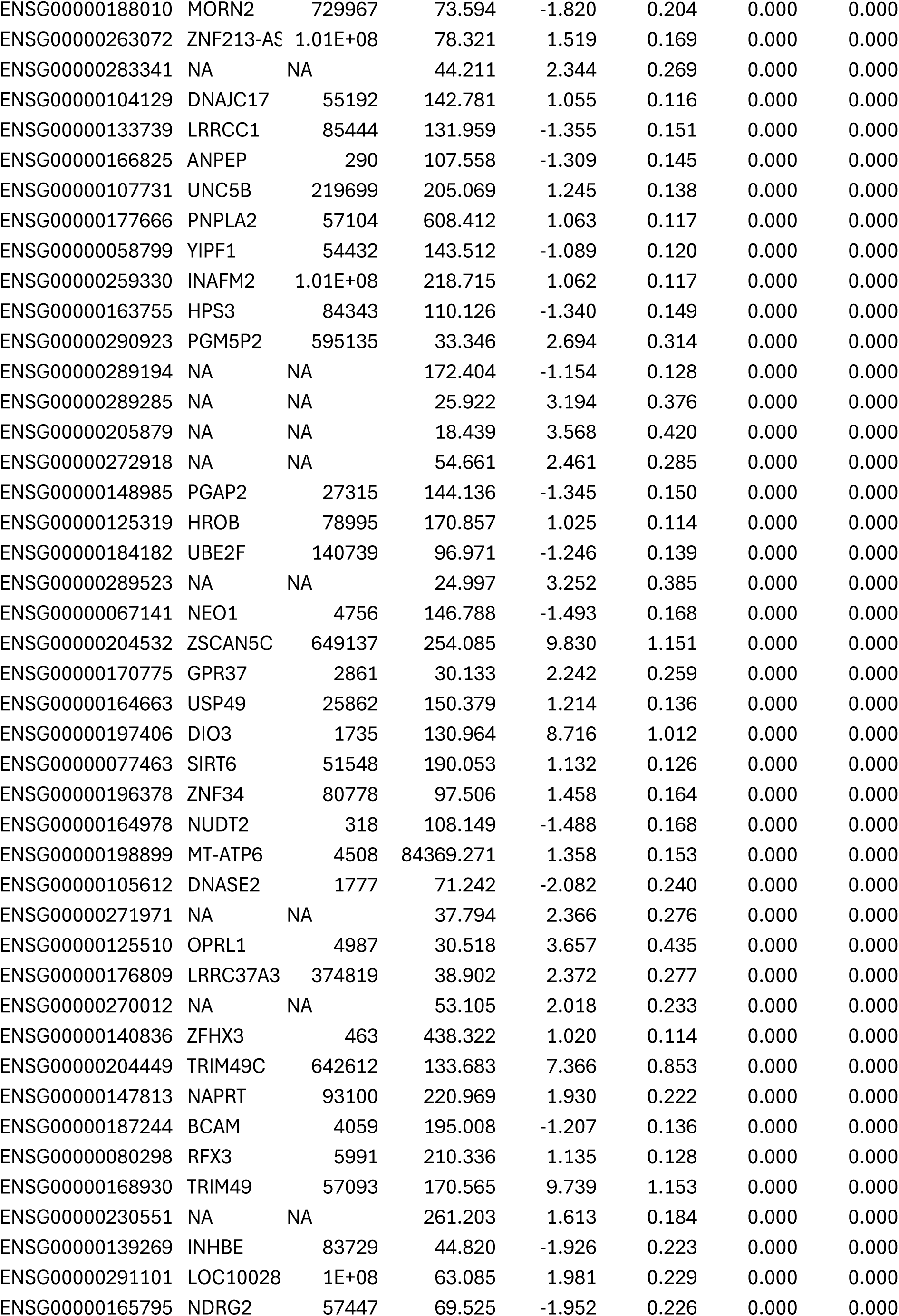
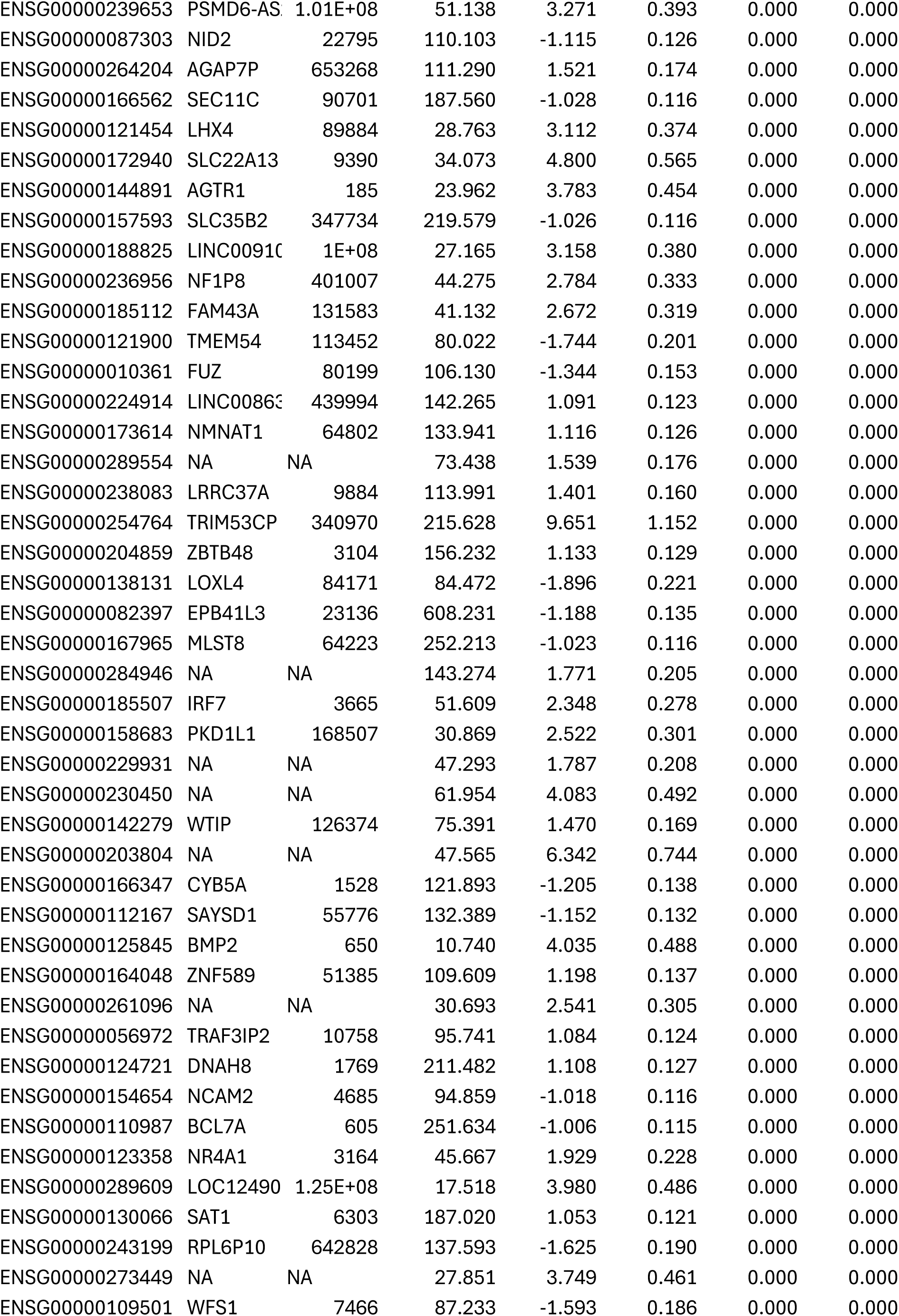
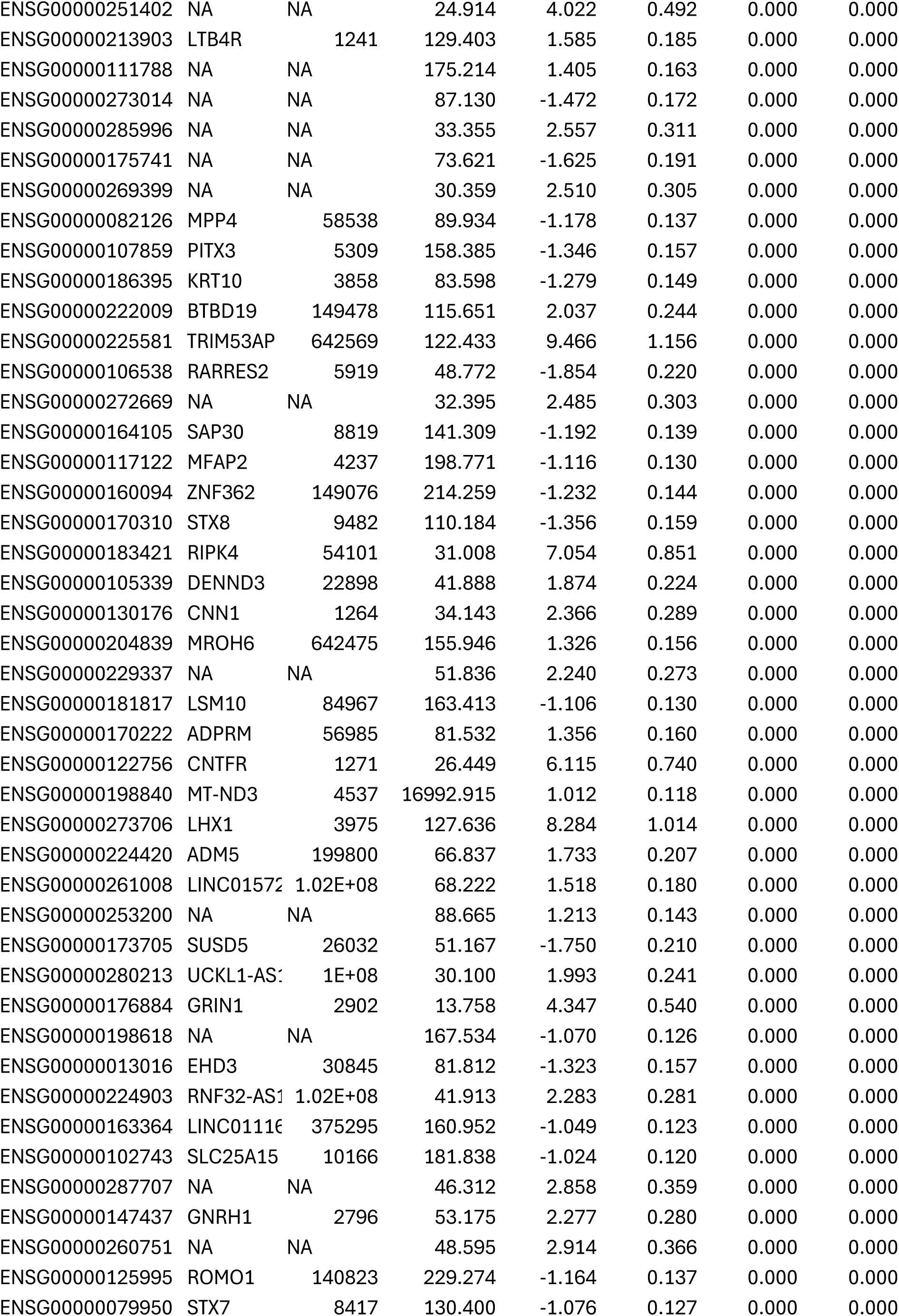
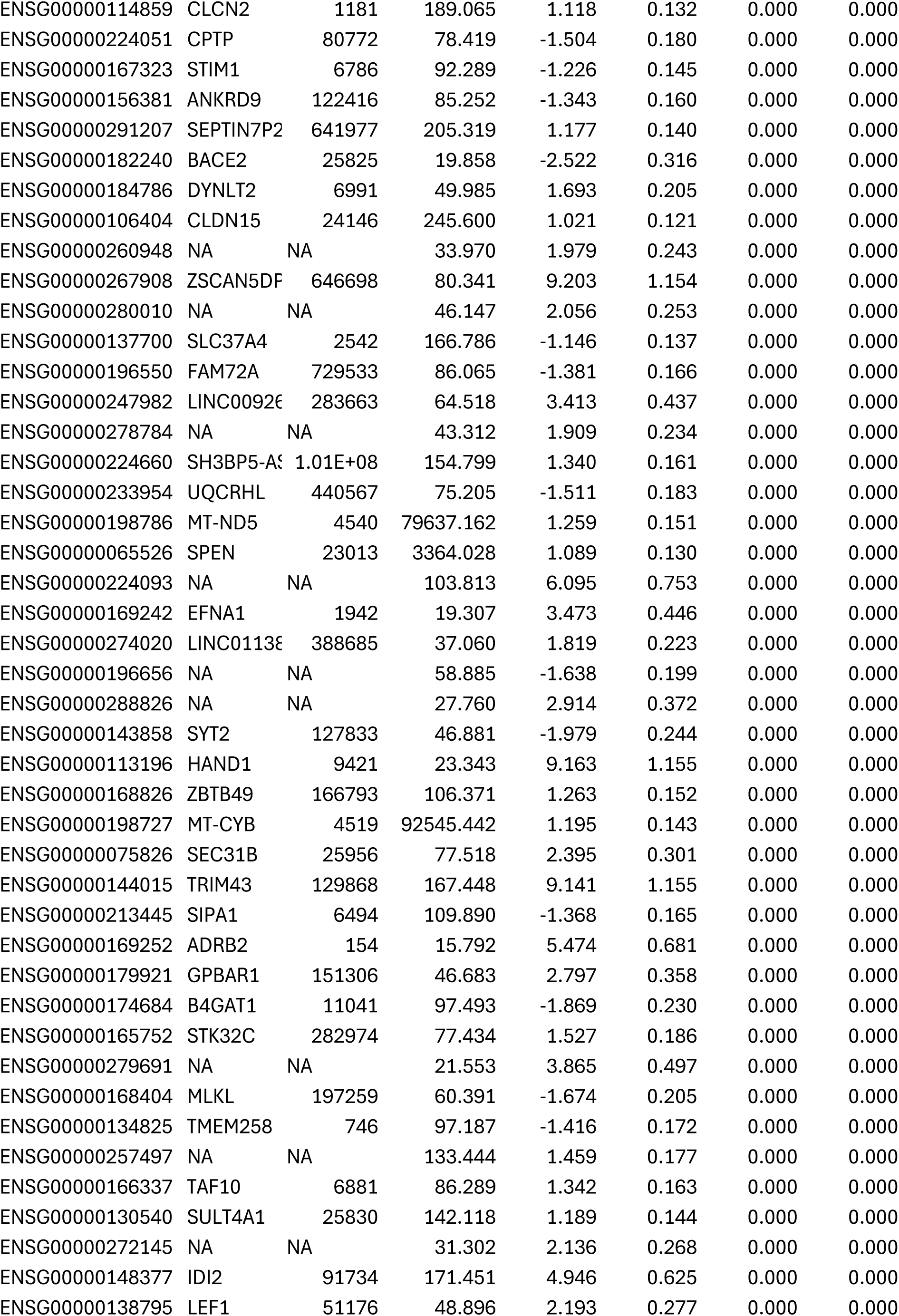
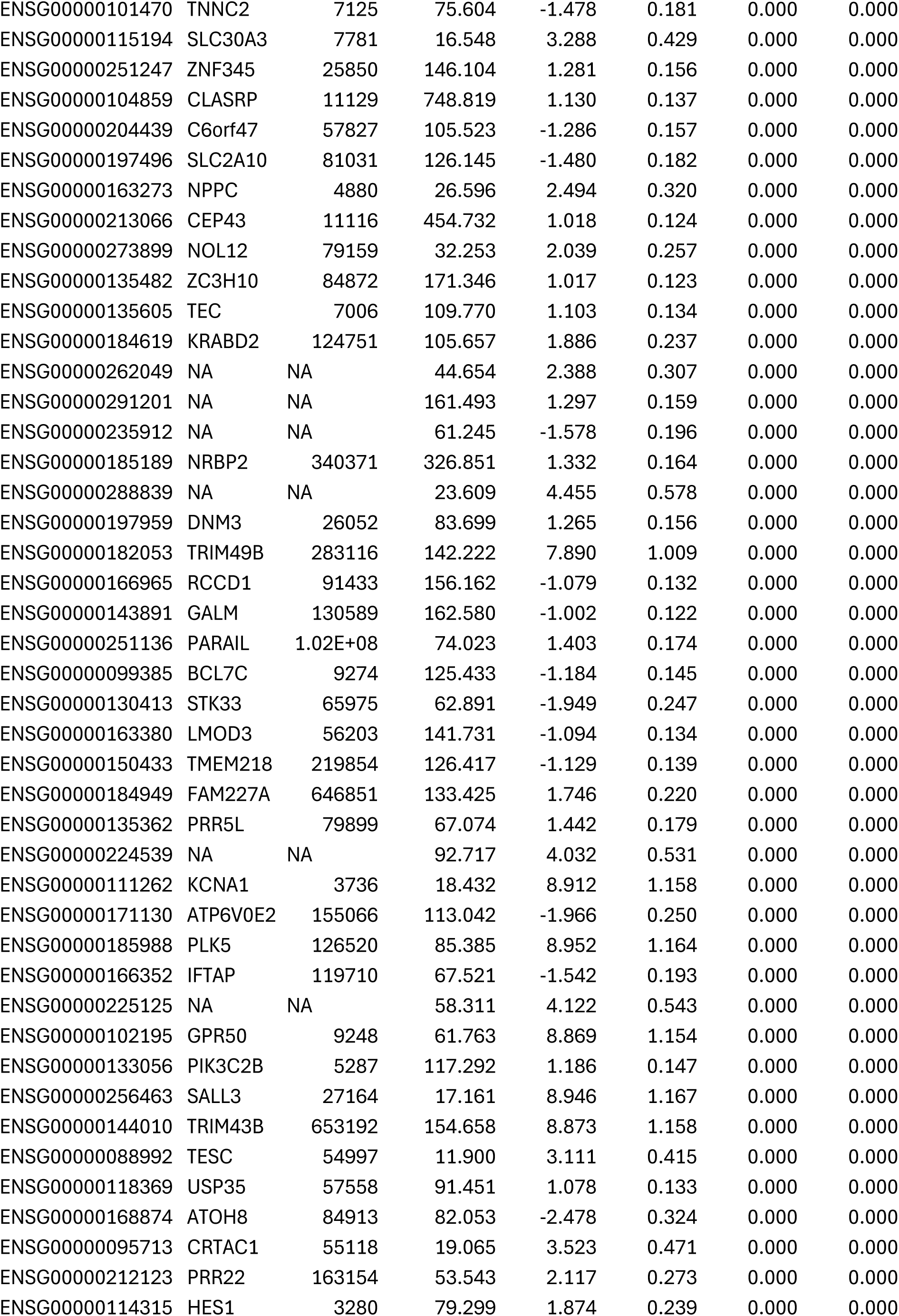
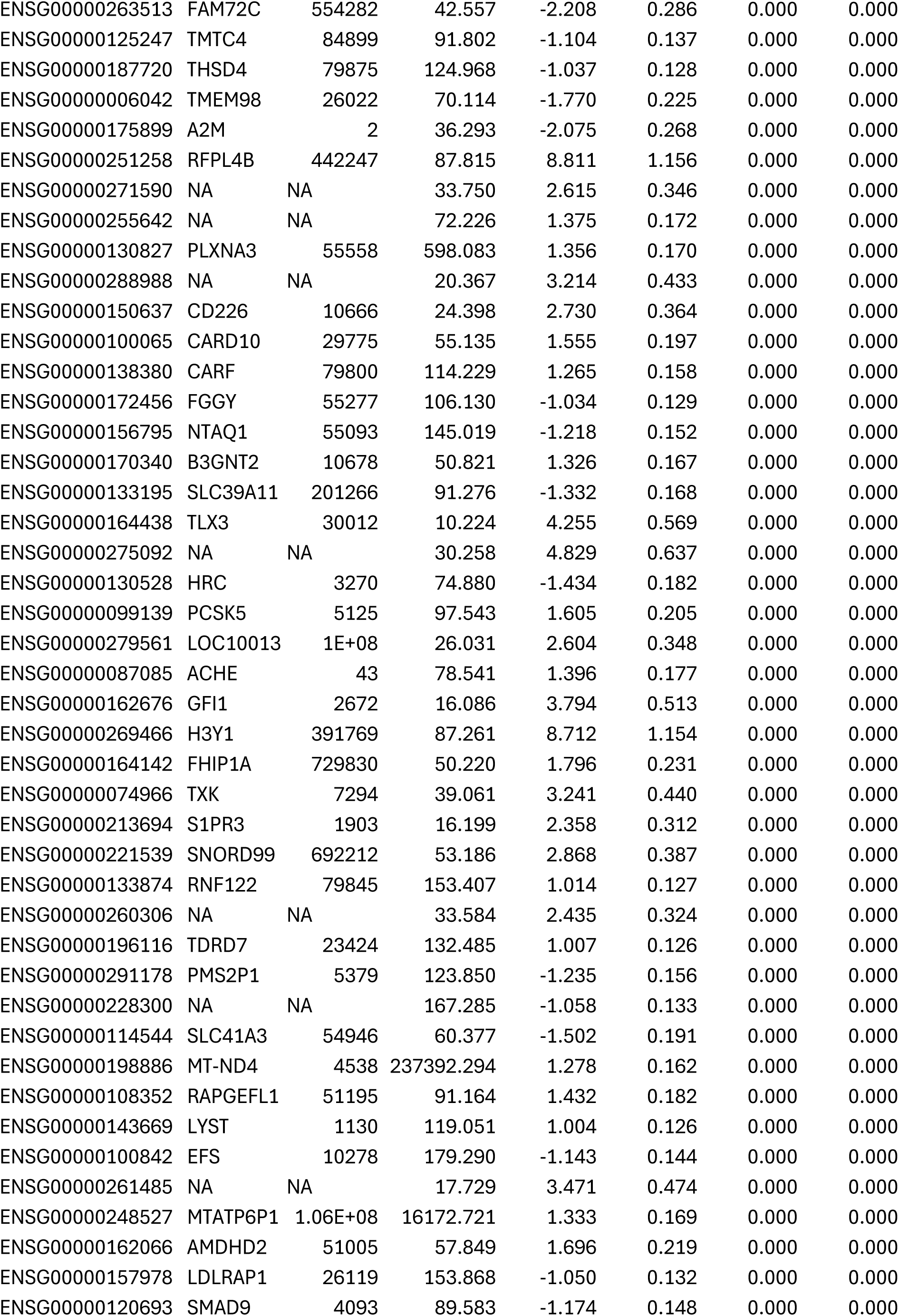
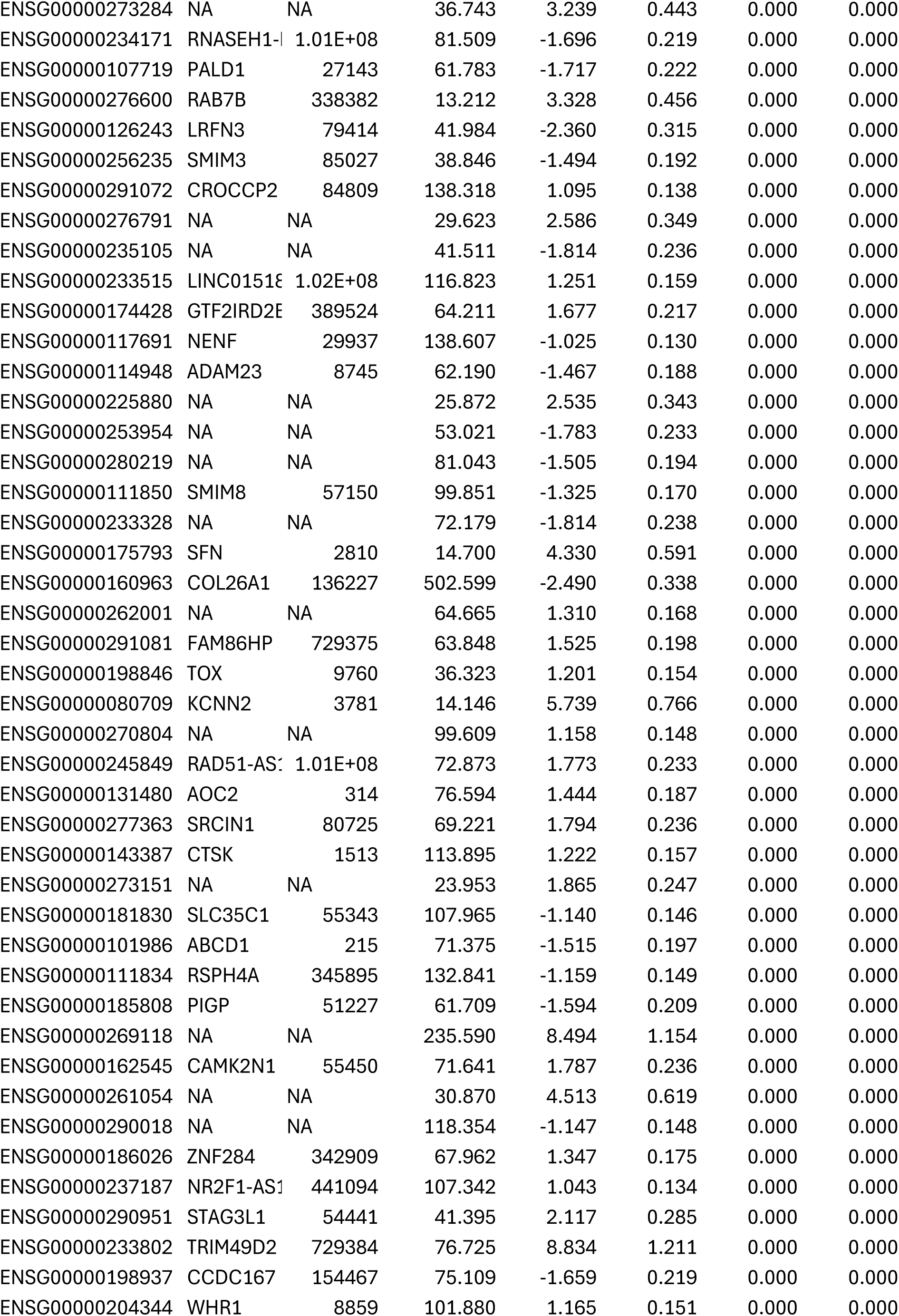
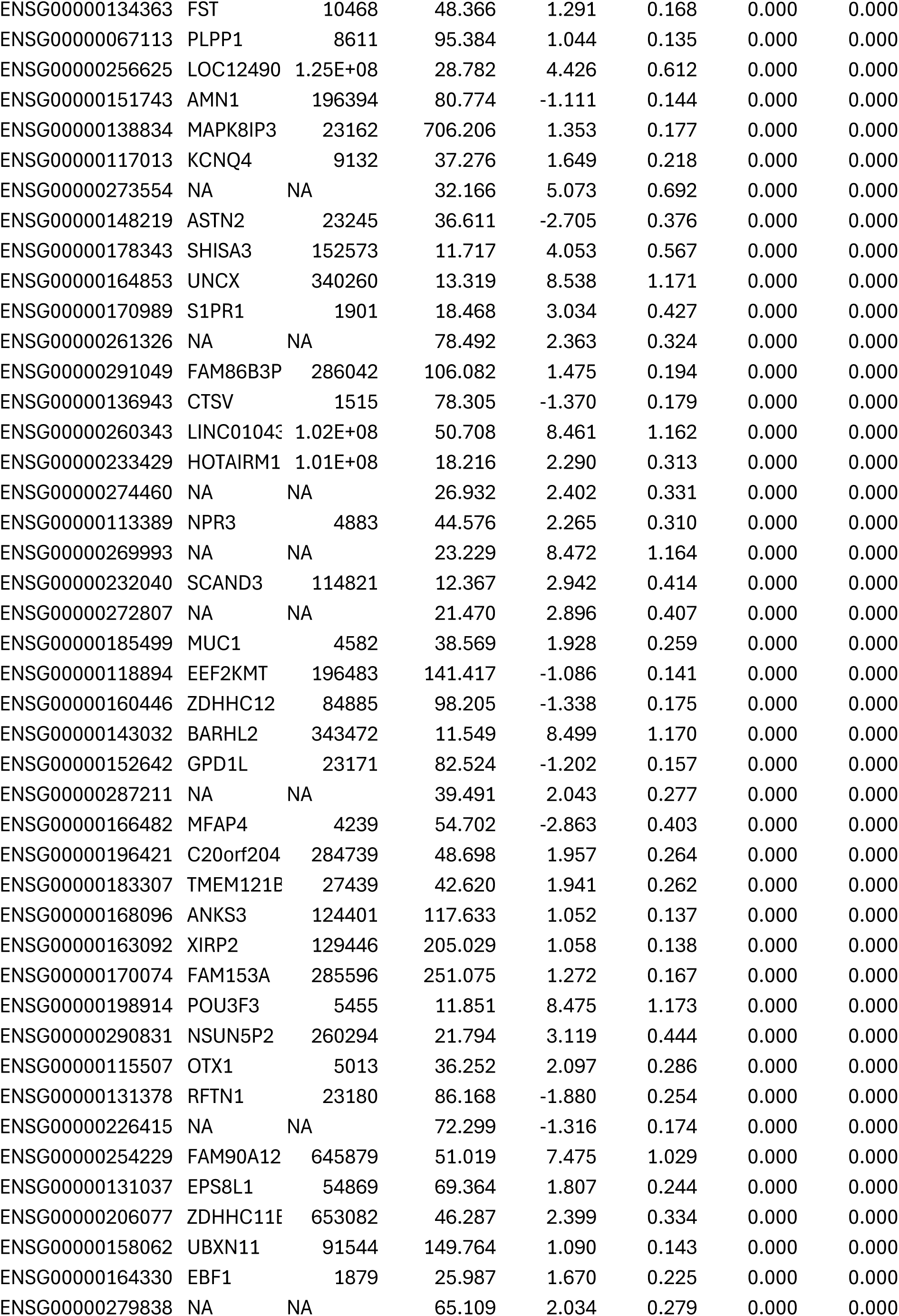
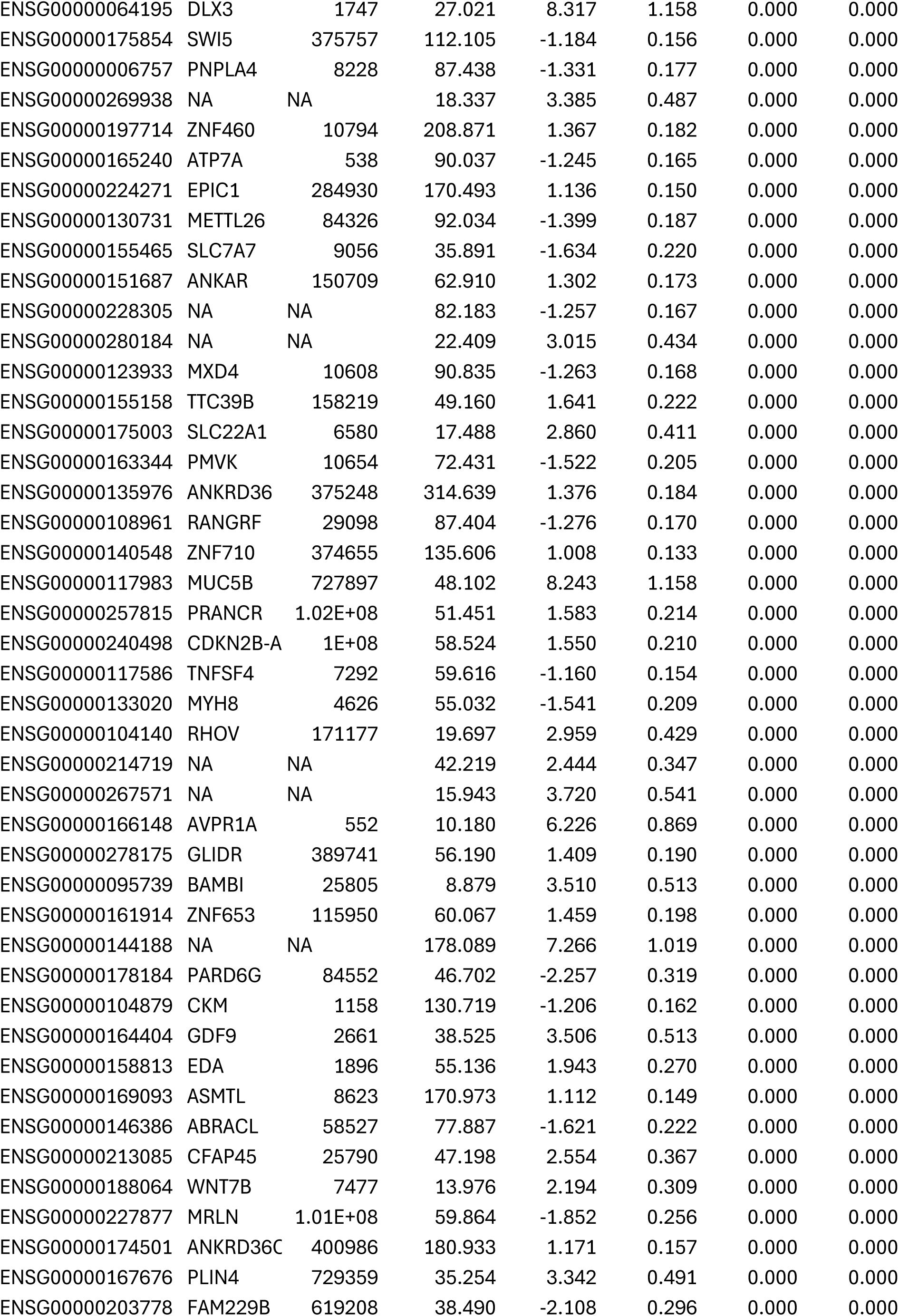
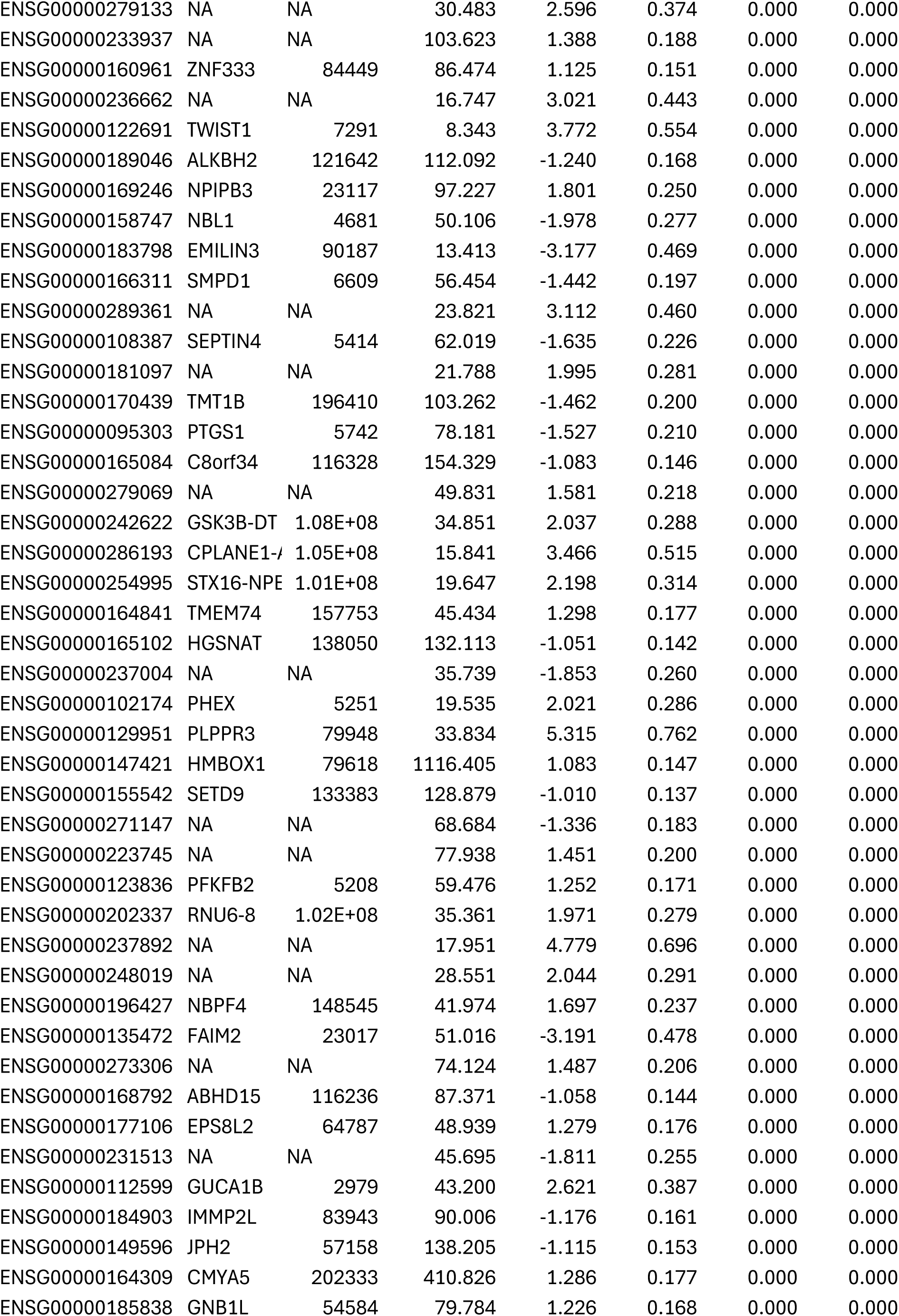
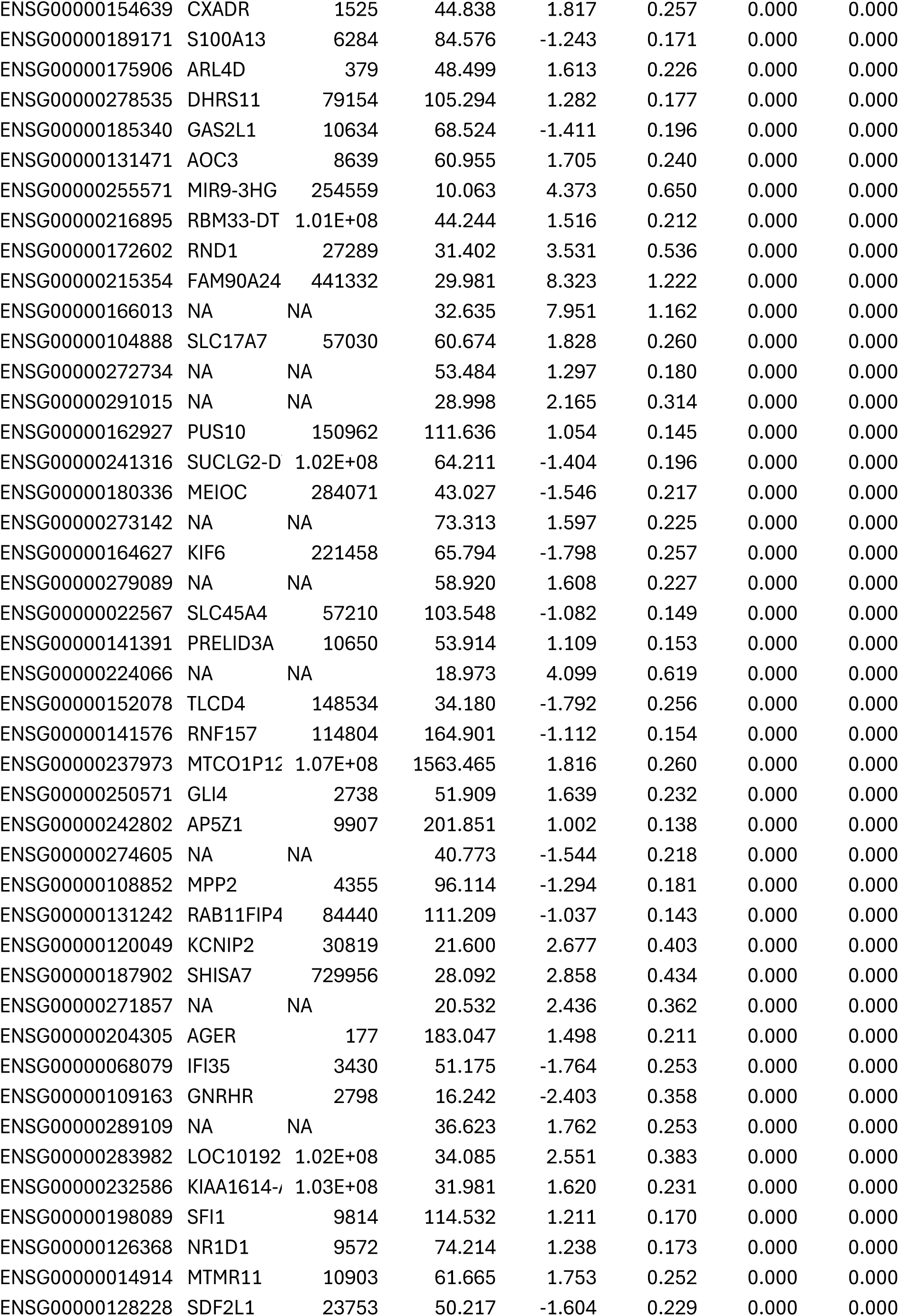
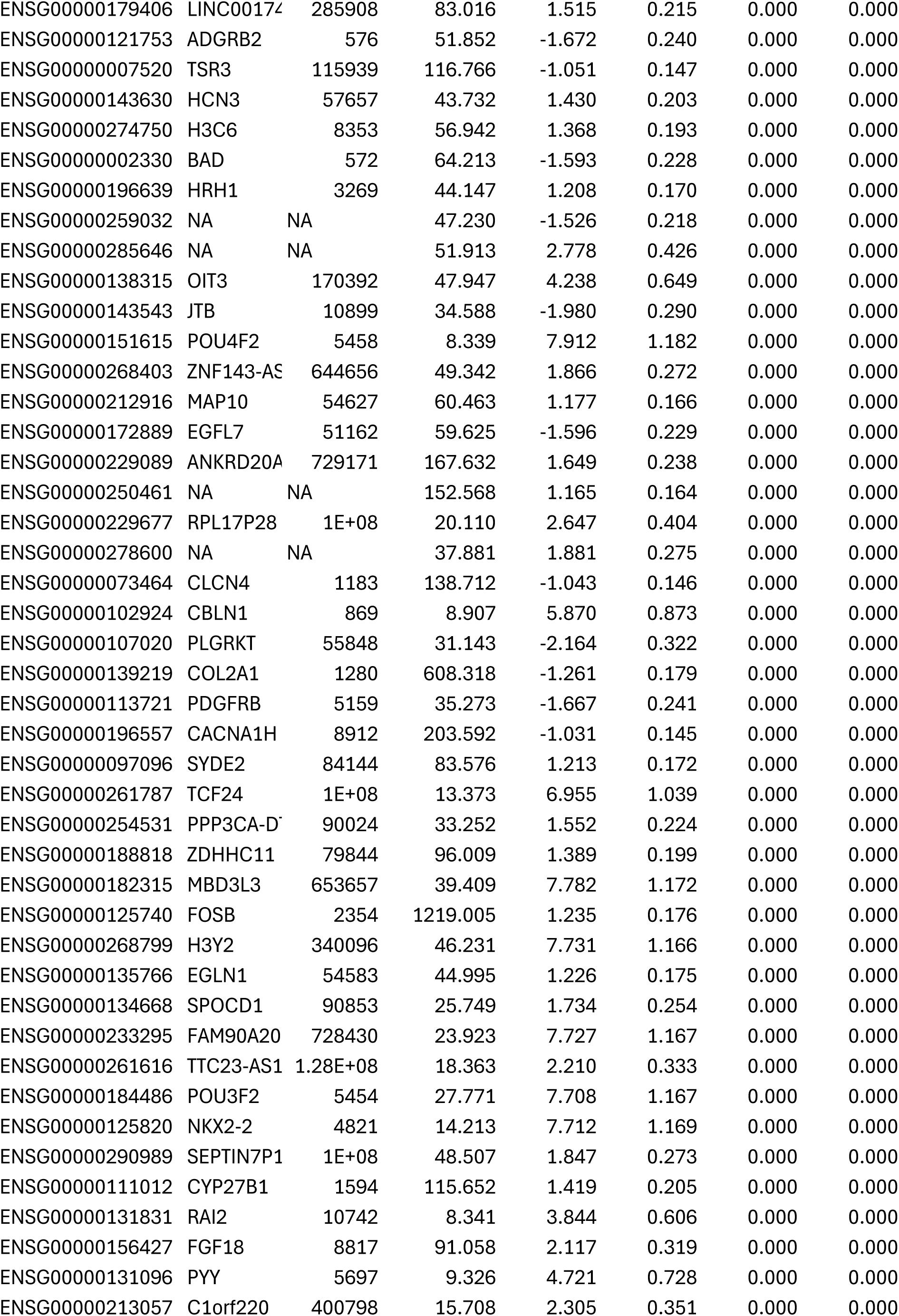
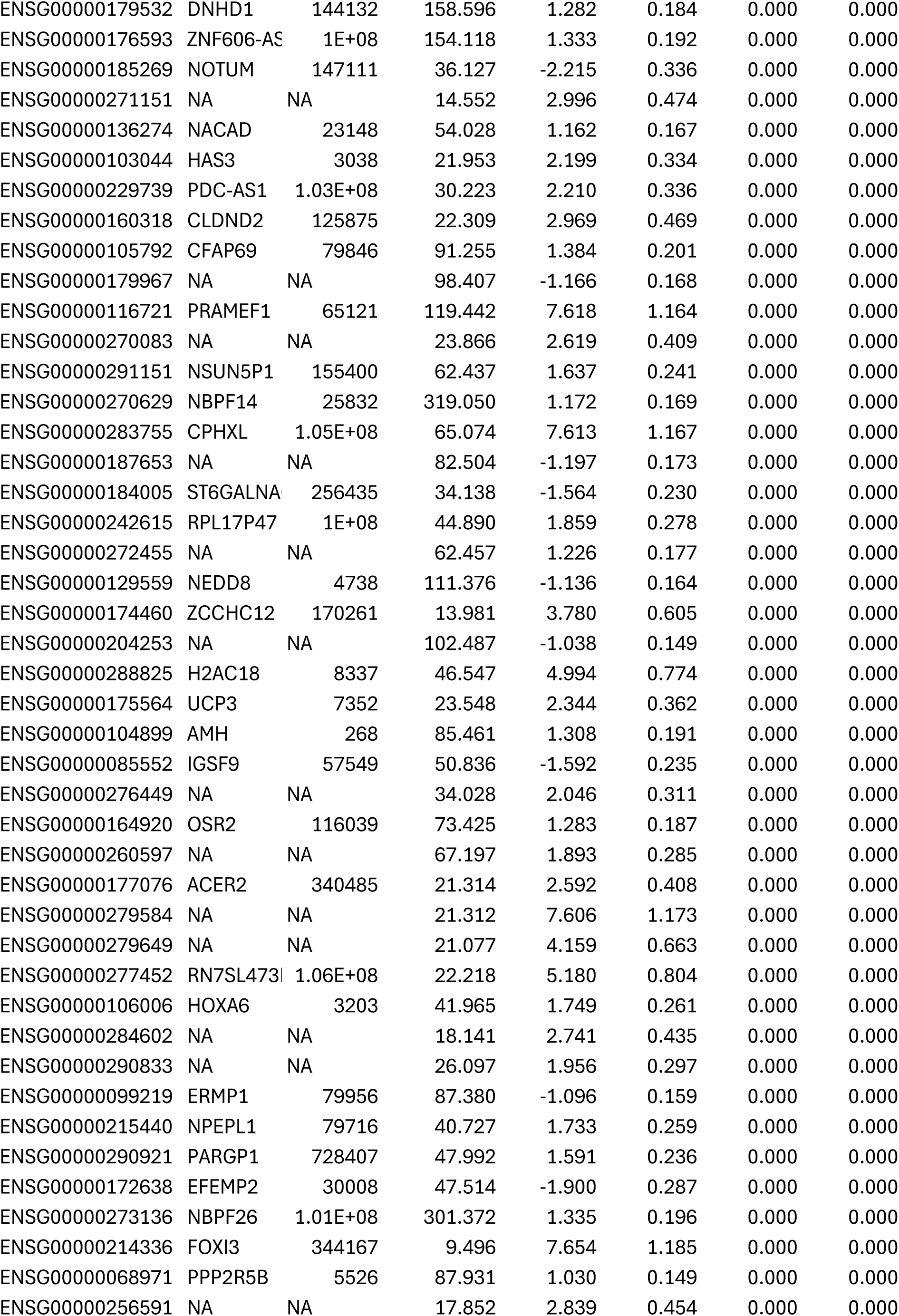
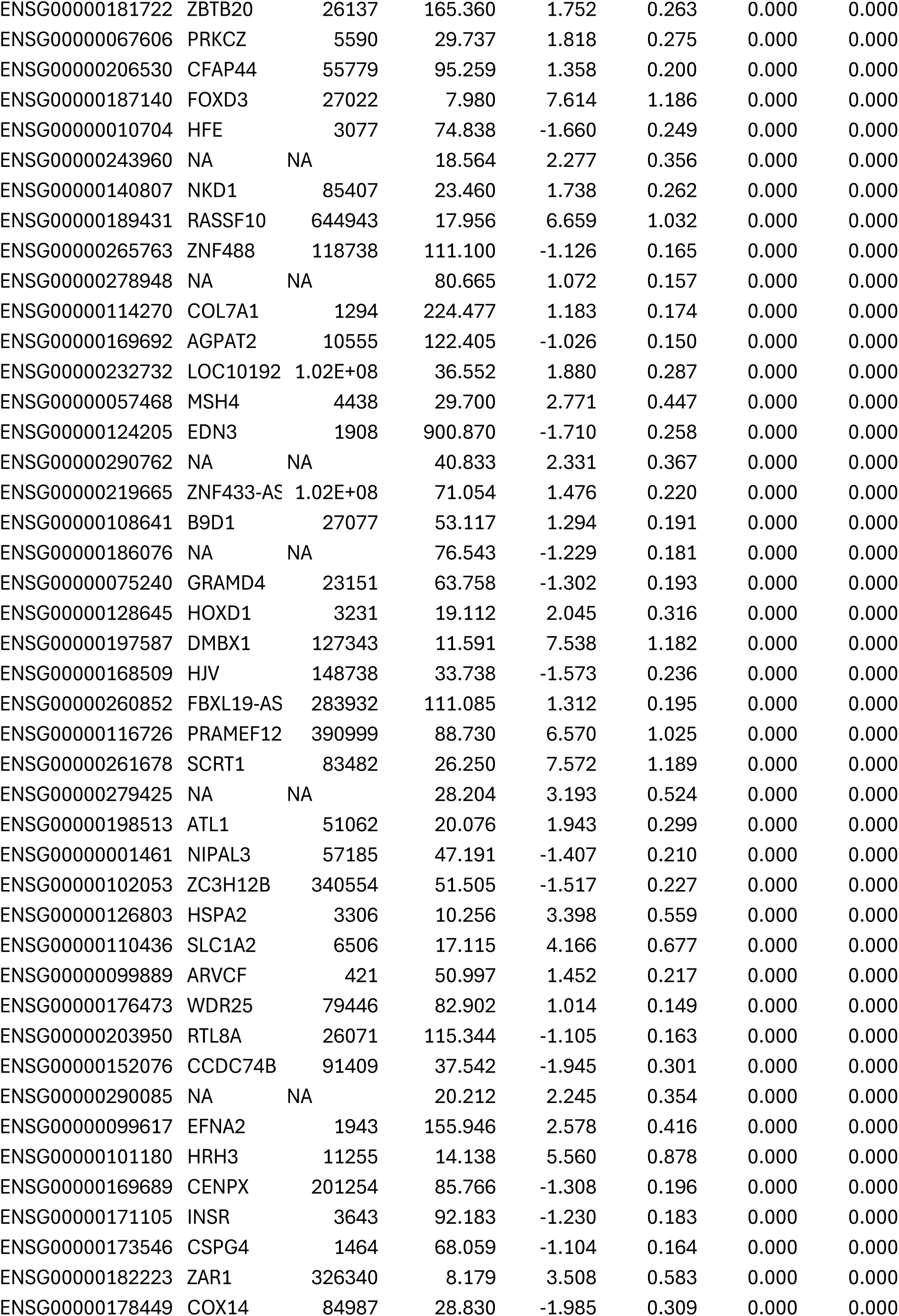
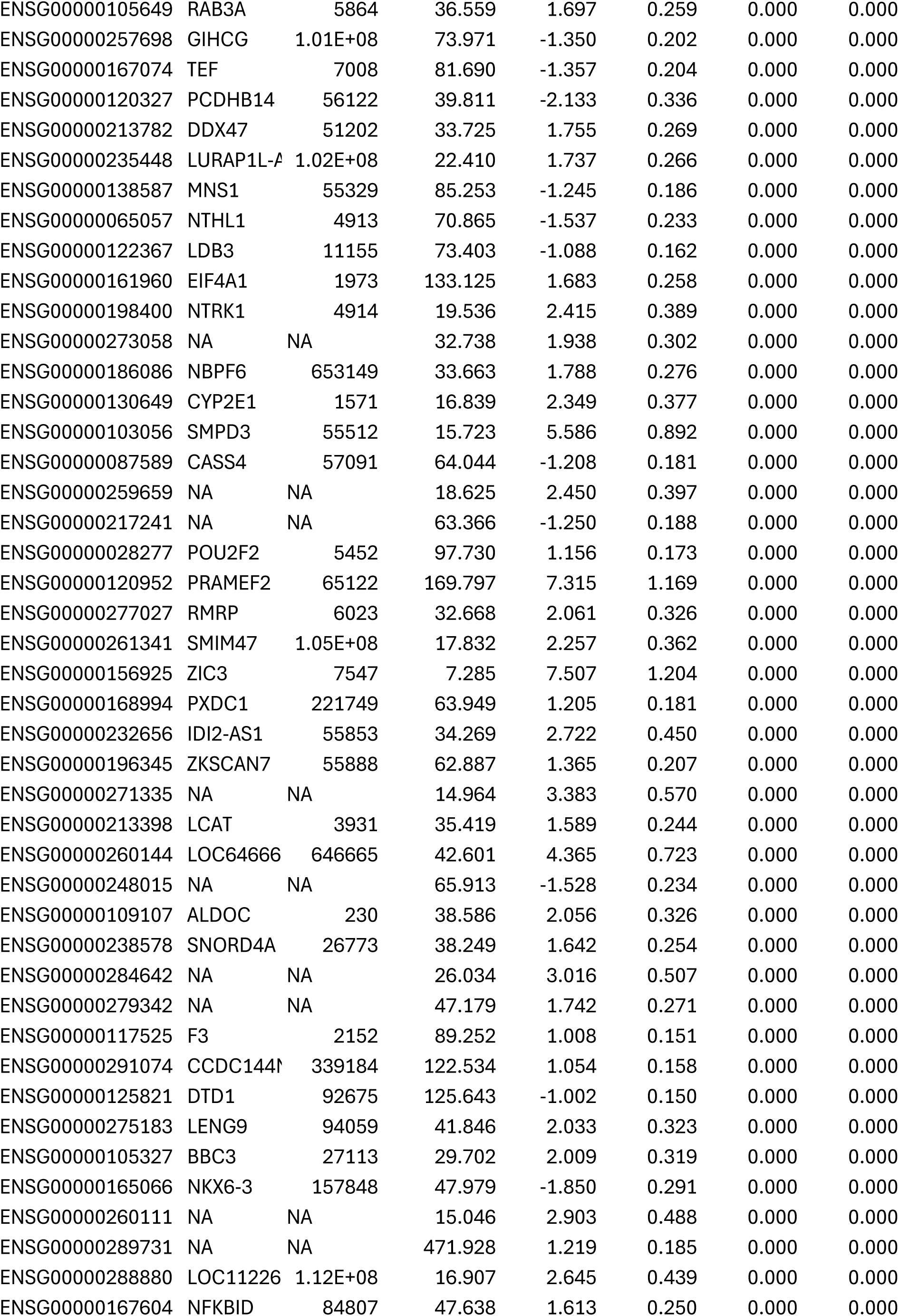
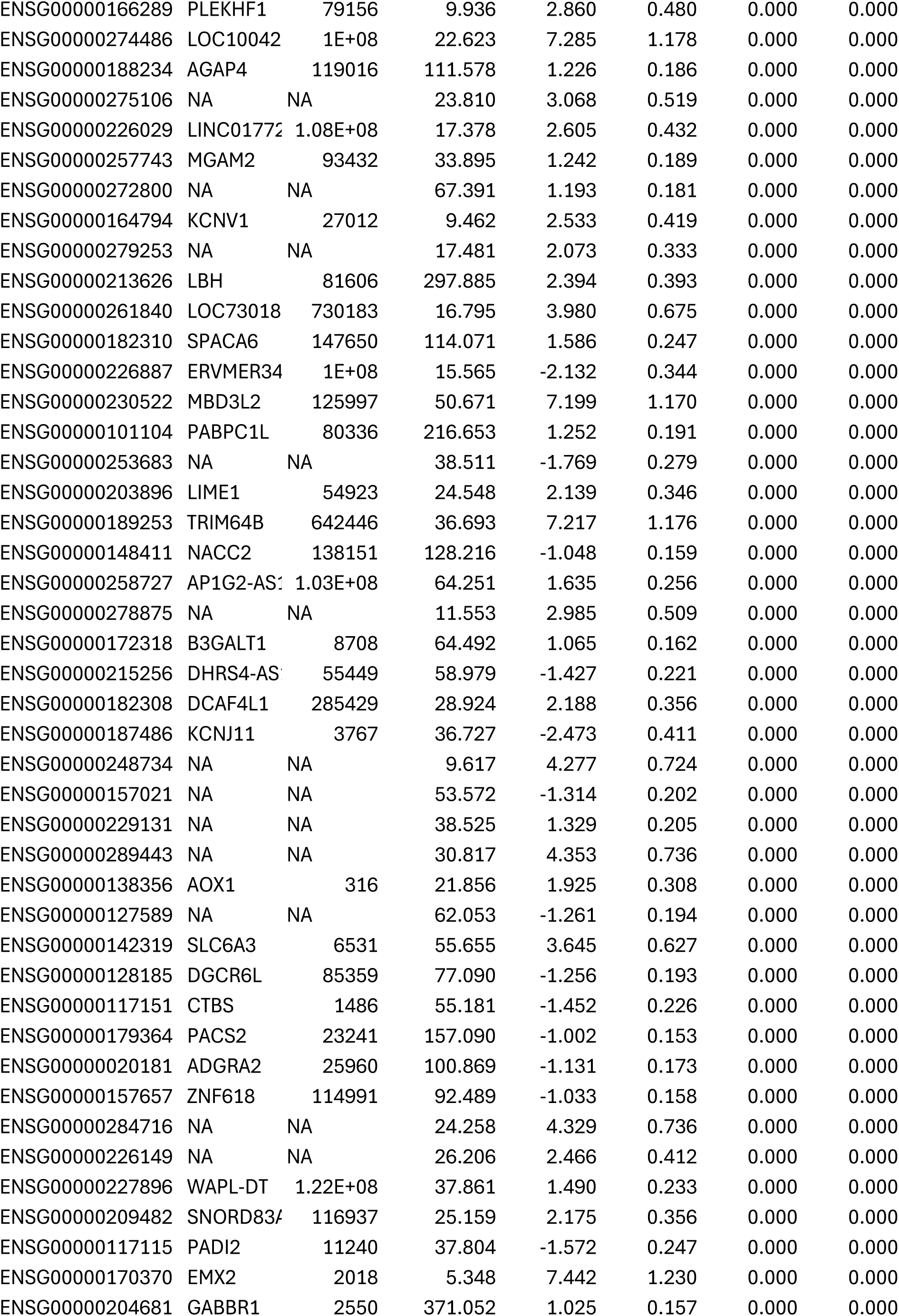
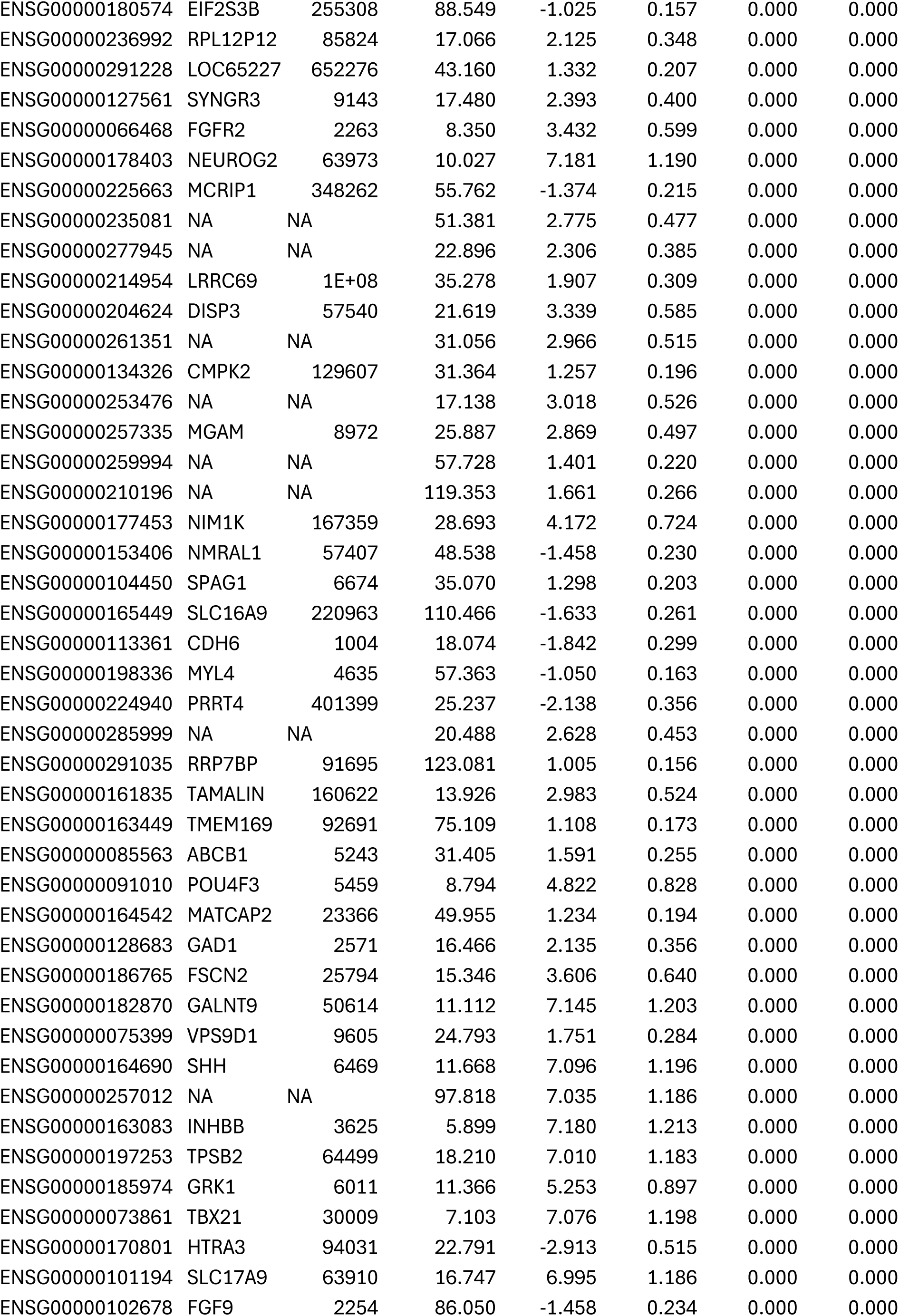
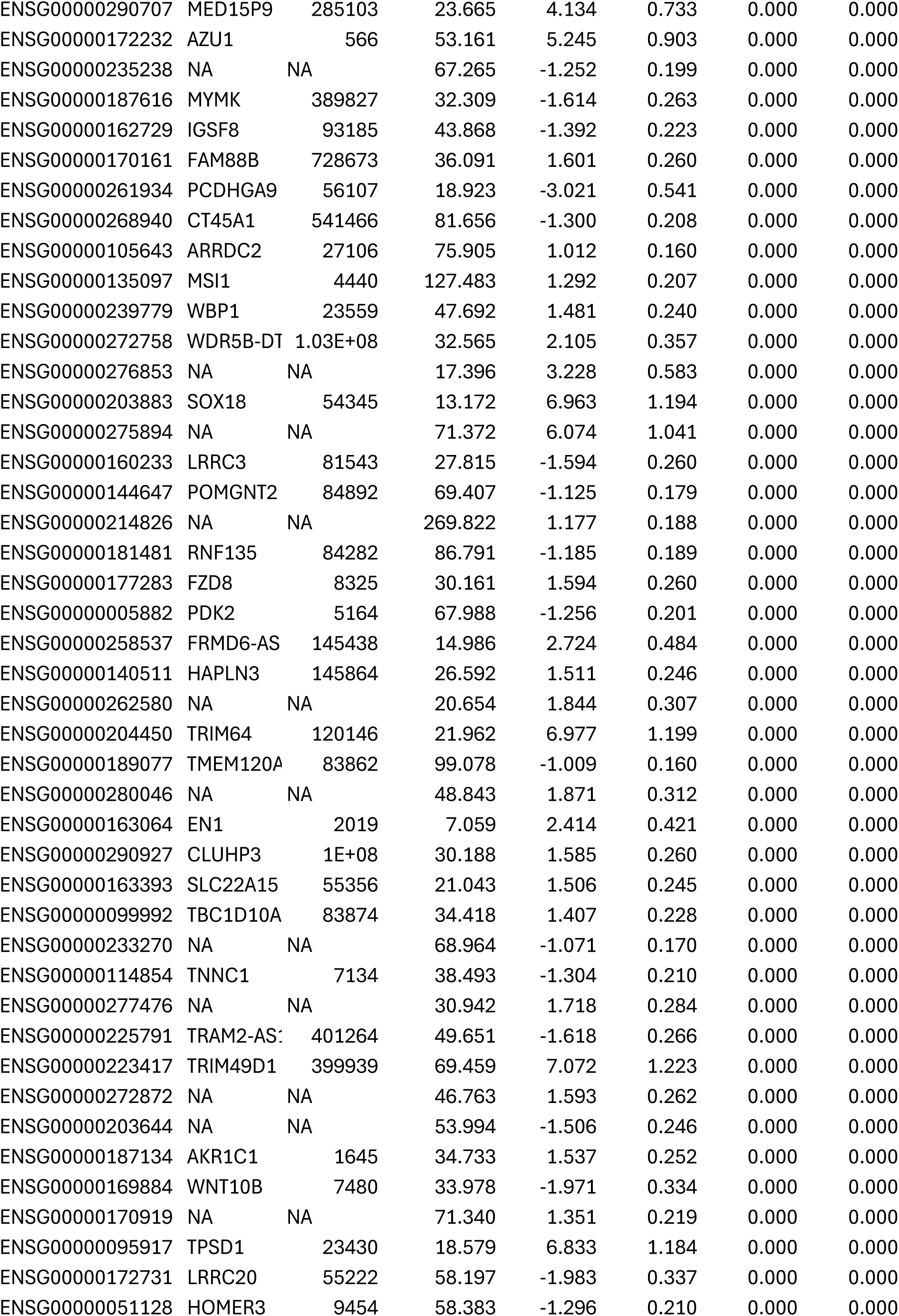
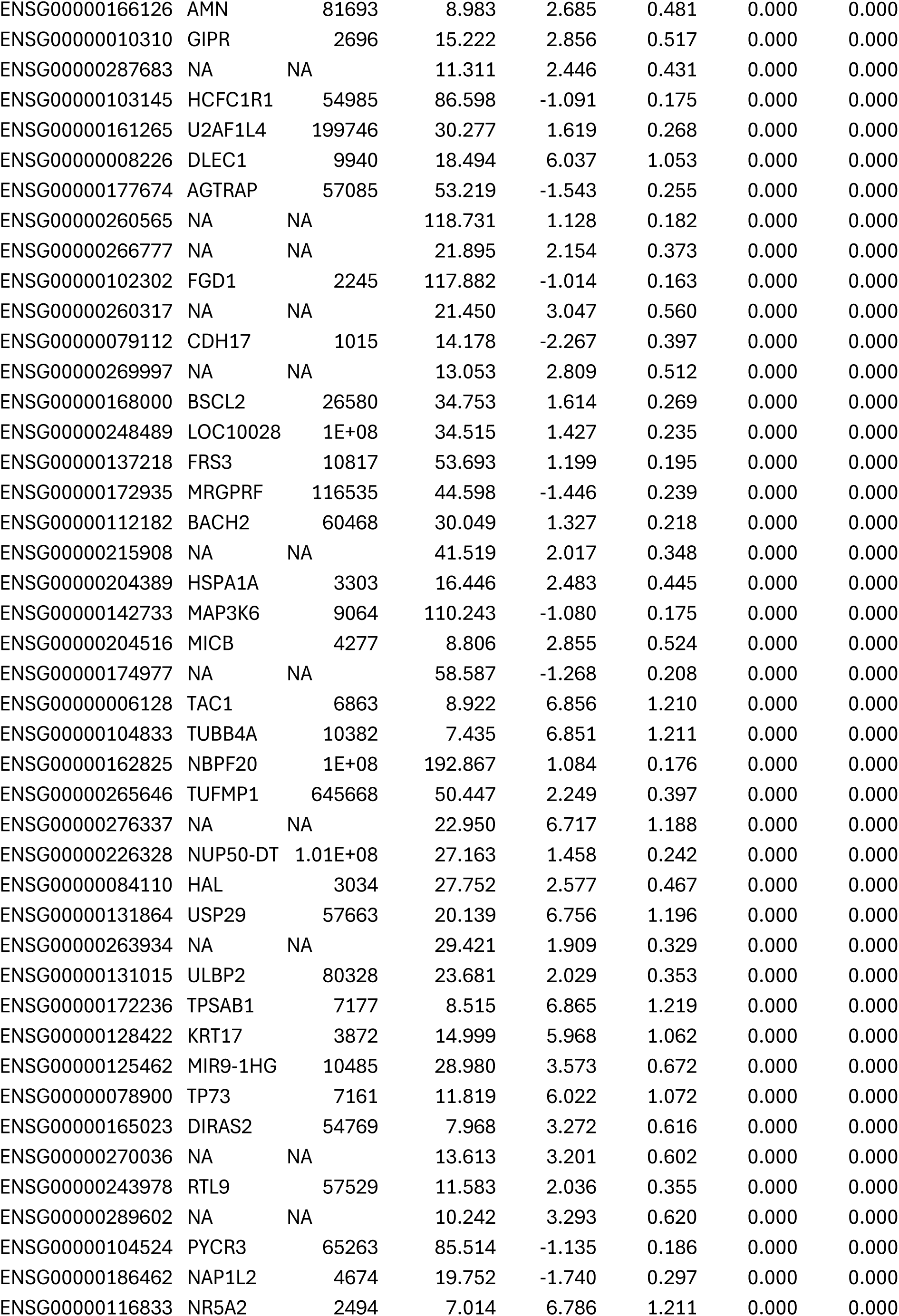
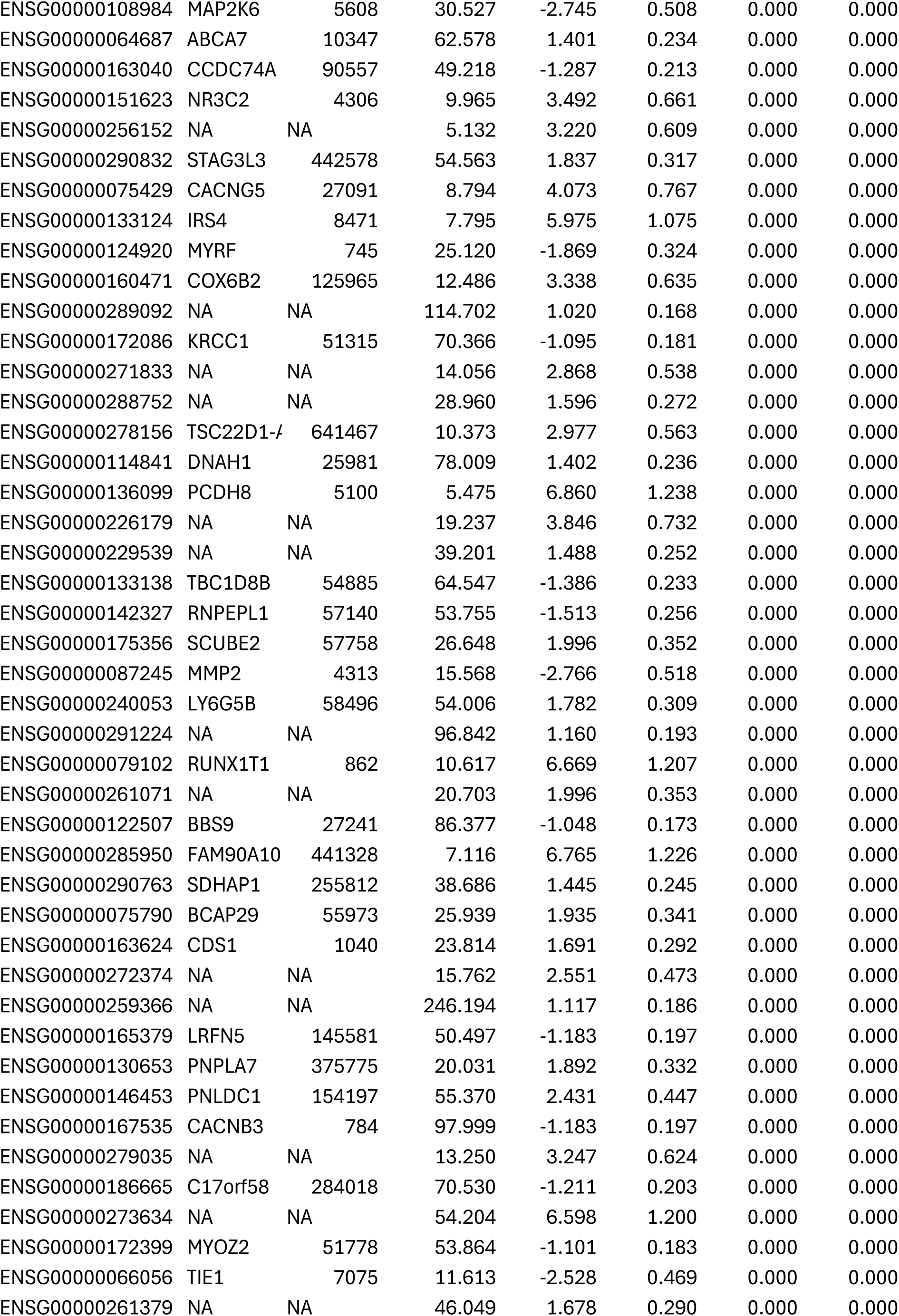
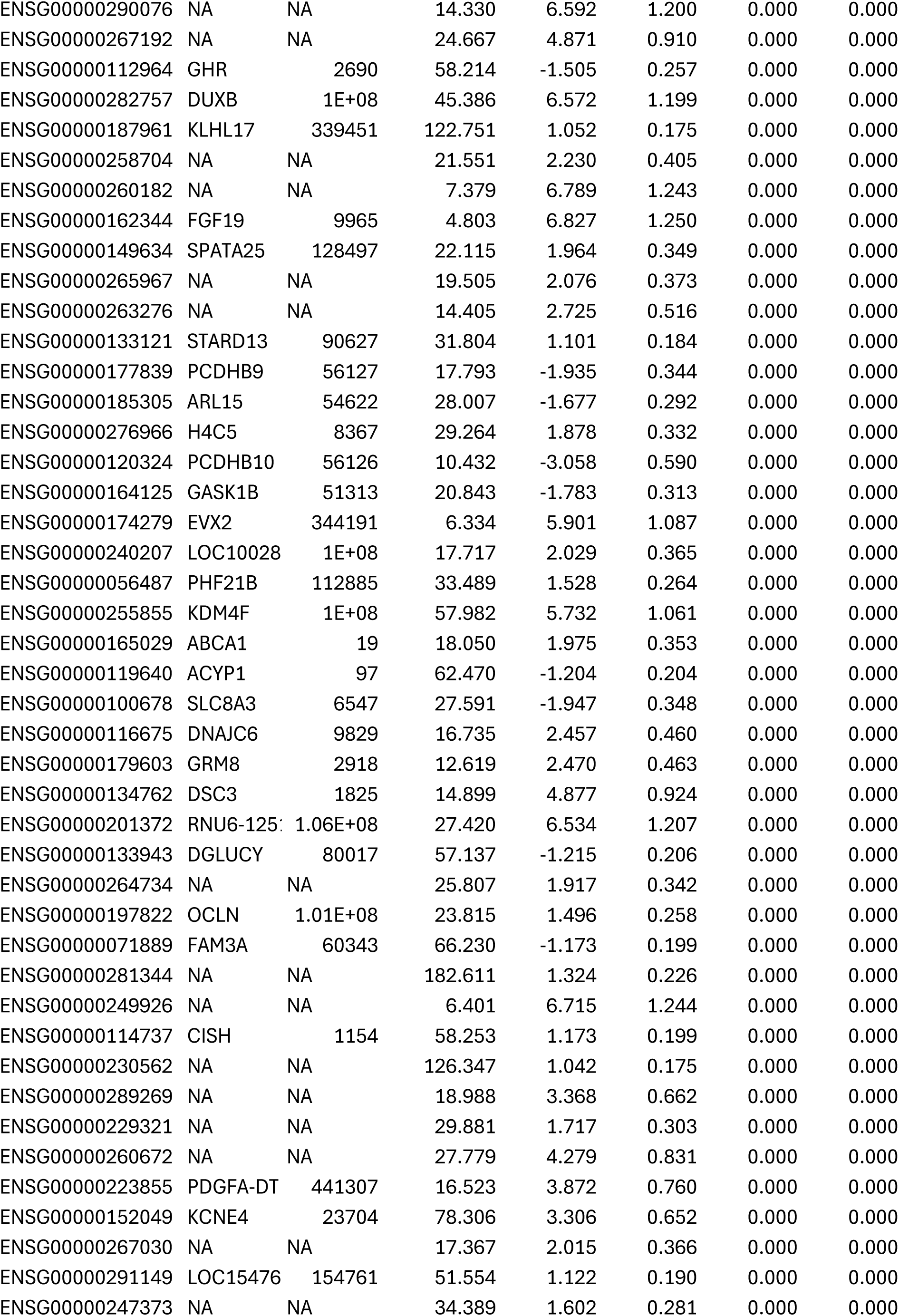
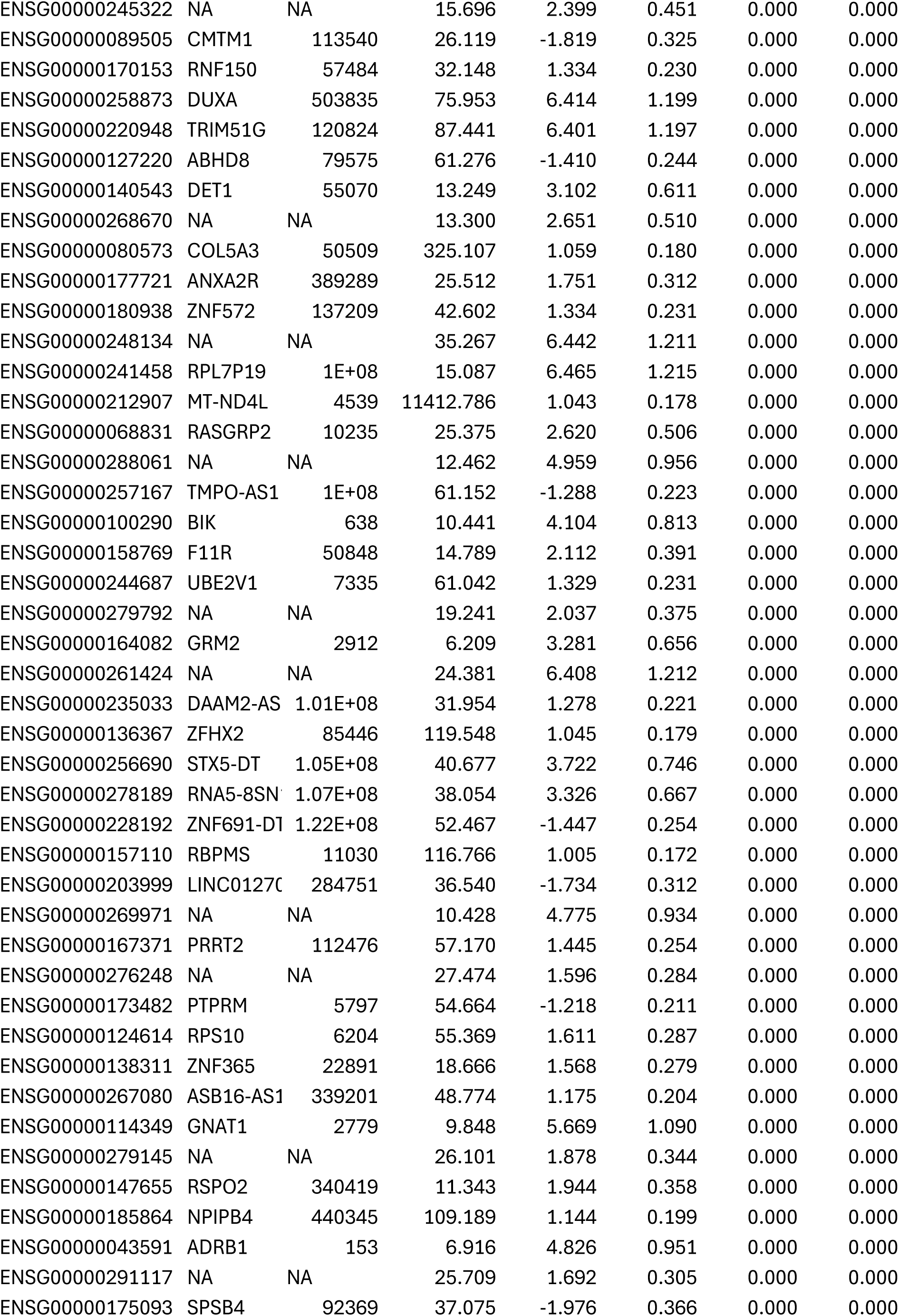
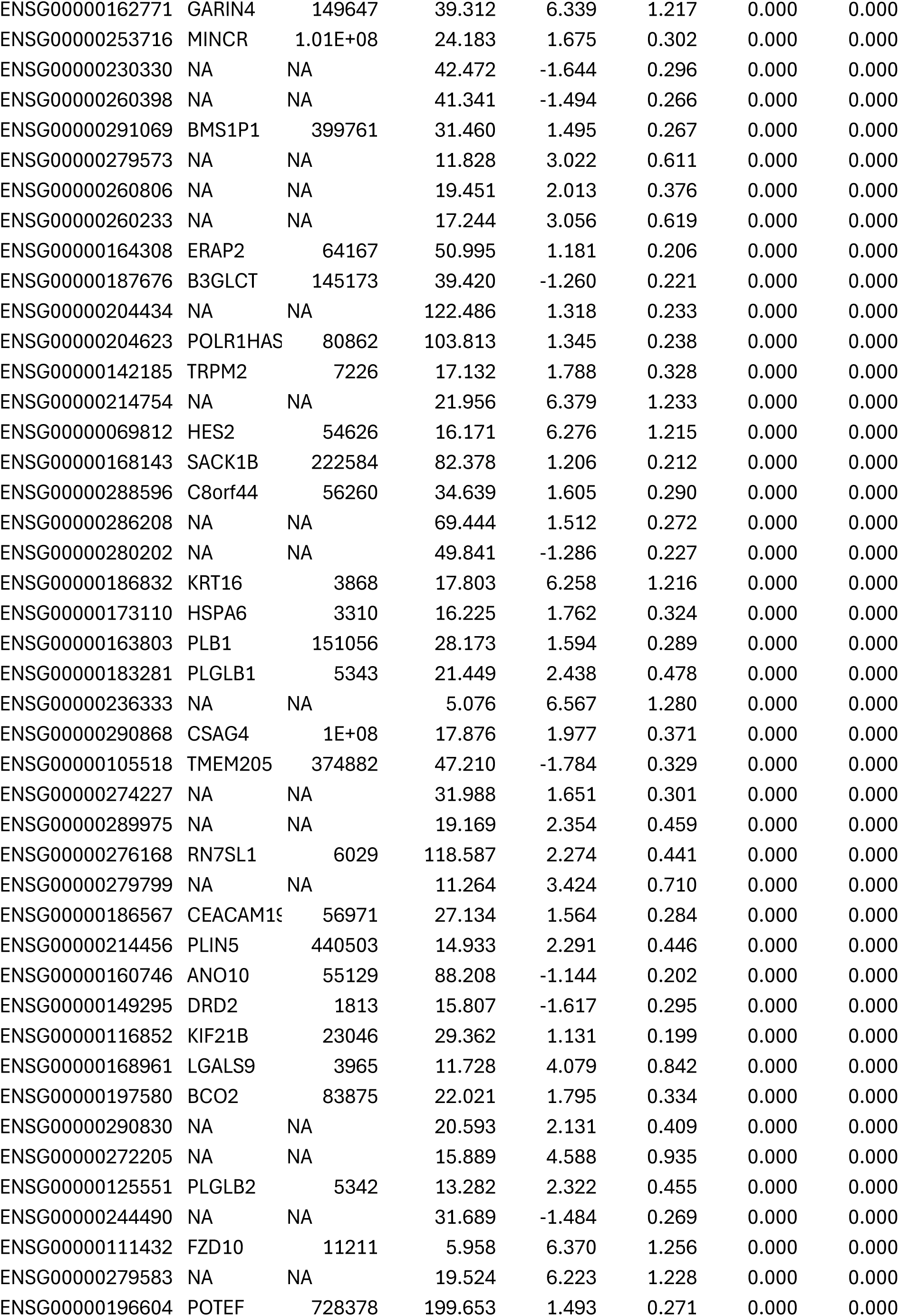
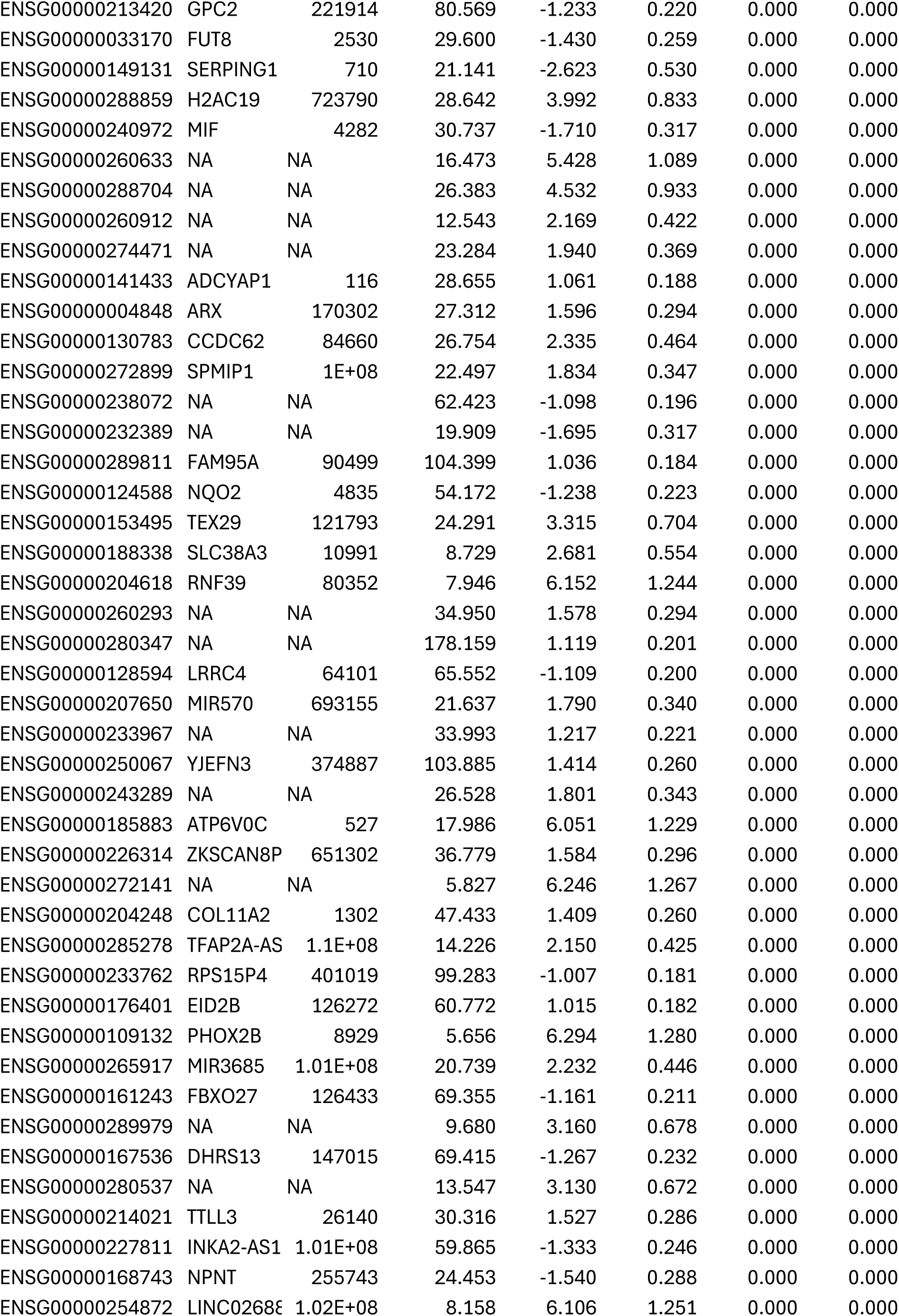
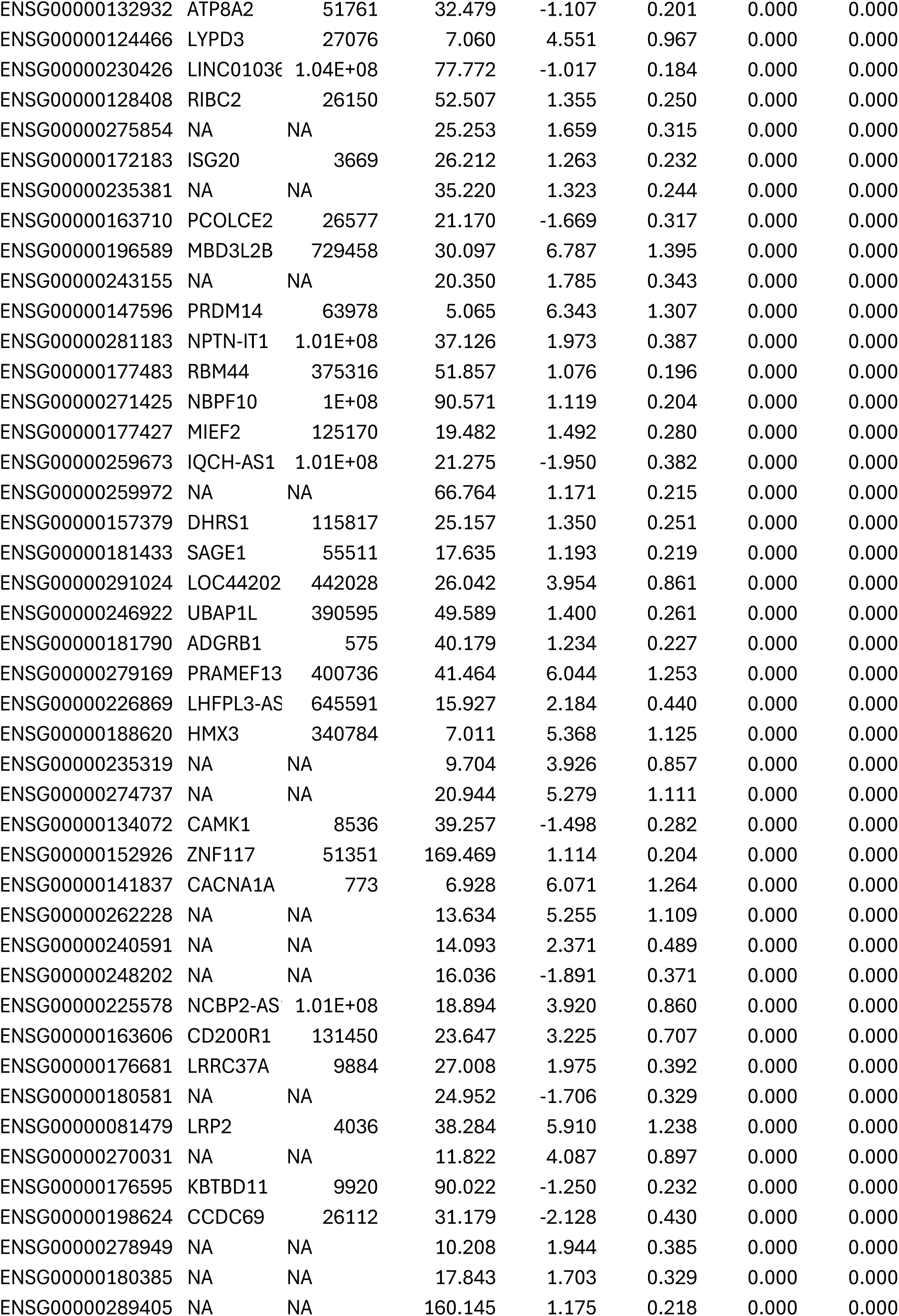
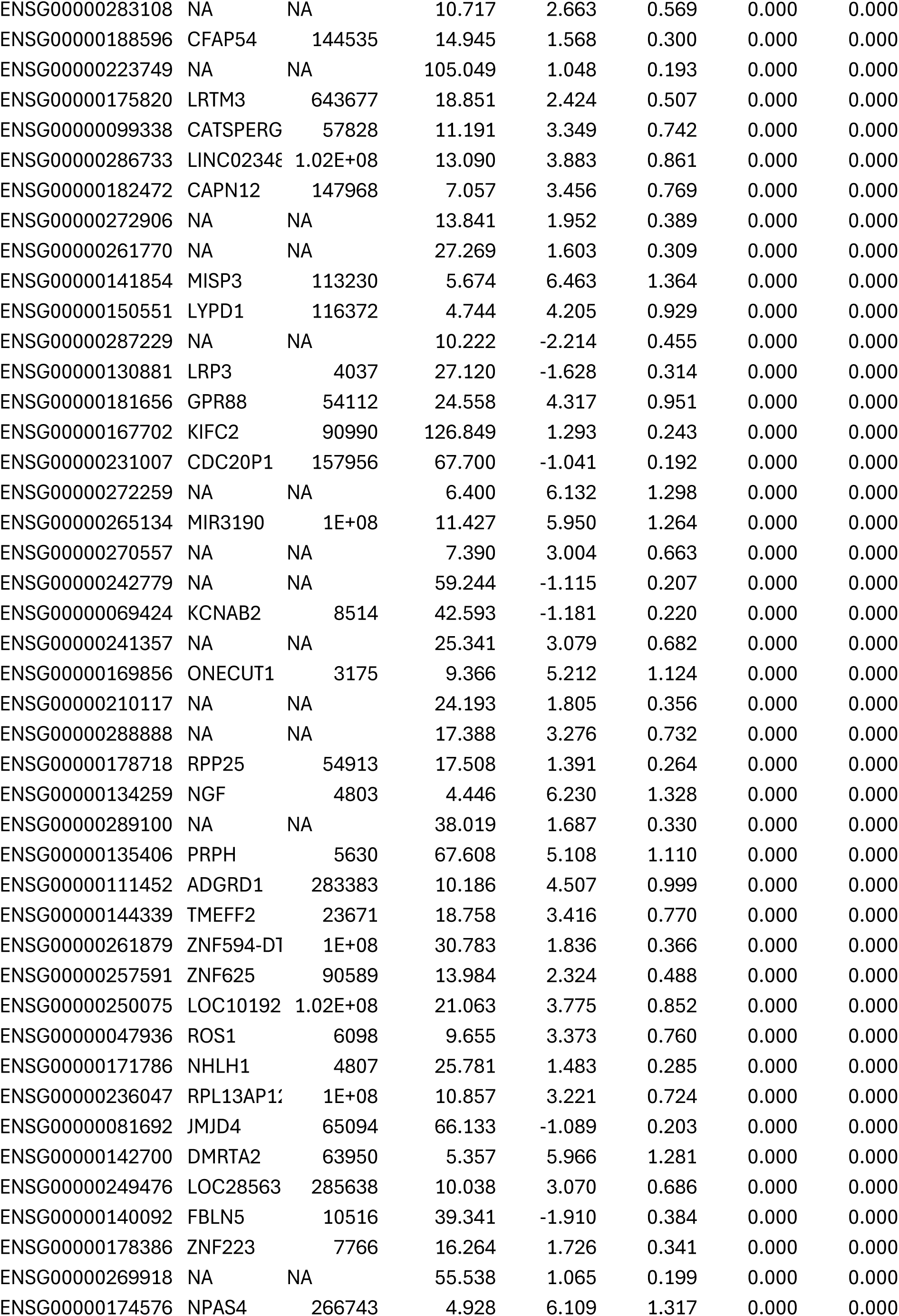
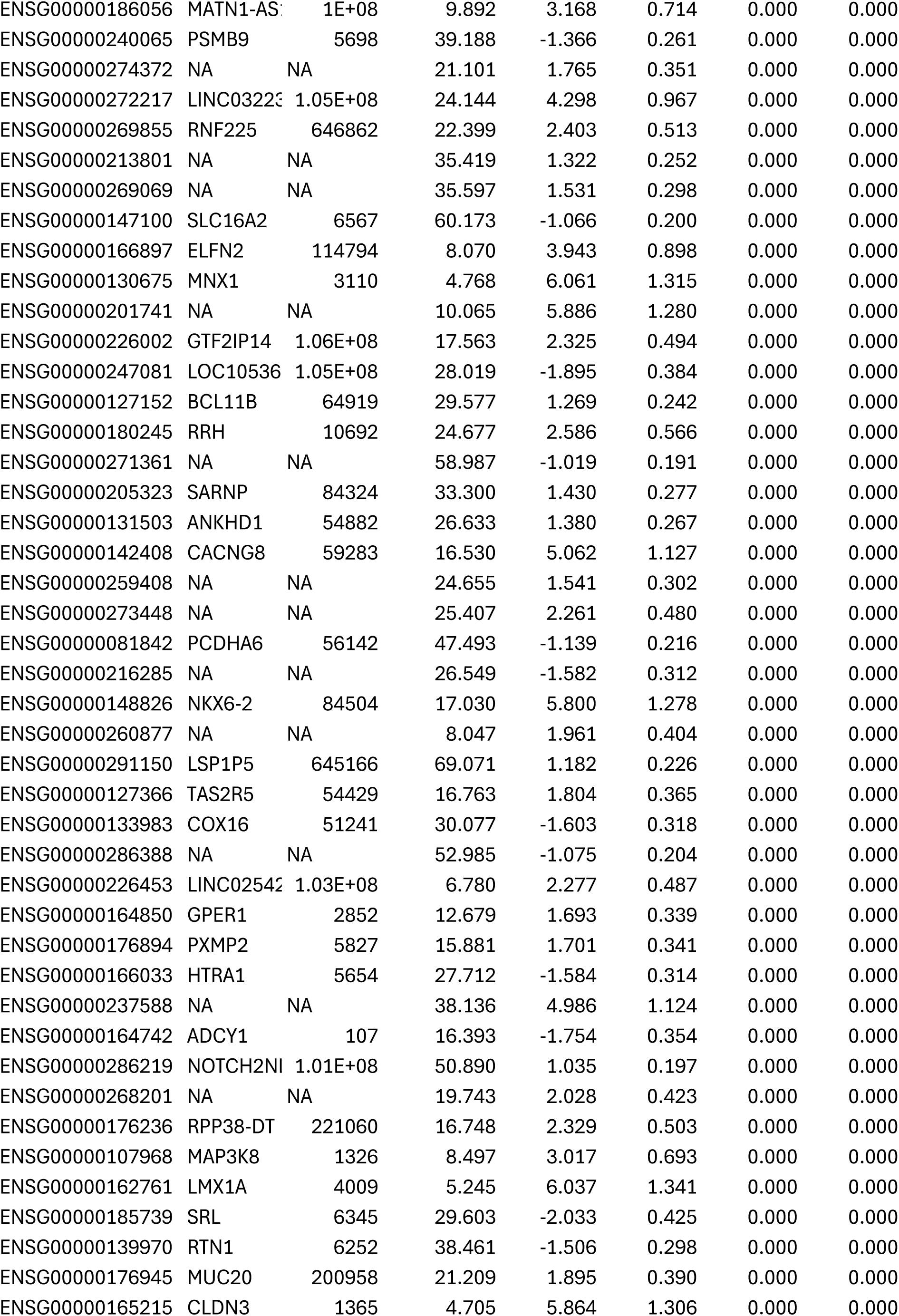
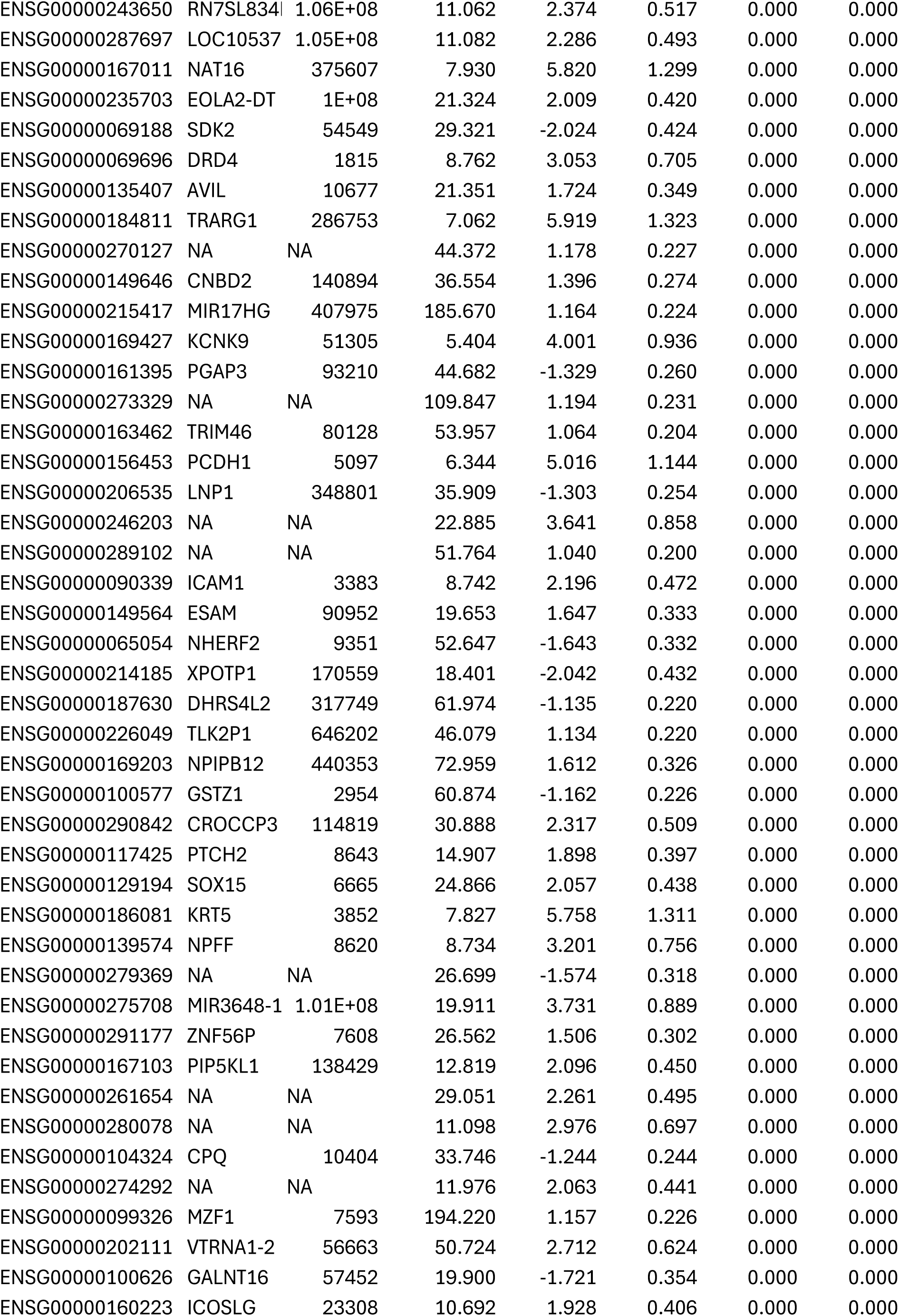
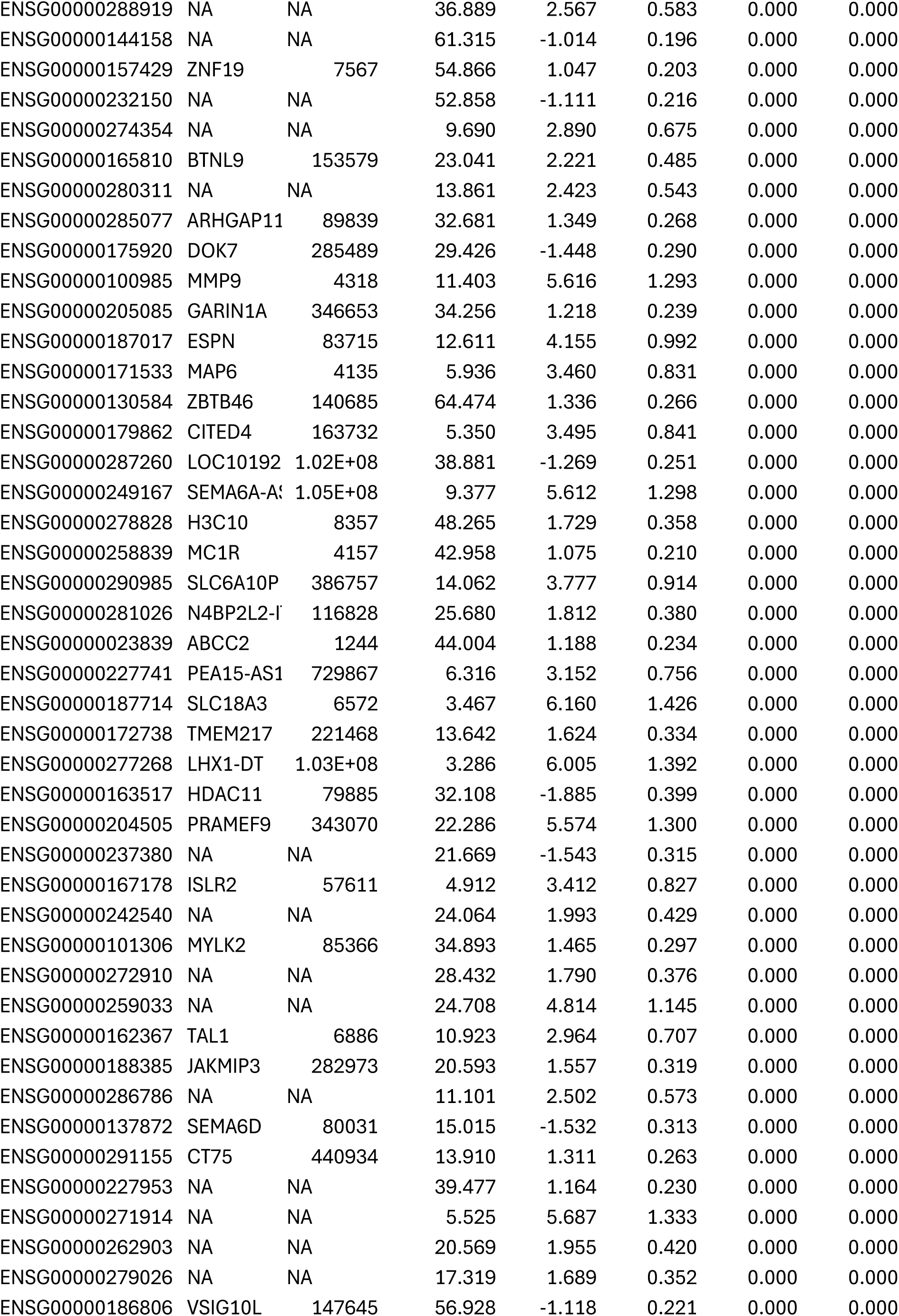
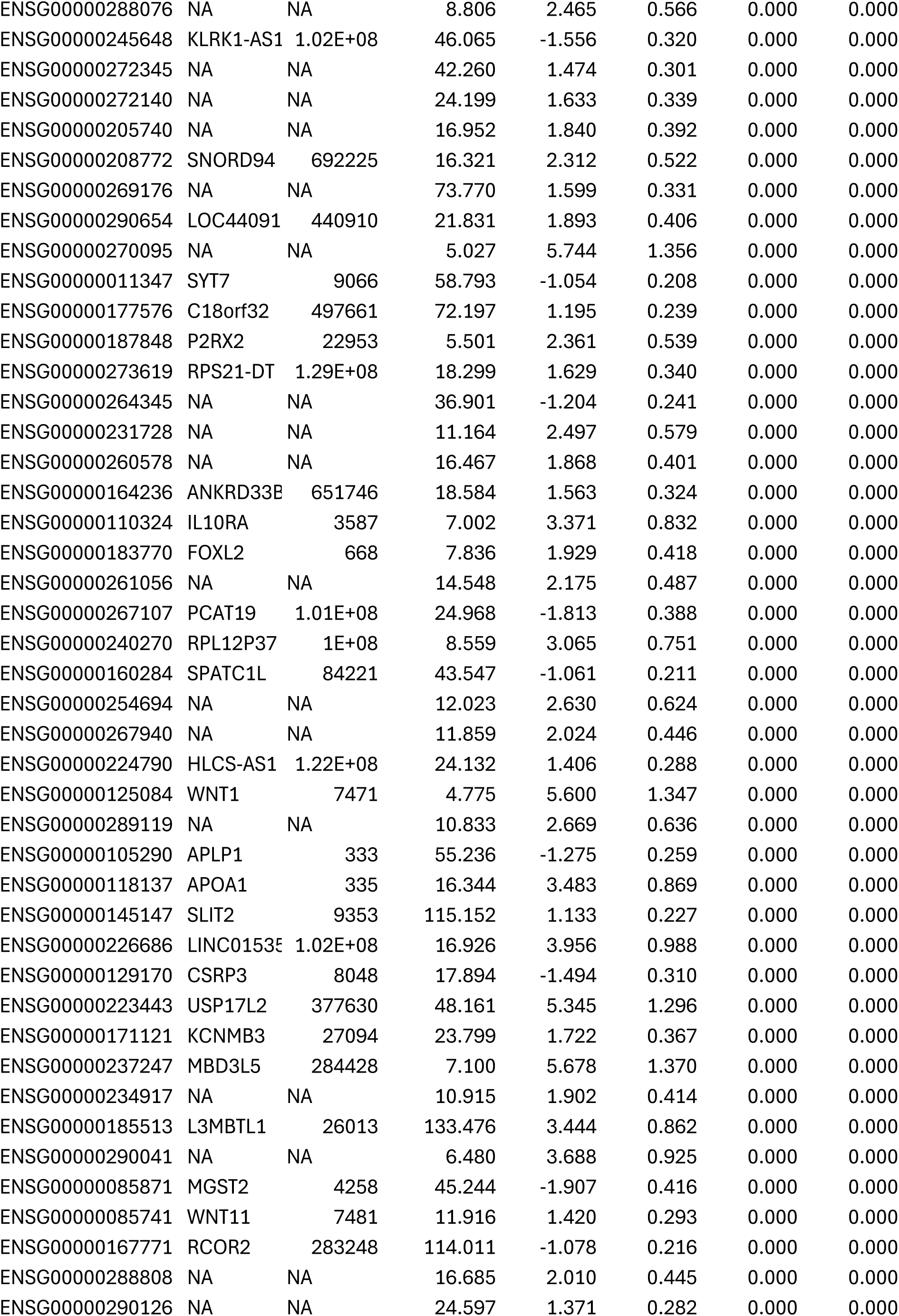
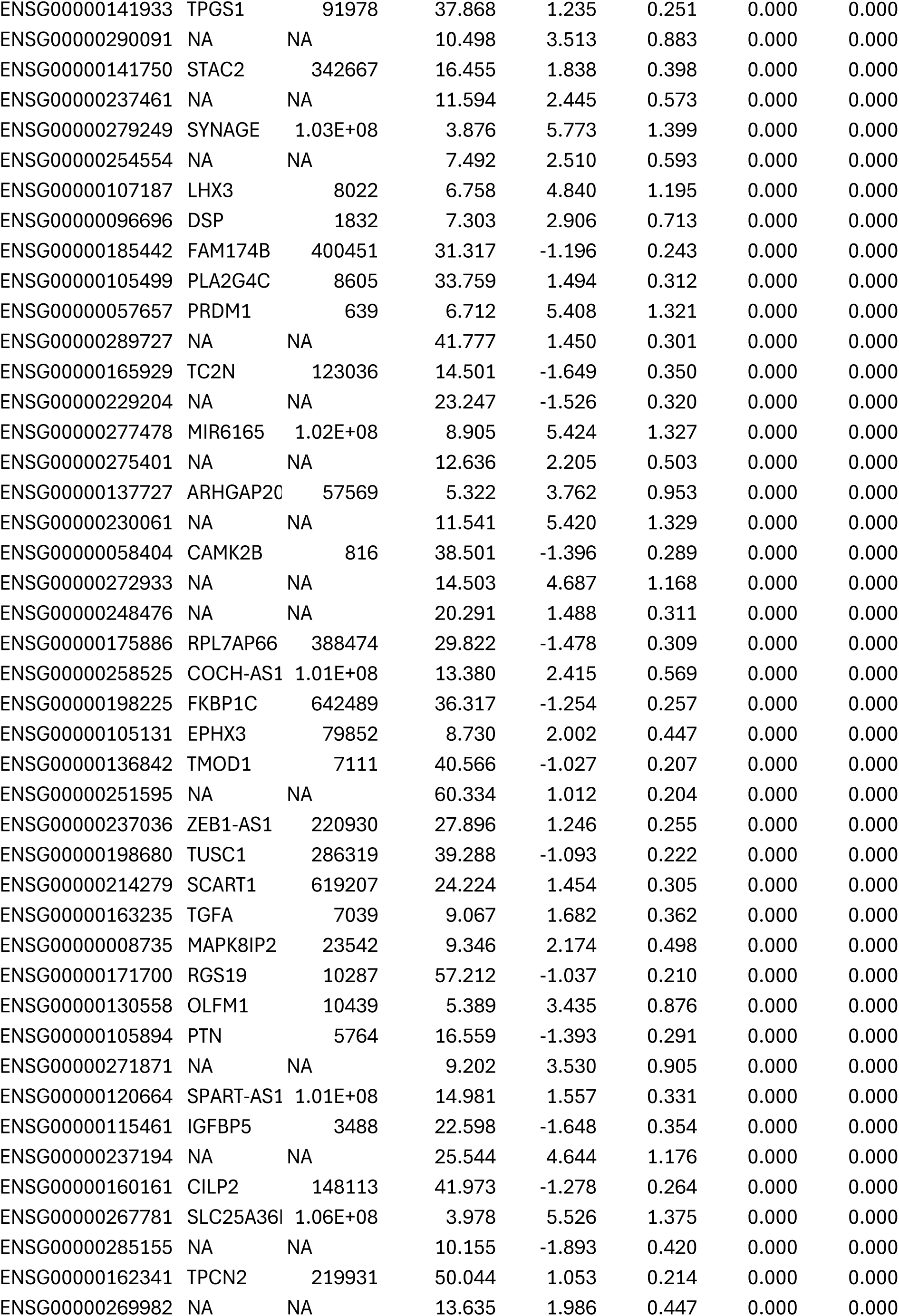
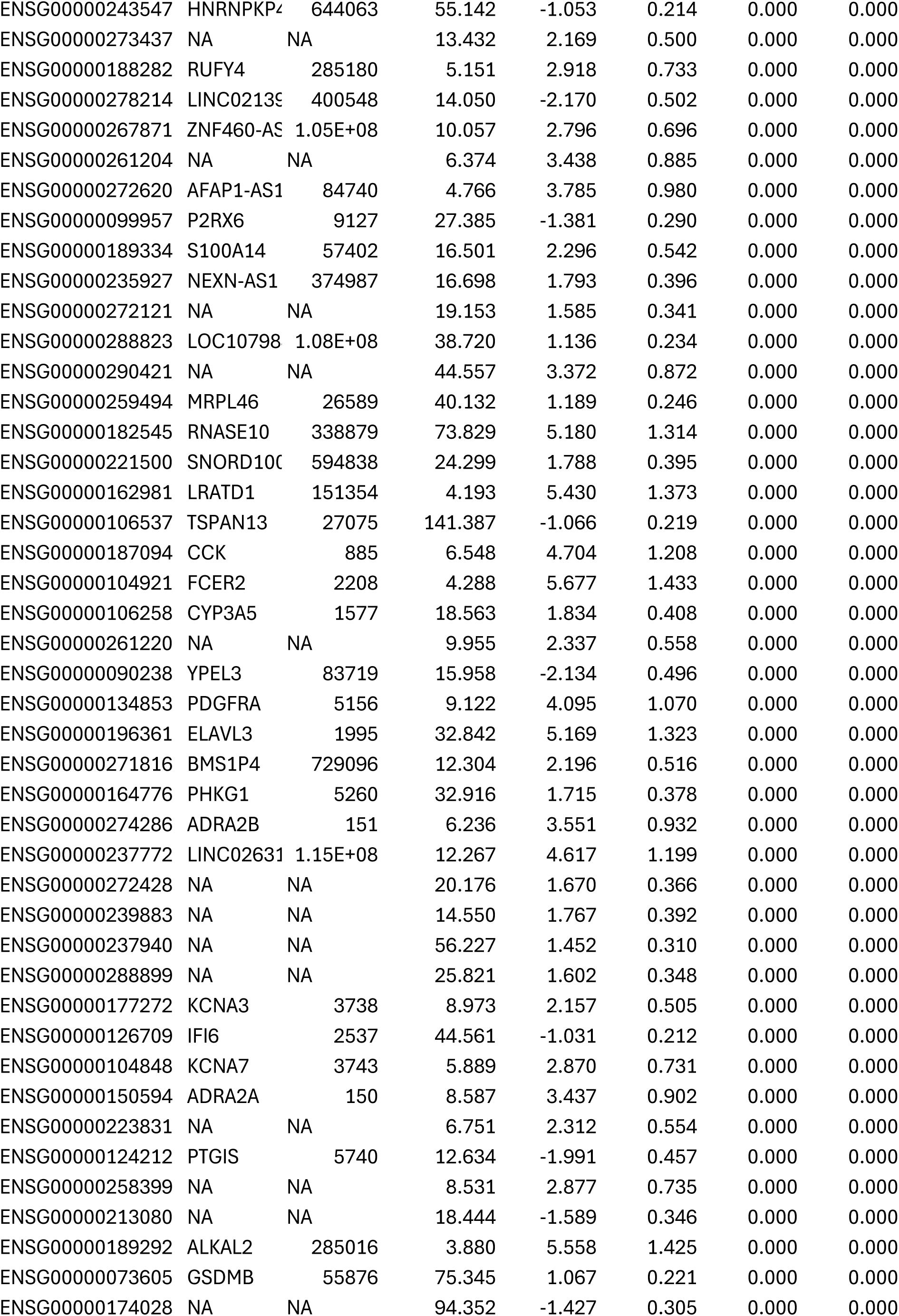
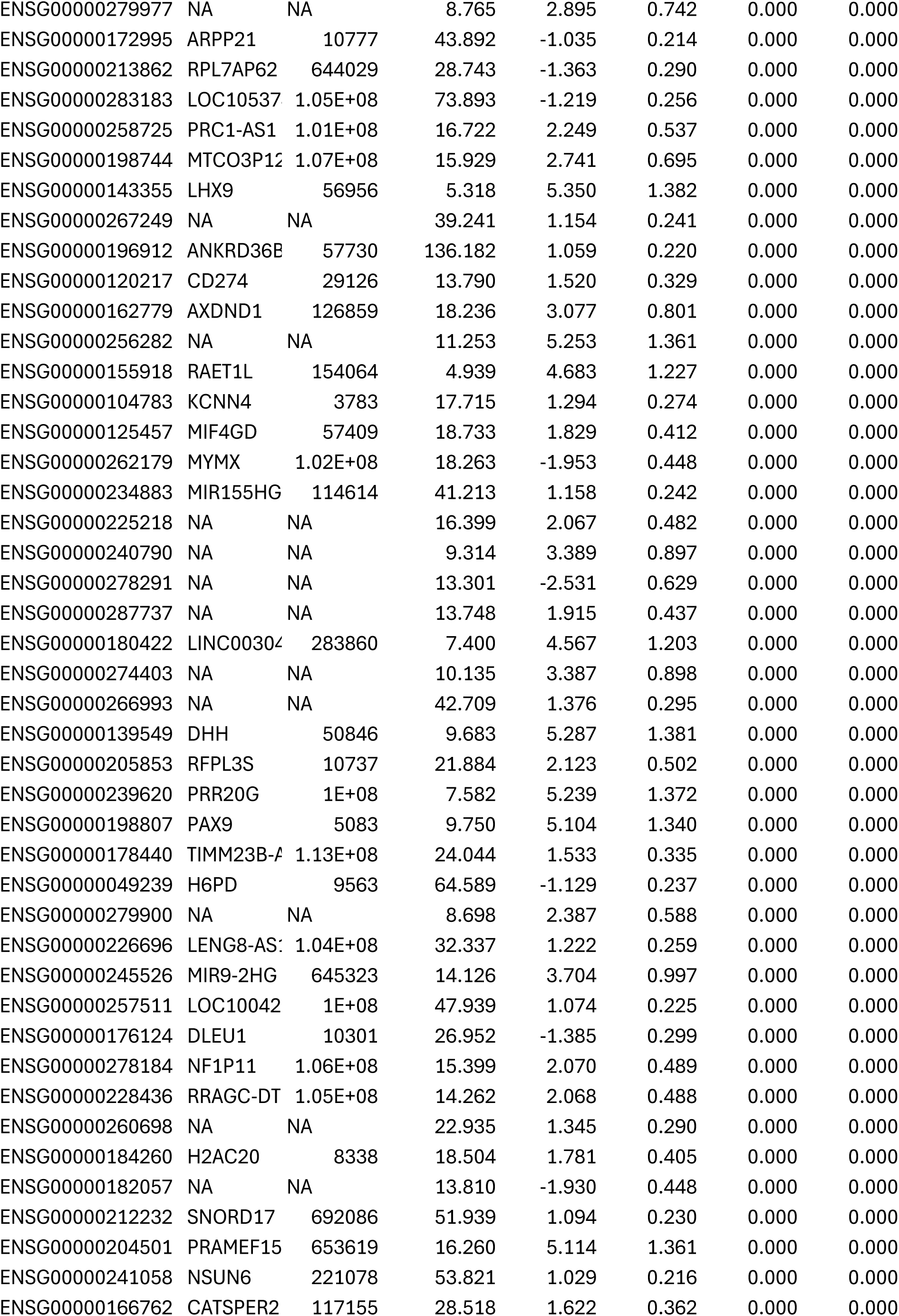
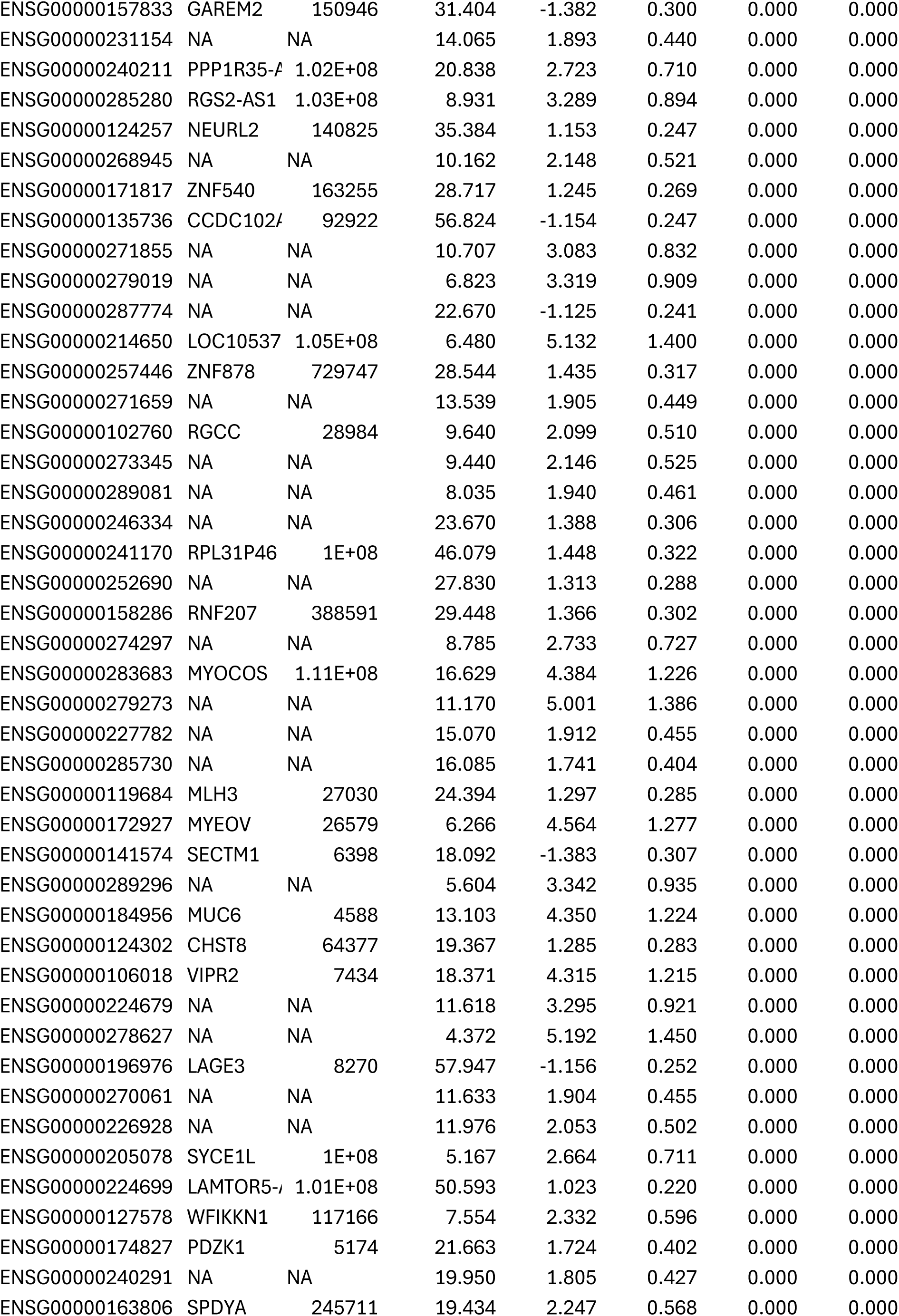
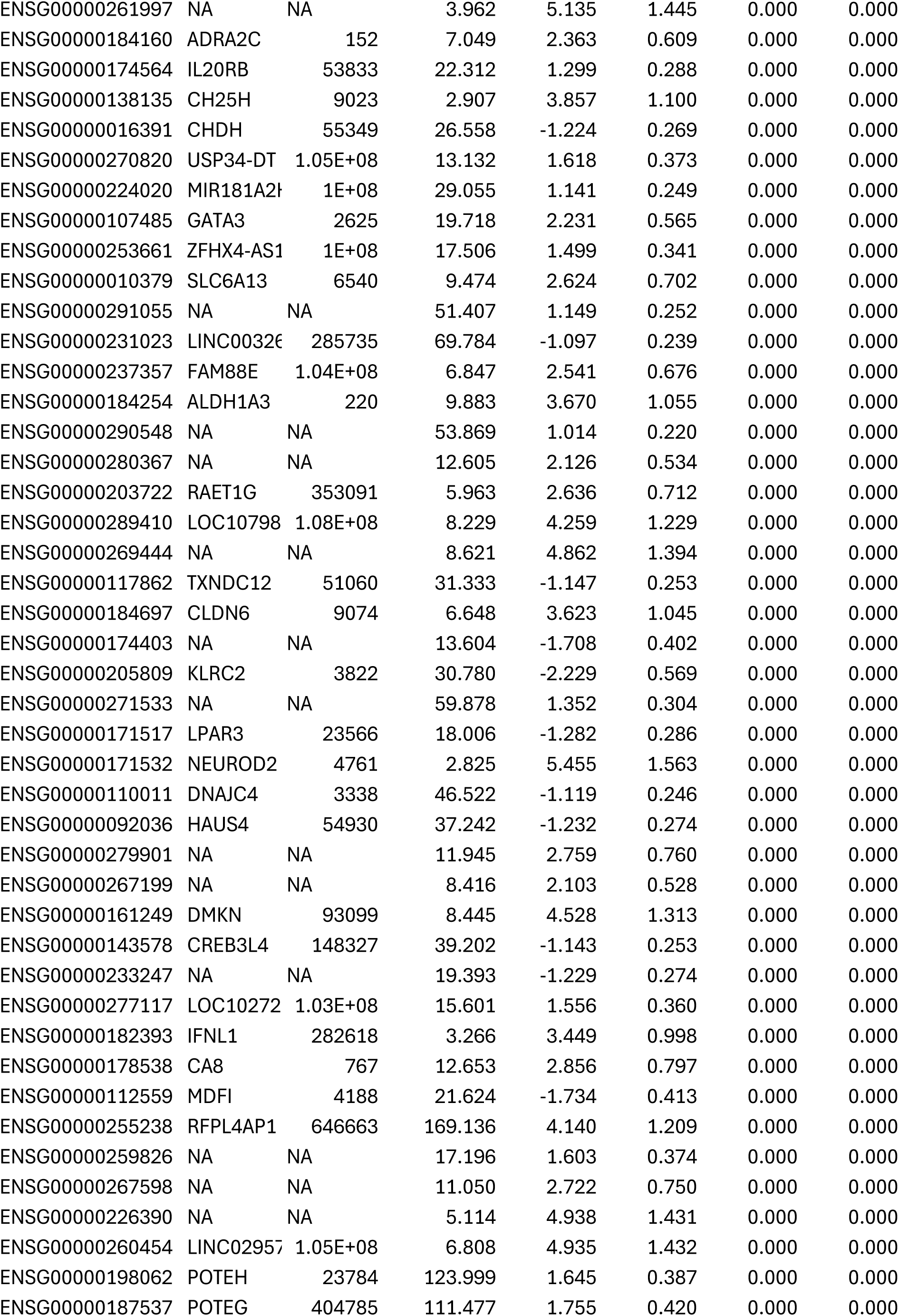
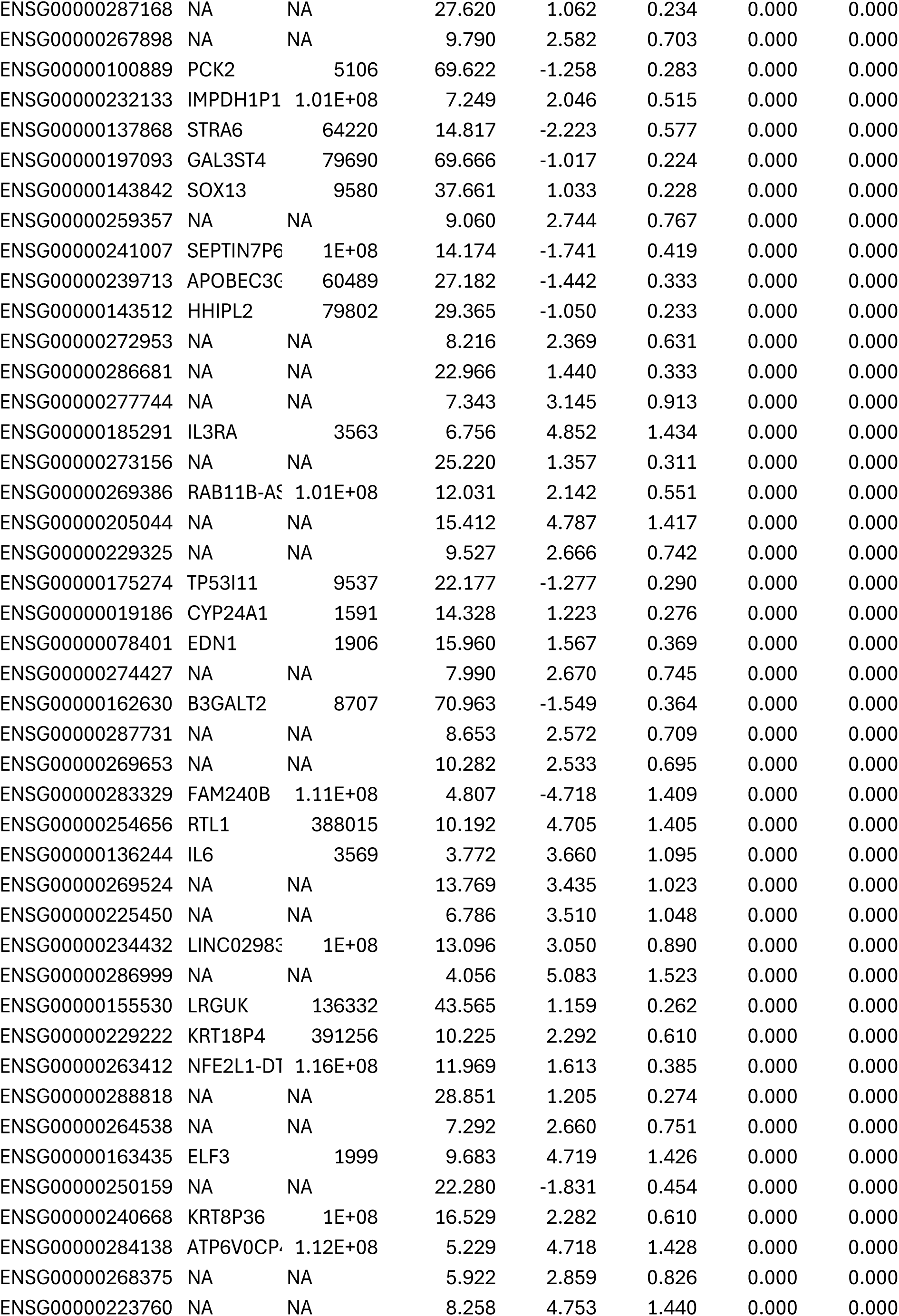
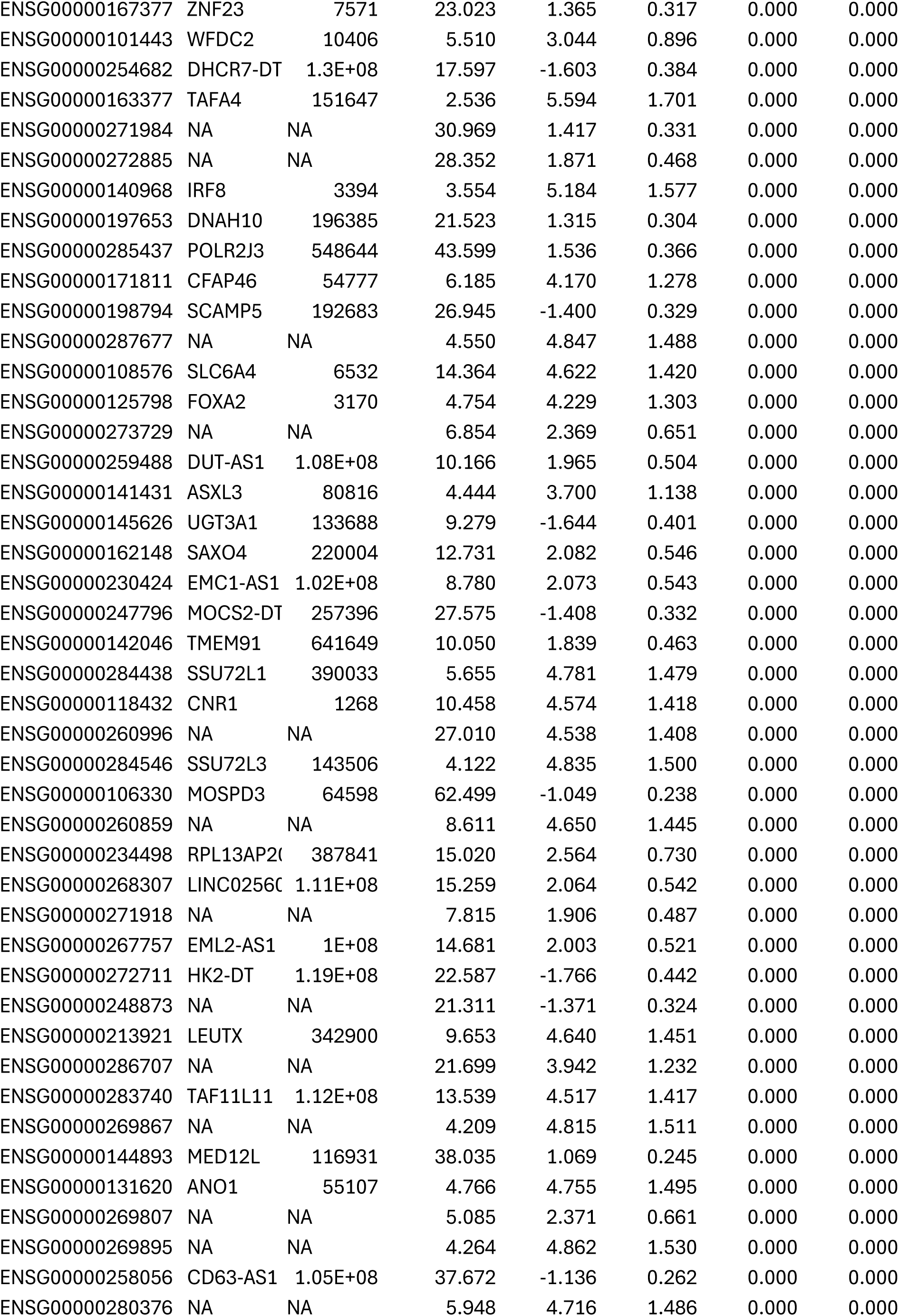
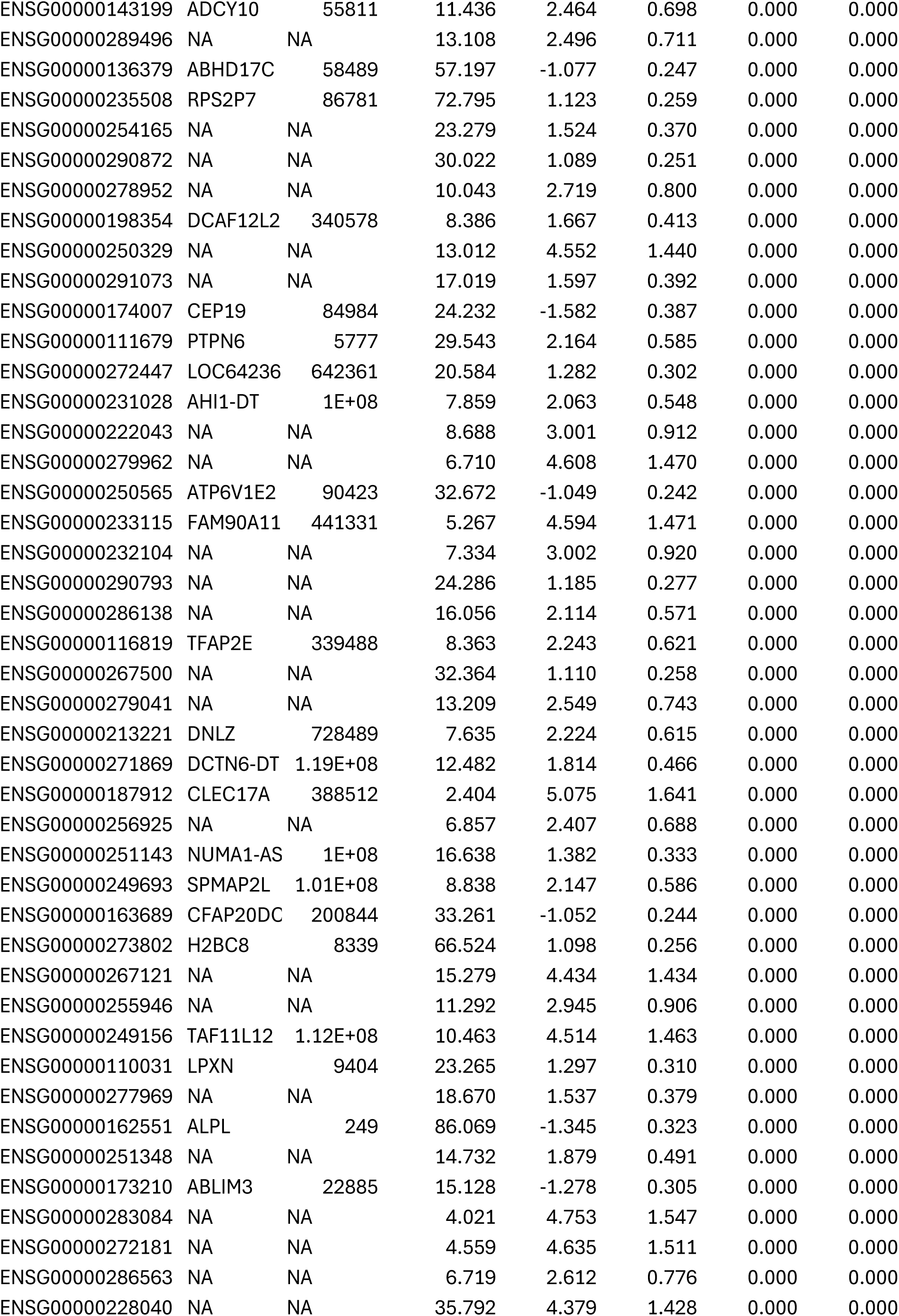
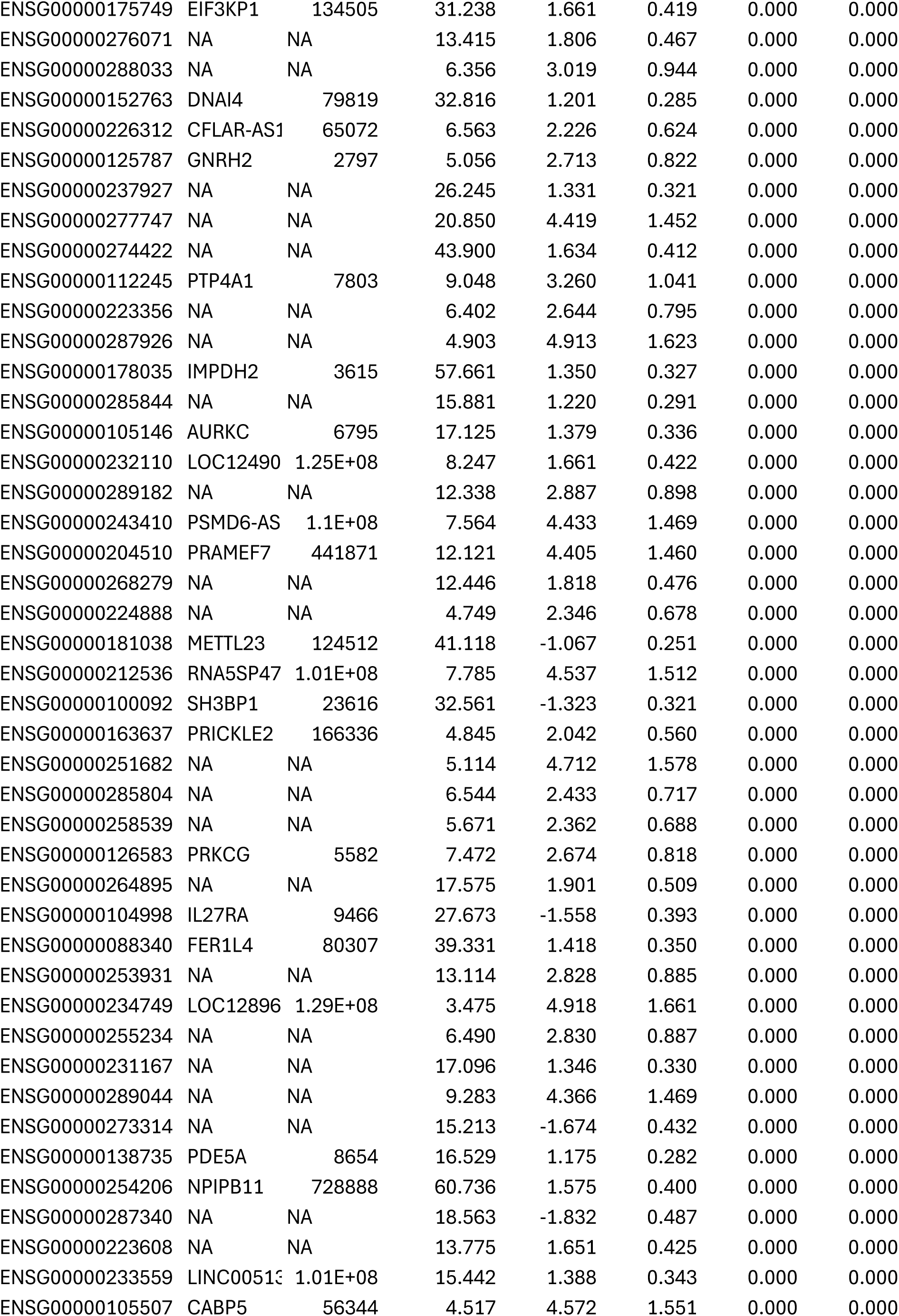
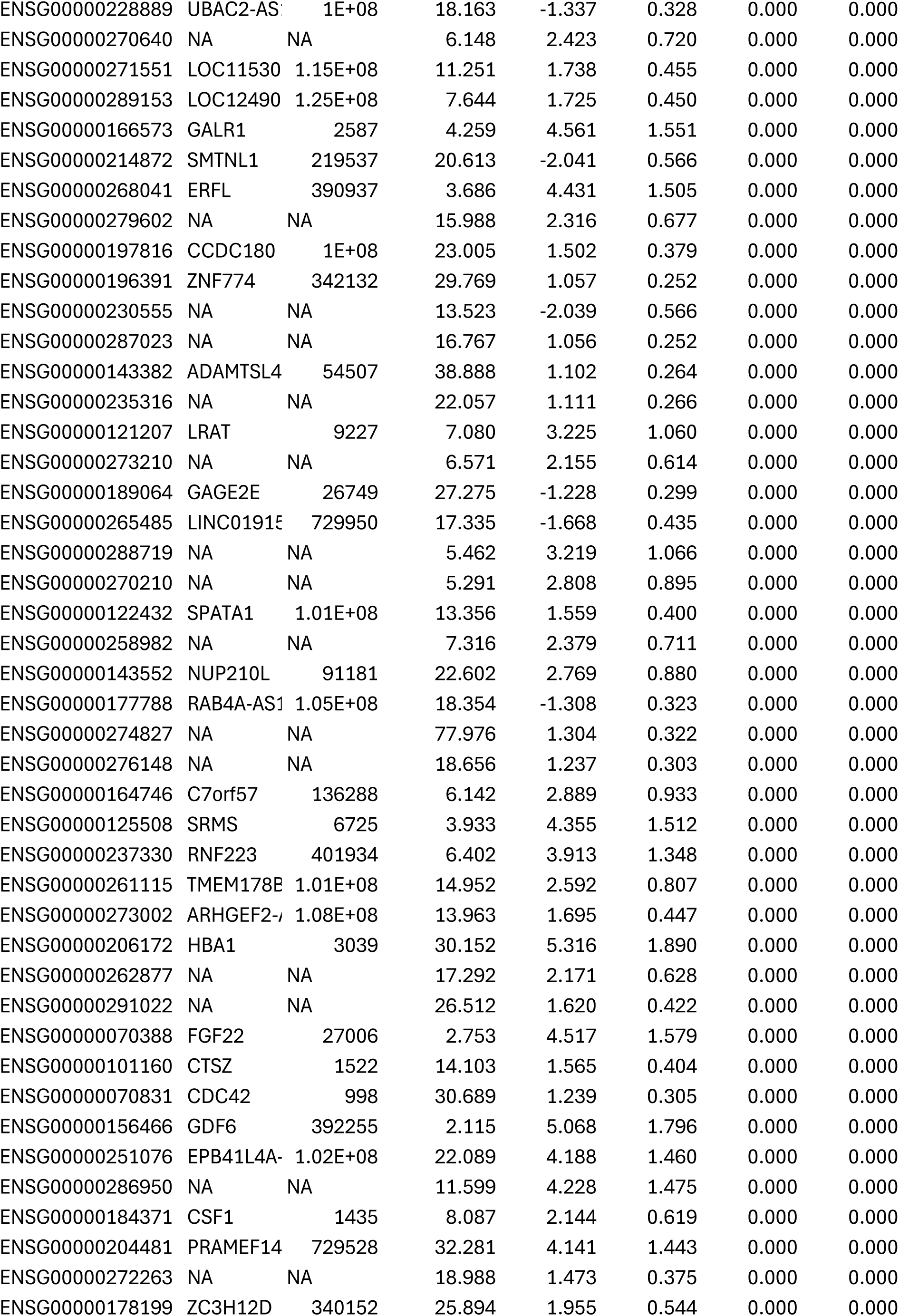
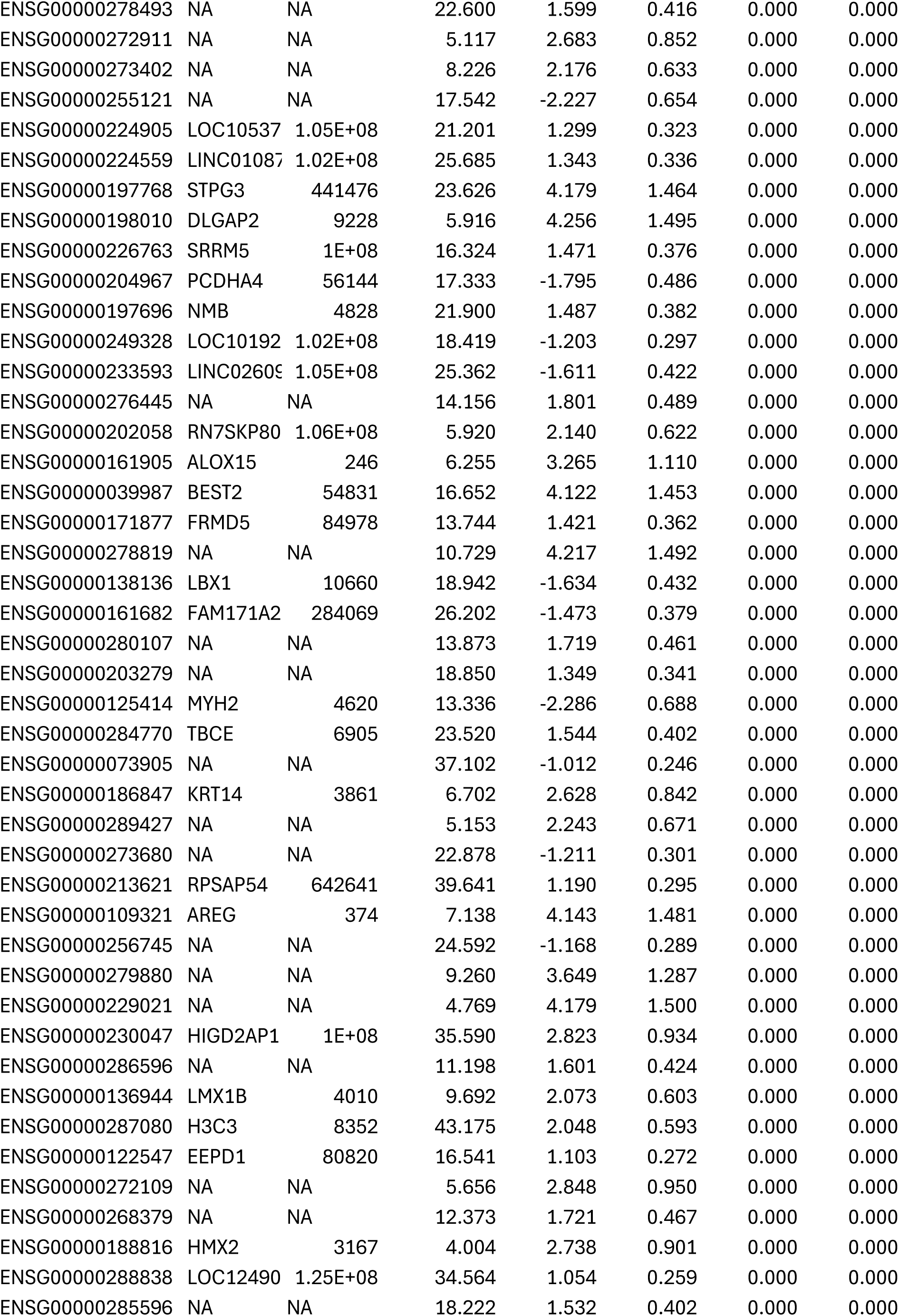
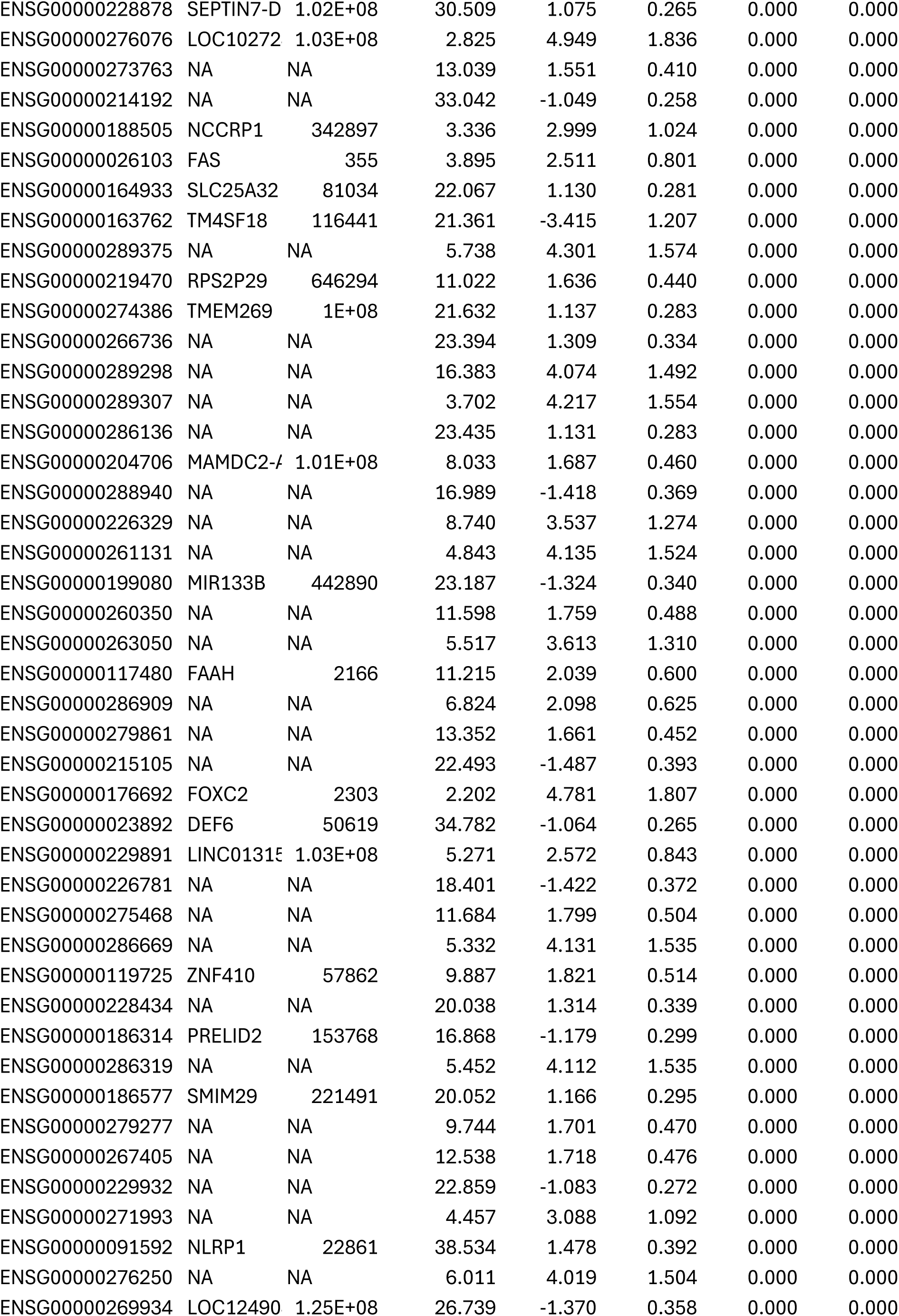
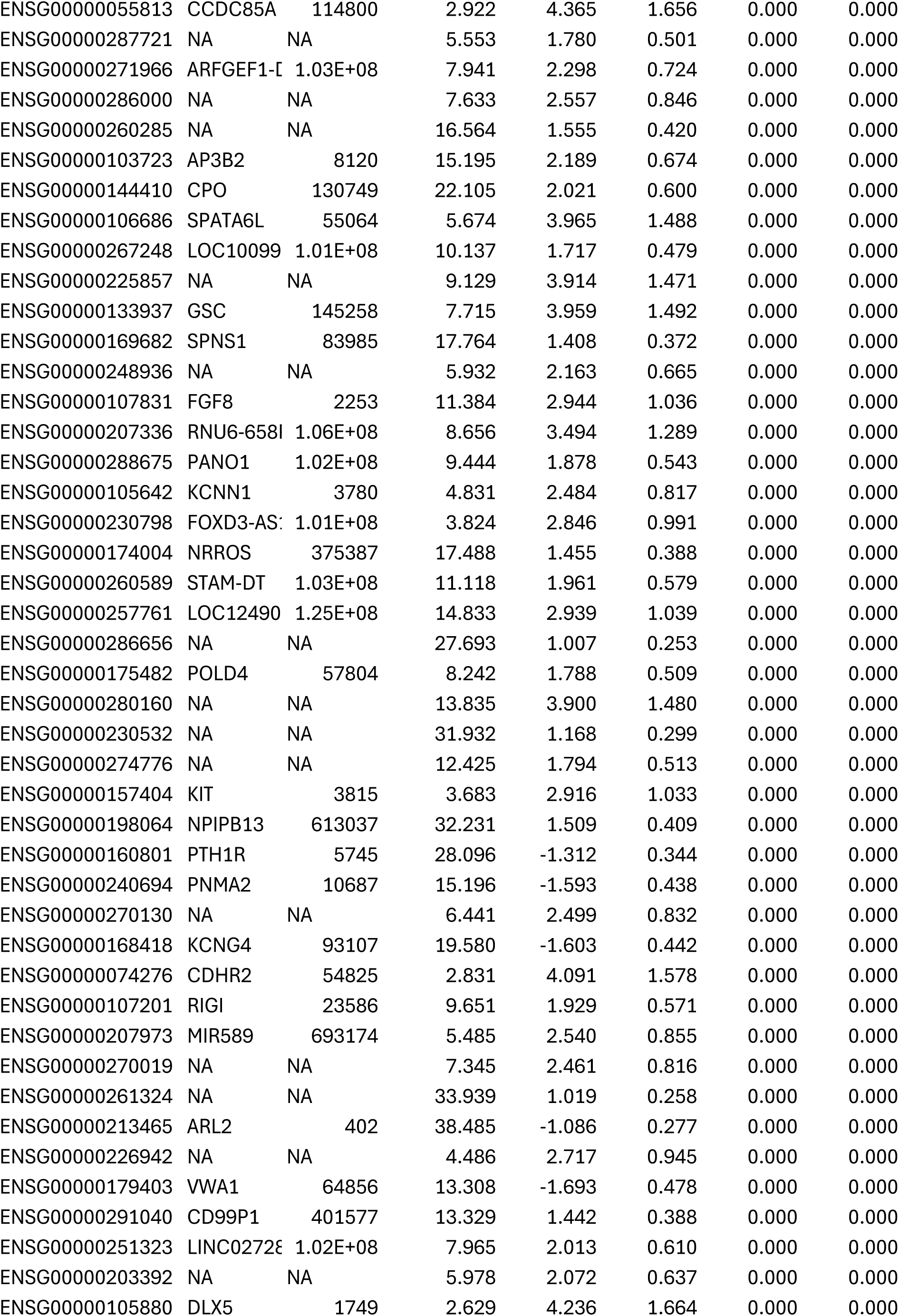
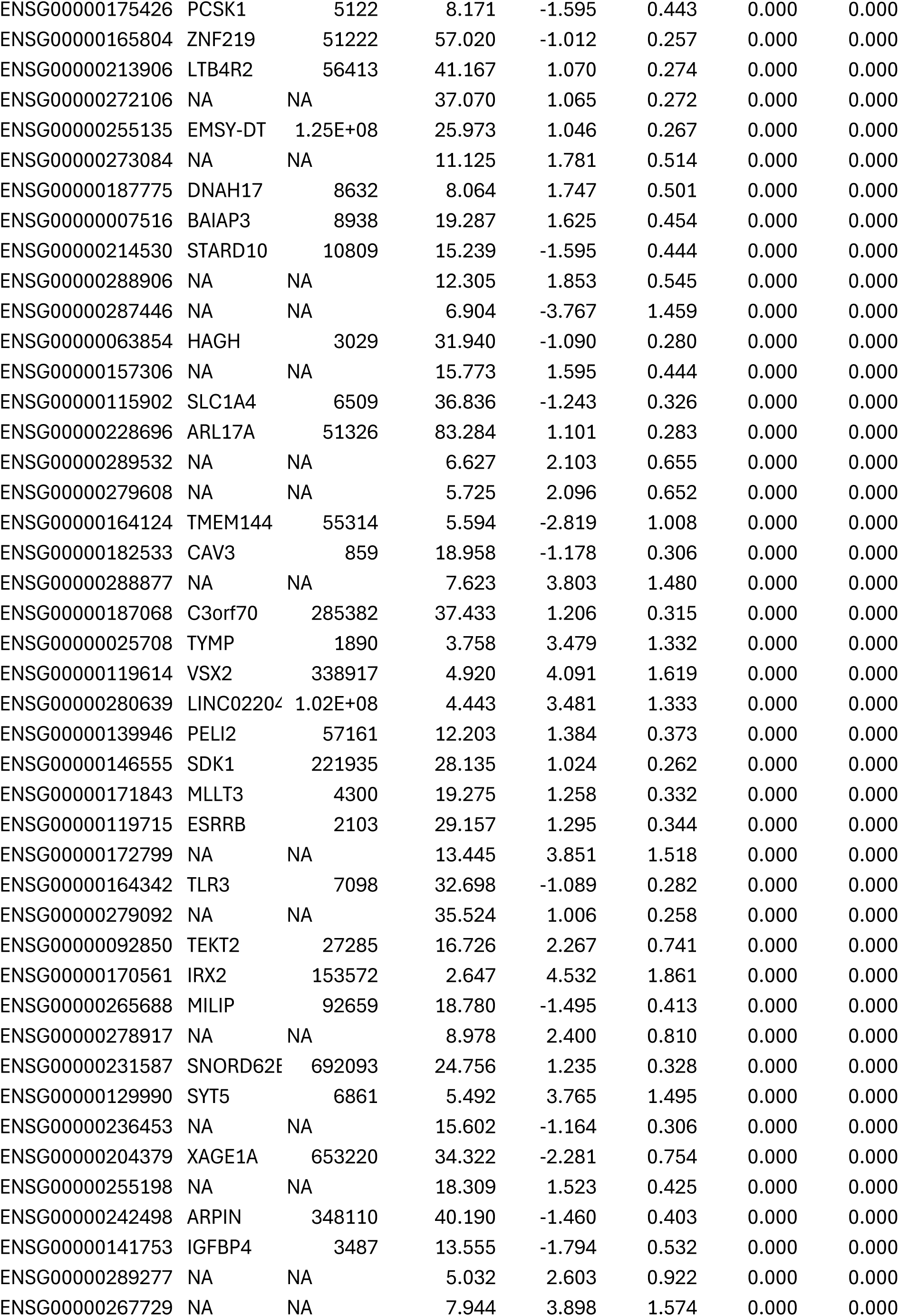
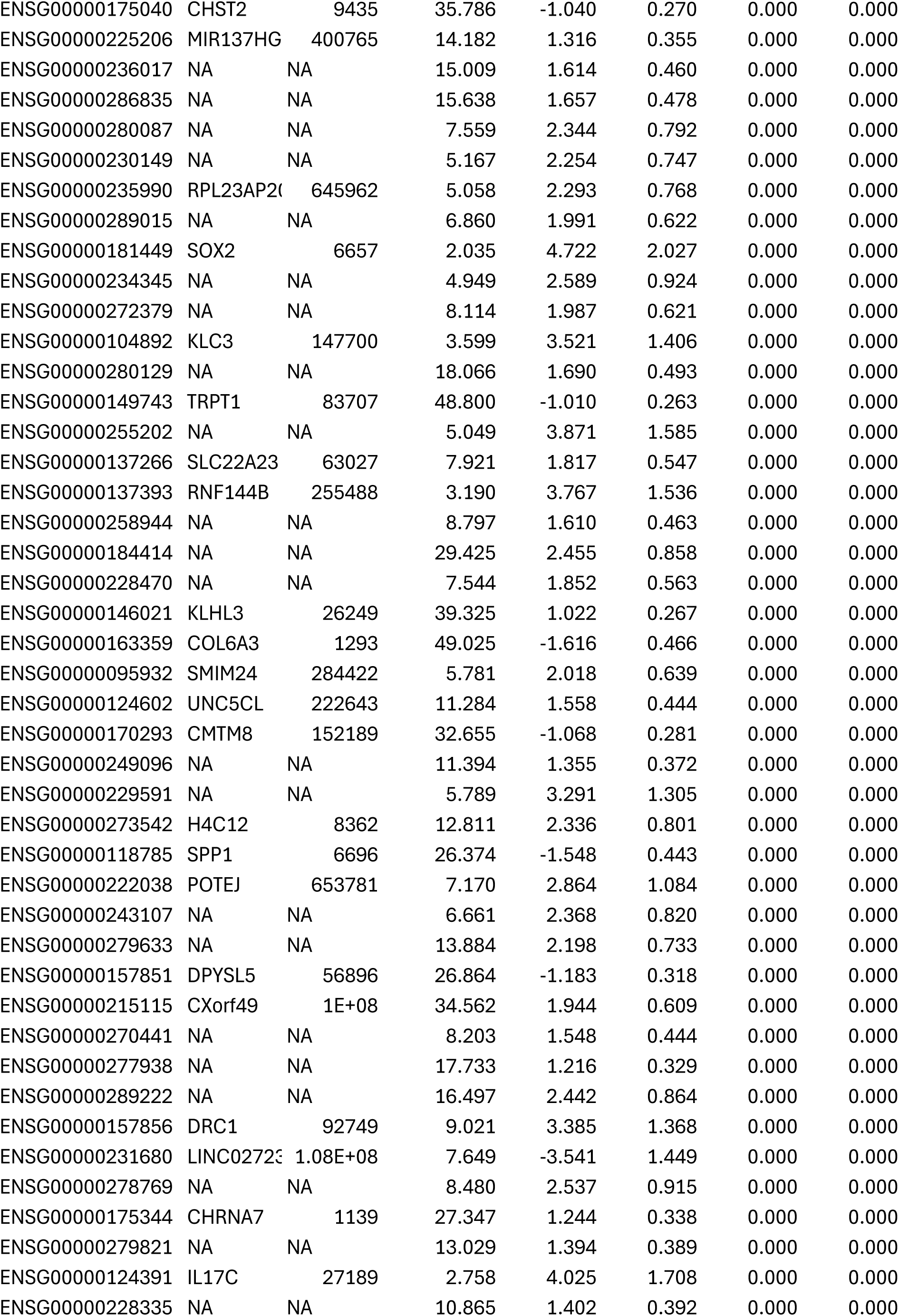
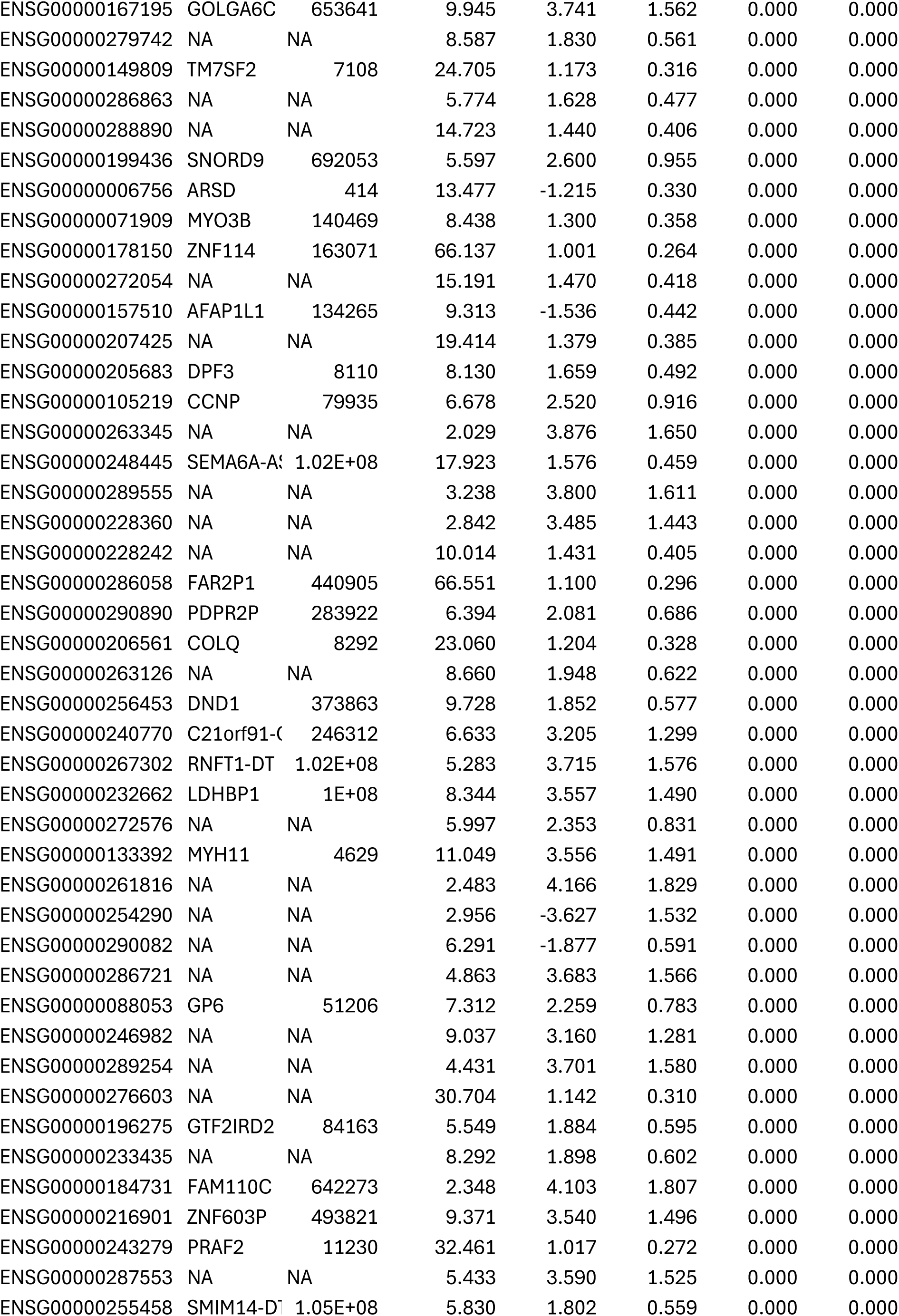
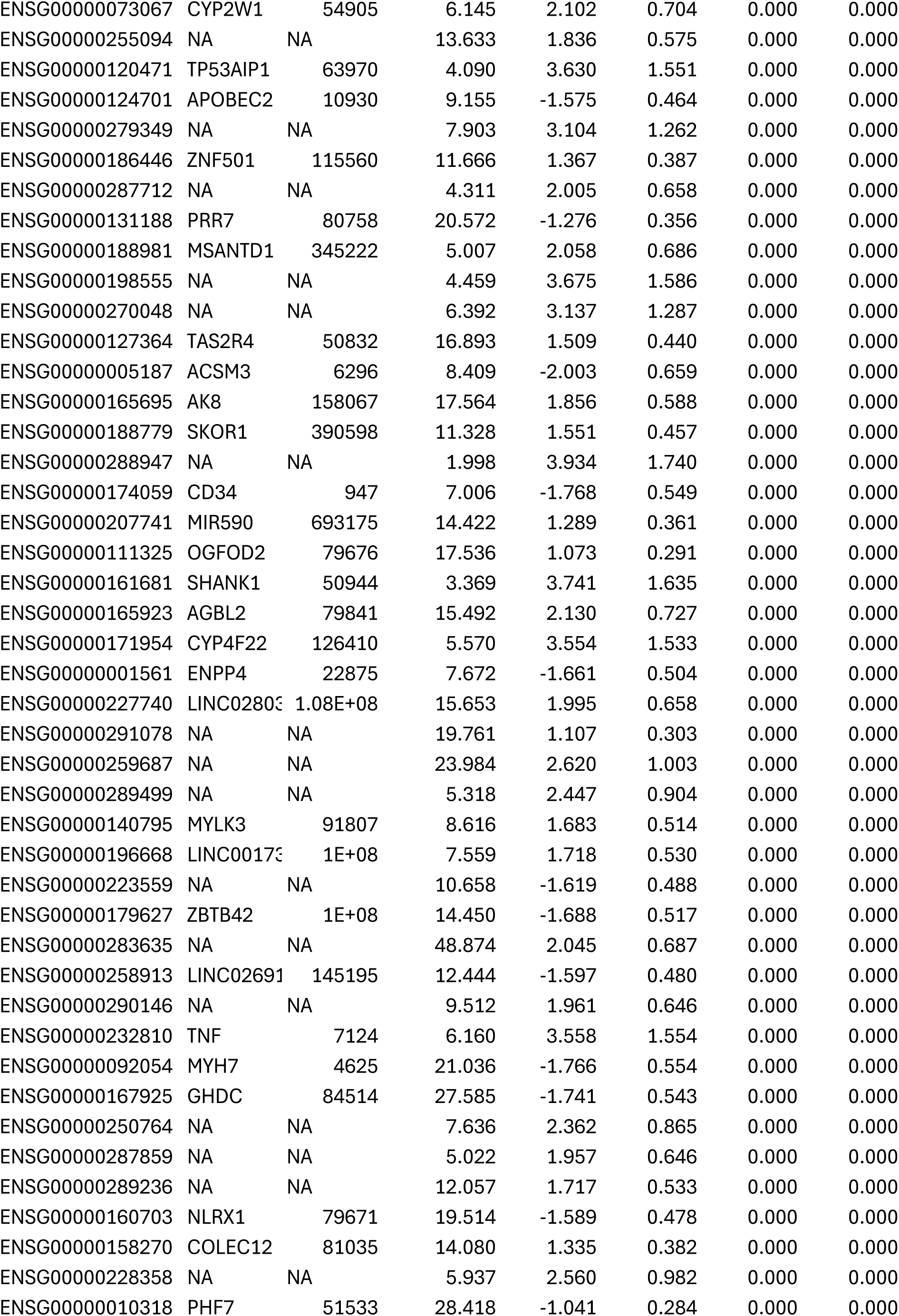
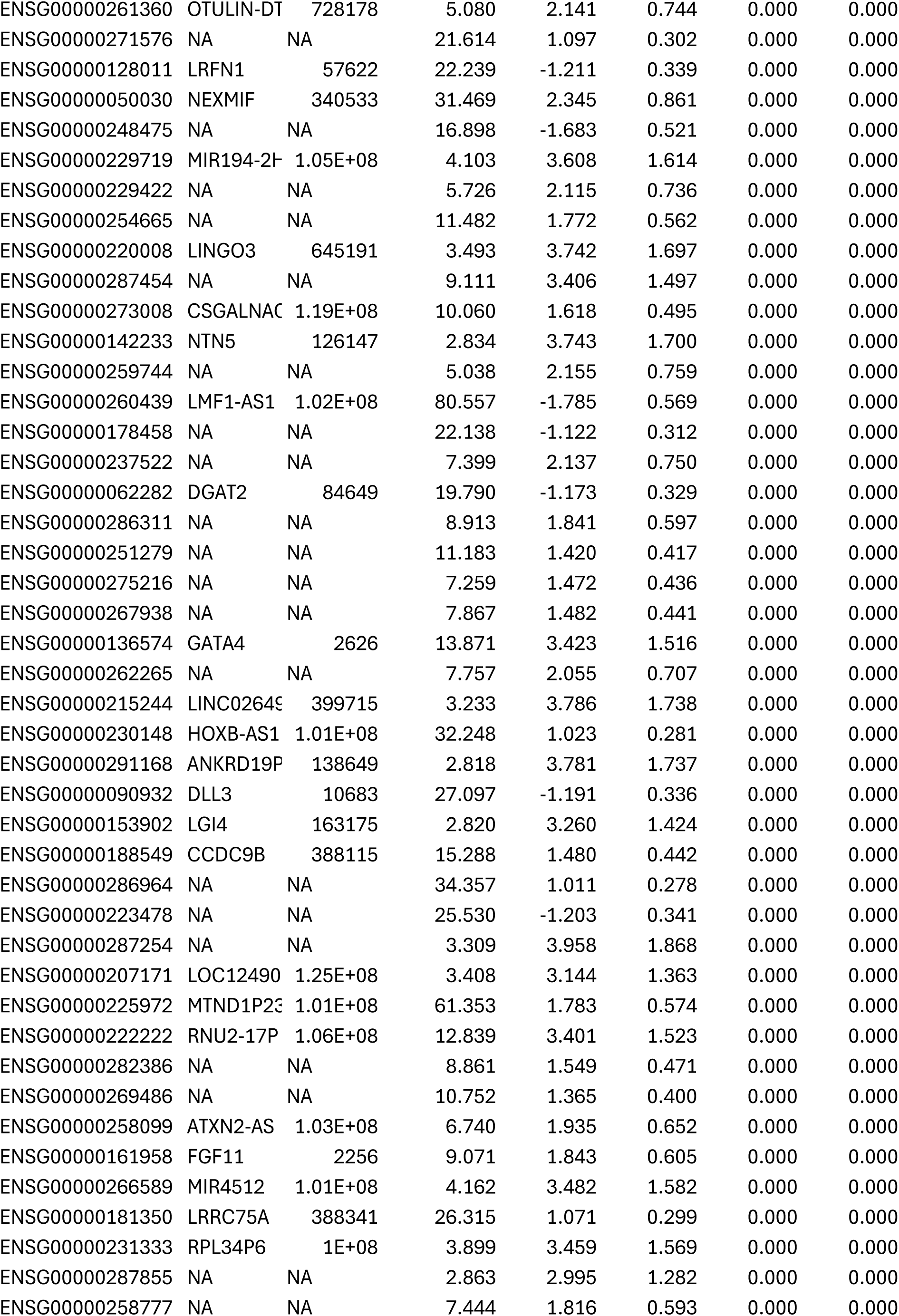
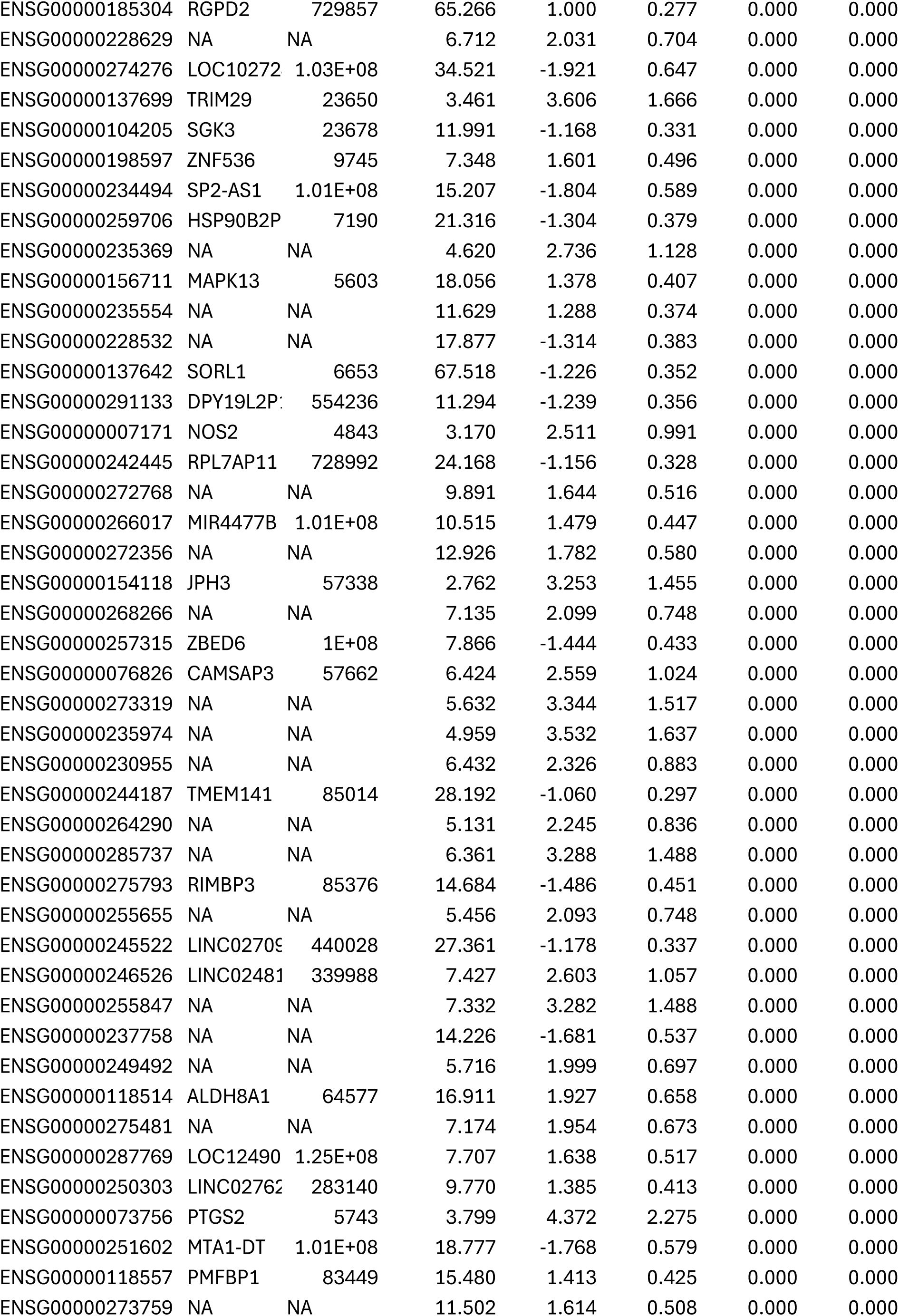
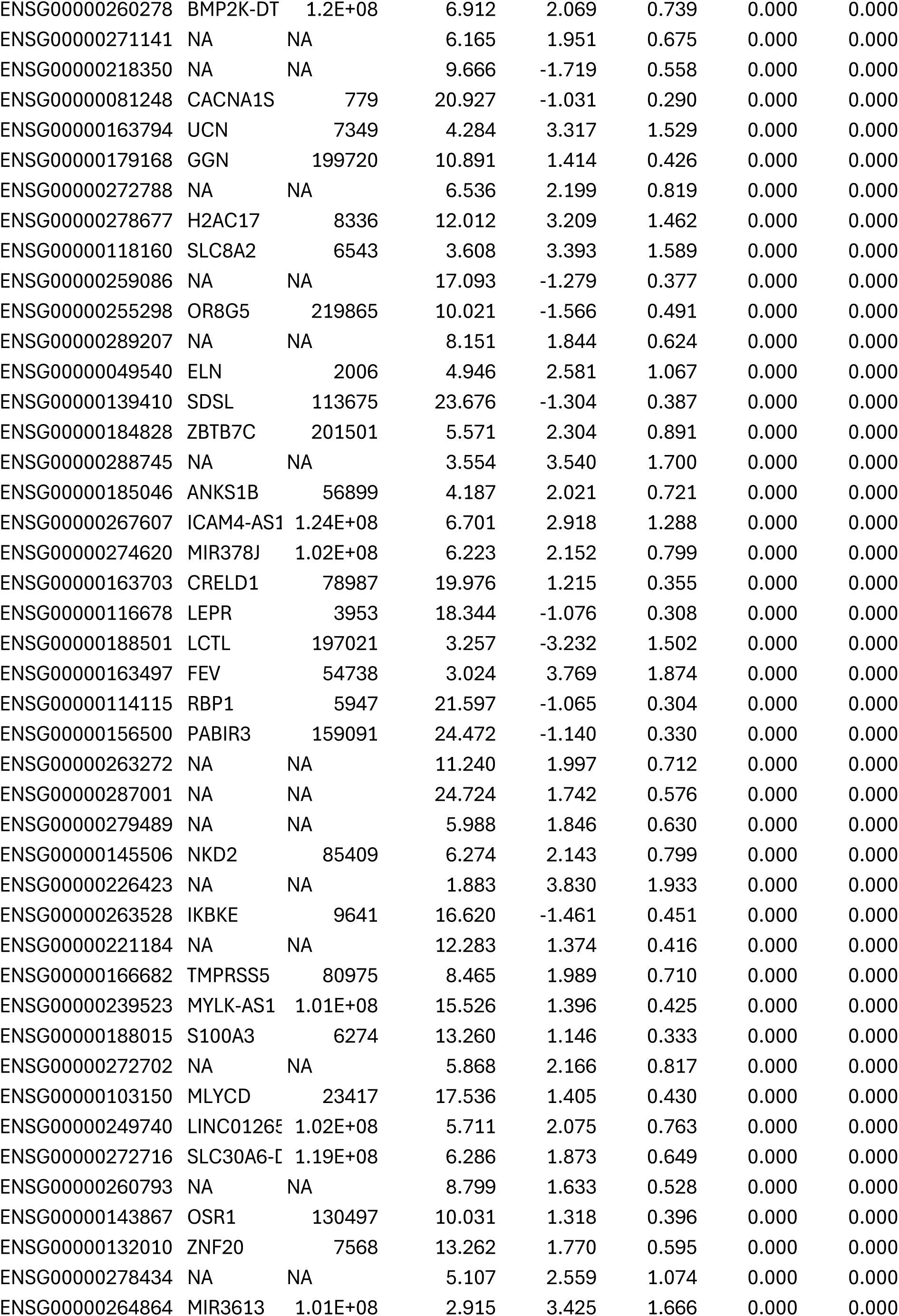
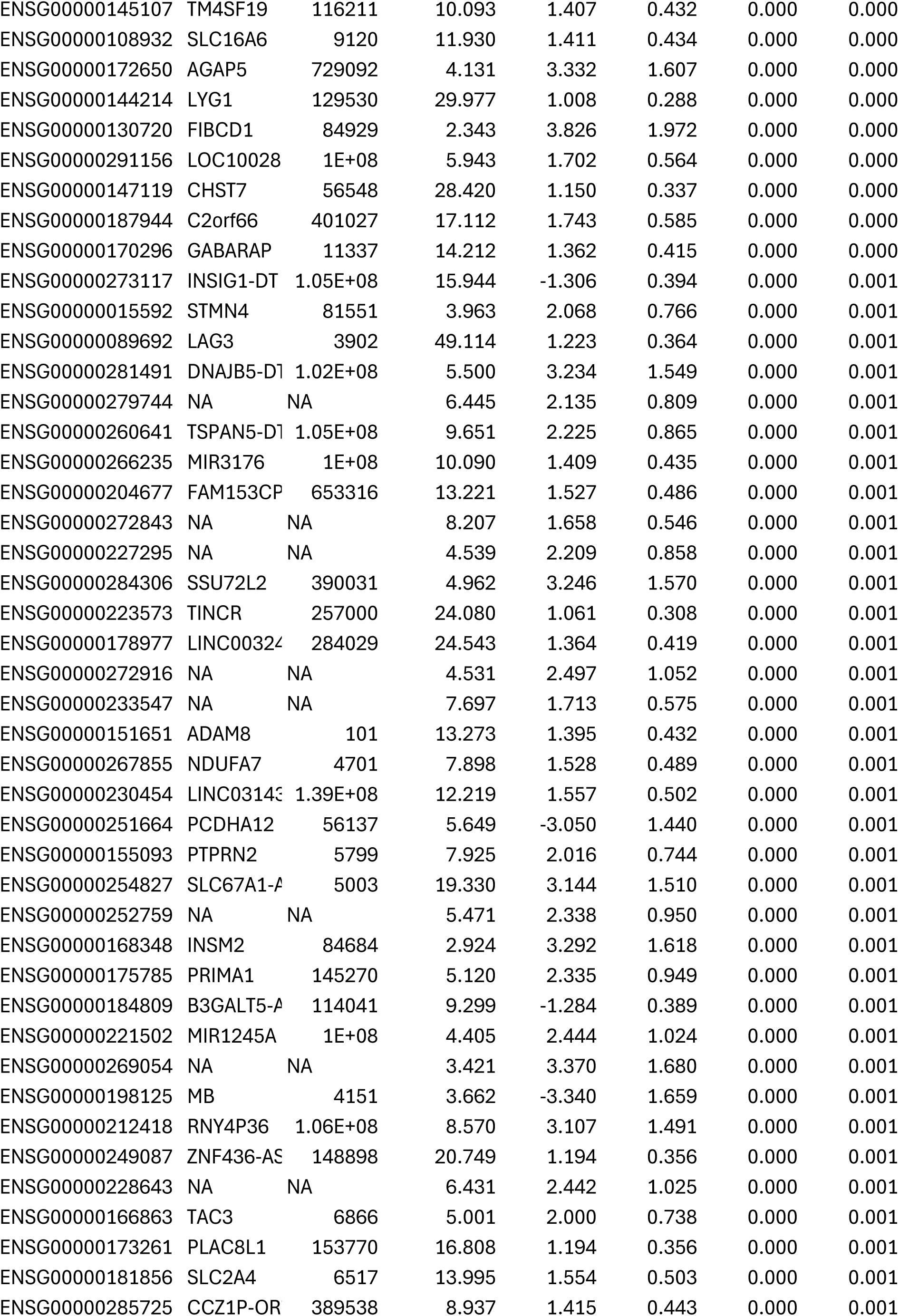
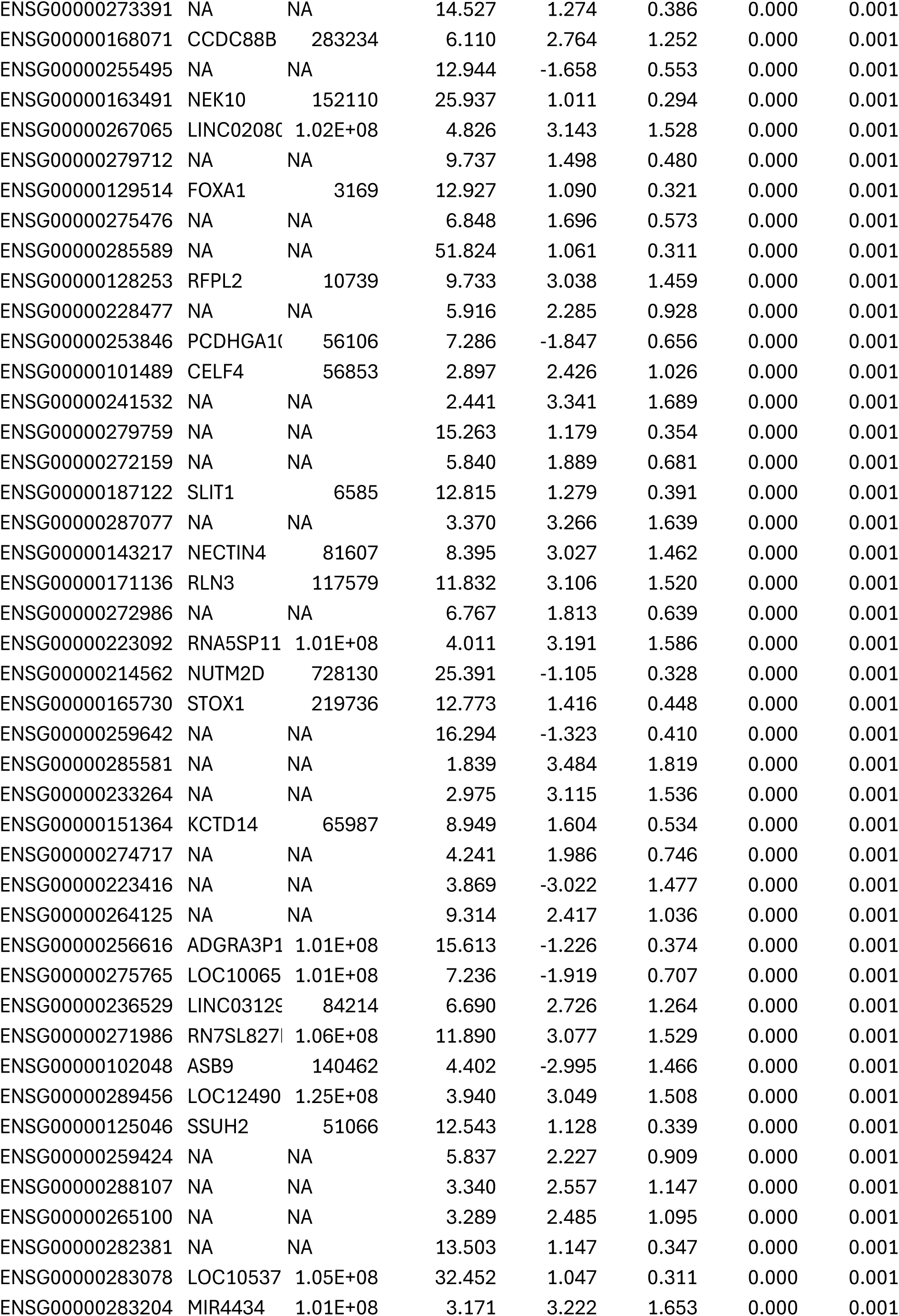
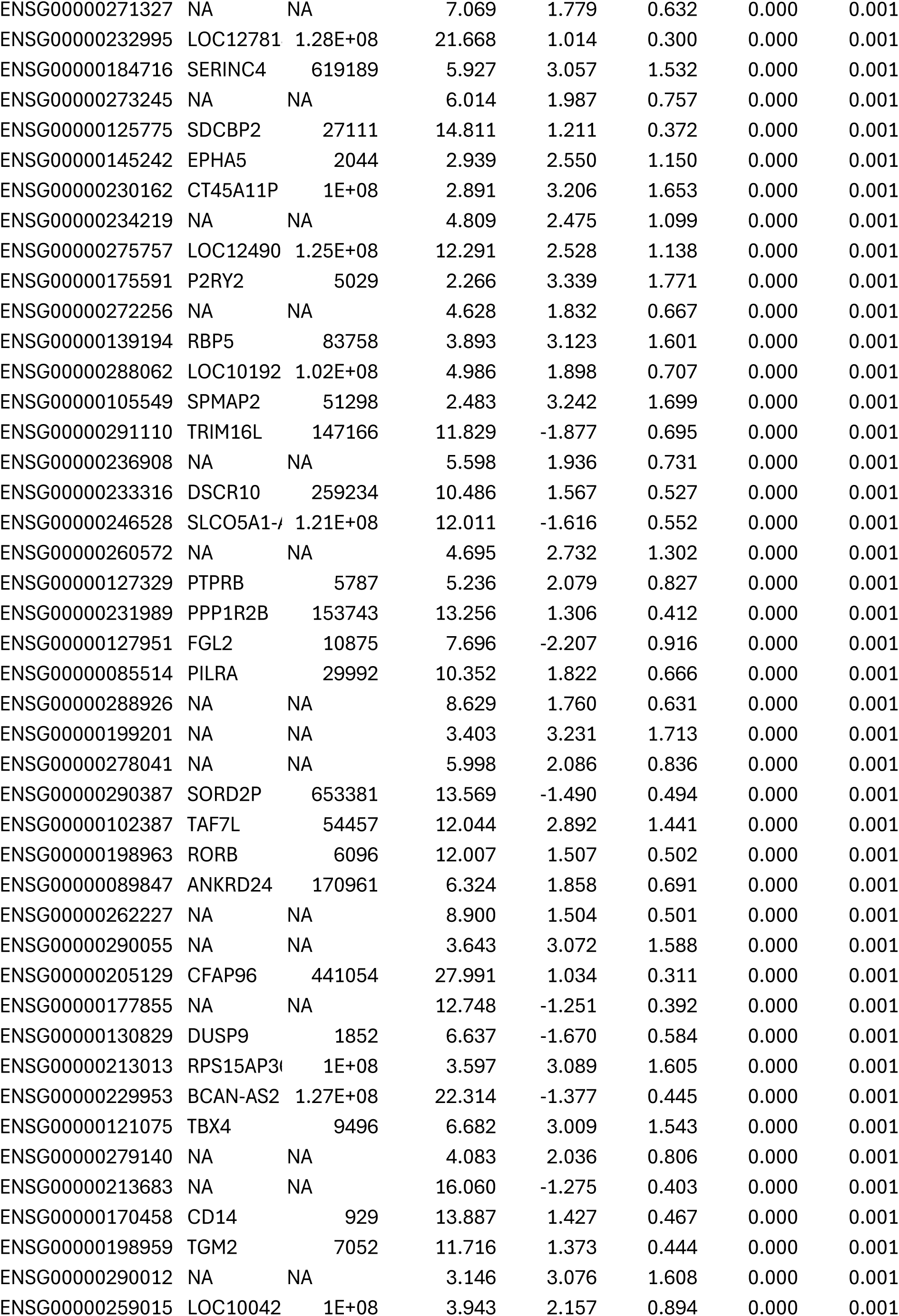
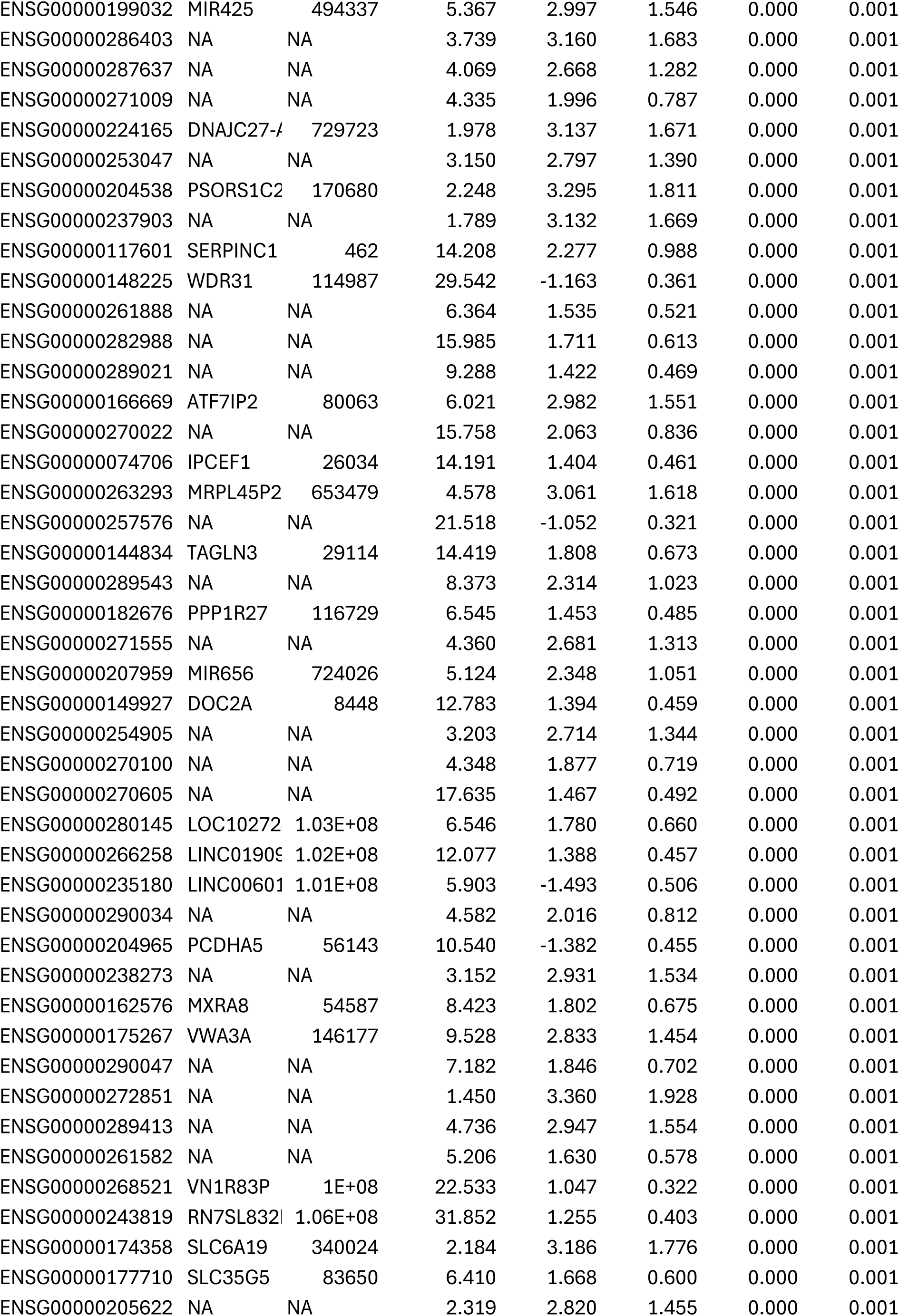
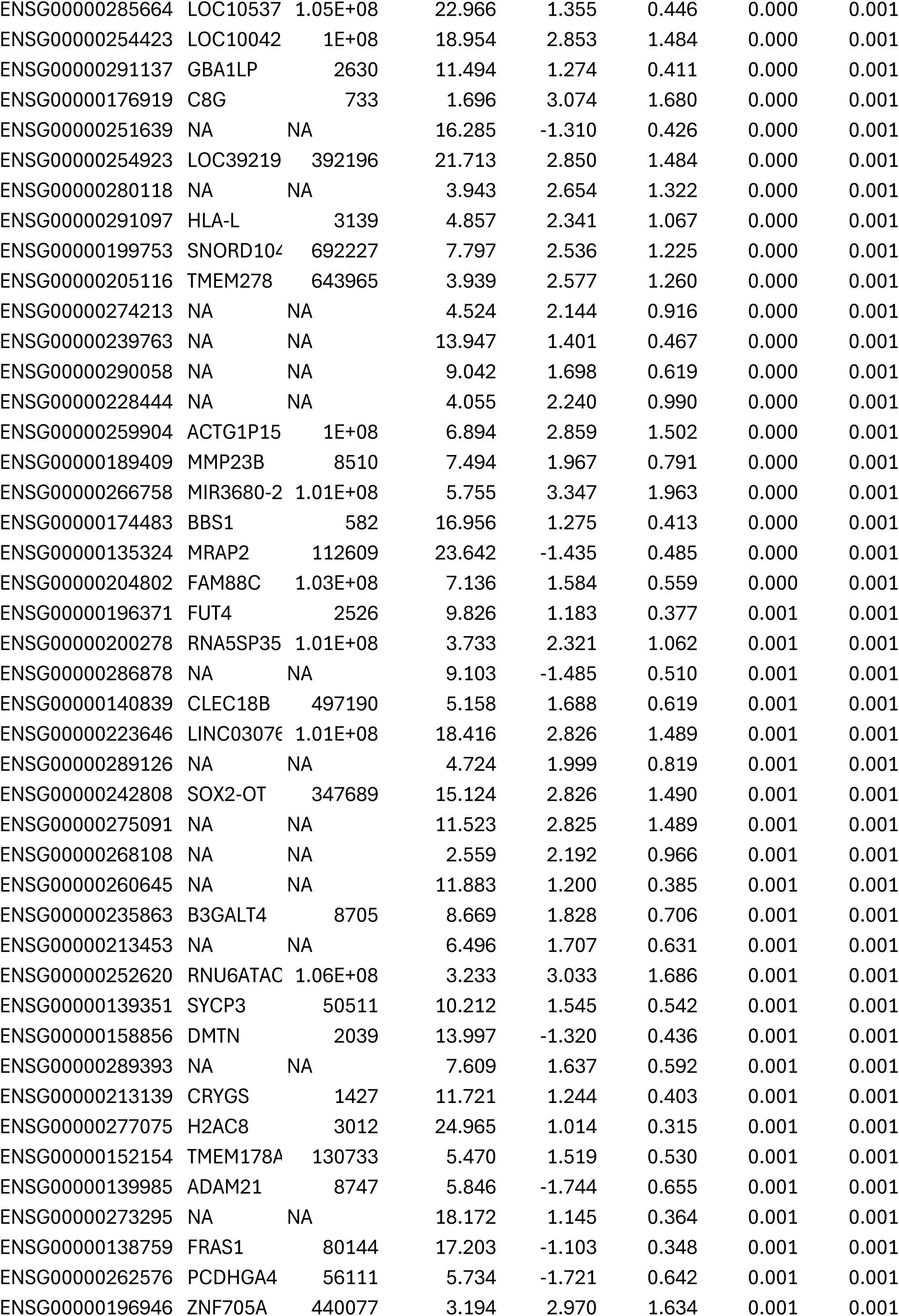
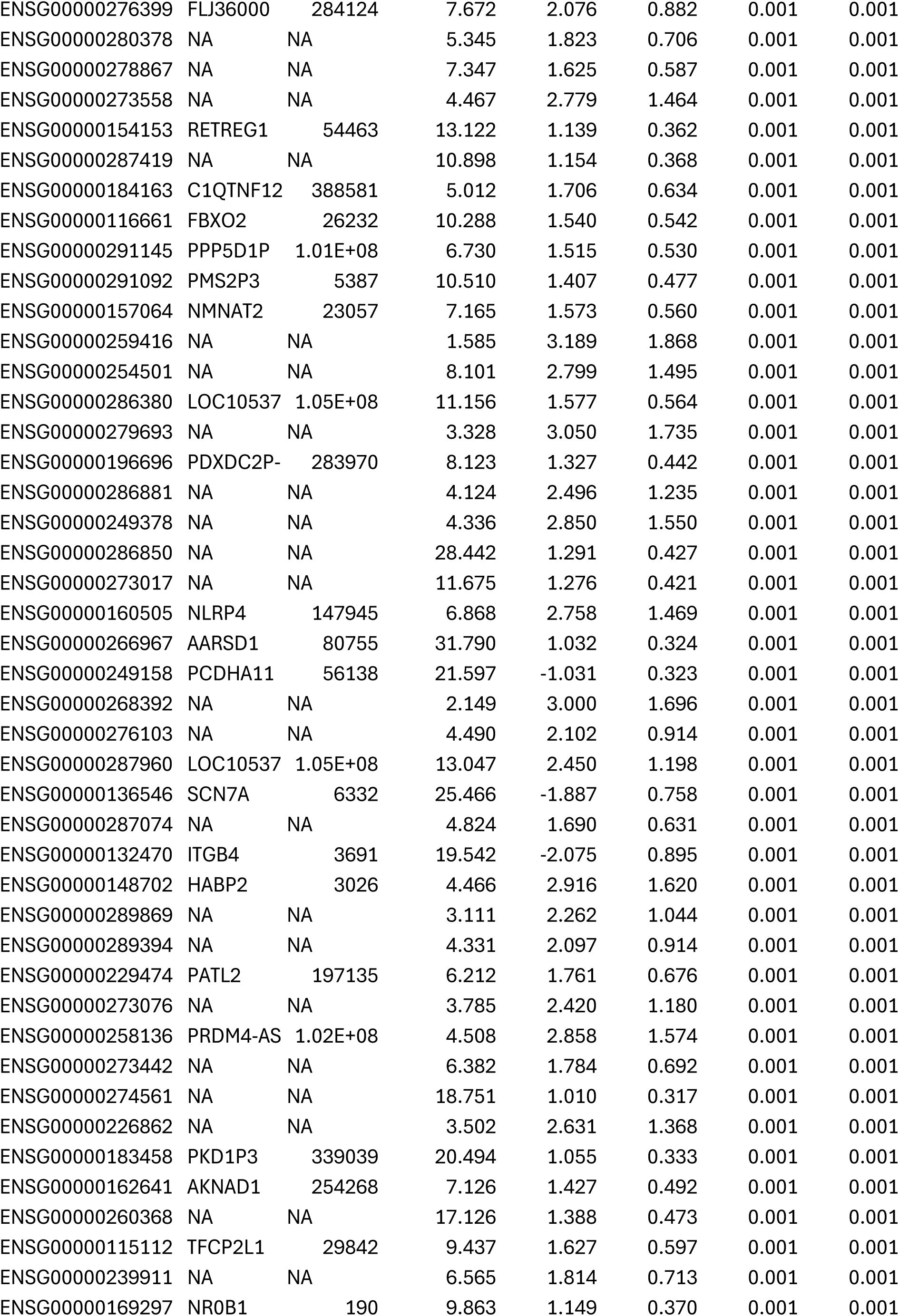
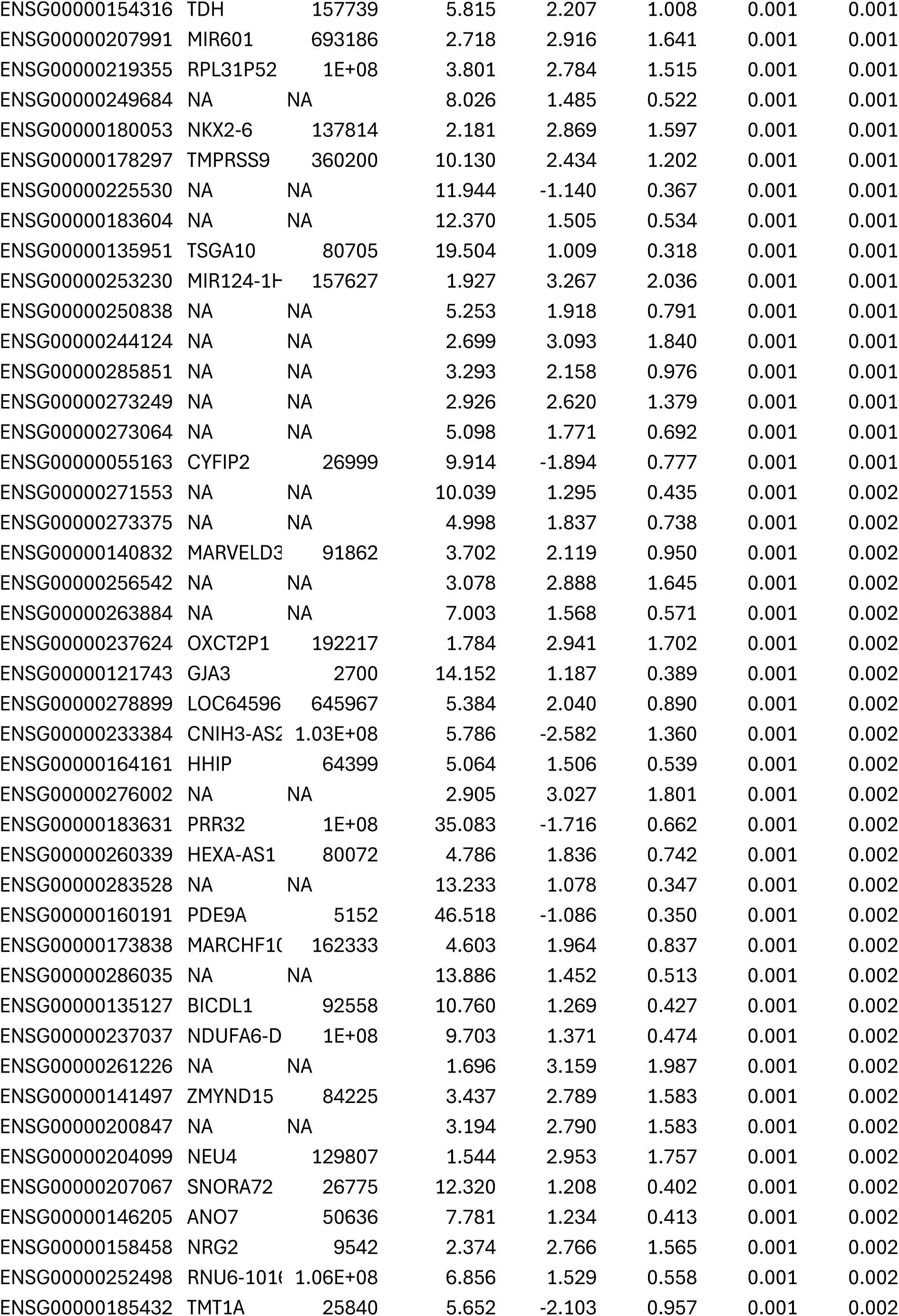
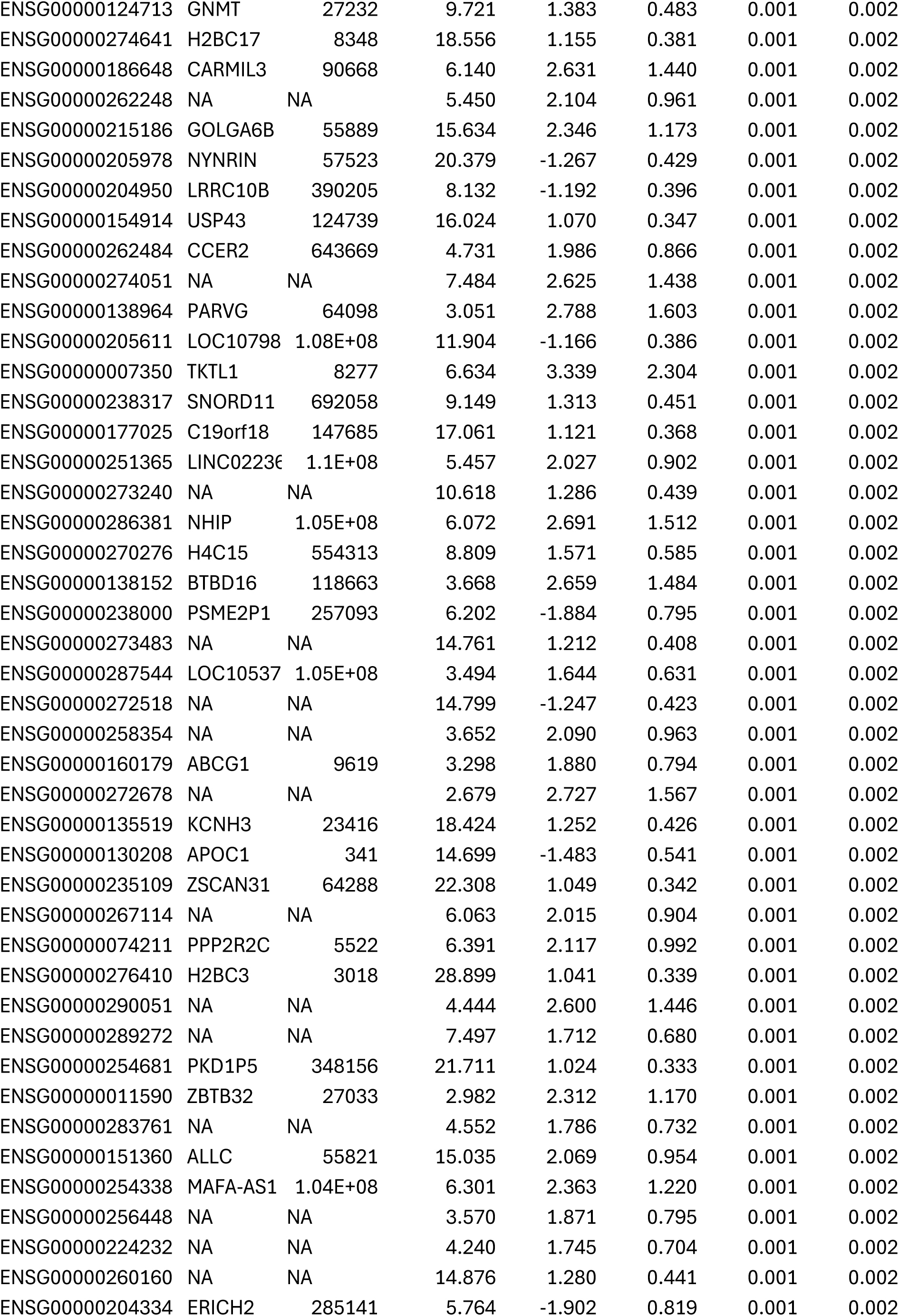
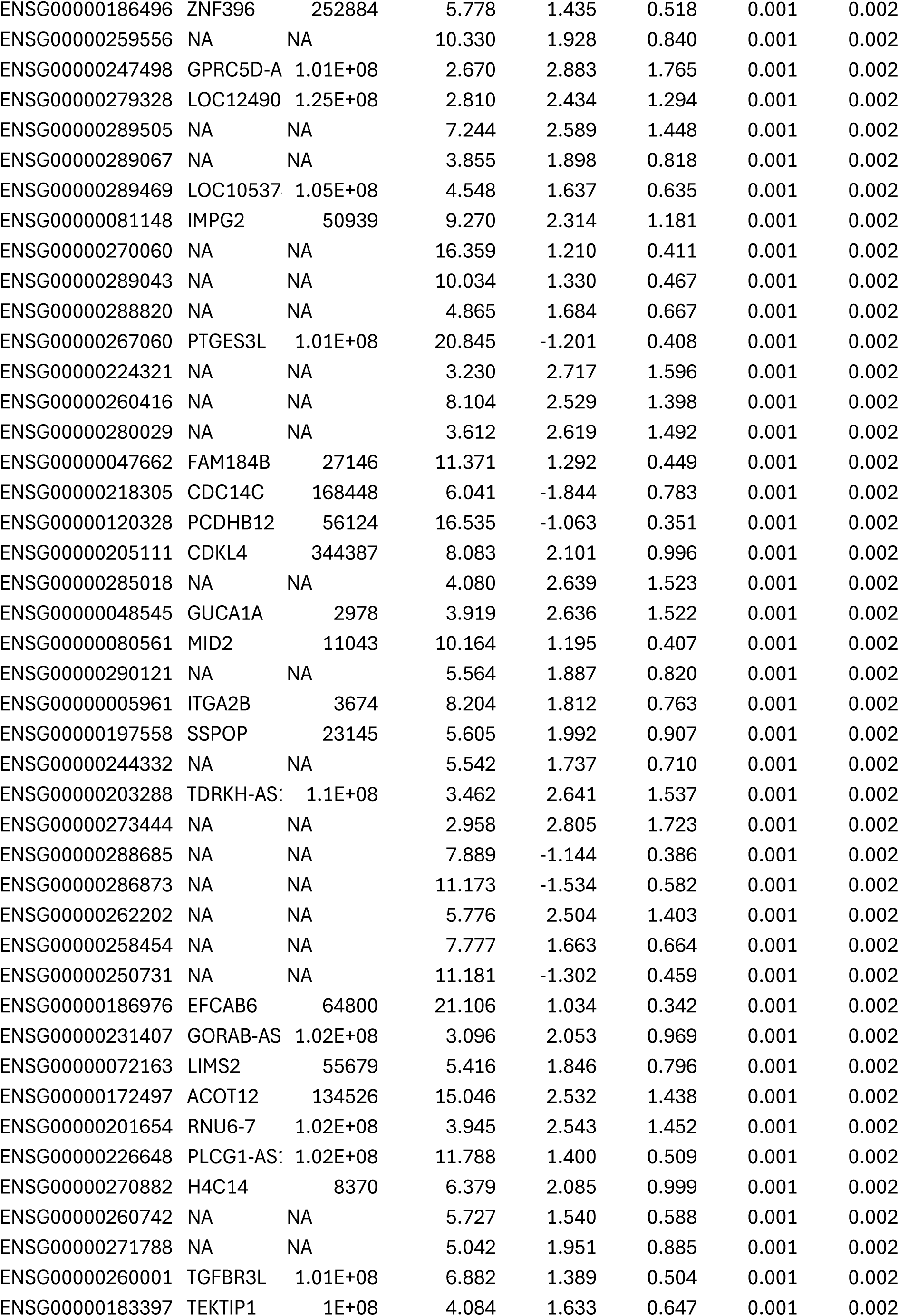
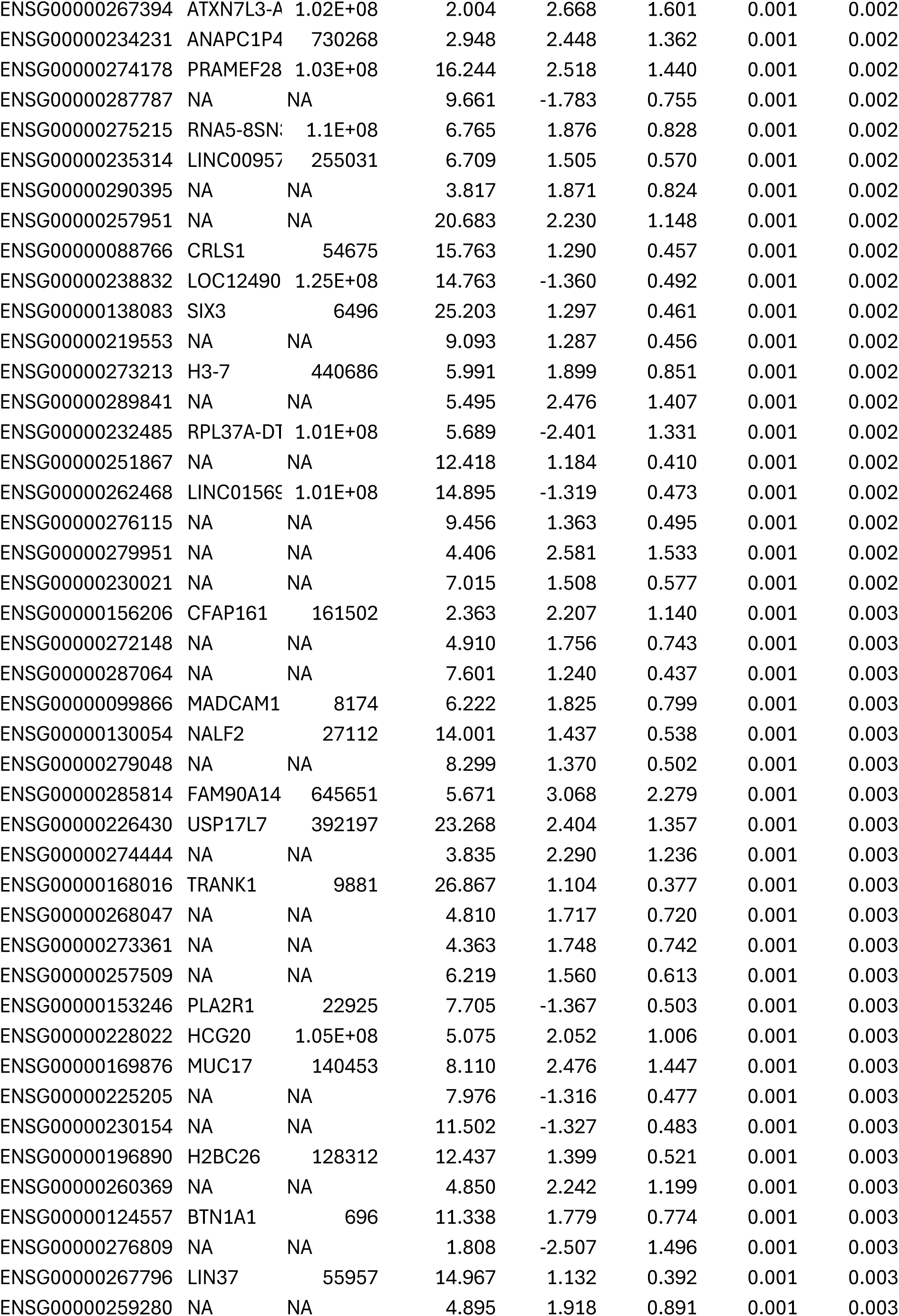
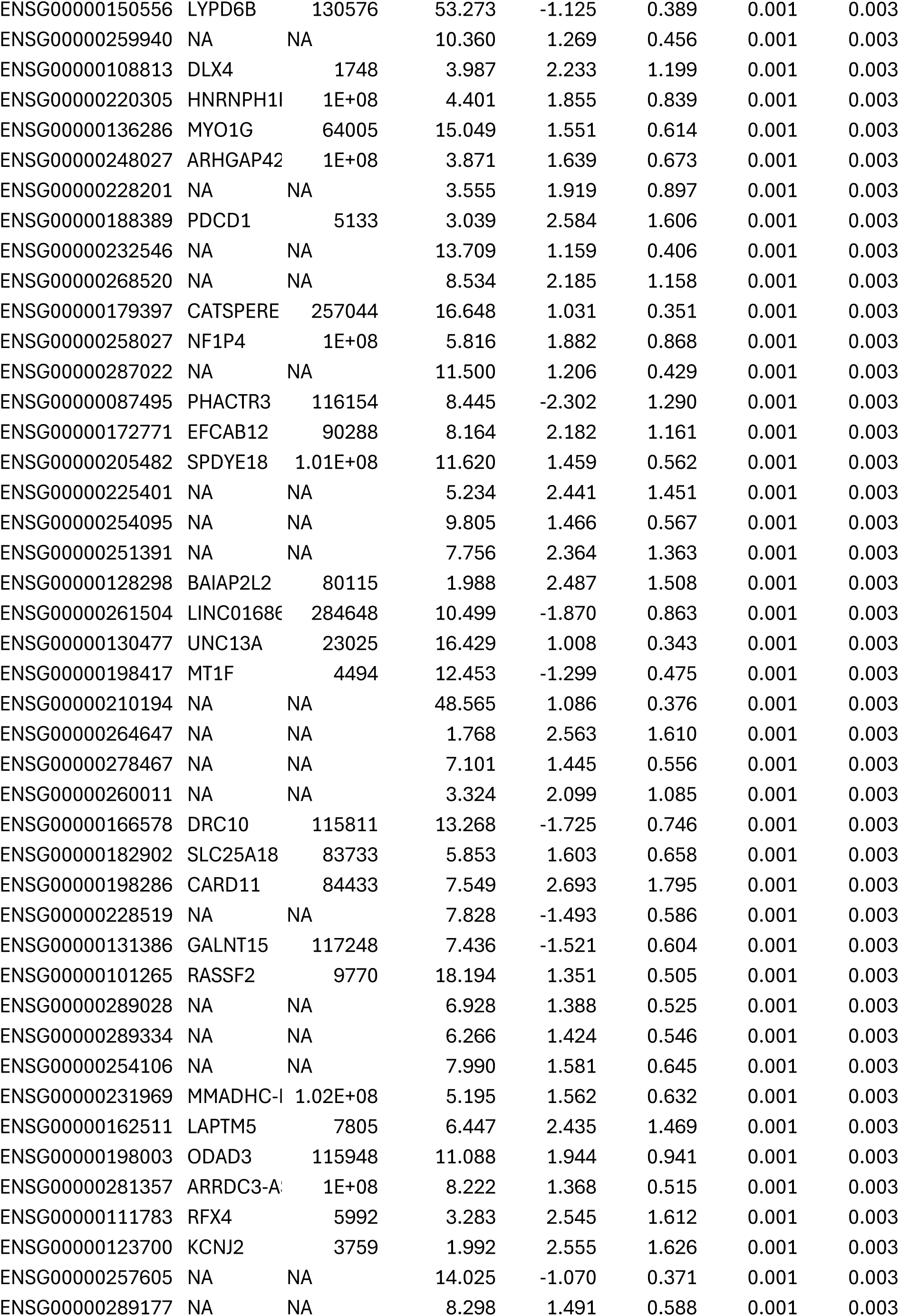
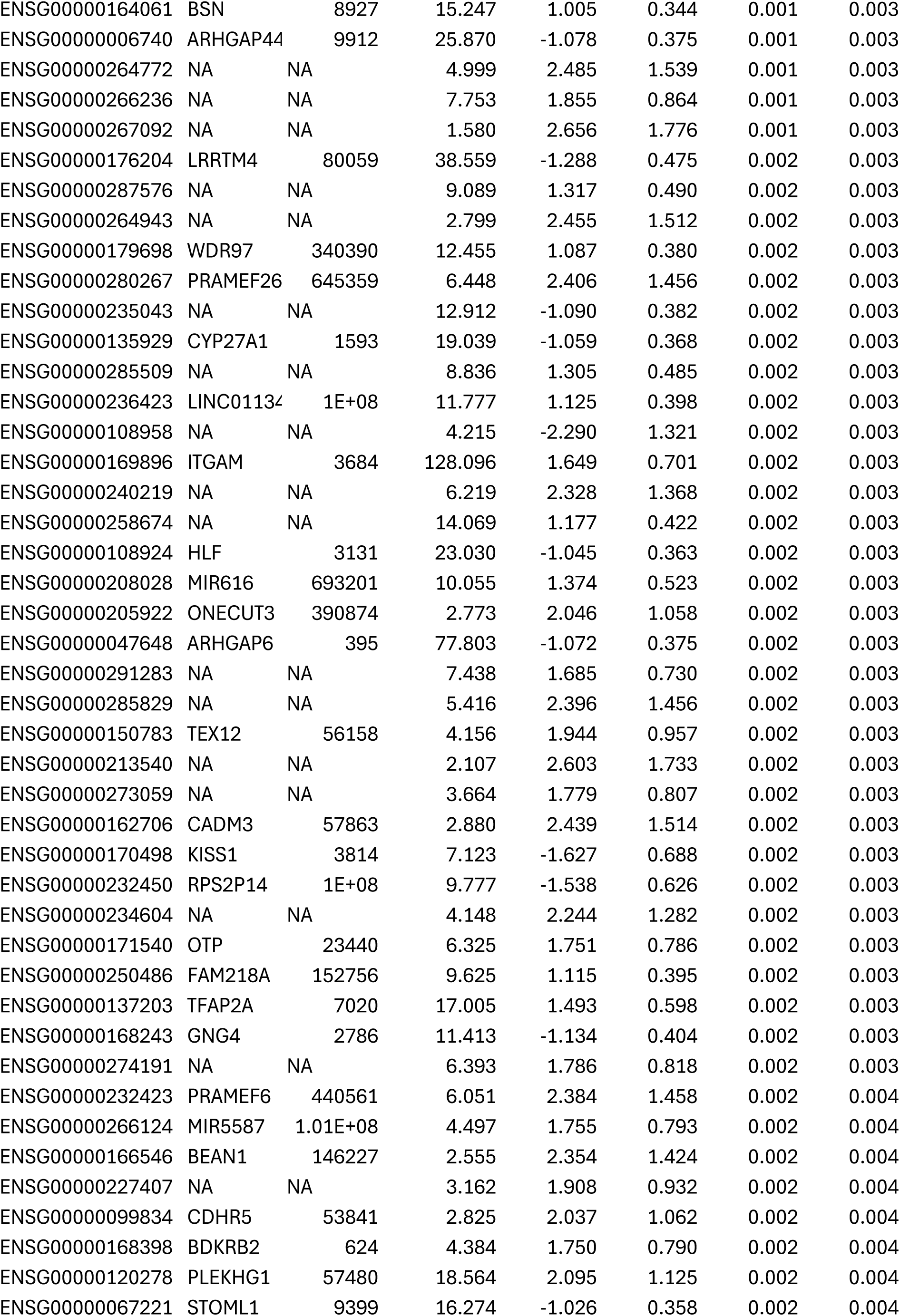
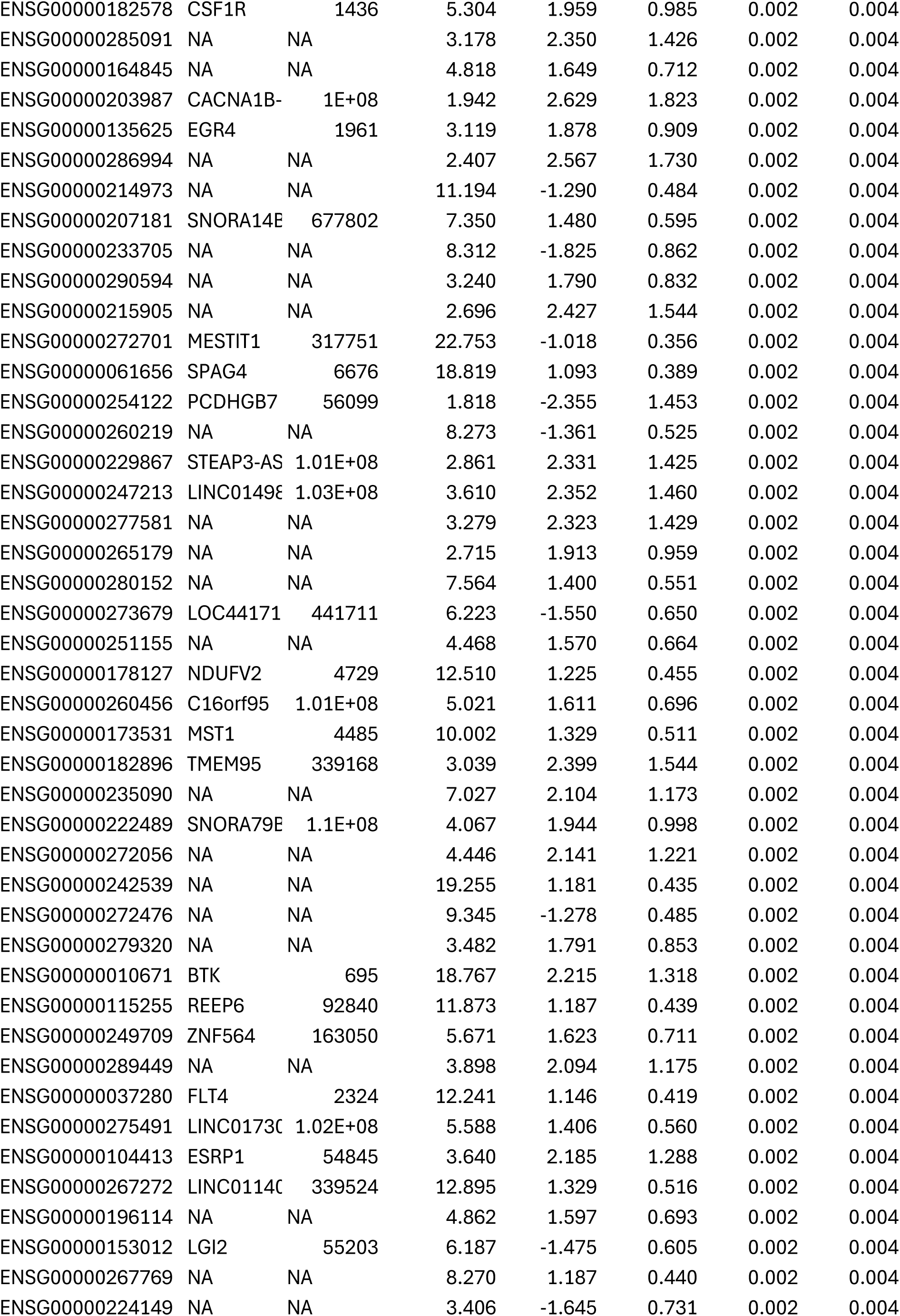
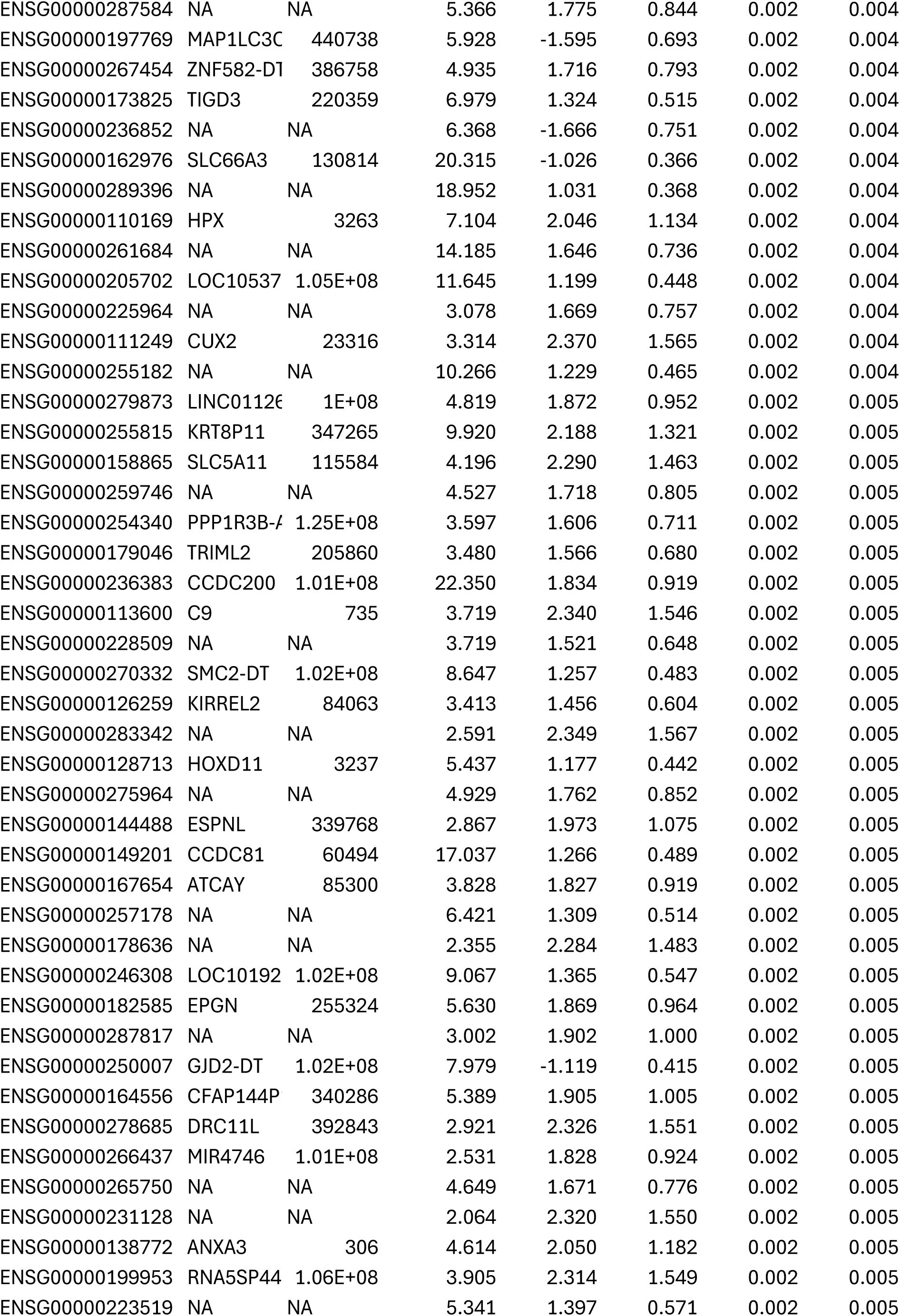
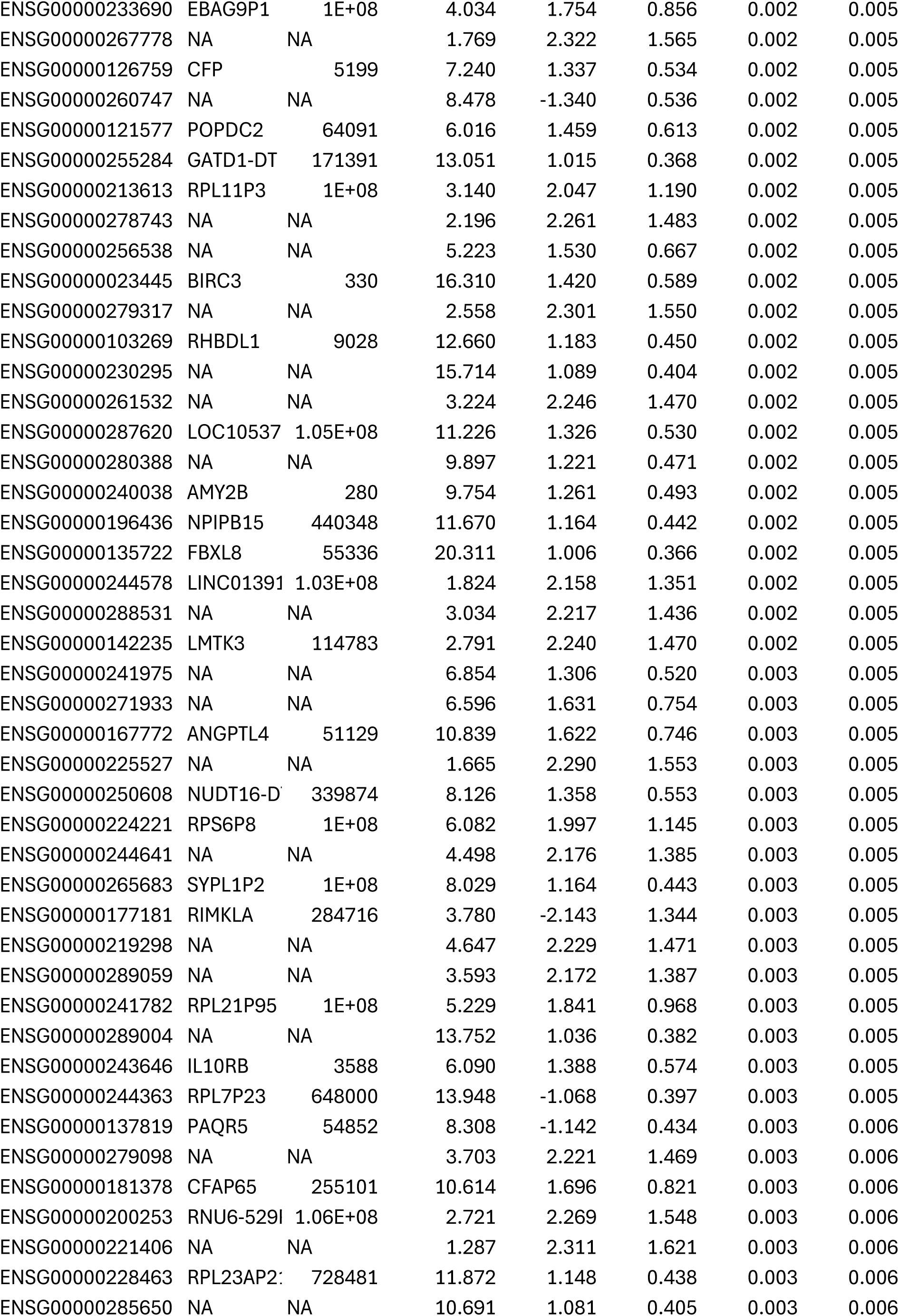
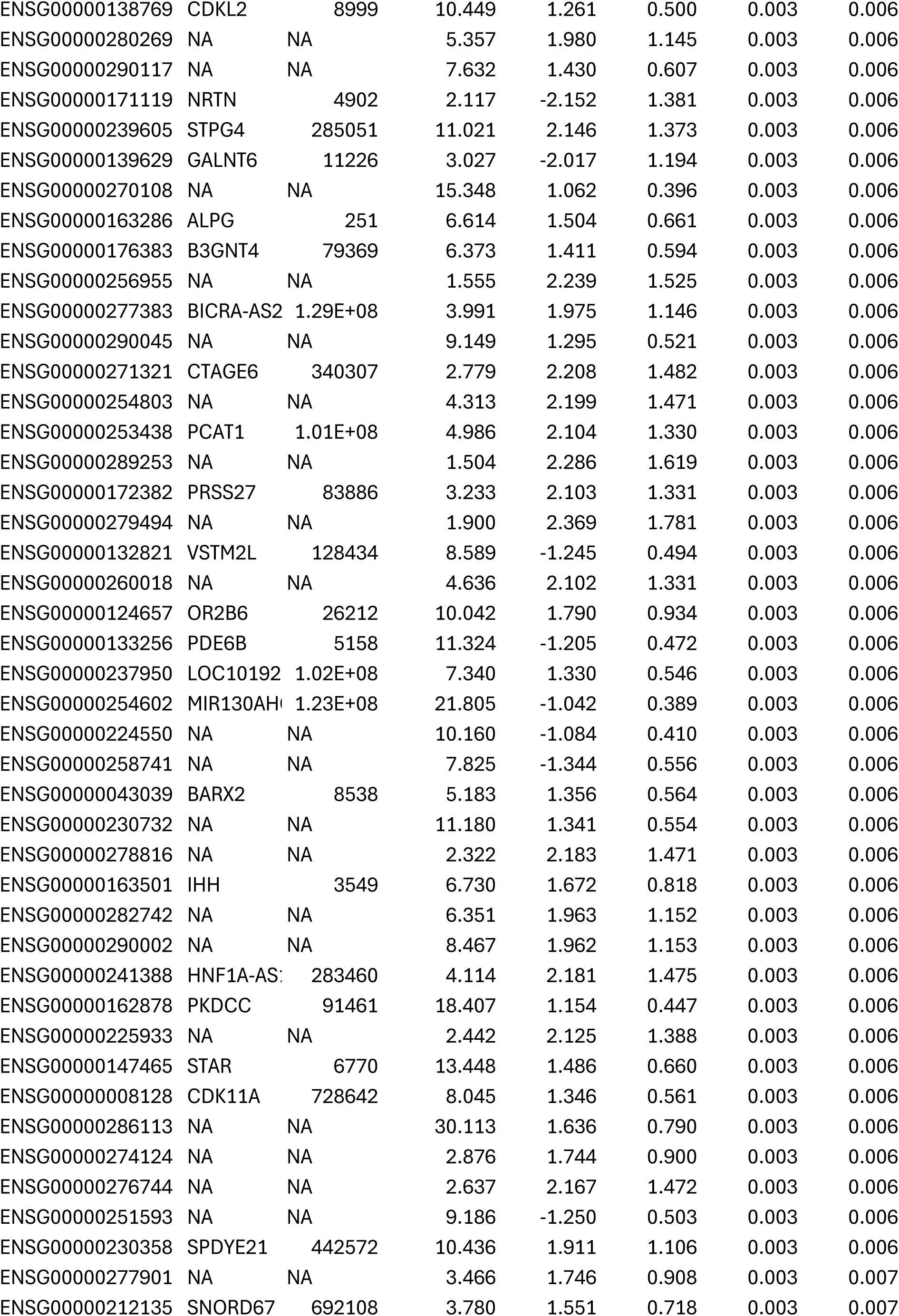
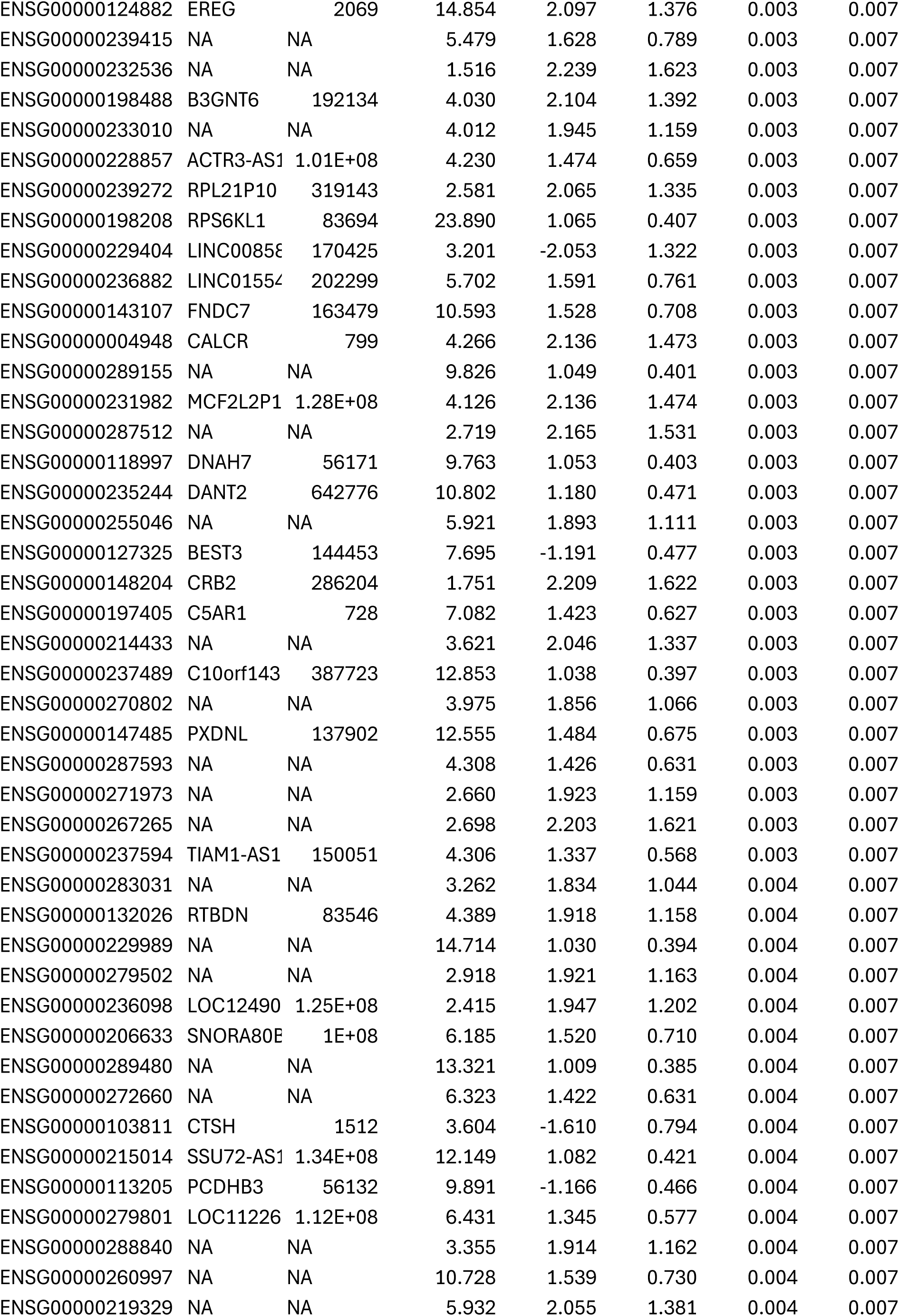
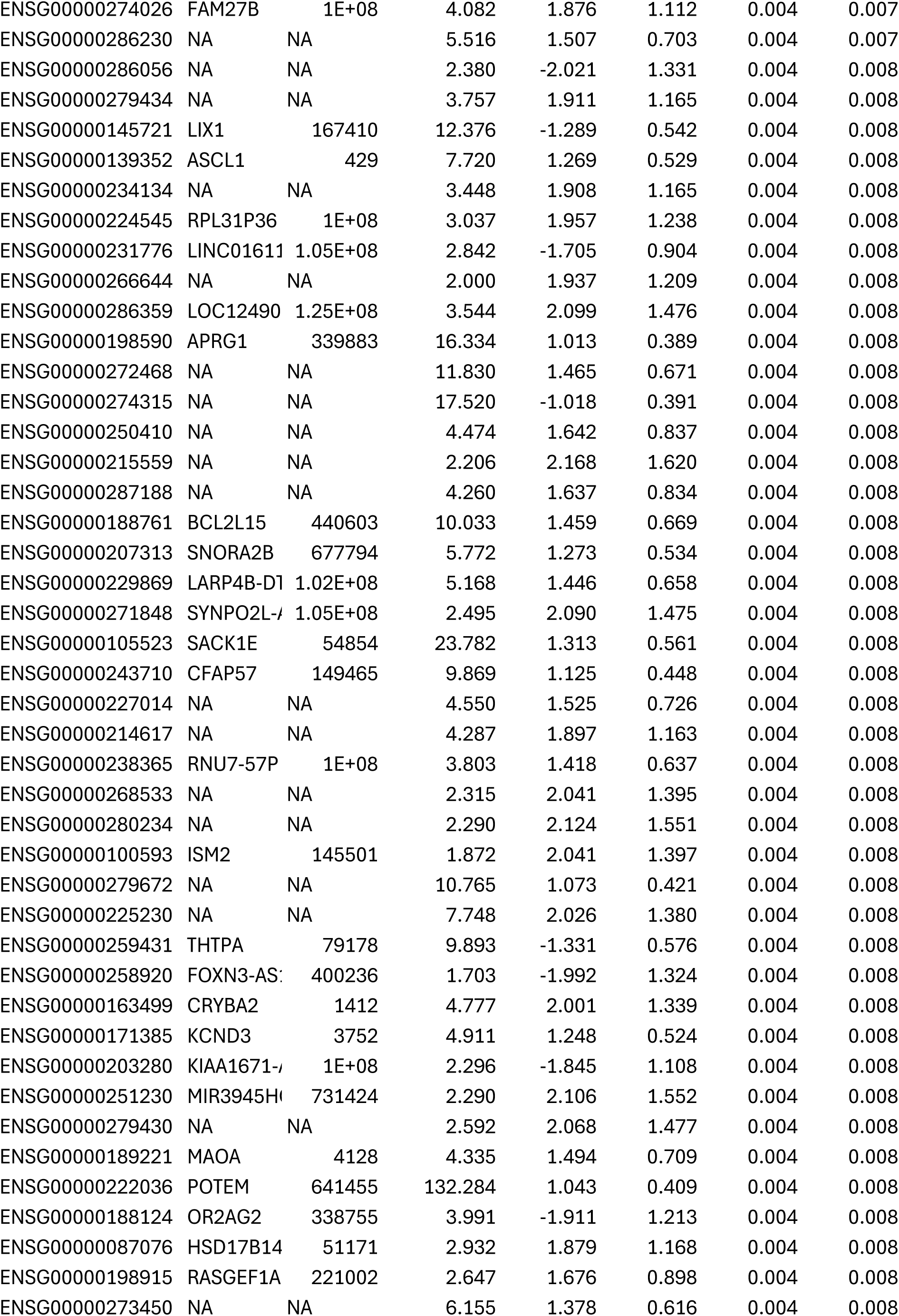
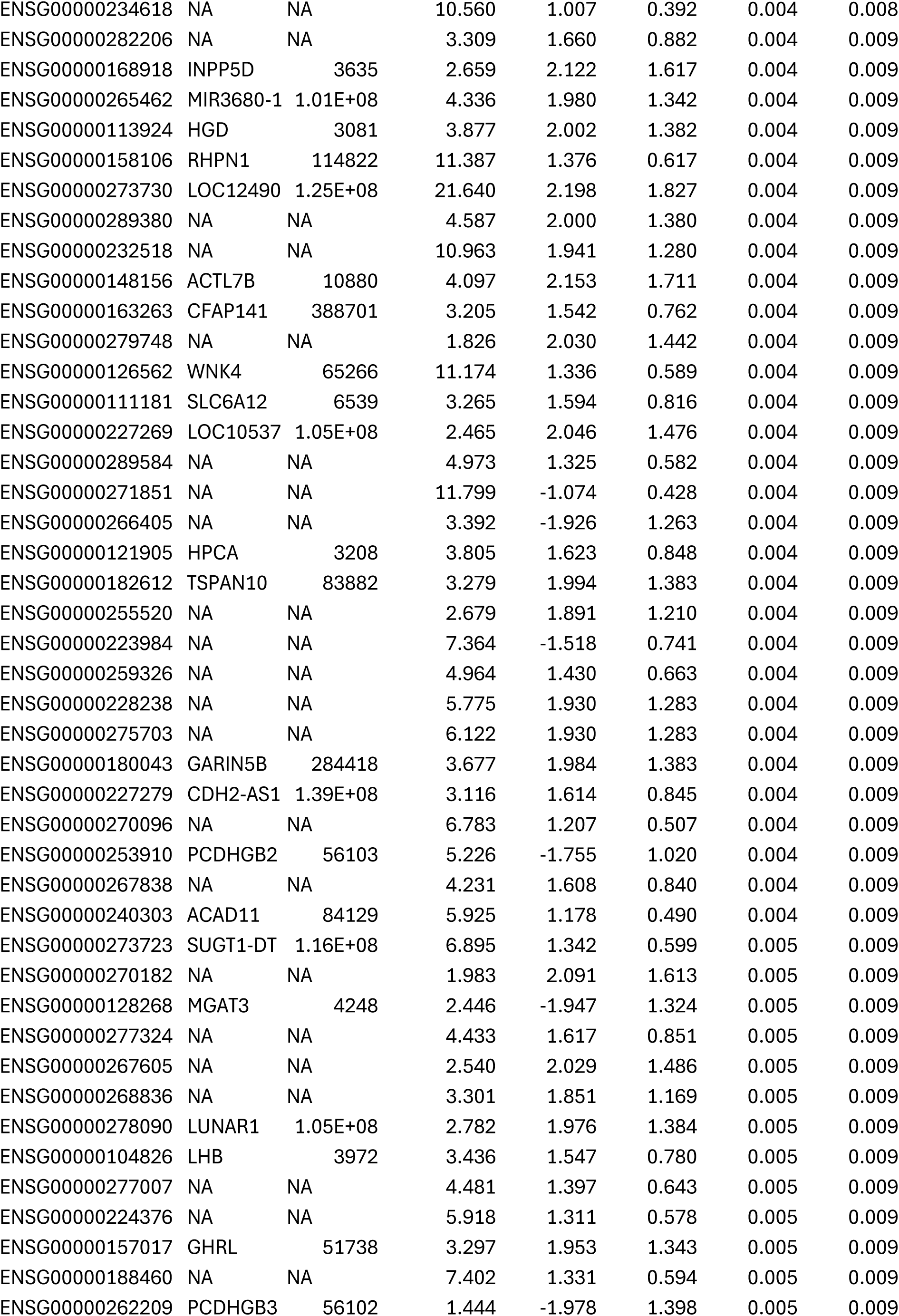
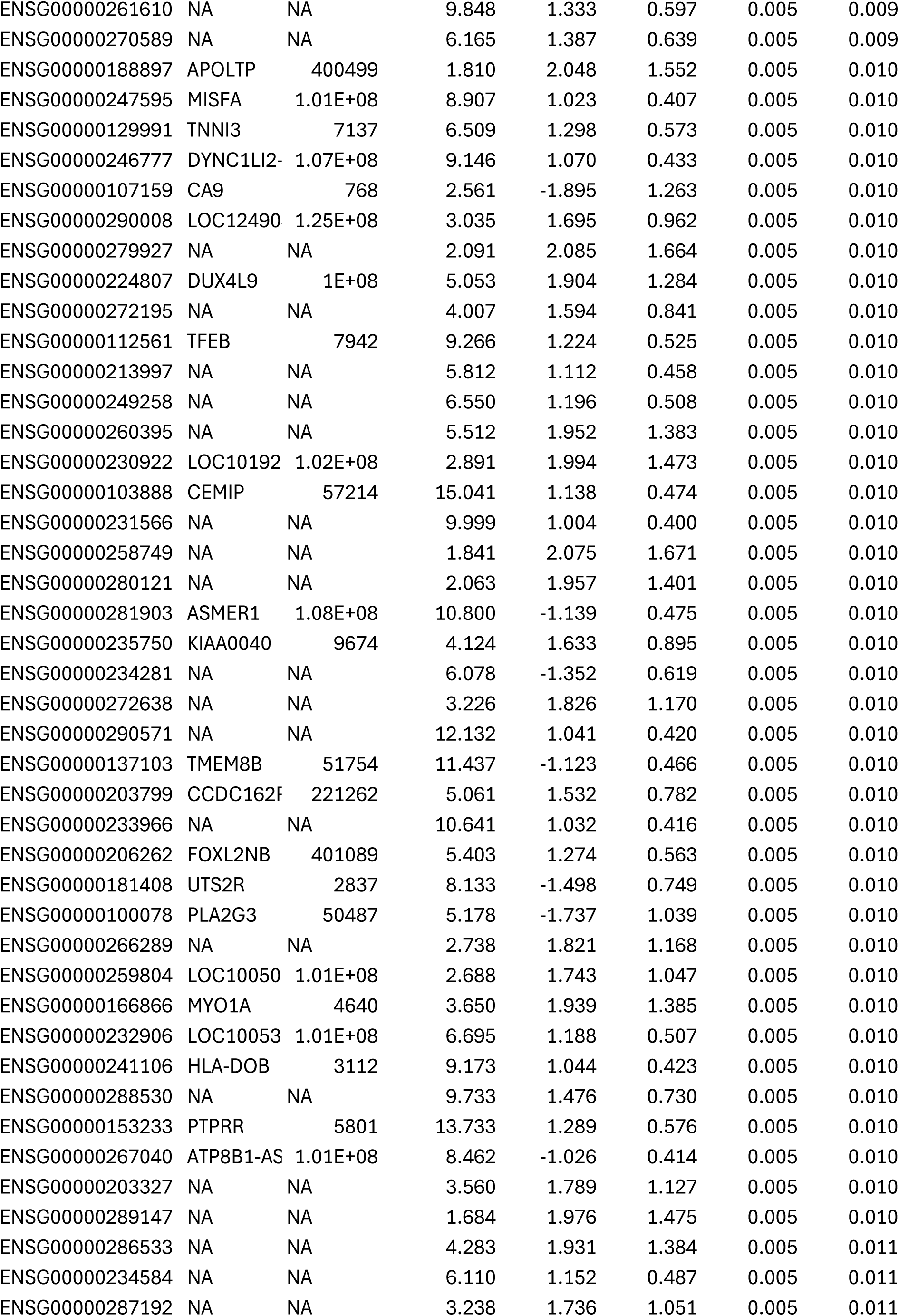
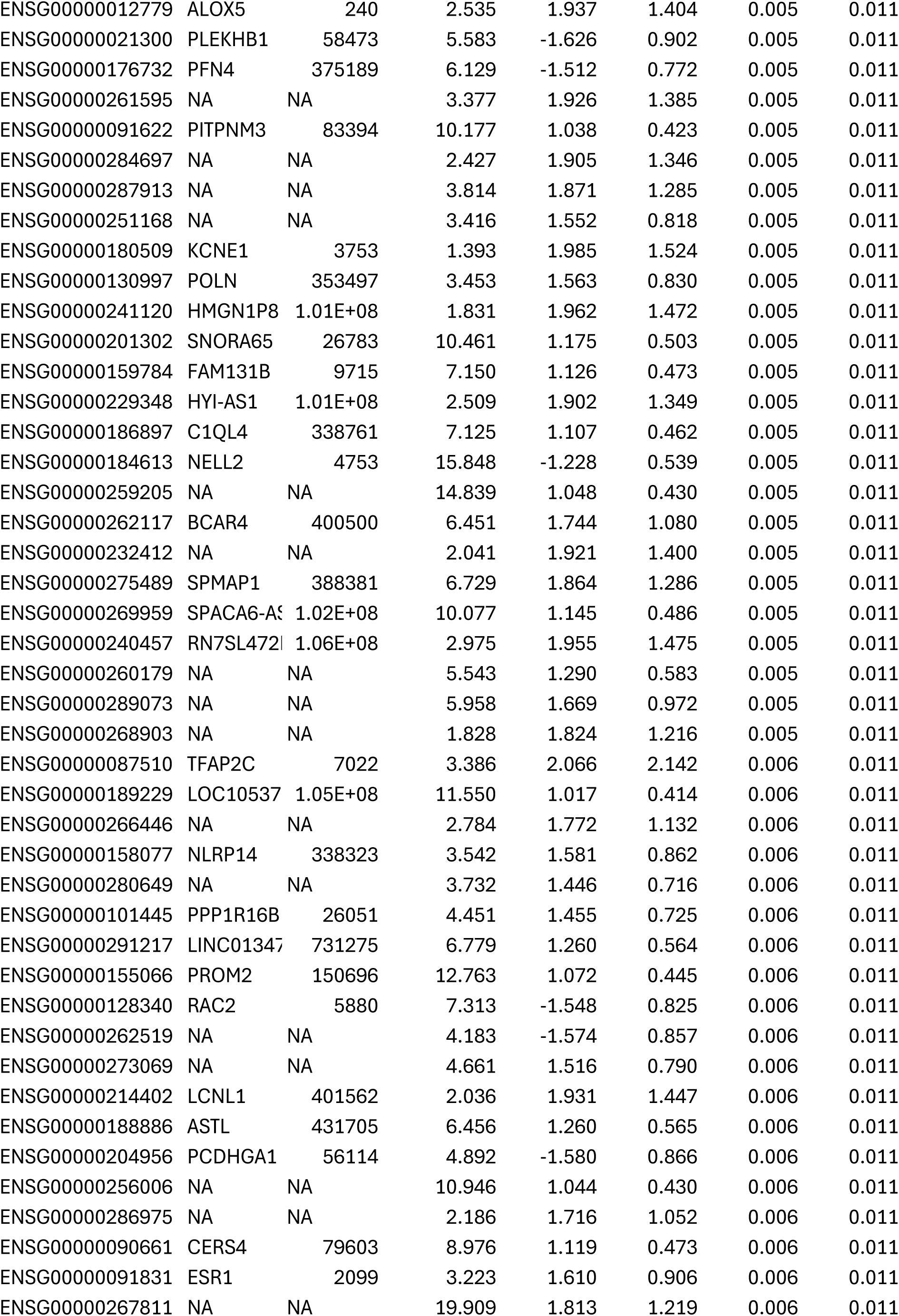
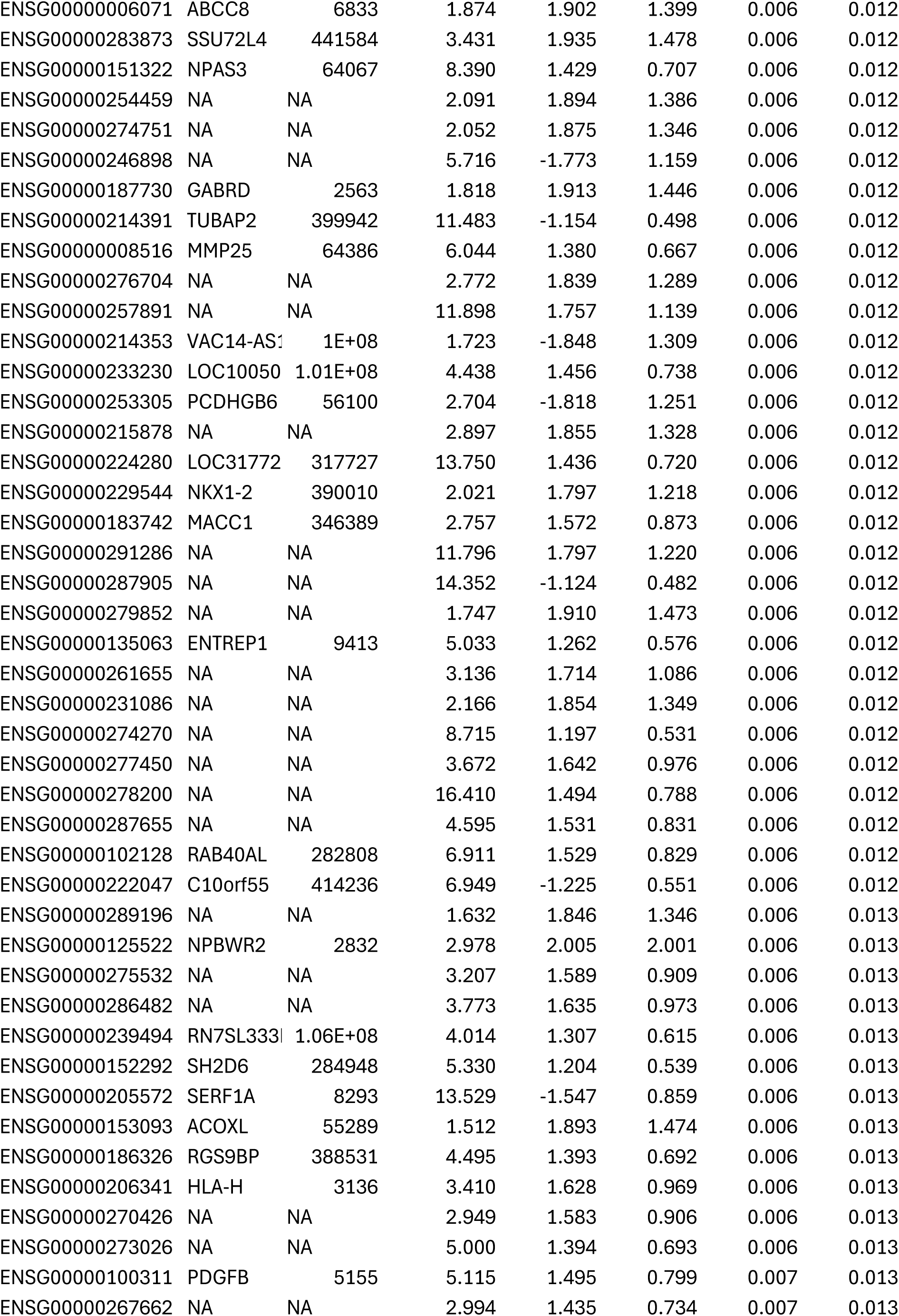
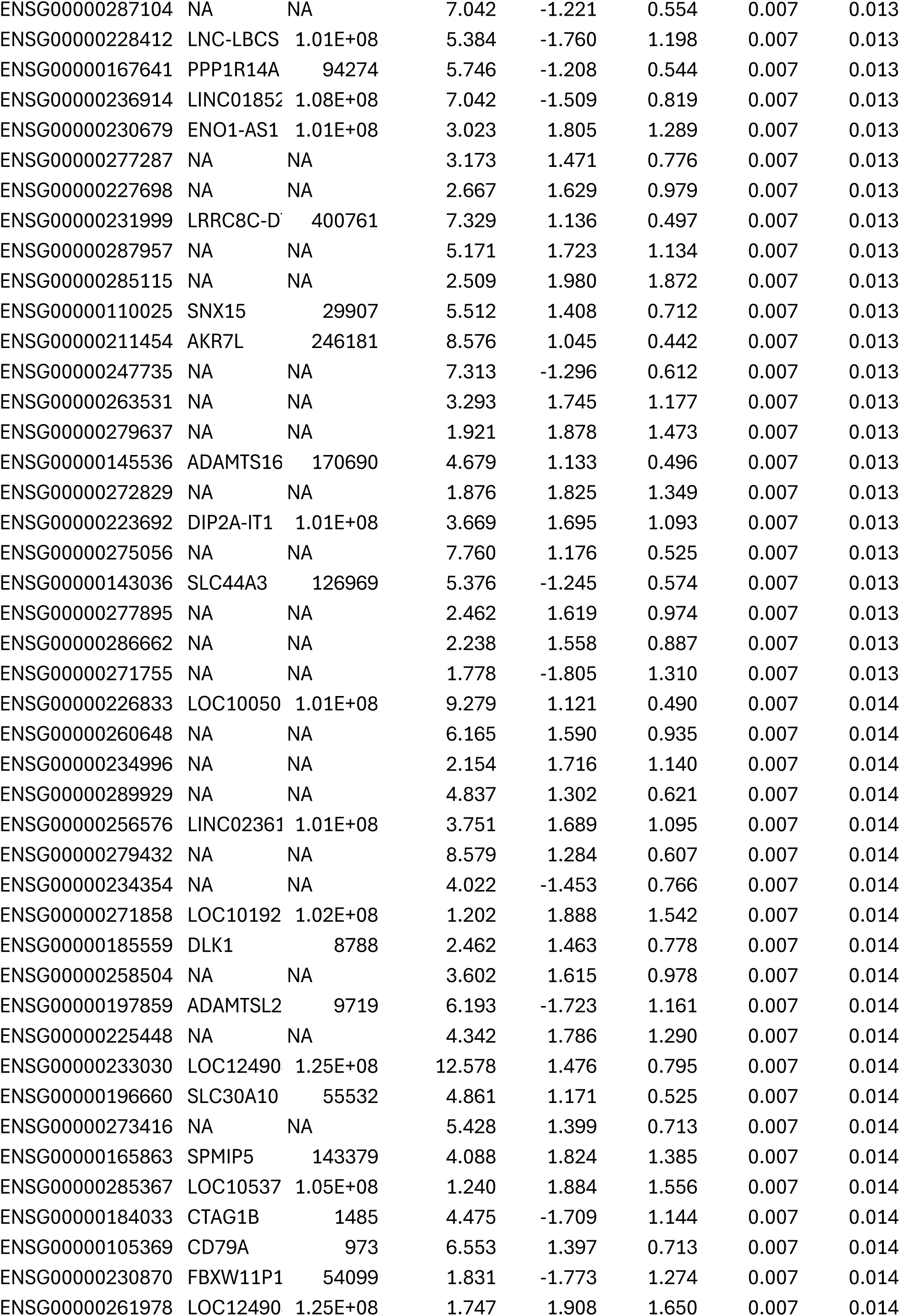
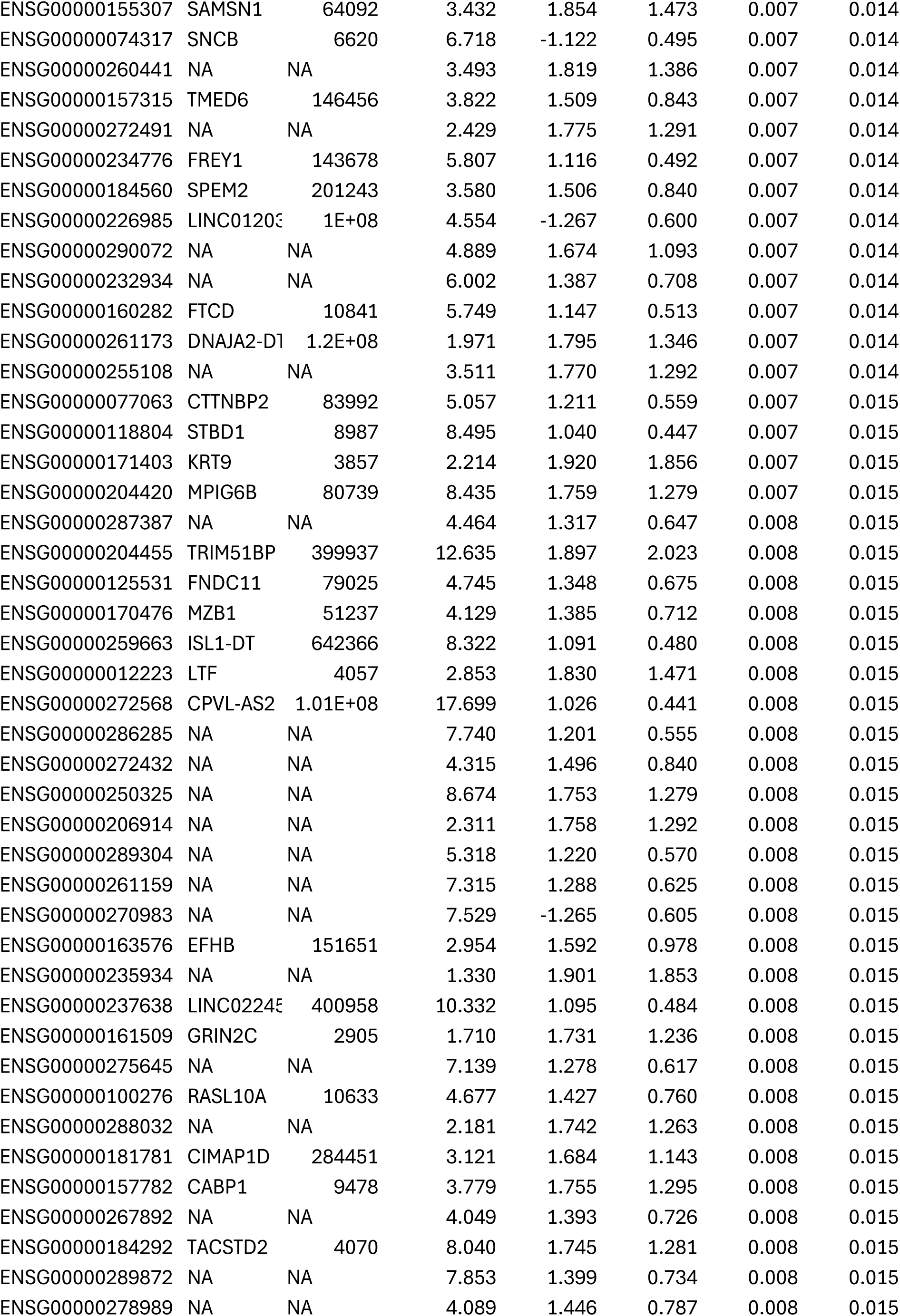
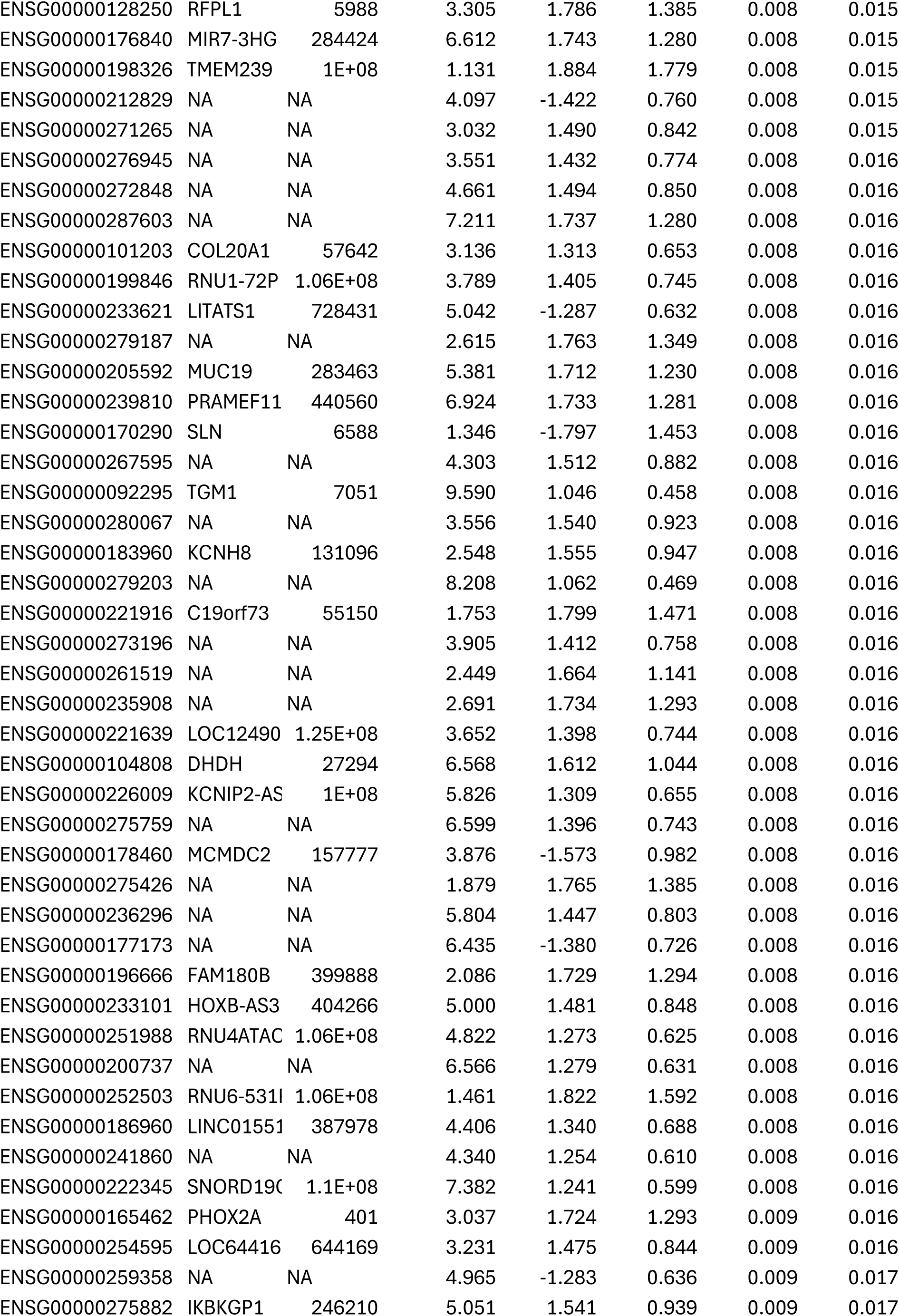
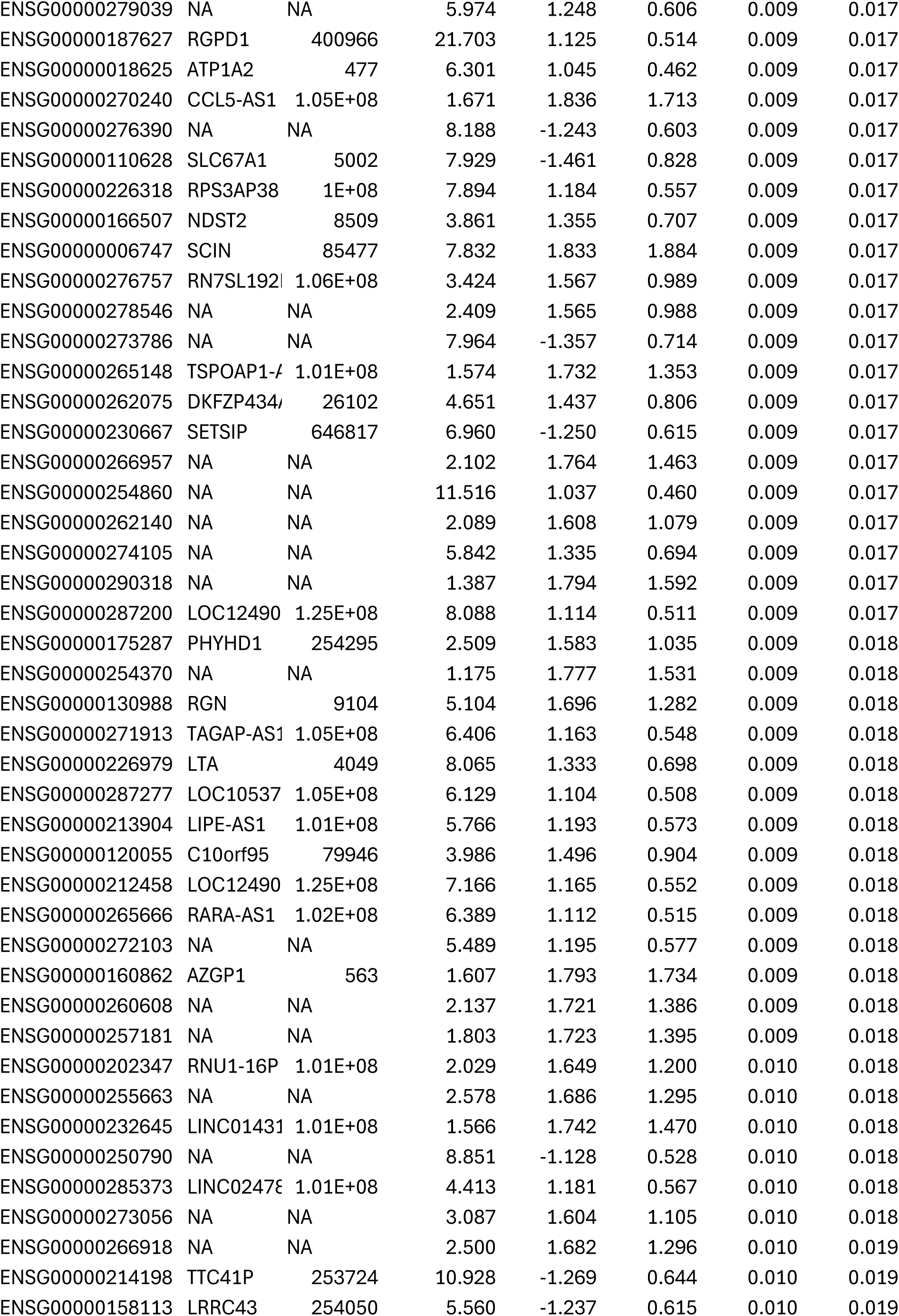
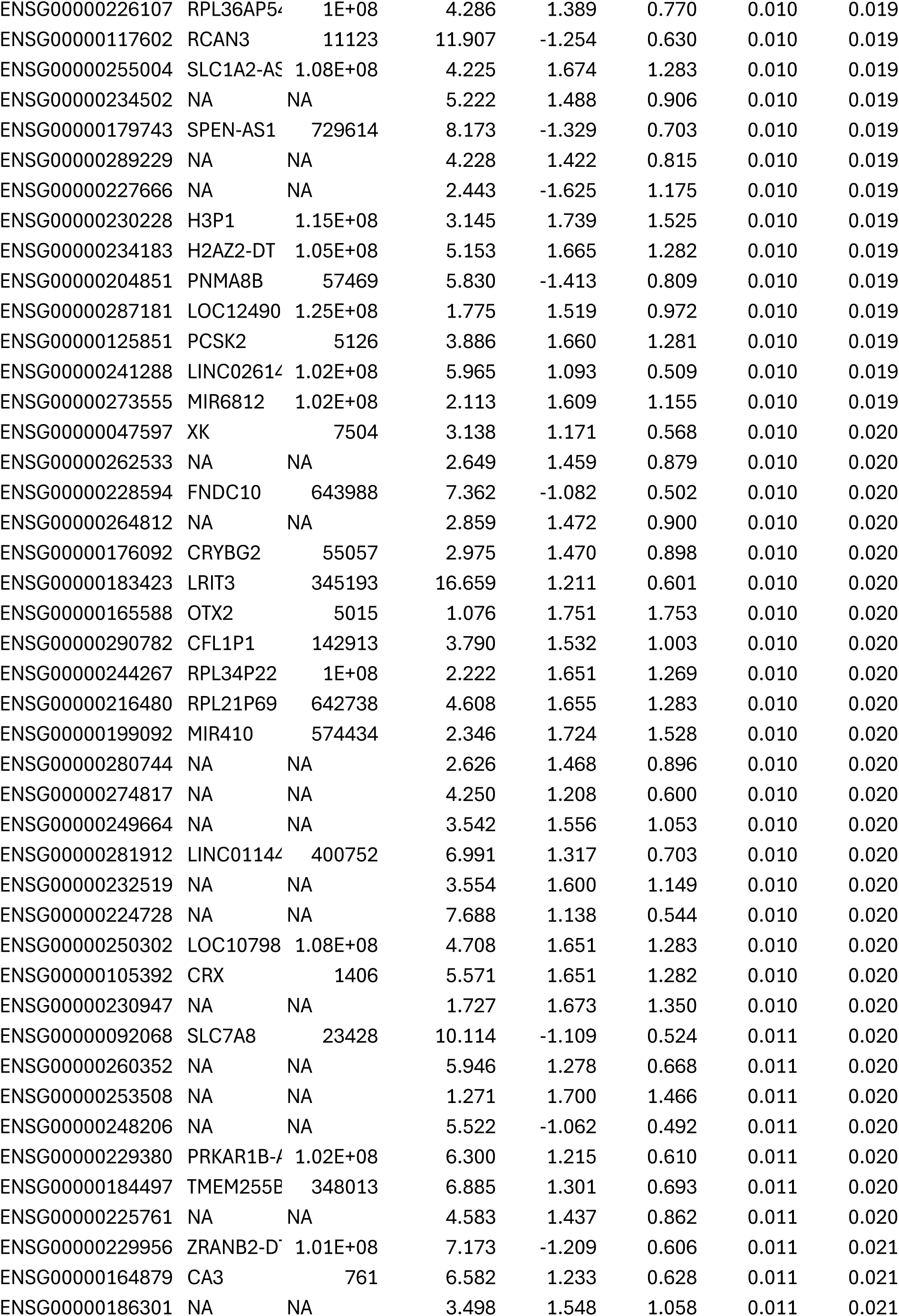
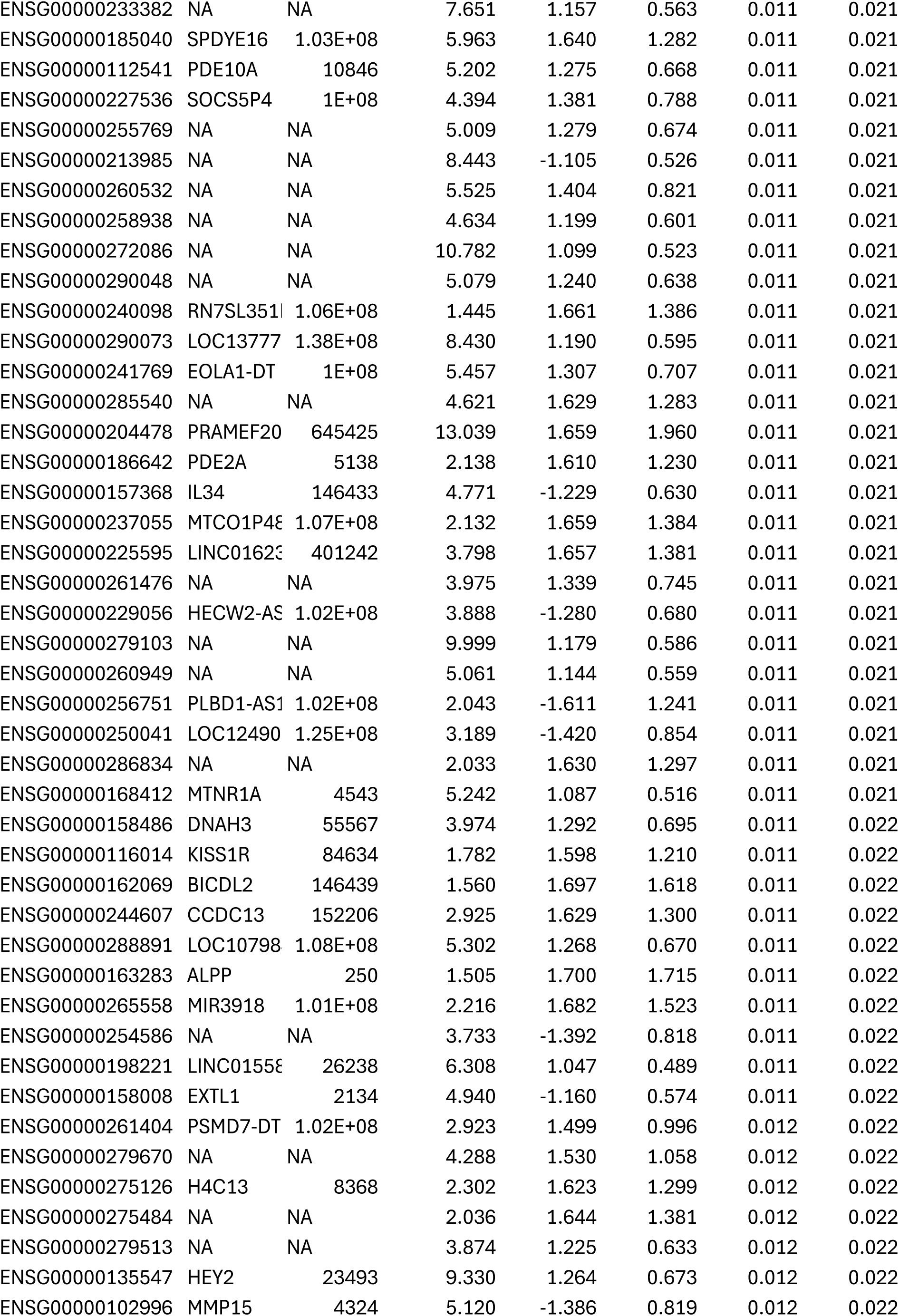
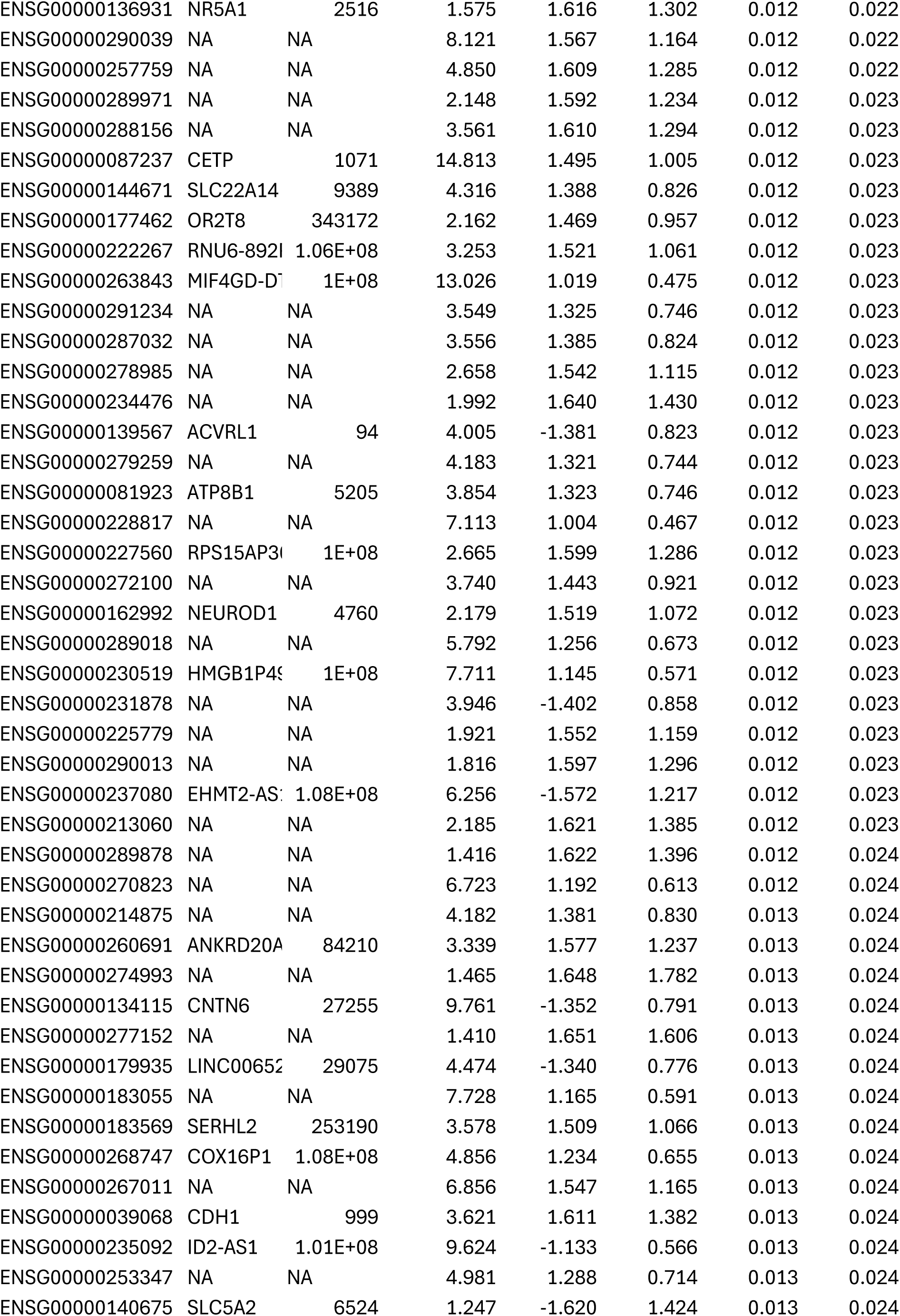
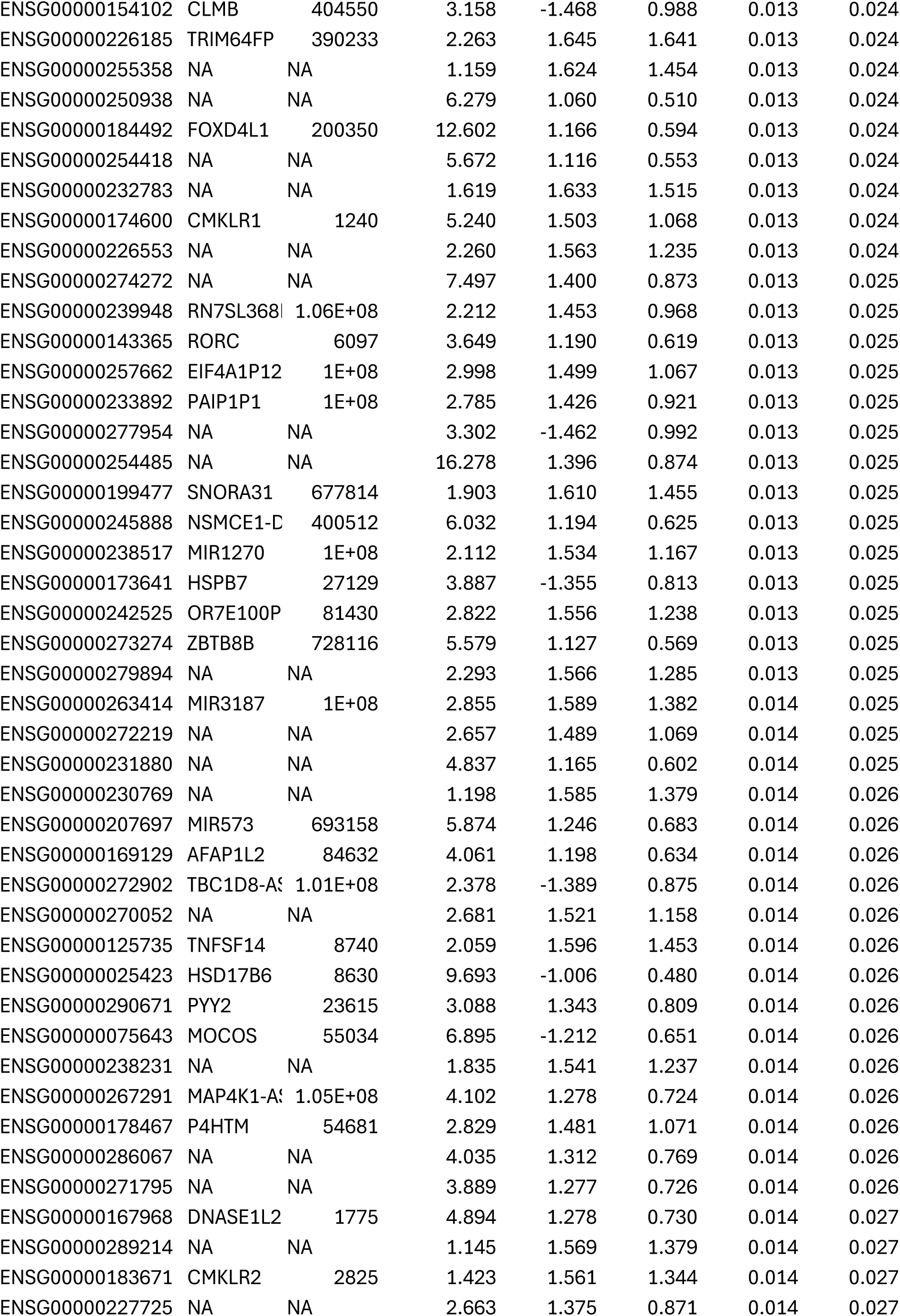
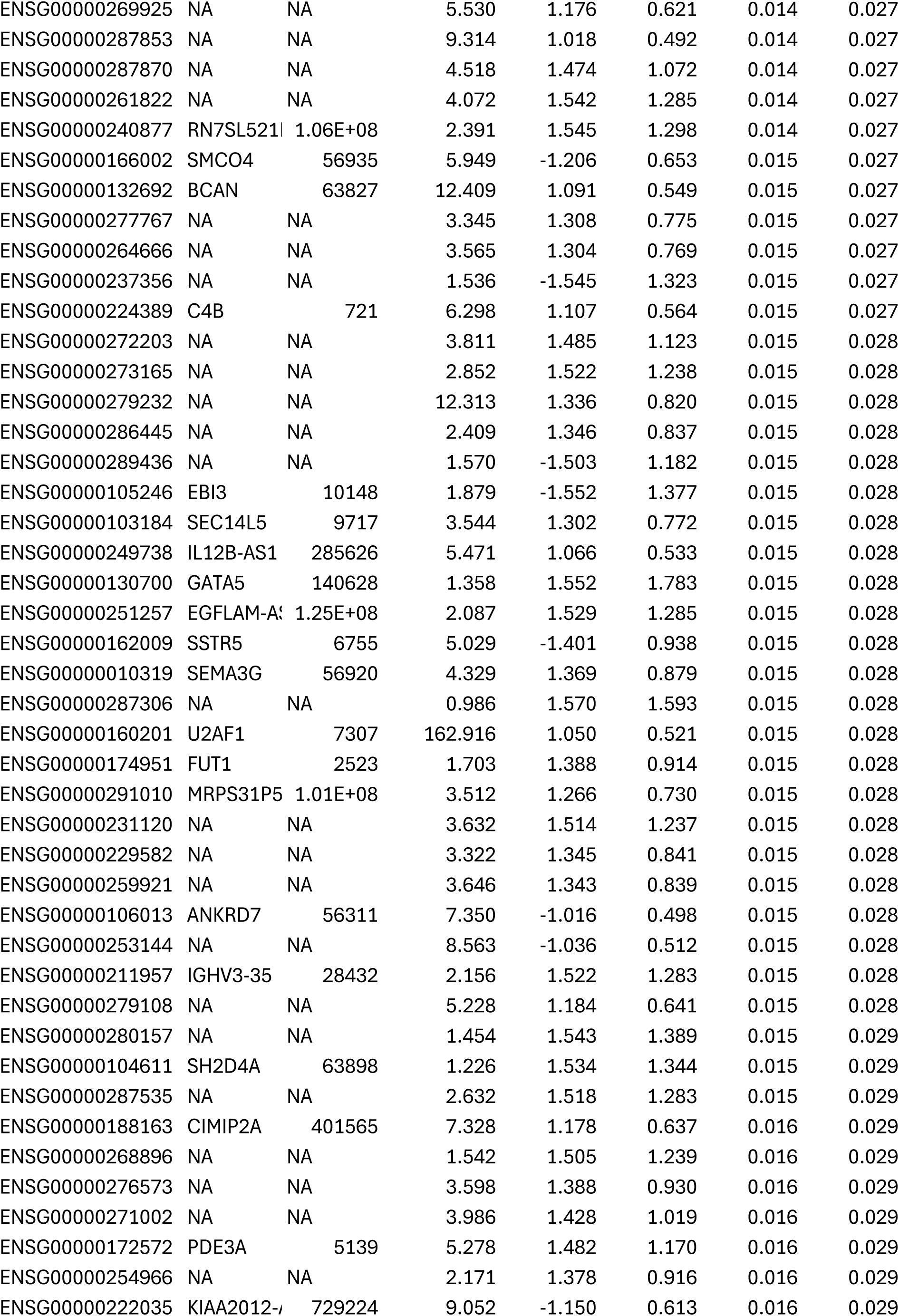
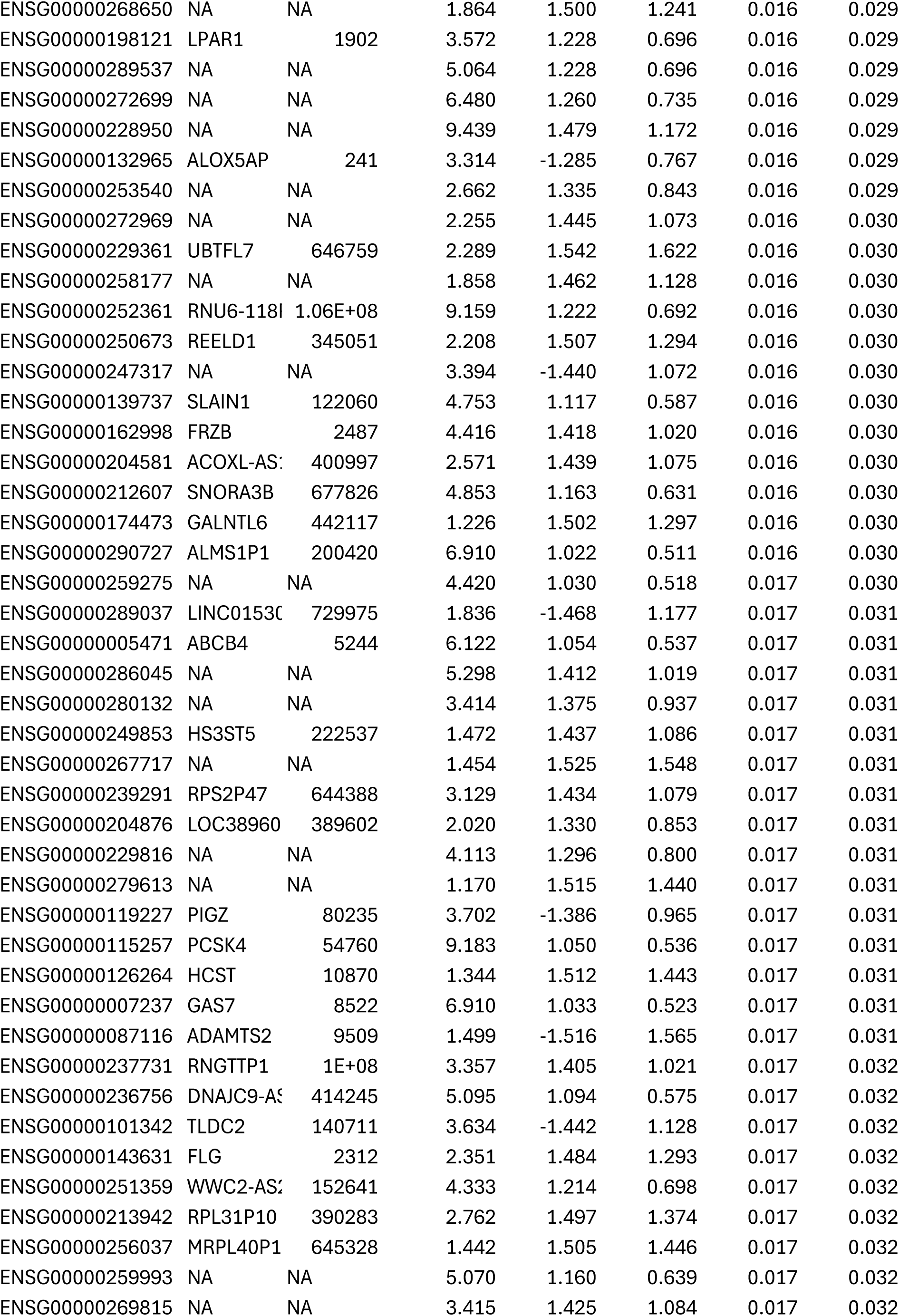
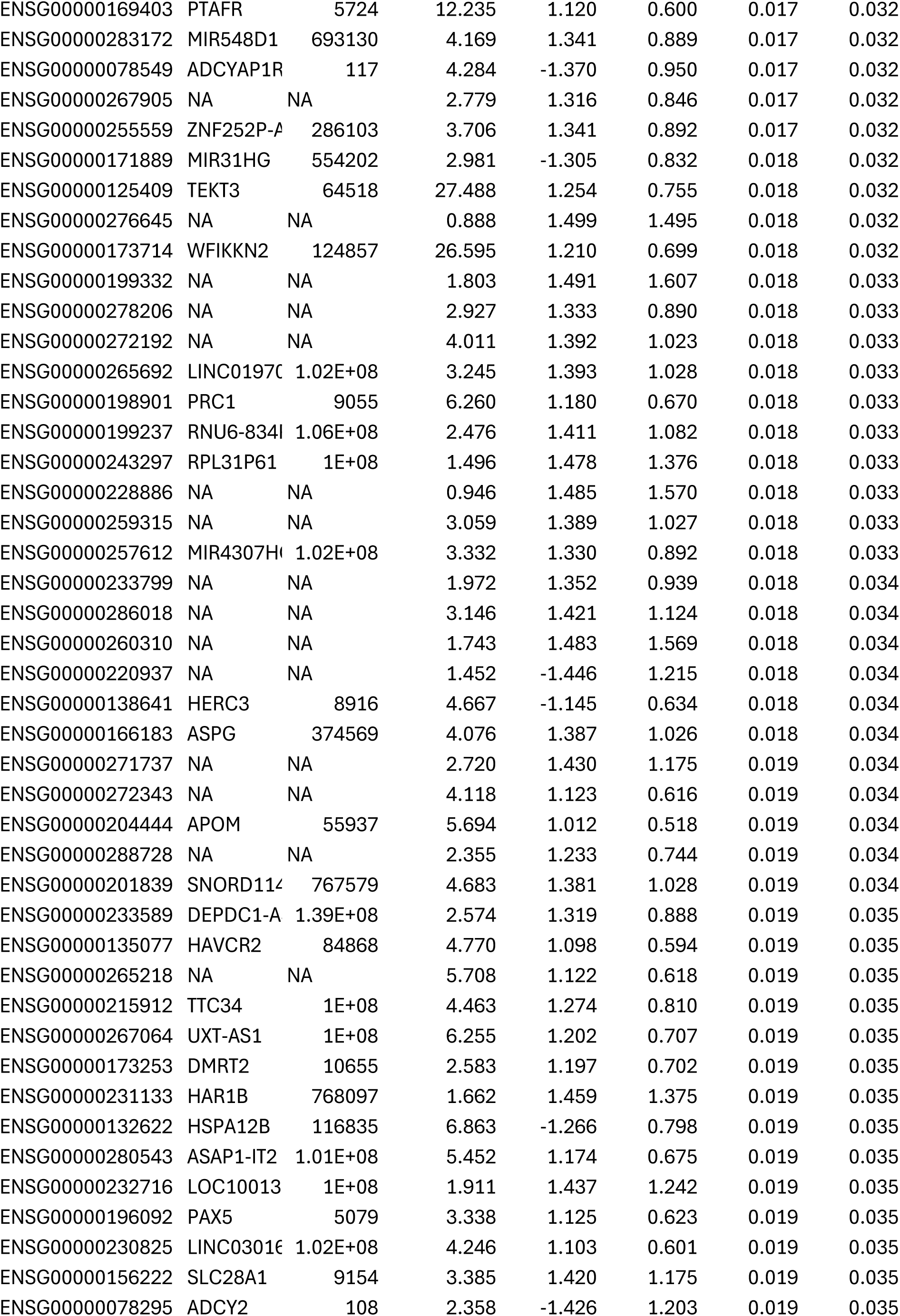
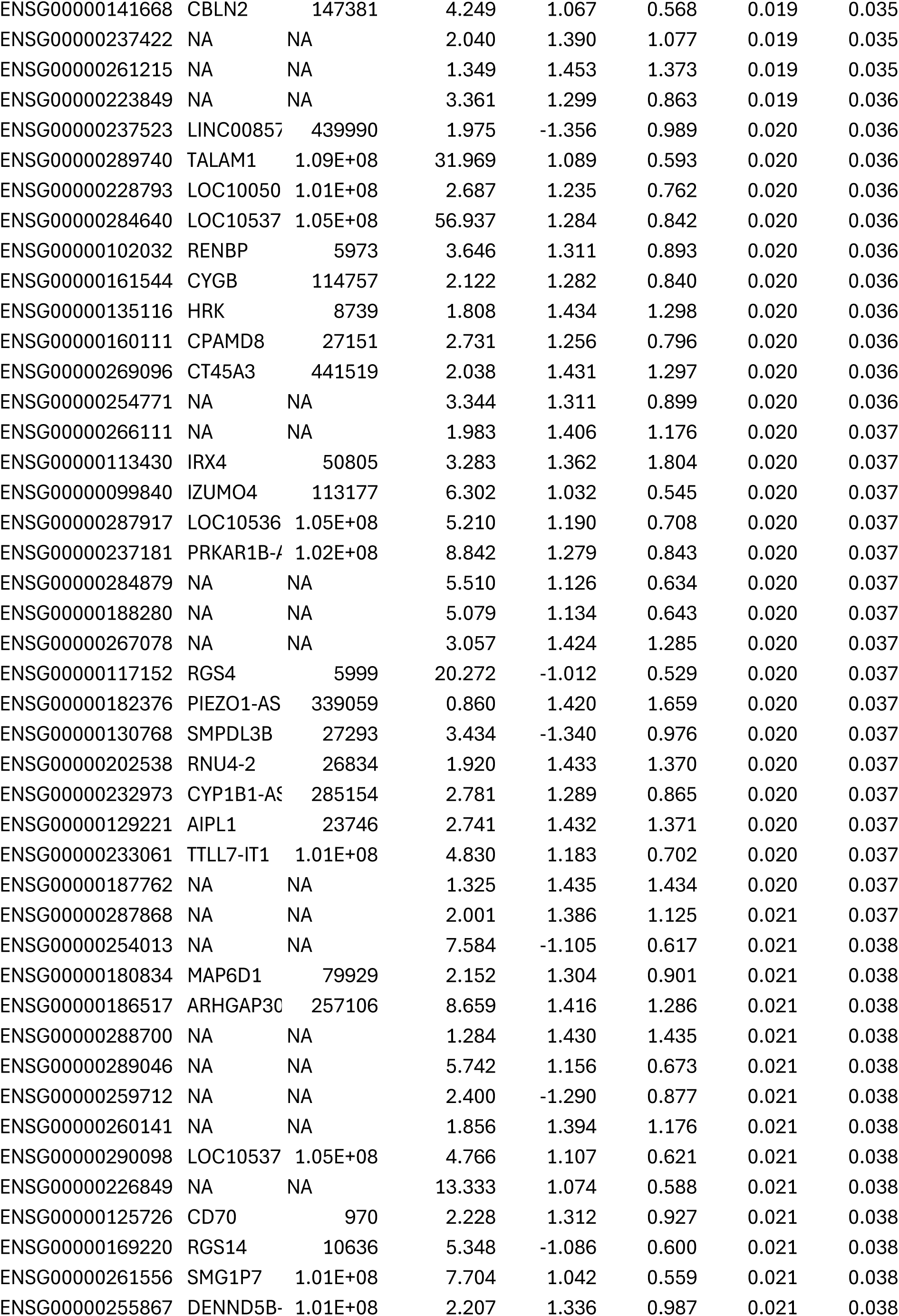
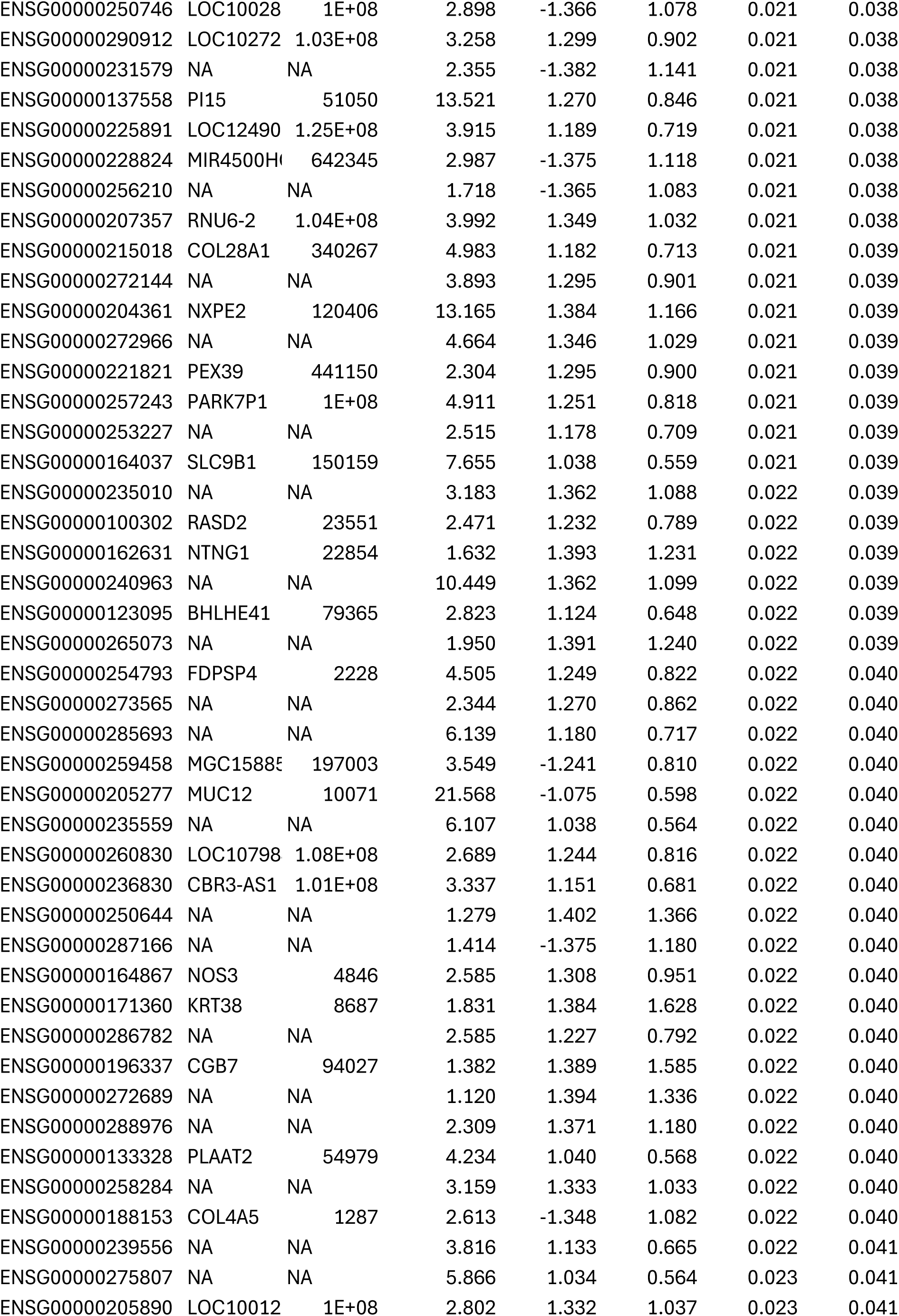
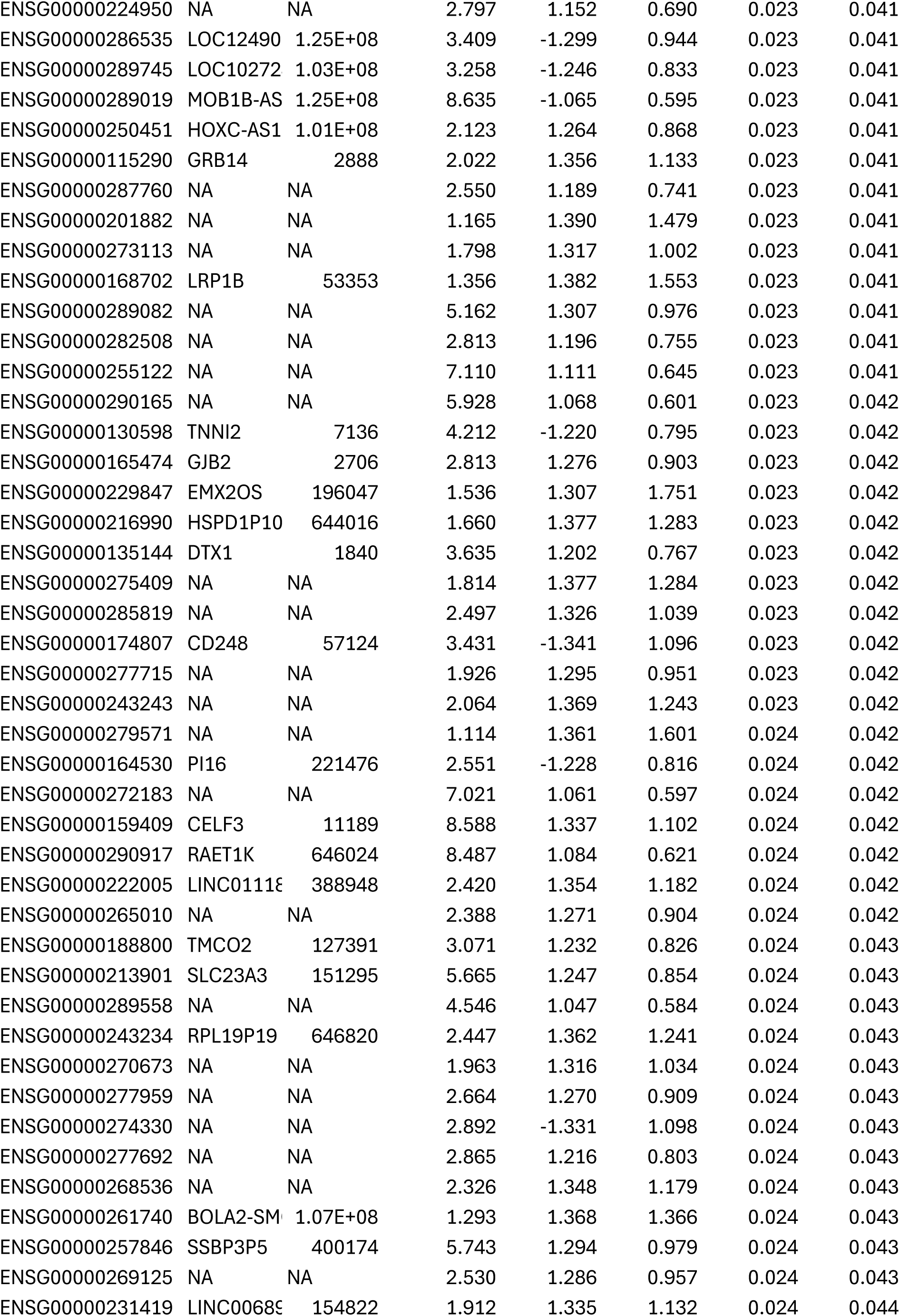
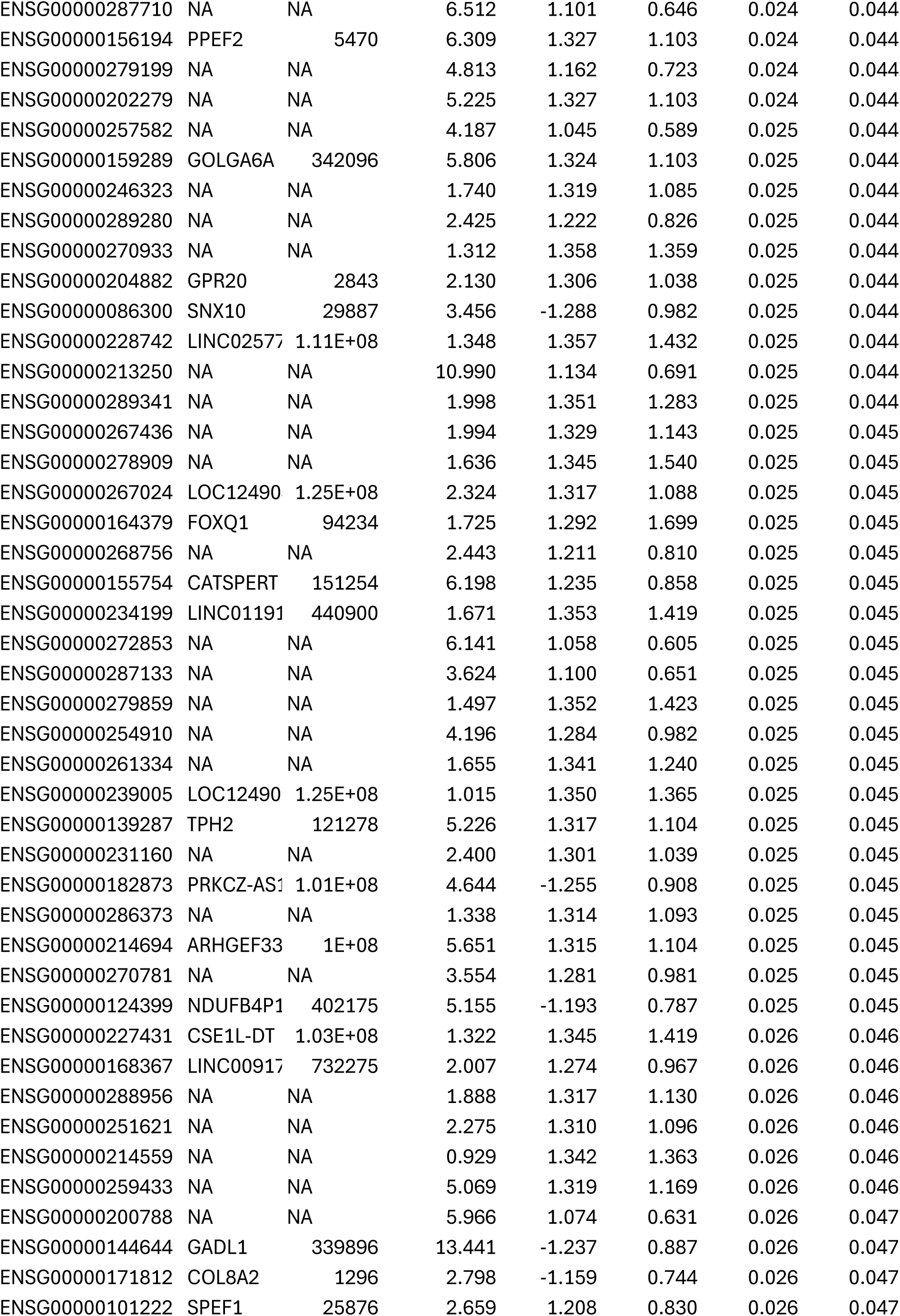
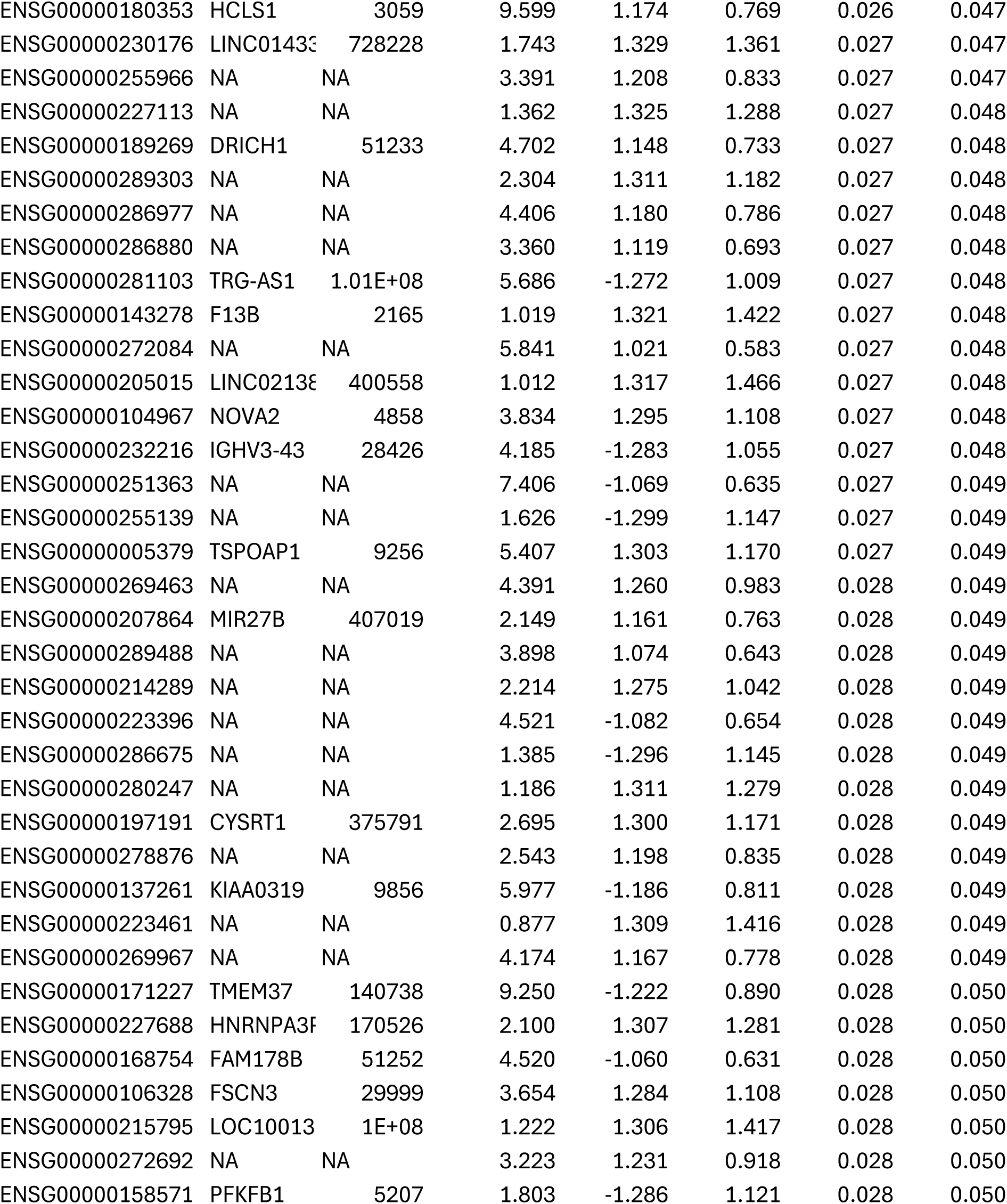
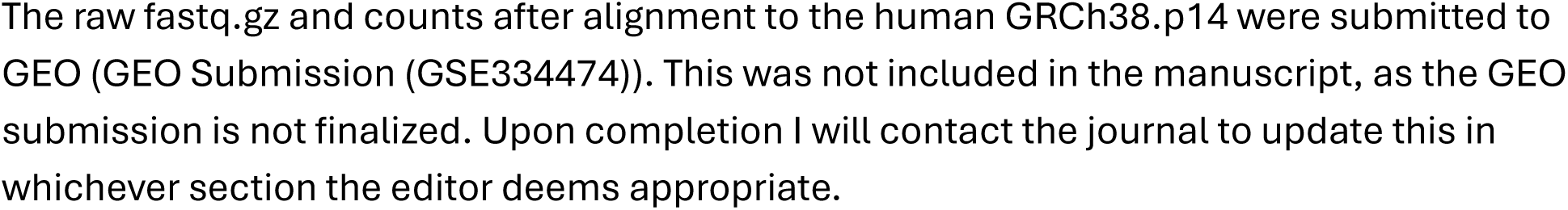

## Notes

### Competing Interest Statement

The authors have declared no competing interest.

## REFERENCES

1. Ognjanovic S, Linabery AM, Charbonneau B, Ross JA. Trends in childhood rhabdomyosarcoma incidence and survival in the United States, 1975-2005. Cancer. 2009;115(18):4218–4226. doi:10.1002/cncr.24465

2. Williamson D, Missiaglia E, De Reyniès A, et al. Fusion gene-negative alveolar rhabdomyosarcoma is clinically and molecularly indistinguishable from embryonal rhabdomyosarcoma. J Clin Oncol. 2010;28(13):2151–2158. doi:10.1200/JCO.2009.26.3814

3. Davicioni E, Anderson MJ, Finckenstein FG, et al. Molecular classification of rhabdomyosarcoma--genotypic and phenotypic determinants of diagnosis: a report from the Children’s Oncology Group. Am J Pathol. 2009;174(2):550–564. doi:10.2353/AJPATH.2009.080631

4. Sorensen PHB, Lynch JC, Qualman SJ, et al. PAX3-FKHR and PAX7-FKHR gene fusions are prognostic indicators in alveolar rhabdomyosarcoma: a report from the children’s oncology group. J Clin Oncol. 2002;20(11):2672–2679. doi:10.1200/JCO.2002.03.137

5. Wei Y, Qin Q, Yan C, et al. Single-cell analysis and functional characterization uncover the stem cell hierarchies and developmental origins of rhabdomyosarcoma. Nat Cancer. 2022;3(8):961–975. doi:10.1038/S43018-022-00414-W

6. Hanna JA, Garcia MR, Lardennois A, et al. PAX3-FOXO1 drives miR-486-5p and represses miR-221 contributing to pathogenesis of alveolar rhabdomyosarcoma. Oncogene. 2018;37(15):1991–2007. doi:10.1038/S41388-017-0081-3

7. Gonda TJ, Ramsay RG. Directly targeting transcriptional dysregulation in cancer. Nat Rev Cancer. 2015;15(11):686–694. doi:10.1038/NRC4018

8. Wachtel M, Schäfer BW. PAX3-FOXO1: Zooming in on an “undruggable” target. Semin Cancer Biol. 2018;50:115–123. doi:10.1016/J.SEMCANCER.2017.11.006

9. Long AH, Morgenstern DA, Leruste A, Bourdeaut F, Davis KL. Checkpoint Immunotherapy in Pediatrics: Here, Gone, and Back Again. American Society of Clinical Oncology Educational Book. 2022;(42):781–794. doi:10.1200/EDBK_349799

10. Casey DL, Wexler LH, Pitter KL, Samstein RM, Slotkin EK, Wolden SL. Genomic Determinants of Clinical Outcomes in Rhabdomyosarcoma. Clin Cancer Res. 2020;26(5):1135–1140. doi:10.1158/1078-0432.CCR-19-2631

11. Todo T, Martuza RL, Rabkin SD, Johnson PA. Oncolytic herpes simplex virus vector with enhanced MHC class I presentation and tumor cell killing. Proc Natl Acad Sci U S A. 2001;98(11):6396–6401. doi:10.1073/PNAS.101136398

12. Wakimoto H, Kesari S, Farrell CJ, et al. Human glioblastoma-derived cancer stem cells: establishment of invasive glioma models and treatment with oncolytic herpes simplex virus vectors. Cancer Res. 2009;69(8):3472–3481. doi:10.1158/0008-5472.CAN-08-3886

13. Mineta T, Rabkin SD, Yazaki T, Hunter WD, Martuza RL. Attenuated multi-mutated herpes simplex virus-1 for the treatment of malignant gliomas. Nat Med. 1995;1(9):938–943. doi:10.1038/NM0995-938

14. Cheema TA, Wakimoto H, Fecci PE, et al. Multifaceted oncolytic virus therapy for glioblastoma in an immunocompetent cancer stem cell model. Proc Natl Acad Sci U S A. 2013;110(29):12006–12011. doi:10.1073/PNAS.1307935110

15. Cao L, Yu Y, Bilke S, et al. Genome-wide identification of PAX3-FKHR binding sites in rhabdomyosarcoma reveals candidate target genes important for development and cancer. Cancer Res. 2010;70(16):6497–6508. doi:10.1158/0008-5472.CAN-10-0582

16. Chalepakis G, Fritsch R, Fickenscher H, Deutsch U, Goulding M, Gruss P. The molecular basis of the undulated/Pax-1 mutation. Cell. 1991;66(5):873–884. doi:10.1016/0092-8674(91)90434-Z

17. Lam PYP, Sublett JE, Hollenbach AD, Roussel MF. The oncogenic potential of the Pax3-FKHR fusion protein requires the Pax3 homeodomain recognition helix but not the Pax3 paired-box DNA binding domain. Mol Cell Biol. 1999;19(1):594–601. doi:10.1128/MCB.19.1.594

18. Anderson MJ, Shelton GD, Cavenee WK, Arden KC. Embryonic expression of the tumor-associated PAX3-FKHR fusion protein interferes with the developmental functions of Pax3. Proc Natl Acad Sci U S A. 2001;98(4):1589–1594. doi:10.1073/PNAS.98.4.1589

19. Bennicelli JL, Edwards RH, Barr FG. Mechanism for transcriptional gain of function resulting from chromosomal translocation in alveolar rhabdomyosarcoma. Proc Natl Acad Sci U S A. 1996;93(11):5455–5459. doi:10.1073/PNAS.93.11.5455

20. Yoshida H, Sato-Dahlman M, Hajeri P, et al. Mutant myogenin promoter-controlled oncolytic adenovirus selectively kills PAX3-FOXO1-positive rhabdomyosarcoma cells. Transl Oncol. 2021;14(2). doi:10.1016/J.TRANON.2020.100997

21. Ullman-Culleré MH, Foltz CJ. Body condition scoring: a rapid and accurate method for assessing health status in mice. Lab Anim Sci. 1999;49(3):319–323. Accessed May 17, 2026. http://www.ncbi.nlm.nih.gov/pubmed/10403450

22. Bankhead P, Loughrey MB, Fernández JA, et al. QuPath: Open source software for digital pathology image analysis. Sci Rep. 2017;7(1). doi:10.1038/S41598-017-17204-5

23. Pandey A, Saggese P, Soto A, Gomez E, Alcaraz M, Scafoglio C. Long Non-Coding RNAs Contribute to Glucose Starvation-Induced Dedifferentiation in Lung Adenocarcinoma. Biomolecules. 2025;15(11):1493. doi:10.3390/BIOM15111493/S1

24. Chen CY, Wang PY, Hutzen B, et al. Cooperation of Oncolytic Herpes Virotherapy and PD-1 Blockade in Murine Rhabdomyosarcoma Models. Sci Rep. 2017;7(1). doi:10.1038/S41598-017-02503-8

25. Gambini E, Reisoli E, Appolloni I, et al. Replication-competent herpes simplex virus retargeted to HER2 as therapy for high-grade glioma. Mol Ther. 2012;20(5):994–1001. doi:10.1038/MT.2012.22

26. Kambara H, Okano H, Chiocca EA, Saeki Y. An oncolytic HSV-1 mutant expressing ICP34.5 under control of a nestin promoter increases survival of animals even when symptomatic from a brain tumor. Cancer Res. 2005;65(7):2832–2839. doi:10.1158/0008-5472.CAN-04-3227

27. Mazzacurati L, Marzulli M, Reinhart B, et al. Use of miRNA response sequences to block off-target replication and increase the safety of an unattenuated, glioblastoma-targeted oncolytic HSV. Mol Ther. 2015;23(1):99–107. doi:10.1038/MT.2014.177

28. Keller C, Arenkiel BR, Coffin CM, El-Bardeesy N, DePinho RA, Capecchi MR. Alveolar rhabdomyosarcomas in conditional Pax3:Fkhr mice: cooperativity of Ink4a/ARF and Trp53 loss of function. Genes Dev. 2004;18(21):2614–2626. doi:10.1101/GAD.1244004

29. Imle R, Blösel D, Kommoss FKF, et al. Somatic gene delivery faithfully recapitulates a molecular spectrum of high-risk sarcomas. Nat Commun. 2025;16(1). doi:10.1038/S41467-025-60519-5

30. Leddon JL, Chen CY, Currier MA, et al. Oncolytic HSV virotherapy in murine sarcomas differentially triggers an antitumor T-cell response in the absence of virus permissivity. Mol Ther Oncolytics. 2015;1. doi:10.1038/MTO.2014.10

31. Meadors JL, Cui Y, Chen Q, et al. Murine rhabdomyosarcoma is immunogenic and responsive to T-cell-based immunotherapy. Pediatr Blood Cancer. 2011;57(6):921–929. doi:10.1002/pbc.23048

32. Cleary MM, Mansoor A, Settelmeyer T, et al. NFκB signaling in alveolar rhabdomyosarcoma. Dis Model Mech. 2017;10(9):1109–1115. doi:10.1242/DMM.030882

33. Huska JD, Doyle KJ, Purkal JJ, et al. Targeting tumor intrinsic TAK1 engages TNF-α-driven cell death through distinct mechanisms and enhances cancer immunotherapy. Cell Death Dis. 2025;16(1). doi:10.1038/S41419-025-08013-0

34. Vanlangenakker N, Vanden Berghe T, Bogaert P, et al. cIAP1 and TAK1 protect cells from TNF-induced necrosis by preventing RIP1/RIP3-dependent reactive oxygen species production. Cell Death Differ. 2011;18(4):656–665. doi:10.1038/CDD.2010.138

35. Struzik J, Szulc-Dąbrowska L. NF-κB Signaling in Targeting Tumor Cells by Oncolytic Viruses-Therapeutic Perspectives. Cancers (Basel). 2018;10(11). doi:10.3390/CANCERS10110426

36. Nabarro S, Himoudi N, Papanastasiou A, et al. Coordinated oncogenic transformation and inhibition of host immune responses by the PAX3-FKHR fusion oncoprotein. J Exp Med. 2005;202(10):1399–1410. doi:10.1084/JEM.20050730

