## Supplementary figures and images for "Novel Synthetic Promoter Armed Oncolytic Herpes Simplex Virus For Treatment of PAX3-FOXO1 Positive Rhabdomyosarcoma"

### Supplemental Data 1

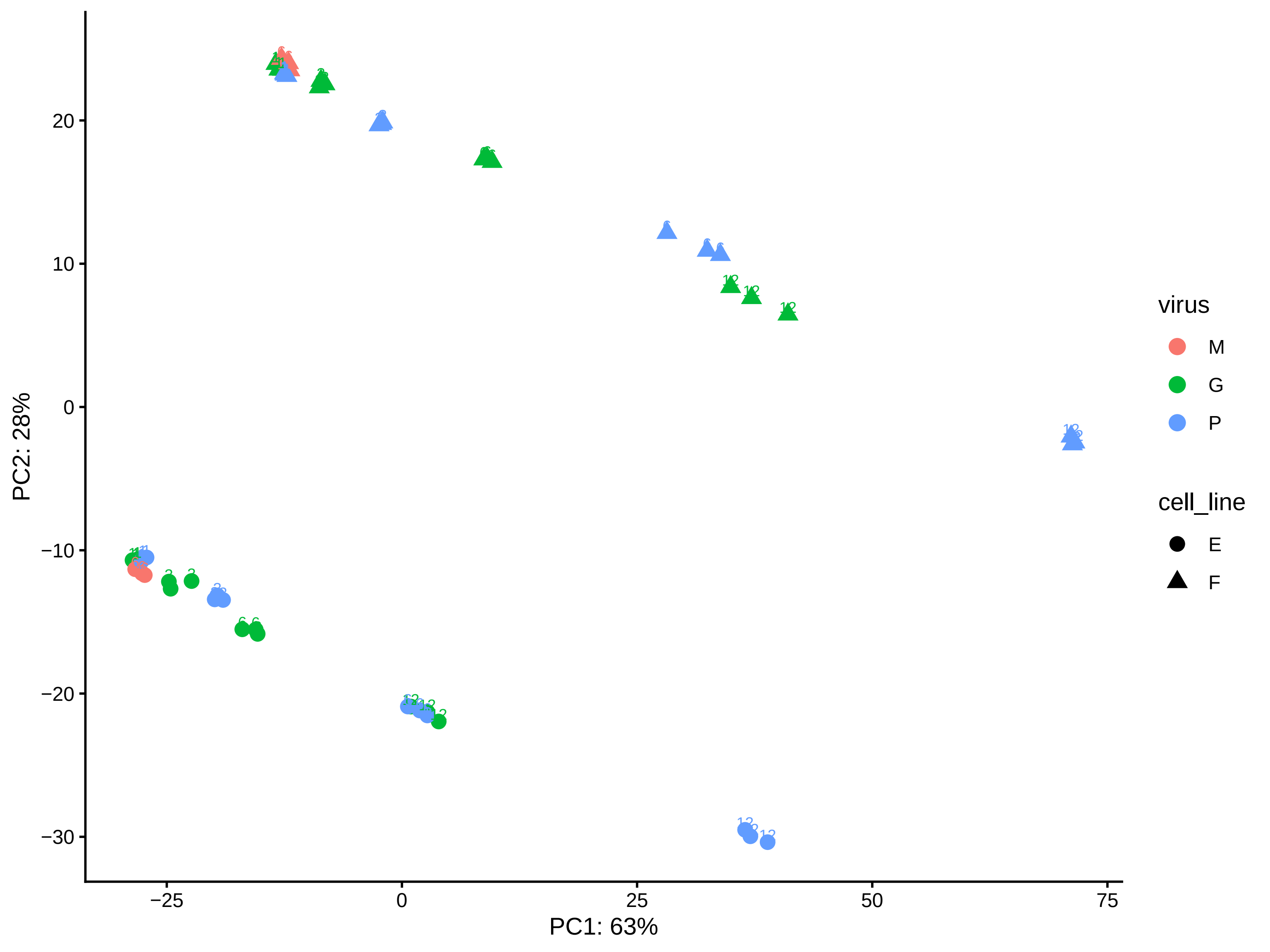
