## Supplemental Data 4 for "Novel Synthetic Promoter Armed Oncolytic Herpes Simplex Virus For Treatment of PAX3-FOXO1 Positive Rhabdomyosarcoma"

**Gene Lists for Heat maps:**

| FIG 7B | NFkB | FIG 7C | NFkB | SFIG7Bi | IFN | SFIG7Bii | IFN | SFIG7Ci | TNF | SFIG7Cii | TNF |
| --- | --- | --- | --- | --- | --- | --- | --- | --- | --- | --- | --- |
| ORDER | GENE | ORDER | GENE | ORDER | GENE | ORDER | GENE | ORDER | GENE | ORDER | GENE |
| 1 | TAB1 | 1 | TNFRSF14 | 1 | CASP9 | 1 | FGF10 | 1 | CASP9 | 1 | NFKB2 |
| 2 | TNFRSF14 | 2 | IKBKG | 2 | RACK1 | 2 | IFNA5 | 2 | DIABLO | 2 | PRKCZ |
| 3 | NFKBIA | 3 | TNFRSF10A | 3 | ACTB | 3 | HLA-DQB1 | 3 | MAP3K14 | 3 | MAP3K8 |
| 4 | RELB | 4 | BIRC3 | 4 | MX2 | 4 | CIITA | 4 | NFKBIB | 4 | RIPK1 |
| 5 | RIPK1 | 5 | CCL2 | 5 | JAK1 | 5 | FGF5 | 5 | JUN | 5 | BTRC |
| 6 | FLOT2 | 6 | CASP8 | 6 | CD70 | 6 | PPP2R5A | 6 | OTUD7B | 6 | SKP1 |
| 7 | HSP90AA1 | 7 | TNF | 7 | RPS27A | 7 | IL18 | 7 | GLUL | 7 | GLUL |
| 8 | UBB | 8 | IKBKB | 8 | TUBB6 | 8 | XAF1 | 8 | PYGL | 8 | OTUD7B |
| 9 | TAB3 | 9 | NFKB2 | 9 | TP53I3 | 9 | LCK | 9 | BCL2L1 | 9 | MAP3K3 |
| 10 | NFKB2 | 10 | REL | 10 | AGFG1 | 10 | TUBA1A | 10 | TXN | 10 | TANK |
| 11 | TRAF2 | 11 | RELA | 11 | TRIM68 | 11 | TNFSF14 | 11 | CASP7 | 11 | TNFAIP3 |
| 12 | IKBKB | 12 | FAS | 12 | CDK1 | 12 | CASP7 | 12 | MAPK8 | 12 | CASP9 |
| 13 | NFKB1 | 13 | BCL3 | 13 | PPM1B | 13 | TRIM46 | 13 | PSMD2 | 13 | KSR1 |
| 14 | XIAP | 14 | NFKBIA | 14 | NUP35 | 14 | PIK3R1 | 14 | TBK1 | 14 | MAP3K1 |
| 15 | NFKBIE | 15 | ITCH | 15 | SMAD7 | 15 | CLK2 | 15 | KRAS | 15 | NFKBIA |
| 16 | STUB1 | 16 | PELI1 | 16 | ABCE1 | 16 | PDE12 | 16 | CUL1 | 16 | IL6 |
| 17 | IKBKG | 17 | NFKBIE | 17 | IFNAR1 | 17 | IFNGR1 | 17 | NSMAF | 17 | MAP3K14 |
| 18 | BIRC3 | 18 | TNFSF10 | 18 | PRMT1 | 18 | IFIT5 | 18 | NCK2 | 18 | JUN |
| 19 | RELA | 19 | TAB3 | 19 | CREM | 19 | FANCE | 19 | MAP3K5 | 19 | MAPK1 |
| 20 | PRKN | 20 | RELB | 20 | PTPN11 | 20 | TRIM68 | 20 | CASP8 | 20 | CASP3 |
| 21 | REL | 21 | RIPK1 | 21 | HINT1 | 21 | PRKCD | 21 | TNFAIP3 | 21 | REL |
| 22 | PDCD6IP | 22 | TAB2 | 22 | CASP7 | 22 | DUS2 | 22 | IL6 | 22 | IKBKB |
| 23 | ITCH | 23 | OGT | 23 | PDCD4 | 23 | TRIM8 | 23 | FADD | 23 | MAP3K5 |
| 24 | TNF | 24 | UBC | 24 | PGGT1B | 24 | PTPRZ1 | 24 | TNFRSF1A | 24 | CASP7 |
| 25 | CCL2 | 25 | TRAF5 | 25 | NDUFB3 | 25 | IRF9 | 25 | CDC37 | 25 | CASP8 |
| 26 | BCL3 | 26 | XIAP | 26 | UBE2N | 26 | IL10RB | 26 | CFLAR | 26 | TNF |
| 27 | FAS | 27 | FADD | 27 | EIF4G2 | 27 | RIGI | 27 | BIRC2 | 27 | TRAF1 |
| 28 | MAP3K7 | 28 | RIPK3 | 28 | HSP90B1 | 28 | TRAIP | 28 | MAPK1 | 28 | BIRC3 |
| 29 | IKBKE | 29 | PRKN | 29 | CD274 | 29 | FNTA | 29 | TAB2 | 29 | CCL2 |
| 30 | CHUK | 30 | CHUK | 30 | HLA-A | 30 | HRK | 30 | MAP3K8 | 30 | IKBKG |
| 31 | TRADD | 31 | MAP3K7 | 31 | GAS6 | 31 | IKBKG | 31 | SOS1 | 31 | KSR2 |
| 32 | BIRC2 | 32 | RPS27A | 32 | PER2 | 32 | CD70 | 32 | CASP3 | 32 | MAP2K3 |
| 33 | TAB2 | 33 | UBB | 33 | ICAM1 | 33 | ERG | 33 | FBXW11 | 33 | TAB2 |
| 34 | TRAF5 | 34 | NFKBIB | 34 | SUMO1 | 34 | IKBKB | 34 | MAP3K7 | 34 | CREBBP |
| 35 | NFKBIB | 35 | CDC37 | 35 | RNASEL | 35 | GRB14 | 35 | MAP2K4 | 35 | RAF1 |
| 36 | UBA52 | 36 | TRADD | 36 | TYK2 | 36 | IL11RA | 36 | RAC1 | 36 | MAP2K7 |
| 37 | TNFRSF10A | 37 | IKBKE | 37 | IL15RA | 37 | MTOR | 37 | BIRC3 | 37 | TAB3 |
| 38 | TNFRSF10B | 38 | FASLG | 38 | IFI44L | 38 | PTPN6 | 38 | BTRC | 38 | TBK1 |
| 39 | PELI1 | 39 | STUB1 | 39 | TUBB3 | 39 | ISG20 | 39 | NRAS | 39 | DIABLO |
| 40 | CASP8 | 40 | TNFRSF1A | 40 | PSMB8 | 40 | GBP3 | 40 | PLK1 | 40 | PPP2CA |
| 41 | UBC | 41 | FLOT2 | 41 | MMP2 | 41 | IFI27 | 41 | IKBKG | 41 | MAP2K4 |
| 42 | MLKL | 42 | MLKL | 42 | OASL | 42 | IRF8 | 42 | MAP2K7 | 42 | MAPK3 |
| 43 | TNFRSF1A | 43 | NFKB1 | 43 | EIF2S3 | 43 | TUBA4A | 43 | CCL2 | 43 | FBXW11 |
| 44 | CFLAR | 44 | TRAF2 | 44 | GBP1 | 44 | ILF2 | 44 | TNF | 44 | CUL1 |
| 45 | CDC37 | 45 | TAB1 | 45 | NFATC2 | 45 | POM121 | 45 | MAP2K6 | 45 | GRB2 |
| 46 | OGT | 46 | HSP90AA1 | 46 | LGALS1 | 46 | NUP160 | 46 | REL | 46 | MAP3K7 |
| 47 | FADD | 47 | UBE2L3 | 47 | TRIM38 | 47 | IRF7 | 47 | BAX | 47 | CSNK2A1 |
| 48 | FASLG | 48 | UBA52 | 48 | PIK3R1 | 48 | FYN | 48 | CHUK | 48 | CHUK |
| 49 | UBE2L3 | 49 | BIRC2 | 49 | TRIM14 | 49 | SFPQ | 49 | MAP4K2 | 49 | MAPK8 |
| 50 | RPS27A | 50 | TNFRSF10B | 50 | UBC | 50 | ACTB | 50 | TRADD | 50 | CDC37 |
| 51 | FLOT1 | 51 | SDCBP | 51 | GBP5 | 51 | RAF1 | 51 | SKP1 | 51 | NFKBIB |
| 52 | SDCBP | 52 | PDCD6IP | 52 | CAMK2D | 52 | JAK2 | 52 | TRAF1 | 52 | RBCK1 |
|  |  | 53 | CFLAR | 53 | MORC3 | 53 | TRIM29 | 53 | PRKCZ | 53 | SMPD2 |
|  |  | 54 | FLOT1 | 54 | IFIH1 | 54 | IFI44 | 54 | TAB1 | 54 | TAB1 |
|  |  |  |  | 55 | B2M | 55 | KPNA4 | 55 | RAF1 | 55 | CFLAR |
|  |  |  |  | 56 | TUBA1C | 56 | MAP2K3 | 56 | MAP2K3 | 56 | SOS1 |
|  |  |  |  | 57 | ATF6 | 57 | PPP2CB | 57 | PPP2CA | 57 | BIRC2 |
|  |  |  |  | 58 | CD44 | 58 | MID1 | 58 | MAPK3 | 58 | NCK2 |
|  |  |  |  | 59 | IFIT2 | 59 | VDR | 59 | MAP3K1 | 59 | PSMD2 |
|  |  |  |  | 60 | HERC5 | 60 | CAMK2A | 60 | NFKBIA | 60 | APAF1 |
|  |  |  |  | 61 | NUP93 | 61 | FAS | 61 | IKBKB | 61 | NSMAF |
|  |  |  |  | 62 | IFI44 | 62 | RAE1 | 62 | MAP3K3 | 62 | RAC1 |
|  |  |  |  | 63 | YBX1 | 63 | ICAM1 | 63 | RBCK1 | 63 | BAD |
|  |  |  |  | 64 | ANXA2 | 64 | KPNA5 | 64 | APAF1 | 64 | HRAS |
|  |  |  |  | 65 | BAG1 | 65 | HSPA2 | 65 | CREBBP | 65 | PYGL |
|  |  |  |  | 66 | TUBA3E | 66 | MX2 | 66 | BAD | 66 | HSP90AA1 |
|  |  |  |  | 67 | TUBA1A | 67 | ISG15 | 67 | TRAP1 | 67 | TXN |
|  |  |  |  | 68 | COX4I1 | 68 | HSPA1A | 68 | AKT1 | 68 | AKT1 |
|  |  |  |  | 69 | ILF2 | 69 | COX6A1 | 69 | RIPK1 | 69 | TNFRSF1B |
|  |  |  |  | 70 | TXN | 70 | FANCA | 70 | BID | 70 | PLK1 |
|  |  |  |  | 71 | OAS2 | 71 | NUP214 | 71 | HSP90AA1 | 71 | NRAS |
|  |  |  |  | 72 | IGFBP6 | 72 | CDKN1C | 72 | TNFRSF1B | 72 | BAX |
|  |  |  |  | 73 | GBP3 | 73 | MAP3K14 | 73 | MADD | 73 | MAPK9 |
|  |  |  |  | 74 | SP100 | 74 | CBL | 74 | MAPK9 | 74 | TRADD |
|  |  |  |  | 75 | VDR | 75 | FOSL2 | 75 | GRB2 | 75 | MAP2K6 |
|  |  |  |  | 76 | S100A2 | 76 | HLA-B | 76 | NCK1 | 76 | KRAS |
|  |  |  |  | 77 | IFIT3 | 77 | DST | 77 | NFKBIE | 77 | TRAF2 |
|  |  |  |  | 78 | SP110 | 78 | NUP35 | 78 | CSNK2A1 | 78 | MAP4K2 |
|  |  |  |  | 79 | BECN1 | 79 | CRKL | 79 | TANK | 79 | BCL2L1 |
|  |  |  |  | 80 | PRDX6 | 80 | PGGT1B | 80 | CYBA | 80 | BID |
|  |  |  |  | 81 | MAPK10 | 81 | TUBB4A | 81 | NFKB2 | 81 | TNFRSF1A |
|  |  |  |  | 82 | JAK2 | 82 | SKP1 | 82 | SMPD2 | 82 | NFKB1 |
|  |  |  |  | 83 | NUP107 | 83 | FLNB | 83 | HRAS | 83 | TRAP1 |
|  |  |  |  | 84 | PRKCA | 84 | POM121C | 84 | TRAF2 | 84 | CYBA |
|  |  |  |  | 85 | ISG20 | 85 | IP6K2 | 85 | TAB3 | 85 | FADD |
|  |  |  |  | 86 | RPLP1 | 86 | SFN | 86 | KSR1 | 86 | NFKBIE |
|  |  |  |  | 87 | PPP2CB | 87 | PIAS1 | 87 | NFKB1 | 87 | MADD |
|  |  |  |  | 88 | TPM2 | 88 | KPNA1 |  |  | 88 | NCK1 |
|  |  |  |  | 89 | DNAJC3 | 89 | ACTG1 |  |  | 89 | RIPK3 |
|  |  |  |  | 90 | HLA-B | 90 | EGR1 |  |  |  |  |
|  |  |  |  | 91 | CRKL | 91 | TP53 |  |  |  |  |
|  |  |  |  | 92 | MID1 | 92 | UBC |  |  |  |  |
|  |  |  |  | 93 | COX6C | 93 | HLA-E |  |  |  |  |
|  |  |  |  | 94 | PIAS3 | 94 | TUBB3 |  |  |  |  |
|  |  |  |  | 95 | NUP188 | 95 | FKBP5 |  |  |  |  |
|  |  |  |  | 96 | IRS1 | 96 | TUBB8 |  |  |  |  |
|  |  |  |  | 97 | KDM5A | 97 | SEC61B |  |  |  |  |
|  |  |  |  | 98 | NUP85 | 98 | IRS1 |  |  |  |  |
|  |  |  |  | 99 | TRIM8 | 99 | TRIM35 |  |  |  |  |
|  |  |  |  | 100 | SNCA | 100 | IFIT1 |  |  |  |  |
|  |  |  |  | 101 | FGF5 | 101 | IFIT3 |  |  |  |  |
|  |  |  |  | 102 | SOCS3 | 102 | OASL |  |  |  |  |
|  |  |  |  | 103 | ADAR | 103 | TRIM62 |  |  |  |  |
|  |  |  |  | 104 | HLA-C | 104 | RGS2 |  |  |  |  |
|  |  |  |  | 105 | RAPGEF1 | 105 | PML |  |  |  |  |
|  |  |  |  | 106 | GBP2 | 106 | IFIT2 |  |  |  |  |
|  |  |  |  | 107 | PPP2R1A | 107 | PER2 |  |  |  |  |
|  |  |  |  | 108 | ISG15 | 108 | TGIF1 |  |  |  |  |
|  |  |  |  | 109 | FURIN | 109 | EIF4A1 |  |  |  |  |
|  |  |  |  | 110 | FTL | 110 | IRF1 |  |  |  |  |
|  |  |  |  | 111 | GSTO1 | 111 | PDGFA |  |  |  |  |
|  |  |  |  | 112 | UBA52 | 112 | NUP42 |  |  |  |  |
|  |  |  |  | 113 | PRDX1 | 113 | TUBB6 |  |  |  |  |
|  |  |  |  | 114 | TGIF1 | 114 | IFNL1 |  |  |  |  |
|  |  |  |  | 115 | TRIM21 | 115 | TRIM3 |  |  |  |  |
|  |  |  |  | 116 | IL18 | 116 | PTAFR |  |  |  |  |
|  |  |  |  | 117 | EIF4A2 | 117 | NUP54 |  |  |  |  |
|  |  |  |  | 118 | HLA-DPB1 | 118 | USP18 |  |  |  |  |
|  |  |  |  | 119 | UBA7 | 119 | MAPK1 |  |  |  |  |
|  |  |  |  | 120 | FOSL2 | 120 | MX1 |  |  |  |  |
|  |  |  |  | 121 | STAT2 | 121 | EIF4A2 |  |  |  |  |
|  |  |  |  | 122 | IRS2 | 122 | IRS2 |  |  |  |  |
|  |  |  |  | 123 | TPR | 123 | IRF4 |  |  |  |  |
|  |  |  |  | 124 | PIK3CD | 124 | TUBB2B |  |  |  |  |
|  |  |  |  | 125 | ENO3 | 125 | MORC3 |  |  |  |  |
|  |  |  |  | 126 | RSAD2 | 126 | HSPA1B |  |  |  |  |
|  |  |  |  | 127 | TUBA3D | 127 | TUBB2A |  |  |  |  |
|  |  |  |  | 128 | FASLG | 128 | ARIH1 |  |  |  |  |
|  |  |  |  | 129 | TP53 | 129 | RSAD2 |  |  |  |  |
|  |  |  |  | 130 | OAS1 | 130 | IFIH1 |  |  |  |  |
|  |  |  |  | 131 | TRIM6 | 131 | HERC5 |  |  |  |  |
|  |  |  |  | 132 | FANCL | 132 | SAMHD1 |  |  |  |  |
|  |  |  |  | 133 | RPGR | 133 | STAT2 |  |  |  |  |
|  |  |  |  | 134 | RPS6KA4 | 134 | TRIM26 |  |  |  |  |
|  |  |  |  | 135 | USP18 | 135 | IGF2 |  |  |  |  |
|  |  |  |  | 136 | IFNAR2 | 136 | NUP153 |  |  |  |  |
|  |  |  |  | 137 | SLIRP | 137 | EIF2AK3 |  |  |  |  |
|  |  |  |  | 138 | FNTB | 138 | SMAD7 |  |  |  |  |
|  |  |  |  | 139 | CAMK2A | 139 | TRIM48 |  |  |  |  |
|  |  |  |  | 140 | NUP62 | 140 | EIF4A3 |  |  |  |  |
|  |  |  |  | 141 | NPM1 | 141 | TUBB4B |  |  |  |  |
|  |  |  |  | 142 | KPNA2 | 142 | RPS2 |  |  |  |  |
|  |  |  |  | 143 | NUP37 | 143 | NFATC2 |  |  |  |  |
|  |  |  |  | 144 | RAC1 | 144 | YBX1 |  |  |  |  |
|  |  |  |  | 145 | XRCC5 | 145 | CDK13 |  |  |  |  |
|  |  |  |  | 146 | NUP153 | 146 | PIM1 |  |  |  |  |
|  |  |  |  | 147 | TRIM48 | 147 | AGFG1 |  |  |  |  |
|  |  |  |  | 148 | TRIM17 | 148 | GAB2 |  |  |  |  |
|  |  |  |  | 149 | MAPK1 | 149 | IRF2 |  |  |  |  |
|  |  |  |  | 150 | TGFBR3 | 150 | SOCS1 |  |  |  |  |
|  |  |  |  | 151 | KPNA3 | 151 | BCL3 |  |  |  |  |
|  |  |  |  | 152 | SPHK1 | 152 | DDIT3 |  |  |  |  |
|  |  |  |  | 153 | NUP54 | 153 | SOCS3 |  |  |  |  |
|  |  |  |  | 154 | IFI30 | 154 | TRIM21 |  |  |  |  |
|  |  |  |  | 155 | UBE2E1 | 155 | REL |  |  |  |  |
|  |  |  |  | 156 | PRKRA | 156 | OAS2 |  |  |  |  |
|  |  |  |  | 157 | DUS2 | 157 | MAP3K1 |  |  |  |  |
|  |  |  |  | 158 | FAS | 158 | EIF4E2 |  |  |  |  |
|  |  |  |  | 159 | IFNE | 159 | NFKBIA |  |  |  |  |
|  |  |  |  | 160 | GBP4 | 160 | RPS6KB1 |  |  |  |  |
|  |  |  |  | 161 | IFI35 | 161 | EIF4E |  |  |  |  |
|  |  |  |  | 162 | TP53BP2 | 162 | CASP9 |  |  |  |  |
|  |  |  |  | 163 | NUP88 | 163 | CD274 |  |  |  |  |
|  |  |  |  | 164 | SEC13 | 164 | IFNE |  |  |  |  |
|  |  |  |  | 165 | CALM2 | 165 | TRIM17 |  |  |  |  |
|  |  |  |  | 166 | RPS6KB1 | 166 | NCK1 |  |  |  |  |
|  |  |  |  | 167 | RAP1A | 167 | BST2 |  |  |  |  |
|  |  |  |  | 168 | NUP58 | 168 | TRIM45 |  |  |  |  |
|  |  |  |  | 169 | PDE12 | 169 | EPHB3 |  |  |  |  |
|  |  |  |  | 170 | HLA-H | 170 | CRK |  |  |  |  |
|  |  |  |  | 171 | HSPA1L | 171 | FANCF |  |  |  |  |
|  |  |  |  | 172 | TRIM45 | 172 | RANBP2 |  |  |  |  |
|  |  |  |  | 173 | NUP205 | 173 | UBA7 |  |  |  |  |
|  |  |  |  | 174 | IRF5 | 174 | IRF5 |  |  |  |  |
|  |  |  |  | 175 | RANBP2 | 175 | TRIM22 |  |  |  |  |
|  |  |  |  | 176 | DST | 176 | IFNLR1 |  |  |  |  |
|  |  |  |  | 177 | NUP210 | 177 | FAAP24 |  |  |  |  |
|  |  |  |  | 178 | TRIM3 | 178 | VCAM1 |  |  |  |  |
|  |  |  |  | 179 | FANCA | 179 | IGFBP3 |  |  |  |  |
|  |  |  |  | 180 | IRF4 | 180 | RNASEL |  |  |  |  |
|  |  |  |  | 181 | EIF2AK3 | 181 | FANCM |  |  |  |  |
|  |  |  |  | 182 | HSPA8 | 182 | MC4R |  |  |  |  |
|  |  |  |  | 183 | DHX9 | 183 | IFNAR2 |  |  |  |  |
|  |  |  |  | 184 | TUBB8B | 184 | SP110 |  |  |  |  |
|  |  |  |  | 185 | PTPRZ1 | 185 | NUP62 |  |  |  |  |
|  |  |  |  | 186 | DDIT3 | 186 | FAAP100 |  |  |  |  |
|  |  |  |  | 187 | TUBA8 | 187 | ENO3 |  |  |  |  |
|  |  |  |  | 188 | COX6A1 | 188 | TUBA8 |  |  |  |  |
|  |  |  |  | 189 | IRF1 | 189 | PIK3CD |  |  |  |  |
|  |  |  |  | 190 | PML | 190 | FAAP20 |  |  |  |  |
|  |  |  |  | 191 | AMPD2 | 191 | IFNGR2 |  |  |  |  |
|  |  |  |  | 192 | MAP3K14 | 192 | RAC1 |  |  |  |  |
|  |  |  |  | 193 | TARBP2 | 193 | IFI6 |  |  |  |  |
|  |  |  |  | 194 | CDKN1C | 194 | TRAF3 |  |  |  |  |
|  |  |  |  | 195 | IL10RB | 195 | E2F1 |  |  |  |  |
|  |  |  |  | 196 | NEDD4 | 196 | ETV4 |  |  |  |  |
|  |  |  |  | 197 | HSPA2 | 197 | PPP2R1A |  |  |  |  |
|  |  |  |  | 198 | TUBB1 | 198 | JAK1 |  |  |  |  |
|  |  |  |  | 199 | TLX1 | 199 | SEC13 |  |  |  |  |
|  |  |  |  | 200 | ZAP70 | 200 | MAPK14 |  |  |  |  |
|  |  |  |  | 201 | TRIM22 | 201 | RPS6KA4 |  |  |  |  |
|  |  |  |  | 202 | IFNL3 | 202 | MT2A |  |  |  |  |
|  |  |  |  | 203 | TRIM29 | 203 | IFI35 |  |  |  |  |
|  |  |  |  | 204 | IGFBP5 | 204 | TRIM2 |  |  |  |  |
|  |  |  |  | 205 | IFNLR1 | 205 | PRKRA |  |  |  |  |
|  |  |  |  | 206 | PIAS1 | 206 | NUP93 |  |  |  |  |
|  |  |  |  | 207 | IKBKG | 207 | BAG1 |  |  |  |  |
|  |  |  |  | 208 | PPP2R5A | 208 | IFITM2 |  |  |  |  |
|  |  |  |  | 209 | SOD1 | 209 | FURIN |  |  |  |  |
|  |  |  |  | 210 | TMSB10 | 210 | NCAM1 |  |  |  |  |
|  |  |  |  | 211 | EIF2AK2 | 211 | ABCE1 |  |  |  |  |
|  |  |  |  | 212 | NUP155 | 212 | COX4I1 |  |  |  |  |
|  |  |  |  | 213 | MT2A | 213 | HINT1 |  |  |  |  |
|  |  |  |  | 214 | BST2 | 214 | NUP43 |  |  |  |  |
|  |  |  |  | 215 | FANCE | 215 | UBE2E1 |  |  |  |  |
|  |  |  |  | 216 | APAF1 | 216 | UBE2I |  |  |  |  |
|  |  |  |  | 217 | MAP2K6 | 217 | EIF4G1 |  |  |  |  |
|  |  |  |  | 218 | REL | 218 | TRIM5 |  |  |  |  |
|  |  |  |  | 219 | KPNA5 | 219 | APAF1 |  |  |  |  |
|  |  |  |  | 220 | EIF4G1 | 220 | MAVS |  |  |  |  |
|  |  |  |  | 221 | HRK | 221 | FGF9 |  |  |  |  |
|  |  |  |  | 222 | STAT3 | 222 | TARBP2 |  |  |  |  |
|  |  |  |  | 223 | TRIM26 | 223 | APOBEC3G | |  |  |  |
|  |  |  |  | 224 | TUBA4B | 224 | TRIM14 |  |  |  |  |
|  |  |  |  | 225 | PTAFR | 225 | CAMK2B |  |  |  |  |
|  |  |  |  | 226 | HSPA5 | 226 | PLCG1 |  |  |  |  |
|  |  |  |  | 227 | BCL2L2 | 227 | CHST15 |  |  |  |  |
|  |  |  |  | 228 | TRAF3 | 228 | SNCA |  |  |  |  |
|  |  |  |  | 229 | EIF2S2 | 229 | MAP2K6 |  |  |  |  |
|  |  |  |  | 230 | PIN1 | 230 | PSMB8 |  |  |  |  |
|  |  |  |  | 231 | FNTA | 231 | IGFBP5 |  |  |  |  |
|  |  |  |  | 232 | PDGFA | 232 | LY6E |  |  |  |  |
|  |  |  |  | 233 | IFNGR2 | 233 | NDC1 |  |  |  |  |
|  |  |  |  | 234 | PPP2R1B | 234 | BRD3 |  |  |  |  |
|  |  |  |  | 235 | NUP160 | 235 | IGFBP6 |  |  |  |  |
|  |  |  |  | 236 | ARIH1 | 236 | SOD1 |  |  |  |  |
|  |  |  |  | 237 | IGFBP3 | 237 | LGALS1 |  |  |  |  |
|  |  |  |  | 238 | E2F1 | 238 | AKT1 |  |  |  |  |
|  |  |  |  | 239 | RXRA | 239 | NUP210 |  |  |  |  |
|  |  |  |  | 240 | IRF3 | 240 | EIF2AK2 |  |  |  |  |
|  |  |  |  | 241 | ETV4 | 241 | TXN |  |  |  |  |
|  |  |  |  | 242 | LY6E | 242 | EIF4EBP1 |  |  |  |  |
|  |  |  |  | 243 | TRIM5 | 243 | BECN1 |  |  |  |  |
|  |  |  |  | 244 | PTPN2 | 244 | SUMO1 |  |  |  |  |
|  |  |  |  | 245 | RPS6 | 245 | HSPA4 |  |  |  |  |
|  |  |  |  | 246 | CRK | 246 | TUBA1B |  |  |  |  |
|  |  |  |  | 247 | IFITM3 | 247 | PRDX1 |  |  |  |  |
|  |  |  |  | 248 | MAPKAPK3 | 248 | COX6C |  |  |  |  |
|  |  |  |  | 249 | TRIM2 | 249 | HSP90B1 |  |  |  |  |
|  |  |  |  | 250 | DNAJA1 | 250 | S100A2 |  |  |  |  |
|  |  |  |  | 251 | EPHB3 | 251 | TRIM25 |  |  |  |  |
|  |  |  |  | 252 | MLST8 | 252 | PDCD4 |  |  |  |  |
|  |  |  |  | 253 | FGF9 | 253 | NDUFB3 |  |  |  |  |
|  |  |  |  | 254 | IFITM2 | 254 | HSPA8 |  |  |  |  |
|  |  |  |  | 255 | FYN | 255 | GSTO1 |  |  |  |  |
|  |  |  |  | 256 | IGF2 | 256 | EIF2S2 |  |  |  |  |
|  |  |  |  | 257 | TUBA1B | 257 | UBE2N |  |  |  |  |
|  |  |  |  | 258 | FTH1 | 258 | TIMP3 |  |  |  |  |
|  |  |  |  | 259 | CHST15 | 259 | MLST8 |  |  |  |  |
|  |  |  |  | 260 | TRIM62 | 260 | TP53I3 |  |  |  |  |
|  |  |  |  | 261 | FANCC | 261 | NUP37 |  |  |  |  |
|  |  |  |  | 262 | MAPT | 262 | MMP2 |  |  |  |  |
|  |  |  |  | 263 | SAMHD1 | 263 | UBE2L6 |  |  |  |  |
|  |  |  |  | 264 | RGS2 | 264 | SEH1L |  |  |  |  |
|  |  |  |  | 265 | TIMP3 | 265 | KPNA3 |  |  |  |  |
|  |  |  |  | 266 | BRD3 | 266 | AAAS |  |  |  |  |
|  |  |  |  | 267 | FKBP5 | 267 | NFKB1 |  |  |  |  |
|  |  |  |  | 268 | UBB | 268 | CENPX |  |  |  |  |
|  |  |  |  | 269 | NUP133 | 269 | FANCG |  |  |  |  |
|  |  |  |  | 270 | UBE2L6 | 270 | RPGR |  |  |  |  |
|  |  |  |  | 271 | TUBA4A | 271 | TUBA3E |  |  |  |  |
|  |  |  |  | 272 | MX1 | 272 | PPP2CA |  |  |  |  |
|  |  |  |  | 273 | NCK1 | 273 | FANCL |  |  |  |  |
|  |  |  |  | 274 | KPNA1 | 274 | HSPA1L |  |  |  |  |
|  |  |  |  | 275 | FANCF | 275 | IL15RA |  |  |  |  |
|  |  |  |  | 276 | STAT5A | 276 | HLA-H |  |  |  |  |
|  |  |  |  | 277 | TRIM25 | 277 | IFI30 |  |  |  |  |
|  |  |  |  | 278 | RPS6KA5 | 278 | ZAP70 |  |  |  |  |
|  |  |  |  | 279 | PIK3CA | 279 | TUBA3D |  |  |  |  |
|  |  |  |  | 280 | ACTG1 | 280 | HLA-F |  |  |  |  |
|  |  |  |  | 281 | NUP50 | 281 | RPS6KA5 |  |  |  |  |
|  |  |  |  | 282 | PIM1 | 282 | TUBB8B |  |  |  |  |
|  |  |  |  | 283 | FANCM | 283 | HLA-A |  |  |  |  |
|  |  |  |  | 284 | KPNB1 | 284 | TUBB1 |  |  |  |  |
|  |  |  |  | 285 | IFNGR1 | 285 | TLX1 |  |  |  |  |
|  |  |  |  | 286 | TUBB2B | 286 | HLA-DPB1 | |  |  |  |
|  |  |  |  | 287 | PRKCD | 287 | EIF4G3 |  |  |  |  |
|  |  |  |  | 288 | EIF4E | 288 | NUP85 |  |  |  |  |
|  |  |  |  | 289 | RPS2 | 289 | NUP88 |  |  |  |  |
|  |  |  |  | 290 | TRIM35 | 290 | KPNB1 |  |  |  |  |
|  |  |  |  | 291 | STAT1 | 291 | TRIM38 |  |  |  |  |
|  |  |  |  | 292 | HSPA4 | 292 | OAS1 |  |  |  |  |
|  |  |  |  | 293 | POM121 | 293 | STAT4 |  |  |  |  |
|  |  |  |  | 294 | TUBB4B | 294 | FTH1 |  |  |  |  |
|  |  |  |  | 295 | CHUK | 295 | FLNA |  |  |  |  |
|  |  |  |  | 296 | NDC1 | 296 | KDM5A |  |  |  |  |
|  |  |  |  | 297 | EIF2S1 | 297 | PTPN1 |  |  |  |  |
|  |  |  |  | 298 | HSPA9 | 298 | NEDD4 |  |  |  |  |
|  |  |  |  | 299 | RAE1 | 299 | GBP1 |  |  |  |  |
|  |  |  |  | 300 | PTPN1 | 300 | PRKCA |  |  |  |  |
|  |  |  |  | 301 | IP6K2 | 301 | SP100 |  |  |  |  |
|  |  |  |  | 302 | EIF4E2 | 302 | IRF3 |  |  |  |  |
|  |  |  |  | 303 | SEC61B | 303 | TRIM6 |  |  |  |  |
|  |  |  |  | 304 | IRF7 | 304 | GBP4 |  |  |  |  |
|  |  |  |  | 305 | RPTOR | 305 | IFNAR1 |  |  |  |  |
|  |  |  |  | 306 | GAB2 | 306 | PPP2R1B |  |  |  |  |
|  |  |  |  | 307 | APOBEC3G | 307 | SPHK1 |  |  |  |  |
|  |  |  |  | 308 | AAAS | 308 | NUP58 |  |  |  |  |
|  |  |  |  | 309 | EGR1 | 309 | HSPA9 |  |  |  |  |
|  |  |  |  | 310 | FAAP24 | 310 | EIF2S1 |  |  |  |  |
|  |  |  |  | 311 | IFIT1 | 311 | NPM1 |  |  |  |  |
|  |  |  |  | 312 | TUBB4A | 312 | NUP133 |  |  |  |  |
|  |  |  |  | 313 | IL5RA | 313 | CENPS |  |  |  |  |
|  |  |  |  | 314 | MAPK3 | 314 | STAT5A |  |  |  |  |
|  |  |  |  | 315 | TUBB2A | 315 | FASLG |  |  |  |  |
|  |  |  |  | 316 | EIF4E3 | 316 | DNAJC3 |  |  |  |  |
|  |  |  |  | 317 | AKT1 | 317 | NUP155 |  |  |  |  |
|  |  |  |  | 318 | IFIT5 | 318 | MAPK10 |  |  |  |  |
|  |  |  |  | 319 | KPNA4 | 319 | TGFBR3 |  |  |  |  |
|  |  |  |  | 320 | CAMK2B | 320 | NUP50 |  |  |  |  |
|  |  |  |  | 321 | SEH1L | 321 | RAP1A |  |  |  |  |
|  |  |  |  | 322 | VCAM1 | 322 | PPARA |  |  |  |  |
|  |  |  |  | 323 | EIF4EBP1 | 323 | TYK2 |  |  |  |  |
|  |  |  |  | 324 | UBE2I | 324 | FANCB |  |  |  |  |
|  |  |  |  | 325 | EIF4A3 | 325 | CHUK |  |  |  |  |
|  |  |  |  | 326 | SKP1 | 326 | B2M |  |  |  |  |
|  |  |  |  | 327 | BCL3 | 327 | CALM2 |  |  |  |  |
|  |  |  |  | 328 | PTPN6 | 328 | NUP107 |  |  |  |  |
|  |  |  |  | 329 | IRF2 | 329 | CDK1 |  |  |  |  |
|  |  |  |  | 330 | PIK3R2 | 330 | DNAJA1 |  |  |  |  |
|  |  |  |  | 331 | IFNA5 | 331 | UBA52 |  |  |  |  |
|  |  |  |  | 332 | FGF10 | 332 | TPM2 |  |  |  |  |
|  |  |  |  | 333 | HLA-F | 333 | RPS6 |  |  |  |  |
|  |  |  |  | 334 | IFNL1 | 334 | MAPKAPK3 | |  |  |  |
|  |  |  |  | 335 | GRB14 | 335 | RXRA |  |  |  |  |
|  |  |  |  | 336 | RIGI | 336 | TPR |  |  |  |  |
|  |  |  |  | 337 | IFI27 | 337 | HSPA5 |  |  |  |  |
|  |  |  |  | 338 | TRAIP | 338 | EIF2S3 |  |  |  |  |
|  |  |  |  | 339 | IRF9 | 339 | GBP2 |  |  |  |  |
|  |  |  |  | 340 | FLNB | 340 | PTPN2 |  |  |  |  |
|  |  |  |  | 341 | HLA-E | 341 | CD44 |  |  |  |  |
|  |  |  |  | 342 | STAT4 | 342 | XRCC5 |  |  |  |  |
|  |  |  |  | 343 | HSPA1A | 343 | STAT1 |  |  |  |  |
|  |  |  |  | 344 | RAF1 | 344 | PRMT1 |  |  |  |  |
|  |  |  |  | 345 | ERG | 345 | TUBA1C |  |  |  |  |
|  |  |  |  | 346 | CIITA | 346 | PRDX6 |  |  |  |  |
|  |  |  |  | 347 | HLA-DRA | 347 | SLIRP |  |  |  |  |
|  |  |  |  | 348 | MAVS | 348 | HLA-C |  |  |  |  |
|  |  |  |  | 349 | HSPA1B | 349 | AMPD2 |  |  |  |  |
|  |  |  |  | 350 | SFPQ | 350 | IFITM3 |  |  |  |  |
|  |  |  |  | 351 | PTPRC | 351 | NCOR1 |  |  |  |  |
|  |  |  |  | 352 | OAS3 | 352 | IFI44L |  |  |  |  |
|  |  |  |  | 353 | CMTR1 | 353 | RPTOR |  |  |  |  |
|  |  |  |  | 354 | EIF4G3 | 354 | CAMK2G |  |  |  |  |
|  |  |  |  | 355 | CBL | 355 | PIK3CA |  |  |  |  |
|  |  |  |  | 356 | POM121C | 356 | RPS27A |  |  |  |  |
|  |  |  |  | 357 | FANCB | 357 | NUP188 |  |  |  |  |
|  |  |  |  | 358 | CDK13 | 358 | UBB |  |  |  |  |
|  |  |  |  | 359 | IFITM1 | 359 | NUP98 |  |  |  |  |
|  |  |  |  | 360 | NCOR1 | 360 | TMSB10 |  |  |  |  |
|  |  |  |  | 361 | NUP98 | 361 | NUP205 |  |  |  |  |
|  |  |  |  | 362 | PPARA | 362 | CREM |  |  |  |  |
|  |  |  |  | 363 | MTOR | 363 | RACK1 |  |  |  |  |
|  |  |  |  | 364 | CENPX | 364 | KPNA2 |  |  |  |  |
|  |  |  |  | 365 | FANCG | 365 | DHX9 |  |  |  |  |
|  |  |  |  | 366 | PLCG1 | 366 | RAPGEF1 |  |  |  |  |
|  |  |  |  | 367 | NFKB1 | 367 | PTPN11 |  |  |  |  |
|  |  |  |  | 368 | FAAP100 | 368 | STAT3 |  |  |  |  |
|  |  |  |  | 369 | IKBKB | 369 | EIF4G2 |  |  |  |  |
|  |  |  |  | 370 | FAAP20 | 370 | ADAR |  |  |  |  |
|  |  |  |  | 371 | MAP3K1 | 371 | CMTR1 |  |  |  |  |
|  |  |  |  | 372 | NFKBIA | 372 | PIAS3 |  |  |  |  |
|  |  |  |  | 373 | MC4R | 373 | CAMK2D |  |  |  |  |
|  |  |  |  | 374 | MAPK14 | 374 | FANCC |  |  |  |  |
|  |  |  |  | 375 | SOCS1 | 375 | FTL |  |  |  |  |
|  |  |  |  | 376 | MAP2K3 | 376 | RPLP1 |  |  |  |  |
|  |  |  |  | 377 | IL11RA | 377 | ANXA2 |  |  |  |  |
|  |  |  |  | 378 | PPP2CA | 378 | ATF6 |  |  |  |  |
|  |  |  |  | 379 | NUP43 | 379 | ILF3 |  |  |  |  |
|  |  |  |  | 380 | CENPS | 380 | MAPT |  |  |  |  |
|  |  |  |  | 381 | NUP214 | 381 | GAS6 |  |  |  |  |
|  |  |  |  | 382 | ILF3 | 382 | IFNL3 |  |  |  |  |
|  |  |  |  | 383 | FLNA | 383 | PPM1B |  |  |  |  |
|  |  |  |  | 384 | TRIM46 | 384 | TP53BP2 |  |  |  |  |
|  |  |  |  | 385 | CLK2 | 385 | OAS3 |  |  |  |  |
|  |  |  |  | 386 | NUP42 | 386 | GBP5 |  |  |  |  |
|  |  |  |  | 387 | IFI6 | 387 | PIN1 |  |  |  |  |
|  |  |  |  | 388 | EIF4A1 | 388 | BCL2L2 |  |  |  |  |
|  |  |  |  | 389 | SFN | 389 | IFITM1 |  |  |  |  |
|  |  |  |  | 390 | CAMK2G | 390 | PIK3R2 |  |  |  |  |
|  |  |  |  | 391 | NCAM1 | 391 | EIF4E3 |  |  |  |  |
|  |  |  |  |  |  | 392 | PTPRC |  |  |  |  |
|  |  |  |  |  |  | 393 | FNTB |  |  |  |  |
|  |  |  |  |  |  | 394 | KPNA7 |  |  |  |  |
|  |  |  |  |  |  | 395 | ICAM2 |  |  |  |  |
|  |  |  |  |  |  | 396 | HLA-DRB1 | |  |  |  |
|  |  |  |  |  |  | 397 | VAV1 |  |  |  |  |
|  |  |  |  |  |  | 398 | TRIM10 |  |  |  |  |
|  |  |  |  |  |  | 399 | MAPK3 |  |  |  |  |
|  |  |  |  |  |  | 400 | TUBA4B |  |  |  |  |
