## Supplemental Data 5 for "Novel Synthetic Promoter Armed Oncolytic Herpes Simplex Virus For Treatment of PAX3-FOXO1 Positive Rhabdomyosarcoma"

The raw fastq.gz and counts after alignment to the human GRCh38.p14 were submitted to GEO (GEO Submission (GSE334474)). This was not included in the manuscript, as the GEO submission is not finalized. Upon completion I will contact the journal to update this in whichever section the editor deems appropriate.
